# High interleukin-3 concentrations prime murine myeloid progenitor cells towards granulocytic differentiation

**DOI:** 10.1101/2024.01.24.576972

**Authors:** Mariam Hakobyan, Hannah Rohdjeß, Daniel B. Lipka

## Abstract

Impaired differentiation a key feature of many hematopoietic malignancies. To decipher the molecular processes underlying malignant transformation, it is important to understand the mechanisms regulating hematopoietic differentiation. Cell lines derived from transformed hematopoietic progenitors or from leukemia patients have proven to be valuable model systems for mechanistic investigations of hematopoiesis. In the present work, we investigated the cell-intrinsic differentiation capacity of the interleukin 3 (IL-3) dependent murine myeloid progenitor cell line 32D. We demonstrated that 32D cells have a cell-intrinsic granulocytic differentiation potential which requires the presence of high IL-3 concentrations in order to induce granulocytic priming. We also show that 32D cells still proliferate at comparable rates but lack granulocytic priming in the presence of intermediate IL-3 concentrations. This phenotypic plasticity is fully reversible and entirely depends on the IL-3 concentrations present in the culture media. RNA-seq analysis revealed that, among other myeloid genes, the expression of *Csf3r* is strongly induced exclusively in the presence of high IL-3 concentrations, likely explaining the granulocytic differentiation in response to G-CSF treatment. Together, our work provides detailed cellular and molecular insights into the phenotypic plasticity of 32D cells driven by different IL-3 concentrations and suggest culture conditions for different experimental set ups.

**Key points:**

- 32D cells represent a murine myeloid cell line with endogenous granulocytic differentiation potential.
- 32D cells present molecular and functional plasticity depending on their culture conditions.
- High IL-3 culture conditions prime 32D cells towards the granulocytic lineage.
- Low IL-3 culture conditions lead to a reversible loss of the granulocytic differentiation potential of 32D cells while fully maintaining their proliferative activity.

## Introduction

Hematopoiesis ensures the production of all mature blood cell types according to the current requirements of the organism. Impaired differentiation is one of the main characteristics of hematopoietic malignancies. To decipher the molecular processes underlying malignant transformation, a detailed understanding of normal hematopoietic differentiation and its control mechanisms is of central importance [1]. While the use of primary bone marrow cells is the most direct way to study hematopoiesis, their isolation, maintenance and manipulation pose significant technical challenges. Therefore, despite their limitations with regard to the direct transferability of findings to the biological situation *in vivo*, cell lines are valuable model systems for the investigation and manipulation of biological processes. The 32D cell line was first described in 1983 and, to date, it is one of to the most frequently used cell line models in hematopoietic research [2]. 32D cells depend on IL-3 for survival and proliferation and can be differentiated into mature granulocytes when stimulated with G-CSF in the absence of IL-3 [2-4]. For the culture of 32D cells, two different options for IL-3 supplementation have been described in the literature: the addition of recombinant IL-3 (rIL-3) or culture in the presence of WEHI-3B conditioned medium (CM). CM consists of filtered cell culture supernatant from the WEHI-3B cell line which secretes IL-3 into the culture medium [2]. While using rIL-3 is the most controlled and reproducible way of culturing 32D cells, the application of CM is more affordable and has, therefore, been used far more frequently. Intriguingly, previous publications subjecting CM-cultured 32D cells to granulocytic differentiation either ectopically expressed the G-CSF receptor (G-CSF-R) or pre-cultured 32D cells for a short period of time in medium supplemented with rIL-3 [4, 5]. However, despite being used over several decades, to date, it remains unclear how culture conditions affect the differentiation potential of 32D cells.

Here, we provide a systematic analysis of the impact of different culture conditions on the cellular and molecular phenotype of 32D cells. We explored how CM and different concentrations of rIL-3 affect cell viability, cell proliferation and, most importantly, granulocytic differentiation. Further, we interrogated transcriptional and epigenetic changes upon switching culture conditions to reveal the molecular changes underlying the observed phenotypic plasticity.

## Results

### Differences in surface marker expression profiles and granulocytic differentiation capacity of 32D cells are dependent on culture conditions

We obtained 32D cells from the German Collection of Microorganisms and Cell Cultures (DSMZ) and cultured the cells in the presence of 10 ng/mL rIL-3 (IL3-32D). We observed that 32D cells differentiated towards granulocytes in the presence of 20 ng/mL G-CSF upon rIL-3 removal (***Suppl. Figure 1A***). Importantly, no ectopic expression of G-CSF receptor was necessary under these conditions. In contrast, when pre-culturing 32D cells in the presence of CM (CM-32D), they failed to survive and to differentiate in the presence of G-CSF upon CM removal (***Suppl. Figure 1B***). Using flow cytometry at steady-state, we observed that IL3-32D cells presented 4-fold higher cell surface expression of the myeloid marker CD11b compared to CM-32D cells (***Figure 1A,B***) [6, 7]. Analysis of additional surface markers showed low levels of cKit expression in both CM- and IL3-32D cells, whereas IL3-32D cells overall expressed higher levels of CD115, Ly6G and Ly6C (***Figure 1C, Suppl. Figure 2***).

**Figure 1:**
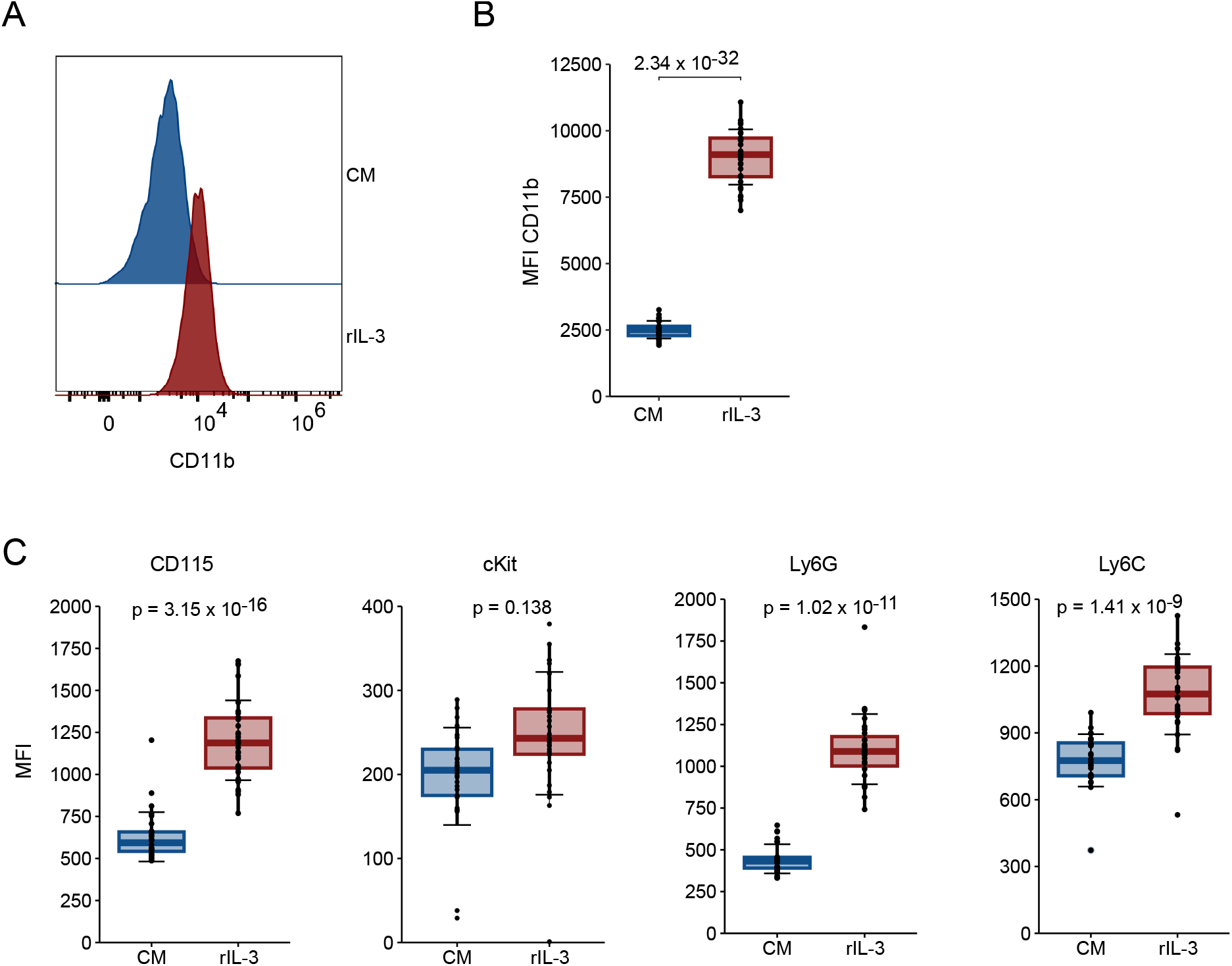
Surface marker expression patterns differs between CM- and IL3-32D cells. CM- and IL3-32D cells were analyzed by flow cytometry at steady-state. (**A**) Histogram of CD11b expression in CM- and IL3-32D cells (n=1). (**B**,**C**) Expression of CD11b (**B**), CD115, cKit, Ly5G, and Ly6C (**C**) on CM- and IL3-32D cells. MFIs were calculated with the FlowJo software and plotted in R. Boxplot horizontal lines denote the mean, edges denote the interquartile range (IQR), whiskers denote 1.5 × IQR and individual data points are represented by dots. N=27 per group. Statistical significance was tested using a one-sided student’s t-test.CM: CM-32D; rIL-3: IL3-32D

### Granulocytic differentiation capacity can be restored by rIL-3 exposure

To understand how the culture media affect the cell surface expression of CD11b and the neutrophil differentiation capacity of 32D cells, a controlled reciprocal exchange of culture media was performed (***Figure 2A***). Briefly, CM-32D and IL3-32D cells were thoroughly washed and split into two groups each – one receiving the original culture medium, the other receiving the alternative medium, resulting in four culture conditions (rIL-3, CM, rIL3->CM, CM->rIL3). Response to media switching was monitored via flow cytometry every 24 hours using CD11b expression as a surrogate marker for differentiation capacity. Additionally, differentiation assays were performed and RNA as well as DNA were collected to study the differentiation capacity, the transcriptional and the epigenetic changes in response to media switching.

**Figure 2:**
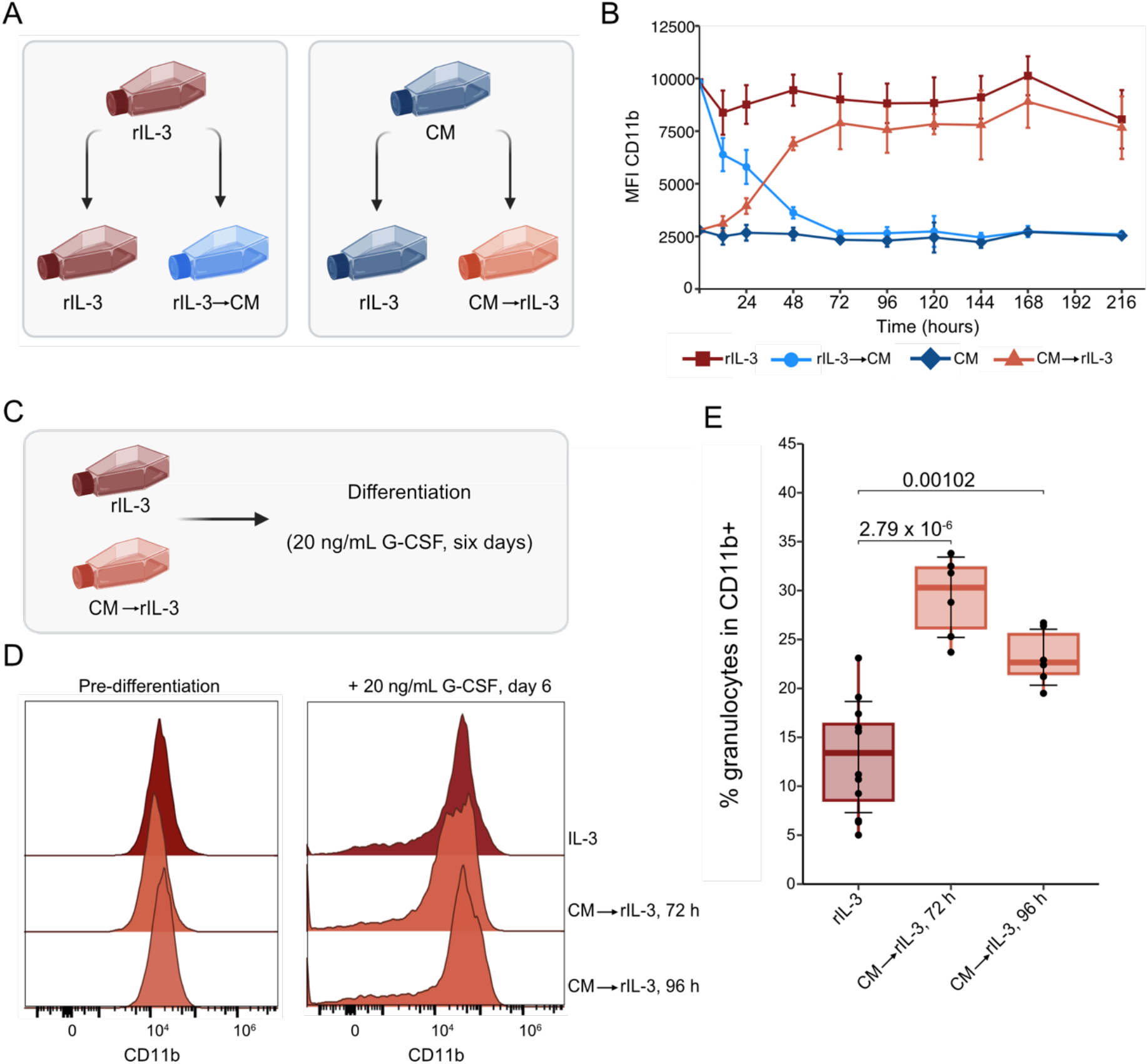
Differentiation potential of CM-32D cells can be restored by exposure to rIL-3. (**A**) Experimental design: 32D cells that had been cultured long-term in either 10% CM or 10 ng/ml rIL-3 (“primary culture”; CM; rIL-3) were washed and two aliquots, each, were resuspended in either of the two media (“secondary culture”; CM->rIL-3; rIL-3->CM). CD11b expression, RNA-seq and DNA methylome analysis were performed at defined timepoints. (**B**) CD11b expression was measured by flow cytometry in the original culture and at defined timepoints after medium switching. MFIs were calculated using the FlowJo software. N=3 per condition. (**C**) Experimental procedure: CM- or IL3-32D cells were washed, resuspended in rIL-3 medium and cultured for 72 or 96 hours. Granulocytic differentiation was initiated with G-CSF and the cells were analyzed after 6 days. (**D**) CD11b expression before and after differentiation with 20 ng/ml G-CSF for 6 days. (**E**) Quantification of the granulocyte proportions within the CD11b+ fractions after differentiation (20 ng/ml G-CSF for 6 days). rIL-3 N=10, CM->rIL-3 12 h N=6, CM->rIL-3 72 h N=6. Boxplot lines denote the mean, edges denote the IQR, whiskers denote 1.5 × IQR and individual data points are represented by dots. Statistical significance was tested by pairwise comparison with one-sided student’s t-test and Bonferroni correction. (**A**) and (**C**) were created using BioRender.

While cells kept in their original culture medium showed stable CD11b expression levels throughout the experiment, rIL3->CM swapped 32D cells showed a significant loss of CD11b expression already after 12 h of exposure to CM and reached a plateau 72 h after the media switch (***Figure 2B***). Conversely, CM->rIL3 cells started to upregulate CD11b expression after 12 h of rIL-3 exposure and reached stable CD11b expression after 72 h. In both media-swapped cultures, the CD11b expression levels achieved were comparable to that of the corresponding steady-state cultures.

Next, we wanted to investigate whether recovery of CD11b expression in CM->rIL3 cells indicates restoration of granulocytic differentiation capacity. Therefore, CM->rIL3 cells were put on differentiation medium (20 ng/mL G-CSF) 72 h or 96 h after start of rIL-3 exposure. Cells in continuous rIL-3 culture served as controls (***Figure 2C***). Levels of CD11b expression were comparable for IL3-32D and CM->rIL3 cells at both timepoints (***Figure 2D***). Nevertheless, we observed more efficient granulocytic differentiation in CM->rIL-3 cells as compared to IL3-32D cells. This effect was statistically significant and most pronounced in CM->rIL3 cells 72 h after the medium switch. Taken together, we observed that the cell surface expression of CD11b and the granulocytic differentiation capacity of 32D cells are correlated phenotypes. Furthermore, we showed that exposure to rIL-3 is sufficient to recover the differentiation potential of CM-32D cells.

### Phenotypic effects in 32D cells are dependent on IL-3 concentration in cell culture media

CM is a non-standardized source of IL-3 and IL-3 concentrations in CM might be variable. Therefore, we sought to investigate if the observed phenotypic differences between CM-32D and IL3-32D cells can be recapitulated by titrating the rIL-3 concentration in the cell culture media. For this experiment, CM-32D cells were washed and subsequently exposed to decreasing rIL-3 concentrations ranging from 10 ng/mL to 4 pg/mL. Cells were counted using a cell counter and viability, CD11b expression and cell size were assessed by flow cytometry 72 h after medium switching (***Figure 3A***). Optimal proliferation conditions for 32D cells were observed at rIL-3 concentrations between 250 pg/mL and1 ng/mL. Cell proliferation only decreased at rIL-3 concentrations < 125 pg/mL, whereas cell viability started to decrease only when rIL-3 was < 16 pg/mL (***Figure 3B***). In contrast to the robust maintenance of proliferation and cell viability across a broad range of rIL-3 concentrations, CD11b expression was closely linked to rIL-3 concentrations present in the media: at 10 ng/mL rIL-3 CD11b expression was highest and showed an almost linear decrease until 31 pg/mL before reaching a plateau at lower concentrations (***Figure 3B***). In summary, these data show that culturing 32D cells at rIL-3 concentrations between 250 pg/mL and 500 pg/mL mimics the phenotypes observed in the presence of CM. To exclude that CM contains factors that actively suppress CD11b upregulation, we next tested whether adding rIL-3 to CM would be sufficient to recover CD11b expression in CM-32D cells (***Figure 3C***). Seventy-two hours after adding rIL-3 to CM, cells significantly upregulated CD11b expression on the cell surface compared to CM-32D cells and the CD11b levels observed were comparable to those observed in CM->rIL3 cells (***Figure 3D***). Taken together, these data suggest that the phenotypic differences between CM-32D and IL3-32D cells can be attributed to different IL-3 concentrations in the cell culture media.

**Figure 3:**
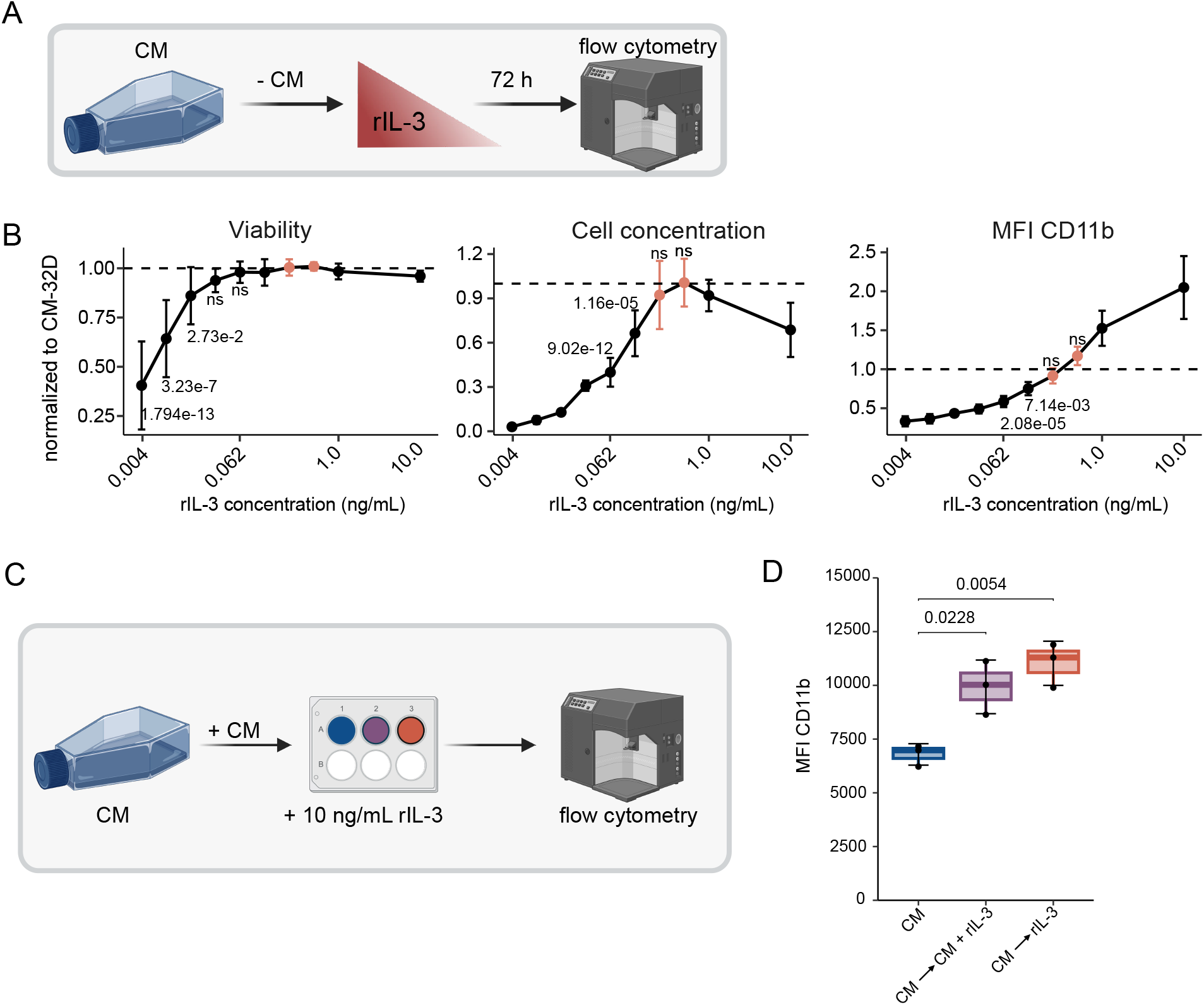
Low rIL-3 concentrations phenocopy CM-32D cells. (**A**) Experimental design: 32D parental cells cultured in 10% CM were washed and seeded in medium containing defined concentrations of rIL-3. A control well was seeded in CM. (**B**) Seventy-two hours post-seeding viability was determined by trypan blue, cells were counted using LUNA-II automated cell counter and stained to measure CD11b expression by flow cytometry. All measurements were normalized to the respective CM control; N=6 per group. For all statistical analyses, analysis of variance with Bonferroni correction was performed. (**C**) Experimental design: CM-32D cells were washed and seeded in medium with the respective supplements: 10% CM (CM), 10% CM + 10 ng/mL rIL-3 (CM->CM+rIL-3), 10 ng/ml rIL-3 (CM->rIL-3). (**D**) Seventy-two hours post-seeding, 1x10^5^ cells were used for staining with anti-CD11b antibody and analyzed by flow cytometry. MFIs were calculated using the FlowJo software. N=3 per group. Boxplot lines denote the mean, edges denote the IQR, whiskers denote 1.5 × IQR and individual data points are represented by dots. Statistical significance was tested by pairwise comparison with one-sided student’s t-test and Bonferroni correction. **(A)** and **(C)** were created using BioRender.

### Transcriptional analysis shows molecular plasticity of 32D cells with myeloid priming in high rIL-3 culture conditions

To better understand the molecular states and dynamic adaptation of IL3-32D and CM-32D cells, we analyzed DNA methylomes and transcriptomes both at steady state and after media switching. While the DNA methylomes remained stable for more than a week (***Suppl. Figure 3***), transcriptional changes were observed already 12 h after media switching (***Figure 4A***). Principal component (PC) analysis based on normalized counts showed grouping of 32D cells, both based on their steady-state as well as on the media switch conditions: while the first PC (PC1) discriminated cells according to their current culture conditions, the second PC (PC2) separated cells according to their original culture conditions. The CM->rIL3-mediated effects on the transcriptome occurred faster than the rIL3->CM-mediated effects (***Figure 4A***). We identified 1202 differentially expressed genes between CM-32D and IL3-32D cells, of which 689 (57%) were upregulated in IL3-32D and 513 (43%) were upregulated in CM-32D cells (***Figure 4B, Suppl. Figure 4***). Several genes from both groups were divided into early- and late-responders (***Figure 4C***). The expression of early-responder genes was quickly adapted upon the medium exchange in both directions (blocks A3, A4 and B1). Late-responder genes presented little to no effect 12 h (or 72 h) after medium exchange (blocks A1, A2 and B2). Further, in line with our experimental observations, genes upregulated in IL3-32D cells were enriched for neutrophil- and immune cell-associated pathways such as “neutrophil degranulation”, “neutrophil mediated immunity”, “cellular response to lipopolysaccharides (LPS)”, and “formation of specific and tertiary granules” (***Suppl. Figure 5***). We analyzed the expression of neutrophil and myeloid lineage-specific genes to assess whether the culture condition affect myeloid gene expression programs (***Figure 4D***). The expression of several neutrophil marker genes was upregulated in IL3-32D cells. For example, *Cebpe*, a key transcription factor involved in neutrophil maturation, as well as *Csf3r, Ly6g* and *Itgam* which encode for the G-CSF receptor and the cell surface markers Ly6G and CD11b, respectively. In contrast, we found the expression of several monocyte and macrophage-associated genes, such *Cd14, Cd93, Cd300a* and *Nav1* to be upregulated in CM-32D cells. The expression of the genes encoding for the IL-3- and G-CSF-receptors, *IL-3ra* and *Csf3r*, respectively, showed an inverse expression pattern (***Figure 4E-F***). IL3-32D cells in steady-state culture presented the lowest expression of *IL-3ra* coupled with the highest expression of *Csf3r*. On the contrary, culturing IL3-32D cells in CM culture led to a gradual increase in *IL-3ra* and a continuous decrease in *Csf3r* expression. Similarly, the reverse phenotype was observed in CM-32D and CM->rIL3 cells.

**Figure 4:**
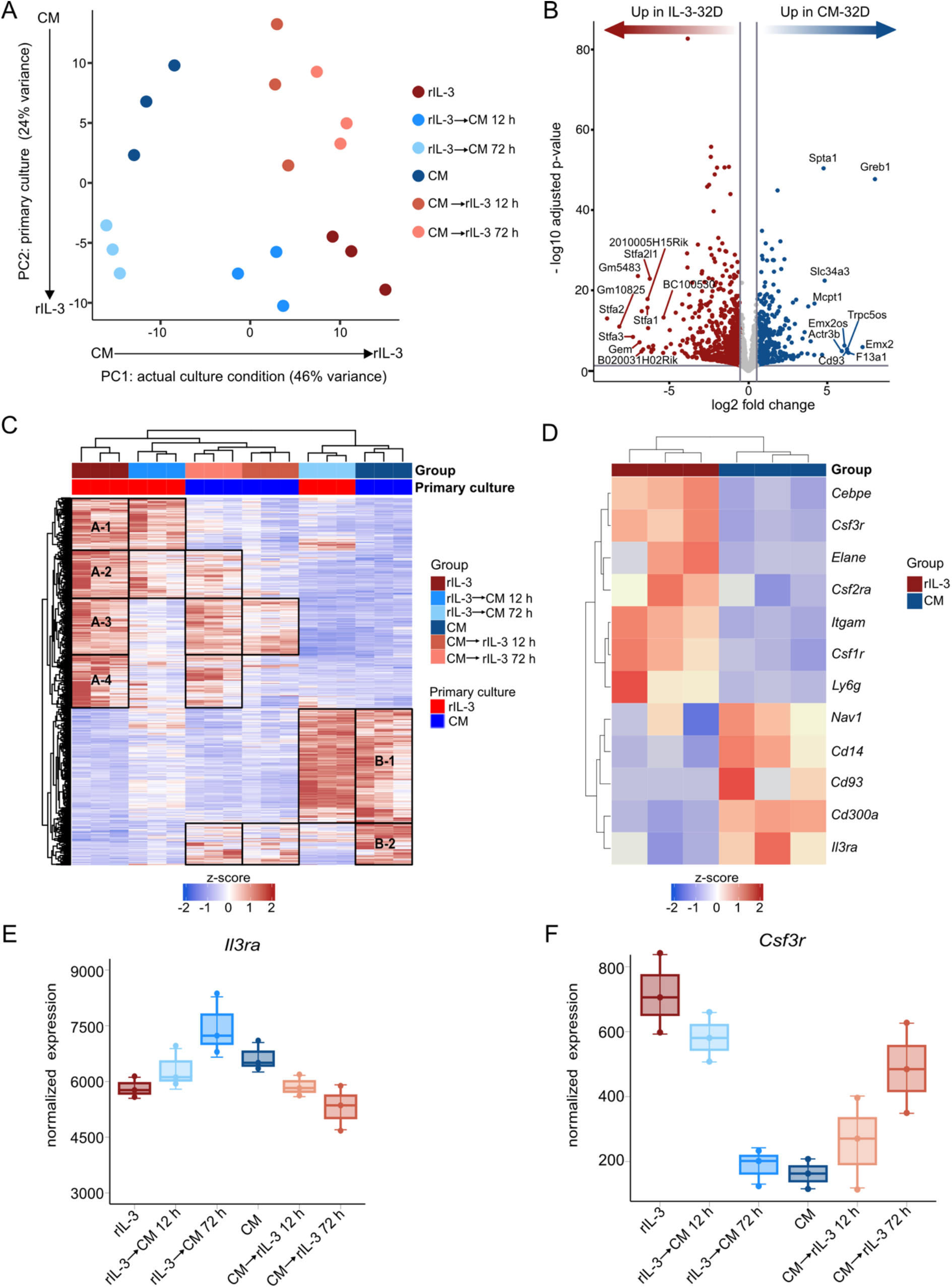
CM-32D cells downregulate neutrophil lineage-specific genes. (**A**) PCA based on normalized and log transformed expression data retrieved from DEseq2. (**B**) Differentially expressed genes (DEGs) between rIL-3 and CM steady state 32D cells. Top 10 genes with the highest fold change per condition are annotated. (**C)** Heatmap showing the expression of all 1202 DEGs between steady-state IL3-32D and CM-32D cells. Rectangles indicate different gene groups based on their early and late response to medium exchange. A1-A4: clusters of genes with high expression in rIL-3; B1-B2: cluster of genes with high expression in CM (**D**) Heatmap showing the expression of neutrophil-specific genes in steady state CM- and IL3-32D cells. (**E, F**) Expression of *Il3ra* (**E**) and *Csf3r* (**F**). Depicted are normalized counts for IL3-32D cells (rIL-3), CM-32D cells (CM) as well as medium switched 32D cells (rIL-3->CM, CM->rIL-3) analyzed 12 and 72 hours after media switching. Boxplot lines denote the mean, edges denote the IQR and whiskers denote 1.5 × IQR and individual data points are represented by dots.

Taken together, these data highlight that the transcriptomes of 32D cells can be dynamically rewired in response to cell culture conditions. We demonstrate that under conditions with high IL-3 exposure 32D cells upregulate neutrophil marker genes, which allows them to differentiate towards mature neutrophils upon G-CSF stimulation.

## Discussion

32D cells depend on IL-3 signaling for survival and proliferation. While clone 3 of 32D cells (32Dcl3) possesses the cell intrinsic potential of granulocytic differentiation and can undergo granulopoiesis upon G-CSF stimulation, clone 10 requires exogenous expression of the G-CSF receptor (32D/G-CSF-R) [4, 5, 8]. IL-3 can be supplemented to the culture medium by using the supernatant of either WEHI-3B cells or of HEK293 cells which overexpress IL-3 [2, 5]. Zjavlovskaja and colleagues have previously described a detailed granulocytic differentiation protocol for 32D/G-CSF-R cells which were cultured in conditioned medium before the start of differentiation [5]. However, we observed that 32Dcl3 cells cultured in conditioned medium do not differentiate but instead undergo cell death upon exposure to G-CSF. Since it has been shown that 32Dcl3 cells possess the cell-intrinsic potential of granulocytic differentiation without the requirement for G-CSF-R overexpression, we systematically tested the effects of different culture conditions on differentiation potential both at the functional and the molecular level. Our data confirmed the endogenous capacity of 32Dcl3 cells to undergo granulocytic differentiation. However, we demonstrated that the granulocytic differentiation potential crucially depends on the IL-3 levels present in culture medium: low IL-3 concentrations such as the ones present in conditioned medium lead to loss of granulocytic differentiation potential. Our flow cytometry and transcriptome data revealed, that CM-32D cells downregulated several cell surface markers, including CD11b and the expression of G-CSF-R (*Csf3r*) but upregulated the expression of IL-3 receptor (*Il3ra*), presumably to compensate for the low availability of IL-3. While we did not quantify IL-3 concentrations in CM, titration experiments of rIL-3 in cell culture medium showed that 32D cells exposed to lower rIL-3 concentrations present cellular phenotypes highly reminiscent of those observed CM-32D cells. Intriguingly, culturing 32D cells in CM or in the presence of low rIL-3 concentrations (≥250 pg/mL) did not affect cell viability nor proliferation whereas CD11b expression was markedly reduced and granulocytic differentiation capacity was completely lost. IL-3 concentrations ≤125 pg/mL eventually decelerated cell growth, but cell viability started to decrease only at IL-3 concentrations ≤16 pg/mL. Together, these data highlight that culturing 32D cells in the presence of low IL-3 concentrations rapidly impairs their differentiation potential, while cellular adaptation mechanisms safeguard cell proliferation and survival. We further demonstrated that 32D cells cultured under low IL-3 conditions (e.g. CM-32D cells) were capable to rapidly and stably re-activate CD11b cell surface expression and to re-gain granulocytic differentiation potential by upregulating *Csf3r* upon exposure to high levels of rIL-3. This effect was observed both when changing the entire culture medium (CM->rIL-3) and upon addition of rIL-3 to CM (CM + rIL-3), further underlining that the IL-3 concentration in culture medium is the critical determinant of the observed cellular phenotypes.

Our data presented here provides systematic evidence for the cell intrinsic capacity of 32Dcl3 cells to differentiate into granulocytes. This requires exposure to high concentrations of IL-3 at least several hours in order to prime 32D cells for granulocytic differentiation and to upregulate the expression of the endogenous G-CSF receptor to make the cells responsive to G-CSF stimulation. Taken together, our data provide molecular insights into granulocytic differentiation and inform cell culture protocols that enable both economic maintenance of 32Dcl3 cells and granulocytic differentiation assays.

## Acknowledgements

We would like to thank the NGS Core Facility and the Omics IT and Data Management Core Facility, German Cancer Research Center (DKFZ), for providing excellent services. We also thank all the members of the Translational Cancer Epigenomics group, the Division of Translational Medical Oncology, and the Division of Applied Functional Genomics for fruitful discussions. This work was in part funded by the DKFZ International PhD Program and the German-Israeli Helmholtz International Research School “Cancer-TRAX” to M.H., and by grants from the “Deutsche José Carreras Leukämie-Stiftung” (DJCLS 14R/2022) and from the “Wilhelm Sander-Stiftung” (# 2022.010.1) to D.B.L.

## Conflict of interest statement

D.B.L. receives honoraria from Infectopharm. All other authors report having no conflicts of interest.

## Data availability statement

RNA-seq and EPIC-array raw data will be made publicly available via GEO (submission #). All other data will be available from the authors upon request.

## Methods

### Cell culture

All cells were cultivated in an incubator at 37°C and 5% CO_2_. All centrifugation steps were carried out at 300 g at room temperature. 32D cells were cultured in conditioned 32D medium or rIL-3 containing 32D medium after thawing (***Suppl. Table 1***). Cells were split every other or third day to an approximate density of 1x10^5^ cells/mL. WEHI-3B cells were cultured in DMEM complete. Prior to splitting, the semi-adherent WEHI-3B cells were scratched from the cell culture flask surface using a cell scraper.

### Production of WEHI supernatant as medium supplement

WEHI-3B cells were thawed and cultured for ten days. Per T175 flask, 1x10^6^ cells/mL were seeded in 25 mL medium. After 24 hours, the supernatant was collected, and 25 mL fresh medium was added. Another 24 hours later, the supernatant was collected again. The cells were counted, washed and 1x10^6^ cells/mL were seeded in 25 mL per T175 flask for a second round of supernatant collection. All collected aliquots were centrifuged at 300 g for 5 minutes at 4°C. The supernatant was pooled, filtered using a 0.22 μm filter, aliquoted and stored at -20°C until further use.

### Flow cytometry

1x10^5^ cells were washed with PBS supplemented with 2% FBS (PBS/FBS) and stained in 50-100 μL staining solution for at least 20 min at 4 °C in the dark. The staining solution was prepared by diluting the respective antibodies in PBS/FBS (***Suppl. Table 2***). Subsequently, the tubes were filled up with PBS/FBS and washed before resuspension in 200 μL PBS/FBS. All centrifugation steps were carried out for 5 – 10 minutes at 300 g at 4°C. Right before flow cytometric analysis, samples were filtered through a 38 μM mesh. A BD FACS Celesta™ SORP was used to measure the samples. FCS3.0 files were exported and analyzed using FlowJo™ v10.8.1 software.

### Differentiation assay

For 32D cell differentiation, cells were collected and washed with 40 ml PBS. 10^5^ cells were seeded in 2 mL RPMI complete per well of a 6 well plate. Differentiation was induced by 20 ng/mL G-CSF (***Suppl. Table 1***). At day 6, samples were stained for surface markers (***Suppl. Table 2***) and analyzed by flow cytometry.

### Nucleic acid extraction

A minimum of 5x10^5^ cells were centrifuged at 300 g for 5 minutes. Pellets were either frozen at -80°C or directly used for nucleic acid extraction. RNA was isolated using the RNeasy plus Mini Kit (Qiagen) and eluted in 15 μL nuclease-free water. For DNA, the DNeasy blood and tissue kit (Qiagen) was used and 30 μL nuclease-free water was used for elution. Nucleic acid concentrations were measured using Nanodrop.

### Interleukin-3 titration

CM-32D cells were washed with 20 mL PBS, and 1x10^5^ cells were resuspended in 1 mL RPMI1640 complete containing different concentrations of rIL-3 in a 12-well plate. After 48 hours, the samples were split 1:3 to prevent overgrowing. Another 24 hours later, samples were counted and 1x10^5^ cells were stained with an anti-CD11b antibody for flow cytometry.

### Interleukin-3 rescue in conditioned medium

CM-32D cells were washed with 20 mL PBS, and 1x10^5^ cells were resuspended in 1 mL RPMI1640 complete, containing either of the following three supplements: 10% CM, 10% CM + 10 ng/mL rIL-3 or pure 10 ng/mL rIL-3. Forty-eight hours later, the samples were split 1:3. Another 24 hours later, samples were counted and 1x10^5^ cells were stained with an anti-CD11b antibody for flow cytometry analysis.

### Differentiation of CM->rIL3 32D cells

CM-32D cells were washed twice with 40 mL PBS. The pellet was resuspended in 32D medium containing 10 ng/mL rIL-3 and cultured for 72h or 96h with splitting after 48h. 1x10^6^ cells were then used to induce differentiation with 20 ng/mL G-CSF, as described above. 1x10^5^ cells were used to stain for differentiation markers pre-differentiation and after incubation with G-CSF for 6 days (***Suppl. Table 2***).

### Sample preparation for mouse methylation array and RNA sequencing

DNA methylation was analyzed using Infinium Mouse Methylation BeadChip (MMBC) Array from Illumina. DNA for MMBC (500 ng in 40 μL nuclease-free water) was submitted to the Microarray Core Facility of the DKFZ and processed according to in house standard procedures. RNA (1200 ng in 35 μL) was submitted to the NGS Core Facility of the DKFZ for RNA sequencing. Sequencing libraries were prepared according to the Illumina TruSeq Stranded protocol with ligation of unique dual indexes. The libraries of all samples were pooled, and 100 bp paired-end sequencing was performed on one lane of the NovaSeq6000 S4 flow cell. A sequencing depth of 139–225 million reads per sample was reached (mean: 167 million reads).

### Processing and analysis of DNA methylation array data

Analysis of DNA methylation array data was performed using the R package *RnBeads* [9]. In brief, IDAT files were imported into *RnBeads* and preprocessed using the ‘rnb.run.preprocessing’ function with the ‘rnb.options’ parameters as follows: ‘normalization.method’ set to ‘scaling’, ‘normalization.background.method’ set to ‘subtraction’, ‘filtering.greedycut.pvalue.threshold’ was set to 0.01, enabled removal of sex chromosomes (‘filtering.sex.chromosomes.removal’ set to TRUE), and all missing values were removed (‘filtering.missing.value.quantile’ set to 0). For unsupervised inspection of the data, principal component analysis (PCA) was computed using ‘pcrcomp’ from the *stats* package using the 5000 most variable CpGs (mvCpGs) across all analyzed 32D cells. The 5000 mvCpGs were defined as the CpGs with the highest standard deviation (SD) across all 32D samples.

### Analysis of RNAseq data

Analysis of gene expression data was performed using the R package *DESeq2* [10]. In brief, the raw expression matrix was loaded into *DESeq2* using the ‘DESeqDataSetFromMatrix’ function and raw counts were normalized by using ‘estimateSizeFactors’. To perform a PCA, the counts from the *DESeq2* data set were log2 transformed with ‘rlog’ and PCA was calculated using ‘plotPCA’. Differentially expressed genes (DEGs) were determined in a pairwise manner using Log2fold shrinkage with the ‘ashr’ method within the function ‘lfcShrink’. P-values were corrected for multiple testing using the Benjamini and Hochberg method. Genes with an adjusted p-value <0.05 and an absolute log2 fold change of >0.58 were considered significant. Gene set enrichment analysis was performed using the R package *enrichr*. The databases ‘GO_Molecular_Function_2018’, ‘GO_Cellular_Component_2018’, and ‘GO_Biological_Process_2018’ were used. GSEA plots were created using the ‘plotEnrichr’ function.

## Data analysis

Flow cytometry data were exported from the Diva software (BD Biosciences) in a FCS3.0 file format. Data was subsequently analyzed and visualized using the FlowJo™ software v10.8.1 (BD Biosciences). For statistical analysis in R [11], the packages *rstatix* (https://github.com/kassambara/rstatix) and *DescTools* (https://github.com/AndriSignorell/DescTools/) were used. Pairwise comparisons were performed using the “pairwise_t_test” function. Analysis of variance was performed using the “aov” function from the package *stats*. Post hoc analysis was performed with the “PostHocTest” function from the package *DescTools* using the method “lsd”. When comparing more than two groups, p-values were adjusted with Bonferroni correction by setting the ‘p.adjust.method’ to ‘bonferroni’ within the function ‘pairwise_t_test’. For data visualization the *ggplot2* package in the R environment was used [12].

