## Supplementary Figures for "High interleukin-3 concentrations prime murine myeloid progenitor cells towards granulocytic differentiation"

### Suppl. Figures

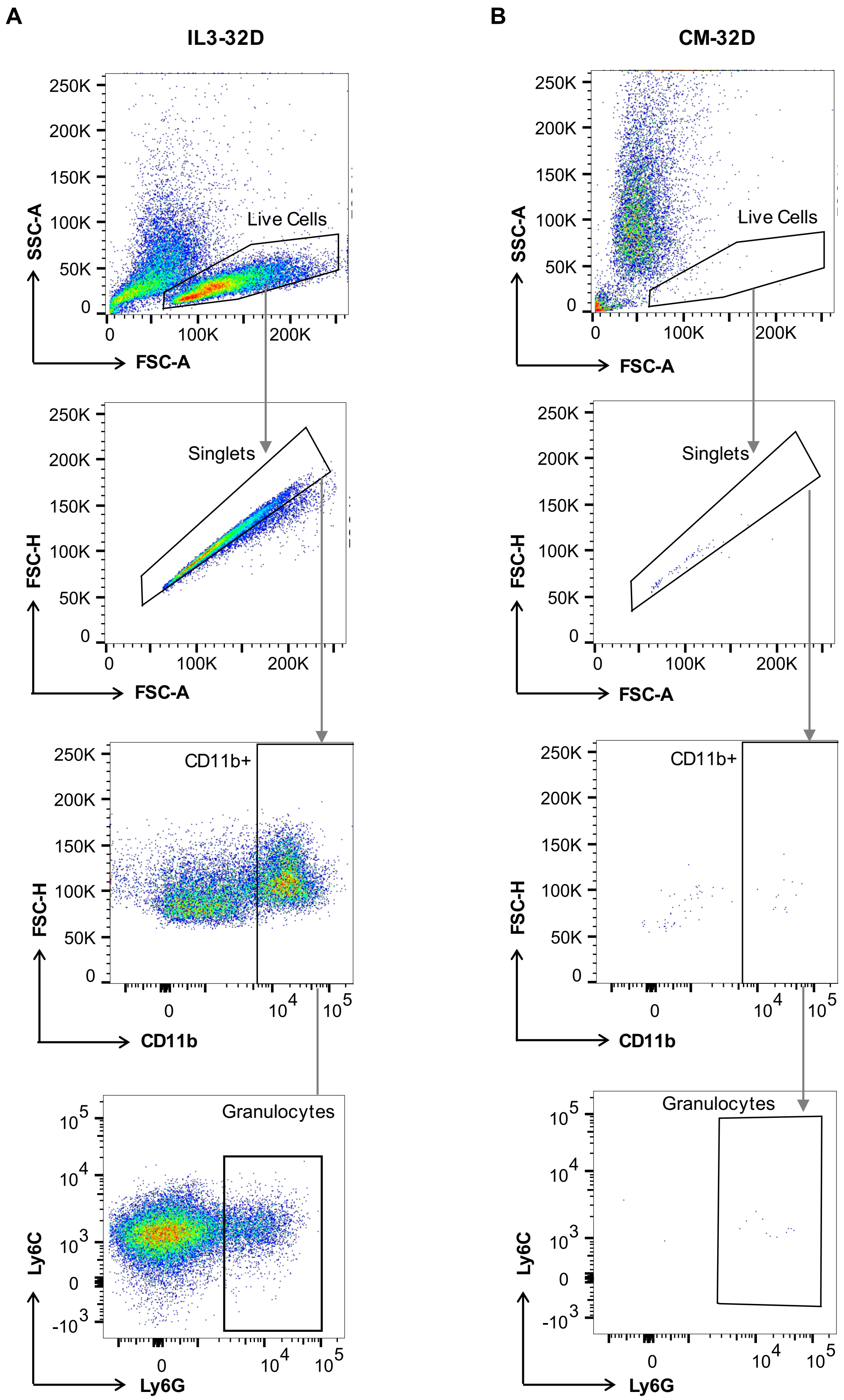


**Figure S1: CM-32D cells fail to differentiate.** (**A**) IL3-32D or (**B**) CM-32D cells were differentiated in IL-3-free medium with 20 ng/ml G-CSF for 6 days. On day 6, cells were stained and analyzed by flow cytometry. Dot plots were generated using the FlowJo software.

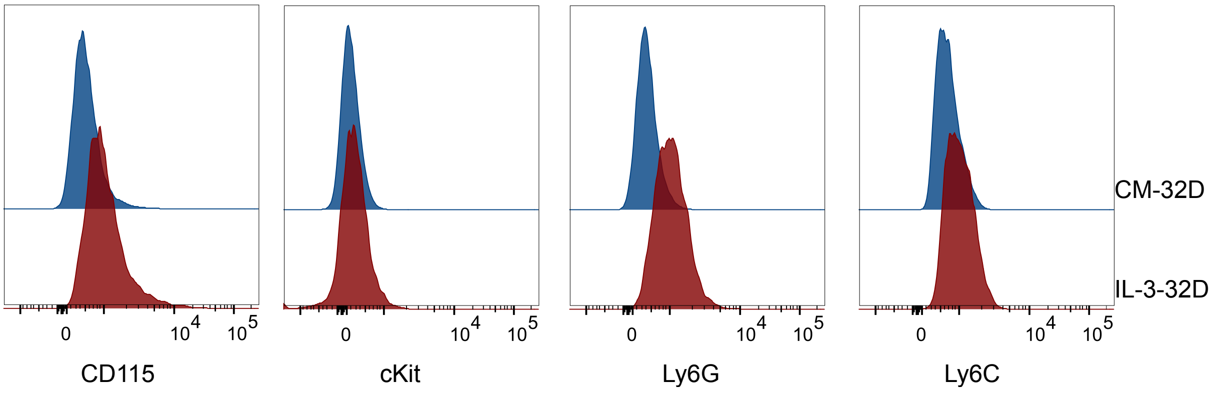


**Figure S2: Surface marker expression of CM- and IL3-32D cells.** Histogram of expression of CD115, cKit, Ly6G and Ly6C in CM- and IL3-32D cells (n=1).

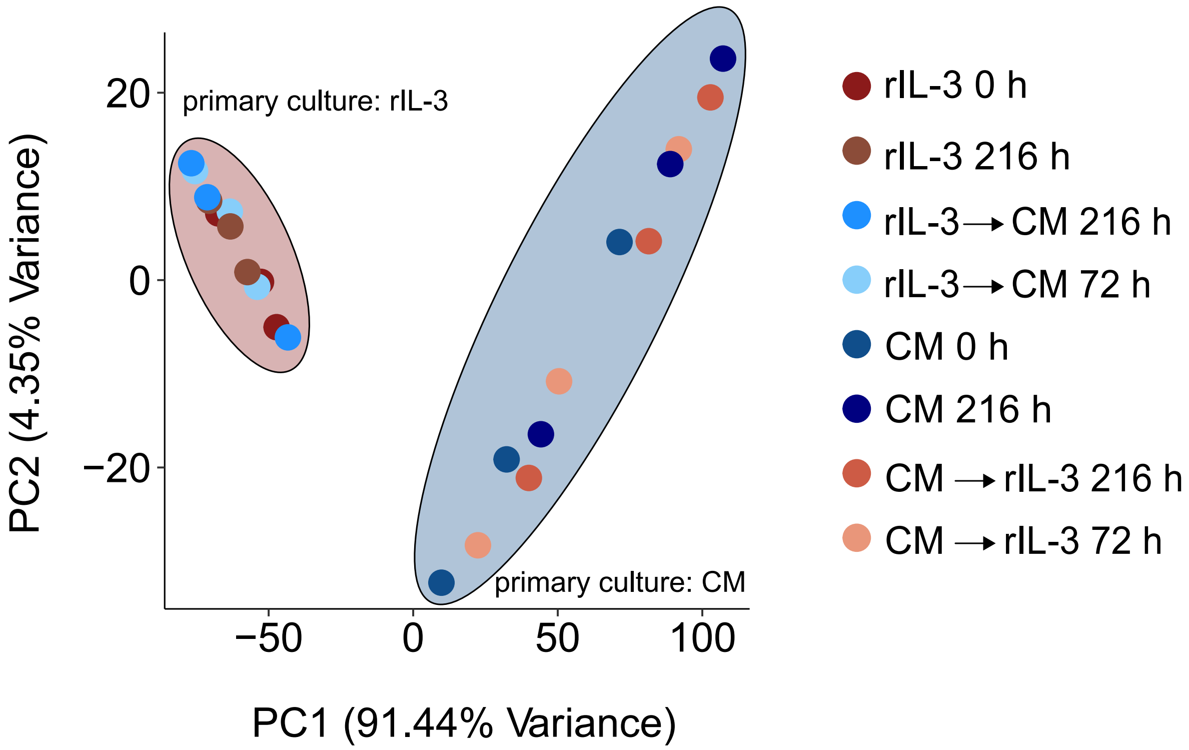


**Figure S3: DNA methylomes remain stable upon media switching.** Principal component analysis of the 5000 most variable CpGs among all measured 32D samples. Depicted are the first two principal components (PC1, PC2).

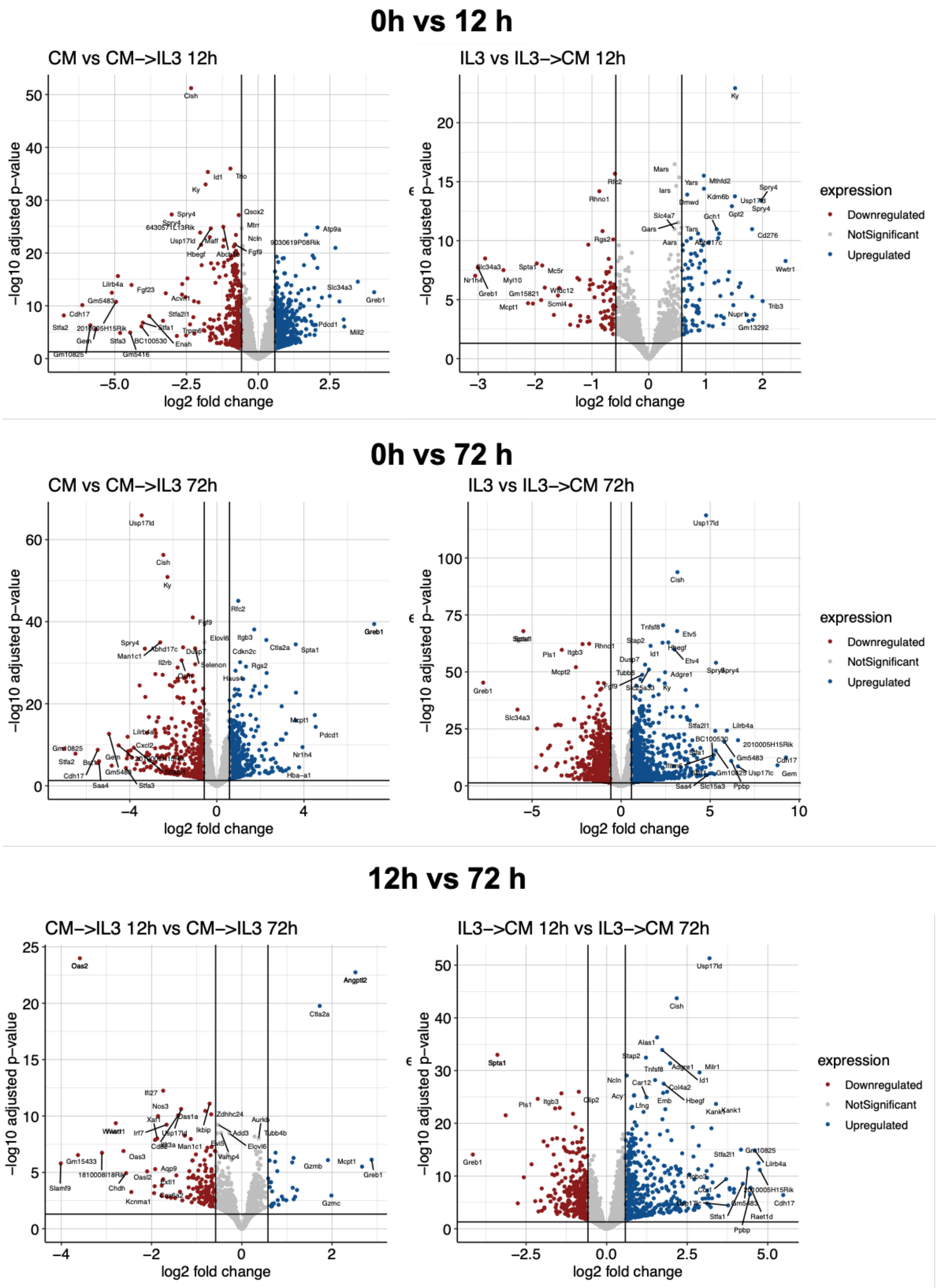


**Figure S4: The transcriptome of 32D cells adapts to the medium switch as early as 12 h after the change.** Pairwise volcano plots presenting differentially up- and downregulated genes between the indicated conditions.

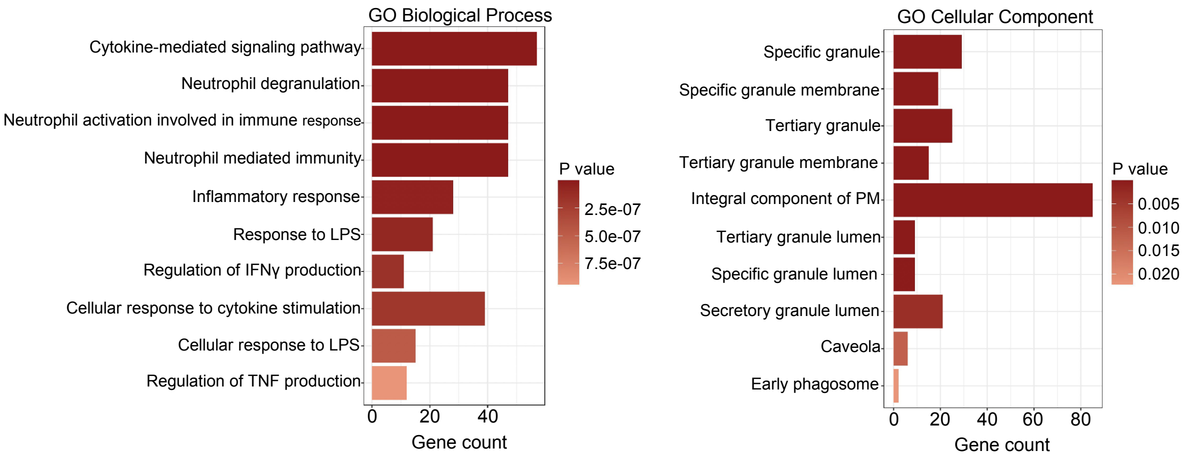


**Figure S5: Enrichment of neutrophil-specific terms in IL-3-32D cells.**
Gene set enrichment analysis (GSEA) was performed using genes differentially upregulated in IL-3-32D compared to CM-32D using enrichR and the annotated GO reference databases. PM = plasma membrane.
