## Supplementary Tables for "High interleukin-3 concentrations prime murine myeloid progenitor cells towards granulocytic differentiation"

### Supplementary Table 1: composition of cell culture media

| Name | Base medium | Supplements |
|---|---|---|
| Conditioned 32D medium (CM) | RPMI1640 | 10% FBS, 1% Penicillin/Streptomycin, 10% WEHI culture supernatant |
| recombinant IL-3 32D medium (rIL-3) | RPMI1640 | 10% FBS, 1% Penicillin/Streptomycin, 10 ng/mL rec. murine IL-3 |
| RPMI complete | RPMI1640 | 10% FBS, 1% Penicillin/Streptomycin |
| DMEM complete | DMEM | 10% FBS, 1% Penicillin/Streptomycin |
| Differentiation medium | RPMI1640 | 10% FBS, 1% Penicillin/Streptomycin, 20 ng/mL rec. murine G-CSF |

### Supplementary Table 2: antibodies used for analyzing cell surface marker expression on 32D cells

| Antibody | Fluorophore | Clone | Company | Dilution |
|---|---|---|---|---|
| CD11b | BV786 | M1/70 | Biolegend | 1:2000 |
| CD115 | BV711 | AFS98 | Biolegend | 1:1000 |
| CD117 (cKit) | APC | 2B8 | Biolegend | 1:1000 |
| Ly6C | BV605 | HK1.4 | Biolegend | 1:500 |
| Ly6G | APC-Cy7 | 1A8 | Biolegend | 1:500 |

**Supplementary Table 3: differentially expressed genes in 32D cells cultured under different media conditions**

| CM-vs-IL3 | | | | | | |
|---|---|---|---|---|---|---|
| | baseMean | log2FoldChange | lfcSE | pvalue | padj | expression |
| Prdm14 | 31 | -6 | 2 | 1,26E-06 | 1,94E-05 | Down |
| Ncoa2 | 12.570 | 0,615411675 | 0,0869404 | 1,77E-14 | 1,27E-12 | Up |
| Paqr8 | 6.633 | 0,799062175 | 0,11748181 | 2,38E-13 | 1,47E-11 | Up |
| Gm38224 | 29 | -0,63678698 | 0,47486617 | 0,00280422 | 0,01444403 | Down |
| 4931428L18Rik | 43 | 2 | 0,37362453 | 7,53E-08 | 1,55E-06 | Up |
| Gm29228 | 301 | -7 | 2 | 2,41E-07 | 4,40E-06 | Down |
| 1700001G17Rik | 11 | 0,683966835 | 0,71712438 | 0,00504881 | 0,02321841 | Up |
| Arid5a | 5.829 | -0,683812898 | 0,11394697 | 2,76E-11 | 1,14E-09 | Down |
| Gm26805 | 11 | -0,609484919 | 0,60300091 | 0,00571224 | 0,0256079 | Down |
| Aff3 | 4.376 | 0,730011042 | 0,2555685 | 4,29E-05 | 0,00041713 | Up |
| Creg2 | 4 | 0,898665303 | 1 | 0,00354252 | 0,01740579 | Up |
| Map4k4 | 1.066 | -1 | 0,28605353 | 7,86E-07 | 1,29E-05 | Down |
| Fhl2 | 25 | 1 | 0,64114525 | 0,00023648 | 0,00180461 | Up |
| Rftn2 | 532 | 0,701071711 | 0,13212455 | 1,53E-09 | 4,55E-08 | Up |
| Ica1l | 338 | 1 | 0,26584554 | 3,26E-09 | 9,16E-08 | Up |
| Ino80dos | 115 | 0,97425108 | 0,24929632 | 9,15E-07 | 1,48E-05 | Up |
| Klf7 | 1.705 | -0,663797465 | 0,12598198 | 1,72E-09 | 5,07E-08 | Down |
| Kansl1l | 2.154 | 0,854119577 | 0,14036657 | 2,52E-11 | 1,05E-09 | Up |
| Tns1 | 44 | 2 | 0,50018879 | 9,65E-07 | 1,54E-05 | Up |
| Stk36 | 70 | 0,650812167 | 0,2946956 | 0,0003466 | 0,00251966 | Up |
| Wnt10a | 66 | 2 | 0,29509232 | 2,49E-12 | 1,27E-10 | Up |
| Resp18 | 26 | -0,759368484 | 0,48543154 | 0,00120135 | 0,00716739 | Down |
| Slc4a3 | 706 | -0,727497747 | 0,24456155 | 2,97E-05 | 0,00030291 | Down |
| Gm15433 | 27 | -2 | 0,7932197 | 8,21E-05 | 0,00073016 | Down |
| Gm29284 | 18 | -0,867938615 | 0,76309888 | 0,00235914 | 0,01252511 | Down |
| Rab17 | 452 | 2 | 0,29868185 | 4,75E-12 | 2,29E-10 | Up |
| Ramp1 | 6.133 | 0,665109232 | 0,17686655 | 1,93E-06 | 2,80E-05 | Up |
| Hes6 | 2.040 | 0,816121185 | 0,10787915 | 1,04E-15 | 8,88E-14 | Up |
| Olfr1414 | 46 | -0,615591138 | 0,37771373 | 0,00164645 | 0,00930542 | Down |
| Otos | 100 | -0,661467085 | 0,20201973 | 1,21E-05 | 0,0001394 | Down |
| Sned1 | 163 | -0,816580826 | 0,24861342 | 9,56E-06 | 0,00011455 | Down |
| Gm7889 | 93 | 0,836685377 | 0,21961464 | 1,49E-06 | 2,24E-05 | Up |
| Gm6086 | 67 | 1 | 0,52831648 | 0,0001966 | 0,00155045 | Up |
| Pdcd1 | 212 | 4 | 0,56937199 | 4,90E-12 | 2,34E-10 | Up |
| 1810006J02Rik | 1.664 | -1 | 0,14803392 | 2,08E-17 | 2,27E-15 | Down |
| Phlpp1 | 2.041 | 0,996339727 | 0,15609075 | 3,44E-12 | 1,71E-10 | Up |
| Tfcp2l1 | 8 | -3 | 1 | 2,00E-05 | 0,00021596 | Down |
| Tmem185b | 9.566 | -0,697176155 | 0,08565272 | 7,08E-18 | 8,29E-16 | Down |
| Map3k19 | 32 | 2 | 0,4411347 | 1,35E-06 | 2,06E-05 | Up |
| Zranb3 | 3.160 | 0,609814682 | 0,12041428 | 4,80E-09 | 1,30E-07 | Up |

| | | | | | | |
|---|---|---|---|---|---|---|
| Sox13 | 60 | 1 | 0,26387583 | 3,94E-07 | 6,92E-06 | Up |
| Lax1 | 1.870 | 0,704067557 | 0,1127668 | 6,62E-12 | 3,05E-10 | Up |
| Btg2 | 1.120 | 1 | 0,1597794 | 4,77E-15 | 3,68E-13 | Up |
| Kdm5b | 7.808 | -0,591051272 | 0,12314046 | 1,88E-08 | 4,51E-07 | Down |
| Lgr6 | 192 | 1 | 0,15709965 | 1,02E-12 | 5,59E-11 | Up |
| 5730559C18Rik | 313 | 1 | 0,91932776 | 0,00128948 | 0,00758507 | Up |
| Rgs2 | 4.395 | 0,885829419 | 0,11586018 | 6,00E-16 | 5,28E-14 | Up |
| Rgs1 | 89 | 2 | 0,32293477 | 6,73E-14 | 4,52E-12 | Up |
| Nmnat2 | 45 | 0,622221722 | 0,36599806 | 0,00136942 | 0,00796569 | Up |
| Lamc2 | 3 | -2 | 2 | 0,0005059 | 0,00347264 | Down |
| BC034090 | 129 | -1 | 0,2451988 | 7,61E-09 | 1,98E-07 | Down |
| Axdnd1 | 17 | 0,708668658 | 0,56239235 | 0,00264131 | 0,0137274 | Up |
| Ankrd45 | 19 | -2 | 0,84052335 | 0,00013109 | 0,00108923 | Down |
| Tnfsf4 | 370 | -0,937138511 | 0,2318021 | 5,70E-07 | 9,63E-06 | Down |
| Sele | 43 | 1 | 0,504774 | 7,45E-05 | 0,00067261 | Up |
| Sell | 30.737 | 2 | 0,17256276 | 3,38E-20 | 5,66E-18 | Up |
| Selp | 1.150 | 0,660454292 | 0,22520315 | 3,87E-05 | 0,00038182 | Up |
| Gm16587 | 15 | -2 | 1 | 0,00053206 | 0,00361751 | Down |
| Ccdc181 | 1.558 | 0,614581246 | 0,1487082 | 4,24E-07 | 7,40E-06 | Up |
| Nme7 | 498 | 0,974240905 | 0,20156763 | 1,89E-08 | 4,51E-07 | Up |
| Tbx19 | 183 | 1 | 0,23778146 | 4,68E-09 | 1,27E-07 | Up |
| Gm7694 | 1.519 | 0,61011079 | 0,16830544 | 3,53E-06 | 4,76E-05 | Up |
| Nos1ap | 689 | 0,702383431 | 0,16421951 | 2,25E-07 | 4,14E-06 | Up |
| Fcrlb | 8 | -3 | 1 | 3,66E-05 | 0,00036399 | Down |
| Fcgr2b | 3.342 | -1 | 0,34763368 | 8,00E-06 | 9,84E-05 | Down |
| Fcgr4 | 33 | -3 | 0,62345309 | 7,12E-09 | 1,86E-07 | Down |
| Fcgr3 | 445 | -4 | 0,49839067 | 1,81E-19 | 2,68E-17 | Down |
| Ly9 | 35 | 0,820418402 | 0,52686636 | 0,00105291 | 0,00642146 | Up |
| Slamf7 | 17 | 0,609900979 | 0,62821001 | 0,00610461 | 0,02716318 | Up |
| A630035G10Rik | 48 | 0,714824144 | 0,39295394 | 0,00075814 | 0,00486857 | Up |
| Slamf1 | 81 | -4 | 0,46921222 | 5,61E-16 | 5,04E-14 | Down |
| Gm37065 | 27 | -0,832551053 | 1 | 0,00423019 | 0,02014642 | Down |
| Igsf8 | 11.645 | -1 | 0,13694748 | 3,77E-18 | 4,50E-16 | Down |
| Slamf9 | 17 | -3 | 0,75377397 | 9,93E-08 | 1,99E-06 | Down |
| Ifi207 | 8 | -5 | 2 | 2,33E-06 | 3,31E-05 | Down |
| Ifi204 | 148 | 0,751862146 | 0,26844256 | 4,87E-05 | 0,00046678 | Up |
| Mndal | 929 | 0,630225416 | 0,21973074 | 5,15E-05 | 0,00049004 | Up |
| Ifi203 | 624 | 0,96347644 | 0,23750998 | 5,26E-07 | 8,97E-06 | Up |
| Spta1 | 352 | 5 | 0,31442777 | 1,57E-54 | 3,74E-51 | Up |
| Olfr417 | 8 | 0,922239993 | 1 | 0,00319074 | 0,0160063 | Up |
| Olfr248 | 16 | 2 | 0,80850482 | 6,55E-05 | 0,00060004 | Up |
| Olfr414 | 17 | 3 | 0,98637482 | 3,02E-06 | 4,15E-05 | Up |
| Cnih3 | 28 | -0,806911116 | 0,45538849 | 0,00066236 | 0,00436232 | Down |
| Enah | 46 | -3 | 0,69223627 | 1,10E-06 | 1,72E-05 | Down |

| | | | | | | |
|---|---|---|---|---|---|---|
| Degs1 | 9.673 | -1 | 0,11677744 | 8,44E-21 | 1,52E-18 | Down |
| 1700112H15Rik | 12 | -0,90418662 | 0,9847294 | 0,00319702 | 0,016033 | Down |
| Gm37800 | 94 | -2 | 0,32202622 | 6,58E-11 | 2,54E-09 | Down |
| Hsd11b1 | 62 | -2 | 0,48981491 | 1,49E-06 | 2,24E-05 | Down |
| Echdc3 | 49 | 1 | 0,30581894 | 4,85E-06 | 6,29E-05 | Up |
| Gm13292 | 42 | -2 | 0,81403321 | 2,16E-05 | 0,0002314 | Down |
| Tmem236 | 7 | -0,714321991 | 0,87639098 | 0,00570227 | 0,02557693 | Down |
| Mrc1 | 96 | -1 | 0,43308414 | 6,93E-05 | 0,00063159 | Down |
| Skida1 | 319 | 0,643999636 | 0,24129393 | 9,34E-05 | 0,00081761 | Up |
| Il1f9 | 11 | -2 | 2 | 0,00051931 | 0,00354866 | Down |
| Slc34a3 | 165 | 5 | 0,48740879 | 1,16E-25 | 3,41E-23 | Up |
| Ptgds | 5 | 2 | 1 | 0,0004149 | 0,00293091 | Up |
| Qsox2 | 6.294 | -0,636621432 | 0,06028384 | 6,46E-28 | 2,40E-25 | Down |
| Fam69b | 4.178 | -0,70906049 | 0,09337388 | 5,64E-16 | 5,05E-14 | Down |
| Lcn4 | 259 | 0,620413341 | 0,40595181 | 0,00201776 | 0,01099779 | Up |
| Slc2a6 | 1.862 | -2 | 0,25866668 | 5,85E-11 | 2,27E-09 | Down |
| Gm13398 | 9 | -1 | 0,93646443 | 0,00170622 | 0,00957532 | Down |
| Sardh | 13 | 2 | 0,68869358 | 0,0001104 | 0,00094361 | Up |
| Spaca9 | 236 | 1 | 0,20996846 | 5,90E-12 | 2,76E-10 | Up |
| Ak8 | 48 | 0,898457146 | 0,47106819 | 0,00039513 | 0,00281502 | Up |
| Zer1 | 3.262 | 0,730361575 | 0,10984239 | 5,29E-13 | 3,08E-11 | Up |
| Hmcn2 | 28 | -2 | 0,65316223 | 1,59E-05 | 0,00017745 | Down |
| Lcn2 | 1.443 | -4 | 0,6424906 | 7,03E-10 | 2,25E-08 | Down |
| Ak1 | 3.729 | -0,660680669 | 0,14177936 | 3,78E-08 | 8,45E-07 | Down |
| Ttc16 | 14 | -0,596763724 | 0,66281849 | 0,00713944 | 0,03085705 | Down |
| C230014O12Rik | 100 | 3 | 0,43217619 | 9,11E-11 | 3,43E-09 | Up |
| Gsn | 13.490 | -0,656902708 | 0,36320776 | 0,00093699 | 0,00581405 | Down |
| Stom | 447 | -1 | 0,87117031 | 0,00046827 | 0,00325164 | Down |
| Gm13496 | 34 | -0,782178731 | 0,41619272 | 0,00054511 | 0,00369179 | Down |
| Fmnl2 | 16 | -2 | 1 | 0,00011265 | 0,00095891 | Down |
| Nr4a2 | 1.421 | 0,939352473 | 0,14312741 | 1,21E-12 | 6,57E-11 | Up |
| Tanc1 | 180 | -1 | 0,70506464 | 0,00081178 | 0,00516739 | Down |
| Galnt3 | 6.168 | -0,910112309 | 0,08678605 | 3,53E-27 | 1,23E-24 | Down |
| Ttc30b | 98 | 0,775831641 | 0,23635095 | 1,01E-05 | 0,00012028 | Up |
| Plekha3 | 4.719 | -0,723970048 | 0,09087441 | 3,28E-17 | 3,49E-15 | Down |
| Tfpi | 172 | -2 | 0,27583838 | 7,81E-15 | 5,94E-13 | Down |
| Prdm11 | 3.830 | 0,623767029 | 0,13348706 | 3,48E-08 | 7,85E-07 | Up |
| Gm13807 | 8 | -1 | 1 | 0,00127924 | 0,00753273 | Down |
| Cd82 | 3.037 | -2 | 0,20074017 | 2,33E-32 | 1,40E-29 | Down |
| D430041D05Rik | 10 | 0,605275092 | 0,74318021 | 0,00786171 | 0,03335845 | Up |
| Inafm2 | 7.460 | 0,606429718 | 0,12339355 | 1,04E-08 | 2,64E-07 | Up |
| Capn3 | 187 | 0,693271547 | 0,2293886 | 2,70E-05 | 0,00027999 | Up |
| Tgm5 | 11 | 3 | 1 | 1,45E-05 | 0,00016318 | Up |
| Ckmt1 | 1.389 | 1 | 0,16101931 | 3,42E-19 | 4,61E-17 | Up |

| | | | | | | |
|---|---|---|---|---|---|---|
| Gm14002 | 34 | 2 | 0,37801031 | 1,60E-08 | 3,89E-07 | Up |
| Fbn1 | 28 | -3 | 0,64906278 | 1,05E-08 | 2,65E-07 | Down |
| Atp8b4 | 4.651 | -0,728442079 | 0,12330276 | 5,56E-11 | 2,17E-09 | Down |
| Ttl | 208 | 0,726600733 | 0,15592519 | 4,08E-08 | 9,04E-07 | Up |
| Slc20a1 | 10.385 | -0,7422249 | 0,08940806 | 2,32E-18 | 2,84E-16 | Down |
| Il1b | 14 | -3 | 1 | 2,73E-05 | 0,00028208 | Down |
| Ap5s1 | 565 | 1 | 0,16143222 | 2,61E-18 | 3,16E-16 | Up |
| Macrod2 | 256 | 0,842775571 | 0,22529189 | 1,91E-06 | 2,77E-05 | Up |
| Cd93 | 363 | 6 | 2 | 1,05E-06 | 1,66E-05 | Up |
| Trib3 | 5.838 | -1 | 0,57251312 | 7,96E-05 | 0,0007118 | Down |
| H13 | 23.153 | -0,698095149 | 0,16489432 | 2,71E-07 | 4,90E-06 | Down |
| Id1 | 13.343 | -2 | 0,13664015 | 2,07E-35 | 1,82E-32 | Down |
| Bcl2l1 | 12.016 | -0,906328598 | 0,10422527 | 1,09E-19 | 1,65E-17 | Down |
| Mylk2 | 5 | -0,926475779 | 1 | 0,00333733 | 0,01657768 | Down |
| Hck | 113 | -2 | 0,23100162 | 7,65E-13 | 4,33E-11 | Down |
| Nol4l | 6.164 | 1 | 0,11544741 | 9,26E-22 | 1,91E-19 | Up |
| Gm21850 | 112 | -0,679657922 | 0,71358322 | 0,00511975 | 0,02344802 | Down |
| Pxmp4 | 2.669 | 0,665418936 | 0,19012383 | 5,27E-06 | 6,78E-05 | Up |
| Ggt7 | 102 | 1 | 0,44171917 | 2,37E-05 | 0,0002494 | Up |
| Sla2 | 10.507 | 1 | 0,1704675 | 3,74E-13 | 2,24E-11 | Up |
| Mroh8 | 32 | 0,783709289 | 0,42638408 | 0,0005995 | 0,00400824 | Up |
| Src | 23 | 0,623603548 | 0,42493246 | 0,00226776 | 0,01210173 | Up |
| Tgm2 | 10.822 | 2 | 0,32216118 | 3,32E-09 | 9,30E-08 | Up |
| D630003M21Rik | 639 | 1 | 0,24370951 | 4,08E-08 | 9,04E-07 | Up |
| Ppp1r16b | 162 | -0,60370214 | 0,27007063 | 0,00037948 | 0,0027209 | Down |
| Wfdc12 | 53 | 2 | 0,37742751 | 4,33E-08 | 9,49E-07 | Up |
| Svs4 | 23 | 2 | 0,75680946 | 1,66E-05 | 0,00018396 | Up |
| Slpi | 1.214 | -2 | 0,46738831 | 2,43E-06 | 3,42E-05 | Down |
| Mmp9 | 25 | -3 | 0,76800928 | 7,93E-06 | 9,75E-05 | Down |
| Sulf2 | 19 | 0,617870384 | 0,52953502 | 0,00419569 | 0,02001566 | Up |
| Gm11469 | 11 | -2 | 1 | 0,00035443 | 0,0025688 | Down |
| Gm14267 | 11 | -2 | 1 | 5,40E-05 | 0,00050935 | Down |
| Gm11473 | 149 | -1 | 0,24941488 | 1,37E-09 | 4,13E-08 | Down |
| Atp9a | 443 | 2 | 0,19118169 | 1,08E-28 | 4,41E-26 | Up |
| Ctsz | 32.777 | -0,605078118 | 0,09781135 | 7,25E-12 | 3,33E-10 | Down |
| Zfp831 | 680 | 0,941317587 | 0,20648877 | 6,78E-08 | 1,42E-06 | Up |
| Edn3 | 84 | 2 | 0,23070276 | 1,15E-17 | 1,31E-15 | Up |
| Gm14393 | 181 | -1 | 0,22357422 | 4,84E-09 | 1,31E-07 | Down |
| Gm14439 | 116 | -0,934144463 | 0,27136762 | 4,93E-06 | 6,38E-05 | Down |
| Gm14387 | 54 | -1 | 0,4503015 | 9,53E-05 | 0,00083044 | Down |
| Lama5 | 288 | -0,594274355 | 0,31628456 | 0,00100011 | 0,00615099 | Down |
| Col9a3 | 254 | 0,963712566 | 0,29470943 | 8,32E-06 | 0,00010188 | Up |
| Slc17a9 | 2.603 | -0,631301806 | 0,16390864 | 1,38E-06 | 2,09E-05 | Down |
| Pde7a | 6.155 | -0,680134558 | 0,07948155 | 1,94E-19 | 2,82E-17 | Down |

| | | | | | | |
|---|---|---|---|---|---|---|
| Cp | 260 | -3 | 0,68846468 | 5,23E-07 | 8,93E-06 | Down |
| 7530428D23Rik | 4 | -0,600335664 | 0,8950329 | 0,00902782 | 0,03728545 | Down |
| Gm2011 | 66 | 0,99143505 | 0,23052247 | 1,92E-07 | 3,61E-06 | Up |
| 5031434O11Rik | 165 | 0,755624162 | 0,17621939 | 2,22E-07 | 4,09E-06 | Up |
| Mgst2 | 3.910 | 0,604050811 | 0,21430559 | 6,52E-05 | 0,00059847 | Up |
| Gm37261 | 10 | 0,879285529 | 0,94623845 | 0,00332016 | 0,016512 | Up |
| Maml3 | 4.177 | 1 | 0,11803107 | 2,10E-19 | 2,98E-17 | Up |
| Gm37170 | 8 | 0,792634235 | 0,91868308 | 0,00440798 | 0,02081478 | Up |
| Wwtr1 | 82 | -3 | 0,42652421 | 4,01E-17 | 4,23E-15 | Down |
| Rnf13 | 3.587 | -0,823833528 | 0,12582186 | 1,35E-12 | 7,16E-11 | Down |
| P2ry14 | 6.297 | -2 | 0,19792509 | 1,36E-16 | 1,34E-14 | Down |
| P2ry13 | 20 | -0,583892662 | 0,55859755 | 0,00587632 | 0,02627308 | Down |
| Gm37696 | 94 | -0,583537323 | 0,29820918 | 0,00086063 | 0,00541862 | Down |
| Gm5540 | 29 | 0,769553952 | 0,45059265 | 0,00084395 | 0,00533341 | Up |
| Tlr2 | 2.739 | -0,767300354 | 0,13505249 | 2,29E-10 | 7,96E-09 | Down |
| Lmna | 40 | -1 | 0,607634 | 0,00022453 | 0,00172779 | Down |
| Rab13 | 172 | 2 | 0,68932539 | 3,98E-05 | 0,00039042 | Up |
| Npr1 | 165 | 2 | 0,30972482 | 1,06E-15 | 8,95E-14 | Up |
| S100a9 | 1.176 | -4 | 2 | 1,75E-05 | 0,00019232 | Down |
| S100a11 | 5.621 | -0,656058922 | 0,17910868 | 2,86E-06 | 3,95E-05 | Down |
| Selenbp1 | 1.431 | 0,676522226 | 0,28006117 | 0,0001804 | 0,00143417 | Up |
| Ctss | 664 | -1 | 0,41473754 | 6,63E-06 | 8,33E-05 | Down |
| Fcgr1 | 5.289 | -1 | 0,14978491 | 2,01E-14 | 1,43E-12 | Down |
| Gm5544 | 8 | -0,748161371 | 0,85779803 | 0,00485795 | 0,02252018 | Down |
| Slc16a1 | 34.988 | -0,585936103 | 0,11308506 | 2,60E-09 | 7,40E-08 | Down |
| Cd53 | 2.648 | -3 | 0,52785805 | 9,98E-10 | 3,09E-08 | Down |
| Prok1 | 7 | -1 | 1 | 0,00101473 | 0,00623177 | Down |
| Csf1 | 1.975 | -1 | 0,23071025 | 1,33E-08 | 3,27E-07 | Down |
| Col11a1 | 20 | 3 | 1 | 3,57E-05 | 0,00035654 | Up |
| Slc44a3 | 5 | 2 | 2 | 0,00083303 | 0,00528061 | Up |
| Gm42776 | 17 | -3 | 0,86146946 | 2,74E-06 | 3,81E-05 | Down |
| 5730508B09Rik | 698 | -0,672493991 | 0,14611808 | 5,02E-08 | 1,08E-06 | Down |
| Gm43254 | 229 | 0,587186735 | 0,21547061 | 9,33E-05 | 0,00081747 | Up |
| Gm43194 | 7 | -0,772068567 | 1 | 0,00512883 | 0,02346981 | Down |
| Dapp1 | 12.758 | 0,692630508 | 0,09187849 | 7,95E-16 | 6,94E-14 | Up |
| Gbp5 | 10 | -0,927098028 | 0,96142496 | 0,00286411 | 0,01469823 | Down |
| Gbp3 | 77 | -0,654407113 | 0,43886851 | 0,00193309 | 0,01062753 | Down |
| 4930519L02Rik | 1.669 | -1 | 0,4548507 | 6,47E-05 | 0,00059493 | Down |
| Ddah1 | 12 | -2 | 0,67793221 | 5,00E-05 | 0,00047735 | Down |
| Dnajb4 | 3.052 | 0,846145902 | 0,15435136 | 7,74E-10 | 2,45E-08 | Up |
| Cth | 7 | -2 | 2 | 0,00040336 | 0,0028639 | Down |
| Tmem68 | 1.813 | -0,585950981 | 0,12156227 | 1,71E-08 | 4,12E-07 | Down |
| A830012C17Rik | 10 | -6 | 2 | 2,43E-07 | 4,43E-06 | Down |
| Chd7 | 176 | -1 | 0,31482493 | 7,02E-08 | 1,46E-06 | Down |

| | | | | | | |
|---|---|---|---|---|---|---|
| Gem | 255 | -7 | 1 | 2,61E-09 | 7,42E-08 | Down |
| Cdh17 | 138 | -6 | 0,96507353 | 3,74E-13 | 2,24E-11 | Down |
| Pdp1 | 2.184 | -2 | 0,22344455 | 1,94E-16 | 1,89E-14 | Down |
| Osgin2 | 3.112 | -0,731801554 | 0,10420673 | 4,12E-14 | 2,86E-12 | Down |
| Epha7 | 24 | -0,853410856 | 0,66485701 | 0,00179351 | 0,00998813 | Down |
| Gm11934 | 122 | -0,681795179 | 0,33807733 | 0,00051763 | 0,00354231 | Down |
| Cfap206 | 10 | 3 | 0,93571045 | 1,81E-05 | 0,00019844 | Up |
| AI464131 | 14 | -1 | 0,85117196 | 0,00032932 | 0,00241085 | Down |
| Dnajb5 | 334 | -0,737448851 | 0,25178911 | 3,37E-05 | 0,00033943 | Down |
| Unc13b | 39 | 0,779939062 | 0,50911309 | 0,00122536 | 0,00728235 | Up |
| Atp8b5 | 129 | 1 | 0,29219914 | 1,56E-06 | 2,33E-05 | Up |
| Cd72 | 402 | 0,818701692 | 0,16031313 | 5,29E-09 | 1,42E-07 | Up |
| Reck | 212 | -0,611931004 | 0,14261059 | 2,10E-07 | 3,89E-06 | Down |
| Glipr2 | 6.208 | -1 | 0,23478158 | 3,02E-09 | 8,52E-08 | Down |
| Fbxo10 | 1.628 | 0,667089154 | 0,09170536 | 5,10E-15 | 3,92E-13 | Up |
| Tnfsf8 | 2.094 | -2 | 0,12921243 | 9,07E-55 | 2,53E-51 | Down |
| Tlr4 | 3.139 | -0,997343112 | 0,18592659 | 1,30E-09 | 3,94E-08 | Down |
| Elavl2 | 987 | -1 | 0,11739134 | 7,03E-23 | 1,57E-20 | Down |
| Nfia | 299 | 0,897259369 | 0,19556181 | 6,16E-08 | 1,30E-06 | Up |
| Dock7 | 11 | -0,912461696 | 0,8267327 | 0,00231933 | 0,01233756 | Down |
| Lepr | 34 | -0,683776966 | 0,49049306 | 0,00215595 | 0,01160125 | Down |
| Pde4b | 1.526 | 0,977695173 | 0,13097417 | 2,00E-15 | 1,64E-13 | Up |
| Acot11 | 38 | 0,593962505 | 0,47484475 | 0,00390629 | 0,0188663 | Up |
| Lrp8 | 9.689 | -0,594359769 | 0,08581915 | 5,07E-14 | 3,47E-12 | Down |
| Cdkn2c | 6.893 | 1 | 0,09111111 | 1,94E-35 | 1,81E-32 | Up |
| Mob3c | 849 | -0,825892495 | 0,11799199 | 6,45E-14 | 4,35E-12 | Down |
| Tmem53 | 5 | -0,819948835 | 1 | 0,00444185 | 0,02093336 | Down |
| Olfr1342 | 762 | 0,775050143 | 0,22687961 | 6,41E-06 | 8,07E-05 | Up |
| Slc2a1 | 11.722 | -0,704070391 | 0,12352297 | 1,74E-10 | 6,19E-09 | Down |
| Rimkla | 286 | -1 | 0,22046246 | 1,36E-10 | 4,96E-09 | Down |
| Rims3 | 830 | -0,897455421 | 0,1439223 | 1,01E-11 | 4,55E-10 | Down |
| Csf3r | 399 | -2 | 0,40957514 | 4,18E-08 | 9,20E-07 | Down |
| Gjb3 | 19 | -2 | 0,59120155 | 1,17E-05 | 0,00013573 | Down |
| Tlr12 | 147 | 2 | 0,25609797 | 2,90E-12 | 1,47E-10 | Up |
| Spocd1 | 103 | 2 | 0,34379842 | 7,07E-09 | 1,86E-07 | Up |
| 1700003M07Rik | 6 | -0,895703923 | 1 | 0,00339219 | 0,01679535 | Down |
| Serinc2 | 1.356 | -0,867185352 | 0,22414141 | 1,17E-06 | 1,82E-05 | Down |
| Selenon | 8.822 | -1 | 0,08078082 | 3,89E-55 | 1,63E-51 | Down |
| Man1c1 | 2.247 | -2 | 0,20211017 | 3,82E-34 | 3,04E-31 | Down |
| Cnr2 | 639 | -1 | 0,24439208 | 6,38E-09 | 1,68E-07 | Down |
| Pla2g2e | 34 | -2 | 0,83538407 | 4,63E-05 | 0,00044565 | Down |
| Iffo2 | 56 | -0,590511632 | 0,32512996 | 0,00118264 | 0,00708613 | Down |
| Rcc2 | 141.539 | 0,608514901 | 0,07999837 | 3,40E-16 | 3,20E-14 | Up |
| Padi6 | 32 | 0,882557395 | 0,48848005 | 0,00051831 | 0,00354327 | Up |

| | | | | | | |
|---|---|---|---|---|---|---|
| Crocc | 1.850 | 0,596048886 | 0,12940623 | 4,87E-08 | 1,05E-06 | Up |
| Tmem51 | 127 | 2 | 0,66860168 | 4,15E-06 | 5,51E-05 | Up |
| Gm13053 | 20 | 2 | 0,51826757 | 1,24E-06 | 1,91E-05 | Up |
| Kazn | 3.609 | 0,904770135 | 0,13433221 | 4,05E-13 | 2,41E-11 | Up |
| Casz1 | 137 | -2 | 0,28934211 | 2,47E-09 | 7,08E-08 | Down |
| Pik3cd | 36.481 | -0,585871064 | 0,09997239 | 5,38E-11 | 2,10E-09 | Down |
| Slc25a33 | 1.909 | -0,899557744 | 0,10165827 | 2,87E-20 | 4,91E-18 | Down |
| Car6 | 5 | -1 | 2 | 0,00199686 | 0,01091194 | Down |
| Gm13091 | 135 | 0,840731074 | 0,40096634 | 0,00027846 | 0,00207757 | Up |
| Isg15 | 35 | -1 | 0,54806441 | 0,00039816 | 0,00283298 | Down |
| Gm26825 | 145 | -0,745800044 | 0,25451143 | 3,31E-05 | 0,00033408 | Down |
| Abcb1b | 4.150 | -1 | 0,11400969 | 1,19E-26 | 3,83E-24 | Down |
| A630072M18Rik | 81 | 0,74436177 | 0,25326447 | 3,23E-05 | 0,00032725 | Up |
| Napepld | 977 | 0,669127185 | 0,09087644 | 2,67E-15 | 2,14E-13 | Up |
| Kcnh2 | 16 | -2 | 0,88545682 | 0,00010973 | 0,0009398 | Down |
| Nos3 | 60 | -2 | 0,27510008 | 1,94E-14 | 1,39E-12 | Down |
| Actr3b | 15 | 6 | 2 | 7,26E-07 | 1,20E-05 | Up |
| Khk | 615 | 2 | 0,13844952 | 5,03E-31 | 2,55E-28 | Up |
| AI839979 | 54 | -2 | 0,55424001 | 1,29E-05 | 0,00014758 | Down |
| Fosl2 | 6.754 | -0,735146326 | 0,16387691 | 9,16E-08 | 1,85E-06 | Down |
| Plb1 | 17 | -0,74763655 | 0,61196984 | 0,00257846 | 0,01345931 | Down |
| Gm15614 | 21 | 2 | 0,60278343 | 2,78E-06 | 3,86E-05 | Up |
| Sh3bp2 | 2.111 | -0,676154582 | 0,07972568 | 3,65E-19 | 4,87E-17 | Down |
| Rgs12 | 21 | -2 | 0,59835096 | 1,00E-06 | 1,59E-05 | Down |
| Sh3tc1 | 913 | -1 | 0,16017733 | 2,44E-12 | 1,25E-10 | Down |
| Ppp2r2c | 9 | 0,690420983 | 0,73911483 | 0,00512733 | 0,02346981 | Up |
| Bst1 | 34 | -6 | 1 | 5,16E-08 | 1,10E-06 | Down |
| Prom1 | 9 | -6 | 2 | 2,81E-07 | 5,07E-06 | Down |
| N4bp2os | 74 | 0,696219954 | 0,28706846 | 0,00016622 | 0,00133545 | Up |
| 9130230L23Rik | 38 | -1 | 0,47348257 | 0,00013874 | 0,00114487 | Down |
| Cnga1 | 150 | 1 | 0,30355813 | 1,54E-08 | 3,76E-07 | Up |
| Gm43417 | 10 | -0,656359133 | 0,7319252 | 0,00598338 | 0,02672322 | Down |
| Stap1 | 5 | -0,714502817 | 1 | 0,00612954 | 0,02720898 | Down |
| 5830473C10Rik | 3.009 | -0,713413962 | 0,09650693 | 2,62E-15 | 2,11E-13 | Down |
| Ppbp | 98 | -4 | 0,93212698 | 1,72E-08 | 4,14E-07 | Down |
| Pf4 | 353 | -2 | 0,33207149 | 8,94E-13 | 4,99E-11 | Down |
| Cxcl3 | 15 | -3 | 2 | 0,00012067 | 0,00101732 | Down |
| Cxcl2 | 58 | -4 | 0,70151798 | 3,32E-10 | 1,12E-08 | Down |
| Mthfd2l | 1.151 | -0,859764928 | 0,10028127 | 3,22E-19 | 4,39E-17 | Down |
| Anxa3 | 8.981 | -1 | 0,28830576 | 1,94E-08 | 4,63E-07 | Down |
| Agpat9 | 20 | -2 | 0,9125543 | 0,00035266 | 0,00255819 | Down |
| Spp1 | 14.166 | -2 | 0,20221369 | 3,30E-25 | 9,05E-23 | Down |
| Gm42518 | 7 | 0,999390472 | 0,93341574 | 0,00211599 | 0,01143398 | Up |
| Gm42517 | 917 | 2 | 0,14343467 | 5,76E-31 | 2,83E-28 | Up |

| | | | | | | |
|---|---|---|---|---|---|---|
| Gm10419 | 2.402 | 1 | 0,10626168 | 6,76E-31 | 3,23E-28 | Up |
| Gm18555 | 102 | 0,764573627 | 0,32583849 | 0,00017352 | 0,00138606 | Up |
| Ulk1 | 5.417 | 0,66024218 | 0,10712464 | 9,46E-12 | 4,27E-10 | Up |
| Ttc28 | 1.355 | 0,65173357 | 0,16943572 | 1,38E-06 | 2,09E-05 | Up |
| Tmem119 | 172 | -0,665976511 | 0,18014816 | 2,48E-06 | 3,48E-05 | Down |
| Ssh1 | 469 | 0,687906519 | 0,26001335 | 8,94E-05 | 0,00078663 | Up |
| Svop | 53 | -0,982859254 | 0,35495134 | 3,64E-05 | 0,00036215 | Down |
| Acacb | 858 | 0,61345136 | 0,19099688 | 1,66E-05 | 0,00018366 | Up |
| Nos1 | 525 | 1 | 0,25601477 | 1,68E-09 | 4,97E-08 | Up |
| Sdsl | 2.168 | 0,667852366 | 0,10718594 | 6,34E-12 | 2,94E-10 | Up |
| Oas2 | 154 | -2 | 0,36222186 | 8,58E-08 | 1,75E-06 | Down |
| Oas3 | 342 | -2 | 0,51548008 | 4,59E-06 | 5,99E-05 | Down |
| P2rx7 | 33 | -2 | 0,83562072 | 0,00021607 | 0,00167714 | Down |
| Tmem120b | 910 | -0,75472511 | 0,12600728 | 3,70E-11 | 1,50E-09 | Down |
| Hcar2 | 29 | -4 | 0,6391225 | 3,21E-10 | 1,09E-08 | Down |
| Clip2 | 3.968 | 0,90722998 | 0,07992845 | 2,74E-31 | 1,48E-28 | Up |
| Syna | 6 | 0,942499773 | 1 | 0,00321538 | 0,01610576 | Up |
| Rfc2 | 25.282 | 0,863771497 | 0,06773211 | 1,23E-38 | 1,37E-35 | Up |
| Fkbp6 | 272 | 0,624578858 | 0,14046104 | 1,02E-07 | 2,04E-06 | Up |
| Cux1 | 58.335 | 0,97407876 | 0,12237984 | 4,40E-17 | 4,60E-15 | Up |
| Gm38082 | 62 | -0,583108354 | 0,49769648 | 0,00471477 | 0,0219616 | Down |
| Myl10 | 156 | 4 | 0,45614026 | 7,05E-19 | 9,15E-17 | Up |
| Tfr2 | 6.268 | 2 | 0,19928791 | 6,72E-23 | 1,52E-20 | Up |
| Zcwpw1 | 1.246 | 0,927997837 | 0,15278241 | 2,64E-11 | 1,09E-09 | Up |
| Pilra | 72 | -3 | 0,36227915 | 1,96E-18 | 2,43E-16 | Down |
| Pilrb1 | 242 | -3 | 0,2801483 | 1,60E-27 | 5,69E-25 | Down |
| Gm36551 | 79 | -3 | 0,28879471 | 9,09E-20 | 1,41E-17 | Down |
| Pilrb2 | 19 | -3 | 0,81011699 | 7,66E-06 | 9,45E-05 | Down |
| Zscan21 | 3.787 | 0,678929528 | 0,07374039 | 5,76E-22 | 1,20E-19 | Up |
| Nxpe5 | 888 | -2 | 0,72355054 | 3,28E-05 | 0,00033162 | Down |
| BC037034 | 9.529 | -0,857848849 | 0,20863958 | 4,62E-07 | 7,98E-06 | Down |
| Prkar1b | 86 | 1 | 0,22076718 | 2,13E-09 | 6,20E-08 | Up |
| Micall2 | 18.999 | 0,802517968 | 0,09932304 | 1,80E-17 | 1,99E-15 | Up |
| Lfng | 13.135 | -0,633338797 | 0,08127915 | 8,63E-17 | 8,80E-15 | Down |
| Mtus2 | 475 | -5 | 1 | 7,87E-09 | 2,04E-07 | Down |
| 2210417A02Rik | 10 | 0,631976895 | 1 | 0,00406287 | 0,01948757 | Up |
| Fry | 10 | 1 | 0,84487075 | 0,00167745 | 0,00944556 | Up |
| 1700028E10Rik | 103 | 0,903537062 | 0,22027365 | 4,66E-07 | 8,05E-06 | Up |
| 2610001J05Rik | 9.643 | -0,704329562 | 0,11248187 | 5,92E-12 | 2,76E-10 | Down |
| Met | 4.940 | -0,618295063 | 0,16909173 | 3,07E-06 | 4,19E-05 | Down |
| St7 | 4.210 | 0,653815937 | 0,09014422 | 5,65E-15 | 4,32E-13 | Up |
| Wnt2 | 11 | 1 | 0,85599305 | 0,00072583 | 0,00469894 | Up |
| Asz1 | 351 | 3 | 0,33497753 | 4,24E-15 | 3,31E-13 | Up |
| Gm20756 | 5 | 2 | 2 | 0,00050094 | 0,00344538 | Up |

| | | | | | | |
|---|---|---|---|---|---|---|
| Flnc | 19 | -1 | 0,75110567 | 0,000208 | 0,00162658 | Down |
| Smo | 355 | 0,915942752 | 0,71725469 | 0,00162122 | 0,00916902 | Up |
| Akr1b7 | 9 | -2 | 2 | 0,00026047 | 0,00195994 | Down |
| Kdm7a | 753 | -0,815251758 | 0,11671189 | 6,99E-14 | 4,66E-12 | Down |
| Slc37a3 | 345 | -0,596105423 | 0,3725664 | 0,00189967 | 0,01046101 | Down |
| Rab19 | 14 | -2 | 1 | 0,00033144 | 0,00242317 | Down |
| Gm10244 | 14 | 0,922385705 | 0,67208429 | 0,00130443 | 0,00765413 | Up |
| Dennd2a | 730 | 1 | 0,28712902 | 7,00E-08 | 1,46E-06 | Up |
| 8030453O22Rik | 369 | 1 | 0,26324223 | 3,09E-08 | 7,01E-07 | Up |
| Clec5a | 2.771 | -2 | 0,32913908 | 4,56E-15 | 3,55E-13 | Down |
| Mgam | 42 | -3 | 0,45911134 | 2,13E-09 | 6,20E-08 | Down |
| Gm43889 | 34 | -3 | 0,71323862 | 1,63E-06 | 2,43E-05 | Down |
| A930035D04Rik | 54 | 0,620756491 | 0,30684317 | 0,00062789 | 0,00416475 | Up |
| Pdia4 | 29.605 | -0,592231623 | 0,14794824 | 7,70E-07 | 1,26E-05 | Down |
| Krba1 | 8 | 1 | 1 | 0,00133356 | 0,00778958 | Up |
| Sspo | 7 | -0,970390204 | 1 | 0,00290038 | 0,01483886 | Down |
| Gimap4 | 1.437 | 0,714643744 | 0,1344415 | 1,50E-09 | 4,48E-08 | Up |
| Gimap6 | 108 | 1 | 0,32053154 | 3,03E-06 | 4,15E-05 | Up |
| Gimap1 | 2.396 | 0,691556323 | 0,12288232 | 2,53E-10 | 8,71E-09 | Up |
| Gimap5 | 1.384 | 1 | 0,16411571 | 7,33E-12 | 3,36E-10 | Up |
| Gm44226 | 119 | 0,583944117 | 0,26782282 | 0,00047668 | 0,00329806 | Up |
| 4833403J16Rik | 77 | 1 | 0,34499694 | 2,43E-05 | 0,00025573 | Up |
| Snx10 | 609 | -0,873828049 | 0,14748987 | 6,56E-11 | 2,54E-09 | Down |
| Nod1 | 107 | -0,921858506 | 0,31767341 | 2,71E-05 | 0,00028009 | Down |
| Ppp1r17 | 12 | -0,609965032 | 0,70672393 | 0,00723958 | 0,03120121 | Down |
| Hpgds | 22 | -2 | 0,65655944 | 6,33E-05 | 0,00058263 | Down |
| Gm6786 | 15 | -2 | 0,86544864 | 9,04E-05 | 0,00079415 | Down |
| Gm1070 | 49 | 2 | 0,46540899 | 7,88E-06 | 9,71E-05 | Up |
| Gm44186 | 18 | -0,604560436 | 0,59542182 | 0,00576036 | 0,02580291 | Down |
| Capg | 17.969 | -0,776953107 | 0,16289801 | 2,60E-08 | 6,03E-07 | Down |
| Tacr1 | 5 | -0,800418576 | 1 | 0,0041617 | 0,01988185 | Down |
| Hk2 | 19.649 | -0,812734269 | 0,10156158 | 3,44E-17 | 3,64E-15 | Down |
| Gm43878 | 56 | -2 | 0,37073151 | 2,16E-08 | 5,10E-07 | Down |
| Wdr54 | 121 | 0,707561372 | 0,45824715 | 0,00145334 | 0,00837046 | Up |
| Mthfd2 | 15.492 | -0,625045046 | 0,11778699 | 1,32E-09 | 4,01E-08 | Down |
| Sfxn5 | 233 | 0,915242653 | 0,16129921 | 2,70E-10 | 9,22E-09 | Up |
| Rab11fip5 | 155 | -2 | 0,41203202 | 6,63E-09 | 1,75E-07 | Down |
| 2310040G24Rik | 489 | 0,687822506 | 0,09302288 | 2,30E-15 | 1,86E-13 | Up |
| Asprv1 | 7 | -2 | 2 | 0,00068592 | 0,00448939 | Down |
| Gm44198 | 184 | -0,90880691 | 0,15263127 | 5,47E-11 | 2,13E-09 | Down |
| Gp9 | 602 | 0,918357097 | 0,28029032 | 8,63E-06 | 0,00010505 | Up |
| Gm26588 | 43 | -1 | 0,3200666 | 2,97E-06 | 4,09E-05 | Down |
| Mgll | 54 | -3 | 1 | 4,55E-05 | 0,00044006 | Down |
| Aldh1l1 | 5 | -1 | 1 | 0,00235665 | 0,01251621 | Down |

| | | | | | | |
|---|---|---|---|---|---|---|
| Frmd4b | 176 | 2 | 0,31475412 | 3,46E-15 | 2,73E-13 | Up |
| Mitf | 799 | -0,718165898 | 0,18820791 | 1,52E-06 | 2,28E-05 | Down |
| Bhlhe40 | 21.345 | -0,696838866 | 0,13153417 | 1,58E-09 | 4,69E-08 | Down |
| Srgap3 | 2.103 | 0,661984422 | 0,09863749 | 2,67E-13 | 1,64E-11 | Up |
| Il17re | 269 | 1 | 0,27178424 | 1,24E-06 | 1,91E-05 | Up |
| Creld1 | 8 | 0,89188888 | 0,83850054 | 0,0025966 | 0,01353062 | Up |
| Mkrn2os | 30 | 0,638348258 | 0,38043217 | 0,00135637 | 0,00790073 | Up |
| Cand2 | 1.624 | 2 | 0,22983246 | 1,47E-17 | 1,66E-15 | Up |
| Fxyd4 | 283 | -0,955183371 | 0,25293369 | 1,51E-06 | 2,27E-05 | Down |
| Cacna2d4 | 145 | 0,794128281 | 0,2259297 | 4,46E-06 | 5,85E-05 | Up |
| Lrtm2 | 28 | 0,676321962 | 0,53570601 | 0,00288723 | 0,01478967 | Up |
| B4galnt3 | 240 | -0,807428416 | 0,3674364 | 0,00023218 | 0,00177585 | Down |
| Tuba8 | 21.444 | -0,924523727 | 0,16608938 | 4,85E-10 | 1,60E-08 | Down |
| Usp18 | 164 | -1 | 0,23459834 | 2,65E-09 | 7,52E-08 | Down |
| Phc1 | 2.602 | 1 | 0,10480056 | 4,06E-24 | 1,00E-21 | Up |
| Slc2a3 | 28.352 | -0,910940576 | 0,1274699 | 2,35E-14 | 1,67E-12 | Down |
| Clec4n | 64 | -3 | 0,87104999 | 1,70E-06 | 2,52E-05 | Down |
| Clec4e | 2.451 | -1 | 0,2229177 | 3,04E-08 | 6,90E-07 | Down |
| C1ra | 714 | -1 | 0,27767202 | 1,40E-07 | 2,70E-06 | Down |
| Ptpn6 | 45.882 | 0,64859817 | 0,19279651 | 8,90E-06 | 0,00010786 | Up |
| Lag3 | 152 | 1 | 0,21817146 | 2,75E-12 | 1,40E-10 | Up |
| Ptms | 5.383 | 0,830023236 | 0,12645671 | 1,23E-12 | 6,62E-11 | Up |
| Fgf23 | 170 | -4 | 0,56493915 | 1,94E-15 | 1,59E-13 | Down |
| Ccnd2 | 58.882 | -0,6020692 | 0,07967281 | 4,91E-16 | 4,45E-14 | Down |
| Gm26770 | 160 | 1 | 0,21823851 | 1,86E-11 | 8,03E-10 | Up |
| Rhno1 | 8.720 | 1 | 0,1069361 | 3,26E-25 | 9,05E-23 | Up |
| Foxm1 | 28.002 | 0,653670169 | 0,08356012 | 7,35E-17 | 7,55E-15 | Up |
| Gm43965 | 94 | 1 | 0,33086486 | 1,36E-06 | 2,07E-05 | Up |
| Cd69 | 4.378 | -2 | 0,28817783 | 1,27E-16 | 1,26E-14 | Down |
| Clec12a | 1.081 | -3 | 0,71303681 | 2,48E-06 | 3,48E-05 | Down |
| Klra2 | 7 | -3 | 2 | 6,76E-05 | 0,0006183 | Down |
| Bcl2l14 | 2.146 | -2 | 0,14114623 | 5,90E-53 | 1,23E-49 | Down |
| Cdkn1b | 2.508 | 0,598666138 | 0,1486326 | 6,84E-07 | 1,13E-05 | Up |
| Emp1 | 7 | -1 | 1 | 0,00224237 | 0,01198814 | Down |
| Caprin2 | 239 | 0,663794128 | 0,17237988 | 1,35E-06 | 2,05E-05 | Up |
| Dennd5b | 69 | -0,955957265 | 0,49921392 | 0,00035112 | 0,00254817 | Down |
| Oscar | 78 | -1 | 0,30213017 | 6,77E-06 | 8,47E-05 | Down |
| Zfp580 | 1.036 | 0,712851423 | 0,12640191 | 2,51E-10 | 8,70E-09 | Up |
| Vmn1r65 | 33 | -0,590754502 | 0,41203018 | 0,00276119 | 0,01425747 | Down |
| Zfp78 | 80 | 1 | 0,27835452 | 2,45E-06 | 3,45E-05 | Up |
| Zfp773 | 110 | 0,637214936 | 0,22359839 | 5,37E-05 | 0,0005076 | Up |
| Pla2g4c | 248 | 2 | 0,35595877 | 3,84E-08 | 8,56E-07 | Up |
| Ehd2 | 211 | -2 | 0,31159792 | 3,04E-12 | 1,53E-10 | Down |
| C5ar1 | 81 | -4 | 0,36022524 | 4,10E-25 | 1,11E-22 | Down |

| | | | | | | |
|---|---|---|---|---|---|---|
| Gm45509 | 71 | -0,644679215 | 0,27873114 | 0,00026531 | 0,00199311 | Down |
| Slc1a5 | 12.791 | -0,640150854 | 0,08800275 | 4,59E-15 | 3,56E-13 | Down |
| Prkd2 | 8.148 | -0,783954217 | 0,09122354 | 2,32E-19 | 3,24E-17 | Down |
| Gng8 | 27 | -1 | 0,70522129 | 0,00053858 | 0,00365493 | Down |
| Ptgir | 1.602 | -2 | 0,20874609 | 3,26E-18 | 3,92E-16 | Down |
| Psg17 | 14 | 4 | 2 | 9,44E-06 | 0,00011346 | Up |
| Mill2 | 27 | 2 | 0,70911266 | 4,59E-05 | 0,00044303 | Up |
| Ccdc61 | 2.143 | 0,59045608 | 0,11581074 | 4,03E-09 | 1,11E-07 | Up |
| Dmwd | 5.860 | -0,769227597 | 0,0836521 | 1,02E-21 | 2,09E-19 | Down |
| Relb | 527 | -1 | 0,23199107 | 3,17E-10 | 1,08E-08 | Down |
| Pvr | 729 | -0,77001827 | 0,18651857 | 4,35E-07 | 7,57E-06 | Down |
| Plaur | 2.253 | -2 | 0,15684292 | 6,35E-27 | 2,17E-24 | Down |
| Cd177 | 3.314 | -0,735853236 | 0,22137606 | 9,18E-06 | 0,00011084 | Down |
| Ccdc97 | 5.506 | 0,599444372 | 0,08925697 | 2,19E-13 | 1,37E-11 | Up |
| Cyp2b10 | 21 | -2 | 0,94792114 | 4,88E-05 | 0,00046752 | Down |
| Ltbp4 | 29 | 1 | 0,50236489 | 9,15E-05 | 0,00080335 | Up |
| Pgam1-ps2 | 138 | -1 | 0,31382821 | 1,67E-06 | 2,47E-05 | Down |
| 9530053A07Rik | 88 | -3 | 0,40704008 | 3,88E-16 | 3,55E-14 | Down |
| Gm45010 | 81 | 0,637434748 | 0,22743407 | 6,24E-05 | 0,00057675 | Up |
| Zfp36 | 1.356 | -1 | 0,19101811 | 1,98E-16 | 1,91E-14 | Down |
| Rasgrp4 | 12 | -2 | 0,91694528 | 0,0001054 | 0,00090639 | Down |
| Zfp383 | 210 | 0,883082933 | 0,15409972 | 2,00E-10 | 7,02E-09 | Up |
| Tyrobp | 8.140 | -0,628718439 | 0,20090382 | 2,14E-05 | 0,00022937 | Down |
| Proser3 | 945 | 0,739607949 | 0,17176612 | 2,03E-07 | 3,79E-06 | Up |
| Zbtb32 | 139 | 1 | 0,23741567 | 1,34E-08 | 3,30E-07 | Up |
| Dmkn | 62 | -0,876986456 | 0,5379681 | 0,00079209 | 0,00506131 | Down |
| Gm2381 | 3 | -0,778176811 | 1 | 0,00423033 | 0,02014642 | Down |
| Siglecf | 2.157 | 2 | 0,31920191 | 1,35E-11 | 5,92E-10 | Up |
| Siglecg | 42 | 2 | 0,29831523 | 1,93E-11 | 8,29E-10 | Up |
| Cd33 | 318 | -2 | 0,59420289 | 1,12E-05 | 0,00013048 | Down |
| Klk1b11 | 5 | -1 | 1 | 0,00251068 | 0,01317951 | Down |
| Gm18905 | 840 | -0,692724424 | 0,29406163 | 0,00020389 | 0,00159816 | Down |
| Napsa | 27.704 | -0,659210745 | 0,12182988 | 7,88E-10 | 2,48E-08 | Down |
| Kcnc3 | 605 | -1 | 0,28581213 | 1,73E-06 | 2,55E-05 | Down |
| Gm44832 | 39 | -1 | 0,395359 | 1,47E-06 | 2,21E-05 | Down |
| Fut1 | 6 | -0,960790956 | 1 | 0,00308822 | 0,01558546 | Down |
| A030001D20Rik | 31 | 0,818816539 | 0,43531793 | 0,00049699 | 0,0034227 | Up |
| Fut2 | 16 | -1 | 0,73720164 | 0,00026535 | 0,00199311 | Down |
| Kcnj14 | 93 | -0,665339145 | 0,36730711 | 0,00090426 | 0,0056502 | Down |
| Saa3 | 231 | 0,702202084 | 0,19542785 | 3,59E-06 | 4,83E-05 | Up |
| Saa4 | 12 | -6 | 1 | 2,23E-08 | 5,24E-07 | Down |
| Ldhc | 154 | 0,697214711 | 0,24906908 | 5,47E-05 | 0,00051456 | Up |
| Gm7336 | 41 | -0,702412545 | 0,36370881 | 0,00060047 | 0,00401313 | Down |
| Gm45001 | 24 | 0,65709324 | 0,52865364 | 0,0031857 | 0,01599541 | Up |

| | | | | | | |
|---|---|---|---|---|---|---|
| Gm34121 | 19 | 0,584679499 | 0,42911425 | 0,00326481 | 0,01629969 | Up |
| 1810008I18Rik | 32 | -4 | 0,56098491 | 5,21E-13 | 3,04E-11 | Down |
| Selenos | 9.797 | -0,689306488 | 0,1770071 | 1,12E-06 | 1,75E-05 | Down |
| Mctp2 | 4 | 1 | 2 | 0,00099101 | 0,00610626 | Up |
| Akap13 | 34.519 | 1 | 0,12710672 | 4,03E-20 | 6,55E-18 | Up |
| Isg20 | 612 | -0,843099322 | 0,56398221 | 0,00115326 | 0,00693491 | Down |
| Plin1 | 55 | -1 | 0,34609567 | 3,41E-06 | 4,61E-05 | Down |
| Zscan2 | 576 | 0,607402455 | 0,12030727 | 5,21E-09 | 1,40E-07 | Up |
| 1700010L04Rik | 61 | -1 | 0,26138155 | 1,02E-06 | 1,63E-05 | Down |
| Abhd17c | 18.929 | -1 | 0,12475957 | 2,14E-33 | 1,49E-30 | Down |
| Arnt2 | 850 | -3 | 0,36741141 | 2,26E-18 | 2,78E-16 | Down |
| Gm44556 | 11 | -2 | 1 | 0,00071109 | 0,00462565 | Down |
| Rab38 | 29.661 | 0,813648632 | 0,09225135 | 3,53E-20 | 5,84E-18 | Up |
| Ankrd42 | 39 | 1 | 0,40492436 | 2,84E-05 | 0,00029251 | Up |
| Kctd14 | 149 | -0,900326738 | 0,28202552 | 1,15E-05 | 0,00013415 | Down |
| Myo7a | 1.301 | -0,685927213 | 0,41003638 | 0,0011657 | 0,00699713 | Down |
| Lrrc32 | 9 | -0,707095862 | 0,75673709 | 0,00490429 | 0,02270983 | Down |
| Gdpd5 | 22 | -1 | 0,54857935 | 0,00015486 | 0,0012562 | Down |
| Xrra1 | 319 | -0,586132975 | 0,25014863 | 0,00029922 | 0,00221569 | Down |
| P2ry6 | 5 | -1 | 1 | 0,00178988 | 0,00997457 | Down |
| Folr1 | 1.261 | -0,624354896 | 0,28173366 | 0,00037128 | 0,00267242 | Down |
| Usp17ld | 816 | -4 | 0,19668535 | 1,16E-87 | 1,94E-83 | Down |
| Usp17lc | 26 | -5 | 0,7939635 | 1,33E-12 | 7,09E-11 | Down |
| Hbb-y | 20 | 0,896034238 | 1 | 0,00362279 | 0,01771696 | Up |
| Trim30d | 92 | -0,781779547 | 0,41398121 | 0,00053386 | 0,00362737 | Down |
| Gm4070 | 373 | -2 | 0,23805841 | 1,76E-14 | 1,27E-12 | Down |
| Trim66 | 1.781 | -0,736224239 | 0,17780635 | 4,09E-07 | 7,17E-06 | Down |
| 4930431P19Rik | 74 | 0,606679904 | 0,28861572 | 0,00053856 | 0,00365493 | Up |
| RP23-462D5,5 | 107 | 0,737566156 | 0,26304547 | 4,96E-05 | 0,00047442 | Up |
| Sbf2 | 14 | 1 | 0,87029767 | 0,00086098 | 0,00541875 | Up |
| Ampd3 | 74 | -0,701662395 | 0,22860812 | 2,28E-05 | 0,00024222 | Down |
| Mrvi1 | 557 | -1 | 0,6036621 | 0,00050147 | 0,00344642 | Down |
| Plekha7 | 1.983 | -0,585450328 | 0,15312507 | 1,67E-06 | 2,46E-05 | Down |
| Dnah3 | 10 | -0,709210852 | 0,73110712 | 0,00459271 | 0,02150502 | Down |
| Gm44901 | 6 | 0,595056074 | 0,71736236 | 0,00796399 | 0,03368992 | Up |
| Gm20274 | 275 | -1 | 0,23382337 | 5,30E-11 | 2,08E-09 | Down |
| Slc5a11 | 765 | -1 | 0,16497469 | 9,77E-15 | 7,30E-13 | Down |
| Il4ra | 15.006 | -0,708409621 | 0,11468769 | 1,02E-11 | 4,58E-10 | Down |
| Apobr | 3.807 | -0,867434777 | 0,11293409 | 4,59E-16 | 4,17E-14 | Down |
| Nupr1 | 765 | -1 | 0,71461023 | 0,00040654 | 0,00288529 | Down |
| Fam57b | 565 | 0,747582678 | 0,13665818 | 7,03E-10 | 2,25E-08 | Up |
| Mylpf | 587 | 0,628898631 | 0,14063656 | 9,08E-08 | 1,84E-06 | Up |
| Itgam | 2.871 | -2 | 0,21482219 | 2,01E-24 | 5,02E-22 | Down |
| Cox6a2 | 154 | -3 | 0,64699641 | 2,83E-07 | 5,10E-06 | Down |

| | | | | | | |
|---|---|---|---|---|---|---|
| Rgs10 | 77 | 0,863656977 | 0,26269279 | 8,92E-06 | 0,00010805 | Up |
| Plekha1 | 6 | -1 | 2 | 0,00210129 | 0,01136403 | Down |
| Gm35147 | 8 | -1 | 0,95185327 | 0,0012452 | 0,00737906 | Down |
| Lrrc27 | 226 | -1 | 0,17952188 | 3,87E-12 | 1,89E-10 | Down |
| Adam8 | 725 | -4 | 0,41065435 | 4,76E-22 | 1,01E-19 | Down |
| Ifitm1 | 142.241 | -1 | 0,16782699 | 5,47E-19 | 7,15E-17 | Down |
| Gm7514 | 58 | -3 | 0,35952488 | 3,03E-19 | 4,16E-17 | Down |
| Ifitm3 | 93.619 | -2 | 0,27519298 | 8,25E-14 | 5,46E-12 | Down |
| Ifitm6 | 396 | -5 | 0,72621047 | 5,76E-16 | 5,13E-14 | Down |
| Lrrc56 | 288 | 1 | 0,16838591 | 1,75E-12 | 9,15E-11 | Up |
| Cdhr5 | 60 | -1 | 0,40322374 | 7,30E-05 | 0,00066066 | Down |
| Muc6 | 11 | 2 | 0,89751283 | 0,00034008 | 0,00247874 | Up |
| Muc5b | 13 | -1 | 0,82212404 | 0,00053164 | 0,00361751 | Down |
| Dusp8 | 14 | -3 | 1 | 1,54E-05 | 0,00017233 | Down |
| Trpm5 | 33 | -1 | 0,40905446 | 8,32E-05 | 0,00073854 | Down |
| Osbpl5 | 160 | 2 | 0,40536476 | 4,82E-08 | 1,04E-06 | Up |
| A430078G23Rik | 2.139 | 0,656103208 | 0,13001643 | 5,49E-09 | 1,47E-07 | Up |
| Arhgef18 | 7.936 | 0,677996398 | 0,1031288 | 7,11E-13 | 4,05E-11 | Up |
| Irs2 | 34 | -2 | 0,43570069 | 4,32E-08 | 9,47E-07 | Down |
| Col4a2 | 1.922 | -3 | 0,16948721 | 2,85E-50 | 4,76E-47 | Down |
| B020031H02Rik | 68 | -7 | 2 | 8,74E-07 | 1,42E-05 | Down |
| Mcf2l | 272 | -2 | 0,78912577 | 4,79E-05 | 0,0004592 | Down |
| F7 | 73 | -3 | 0,58348589 | 3,37E-07 | 6,01E-06 | Down |
| Adprhl1 | 12 | 1 | 0,94073108 | 0,00082788 | 0,00525391 | Up |
| 9530085L11Rik | 133 | 0,648527143 | 0,17133545 | 1,77E-06 | 2,60E-05 | Up |
| Gm7676 | 30 | -2 | 0,39963093 | 9,87E-08 | 1,98E-06 | Down |
| Rbpms | 4.998 | 1 | 0,1692943 | 1,23E-15 | 1,03E-13 | Up |
| Jund | 3.265 | -0,803532812 | 0,22588289 | 3,81E-06 | 5,09E-05 | Down |
| Il12rb1 | 432 | -0,72127772 | 0,17811724 | 5,95E-07 | 1,00E-05 | Down |
| Abhd8 | 2.098 | 0,836946075 | 0,11771694 | 3,02E-14 | 2,11E-12 | Up |
| Nwd1 | 3.134 | 1 | 0,33750883 | 9,55E-06 | 0,00011455 | Up |
| F2rl3 | 624 | -0,762094514 | 0,18213098 | 3,45E-07 | 6,13E-06 | Down |
| Smad1 | 116 | 2 | 0,29281515 | 3,21E-13 | 1,95E-11 | Up |
| Il15 | 77 | 0,804848047 | 0,45149872 | 0,00064907 | 0,00428997 | Up |
| Gm10644 | 47 | 2 | 0,33048226 | 1,12E-08 | 2,80E-07 | Up |
| Palm3 | 106 | -0,618667252 | 0,3354737 | 0,00098886 | 0,0060975 | Down |
| Klf1 | 813 | 1 | 0,12688198 | 6,11E-34 | 4,65E-31 | Up |
| Junb | 6.396 | -0,658702356 | 0,10665594 | 8,70E-12 | 3,94E-10 | Down |
| Gpt2 | 11.447 | -0,776752068 | 0,20081784 | 1,23E-06 | 1,89E-05 | Down |
| Gm45425 | 36 | -2 | 0,36554258 | 1,34E-09 | 4,05E-08 | Down |
| Bbs2 | 490 | 0,728094618 | 0,11454655 | 3,54E-12 | 1,75E-10 | Up |
| Adgrg1 | 224.935 | 0,771972543 | 0,10047011 | 3,71E-16 | 3,43E-14 | Up |
| Drc7 | 87 | 2 | 0,40832649 | 7,29E-07 | 1,20E-05 | Up |
| Kifc3 | 1.942 | 0,918218372 | 0,20954786 | 1,48E-07 | 2,83E-06 | Up |

| | | | | | | |
|---|---|---|---|---|---|---|
| Cmtm3 | 208 | 0,802553414 | 0,18622874 | 2,06E-07 | 3,83E-06 | Up |
| Ces2g | 341 | -2 | 0,54948673 | 6,49E-07 | 1,08E-05 | Down |
| Lrrc29 | 453 | 0,602437951 | 0,0966044 | 5,26E-12 | 2,50E-10 | Up |
| Gm38250 | 55 | 1 | 0,32275689 | 5,66E-06 | 7,22E-05 | Up |
| Nqo1 | 89 | 2 | 0,32593903 | 6,25E-09 | 1,65E-07 | Up |
| Hp | 192 | -0,981116153 | 0,69014571 | 0,00106517 | 0,00647969 | Down |
| 1700030J22Rik | 40 | 0,695768258 | 0,36039946 | 0,00061398 | 0,00408949 | Up |
| Osgin1 | 363 | -0,971854327 | 0,23436202 | 3,66E-07 | 6,46E-06 | Down |
| Cotl1 | 639 | -1 | 0,24157994 | 4,79E-08 | 1,04E-06 | Down |
| Gse1 | 29.887 | 0,814205187 | 0,11461536 | 3,03E-14 | 2,11E-12 | Up |
| 1700016A09Rik | 38 | 2 | 0,37628623 | 2,53E-07 | 4,58E-06 | Up |
| Gm26812 | 26 | -0,774492346 | 0,49940474 | 0,00119089 | 0,007119 | Down |
| Slc7a5 | 89.433 | -0,651024428 | 0,12408908 | 1,90E-09 | 5,58E-08 | Down |
| Gm27011 | 35 | 0,682712247 | 0,39664076 | 0,00105423 | 0,00642715 | Up |
| Gm17709 | 80 | -0,726633665 | 0,29617508 | 0,00014194 | 0,0011674 | Down |
| Zfpm1 | 21.568 | 0,84288987 | 0,11995037 | 5,45E-14 | 3,73E-12 | Up |
| Gm45353 | 43 | 3 | 0,37669018 | 4,37E-13 | 2,58E-11 | Up |
| Cyba | 21.003 | -0,817871913 | 0,14779932 | 5,70E-10 | 1,85E-08 | Down |
| RP24-94F12,3 | 3 | 0,789326389 | 1 | 0,0043637 | 0,02064064 | Up |
| Spata33 | 98 | 0,637287229 | 0,28962848 | 0,0003706 | 0,00266979 | Up |
| Rab4a | 2.004 | 0,587521287 | 0,08243139 | 1,19E-14 | 8,83E-13 | Up |
| RP23-312B17,1 | 6 | -0,611278842 | 0,81564683 | 0,00829989 | 0,03484664 | Down |
| Mmp8 | 20 | -2 | 1 | 0,00015109 | 0,00123278 | Down |
| Maml2 | 1.578 | 1 | 0,15521233 | 2,22E-12 | 1,14E-10 | Up |
| Icam1 | 2.168 | -0,795640512 | 0,12837184 | 1,16E-11 | 5,14E-10 | Down |
| RP24-496O17,7 | 26 | 0,586180111 | 0,39652938 | 0,00256656 | 0,01341392 | Up |
| AB124611 | 2.151 | -0,906668924 | 0,1877628 | 2,05E-08 | 4,88E-07 | Down |
| 9530077C05Rik | 71 | 1 | 0,32772673 | 4,81E-06 | 6,25E-05 | Up |
| Cdon | 154 | 0,922633749 | 0,68719707 | 0,00139169 | 0,00807838 | Up |
| Ddx25 | 19 | 1 | 1 | 0,00139353 | 0,00808621 | Up |
| Robo3 | 30 | -3 | 0,4681975 | 4,56E-12 | 2,21E-10 | Down |
| Vsig2 | 49 | 1 | 0,3899427 | 2,22E-05 | 0,00023715 | Up |
| Nrgn | 5.177 | 1 | 0,17345986 | 3,77E-11 | 1,52E-09 | Up |
| Hyou1 | 86.610 | -0,615598347 | 0,15839825 | 1,21E-06 | 1,88E-05 | Down |
| RP24-175C20,15 | 72 | -1 | 0,41642665 | 1,30E-05 | 0,00014845 | Down |
| RP24-175C20,19 | 352 | 0,591742328 | 0,17497026 | 9,37E-06 | 0,00011276 | Up |
| Nxpe2 | 32 | 1 | 0,6510448 | 0,00050378 | 0,0034595 | Up |
| Tmprss5 | 6 | -0,755699327 | 0,8949017 | 0,00494053 | 0,02285234 | Down |
| 4833427G06Rik | 26 | -1 | 0,56397227 | 0,00040988 | 0,00290282 | Down |
| Dmxl2 | 12.821 | -0,689682198 | 0,16622515 | 3,89E-07 | 6,84E-06 | Down |
| Gm10658 | 113 | 0,685984263 | 0,19660875 | 5,36E-06 | 6,88E-05 | Up |
| Lman1l | 23 | -1 | 0,47116883 | 0,0001647 | 0,00132449 | Down |
| Cd276 | 489 | -2 | 0,25040306 | 1,56E-21 | 3,08E-19 | Down |

| | | | | | | |
|---|---|---|---|---|---|---|
| Neo1 | 877 | -2 | 0,9284427 | 0,00023209 | 0,00177585 | Down |
| Tmem202 | 7 | -0,701511376 | 0,79040546 | 0,00535848 | 0,0242756 | Down |
| Celf6 | 136 | 1 | 0,22634897 | 2,12E-08 | 5,01E-07 | Up |
| Gramd2 | 16 | -1 | 0,73446384 | 0,00054872 | 0,00370724 | Down |
| Uaca | 471 | -0,782127092 | 0,4514651 | 0,00077194 | 0,00494387 | Down |
| RP23-9A12,4 | 6 | -0,900349535 | 1 | 0,0033606 | 0,01667346 | Down |
| RP23-116C19,1 | 234 | -1 | 0,3343005 | 1,33E-06 | 2,03E-05 | Down |
| Tle3 | 102 | -3 | 0,35170735 | 1,31E-18 | 1,64E-16 | Down |
| Slc24a1 | 31 | 0,848740804 | 0,4949818 | 0,00068956 | 0,0045077 | Up |
| Parp16 | 266 | 1 | 0,14755333 | 2,10E-13 | 1,33E-11 | Up |
| Car12 | 3.838 | -0,670168145 | 0,11928676 | 2,53E-10 | 8,71E-09 | Down |
| Aqp9 | 2.236 | -3 | 0,38424958 | 9,93E-18 | 1,15E-15 | Down |
| RP23-2F19,2 | 3 | -1 | 2 | 0,00115708 | 0,00695289 | Down |
| Gm37234 | 52 | -0,654001879 | 0,50035416 | 0,00283273 | 0,01456397 | Down |
| Ccpg1os | 185 | 0,687230556 | 0,14566481 | 2,95E-08 | 6,73E-07 | Up |
| Gsta4 | 475 | 2 | 0,15283132 | 5,35E-28 | 2,03E-25 | Up |
| Sh3bgrl2 | 23 | -3 | 0,63904078 | 5,85E-08 | 1,24E-06 | Down |
| Bcl2a1d | 151 | -3 | 0,27623632 | 2,84E-25 | 8,05E-23 | Down |
| Bcl2a1a | 3.219 | -2 | 0,31161556 | 1,49E-14 | 1,08E-12 | Down |
| Bcl2a1b | 1.443 | -2 | 0,23676723 | 6,14E-23 | 1,41E-20 | Down |
| Gm29094 | 4 | -2 | 2 | 0,00048434 | 0,0033452 | Down |
| Pcolce2 | 191 | 1 | 0,25671751 | 1,10E-08 | 2,77E-07 | Up |
| Pls1 | 3.170 | 2 | 0,20003257 | 3,04E-31 | 1,59E-28 | Up |
| 1700065D16Rik | 152 | 2 | 0,2317068 | 9,15E-19 | 1,17E-16 | Up |
| Rasa2 | 3.209 | -0,581023138 | 0,12802976 | 6,89E-08 | 1,44E-06 | Down |
| Gm26767 | 320 | -3 | 1 | 7,13E-05 | 0,0006467 | Down |
| Ky | 967 | -2 | 0,14578733 | 1,98E-60 | 1,65E-56 | Down |
| Ryk | 48 | 2 | 0,35741481 | 1,12E-08 | 2,80E-07 | Up |
| Srprb | 8.688 | -0,808699849 | 0,16963195 | 2,71E-08 | 6,24E-07 | Down |
| Bfsp2 | 56 | -2 | 0,29230583 | 3,18E-16 | 3,02E-14 | Down |
| Nphp3 | 37 | -1 | 0,76601322 | 0,0005215 | 0,00356219 | Down |
| Tlr9 | 120 | -3 | 0,47826565 | 2,49E-09 | 7,12E-08 | Down |
| Alas1 | 5.365 | -1 | 0,12099768 | 2,97E-33 | 1,99E-30 | Down |
| Dusp7 | 2.854 | -0,780208635 | 0,08609854 | 3,64E-21 | 6,99E-19 | Down |
| Rrp9 | 13.276 | -0,589575134 | 0,10271522 | 1,10E-10 | 4,07E-09 | Down |
| Mapkapk3 | 18.250 | -2 | 0,17564776 | 6,58E-20 | 1,03E-17 | Down |
| Cish | 14.897 | -2 | 0,15051307 | 9,39E-58 | 5,24E-54 | Down |
| Hemk1 | 5.617 | -1 | 0,15495873 | 1,48E-17 | 1,67E-15 | Down |
| 6430571L13Rik | 341 | -2 | 0,19068974 | 2,01E-31 | 1,12E-28 | Down |
| Sema3b | 10 | 3 | 0,91047351 | 1,94E-05 | 0,00021122 | Up |
| Gnat1 | 18 | -0,980825822 | 1 | 0,00292003 | 0,01493026 | Down |
| Nicn1 | 1.483 | 0,668863507 | 0,11147383 | 2,65E-11 | 1,10E-09 | Up |
| Camp | 6 | -0,860038958 | 1 | 0,00366179 | 0,01788154 | Down |
| Gm42432 | 32 | -0,969336113 | 0,60032723 | 0,00069972 | 0,00456687 | Down |

| | | | | | | |
|---|---|---|---|---|---|---|
| Gm42762 | 74 | 0,739818354 | 0,38217765 | 0,0005322 | 0,00361751 | Up |
| Ltf | 279 | -3 | 0,3447683 | 1,83E-19 | 2,69E-17 | Down |
| Bcl2a1c | 8 | -1 | 1 | 0,00103769 | 0,00634483 | Down |
| Ccr4 | 3.470 | -0,78884583 | 0,17528667 | 8,99E-08 | 1,82E-06 | Down |
| Csrnp1 | 4.758 | -0,762316953 | 0,12671556 | 3,23E-11 | 1,32E-09 | Down |
| Pomgnt2 | 40 | 1 | 0,44033324 | 3,17E-05 | 0,00032217 | Up |
| Ano10 | 47 | -2 | 0,52176696 | 4,74E-07 | 8,17E-06 | Down |
| Abhd5 | 4.996 | -0,73221535 | 0,12084709 | 2,28E-11 | 9,69E-10 | Down |
| Zkscan7 | 38 | 0,614518432 | 0,40801812 | 0,00216344 | 0,01162657 | Up |
| Ccr1 | 520 | -3 | 0,60521017 | 1,51E-10 | 5,47E-09 | Down |
| Ccr2 | 81 | -2 | 0,98760767 | 0,00022083 | 0,00170536 | Down |
| Ccr5 | 30 | -1 | 0,92148613 | 0,00071271 | 0,00462835 | Down |
| Grm1 | 15 | 0,621406797 | 0,6485877 | 0,00601242 | 0,02683143 | Up |
| Aig1 | 6 | -0,634214452 | 1 | 0,00759245 | 0,03246286 | Down |
| Cited2 | 55.471 | 1 | 0,18428533 | 3,84E-12 | 1,89E-10 | Up |
| Nhsl1 | 1.298 | -0,757051452 | 0,13832657 | 7,09E-10 | 2,26E-08 | Down |
| Arfgef3 | 2.395 | -0,683162239 | 0,14718771 | 4,22E-08 | 9,27E-07 | Down |
| Il22ra2 | 15 | 5 | 2 | 8,12E-06 | 9,97E-05 | Up |
| Aldh8a1 | 263 | 4 | 1 | 3,80E-06 | 5,08E-05 | Up |
| Gm34267 | 26 | -0,981582933 | 0,5355054 | 0,00041106 | 0,00290869 | Down |
| Gm26581 | 5 | -2 | 2 | 0,00072728 | 0,00470586 | Down |
| C920009B18Rik | 1.010 | -0,669425891 | 0,13145798 | 4,42E-09 | 1,20E-07 | Down |
| Raet1d | 36 | -4 | 0,7727233 | 3,42E-09 | 9,52E-08 | Down |
| Gm10825 | 50 | -8 | 1 | 1,47E-13 | 9,40E-12 | Down |
| Stx7 | 9.119 | -0,625093266 | 0,0992129 | 3,62E-12 | 1,79E-10 | Down |
| RP23-331E5,10 | 95 | 0,734171477 | 0,30875093 | 0,00017175 | 0,00137394 | Up |
| Traf3ip2 | 784 | -0,690170775 | 0,11782425 | 6,65E-11 | 2,56E-09 | Down |
| Cdk19 | 4.869 | 0,672050785 | 0,11625104 | 9,90E-11 | 3,67E-09 | Up |
| Scml4 | 62 | 3 | 0,41090396 | 2,91E-14 | 2,04E-12 | Up |
| Lilrb4a | 4.830 | -5 | 0,5931201 | 3,01E-17 | 3,23E-15 | Down |
| Cdh23 | 246 | 0,903865256 | 0,3725336 | 0,00010507 | 0,00090449 | Up |
| Vsir | 3.909 | -1 | 0,36960675 | 4,39E-06 | 5,78E-05 | Down |
| Sgpl1 | 5.661 | -0,747405539 | 0,12534418 | 4,25E-11 | 1,69E-09 | Down |
| Hkdc1 | 600 | -1 | 0,24060895 | 9,03E-11 | 3,40E-09 | Down |
| Tet1 | 377 | 0,762444788 | 0,34615688 | 0,00025537 | 0,00192854 | Up |
| Rhobtb1 | 962 | -0,740135226 | 0,09394211 | 7,12E-17 | 7,36E-15 | Down |
| Adora2a | 4.036 | 0,845734129 | 0,13476828 | 7,82E-12 | 3,55E-10 | Up |
| Ggt5 | 50 | 1 | 0,35715467 | 1,02E-05 | 0,00012128 | Up |
| Gm10941 | 15 | 1 | 0,6148968 | 0,00062899 | 0,0041688 | Up |
| Itgb2 | 23.013 | -0,70840154 | 0,09902288 | 1,47E-14 | 1,07E-12 | Down |
| Trpm2 | 18 | -2 | 0,6140113 | 8,33E-06 | 0,00010192 | Down |
| Icosl | 2.230 | -0,600448593 | 0,08443523 | 1,36E-14 | 9,96E-13 | Down |
| Agpat3 | 3.053 | -0,89722398 | 0,12936378 | 1,04E-13 | 6,79E-12 | Down |
| 2610008E11Rik | 616 | 0,700024058 | 0,1150172 | 1,73E-11 | 7,53E-10 | Up |

| | | | | | | |
|---|---|---|---|---|---|---|
| Plpp2 | 1.736 | -0,602842384 | 0,09357726 | 1,39E-12 | 7,31E-11 | Down |
| Plppr3 | 5.056 | -1 | 0,20526162 | 2,56E-10 | 8,78E-09 | Down |
| Gamt | 54 | 0,946623202 | 0,31570604 | 1,96E-05 | 0,00021284 | Up |
| Apc2 | 209 | 0,864236046 | 0,26376315 | 9,24E-06 | 0,00011146 | Up |
| Btbd2 | 3.934 | 0,641372213 | 0,14521497 | 1,17E-07 | 2,31E-06 | Up |
| Tjp3 | 624 | 1 | 0,27685728 | 5,71E-08 | 1,21E-06 | Up |
| RP23-132J19,1 | 72 | -2 | 0,32817512 | 1,51E-13 | 9,59E-12 | Down |
| A230060F14Rik | 82 | 2 | 0,32118616 | 7,46E-13 | 4,23E-11 | Up |
| Fbxo7 | 10.180 | 1 | 0,1862332 | 8,56E-13 | 4,79E-11 | Up |
| Syn3 | 41 | 2 | 0,47834082 | 4,86E-08 | 1,05E-06 | Up |
| Gm26546 | 34 | 0,662955443 | 0,41229337 | 0,00145552 | 0,00837046 | Up |
| Ano4 | 5 | 2 | 2 | 0,00073935 | 0,00477356 | Up |
| Gas2l3 | 16.653 | 0,927091421 | 0,19463007 | 2,74E-08 | 6,31E-07 | Up |
| Nr1h4 | 82 | 3 | 0,6083623 | 2,63E-10 | 9,02E-09 | Up |
| Slc17a8 | 23 | 3 | 0,92342321 | 1,02E-05 | 0,00012128 | Up |
| 1500026H17Rik | 15 | 0,857849495 | 0,75902613 | 0,00244001 | 0,01287718 | Up |
| Anks1b | 53 | -2 | 0,37311566 | 1,10E-07 | 2,17E-06 | Down |
| RP23-381H23,1 | 442 | 0,707309837 | 0,18849533 | 1,95E-06 | 2,82E-05 | Up |
| Amdhd1 | 240 | -0,768934911 | 0,22690282 | 7,08E-06 | 8,81E-05 | Down |
| Ccdc38 | 1.049 | -0,802795022 | 0,24616755 | 1,05E-05 | 0,00012423 | Down |
| Gm15915 | 150 | -2 | 0,20194173 | 5,79E-16 | 5,13E-14 | Down |
| Cep83os | 549 | 0,584131327 | 0,22983838 | 0,00016749 | 0,00134387 | Up |
| Plxnc1 | 719 | -2 | 0,18572068 | 5,95E-18 | 7,02E-16 | Down |
| Cradd | 420 | 0,647425375 | 0,12840653 | 5,55E-09 | 1,48E-07 | Up |
| Dusp6 | 5.131 | -0,983739345 | 0,14918147 | 9,04E-13 | 5,03E-11 | Down |
| Mgat4c | 93 | 0,917580933 | 0,3539002 | 6,55E-05 | 0,00060004 | Up |
| Tmtc2 | 3.411 | -0,642647808 | 0,11573152 | 3,45E-10 | 1,16E-08 | Down |
| Gm15663 | 430 | -0,914425147 | 0,14264665 | 3,35E-12 | 1,67E-10 | Down |
| Tph2 | 12 | 2 | 0,74023286 | 1,11E-05 | 0,00013 | Up |
| Avil | 248 | -3 | 0,86172399 | 5,42E-06 | 6,95E-05 | Down |
| Kif5a | 352 | 0,658459509 | 0,20667019 | 1,65E-05 | 0,00018352 | Up |
| Ddit3 | 1.598 | -0,593047517 | 0,33554434 | 0,00131433 | 0,00770141 | Down |
| Lrp1 | 58 | -2 | 0,31979571 | 1,43E-11 | 6,26E-10 | Down |
| Il23a | 78 | -0,59239842 | 0,32850125 | 0,00121028 | 0,00721299 | Down |
| Mir8105 | 15 | 0,719745578 | 0,59142064 | 0,00279666 | 0,01441837 | Up |
| Mettl7b | 12 | 3 | 2 | 0,00010452 | 0,00090066 | Up |
| Tespa1 | 1.680 | -0,612134267 | 0,13231112 | 4,37E-08 | 9,55E-07 | Down |
| Ccdc157 | 240 | 0,812537904 | 0,19899586 | 5,24E-07 | 8,93E-06 | Up |
| Gatsl3 | 235 | -2 | 0,2207362 | 4,71E-26 | 1,41E-23 | Down |
| Osm | 2.398 | -2 | 0,1367621 | 6,57E-30 | 2,89E-27 | Down |
| Lif | 96 | -2 | 0,28931552 | 5,02E-12 | 2,39E-10 | Down |
| Gm11963 | 138 | -1 | 0,25669289 | 2,67E-08 | 6,15E-07 | Down |
| Xbp1 | 46.770 | -1 | 0,10678206 | 2,78E-25 | 8,03E-23 | Down |
| Polm | 1.100 | 1 | 0,16070619 | 9,63E-13 | 5,34E-11 | Up |

| | | | | | | |
|---|---|---|---|---|---|---|
| Gck | 66 | 1 | 0,28597769 | 6,34E-07 | 1,06E-05 | Up |
| H2afv | 5.771 | 0,594830226 | 0,15313716 | 1,27E-06 | 1,95E-05 | Up |
| Upp1 | 1.663 | 0,836142894 | 0,20422888 | 4,99E-07 | 8,56E-06 | Up |
| Abca13 | 259 | -0,862152721 | 0,27991578 | 1,73E-05 | 0,00019053 | Down |
| 1700042O10Rik | 212 | -0,918637993 | 0,18417092 | 9,56E-09 | 2,42E-07 | Down |
| Grb10 | 261 | -0,950568218 | 0,63331065 | 0,00093477 | 0,00580458 | Down |
| Plek | 48.649 | 1 | 0,1585463 | 4,25E-19 | 5,60E-17 | Up |
| Gm12031 | 350 | -0,668198522 | 0,19639894 | 7,53E-06 | 9,31E-05 | Down |
| Wdpcp | 418 | 0,889146528 | 0,16073366 | 5,98E-10 | 1,93E-08 | Up |
| Hba-x | 4 | 2 | 2 | 0,00085624 | 0,00539296 | Up |
| Olfr56 | 110 | -1 | 0,24545764 | 9,81E-09 | 2,48E-07 | Down |
| Tgtp2 | 69 | -2 | 0,318579 | 9,78E-11 | 3,64E-09 | Down |
| Ifi47 | 1.481 | -0,982899029 | 0,14719638 | 5,22E-13 | 3,04E-11 | Down |
| Rasgef1c | 7 | 0,582209482 | 0,78556326 | 0,00922083 | 0,0379328 | Up |
| Ltc4s | 612 | -0,824216623 | 0,16004732 | 4,29E-09 | 1,17E-07 | Down |
| D930048N14Rik | 299 | 0,71827462 | 0,20613423 | 5,29E-06 | 6,80E-05 | Up |
| Tcf7 | 8 | 1 | 0,87600151 | 0,00120843 | 0,00720616 | Up |
| Sep 08 | 18.118 | 0,589920447 | 0,0791286 | 1,04E-15 | 8,88E-14 | Up |
| Acsl6 | 220 | 0,82831446 | 0,23391465 | 3,95E-06 | 5,26E-05 | Up |
| Slc36a3 | 37 | -0,657679726 | 0,48247691 | 0,00249819 | 0,01311805 | Down |
| Slc36a1os | 60 | 1 | 0,30697795 | 8,27E-06 | 0,00010129 | Up |
| Fat2 | 186 | -2 | 0,32743402 | 7,94E-15 | 6,01E-13 | Down |
| Gm12242 | 16 | 0,917886326 | 0,67245689 | 0,00133615 | 0,00779653 | Up |
| Gm12247 | 49 | 1 | 0,3843332 | 5,19E-05 | 0,00049341 | Up |
| Trim58 | 2.092 | 1 | 0,12768083 | 3,58E-22 | 7,77E-20 | Up |
| Natd1 | 990 | 1 | 0,15866365 | 4,71E-14 | 3,24E-12 | Up |
| Map2k3 | 20.175 | 0,715225582 | 0,11239725 | 3,23E-12 | 1,63E-10 | Up |
| Specc1 | 7.088 | 0,735476598 | 0,13724596 | 1,25E-09 | 3,79E-08 | Up |
| Trpv2 | 5.777 | -0,638691695 | 0,18375443 | 5,98E-06 | 7,56E-05 | Down |
| Trim16 | 1.371 | -2 | 0,48036344 | 2,67E-06 | 3,73E-05 | Down |
| Hs3st3b1 | 2.950 | -0,857893548 | 0,09793319 | 6,38E-20 | 1,01E-17 | Down |
| Pik3r5 | 3.647 | -0,610484274 | 0,14928249 | 5,15E-07 | 8,82E-06 | Down |
| Alox8 | 111 | -1 | 0,25707567 | 1,60E-08 | 3,89E-07 | Down |
| Tmem88 | 18 | -0,782888704 | 0,66810105 | 0,00262514 | 0,01365614 | Down |
| Kdm6b | 6.599 | -1 | 0,11890212 | 1,19E-24 | 3,08E-22 | Down |
| Cd68 | 1.730 | -2 | 0,24051458 | 4,14E-12 | 2,02E-10 | Down |
| Tnfsf13os | 9 | 1 | 0,94321019 | 0,00047116 | 0,00326899 | Up |
| Tnfsfm13 | 4 | 0,818350402 | 1 | 0,00406162 | 0,01948757 | Up |
| Tnfsf13 | 63 | 1 | 0,32105172 | 9,05E-07 | 1,47E-05 | Up |
| Zbtb4 | 22 | 1 | 0,58107863 | 0,00035579 | 0,00257645 | Up |
| Mgl2 | 18 | 3 | 3 | 0,00015514 | 0,00125786 | Up |
| Arrb2 | 15.837 | -0,700829902 | 0,09130296 | 2,89E-16 | 2,76E-14 | Down |
| Gp1ba | 496 | 0,794980194 | 0,20274636 | 9,94E-07 | 1,58E-05 | Up |
| Rnf167 | 5.109 | 0,813331547 | 0,0915795 | 2,05E-20 | 3,57E-18 | Up |

| | | | | | | |
|---|---|---|---|---|---|---|
| Gm12319 | 87 | 0,640869778 | 0,22914737 | 6,30E-05 | 0,00058063 | Up |
| Scimp | 8 | -0,605148227 | 0,76527803 | 0,00810091 | 0,03413968 | Down |
| Nlrp1a | 85 | -0,59465971 | 0,23869315 | 0,00018754 | 0,00148601 | Down |
| Aipl1 | 423 | 0,590439862 | 0,22456482 | 0,00012488 | 0,00104804 | Up |
| Tekt1 | 6 | 0,725310266 | 1 | 0,00527843 | 0,02399091 | Up |
| Gm45606 | 553 | 0,968772163 | 0,16526397 | 8,69E-11 | 3,28E-09 | Up |
| Inpp5k | 2.880 | 0,743912459 | 0,10866124 | 1,47E-13 | 9,40E-12 | Up |
| Rph3al | 785 | -2 | 0,281794 | 5,58E-12 | 2,63E-10 | Down |
| Gm12341 | 96 | -2 | 0,21195767 | 2,48E-17 | 2,70E-15 | Down |
| Tmem97 | 13.702 | -0,607651535 | 0,09274154 | 6,73E-13 | 3,86E-11 | Down |
| Lgals9 | 16.272 | 0,611745078 | 0,20086215 | 2,98E-05 | 0,00030404 | Up |
| Rab11fip4 | 393 | -0,908230186 | 0,3032474 | 2,09E-05 | 0,00022496 | Down |
| Gm11417 | 103 | 0,865308807 | 0,20886834 | 4,06E-07 | 7,12E-06 | Up |
| Slfn5 | 588 | -1 | 0,2663885 | 2,63E-10 | 9,02E-09 | Down |
| Slfn10-ps | 1.498 | -0,671413046 | 0,09681985 | 5,94E-14 | 4,02E-12 | Down |
| Slfn2 | 1.046 | -4 | 0,33837932 | 8,60E-33 | 5,53E-30 | Down |
| Slfn1 | 9 | -4 | 2 | 3,80E-05 | 0,00037589 | Down |
| Ccl6 | 1.120 | -4 | 0,79373776 | 8,95E-10 | 2,81E-08 | Down |
| Ccl3 | 442 | -3 | 0,44638662 | 3,85E-12 | 1,89E-10 | Down |
| Ccl4 | 335 | -4 | 0,36003457 | 4,43E-29 | 1,85E-26 | Down |
| Wfdc21 | 16 | -2 | 0,97465711 | 4,20E-05 | 0,0004095 | Down |
| Epx | 30 | -4 | 1 | 6,55E-07 | 1,09E-05 | Down |
| Wfikkn2 | 11 | -2 | 0,65128178 | 8,13E-06 | 9,98E-05 | Down |
| Abcc3 | 884 | -1 | 0,15453705 | 3,41E-15 | 2,70E-13 | Down |
| Acsf2 | 171 | 1 | 0,22248222 | 6,99E-11 | 2,68E-09 | Up |
| Sgca | 19 | -0,913895347 | 0,60280606 | 0,0009605 | 0,0059379 | Down |
| Dlx4os | 173 | -0,58855321 | 0,19496561 | 3,50E-05 | 0,00035016 | Down |
| Tbkbp1 | 9.541 | -0,791249966 | 0,10282596 | 3,62E-16 | 3,39E-14 | Down |
| Plxdc1 | 19 | 1 | 0,58480013 | 0,0003572 | 0,00258553 | Up |
| Cacnb1 | 131 | 0,601153768 | 0,28279039 | 0,00051633 | 0,00353553 | Up |
| Rara | 1.249 | -1 | 0,13099992 | 2,59E-28 | 1,03E-25 | Down |
| Tns4 | 360 | -2 | 0,32974016 | 5,59E-10 | 1,82E-08 | Down |
| Ttc25 | 970 | -0,945643816 | 0,16478388 | 1,82E-10 | 6,45E-09 | Down |
| Etv4 | 842 | -3 | 0,17844692 | 9,25E-50 | 1,41E-46 | Down |
| Gm20511 | 9 | 0,621322739 | 0,82129683 | 0,00798074 | 0,0337352 | Up |
| Myl4 | 105 | 1 | 0,24110709 | 6,02E-08 | 1,27E-06 | Up |
| Itgb3 | 28.796 | 2 | 0,12769036 | 8,53E-49 | 1,19E-45 | Up |
| Icam2 | 2.299 | -0,729993664 | 0,11072864 | 7,69E-13 | 4,33E-11 | Down |
| Milr1 | 749 | -3 | 0,24304618 | 2,74E-26 | 8,50E-24 | Down |
| Slc16a6 | 11.529 | -0,962477767 | 0,13042988 | 3,95E-15 | 3,10E-13 | Down |
| Gm25540 | 31 | -0,719215313 | 0,43251387 | 0,00107604 | 0,00653631 | Down |
| Cd300a | 306 | 2 | 0,25821549 | 1,47E-11 | 6,41E-10 | Up |
| Cd300ld | 194 | -0,92409062 | 0,14756172 | 8,40E-12 | 3,81E-10 | Down |
| Cd300ld2 | 46 | -1 | 0,35970781 | 2,81E-05 | 0,00028942 | Down |

| | | | | | | |
|---|---|---|---|---|---|---|
| Rab37 | 416 | 0,767055115 | 0,21962835 | 4,94E-06 | 6,38E-05 | Up |
| Cd300lf | 3.294 | -0,96430918 | 0,10738573 | 7,67E-21 | 1,41E-18 | Down |
| Smim5 | 8.824 | 1 | 0,13386106 | 1,52E-16 | 1,49E-14 | Up |
| Itgb4 | 428 | -0,911385783 | 0,2113941 | 1,99E-07 | 3,73E-06 | Down |
| Trim47 | 130 | 0,791614731 | 0,21053002 | 1,84E-06 | 2,68E-05 | Up |
| Cdk3-ps | 19 | 0,724224329 | 0,685451 | 0,00378106 | 0,0183515 | Up |
| Rhbdf2 | 7.966 | -0,8482432 | 0,12130258 | 6,96E-14 | 4,66E-12 | Down |
| Tha1 | 1.142 | 0,63869094 | 0,16252996 | 9,90E-07 | 1,58E-05 | Up |
| Lgals3bp | 8.464 | -0,837764729 | 0,44021277 | 0,0004515 | 0,00315483 | Down |
| C1qtnf1 | 10 | -0,956654392 | 0,81493868 | 0,00187352 | 0,01034952 | Down |
| Engase | 48 | 1 | 0,83770987 | 0,00093621 | 0,00581137 | Up |
| Sgsh | 2.347 | 0,900588752 | 0,23838954 | 1,59E-06 | 2,37E-05 | Up |
| Slc26a11 | 69 | 1 | 0,46116348 | 5,97E-05 | 0,00055501 | Up |
| Narf | 2.990 | 0,751765968 | 0,18402482 | 5,17E-07 | 8,85E-06 | Up |
| Zfp750 | 45 | -1 | 0,35509972 | 6,89E-06 | 8,59E-05 | Down |
| Metrnl | 4.888 | -0,954355709 | 0,10822801 | 3,36E-20 | 5,66E-18 | Down |
| Gm12589 | 34 | -1 | 0,44898392 | 1,53E-05 | 0,00017173 | Down |
| 1110002L01Rik | 455 | -0,653564101 | 0,14124963 | 4,40E-08 | 9,60E-07 | Down |
| Fkbp1b | 190 | 0,947951722 | 0,21793177 | 1,65E-07 | 3,14E-06 | Up |
| Mycn | 8.715 | -0,815956575 | 0,2185633 | 2,00E-06 | 2,89E-05 | Down |
| Greb1 | 5.300 | 8 | 0,53895546 | 9,92E-52 | 1,85E-48 | Up |
| Gm4419 | 82 | 0,9013856 | 0,2806439 | 1,08E-05 | 0,00012745 | Up |
| Rrm2 | 48.360 | -0,607461411 | 0,11874944 | 3,67E-09 | 1,02E-07 | Down |
| Id2 | 4.320 | -0,640561097 | 0,14838027 | 1,85E-07 | 3,48E-06 | Down |
| Pik3cg | 41.780 | -0,586328178 | 0,16725316 | 5,94E-06 | 7,51E-05 | Down |
| Hdac9 | 276 | -2 | 0,2836268 | 1,02E-12 | 5,59E-11 | Down |
| Ifrd1 | 11.871 | -0,944072755 | 0,15494668 | 2,31E-11 | 9,80E-10 | Down |
| Nrcam | 67 | -0,872785919 | 0,55247982 | 0,00089654 | 0,00561091 | Down |
| Abhd12b | 5 | -1 | 1 | 0,00261224 | 0,01359745 | Down |
| Rtn1 | 325 | 1 | 0,16102682 | 1,46E-21 | 2,90E-19 | Up |
| Prkch | 22.396 | 0,748363614 | 0,08056067 | 3,89E-22 | 8,35E-20 | Up |
| Rhoj | 8.707 | -2 | 0,47109142 | 1,97E-07 | 3,70E-06 | Down |
| Sipa1l1 | 98.257 | 0,806812521 | 0,1156336 | 7,24E-14 | 4,81E-12 | Up |
| Prox2 | 519 | 0,645940755 | 0,10772288 | 2,55E-11 | 1,06E-09 | Up |
| Tgfb3 | 215 | 3 | 0,32071971 | 6,35E-20 | 1,01E-17 | Up |
| Ifi27 | 6.938 | -0,945536933 | 0,24148479 | 9,17E-07 | 1,48E-05 | Down |
| Serpina3f | 223 | -3 | 0,34179107 | 2,35E-20 | 4,05E-18 | Down |
| Serpina3g | 12.714 | -1 | 0,36172319 | 2,46E-05 | 0,0002581 | Down |
| Hhipl1 | 4 | -2 | 2 | 0,00085538 | 0,0053896 | Down |
| Evl | 138 | -1 | 1 | 0,00084002 | 0,00531884 | Down |
| Cdc42bpb | 67 | 2 | 1 | 8,61E-05 | 0,00076134 | Up |
| Pld4 | 17.834 | -0,620004058 | 0,17772846 | 5,88E-06 | 7,45E-05 | Down |
| Ahnak2 | 118 | -0,754981538 | 0,34664229 | 0,00027749 | 0,00207129 | Down |
| Gpr132 | 72 | -2 | 0,31844795 | 5,39E-12 | 2,55E-10 | Down |

| | | | | | | |
|---|---|---|---|---|---|---|
| Nudt14 | 1.735 | 0,648154771 | 0,11427581 | 1,76E-10 | 6,26E-09 | Up |
| Ighm | 1.458 | 1 | 0,25756973 | 1,01E-09 | 3,11E-08 | Up |
| Akr1c14 | 101 | 3 | 1 | 2,48E-05 | 0,00025941 | Up |
| Idi1 | 7.543 | 1 | 0,20053028 | 2,00E-11 | 8,54E-10 | Up |
| Idi2 | 7 | 0,766319515 | 0,9132813 | 0,00483954 | 0,02244724 | Up |
| Dip2c | 16 | -0,895949592 | 1 | 0,00360579 | 0,01765446 | Down |
| Lyst | 43 | -3 | 0,68223555 | 7,43E-09 | 1,94E-07 | Down |
| A530099J19Rik | 5 | -1 | 1 | 0,00172345 | 0,00965676 | Down |
| Gpr141 | 1.957 | -2 | 0,49512999 | 4,63E-08 | 1,01E-06 | Down |
| Zkscan4 | 181 | 0,981413588 | 0,22661681 | 1,72E-07 | 3,27E-06 | Up |
| Zfp184 | 128 | 0,667291426 | 0,23536585 | 5,23E-05 | 0,00049646 | Up |
| Hist1h3c | 151 | 0,604168124 | 0,22158709 | 8,77E-05 | 0,00077483 | Up |
| Hist1h4b | 40 | 0,598319643 | 0,33458209 | 0,00122459 | 0,00728235 | Up |
| Serpinb6b | 1.802 | 0,628650317 | 0,15240097 | 4,34E-07 | 7,56E-06 | Up |
| Serpinb9 | 774 | 0,832803718 | 0,18031478 | 5,42E-08 | 1,16E-06 | Up |
| Serpinb9g | 19 | -2 | 0,68495392 | 2,66E-05 | 0,0002766 | Down |
| Slc22a23 | 6.460 | 0,787252394 | 0,12232578 | 2,57E-12 | 1,31E-10 | Up |
| Ppp1r3g | 145 | -2 | 0,46266938 | 1,20E-07 | 2,35E-06 | Down |
| F13a1 | 34 | 6 | 2 | 2,20E-06 | 3,14E-05 | Up |
| Eef1e1 | 7.193 | -0,720539299 | 0,10198226 | 2,91E-14 | 2,04E-12 | Down |
| Gmpr | 96 | -0,604448803 | 0,26640852 | 0,00034471 | 0,00250704 | Down |
| Nfil3 | 7.839 | -0,663142165 | 0,11175711 | 3,90E-11 | 1,57E-09 | Down |
| Hrh2 | 19 | -2 | 0,84148186 | 6,66E-05 | 0,00060908 | Down |
| Unc5a | 834 | 1 | 0,21227997 | 1,00E-12 | 5,51E-11 | Up |
| Hk3 | 26.801 | -2 | 0,19629335 | 2,08E-23 | 4,96E-21 | Down |
| Slc34a1 | 1.097 | -0,963086324 | 0,28332504 | 5,49E-06 | 7,03E-05 | Down |
| Prr7 | 516 | -1 | 0,12693259 | 1,33E-22 | 2,93E-20 | Down |
| Dbn1 | 418 | -0,605679616 | 0,29338092 | 0,00059685 | 0,0039937 | Down |
| 4930451E10Rik | 18 | 0,653048883 | 0,48742841 | 0,00266027 | 0,01380881 | Up |
| Ctla2b | 974 | 2 | 0,21503713 | 3,37E-16 | 3,18E-14 | Up |
| Ctla2a | 2.672 | 1 | 0,18011465 | 9,13E-15 | 6,85E-13 | Up |
| Fbp2 | 13.397 | -2 | 0,18781281 | 1,52E-30 | 7,05E-28 | Down |
| Fbp1 | 507 | -3 | 0,30430152 | 3,67E-19 | 4,87E-17 | Down |
| Fancc | 5.048 | 1 | 0,1504599 | 2,46E-21 | 4,79E-19 | Up |
| Gm28557 | 188 | 0,704420898 | 0,1606461 | 1,40E-07 | 2,70E-06 | Up |
| Slc12a7 | 1.359 | -0,602420767 | 0,12521952 | 1,77E-08 | 4,24E-07 | Down |
| Mctp1 | 306 | 0,817454656 | 0,33079141 | 0,00010988 | 0,00094052 | Up |
| F2r | 100.570 | 0,849568692 | 0,23490624 | 2,97E-06 | 4,09E-05 | Up |
| Sv2c | 43 | 1 | 0,67290209 | 0,00040026 | 0,00284675 | Up |
| Tmem171 | 66 | -1 | 0,35034267 | 4,37E-07 | 7,60E-06 | Down |
| Gm807 | 5 | -1 | 1 | 0,002467 | 0,01299506 | Down |
| 4933425L06Rik | 18 | 0,865498582 | 0,58191464 | 0,00112104 | 0,00676547 | Up |
| Rnf180 | 15 | 4 | 0,75919675 | 1,46E-09 | 4,36E-08 | Up |
| Hspb3 | 3 | 0,712374609 | 1 | 0,00413214 | 0,01975751 | Up |

| | | | | | | |
|---|---|---|---|---|---|---|
| Emb | 15.484 | -1 | 0,16387495 | 1,32E-14 | 9,76E-13 | Down |
| Ccl28 | 5 | -0,635284684 | 0,83529361 | 0,00760543 | 0,03251005 | Down |
| Nim1k | 633 | 0,786750094 | 0,22875138 | 5,85E-06 | 7,42E-05 | Up |
| Acox2 | 9 | -1 | 2 | 0,00106129 | 0,00645854 | Down |
| Slc4a7 | 6.191 | -0,695214902 | 0,06242985 | 1,70E-30 | 7,70E-28 | Down |
| Ndst2 | 7.588 | 0,936106883 | 0,10757747 | 9,83E-20 | 1,51E-17 | Up |
| Gm10248 | 75 | -0,846688375 | 0,30673299 | 4,61E-05 | 0,00044448 | Down |
| Kcnma1 | 69 | -3 | 1 | 3,43E-05 | 0,00034413 | Down |
| Erc2 | 101 | 2 | 0,53621414 | 9,57E-06 | 0,00011461 | Up |
| Itih4 | 64 | 1 | 0,65664315 | 0,00054291 | 0,00367841 | Up |
| Stab1 | 573 | -1 | 0,52089365 | 0,00012403 | 0,00104146 | Down |
| Sh3bp5 | 733 | -1 | 0,22451359 | 2,42E-09 | 6,95E-08 | Down |
| Eaf1 | 14.776 | -0,703164534 | 0,10604267 | 5,38E-13 | 3,11E-11 | Down |
| Colq | 47 | -1 | 0,32745868 | 4,45E-06 | 5,84E-05 | Down |
| Gprin2 | 16 | 2 | 0,73031028 | 2,54E-05 | 0,00026581 | Up |
| Gm26782 | 363 | -0,659113071 | 0,16745878 | 9,53E-07 | 1,53E-05 | Down |
| Rnase10 | 9 | -5 | 2 | 3,29E-06 | 4,48E-05 | Down |
| Rnase4 | 47 | -1 | 0,51943159 | 0,00022248 | 0,00171494 | Down |
| B230359F08Rik | 16 | 1 | 0,83496778 | 0,0013041 | 0,00765413 | Up |
| Trdc | 4 | 0,586877336 | 1 | 0,00891269 | 0,03689575 | Up |
| Traj59 | 3 | 1 | 2 | 0,00153409 | 0,0087353 | Up |
| Trac | 148 | 3 | 0,33816339 | 1,09E-20 | 1,92E-18 | Up |
| Abhd4 | 11.073 | 0,901107533 | 0,11151044 | 1,94E-17 | 2,13E-15 | Up |
| Slc7a7 | 7.431 | 2 | 0,17528267 | 8,02E-21 | 1,46E-18 | Up |
| Haus4 | 5.871 | 0,868928405 | 0,07539517 | 3,86E-32 | 2,22E-29 | Up |
| 4931414P19Rik | 1.814 | 0,607901014 | 0,09374407 | 1,05E-12 | 5,75E-11 | Up |
| Cebpe | 1.625 | -0,915727246 | 0,4027536 | 0,00015344 | 0,0012459 | Down |
| Gm37271 | 94 | -1 | 0,40117342 | 2,16E-05 | 0,0002311 | Down |
| Slc22a17 | 1.544 | 0,997107827 | 0,11047287 | 4,49E-21 | 8,54E-19 | Up |
| Myh7 | 39 | -1 | 0,53112939 | 0,00020982 | 0,00163776 | Down |
| Cpne6 | 34 | -1 | 0,53444496 | 9,50E-05 | 0,00082924 | Down |
| Adcy4 | 352 | -0,893663732 | 0,1566614 | 2,31E-10 | 8,01E-09 | Down |
| Ripk3 | 124 | -2 | 0,29989742 | 7,57E-18 | 8,80E-16 | Down |
| Mcpt1 | 70 | 4 | 0,49426378 | 1,15E-19 | 1,72E-17 | Up |
| Mcpt2 | 338 | 2 | 0,16478559 | 5,78E-36 | 5,69E-33 | Up |
| Gzmc | 159 | 2 | 1 | 0,00037264 | 0,00268109 | Up |
| Cryl1 | 1.527 | 0,819952075 | 0,07703849 | 6,66E-28 | 2,42E-25 | Up |
| Fgf9 | 1.435 | -1 | 0,08026475 | 7,54E-48 | 9,70E-45 | Down |
| Sgcg | 7 | 1 | 0,94293453 | 0,00079347 | 0,00506628 | Up |
| Gm15918 | 25 | 1 | 0,63174733 | 0,00061862 | 0,00411474 | Up |
| Nuggc | 872 | 0,89357325 | 0,20668741 | 1,92E-07 | 3,60E-06 | Up |
| Slc39a14 | 6.858 | -0,697730645 | 0,11223597 | 7,67E-12 | 3,50E-10 | Down |
| Piwil2 | 1.040 | -0,987497987 | 0,12853407 | 3,71E-16 | 3,43E-14 | Down |
| Dok2 | 12.680 | -0,673693336 | 0,09103061 | 2,06E-15 | 1,67E-13 | Down |

| | | | | | | |
|---|---|---|---|---|---|---|
| Olfm4 | 17 | -4 | 1 | 4,44E-06 | 5,84E-05 | Down |
| Lmo7 | 9 | -0,60030795 | 0,78157747 | 0,00844428 | 0,03534644 | Down |
| Acod1 | 38 | -2 | 0,4604674 | 2,85E-07 | 5,12E-06 | Down |
| Dzip1 | 190 | 2 | 0,19434867 | 1,10E-16 | 1,11E-14 | Up |
| Farp1 | 33 | -1 | 0,54657633 | 0,0003855 | 0,00275701 | Down |
| Mtmr12 | 10.351 | 0,702403125 | 0,09310391 | 7,96E-16 | 6,94E-14 | Up |
| Pdzd2 | 426 | 2 | 0,30862831 | 3,41E-09 | 9,50E-08 | Up |
| Gm2606 | 51 | 1 | 0,92391372 | 0,00145413 | 0,00837046 | Up |
| Basp1 | 21 | 0,886291761 | 0,83027331 | 0,002606 | 0,01357005 | Up |
| Trio | 13.128 | -0,863331527 | 0,07454405 | 2,00E-32 | 1,24E-29 | Down |
| Gm3362 | 71 | -0,587081731 | 0,24520364 | 0,00025706 | 0,00193781 | Down |
| Lrp12 | 26.105 | -1 | 0,15157969 | 2,22E-13 | 1,38E-11 | Down |
| Nov | 33 | -3 | 0,632821 | 9,21E-07 | 1,48E-05 | Down |
| Gsdmc4 | 21 | -0,876554099 | 0,49767167 | 0,00058398 | 0,00391385 | Down |
| Sla | 19.085 | 1 | 0,15493415 | 1,36E-14 | 9,96E-13 | Up |
| Ndrg1 | 5.030 | -0,9513611 | 0,15812882 | 3,61E-11 | 1,47E-09 | Down |
| St3gal1 | 9.613 | 0,860880672 | 0,11422458 | 1,37E-15 | 1,14E-13 | Up |
| Ptp4a3 | 19.190 | -0,692399835 | 0,13869333 | 7,68E-09 | 1,99E-07 | Down |
| Ly6a | 28 | -1 | 0,83839651 | 0,00037708 | 0,00270598 | Down |
| Ly6g | 18 | -4 | 1 | 2,83E-06 | 3,92E-05 | Down |
| 9030619P08Rik | 3.853 | 2 | 0,16138143 | 1,31E-24 | 3,32E-22 | Up |
| Il2rb | 2.124 | -2 | 0,15500311 | 1,04E-33 | 7,55E-31 | Down |
| Mfng | 9.837 | -0,605733178 | 0,08949922 | 1,55E-13 | 9,81E-12 | Down |
| Maff | 560 | -2 | 0,16576005 | 3,17E-23 | 7,46E-21 | Down |
| Pdgfb | 23 | -0,59553339 | 0,49965234 | 0,00433629 | 0,02053998 | Down |
| Syngr1 | 1.759 | 0,636663199 | 0,08170988 | 8,76E-17 | 8,89E-15 | Up |
| Grap2 | 45 | 2 | 0,53764334 | 1,10E-05 | 0,00012848 | Up |
| Parvb | 447 | -2 | 0,30660762 | 3,39E-09 | 9,46E-08 | Down |
| 5031439G07Rik | 24.803 | 0,799840183 | 0,15199542 | 2,35E-09 | 6,79E-08 | Up |
| Odf3b | 398 | 0,837488701 | 0,14498722 | 1,51E-10 | 5,47E-09 | Up |
| Shank3 | 692 | -2 | 0,27038446 | 3,63E-11 | 1,47E-09 | Down |
| Prickle1 | 609 | 0,931642764 | 0,24120954 | 1,13E-06 | 1,77E-05 | Up |
| Rnd1 | 1.087 | -0,588811681 | 0,2177632 | 9,93E-05 | 0,00086126 | Down |
| Mettl7a1 | 3.337 | 1 | 0,2394024 | 2,33E-11 | 9,87E-10 | Up |
| Galnt6 | 13.781 | -1 | 0,09643067 | 2,32E-29 | 9,94E-27 | Down |
| Acvrl1 | 417 | -4 | 0,44055827 | 7,42E-19 | 9,55E-17 | Down |
| Csad | 543 | 0,798739092 | 0,1285623 | 1,07E-11 | 4,76E-10 | Up |
| Rarg | 671 | -0,604264423 | 0,20448819 | 4,15E-05 | 0,00040503 | Down |
| Calcoco1 | 1.748 | 0,642681077 | 0,16950163 | 1,74E-06 | 2,56E-05 | Up |
| Gpr84 | 315 | -3 | 0,30492453 | 2,01E-19 | 2,91E-17 | Down |
| Zfp385a | 4.709 | 1 | 0,15453175 | 1,16E-16 | 1,16E-14 | Up |
| Coro7 | 8.024 | -0,829264158 | 0,13907888 | 5,09E-11 | 2,00E-09 | Down |
| Ciita | 35 | -0,758327357 | 0,51138216 | 0,00144251 | 0,00831857 | Down |
| Socs1 | 379 | -2 | 0,18276976 | 1,20E-23 | 2,91E-21 | Down |

| | | | | | | |
|---|---|---|---|---|---|---|
| Snx29 | 529 | 0,738131566 | 0,18391033 | 6,91E-07 | 1,14E-05 | Up |
| Mkl2 | 9.248 | 0,796392124 | 0,12561797 | 4,83E-12 | 2,31E-10 | Up |
| Myh11 | 104 | -2 | 0,85394157 | 0,00010786 | 0,00092468 | Down |
| Cebpd | 1.387 | -0,833228327 | 0,13178248 | 5,74E-12 | 2,69E-10 | Down |
| Vpreb1 | 22 | 0,782787813 | 0,56157295 | 0,00164765 | 0,00930905 | Up |
| Rimbp3 | 353 | 0,762979064 | 0,16757342 | 6,96E-08 | 1,45E-06 | Up |
| Vpreb2 | 4 | 0,720013853 | 1 | 0,00554279 | 0,02499558 | Up |
| Etv5 | 5.632 | -2 | 0,17700804 | 7,54E-25 | 1,97E-22 | Down |
| St6gal1 | 834 | -3 | 0,4990534 | 4,80E-12 | 2,31E-10 | Down |
| Nrros | 21.693 | -0,66334897 | 0,0835311 | 2,99E-17 | 3,23E-15 | Down |
| Gm15657 | 116 | 0,691367627 | 0,31482817 | 0,00031576 | 0,00232377 | Up |
| Hacd2 | 5.490 | -0,598471968 | 0,1518164 | 9,87E-07 | 1,57E-05 | Down |
| Stfa2l1 | 107 | -6 | 0,61649707 | 3,79E-26 | 1,15E-23 | Down |
| Gm5483 | 453 | -7 | 0,67578949 | 7,25E-27 | 2,43E-24 | Down |
| Gm5416 | 23 | -5 | 1 | 7,70E-08 | 1,57E-06 | Down |
| 2010005H15Rik | 295 | -6 | 0,71973831 | 7,14E-21 | 1,33E-18 | Down |
| Stfa1 | 435 | -6 | 0,77327099 | 1,44E-18 | 1,80E-16 | Down |
| BC117090 | 5 | -2 | 2 | 0,00074388 | 0,00479577 | Down |
| BC100530 | 158 | -7 | 0,84038559 | 1,22E-17 | 1,39E-15 | Down |
| Stfa2 | 67 | -9 | 1 | 1,05E-15 | 8,88E-14 | Down |
| Stfa3 | 23 | -7 | 1 | 9,36E-11 | 3,50E-09 | Down |
| Cd80 | 94 | -3 | 0,31236351 | 2,08E-19 | 2,97E-17 | Down |
| B4galt4 | 217 | -0,71621596 | 0,15389976 | 4,15E-08 | 9,17E-07 | Down |
| Sidt1 | 14 | 0,583830871 | 0,5453638 | 0,00562692 | 0,02528293 | Up |
| Cd200r2 | 31 | -0,629764023 | 0,67997988 | 0,0061759 | 0,0273857 | Down |
| Zbtb11os1 | 261 | 0,583203079 | 0,23148105 | 0,00017973 | 0,00142956 | Up |
| Gpr15 | 255 | -0,976807108 | 0,33765629 | 2,57E-05 | 0,00026877 | Down |
| Gm9027 | 91 | -0,596648871 | 0,23119014 | 0,00014196 | 0,0011674 | Down |
| Krtap13 | 7 | -0,608599468 | 0,73088296 | 0,00759155 | 0,03246286 | Down |
| 2310034C09Rik | 10 | -0,821249296 | 0,77957532 | 0,00304524 | 0,01540105 | Down |
| Tmem50b | 3.533 | -0,612728452 | 0,13397684 | 5,64E-08 | 1,20E-06 | Down |
| 1600002D24Rik | 11 | -0,696699147 | 0,61644513 | 0,00353231 | 0,01736919 | Down |
| Brwd1 | 12 | 2 | 2 | 0,00032287 | 0,00236878 | Up |
| Fam3b | 42 | -4 | 0,60387031 | 6,11E-11 | 2,37E-09 | Down |
| Tmem181a | 244 | -0,941482903 | 0,21121868 | 1,05E-07 | 2,09E-06 | Down |
| Tmem181b-ps | 217 | -0,904348172 | 0,32319925 | 3,76E-05 | 0,00037269 | Down |
| Sytl3 | 34 | -2 | 0,46716858 | 2,48E-07 | 4,50E-06 | Down |
| Rsph3b | 10 | 0,790583223 | 0,68792006 | 0,00270026 | 0,01399033 | Up |
| Gm16702 | 10 | -0,976926716 | 0,78567422 | 0,00157398 | 0,00894112 | Down |
| Igf2r | 1.450 | -3 | 0,43876894 | 5,23E-10 | 1,71E-08 | Down |
| Abca3 | 76 | 0,652481465 | 0,31131679 | 0,0004648 | 0,00323564 | Up |
| D330041H03Rik | 116 | 0,804834298 | 0,29949038 | 6,15E-05 | 0,00056834 | Up |
| Eci1 | 2.386 | 0,590255263 | 0,1018596 | 7,97E-11 | 3,04E-09 | Up |
| Slc9a3r2 | 18 | -2 | 0,54336274 | 1,98E-05 | 0,00021469 | Down |

| | | | | | | |
|---|---|---|---|---|---|---|
| Igfals | 347 | 1 | 0,24213895 | 2,91E-10 | 9,92E-09 | Up |
| Tpsab1 | 9 | -3 | 1 | 0,000101 | 0,0008749 | Down |
| Fbxl16 | 36 | -2 | 1 | 0,00023569 | 0,00180023 | Down |
| Wdr90 | 3.891 | 0,634732816 | 0,18646697 | 7,87E-06 | 9,69E-05 | Up |
| Rab11fip3 | 208 | 1 | 0,26955093 | 6,85E-08 | 1,43E-06 | Up |
| Syngap1 | 325 | 2 | 0,81886018 | 0,0001967 | 0,00155056 | Up |
| Pacsin1 | 99 | -0,999520476 | 0,49030201 | 0,00024313 | 0,00184522 | Down |
| Cdkn1a | 27.393 | -0,97731001 | 0,13621595 | 1,69E-14 | 1,22E-12 | Down |
| Pim1 | 35.422 | -0,886976214 | 0,10195422 | 1,08E-19 | 1,64E-17 | Down |
| 1700097N02Rik | 188 | 2 | 0,36841296 | 5,82E-09 | 1,55E-07 | Up |
| Dnah8 | 446 | 1 | 0,28506836 | 2,05E-09 | 6,01E-08 | Up |
| Umodl1 | 10 | 2 | 1 | 0,00069375 | 0,00453157 | Up |
| Ubash3a | 5.057 | -0,717046894 | 0,35948253 | 0,00049124 | 0,00338864 | Down |
| Cyp4f40 | 5 | -2 | 2 | 0,00025782 | 0,00194266 | Down |
| Myo1f | 3.980 | -0,669920669 | 0,20752809 | 1,40E-05 | 0,00015875 | Down |
| Psmb9 | 6.708 | 0,736667143 | 0,15204441 | 1,72E-08 | 4,13E-07 | Up |
| Tap1 | 9.559 | 0,781834918 | 0,13700378 | 2,05E-10 | 7,19E-09 | Up |
| Tap2 | 24.397 | 0,864910262 | 0,15663904 | 6,31E-10 | 2,03E-08 | Up |
| Gm15821 | 68 | 2 | 0,51443066 | 1,10E-06 | 1,73E-05 | Up |
| H2-Ob | 189 | 1 | 0,41231027 | 7,30E-06 | 9,05E-05 | Up |
| Btnl2 | 13 | -4 | 2 | 3,08E-05 | 0,00031293 | Down |
| Vwa7 | 200 | 0,592243166 | 0,18038395 | 1,35E-05 | 0,00015392 | Up |
| Lst1 | 1.435 | -0,768530615 | 0,22546426 | 6,62E-06 | 8,31E-05 | Down |
| Ltb | 2.723 | -2 | 0,33139235 | 4,76E-08 | 1,04E-06 | Down |
| Tnf | 576 | -2 | 0,38616104 | 2,36E-09 | 6,80E-08 | Down |
| Gm20443 | 18 | 2 | 0,54231741 | 2,79E-06 | 3,87E-05 | Up |
| H2-M3 | 533 | -0,814209474 | 0,27081439 | 2,33E-05 | 0,00024679 | Down |
| Crisp1 | 11 | -3 | 2 | 5,03E-05 | 0,00048023 | Down |
| A530064D06Rik | 15 | -3 | 0,8945869 | 1,41E-06 | 2,13E-05 | Down |
| Trem3 | 390 | -3 | 0,27436157 | 4,02E-28 | 1,56E-25 | Down |
| Treml4 | 45 | -2 | 0,43726385 | 3,45E-08 | 7,80E-07 | Down |
| Trem2 | 70 | -1 | 0,97073686 | 0,00154058 | 0,00876626 | Down |
| Apobec2 | 31 | 2 | 0,39052035 | 5,84E-08 | 1,24E-06 | Up |
| Rftn1 | 5 | 2 | 2 | 0,0007058 | 0,00459951 | Up |
| Stap2 | 1.615 | -2 | 0,09903891 | 6,93E-55 | 2,32E-51 | Down |
| Mpnd | 8.048 | -0,905124642 | 0,10103755 | 1,07E-20 | 1,91E-18 | Down |
| Kdm4b | 2.232 | 0,581440163 | 0,12335799 | 2,93E-08 | 6,70E-07 | Up |
| Vmac | 531 | 0,726553603 | 0,13464744 | 1,00E-09 | 3,09E-08 | Up |
| Rfx2 | 198 | 2 | 0,22514328 | 5,24E-17 | 5,45E-15 | Up |
| Tnfsf14 | 2.355 | -0,896171082 | 0,1381659 | 2,09E-12 | 1,08E-10 | Down |
| C3 | 3.833 | -3 | 0,37819806 | 9,96E-14 | 6,51E-12 | Down |
| Adgre1 | 31.351 | -2 | 0,16152288 | 1,57E-43 | 1,88E-40 | Down |
| Emilin2 | 72.153 | -0,624291591 | 0,18142581 | 6,98E-06 | 8,69E-05 | Down |
| Lbh | 2.209 | 0,650792142 | 0,16122029 | 6,27E-07 | 1,05E-05 | Up |

| | | | | | | |
|---|---|---|---|---|---|---|
| Xdh | 29 | 1 | 0,71964725 | 0,00057369 | 0,00384954 | Up |
| Rasgrp3 | 129 | -0,754227407 | 0,81799072 | 0,00444595 | 0,0209445 | Down |
| Prkd3 | 7 | -0,614562057 | 0,93648822 | 0,00858057 | 0,03577377 | Down |
| Map4k3 | 32 | 2 | 0,41863965 | 2,05E-06 | 2,95E-05 | Up |
| Zfp36l2 | 36.259 | 0,611048575 | 0,10504546 | 7,07E-11 | 2,70E-09 | Up |
| Dync2li1 | 42 | 1 | 0,98194035 | 0,00090294 | 0,00564462 | Up |
| 1110020A21Rik | 81 | 0,942233704 | 0,29084305 | 9,41E-06 | 0,00011316 | Up |
| Rhoq | 4.659 | -0,766804893 | 0,1524769 | 7,30E-09 | 1,91E-07 | Down |
| Mcfd2 | 6.635 | -0,847044354 | 0,13391054 | 5,73E-12 | 2,69E-10 | Down |
| 4833418N02Rik | 78 | -0,948108892 | 0,26481167 | 3,04E-06 | 4,16E-05 | Down |
| Ppp1r21 | 390 | -0,723287611 | 0,5781302 | 0,00258534 | 0,01348683 | Down |
| Cables1 | 447 | 2 | 0,256974 | 2,68E-11 | 1,10E-09 | Up |
| Lama3 | 18 | -0,989716843 | 0,74048336 | 0,00126688 | 0,00748367 | Down |
| Cabyr | 88 | 0,835733636 | 0,35597448 | 0,00014741 | 0,00120515 | Up |
| Slc39a6 | 7.024 | -0,906279664 | 0,08796103 | 2,42E-26 | 7,64E-24 | Down |
| Celf4 | 147 | -3 | 0,59650858 | 6,87E-09 | 1,81E-07 | Down |
| Bin1 | 1.946 | 0,707230206 | 0,13577686 | 2,59E-09 | 7,40E-08 | Up |
| Gypc | 10.769 | 0,618356868 | 0,10879136 | 1,56E-10 | 5,64E-09 | Up |
| Cxxc5 | 8.768 | 0,660360625 | 0,09958417 | 4,56E-13 | 2,68E-11 | Up |
| Hbegf | 1.402 | -2 | 0,15321349 | 7,56E-37 | 7,91E-34 | Down |
| Cd14 | 160 | 0,676325117 | 0,27506642 | 0,00015957 | 0,00129007 | Up |
| Spry4 | 227 | -3 | 0,26601824 | 4,33E-35 | 3,62E-32 | Down |
| Ap3s1 | 11.592 | -0,608438073 | 0,1658532 | 2,98E-06 | 4,11E-05 | Down |
| Arl14epl | 215 | -1 | 0,2844761 | 3,62E-07 | 6,41E-06 | Down |
| Cd74 | 1.766 | -3 | 0,45126403 | 3,90E-13 | 2,33E-11 | Down |
| Camk2a | 29 | -0,613398734 | 0,42199075 | 0,00242591 | 0,01282301 | Down |
| Csf1r | 782 | -1 | 0,40489721 | 3,37E-05 | 0,00033928 | Down |
| Afap1l1 | 2.169 | -0,809108106 | 0,14062244 | 1,65E-10 | 5,95E-09 | Down |
| Tubb6 | 18.990 | -0,774315511 | 0,07455933 | 8,76E-27 | 2,87E-24 | Down |
| Mc5r | 1.407 | 2 | 0,31178109 | 3,33E-12 | 1,66E-10 | Up |
| Smad7 | 2.163 | 0,874620968 | 0,15218738 | 1,82E-10 | 6,45E-09 | Up |
| Katnal2 | 143 | 0,593492714 | 0,16269598 | 3,34E-06 | 4,52E-05 | Up |
| A330094K24Rik | 13 | 2 | 0,68611029 | 8,55E-06 | 0,00010418 | Up |
| Tshz1 | 11.997 | 0,587801773 | 0,09401207 | 4,69E-12 | 2,27E-10 | Up |
| Aldh3b1 | 123 | -2 | 0,36724301 | 1,07E-08 | 2,68E-07 | Down |
| Unc93b1 | 4.615 | -0,86303926 | 0,15806839 | 8,79E-10 | 2,76E-08 | Down |
| Pcx | 5.514 | 0,645185394 | 0,15843796 | 5,40E-07 | 9,20E-06 | Up |
| Tsga10ip | 1.121 | 0,718573058 | 0,14334152 | 7,26E-09 | 1,90E-07 | Up |
| Cdc42bpg | 103 | 0,663596874 | 0,30866066 | 0,00038879 | 0,00277578 | Up |
| Pygm | 82 | 3 | 0,43676042 | 3,56E-10 | 1,19E-08 | Up |
| Slc22a12 | 11 | 2 | 0,99285276 | 0,00038023 | 0,00272276 | Up |
| Gm6445 | 648 | -0,783340881 | 0,24899618 | 1,57E-05 | 0,00017554 | Down |
| Cd5 | 7 | 0,779080535 | 0,82592338 | 0,00405237 | 0,01945951 | Up |
| Slc15a3 | 18 | -3 | 1 | 2,29E-05 | 0,00024304 | Down |

| | | | | | | |
|---|---|---|---|---|---|---|
| Ms4a6b | 21 | 1 | 0,88081431 | 0,00142774 | 0,0082476 | Up |
| Ms4a2 | 8 | -3 | 1 | 3,58E-05 | 0,00035704 | Down |
| Ms4a3 | 1.878 | -4 | 0,45572877 | 1,07E-18 | 1,35E-16 | Down |
| Oosp1 | 5 | -3 | 2 | 0,00017474 | 0,0013945 | Down |
| Pfpl | 23 | -2 | 0,70223237 | 2,27E-05 | 0,00024135 | Down |
| Mpeg1 | 526 | -0,857132461 | 0,27621378 | 1,62E-05 | 0,00018063 | Down |
| Glyat | 7 | -3 | 2 | 0,00018222 | 0,00144799 | Down |
| Gna14 | 108 | 1 | 0,48178554 | 5,25E-05 | 0,00049753 | Up |
| Trpm6 | 13 | -3 | 0,95741116 | 2,32E-06 | 3,30E-05 | Down |
| Mamdc2 | 17 | -1 | 0,75750182 | 0,00024956 | 0,00188975 | Down |
| Kank1 | 118 | -3 | 0,30853833 | 6,08E-23 | 1,41E-20 | Down |
| Slc1a1 | 78 | -3 | 0,9104474 | 2,09E-06 | 3,00E-05 | Down |
| 4430402I18Rik | 90 | 0,996057369 | 0,2498303 | 6,63E-07 | 1,10E-05 | Up |
| Cd274 | 1.914 | -0,673516538 | 0,11184954 | 2,36E-11 | 9,98E-10 | Down |
| Pdcd1lg2 | 272 | -1 | 0,29269794 | 6,68E-09 | 1,76E-07 | Down |
| Ankrd22 | 50 | -3 | 0,72395026 | 5,42E-07 | 9,21E-06 | Down |
| Stambpl1 | 4.520 | -0,62606922 | 0,06871155 | 1,07E-21 | 2,16E-19 | Down |
| Acta2 | 609 | -0,588665536 | 0,22463435 | 0,00012912 | 0,00107668 | Down |
| Ifit1 | 35 | -1 | 0,52486417 | 0,00031289 | 0,00230567 | Down |
| Pdlim1 | 4 | -3 | 2 | 0,00013238 | 0,00109942 | Down |
| Frat2 | 3.582 | 0,996358018 | 0,15036929 | 7,10E-13 | 4,05E-11 | Up |
| Cpn1 | 84 | 1 | 0,22584942 | 2,15E-09 | 6,25E-08 | Up |
| Obfc1 | 1.183 | 0,65296955 | 0,10705539 | 1,38E-11 | 6,04E-10 | Up |
| Gm19557 | 54 | 0,702789668 | 0,41209755 | 0,00102597 | 0,00628584 | Up |
| Pdcd4 | 1.961 | 1 | 0,17317571 | 8,65E-11 | 3,27E-09 | Up |
| Ablim1 | 17.996 | -1 | 0,1866049 | 2,84E-11 | 1,16E-09 | Down |
| Shtn1 | 51 | 0,599667988 | 0,40426864 | 0,00240492 | 0,01272006 | Up |
| Emx2os | 21 | 6 | 1 | 2,09E-08 | 4,95E-07 | Up |
| Emx2 | 40 | 7 | 2 | 5,59E-08 | 1,19E-06 | Up |
| Csf2ra | 800 | -0,823117757 | 0,31506948 | 7,34E-05 | 0,00066419 | Down |
| Ccdc120 | 1.294 | 0,720070615 | 0,09538786 | 8,35E-16 | 7,24E-14 | Up |
| Pim2 | 2.848 | -0,95646088 | 0,1162635 | 5,28E-18 | 6,27E-16 | Down |
| Glod5 | 23 | 3 | 0,51132375 | 3,22E-09 | 9,06E-08 | Up |
| Cfp | 7.659 | -2 | 0,2703938 | 3,61E-20 | 5,92E-18 | Down |
| Gm4764 | 16 | -2 | 0,66828252 | 1,11E-05 | 0,00012944 | Down |
| Gm7598 | 25 | -0,605405305 | 0,88743308 | 0,0088409 | 0,03666109 | Down |
| Ap3s1-ps1 | 33 | -1 | 0,4804549 | 9,96E-05 | 0,00086366 | Down |
| Gria3 | 444 | 0,9269923 | 0,76761843 | 0,00183942 | 0,01019296 | Up |
| Plac1 | 8 | -3 | 1 | 1,89E-05 | 0,00020624 | Down |
| 3830403N18Rik | 1.450 | -1 | 0,22594949 | 7,95E-09 | 2,05E-07 | Down |
| Gm364 | 51 | -1 | 0,43899397 | 1,37E-05 | 0,00015578 | Down |
| L1cam | 1.191 | -0,877860759 | 0,27206322 | 1,07E-05 | 0,00012578 | Down |
| 4933407K13Rik | 226 | 1 | 0,43086857 | 0,00010232 | 0,00088451 | Up |
| Gm14760 | 51 | -0,625487054 | 0,28922009 | 0,00042747 | 0,00300826 | Down |

| | | | | | | |
|---|---|---|---|---|---|---|
| Pcyt1b | 238 | 0,857754149 | 0,2877259 | 2,36E-05 | 0,00024868 | Up |
| F630028O10Rik | 29 | -1 | 1 | 0,00201914 | 0,01099945 | Down |
| Eda2r | 159 | -1 | 0,24384595 | 1,16E-10 | 4,26E-09 | Down |
| Nlgn3 | 7 | -1 | 0,97019071 | 0,001518 | 0,00865838 | Down |
| Nhsl2 | 4.186 | -1 | 0,1526495 | 3,40E-13 | 2,05E-11 | Down |
| C77370 | 8 | 0,68545924 | 0,7912024 | 0,00577282 | 0,02584486 | Up |
| Tlr13 | 26 | -3 | 0,67941527 | 4,40E-07 | 7,64E-06 | Down |
| Hmgn5 | 10.157 | 0,582502289 | 0,06549914 | 6,87E-21 | 1,29E-18 | Up |
| Diaph2 | 7.804 | -1 | 0,21760373 | 1,34E-08 | 3,29E-07 | Down |
| Nox1 | 51 | -0,770966226 | 0,50134184 | 0,00123697 | 0,00734546 | Down |
| AV320801 | 3 | -1 | 2 | 0,00151168 | 0,00863118 | Down |
| Nxf7 | 20 | -1 | 0,81438079 | 0,00119403 | 0,0071339 | Down |
| Trpc5os | 22 | 6 | 2 | 2,28E-07 | 4,19E-06 | Up |
| Mir3473a | 689 | -0,783061866 | 0,1581468 | 1,10E-08 | 2,76E-07 | Down |
| Bmx | 4 | 1 | 2 | 0,00184639 | 0,01022482 | Up |
| Gpm6b | 625 | -1 | 0,22252922 | 3,04E-13 | 1,86E-11 | Down |
| Gm8817 | 5 | -1 | 1 | 0,00199502 | 0,01091194 | Down |
| Arhgap6 | 8.356 | 0,968333904 | 0,219221 | 1,21E-07 | 2,37E-06 | Up |
| mt-Tm | 72 | -0,722485245 | 0,48531394 | 0,00157022 | 0,00892281 | Down |

| **CM-vs-CM-IL3-12** | | | | | | |
|---|---|---|---|---|---|---|
| | **baseMean** | **log2FoldChange** | **lfcSE** | **pvalue** | **padj** | **expression** |
| Efhc1 | 16,97640602 | 0,645447403 | 0,458595671 | 0,001841016 | 0,009357408 | Up |
| Tmem14a | 317,0076962 | 0,842815865 | 0,176599707 | 2,23E-08 | 5,56E-07 | Up |
| Fer1l5 | 587,7243083 | 0,591543711 | 0,110141897 | 2,76E-09 | 9,28E-08 | Up |
| Zap70 | 14,04594533 | 0,609984996 | 0,616289047 | 0,002669754 | 0,012709607 | Up |
| 2010300C02Rik | 49,72523763 | 0,593978889 | 0,379174048 | 0,002125469 | 0,010513945 | Up |
| Aff3 | 4375,805027 | 0,792309487 | 0,235285872 | 8,18E-06 | 9,12E-05 | Up |
| Il18r1 | 6743,735514 | 0,777074731 | 0,113143119 | 1,67E-13 | 1,64E-11 | Up |
| Fhl2 | 24,97946615 | 0,688150326 | 0,494131353 | 0,001455545 | 0,007647739 | Up |
| Wdr75 | 13568,03793 | -0,592372057 | 0,093911101 | 1,06E-11 | 6,84E-10 | Down |
| Gm5526 | 377,8448003 | -0,590816159 | 0,306306559 | 0,001202896 | 0,006520876 | Down |
| Rftn2 | 532,1447096 | 0,775528901 | 0,127339085 | 2,61E-11 | 1,52E-09 | Up |
| Ino80dos | 114,8376389 | 0,643966044 | 0,233326027 | 0,000117686 | 0,00091303 | Up |
| Idh1 | 5830,96915 | 0,701344485 | 0,184417272 | 2,79E-06 | 3,54E-05 | Up |
| Kansl1l | 2153,789982 | 0,735138659 | 0,137886969 | 2,35E-09 | 8,04E-08 | Up |
| Slc11a1 | 107,6252626 | -0,590073018 | 0,685463953 | 0,002639975 | 0,012620386 | Down |
| Stk36 | 69,79589087 | 0,819464168 | 0,279722068 | 2,74E-05 | 0,000264287 | Up |
| Itm2c | 20465,34958 | 0,594817158 | 0,129529175 | 1,44E-07 | 2,75E-06 | Up |
| Rab17 | 452,0251321 | 0,661764066 | 0,303113072 | 0,000451982 | 0,002852358 | Up |
| Ramp1 | 6132,934336 | 0,960941324 | 0,166364124 | 5,09E-11 | 2,75E-09 | Up |
| Hes6 | 2039,661949 | 0,909515709 | 0,1072594 | 2,55E-19 | 6,90E-17 | Up |
| D2hgdh | 1369,304957 | 0,646633107 | 0,093231196 | 1,54E-13 | 1,54E-11 | Up |
| Gm7889 | 93,02605462 | 0,729445663 | 0,208305634 | 7,24E-06 | 8,17E-05 | Up |

| | | | | | | |
|---|---|---|---|---|---|---|
| Gm6086 | 66,53827624 | 0,749527933 | 0,44342701 | 0,000695037 | 0,004112685 | Up |
| Pdcd1 | 212,3635005 | 2,97934959 | 0,625948173 | 1,03E-09 | 3,91E-08 | Up |
| Gm28188 | 183,1912916 | 0,799894467 | 0,273086033 | 3,05E-05 | 0,000289406 | Up |
| Phlpp1 | 2041,35478 | 0,941001371 | 0,155501974 | 1,13E-11 | 7,13E-10 | Up |
| Tmem185b | 9566,073431 | -0,809941691 | 0,083947222 | 1,29E-23 | 8,72E-21 | Down |
| Ddx18 | 21063,86072 | -0,607797035 | 0,085351138 | 4,21E-14 | 4,94E-12 | Down |
| Map3k19 | 32,01209998 | 2,014216012 | 0,44697992 | 1,57E-08 | 4,10E-07 | Up |
| Cxcr4 | 7686,079561 | 0,80758643 | 0,115148738 | 5,10E-14 | 5,95E-12 | Up |
| Btg2 | 1120,205726 | 1,308424745 | 0,161616766 | 8,83E-19 | 2,17E-16 | Up |
| Ppfia4 | 21,39133961 | 0,627975202 | 0,456173782 | 0,002115425 | 0,010473838 | Up |
| Lgr6 | 191,7070607 | 0,625248433 | 0,153470226 | 1,30E-06 | 1,83E-05 | Up |
| Elf3 | 18,41098577 | 0,808768871 | 0,640173194 | 0,000889554 | 0,005060664 | Up |
| Timm17a | 13053,33836 | -0,673717955 | 0,097527247 | 1,80E-13 | 1,74E-11 | Down |
| Gm15850 | 27,76454983 | 0,599732159 | 0,444404664 | 0,00260008 | 0,012444337 | Up |
| Rgs2 | 4394,600716 | 0,767659603 | 0,114689086 | 5,73E-13 | 5,05E-11 | Up |
| Rgs1 | 88,76523314 | 1,664643738 | 0,338358413 | 2,00E-09 | 6,90E-08 | Up |
| Rgl1 | 163,2359952 | 0,904548671 | 0,211042919 | 1,38E-07 | 2,65E-06 | Up |
| Rgsl1 | 16,14664771 | 0,84151401 | 0,576207015 | 0,000627627 | 0,003771571 | Up |
| Mr1 | 595,2540975 | 0,638866912 | 0,18647038 | 1,47E-05 | 0,000152186 | Up |
| Axdnd1 | 17,10151949 | 0,615950282 | 0,472117654 | 0,002428672 | 0,011790905 | Up |
| Nme7 | 497,7444183 | 1,025754234 | 0,200764007 | 1,36E-09 | 4,99E-08 | Up |
| Tbx19 | 183,1212648 | 2,065659995 | 0,235031546 | 1,97E-20 | 6,65E-18 | Up |
| Mgst3 | 1096,111012 | 0,635786405 | 0,198187516 | 3,12E-05 | 0,0002945 | Up |
| Dusp12 | 1364,685485 | -0,587019598 | 0,11672998 | 1,71E-08 | 4,41E-07 | Down |
| Fcgr4 | 32,96658856 | -0,599462263 | 0,577381252 | 0,002918829 | 0,013641652 | Down |
| Fcgr3 | 445,2897942 | -2,372547062 | 0,533285893 | 1,23E-08 | 3,32E-07 | Down |
| Cd244 | 2297,6769 | 0,594703163 | 0,160953302 | 6,63E-06 | 7,60E-05 | Up |
| Ly9 | 34,8917493 | 0,728379983 | 0,45647302 | 0,000901062 | 0,00511001 | Up |
| Igsf8 | 11644,53898 | -1,126985207 | 0,137960542 | 7,21E-19 | 1,80E-16 | Down |
| Ifi204 | 148,3465776 | 1,014396717 | 0,256658779 | 3,04E-07 | 5,28E-06 | Up |
| Mndal | 929,3119738 | 1,070651059 | 0,207449087 | 8,14E-10 | 3,17E-08 | Up |
| Ifi211 | 27,8513269 | 0,940809722 | 0,470313351 | 0,000160686 | 0,001190467 | Up |
| Ifi203 | 624,3994017 | 1,349933828 | 0,235199275 | 1,55E-11 | 9,47E-10 | Up |
| Spta1 | 352,3317916 | 2,693161519 | 0,28242551 | 1,46E-24 | 1,13E-21 | Up |
| Olfr417 | 8,060975743 | 0,655993638 | 1,009104036 | 0,001184909 | 0,006444903 | Up |
| Olfr248 | 16,25454946 | 1,037498775 | 0,729535898 | 0,000310904 | 0,002083609 | Up |
| Olfr414 | 16,69962429 | 1,061418899 | 0,786593591 | 0,000309586 | 0,002076494 | Up |
| H3f3aos | 23,41425315 | 1,016087188 | 0,452733078 | 6,97E-05 | 0,000584238 | Up |
| Enah | 45,52905557 | -3,79219294 | 0,724615923 | 1,84E-10 | 8,70E-09 | Down |
| Degs1 | 9673,181637 | -0,881875603 | 0,116210253 | 4,40E-16 | 7,37E-14 | Down |
| Ptpn14 | 111,6487888 | 0,699022239 | 0,393309145 | 0,000796614 | 0,004596394 | Up |
| Slc30a1 | 697,9631187 | -0,64918173 | 0,12739309 | 1,06E-08 | 2,93E-07 | Down |
| Rpp38 | 1625,662356 | -0,624796764 | 0,099313684 | 1,15E-11 | 7,27E-10 | Down |
| Echdc3 | 48,56900599 | 1,236514132 | 0,30939888 | 1,19E-07 | 2,34E-06 | Up |
| Il2ra | 1536,200824 | -0,880551491 | 0,474963658 | 0,000277394 | 0,00189424 | Down |
| Gm13375 | 364,7283636 | 1,011921261 | 0,17226793 | 2,02E-11 | 1,20E-09 | Up |
| Slc34a3 | 164,5404617 | 3,47304878 | 0,456799082 | 1,29E-17 | 2,82E-15 | Up |
| Dpp7 | 766,4621852 | 0,60832478 | 0,198554789 | 5,72E-05 | 0,000493257 | Up |
| Clic3 | 77,12869887 | 0,752031911 | 0,33060168 | 0,000216674 | 0,001532153 | Up |
| Ptgds | 5,043273106 | 1,024834004 | 1,234399139 | 0,000382164 | 0,002471345 | Up |
| Qsox2 | 6293,675937 | -0,677081585 | 0,059449582 | 2,27E-31 | 6,13E-28 | Down |

| | | | | | | |
|---|---|---|---|---|---|---|
| Lcn4 | 258,9932678 | 0,837993526 | 0,397217136 | 0,000199246 | 0,001422872 | Up |
| Stkld1 | 148,4656588 | 0,728281925 | 0,292849694 | 0,000149526 | 0,001118 | Up |
| Spaca9 | 236,2926072 | 1,951818201 | 0,209758234 | 1,78E-22 | 9,01E-20 | Up |
| Ak8 | 47,89398858 | 0,942286195 | 0,440669429 | 0,00011794 | 0,000913398 | Up |
| Cercam | 1490,254638 | 0,655869856 | 0,186379433 | 9,75E-06 | 0,00010665 | Up |
| Zer1 | 3262,2588 | 0,651519451 | 0,107600538 | 4,84E-11 | 2,64E-09 | Up |
| Qrfp | 115,3309365 | 1,203709402 | 0,264663781 | 1,09E-08 | 2,98E-07 | Up |
| Fibcd1 | 34,55131613 | 0,836369704 | 0,456310495 | 0,000351782 | 0,002309858 | Up |
| St6galnac4 | 10520,07696 | -0,633118206 | 0,090304412 | 9,16E-14 | 9,71E-12 | Down |
| Angptl2 | 299,2275307 | -0,871644879 | 0,242511169 | 2,45E-06 | 3,17E-05 | Down |
| Nek6 | 9623,426214 | -0,76235653 | 0,085309854 | 1,33E-20 | 4,58E-18 | Down |
| Gm13502 | 42,7656329 | 0,872701371 | 0,485471464 | 0,000319293 | 0,002124908 | Up |
| Nr4a2 | 1421,172583 | 0,718082395 | 0,14155046 | 9,59E-09 | 2,70E-07 | Up |
| Galnt3 | 6167,877876 | -0,743551881 | 0,086431382 | 2,74E-19 | 7,16E-17 | Down |
| Plekha3 | 4718,743872 | -0,785578626 | 0,089149022 | 3,55E-20 | 1,11E-17 | Down |
| Ube2l6 | 580,9973168 | 0,865041474 | 0,209361056 | 3,28E-07 | 5,62E-06 | Up |
| Timm10 | 2883,452495 | -0,613464262 | 0,12042685 | 1,16E-08 | 3,13E-07 | Down |
| Olfr1250 | 53,95879206 | 0,600028956 | 0,316533672 | 0,00120409 | 0,00652517 | Up |
| Rapsn | 937,8298717 | 0,629167144 | 0,134491771 | 8,74E-08 | 1,79E-06 | Up |
| Accs | 1531,17446 | 0,721153328 | 0,123386096 | 1,42E-10 | 6,89E-09 | Up |
| Nat10 | 18285,97154 | -0,598062759 | 0,097214757 | 2,83E-11 | 1,65E-09 | Down |
| Bmf | 671,855034 | 0,966855229 | 0,162894211 | 1,90E-11 | 1,13E-09 | Up |
| Spint1 | 253,6176392 | 0,936673982 | 0,243893488 | 6,98E-07 | 1,08E-05 | Up |
| Ganc | 2970,092092 | 0,613633271 | 0,119823028 | 1,00E-08 | 2,80E-07 | Up |
| Epb42 | 836,6632083 | 0,742005754 | 0,202581186 | 3,84E-06 | 4,70E-05 | Up |
| Tgm5 | 10,7156236 | 0,610829801 | 0,717069912 | 0,002281631 | 0,011170627 | Up |
| Ckmt1 | 1388,774942 | 0,692706361 | 0,15905946 | 3,07E-07 | 5,32E-06 | Up |
| Gm14002 | 33,59088708 | 1,378585924 | 0,403324564 | 9,12E-07 | 1,36E-05 | Up |
| Dusp2 | 2214,838787 | -0,669408959 | 0,131761745 | 1,08E-08 | 2,98E-07 | Down |
| Ttl | 207,8065823 | 0,686482427 | 0,14965809 | 1,12E-07 | 2,22E-06 | Up |
| Slc20a1 | 10384,98151 | -0,708555572 | 0,088046629 | 3,21E-17 | 6,61E-15 | Down |
| Fastkd5 | 1720,723459 | -0,587589136 | 0,097724359 | 6,75E-11 | 3,57E-09 | Down |
| Ap5s1 | 565,1535008 | 1,045519142 | 0,161435568 | 3,62E-13 | 3,30E-11 | Up |
| Trmt6 | 8511,311653 | -0,588513141 | 0,110873604 | 3,91E-09 | 1,24E-07 | Down |
| Macrod2 | 256,2804725 | 0,597890871 | 0,210879083 | 0,000122292 | 0,000943956 | Up |
| Id1 | 13342,87747 | -1,751695601 | 0,136332296 | 8,69E-40 | 4,70E-36 | Down |
| Cox4i2 | 37,51901582 | -0,726852555 | 0,320577933 | 0,000254245 | 0,00176212 | Down |
| Bcl2l1 | 12015,67863 | -0,890544187 | 0,103895923 | 1,36E-19 | 3,86E-17 | Down |
| Pxmp4 | 2668,970236 | 0,848087149 | 0,176695494 | 1,90E-08 | 4,82E-07 | Up |
| Ggt7 | 101,5361539 | 0,928424179 | 0,415389061 | 9,98E-05 | 0,000796537 | Up |
| Acss2 | 1823,739378 | 0,669917415 | 0,203474468 | 1,99E-05 | 0,000198268 | Up |
| Sla2 | 10507,01329 | 0,666455702 | 0,169093706 | 1,90E-06 | 2,54E-05 | Up |
| Mroh8 | 31,87457383 | 0,936130445 | 0,41227297 | 8,89E-05 | 0,000720395 | Up |
| D630003M21Rik | 639,0190827 | 1,266007427 | 0,245347887 | 4,32E-10 | 1,83E-08 | Up |
| Fitm2 | 2813,975891 | -0,610626464 | 0,089414151 | 3,29E-13 | 3,01E-11 | Down |
| Sulf2 | 19,01417355 | 1,4258727 | 0,655271471 | 2,95E-05 | 0,000281423 | Up |
| Gm11473 | 149,3113433 | -1,361740039 | 0,251879239 | 1,06E-10 | 5,34E-09 | Down |
| Fam65c | 123,4731344 | 0,844681361 | 0,243108276 | 4,30E-06 | 5,14E-05 | Up |
| Gm14236 | 22,32851066 | 1,397607397 | 0,563375571 | 1,51E-05 | 0,000156326 | Up |
| Kcng1 | 18,44932235 | 0,896957895 | 0,723190136 | 0,000617742 | 0,003719058 | Up |
| Atp9a | 443,3186064 | 2,068713783 | 0,191901407 | 7,13E-29 | 1,45E-25 | Up |

| | | | | | | |
|---|---|---|---|---|---|---|
| Pfdn4 | 3611,725278 | -0,668760497 | 0,093371206 | 3,01E-14 | 3,67E-12 | Down |
| Ppp4r1l-ps | 511,7770638 | 0,66091885 | 0,19287021 | 1,32E-05 | 0,000138783 | Up |
| Zfp831 | 679,8420958 | 0,925390578 | 0,20222597 | 3,38E-08 | 7,97E-07 | Up |
| Edn3 | 84,35801001 | 1,478304432 | 0,233807443 | 5,07E-13 | 4,52E-11 | Up |
| Col9a3 | 253,833036 | 1,222037099 | 0,287897959 | 4,20E-08 | 9,59E-07 | Up |
| E2f5 | 524,6423764 | -0,62767118 | 0,139674406 | 2,07E-07 | 3,77E-06 | Down |
| Cpa3 | 212,7372238 | 0,872759752 | 0,346715669 | 6,43E-05 | 0,00054611 | Up |
| Bbs12 | 659,7257054 | -0,710720249 | 0,194075232 | 4,50E-06 | 5,37E-05 | Down |
| 1700017G19Rik | 81,01020673 | 1,101477312 | 0,247006064 | 2,31E-08 | 5,71E-07 | Up |
| Gm2011 | 65,66743088 | 1,127196227 | 0,236300955 | 4,71E-09 | 1,45E-07 | Up |
| 5031434O11Rik | 165,0372201 | 0,737529372 | 0,168467225 | 2,30E-07 | 4,10E-06 | Up |
| Mgst2 | 3909,610169 | 0,827274916 | 0,197927574 | 3,34E-07 | 5,69E-06 | Up |
| P2ry14 | 6297,454053 | -0,851336507 | 0,199145874 | 1,96E-07 | 3,61E-06 | Down |
| Gm5540 | 29,31223885 | 1,032130674 | 0,451873448 | 5,99E-05 | 0,000513715 | Up |
| Lxn | 1989,428591 | 0,685157417 | 0,113822148 | 5,61E-11 | 3,00E-09 | Up |
| Ppid | 11918,65474 | -0,616469418 | 0,143277966 | 5,04E-07 | 8,15E-06 | Down |
| Ctso | 607,02333 | 0,605621958 | 0,206188862 | 8,62E-05 | 0,000701191 | Up |
| Sh2d2a | 1282,547855 | 0,661005529 | 0,153070761 | 4,05E-07 | 6,78E-06 | Up |
| Gpatch4 | 8405,066357 | -0,581881313 | 0,101739379 | 3,91E-10 | 1,67E-08 | Down |
| Pmvk | 4873,250219 | 0,594570896 | 0,132377896 | 2,30E-07 | 4,10E-06 | Up |
| Il6ra | 221,0480683 | -0,723653729 | 0,177859403 | 8,91E-07 | 1,33E-05 | Down |
| Selenbp2 | 107,7579436 | 1,281029729 | 0,330236784 | 1,78E-07 | 3,31E-06 | Up |
| Selenbp1 | 1431,108593 | 1,354808228 | 0,264084179 | 4,75E-10 | 1,99E-08 | Up |
| 4930481B07Rik | 48,0674704 | 0,699186557 | 0,293531636 | 0,000228703 | 0,001607059 | Up |
| Gm4349 | 608,645391 | 0,614193429 | 0,169340075 | 7,91E-06 | 8,85E-05 | Up |
| Hist2h2be | 164,9131176 | 0,584320076 | 0,184298247 | 4,56E-05 | 0,000406266 | Up |
| Fcgr1 | 5289,287716 | -0,679590012 | 0,148994485 | 1,29E-07 | 2,51E-06 | Down |
| Pias3 | 2678,101871 | 0,642957715 | 0,084490283 | 1,11E-15 | 1,78E-13 | Up |
| Ptpn22 | 2734,746155 | 0,701946119 | 0,152554195 | 9,87E-08 | 1,98E-06 | Up |
| Cd53 | 2647,867443 | -1,02818766 | 0,62930081 | 0,000237537 | 0,001661448 | Down |
| Bank1 | 390,3070533 | -0,657901309 | 0,270599814 | 0,000259359 | 0,001790679 | Down |
| Dnajb4 | 3052,467561 | 0,718617844 | 0,15090882 | 4,38E-08 | 9,94E-07 | Up |
| Cyp7a1 | 157,4428304 | 0,675714812 | 0,216150243 | 3,31E-05 | 0,00031018 | Up |
| Gem | 254,5110708 | -5,651208963 | 1,484901548 | 1,30E-07 | 2,53E-06 | Down |
| Cdh17 | 137,6528549 | -6,128922189 | 0,910282451 | 8,54E-13 | 7,18E-11 | Down |
| Pdp1 | 2184,299773 | -1,969095897 | 0,220525083 | 5,50E-21 | 2,08E-18 | Down |
| Ndufaf4 | 4332,278384 | -0,667484949 | 0,121057757 | 1,09E-09 | 4,11E-08 | Down |
| Mdn1 | 16966,43841 | -0,795639608 | 0,207631413 | 1,56E-06 | 2,15E-05 | Down |
| 4933421O10Rik | 340,2676421 | 0,715456995 | 0,154747556 | 8,46E-08 | 1,75E-06 | Up |
| Pnrc1 | 2319,610728 | 0,728751375 | 0,202880424 | 5,29E-06 | 6,21E-05 | Up |
| Cfap206 | 10,11146459 | 1,297095352 | 0,912196696 | 0,000143382 | 0,001080524 | Up |
| Dnaic1 | 45,39712919 | 0,632986913 | 0,376528606 | 0,001394697 | 0,007383023 | Up |
| Cd72 | 402,1646575 | 1,080734708 | 0,160615012 | 5,31E-14 | 6,07E-12 | Up |
| Hint2 | 2775,708215 | 0,58790065 | 0,089638853 | 2,07E-12 | 1,60E-10 | Up |
| Tmem8b | 62,56912016 | 0,738568788 | 0,380029394 | 0,000469876 | 0,002945803 | Up |
| Ptgr1 | 4306,690766 | 0,688418371 | 0,133231149 | 6,57E-09 | 1,92E-07 | Up |
| Gm12527 | 24,34409068 | -0,634377026 | 0,379773429 | 0,001405047 | 0,00742328 | Down |
| Tnfsf8 | 2093,575493 | -1,182845256 | 0,133503967 | 1,44E-21 | 6,33E-19 | Down |
| C630043F03Rik | 328,7396824 | -0,665687939 | 0,200334283 | 1,83E-05 | 0,000184911 | Down |
| Hacd4 | 6567,581851 | 0,867751324 | 0,234674787 | 1,73E-06 | 2,35E-05 | Up |
| Mtap | 12636,51083 | -0,612759018 | 0,13180897 | 1,05E-07 | 2,10E-06 | Down |

| | | | | | | |
|---|---|---|---|---|---|---|
| Elavl2 | 986,5023764 | -0,874351098 | 0,117063838 | 1,16E-15 | 1,84E-13 | Down |
| Nfia | 299,0884067 | 0,751853672 | 0,189330937 | 1,15E-06 | 1,65E-05 | Up |
| Lepr | 34,45425309 | -0,857729813 | 0,470812609 | 0,000325916 | 0,002164548 | Down |
| Pde4b | 1526,193179 | 0,843108921 | 0,129883493 | 1,40E-12 | 1,13E-10 | Up |
| Glis1 | 22,5868593 | -0,693341009 | 0,451619077 | 0,001191522 | 0,006472193 | Down |
| Lrp8 | 9688,957132 | -0,684797902 | 0,083116594 | 7,05E-18 | 1,57E-15 | Down |
| Ttc39a | 37,99889944 | 0,663994857 | 0,392496398 | 0,001129518 | 0,006189269 | Up |
| Cdkn2c | 6893,065099 | 0,707937422 | 0,090313464 | 1,70E-16 | 3,00E-14 | Up |
| Hectd3 | 4740,445965 | 0,587925611 | 0,067316509 | 1,03E-19 | 3,00E-17 | Up |
| Eif2b3 | 7067,412782 | -0,681733897 | 0,101911828 | 7,90E-13 | 6,68E-11 | Down |
| Plk3 | 911,0619477 | -0,697227493 | 0,222855589 | 2,93E-05 | 0,000279749 | Down |
| Olfr1342 | 762,4197633 | 1,122832885 | 0,217596694 | 6,42E-10 | 2,57E-08 | Up |
| Hivep3 | 636,3806459 | -0,777483862 | 0,177037501 | 1,78E-07 | 3,31E-06 | Down |
| Yrdc | 6107,586385 | -0,661758456 | 0,10130694 | 2,32E-12 | 1,79E-10 | Down |
| Tmem35b | 1067,155059 | 0,648874915 | 0,10403996 | 1,58E-11 | 9,59E-10 | Up |
| Gjb3 | 19,38929696 | -1,105671613 | 0,623099057 | 0,000145177 | 0,001092025 | Down |
| Tlr12 | 147,3951441 | 1,299110691 | 0,266396653 | 1,83E-09 | 6,41E-08 | Up |
| Zfp362 | 5602,70999 | 0,583029844 | 0,123180032 | 7,57E-08 | 1,59E-06 | Up |
| Azin2 | 304,7513812 | 0,733522679 | 0,16745037 | 2,31E-07 | 4,12E-06 | Up |
| Lck | 92,78525319 | 0,753262797 | 0,444960642 | 0,000678632 | 0,004025886 | Up |
| Ccdc28b | 668,8239146 | 0,779098041 | 0,11508987 | 3,24E-13 | 2,99E-11 | Up |
| Spocd1 | 103,4212202 | 2,49221396 | 0,359449273 | 1,07E-14 | 1,43E-12 | Up |
| Matn1 | 423,3702018 | 0,616066132 | 0,261333934 | 0,000400905 | 0,002574464 | Up |
| Sytl1 | 1608,937679 | 0,90378857 | 0,255490633 | 2,52E-06 | 3,25E-05 | Up |
| Selenon | 8822,329824 | -0,649764485 | 0,080634137 | 3,22E-17 | 6,61E-15 | Down |
| Man1c1 | 2246,77068 | -1,12430737 | 0,216010838 | 5,02E-10 | 2,07E-08 | Down |
| Ece1 | 103,7080613 | 0,797235 | 0,33108741 | 0,000123774 | 0,00095267 | Up |
| Mrto4 | 19176,00783 | -0,600694575 | 0,124913458 | 5,00E-08 | 1,11E-06 | Down |
| Padi6 | 31,51235846 | 0,895048454 | 0,454596152 | 0,000207096 | 0,001471164 | Up |
| Fblim1 | 3942,264353 | 0,684742254 | 0,116882007 | 1,46E-10 | 7,03E-09 | Up |
| Tmem51 | 126,582011 | 0,80969168 | 0,650788616 | 0,000897552 | 0,005099011 | Up |
| Tnfrsf8 | 485,8381071 | 0,667792482 | 0,222607524 | 5,10E-05 | 0,000447582 | Up |
| Srm | 41624,26871 | -0,650186891 | 0,085523206 | 1,17E-15 | 1,84E-13 | Down |
| Slc25a33 | 1909,241325 | -0,940075727 | 0,101640759 | 2,00E-22 | 9,82E-20 | Down |
| Gm13091 | 135,4061648 | 0,760549297 | 0,356706187 | 0,000283274 | 0,001927682 | Up |
| Tnfrsf9 | 961,1425314 | -0,792541453 | 0,185671239 | 2,79E-07 | 4,89E-06 | Down |
| Chd5 | 261,9581367 | 1,14828093 | 0,566829671 | 7,70E-05 | 0,000635832 | Up |
| Tas1r3 | 110,3908491 | 0,621894884 | 0,240578087 | 0,000216333 | 0,001531413 | Up |
| 4833413G10Rik | 94,35688259 | 0,61768577 | 0,227573384 | 0,000155409 | 0,001156238 | Up |
| Abcb1b | 4149,851263 | -1,215469592 | 0,114643636 | 4,63E-29 | 1,07E-25 | Down |
| Crot | 3100,332009 | 0,644095516 | 0,12126093 | 3,45E-09 | 1,12E-07 | Up |
| 5031425E22Rik | 1281,159012 | 0,634992003 | 0,095022452 | 8,82E-13 | 7,34E-11 | Up |
| Adgrf3 | 179,9646314 | 0,732170762 | 0,245267135 | 3,76E-05 | 0,00034544 | Up |
| Khk | 614,7936558 | 0,837750692 | 0,13564363 | 1,08E-11 | 6,89E-10 | Up |
| Slc5a6 | 1836,307114 | -0,706775986 | 0,124275445 | 3,70E-10 | 1,59E-08 | Down |
| Mxd4 | 4469,061136 | 1,230751042 | 0,444544208 | 9,32E-06 | 0,000102565 | Up |
| Acox3 | 2910,678591 | 0,73329394 | 0,228748555 | 1,90E-05 | 0,000190217 | Up |
| Sh3tc1 | 912,8728443 | -1,010985449 | 0,160557967 | 1,47E-12 | 1,18E-10 | Down |
| Wdr19 | 376,459108 | 0,670032666 | 0,165772001 | 1,25E-06 | 1,77E-05 | Up |
| Apbb2 | 43,3166444 | 1,257935976 | 0,527751629 | 2,52E-05 | 0,000245654 | Up |
| Cnga1 | 150,1498631 | 1,122984789 | 0,311963686 | 7,72E-07 | 1,17E-05 | Up |

| | | | | | | |
|---|---|---|---|---|---|---|
| 5830473C10Rik | 3008,949964 | -0,83085748 | 0,094368869 | 2,82E-20 | 8,97E-18 | Down |
| Pf4 | 352,6255095 | -0,921798424 | 0,374661992 | 5,92E-05 | 0,000508772 | Down |
| Cxcl3 | 14,84452677 | -0,64856971 | 1,049197789 | 0,001133376 | 0,006204127 | Down |
| Cxcl2 | 57,55468698 | -1,673824647 | 0,921556244 | 3,59E-05 | 0,000332248 | Down |
| Mthfd2l | 1151,184745 | -0,840373256 | 0,099729498 | 6,98E-19 | 1,77E-16 | Down |
| Sdad1 | 12175,14052 | -0,608754026 | 0,102070284 | 8,87E-11 | 4,53E-09 | Down |
| Gm43173 | 16,72317913 | -0,631568078 | 0,42790551 | 0,001859921 | 0,009427435 | Down |
| Ccng2 | 1329,31746 | 0,926753964 | 0,223974775 | 2,26E-07 | 4,04E-06 | Up |
| Gm7993 | 11509,51679 | -0,797634351 | 0,090307836 | 2,75E-20 | 8,97E-18 | Down |
| Bmp2k | 5934,617466 | 0,591065892 | 0,124369892 | 6,75E-08 | 1,43E-06 | Up |
| Spp1 | 14166,01247 | -0,875479941 | 0,210523761 | 2,77E-07 | 4,88E-06 | Down |
| Gm43819 | 34,74630871 | 1,286045968 | 0,576263329 | 3,45E-05 | 0,000321366 | Up |
| Gm10419 | 2401,713468 | 0,671054828 | 0,104965606 | 5,65E-12 | 3,93E-10 | Up |
| Gm18555 | 102,3025521 | 0,605118093 | 0,283383348 | 0,000717156 | 0,004211311 | Up |
| Hscb | 2015,219503 | 0,581550681 | 0,08226819 | 6,12E-14 | 6,81E-12 | Up |
| Tpst2 | 5661,792817 | -0,727564372 | 0,075457342 | 2,11E-23 | 1,37E-20 | Down |
| Ssh1 | 468,8903113 | 1,105967103 | 0,246567776 | 2,01E-08 | 5,08E-07 | Up |
| Rbm19 | 8907,972912 | -0,718497889 | 0,124115304 | 1,98E-10 | 9,25E-09 | Down |
| Sdsl | 2168,067081 | 0,715997178 | 0,103668075 | 1,65E-13 | 1,63E-11 | Up |
| Acad12 | 811,1184419 | 0,625633355 | 0,140739906 | 2,59E-07 | 4,58E-06 | Up |
| Tmem120b | 909,7917846 | -0,642184816 | 0,123664836 | 6,49E-09 | 1,90E-07 | Down |
| Sumf2 | 6590,913562 | -0,609578025 | 0,087906284 | 1,58E-13 | 1,58E-11 | Down |
| Zbed5 | 301,228206 | -0,787855269 | 0,188998076 | 4,33E-07 | 7,11E-06 | Down |
| Clip2 | 3968,004608 | 0,637825056 | 0,079726797 | 5,16E-17 | 1,02E-14 | Up |
| Rfc2 | 25282,33014 | 0,643442746 | 0,067633294 | 8,30E-23 | 4,65E-20 | Up |
| Wbscr27 | 680,0789105 | 0,739820726 | 0,178303076 | 5,96E-07 | 9,39E-06 | Up |
| Ssc4d | 446,6429888 | 0,624235919 | 0,142669447 | 3,56E-07 | 6,02E-06 | Up |
| Myl10 | 156,465505 | 0,900841707 | 0,465509129 | 0,000216564 | 0,001532153 | Up |
| Tfr2 | 6268,32924 | 1,506143607 | 0,206451619 | 6,82E-16 | 1,12E-13 | Up |
| Nyap1 | 219,7785756 | 0,776479838 | 0,212589389 | 3,33E-06 | 4,17E-05 | Up |
| Pilra | 72,16270277 | -0,652325153 | 0,38917686 | 0,001243497 | 0,006705133 | Down |
| Pilrb1 | 241,8391833 | -1,264449446 | 0,31842136 | 1,26E-07 | 2,46E-06 | Down |
| Gm36551 | 78,80995591 | -0,8473178 | 0,331804362 | 6,64E-05 | 0,000560127 | Down |
| Prkar1b | 85,97825731 | 1,182373933 | 0,225958439 | 3,51E-10 | 1,52E-08 | Up |
| Micall2 | 18999,42081 | 0,643316066 | 0,098618663 | 2,52E-12 | 1,93E-10 | Up |
| Lfng | 13135,06574 | -0,772279473 | 0,07914095 | 5,56E-24 | 3,92E-21 | Down |
| Zfp853 | 12,00776943 | -0,658692455 | 0,527922019 | 0,001948814 | 0,009801458 | Down |
| Hsph1 | 59171,53977 | -0,587562551 | 0,134370428 | 4,04E-07 | 6,76E-06 | Down |
| 2610001J05Rik | 9642,694611 | -0,60596135 | 0,110111371 | 1,30E-09 | 4,78E-08 | Down |
| Wnt2 | 10,91640668 | 0,747098851 | 0,651449328 | 0,001266463 | 0,006813116 | Up |
| Asz1 | 351,0637868 | 1,347382055 | 0,379049126 | 5,76E-07 | 9,13E-06 | Up |
| Hilpda | 1219,348805 | -0,608606632 | 0,103245217 | 1,35E-10 | 6,58E-09 | Down |
| Cpa5 | 32,44097674 | 0,943323774 | 0,459262149 | 0,000141714 | 0,001069444 | Up |
| Clec5a | 2770,782305 | -0,929069603 | 0,371919799 | 5,21E-05 | 0,000454499 | Down |
| Gimap8 | 581,8541394 | 0,61261466 | 0,243021101 | 0,000270887 | 0,001856838 | Up |
| Gimap4 | 1437,386556 | 0,608509922 | 0,130460382 | 9,86E-08 | 1,98E-06 | Up |
| Gimap5 | 1384,344771 | 0,622216498 | 0,16310847 | 3,73E-06 | 4,60E-05 | Up |
| Gimap3 | 174,8603122 | 1,147908653 | 0,49311918 | 3,78E-05 | 0,000347478 | Up |
| Tmem176a | 11316,0182 | 0,587724868 | 0,191309792 | 6,19E-05 | 0,000528728 | Up |
| Gm7932 | 19,05910747 | 0,950681588 | 0,574633104 | 0,000301616 | 0,002029745 | Up |
| 9430076C15Rik | 3288,824048 | 0,648340618 | 0,174723449 | 5,00E-06 | 5,91E-05 | Up |

| | | | | | | |
|---|---|---|---|---|---|---|
| Ggct | 684,1492907 | -0,632803175 | 0,1401927 | 1,86E-07 | 3,44E-06 | Down |
| Gm6786 | 14,5831849 | -0,709104024 | 0,66735385 | 0,001547034 | 0,008065752 | Down |
| Igkv14-111 | 84,3055212 | -1,265231237 | 0,342400842 | 3,78E-07 | 6,38E-06 | Down |
| Gm25436 | 45,25769782 | -1,635202176 | 0,49608094 | 1,17E-06 | 1,68E-05 | Down |
| Polr1a | 20692,40902 | -0,591904939 | 0,110716422 | 3,16E-09 | 1,03E-07 | Down |
| Tmem150a | 223,9175703 | -0,668887929 | 0,165069181 | 1,20E-06 | 1,72E-05 | Down |
| Tacr1 | 5,078795076 | -0,637341696 | 1,194019898 | 0,00089971 | 0,005105911 | Down |
| Hk2 | 19649,01962 | -0,892293694 | 0,100709189 | 1,06E-20 | 3,72E-18 | Down |
| Gm17034 | 20,10568958 | 0,858569298 | 0,590272963 | 0,000587528 | 0,003560832 | Up |
| Wbp1 | 1559,880948 | 0,75888939 | 0,140818847 | 1,56E-09 | 5,58E-08 | Up |
| Smyd5 | 18393,46982 | -0,677776825 | 0,104196623 | 2,70E-12 | 2,05E-10 | Down |
| Fam136a | 15246,30518 | -0,592147874 | 0,103982929 | 4,45E-10 | 1,87E-08 | Down |
| Gp9 | 601,7823474 | 0,795744685 | 0,262634279 | 2,32E-05 | 0,000227495 | Up |
| Chchd4 | 12949,78282 | -0,645493324 | 0,099160321 | 2,75E-12 | 2,07E-10 | Down |
| Frmd4b | 176,0736522 | 1,166507898 | 0,349718094 | 1,76E-06 | 2,39E-05 | Up |
| Il17re | 269,1383942 | 1,953394246 | 0,255879777 | 2,20E-16 | 3,85E-14 | Up |
| Irak2 | 1033,048835 | 0,607047468 | 0,166905768 | 7,85E-06 | 8,79E-05 | Up |
| Cand2 | 1623,639553 | 1,747009592 | 0,233159132 | 3,80E-16 | 6,50E-14 | Up |
| Ift122 | 6828,286105 | 0,61364496 | 0,227225775 | 0,000165253 | 0,001214873 | Up |
| Gm8213 | 184,2105679 | 0,772956949 | 0,199936269 | 1,54E-06 | 2,13E-05 | Up |
| Rasgef1a | 67,70702736 | 0,837700442 | 0,330651457 | 7,34E-05 | 0,000611063 | Up |
| Cacna2d4 | 144,6023226 | 0,918070529 | 0,216457391 | 1,54E-07 | 2,93E-06 | Up |
| Wnt5b | 79,42463244 | 0,653744779 | 0,313588642 | 0,000586373 | 0,003556602 | Up |
| Tuba8 | 21444,23503 | -0,844106493 | 0,163290173 | 3,12E-09 | 1,02E-07 | Down |
| Phc1 | 2601,639079 | 0,697655283 | 0,103939305 | 6,54E-13 | 5,65E-11 | Up |
| Slc2a3 | 28352,06815 | -0,64435676 | 0,127168852 | 1,24E-08 | 3,33E-07 | Down |
| C1ra | 714,1074443 | -0,797417586 | 0,271271248 | 3,00E-05 | 0,000285382 | Down |
| Lag3 | 151,5211733 | 1,092937125 | 0,219356418 | 1,86E-09 | 6,49E-08 | Up |
| Ptms | 5382,522805 | 0,97371456 | 0,125621806 | 5,94E-17 | 1,15E-14 | Up |
| Nop2 | 29253,70513 | -0,599276245 | 0,129224728 | 1,15E-07 | 2,28E-06 | Down |
| Fgf23 | 169,6018244 | -4,413776029 | 0,559071299 | 5,81E-17 | 1,14E-14 | Down |
| Tigar | 2775,434518 | -0,808953096 | 0,134698889 | 3,63E-11 | 2,06E-09 | Down |
| Gm26770 | 159,7417486 | 0,91795875 | 0,213594495 | 1,21E-07 | 2,39E-06 | Up |
| Rhno1 | 8719,928351 | 0,620582807 | 0,106830966 | 2,20E-10 | 1,01E-08 | Up |
| Gm43965 | 93,62503784 | 1,201817239 | 0,332779167 | 5,92E-07 | 9,36E-06 | Up |
| Cd69 | 4378,454413 | -1,544174941 | 0,313168574 | 1,56E-09 | 5,58E-08 | Down |
| Bcl2l14 | 2146,088438 | -0,635953043 | 0,145548962 | 3,53E-07 | 5,97E-06 | Down |
| Cdkn1b | 2508,115179 | 0,669849379 | 0,139777822 | 4,51E-08 | 1,02E-06 | Up |
| Emp1 | 7,454085829 | -0,618714026 | 0,990703079 | 0,001338906 | 0,007124876 | Down |
| Fam60a | 3785,475364 | -0,704856695 | 0,080591225 | 8,92E-20 | 2,63E-17 | Down |
| Gm44160 | 2666,084895 | -0,762156703 | 0,234665157 | 1,42E-05 | 0,000148136 | Down |
| Leng8 | 5824,63832 | 0,647292698 | 0,226534498 | 8,73E-05 | 0,000709111 | Up |
| Rdh13 | 4023,991193 | -0,70159073 | 0,133856338 | 4,28E-09 | 1,34E-07 | Down |
| Tmem150b | 60,43056541 | 0,625877089 | 0,31922419 | 0,000898183 | 0,00510081 | Up |
| Sbk2 | 129,9005395 | 0,767703354 | 0,246647658 | 2,11E-05 | 0,000208319 | Up |
| Zfp580 | 1035,542264 | 0,58067847 | 0,123350253 | 8,62E-08 | 1,77E-06 | Up |
| Ehd2 | 210,8835274 | -2,078869747 | 0,312871227 | 2,29E-13 | 2,16E-11 | Down |
| C5ar1 | 80,99254291 | -1,654102845 | 0,414129231 | 1,06E-07 | 2,11E-06 | Down |
| Prkd2 | 8148,275064 | -0,773983802 | 0,090114937 | 2,67E-19 | 7,10E-17 | Down |
| Gng8 | 26,98188007 | -0,850178508 | 0,593489508 | 0,000626091 | 0,003763739 | Down |
| Ptgir | 1602,251641 | -0,985360467 | 0,216599205 | 2,70E-08 | 6,52E-07 | Down |

| | | | | | | |
|---|---|---|---|---|---|---|
| Psg17 | 14,23180398 | 1,073678784 | 1,265048722 | 0,000335838 | 0,002220396 | Up |
| Mill2 | 26,68541156 | 3,003225746 | 0,763598742 | 3,69E-08 | 8,62E-07 | Up |
| Gm45138 | 8,552773256 | 0,868597411 | 0,904290714 | 0,000701997 | 0,004141782 | Up |
| Dmwd | 5860,363685 | -0,747436532 | 0,08271588 | 5,77E-21 | 2,13E-18 | Down |
| Pvr | 729,3870502 | -0,783198157 | 0,176761221 | 1,46E-07 | 2,78E-06 | Down |
| Plaur | 2252,858594 | -1,407565099 | 0,15994344 | 2,47E-21 | 1,03E-18 | Down |
| Rab4b | 333,2527098 | 0,647451383 | 0,131426817 | 2,50E-08 | 6,11E-07 | Up |
| Numbl | 178,9177678 | 0,983172604 | 0,358574513 | 2,15E-05 | 0,000212015 | Up |
| Sertad3 | 879,2062655 | 0,644376331 | 0,11152334 | 2,55E-10 | 1,15E-08 | Up |
| Prx | 249,3020605 | 0,610073574 | 0,190735896 | 3,66E-05 | 0,000337717 | Up |
| 9530053A07Rik | 88,21856048 | -1,587065862 | 0,480447441 | 1,17E-06 | 1,69E-05 | Down |
| Gm45010 | 81,40064451 | 0,82938942 | 0,216158895 | 1,27E-06 | 1,80E-05 | Up |
| Zfp36 | 1356,414366 | -1,488689587 | 0,192147785 | 2,09E-17 | 4,48E-15 | Down |
| Zfp14 | 96,41452789 | 0,660655676 | 0,192712579 | 1,31E-05 | 0,000138446 | Up |
| Proser3 | 945,1244147 | 0,643891211 | 0,163855441 | 2,18E-06 | 2,87E-05 | Up |
| Ffar2 | 2584,762228 | 1,166153595 | 0,54922598 | 5,90E-05 | 0,000507031 | Up |
| Tdrd12 | 39,2821607 | -0,703601044 | 0,326604676 | 0,000373868 | 0,002426403 | Down |
| Siglecf | 2157,366372 | 2,220151454 | 0,318822425 | 2,29E-14 | 2,84E-12 | Up |
| Siglecg | 42,41160238 | 1,147025586 | 0,304503168 | 3,77E-07 | 6,36E-06 | Up |
| Nkg7 | 1081,063626 | 0,690382281 | 0,160563226 | 3,93E-07 | 6,61E-06 | Up |
| Ptov1 | 8330,207477 | 0,649084036 | 0,090642219 | 3,11E-14 | 3,73E-12 | Up |
| Ntf5 | 118,2839326 | 0,667563625 | 0,222725083 | 5,15E-05 | 0,000450648 | Up |
| Hsd17b14 | 265,8875227 | 1,027474668 | 0,228058228 | 2,63E-08 | 6,40E-07 | Up |
| A030001D20Rik | 31,10105575 | 1,064971785 | 0,432082339 | 3,39E-05 | 0,00031722 | Up |
| Grwd1 | 15240,31023 | -0,611010783 | 0,134933156 | 1,85E-07 | 3,43E-06 | Down |
| Syngr4 | 51,83139779 | 0,735499355 | 0,266875148 | 7,10E-05 | 0,000593421 | Up |
| Nomo1 | 42213,93297 | -0,740840559 | 0,110476928 | 5,94E-13 | 5,15E-11 | Down |
| Saa3 | 230,6534583 | 0,81421586 | 0,184029725 | 1,27E-07 | 2,48E-06 | Up |
| Saa4 | 11,52120595 | -1,429746909 | 1,718745781 | 0,000150147 | 0,001121607 | Down |
| Gtf2h1 | 19111,93618 | -0,604315788 | 0,118844102 | 1,23E-08 | 3,32E-07 | Down |
| Ldhc | 153,8158164 | 0,847815539 | 0,233025492 | 2,38E-06 | 3,10E-05 | Up |
| Gm2788 | 162,2014534 | 0,676877556 | 0,273295652 | 0,000206896 | 0,001470387 | Up |
| Tjp1 | 79,21729523 | 0,68071988 | 0,482025317 | 0,001486961 | 0,007800175 | Up |
| Aen | 10887,20713 | -0,722728799 | 0,12198416 | 8,83E-11 | 4,52E-09 | Down |
| Zscan2 | 576,059414 | 0,600332317 | 0,115832253 | 7,46E-09 | 2,15E-07 | Up |
| Alpk3 | 1330,12285 | 0,639627666 | 0,155317691 | 1,03E-06 | 1,49E-05 | Up |
| Abhd17c | 18929,1166 | -1,218748924 | 0,126187029 | 7,30E-25 | 5,92E-22 | Down |
| Arnt2 | 849,7298932 | -2,655122696 | 0,383109474 | 5,16E-15 | 7,35E-13 | Down |
| Gm44556 | 10,50485091 | -1,331876686 | 1,180543732 | 0,000167812 | 0,001227013 | Down |
| Rab38 | 29661,05411 | 0,853158557 | 0,091531041 | 2,13E-22 | 1,02E-19 | Up |
| Usp35 | 183,9917536 | 0,590734775 | 0,240395278 | 0,000364506 | 0,002380404 | Up |
| Aqp11 | 254,6430333 | 0,866945983 | 0,145008512 | 2,96E-11 | 1,71E-09 | Up |
| Stard10 | 87,27593361 | 0,75784643 | 0,302375477 | 0,000120505 | 0,000930604 | Up |
| Usp17ld | 816,2852816 | -1,988712377 | 0,199419847 | 3,04E-25 | 2,90E-22 | Down |
| Usp17lc | 25,65547678 | -1,12232981 | 0,962992826 | 0,000287655 | 0,001953623 | Down |
| Trim34b | 146,991542 | 0,75934127 | 0,180707689 | 4,36E-07 | 7,15E-06 | Up |
| Gm4070 | 372,7722627 | -1,117437295 | 0,252180784 | 2,47E-08 | 6,05E-07 | Down |
| 4930431P19Rik | 74,04228373 | 0,694854246 | 0,269437012 | 0,000143522 | 0,001081083 | Up |
| RP23-462D5,5 | 106,5448752 | 0,635050759 | 0,23868878 | 0,000163299 | 0,00120433 | Up |
| Lyve1 | 54,82085729 | 0,77665662 | 0,407676558 | 0,000411797 | 0,002632548 | Up |
| Gm45355 | 192,1436969 | -0,658521821 | 0,173064592 | 3,34E-06 | 4,17E-05 | Down |

| | | | | | | |
|---|---|---|---|---|---|---|
| Il4ra | 15006,45332 | -0,735379158 | 0,111028655 | 1,05E-12 | 8,66E-11 | Down |
| Mapk3 | 9715,580758 | 0,596369654 | 0,099582948 | 7,74E-11 | 4,04E-09 | Up |
| Ypel3 | 612,7755544 | 0,663936922 | 0,378130873 | 0,001019134 | 0,005668649 | Up |
| Fam57b | 565,1497405 | 0,582718003 | 0,133451975 | 4,21E-07 | 6,95E-06 | Up |
| Qprt | 327,3661563 | 0,796560972 | 0,156228992 | 5,87E-09 | 1,75E-07 | Up |
| Mylpf | 586,579511 | 0,726979831 | 0,133269729 | 1,25E-09 | 4,66E-08 | Up |
| Gm44728 | 15,19890685 | 0,684393155 | 0,507183571 | 0,001551992 | 0,008086405 | Up |
| Itgam | 2870,78632 | -0,801642154 | 0,225192501 | 4,04E-06 | 4,89E-05 | Down |
| Adam8 | 724,7217499 | -2,259569943 | 0,424769557 | 3,29E-10 | 1,43E-08 | Down |
| Gm7514 | 58,25403277 | -1,728838241 | 0,408136773 | 4,30E-08 | 9,77E-07 | Down |
| Ifitm6 | 396,4509885 | -1,057169528 | 0,846628019 | 0,000339773 | 0,002241895 | Down |
| Lrrc56 | 288,3536262 | 1,057071869 | 0,169976342 | 1,80E-12 | 1,41E-10 | Up |
| Lmntd2 | 26,60692556 | 0,799669114 | 0,568752522 | 0,000810215 | 0,004661608 | Up |
| Slc25a22 | 4130,3119 | -0,615597181 | 0,143752679 | 5,52E-07 | 8,83E-06 | Down |
| Cracr2b | 146,9201885 | 1,055880935 | 0,245405379 | 5,71E-08 | 1,24E-06 | Up |
| Tspan4 | 10535,26859 | -0,804267556 | 0,098262277 | 6,75E-18 | 1,52E-15 | Down |
| Muc6 | 10,9192414 | 0,593562187 | 0,56700443 | 0,003039536 | 0,014082374 | Up |
| Muc5b | 13,40767895 | -1,288641089 | 0,767535545 | 0,000103921 | 0,000823301 | Down |
| E230032D23Rik | 70,81691303 | 0,694147991 | 0,338709101 | 0,000492582 | 0,003065649 | Up |
| Osbpl5 | 159,6143773 | 1,473887287 | 0,447823527 | 1,30E-06 | 1,84E-05 | Up |
| Dhcr7 | 10305,01746 | 0,620006284 | 0,170247852 | 7,30E-06 | 8,24E-05 | Up |
| Col4a2 | 1922,424638 | -0,990464883 | 0,176962122 | 1,18E-10 | 5,91E-09 | Down |
| Adprhl1 | 11,71580451 | 1,470887475 | 1,052112586 | 9,70E-05 | 0,000776638 | Up |
| RP23-106I21,5 | 45,37578159 | 0,59262389 | 0,317351384 | 0,001332839 | 0,007099577 | Up |
| Adgra2 | 117,7847931 | 0,831088487 | 0,229510567 | 2,78E-06 | 3,54E-05 | Up |
| Rbpms | 4998,03979 | 0,685286301 | 0,167583573 | 9,69E-07 | 1,43E-05 | Up |
| Mtmr7 | 205,1509491 | 0,797675353 | 0,222892124 | 3,87E-06 | 4,72E-05 | Up |
| Tma16 | 3261,515626 | -0,714410681 | 0,131291374 | 1,41E-09 | 5,12E-08 | Down |
| Slc18a1 | 40,47516133 | 0,861822007 | 0,36425693 | 9,87E-05 | 0,000788571 | Up |
| Abhd8 | 2097,592191 | 0,616283464 | 0,117113593 | 4,75E-09 | 1,46E-07 | Up |
| Slc27a1 | 1131,808883 | 0,62521067 | 0,140971733 | 2,72E-07 | 4,80E-06 | Up |
| Calr3 | 350,2557581 | -0,595404946 | 0,154147998 | 3,43E-06 | 4,26E-05 | Down |
| Gm17435 | 245,4372985 | -0,607634752 | 0,142536036 | 6,16E-07 | 9,67E-06 | Down |
| Nwd1 | 3134,380359 | 1,61225427 | 0,313500274 | 6,07E-10 | 2,44E-08 | Up |
| Hmox1 | 11159,22222 | -0,678921158 | 0,20760609 | 2,05E-05 | 0,000202997 | Down |
| Smad1 | 116,2128646 | 1,518015903 | 0,310384195 | 1,84E-09 | 6,44E-08 | Up |
| Ier2 | 10364,82877 | -0,596294097 | 0,106524618 | 7,72E-10 | 3,03E-08 | Down |
| G430095P16Rik | 558,6620452 | 0,623121305 | 0,107220761 | 2,16E-10 | 9,97E-09 | Up |
| Klf1 | 812,5738791 | 0,884724156 | 0,124733239 | 1,69E-14 | 2,18E-12 | Up |
| Junb | 6396,057393 | -0,818353264 | 0,102776423 | 3,75E-17 | 7,60E-15 | Down |
| Nod2 | 319,3092233 | 1,153359127 | 0,151949148 | 6,70E-17 | 1,28E-14 | Up |
| Drc7 | 87,03787323 | 1,500073556 | 0,425677999 | 5,83E-07 | 9,24E-06 | Up |
| Lrrc29 | 453,0647245 | 0,89530657 | 0,096222963 | 1,75E-22 | 9,01E-20 | Up |
| Slc9a5 | 593,7840745 | 0,828850212 | 0,149981045 | 5,10E-10 | 2,09E-08 | Up |
| Lrrc36 | 24,50060593 | 1,058911671 | 0,43946008 | 4,00E-05 | 0,000363438 | Up |
| Fam65a | 21843,04541 | 0,782891307 | 0,153258063 | 6,00E-09 | 1,78E-07 | Up |
| Cirh1a | 18575,52838 | -0,629537292 | 0,090229495 | 1,16E-13 | 1,22E-11 | Down |
| Rpl10-ps2 | 16,17777613 | -1,68115022 | 0,633741779 | 6,55E-06 | 7,52E-05 | Down |
| Nip7 | 12254,68966 | -0,623552575 | 0,1082922 | 2,96E-10 | 1,29E-08 | Down |
| Tmem231 | 119,1414602 | 0,655256774 | 0,244702244 | 0,000138686 | 0,001049813 | Up |
| 6430548M08Rik | 31,95152801 | 0,581183991 | 0,459524844 | 0,003135482 | 0,014403509 | Up |

| | | | | | | |
|---|---|---|---|---|---|---|
| 1700016A09Rik | 37,75277305 | 1,075147775 | 0,384078256 | 1,29E-05 | 0,000136954 | Up |
| Gm27011 | 34,78370778 | 1,086321261 | 0,411630565 | 1,98E-05 | 0,0001972 | Up |
| Gm17709 | 80,27379135 | -0,924469469 | 0,273393377 | 3,88E-06 | 4,72E-05 | Down |
| Gm45353 | 43,3470747 | 1,852264584 | 0,379433026 | 3,07E-09 | 1,01E-07 | Up |
| Rab4a | 2004,414781 | 0,624336784 | 0,080491176 | 3,57E-16 | 6,16E-14 | Up |
| Fam89a | 27,71188534 | -0,898406274 | 0,410870248 | 0,00012705 | 0,000974652 | Down |
| Maml2 | 1577,826102 | 0,737180502 | 0,153364916 | 3,28E-08 | 7,80E-07 | Up |
| 9530077C05Rik | 70,95159107 | 0,671421694 | 0,295661263 | 0,000350079 | 0,002302407 | Up |
| Robo3 | 29,5167106 | -0,964857906 | 0,57355269 | 0,000272898 | 0,001868261 | Down |
| Esam | 16,95828264 | 0,702921835 | 0,575034715 | 0,001544263 | 0,008053894 | Up |
| Vsig2 | 48,79650489 | 1,191270218 | 0,383983104 | 3,63E-06 | 4,48E-05 | Up |
| Nrgn | 5176,970255 | 0,796554427 | 0,171131857 | 5,00E-08 | 1,11E-06 | Up |
| RP23-181A8,7 | 61,65373114 | 1,250672589 | 0,478299338 | 1,39E-05 | 0,000145143 | Up |
| H2afx | 11103,76724 | 0,588086128 | 0,204310587 | 0,000113338 | 0,000884094 | Up |
| RP24-175C20,15 | 71,69740683 | -0,708602532 | 0,374428984 | 0,000613698 | 0,003697455 | Down |
| RP24-282C4,11 | 92,43994388 | 0,743708104 | 0,545820721 | 0,001115189 | 0,006127307 | Up |
| Acsbg1 | 13,55163834 | -1,058930821 | 0,702215258 | 0,000259709 | 0,001792337 | Down |
| Gm10658 | 113,396517 | 0,945715251 | 0,190743385 | 4,79E-09 | 1,47E-07 | Up |
| Neil1 | 1084,607884 | 0,594443056 | 0,116847133 | 1,24E-08 | 3,34E-07 | Up |
| Islr | 32989,16992 | 0,791824558 | 0,151580745 | 3,17E-09 | 1,03E-07 | Up |
| Cd276 | 489,4175603 | -1,377629532 | 0,274213049 | 8,32E-10 | 3,23E-08 | Down |
| Tle3 | 102,443084 | -0,974836817 | 0,417596851 | 6,52E-05 | 0,000551927 | Down |
| Slc24a1 | 31,36208682 | 0,884495938 | 0,459272889 | 0,0002366 | 0,00165681 | Up |
| Parp16 | 265,9383329 | 0,661897297 | 0,144791987 | 1,30E-07 | 2,53E-06 | Up |
| Car12 | 3838,001602 | -1,048801076 | 0,115734975 | 4,66E-22 | 2,16E-19 | Down |
| Aph1b | 597,931682 | 0,775901077 | 0,288179282 | 6,73E-05 | 0,000566479 | Up |
| Ccnb2 | 22378,82777 | 0,661523405 | 0,09291729 | 4,13E-14 | 4,89E-12 | Up |
| Fam63b | 5455,108403 | 0,603260766 | 0,188070003 | 3,67E-05 | 0,000338596 | Up |
| Gm10642 | 31,73537749 | 1,158534691 | 0,62381924 | 0,000106471 | 0,00084145 | Up |
| Ccpg1os | 184,6730959 | 0,652074171 | 0,139897806 | 8,87E-08 | 1,81E-06 | Up |
| Gsta4 | 475,055393 | 0,90572677 | 0,150579274 | 1,81E-11 | 1,08E-09 | Up |
| Filip1 | 24,89787585 | 0,619475575 | 0,405070586 | 0,001883052 | 0,009523853 | Up |
| Bcl2a1d | 151,2978307 | -0,717440091 | 0,30425496 | 0,000217597 | 0,001537679 | Down |
| Bcl2a1a | 3219,401871 | -0,701887489 | 0,33008146 | 0,000400856 | 0,002574464 | Down |
| Bcl2a1b | 1443,482025 | -0,638798167 | 0,24743407 | 0,000197633 | 0,001413219 | Down |
| Pcolce2 | 191,0624723 | 1,165452713 | 0,260165965 | 1,67E-08 | 4,34E-07 | Up |
| Trpc1 | 11,72276136 | 0,795140412 | 0,800290586 | 0,000983118 | 0,005507974 | Up |
| Pls1 | 3170,196103 | 1,899563813 | 0,201564551 | 5,32E-23 | 3,20E-20 | Up |
| 1700065D16Rik | 152,2227638 | 1,066939157 | 0,236275874 | 2,07E-08 | 5,20E-07 | Up |
| Ky | 967,2072409 | -1,831432155 | 0,147434005 | 2,63E-37 | 1,07E-33 | Down |
| Bfsp2 | 56,3096011 | -0,718960238 | 0,328771319 | 0,000318422 | 0,002121722 | Down |
| Alas1 | 5364,66239 | -0,956784686 | 0,121115827 | 2,10E-17 | 4,48E-15 | Down |
| Dusp7 | 2854,224814 | -0,846068447 | 0,085278458 | 6,70E-25 | 5,72E-22 | Down |
| Rrp9 | 13275,934 | -0,927497496 | 0,098753604 | 5,80E-23 | 3,36E-20 | Down |
| Mapkapk3 | 18249,63982 | -1,089291789 | 0,179097041 | 3,56E-12 | 2,58E-10 | Down |
| Cish | 14896,70043 | -2,33745174 | 0,150899592 | 3,93E-56 | 6,37E-52 | Down |
| Hemk1 | 5617,126081 | -1,049290821 | 0,155466417 | 5,58E-14 | 6,25E-12 | Down |
| 6430571L13Rik | 340,745674 | -2,021531264 | 0,191812199 | 9,34E-28 | 1,38E-24 | Down |
| Hyal2 | 3828,837013 | -0,600438553 | 0,094022312 | 6,38E-12 | 4,33E-10 | Down |
| Hyal1 | 77,53824491 | 0,718448913 | 0,340873899 | 0,000378445 | 0,002451405 | Up |
| Uba7 | 6430,153944 | 0,743995095 | 0,19307936 | 1,88E-06 | 2,52E-05 | Up |

| | | | | | | |
|---|---|---|---|---|---|---|
| Nicn1 | 1482,958208 | 0,680302019 | 0,107831619 | 9,48E-12 | 6,18E-10 | Up |
| Gm37475 | 14,36108092 | 0,622484329 | 0,5054217 | 0,002446744 | 0,01185293 | Up |
| 4833445I07Rik | 115,5348381 | 0,706719871 | 0,241810529 | 5,20E-05 | 0,000453679 | Up |
| 3000002C10Rik | 450,4893725 | -0,710186637 | 0,203439258 | 8,29E-06 | 9,23E-05 | Down |
| Nme6 | 4117,500967 | -0,645253174 | 0,087952247 | 8,66E-15 | 1,18E-12 | Down |
| Gm42432 | 31,82440315 | -0,888068759 | 0,532015615 | 0,000376086 | 0,002439823 | Down |
| Gm42762 | 74,11714831 | 1,147375187 | 0,380878584 | 5,43E-06 | 6,35E-05 | Up |
| Gm43814 | 11,2666907 | 0,600711211 | 0,534513676 | 0,00291564 | 0,013639156 | Up |
| Ccrl2 | 579,18791 | 0,719512271 | 0,249583794 | 5,43E-05 | 0,000471778 | Up |
| Ccr4 | 3470,273775 | -0,641954491 | 0,168805404 | 3,61E-06 | 4,46E-05 | Down |
| Csrnp1 | 4757,737723 | -0,605800129 | 0,124730678 | 3,90E-08 | 9,02E-07 | Down |
| Pomgnt2 | 40,24995299 | 1,675187877 | 0,425208001 | 1,31E-07 | 2,53E-06 | Up |
| Ccr1 | 520,2067066 | -1,137491075 | 0,734090063 | 0,000191702 | 0,001375053 | Down |
| Grm1 | 15,35233286 | 1,334959758 | 0,89400083 | 0,000119803 | 0,000926503 | Up |
| Gm8355 | 611,5442359 | -0,688869667 | 0,098349825 | 8,94E-14 | 9,61E-12 | Down |
| Ccdc28a | 620,3424095 | 0,777517313 | 0,12577118 | 1,47E-11 | 9,02E-10 | Up |
| Gm10825 | 50,07323976 | -5,848273284 | 1,256148891 | 1,90E-08 | 4,82E-07 | Down |
| Arhgap18 | 4949,269253 | 0,748992923 | 0,157222314 | 3,81E-08 | 8,85E-07 | Up |
| RP23-331E5,10 | 95,31111914 | 0,913111561 | 0,29148119 | 9,20E-06 | 0,00010159 | Up |
| Cdk19 | 4869,431134 | 0,753250573 | 0,111741413 | 4,45E-13 | 4,01E-11 | Up |
| Smpd2 | 190,4245367 | 0,938623775 | 0,187855425 | 4,13E-09 | 1,30E-07 | Up |
| Scml4 | 61,52170318 | 1,897982591 | 0,416086107 | 1,26E-08 | 3,38E-07 | Up |
| Lilrb4a | 4829,529236 | -4,888676661 | 0,572279022 | 9,49E-19 | 2,30E-16 | Down |
| Cdh23 | 245,9054833 | 1,148107459 | 0,353582813 | 2,51E-06 | 3,24E-05 | Up |
| Ppa1 | 20164,62417 | -0,62645944 | 0,122874605 | 1,10E-08 | 3,01E-07 | Down |
| Stox1 | 662,4546005 | 0,665443621 | 0,123980604 | 2,44E-09 | 8,27E-08 | Up |
| Tet1 | 377,1038207 | 0,811583595 | 0,316670318 | 7,79E-05 | 0,00064229 | Up |
| Egr2 | 28,52885209 | -0,685131005 | 0,368678993 | 0,000750372 | 0,004362849 | Down |
| Rhobtb1 | 961,7149619 | -0,67097854 | 0,092835188 | 1,87E-14 | 2,35E-12 | Down |
| Adora2a | 4036,259336 | 0,871839718 | 0,132802893 | 7,07E-13 | 6,01E-11 | Up |
| Ggt5 | 50,42020708 | 0,899116713 | 0,338505172 | 3,93E-05 | 0,000359037 | Up |
| Gstt3 | 2123,004728 | 0,737776446 | 0,152636978 | 2,88E-08 | 6,91E-07 | Up |
| Slc19a1 | 5856,627741 | -0,658001562 | 0,113782568 | 2,41E-10 | 1,10E-08 | Down |
| Dnmt3l | 1127,374534 | -0,75409481 | 0,211746653 | 5,18E-06 | 6,10E-05 | Down |
| Pwp2 | 12485,74247 | -0,711260595 | 0,112288179 | 7,45E-12 | 4,96E-10 | Down |
| Gamt | 53,96634743 | 0,852540162 | 0,296685953 | 2,71E-05 | 0,000261735 | Up |
| Apc2 | 208,6627662 | 0,735095343 | 0,245603441 | 3,61E-05 | 0,000333836 | Up |
| Reep6 | 194,4615902 | -1,069495528 | 0,153182135 | 9,61E-15 | 1,30E-12 | Down |
| Btbd2 | 3933,672217 | 0,580110069 | 0,138484066 | 9,31E-07 | 1,38E-05 | Up |
| Zfr2 | 68,36845577 | 0,674944029 | 0,373012247 | 0,000870044 | 0,004968833 | Up |
| Tjp3 | 623,5626684 | 1,396332815 | 0,277544632 | 8,07E-10 | 3,15E-08 | Up |
| Smim24 | 924,2397926 | 1,092322791 | 0,210656693 | 6,43E-10 | 2,57E-08 | Up |
| Celf5 | 35,77832373 | 0,703970137 | 0,394638989 | 0,000765406 | 0,004438423 | Up |
| Tle2 | 36,50292604 | 0,878824387 | 0,420492178 | 0,000171902 | 0,001251286 | Up |
| Eid3 | 30,58930519 | -0,748692837 | 0,392053124 | 0,000473988 | 0,002968322 | Down |
| Slc41a2 | 107,4646155 | 0,990980716 | 0,265297017 | 7,91E-07 | 1,20E-05 | Up |
| RP23-132J19,1 | 72,17940298 | -2,236215589 | 0,33394323 | 1,31E-13 | 1,35E-11 | Down |
| A230060F14Rik | 81,9251504 | 1,819420988 | 0,329626673 | 1,29E-10 | 6,34E-09 | Up |
| Fbxo7 | 10180,18259 | 1,083522629 | 0,187372816 | 2,29E-11 | 1,35E-09 | Up |
| Syn3 | 40,9513894 | 1,101260944 | 0,510188392 | 6,51E-05 | 0,000551336 | Up |
| Gm26546 | 34,28311744 | 0,628577173 | 0,363632963 | 0,001331946 | 0,007097151 | Up |

| | | | | | | |
|---|---|---|---|---|---|---|
| Utp20 | 17353,32244 | -0,753614822 | 0,169453245 | 1,58E-07 | 2,98E-06 | Down |
| Gas2l3 | 16653,24652 | 0,648795578 | 0,189981782 | 1,45E-05 | 0,000151156 | Up |
| Nr1h4 | 81,58779119 | 1,964534318 | 0,682206096 | 2,62E-06 | 3,36E-05 | Up |
| 1500026H17Rik | 14,69804799 | 0,610298397 | 0,578056864 | 0,002737061 | 0,012975911 | Up |
| Elk3 | 7839,28634 | 0,753800985 | 0,125125736 | 4,31E-11 | 2,39E-09 | Up |
| RP23-381H23,1 | 442,2506705 | 1,597862844 | 0,1817236 | 5,42E-21 | 2,08E-18 | Up |
| Gm3571 | 49,55293037 | 0,617686047 | 0,309758982 | 0,000887038 | 0,005051664 | Up |
| Cep83os | 549,2854676 | 0,993511996 | 0,217433331 | 2,35E-08 | 5,79E-07 | Up |
| Plxnc1 | 718,5796189 | -0,610919308 | 0,187964284 | 3,06E-05 | 0,000289899 | Down |
| Dusp6 | 5131,240224 | -0,625665057 | 0,149098951 | 7,75E-07 | 1,18E-05 | Down |
| Tmtc2 | 3411,412567 | -0,842357945 | 0,110689425 | 5,03E-16 | 8,32E-14 | Down |
| Gm15663 | 429,8583652 | -0,731048571 | 0,141758578 | 5,95E-09 | 1,77E-07 | Down |
| Phlda1 | 371,3231829 | -0,714658391 | 0,169509985 | 5,07E-07 | 8,18E-06 | Down |
| Kif5a | 352,4981487 | 1,313807192 | 0,200158303 | 8,41E-14 | 9,16E-12 | Up |
| Gli1 | 31,8384981 | 0,782189901 | 0,401730783 | 0,000368864 | 0,002402105 | Up |
| Lrp1 | 58,39551314 | -1,245147127 | 0,367044834 | 1,20E-06 | 1,72E-05 | Down |
| Apof | 33,50506624 | 0,770434317 | 0,527701456 | 0,000873221 | 0,004983471 | Up |
| Stat2 | 5653,784203 | 0,834936639 | 0,134387065 | 8,72E-12 | 5,70E-10 | Up |
| Suox | 2564,30032 | 0,654508904 | 0,086603937 | 1,64E-15 | 2,56E-13 | Up |
| Dgka | 2076,944143 | 0,804298879 | 0,170703565 | 3,69E-08 | 8,62E-07 | Up |
| Ccdc157 | 239,7664646 | 0,873952593 | 0,191263499 | 4,74E-08 | 1,06E-06 | Up |
| Gatsl3 | 234,9136018 | -1,138196224 | 0,242945085 | 6,87E-09 | 2,00E-07 | Down |
| Osm | 2398,355282 | -0,979797539 | 0,137902125 | 7,41E-15 | 1,03E-12 | Down |
| Znrf3 | 2485,085134 | -0,591164738 | 0,091250952 | 3,52E-12 | 2,56E-10 | Down |
| Polm | 1099,511717 | 1,0574652 | 0,161850381 | 2,30E-13 | 2,16E-11 | Up |
| H2afv | 5771,250671 | 0,779901827 | 0,142824653 | 9,60E-10 | 3,67E-08 | Up |
| Nacad | 199,1895544 | 0,584770602 | 0,212984692 | 0,000171224 | 0,001247469 | Up |
| Upp1 | 1663,281966 | 1,037338332 | 0,198336221 | 6,73E-10 | 2,67E-08 | Up |
| Pno1 | 8850,649921 | -0,727787996 | 0,100373416 | 1,38E-14 | 1,81E-12 | Down |
| Wdpcp | 417,7205719 | 0,799583212 | 0,15757645 | 6,53E-09 | 1,91E-07 | Up |
| Havcr2 | 694,1999369 | 0,78046932 | 0,159484708 | 1,76E-08 | 4,53E-07 | Up |
| Olfr56 | 109,8057391 | -1,058405912 | 0,24747463 | 6,33E-08 | 1,35E-06 | Down |
| BC049762 | 33,06231978 | 0,659188078 | 0,382175419 | 0,001103922 | 0,006069514 | Up |
| D930048N14Rik | 299,0755494 | 0,679916047 | 0,192495217 | 8,32E-06 | 9,25E-05 | Up |
| Acsl6 | 220,3004438 | 1,275182611 | 0,231005374 | 5,77E-11 | 3,08E-09 | Up |
| Fat2 | 185,720143 | -0,957533708 | 0,375788917 | 4,05E-05 | 0,000367361 | Down |
| Gm12247 | 49,05827899 | 1,005681517 | 0,368307155 | 2,03E-05 | 0,000201306 | Up |
| Gm12258 | 129,3155738 | 0,732797875 | 0,228866825 | 1,92E-05 | 0,000192776 | Up |
| Natd1 | 989,8476047 | 1,064401556 | 0,159697613 | 9,06E-14 | 9,67E-12 | Up |
| Map2k3os | 374,2703328 | -0,62321396 | 0,182707716 | 1,66E-05 | 0,000169527 | Down |
| Lrrc75a | 1127,904021 | 0,882064988 | 0,173021623 | 3,54E-09 | 1,14E-07 | Up |
| Hs3st3b1 | 2950,344531 | -0,83578842 | 0,097322462 | 1,81E-19 | 5,07E-17 | Down |
| Slc25a35 | 67,47216633 | 1,367397759 | 0,568582878 | 1,93E-05 | 0,000193071 | Up |
| Alox8 | 110,5484431 | -0,682309095 | 0,253734261 | 0,000115153 | 0,000896956 | Down |
| Kdm6b | 6599,126144 | -1,193539742 | 0,119628251 | 3,26E-26 | 3,53E-23 | Down |
| Cd68 | 1730,234601 | -0,760600026 | 0,24214654 | 2,00E-05 | 0,000199504 | Down |
| Tnfsf13os | 9,098577115 | 0,598265892 | 0,591141808 | 0,002911523 | 0,013633323 | Up |
| Tnfsf13 | 62,71437089 | 0,806919956 | 0,305005551 | 6,44E-05 | 0,000547003 | Up |
| Zbtb4 | 21,65918398 | 1,796955736 | 0,587137911 | 2,04E-06 | 2,70E-05 | Up |
| Nlgn2 | 91,9016118 | 0,588050712 | 0,233557332 | 0,000316281 | 0,002111793 | Up |
| 2810408A11Rik | 34,14067 | 0,717125436 | 0,341698982 | 0,0003886 | 0,002506975 | Up |

| | | | | | | |
|---|---|---|---|---|---|---|
| Arrb2 | 15836,62708 | -0,709483976 | 0,089335161 | 7,48E-17 | 1,40E-14 | Down |
| Gp1ba | 496,4015227 | 1,400253098 | 0,198398272 | 2,92E-15 | 4,27E-13 | Up |
| Rnf167 | 5108,691452 | 0,655260526 | 0,090993515 | 2,31E-14 | 2,84E-12 | Up |
| Aipl1 | 422,8715118 | 1,06527889 | 0,214026161 | 2,18E-09 | 7,49E-08 | Up |
| Fam64a | 7453,227251 | 0,760539271 | 0,167144813 | 9,65E-08 | 1,95E-06 | Up |
| Spns3 | 7776,20801 | 0,768227836 | 0,319884575 | 0,000147102 | 0,001103432 | Up |
| Itgae | 446,2489257 | 0,631601793 | 0,152850866 | 9,96E-07 | 1,46E-05 | Up |
| Serpinf1 | 509,0275843 | 1,140015409 | 0,310576632 | 5,59E-07 | 8,91E-06 | Up |
| Gm45606 | 553,0133456 | 0,734087948 | 0,162659044 | 1,29E-07 | 2,51E-06 | Up |
| Inpp5k | 2879,57959 | 0,631108095 | 0,106919601 | 1,25E-10 | 6,16E-09 | Up |
| Lgals9 | 16272,47577 | 0,893037214 | 0,185972964 | 1,43E-08 | 3,78E-07 | Up |
| Gm11417 | 102,8043685 | 1,199897094 | 0,213695078 | 3,86E-11 | 2,16E-09 | Up |
| Slfn10-ps | 1497,875807 | -0,616284383 | 0,094927705 | 3,17E-12 | 2,34E-10 | Down |
| Slfn2 | 1045,664031 | -2,646125118 | 0,348086933 | 3,80E-17 | 7,60E-15 | Down |
| Slfn1 | 8,722753994 | -0,915712092 | 1,360572994 | 0,000449501 | 0,00283891 | Down |
| Ccl3 | 441,8749966 | -1,901865454 | 0,500469074 | 2,11E-07 | 3,82E-06 | Down |
| Ccl4 | 335,2517241 | -2,532303471 | 0,370563528 | 1,77E-14 | 2,26E-12 | Down |
| Myo19 | 4985,160782 | -0,759366354 | 0,159333047 | 3,60E-08 | 8,44E-07 | Down |
| Utp18 | 8967,047736 | -0,58757951 | 0,086325069 | 3,87E-13 | 3,51E-11 | Down |
| Acsf2 | 171,4211119 | 0,727024237 | 0,215369169 | 1,13E-05 | 0,000120998 | Up |
| Tbkbp1 | 9540,603635 | -0,700387928 | 0,101494186 | 1,79E-13 | 1,74E-11 | Down |
| Plxdc1 | 19,07902451 | 1,188674027 | 0,573735089 | 6,26E-05 | 0,000533865 | Up |
| Cacnb1 | 130,9913445 | 0,599771101 | 0,256790965 | 0,000467613 | 0,002933881 | Up |
| Fbxl20 | 4941,19867 | 0,773204221 | 0,166325362 | 5,80E-08 | 1,25E-06 | Up |
| Thra | 1458,433579 | 0,635081408 | 0,144405706 | 3,12E-07 | 5,39E-06 | Up |
| Rara | 1249,45101 | -0,97463809 | 0,131638341 | 8,52E-16 | 1,38E-13 | Down |
| Jup | 235,9404248 | 1,284267606 | 0,454888002 | 7,14E-06 | 8,09E-05 | Up |
| Cnp | 6752,069168 | 0,653276346 | 0,136331995 | 4,74E-08 | 1,06E-06 | Up |
| Stat5a | 4232,933386 | 0,781513613 | 0,124360423 | 7,63E-12 | 5,05E-10 | Up |
| Plekhh3 | 34,01354151 | -1,500752667 | 0,529125746 | 5,07E-06 | 5,99E-05 | Down |
| Cntnap1 | 15,48734515 | -0,751354302 | 0,599218022 | 0,001177971 | 0,006415768 | Down |
| Etv4 | 842,4524741 | -1,641174357 | 0,183923116 | 2,07E-21 | 8,82E-19 | Down |
| Acbd4 | 2574,740235 | 0,690969105 | 0,12786736 | 1,87E-09 | 6,50E-08 | Up |
| Gm8738 | 127,3668527 | -0,760551707 | 0,272812236 | 5,65E-05 | 0,000488244 | Down |
| Itgb3 | 28796,22896 | 1,017040769 | 0,130070973 | 2,48E-17 | 5,23E-15 | Up |
| Ftsj3 | 39137,72399 | -0,612604368 | 0,088112632 | 1,39E-13 | 1,42E-11 | Down |
| Tcam1 | 48,45755747 | 0,582084156 | 0,347371681 | 0,001974436 | 0,009902697 | Up |
| Cd79b | 160,9966712 | 1,29808273 | 0,368692259 | 6,93E-07 | 1,07E-05 | Up |
| Milr1 | 749,3495395 | -1,093501334 | 0,266761784 | 1,17E-07 | 2,32E-06 | Down |
| Slc16a6 | 11528,65552 | -0,818424325 | 0,129295908 | 4,64E-12 | 3,30E-10 | Down |
| Cd300a | 305,8562464 | 0,839643186 | 0,260436062 | 1,01E-05 | 0,000109689 | Up |
| Cd300ld | 194,4014277 | -0,607054049 | 0,149908699 | 1,53E-06 | 2,13E-05 | Down |
| Rab37 | 416,2420846 | 0,824674112 | 0,206119034 | 6,95E-07 | 1,08E-05 | Up |
| Cd300lf | 3294,058118 | -0,624665243 | 0,107812644 | 2,39E-10 | 1,09E-08 | Down |
| Fads6 | 493,9930731 | 0,80380508 | 0,204888285 | 1,05E-06 | 1,53E-05 | Up |
| Hid1 | 20,88227253 | 1,153774844 | 0,801458753 | 0,000210307 | 0,001491362 | Up |
| Smim5 | 8823,774637 | 0,766381961 | 0,132489206 | 1,67E-10 | 7,98E-09 | Up |
| Rhbdf2 | 7965,901881 | -0,705509689 | 0,11992607 | 1,20E-10 | 5,98E-09 | Down |
| Usp36 | 15953,13352 | -0,659864006 | 0,097464242 | 4,74E-13 | 4,25E-11 | Down |
| Ccdc40 | 263,1998119 | 0,610782362 | 0,209621752 | 9,00E-05 | 0,000728306 | Up |
| Sgsh | 2347,24797 | 1,129118833 | 0,232433223 | 3,02E-09 | 9,98E-08 | Up |

| | | | | | | |
|---|---|---|---|---|---|---|
| Slc26a11 | 69,14630225 | 1,271943802 | 0,447402784 | 6,89E-06 | 7,85E-05 | Up |
| 0610009L18Rik | 162,2274992 | 0,606185398 | 0,221284856 | 0,000154446 | 0,001150542 | Up |
| Tspan10 | 41,90212819 | 0,962180116 | 0,411618422 | 6,80E-05 | 0,000571978 | Up |
| Ccdc137 | 5048,72837 | -0,61083181 | 0,238300537 | 0,000244701 | 0,001704714 | Down |
| Ppp1r27 | 19,09673643 | 0,644858365 | 0,512169495 | 0,002099341 | 0,010403727 | Up |
| Tex19,2 | 27,7198312 | -1,122460869 | 0,462578154 | 3,15E-05 | 0,00029738 | Down |
| Narf | 2989,659917 | 0,969787262 | 0,175790207 | 2,12E-10 | 9,84E-09 | Up |
| 1110002L01Rik | 454,7787592 | -0,946186489 | 0,133693159 | 1,17E-14 | 1,56E-12 | Down |
| Fkbp1b | 190,2514802 | 1,179242058 | 0,220152438 | 1,79E-10 | 8,52E-09 | Up |
| Mycn | 8714,752032 | -0,946797449 | 0,208018671 | 3,33E-08 | 7,91E-07 | Down |
| Lpin1 | 5191,723591 | 0,747904404 | 0,217025795 | 7,96E-06 | 8,91E-05 | Up |
| Greb1 | 5300,365898 | 4,043740443 | 0,557993898 | 1,72E-15 | 2,67E-13 | Up |
| Pqlc3 | 5114,398989 | 0,586048599 | 0,141929824 | 1,18E-06 | 1,69E-05 | Up |
| Odc1 | 90651,84054 | -0,706569263 | 0,115108157 | 2,57E-11 | 1,50E-09 | Down |
| Gm22748 | 19,38472674 | -0,603481903 | 0,509782584 | 0,00282075 | 0,013308132 | Down |
| Gm4419 | 81,5986721 | 0,715046385 | 0,259194199 | 7,90E-05 | 0,000650062 | Up |
| Id2 | 4319,623589 | -0,71774663 | 0,139554891 | 6,72E-09 | 1,96E-07 | Down |
| Hdac9 | 275,840004 | -1,320218648 | 0,309942099 | 3,38E-08 | 7,97E-07 | Down |
| Ifrd1 | 11870,87126 | -0,673010775 | 0,153721809 | 2,99E-07 | 5,20E-06 | Down |
| Egln3 | 62193,80708 | -0,677708493 | 0,126613278 | 2,55E-09 | 8,61E-08 | Down |
| 9330151L19Rik | 239,7470378 | 0,624752822 | 0,211615876 | 7,43E-05 | 0,000616854 | Up |
| Atl1 | 92,87026523 | 0,585090855 | 0,25819713 | 0,000605947 | 0,003656937 | Up |
| Prkch | 22395,79984 | 0,589281705 | 0,080199007 | 8,01E-15 | 1,10E-12 | Up |
| Rhoj | 8706,867025 | -0,960281758 | 0,525564563 | 0,000212431 | 0,00150445 | Down |
| Hspa2 | 331,253434 | 0,80046992 | 0,161690091 | 1,20E-08 | 3,25E-07 | Up |
| Rdh12 | 730,9090474 | 0,720161976 | 0,185106493 | 1,83E-06 | 2,46E-05 | Up |
| 2310015A10Rik | 599,3068281 | 0,83357878 | 0,113260955 | 3,53E-15 | 5,11E-13 | Up |
| Slc8a3 | 59,93000479 | 0,753814136 | 0,284566716 | 8,54E-05 | 0,000695582 | Up |
| Sipa1l1 | 98257,06086 | 0,663074488 | 0,114272675 | 2,13E-10 | 9,86E-09 | Up |
| Zc2hc1c | 52,60094833 | 0,604432669 | 0,278913053 | 0,000671734 | 0,003995388 | Up |
| Tgfb3 | 214,763744 | 1,650585621 | 0,334659095 | 1,84E-09 | 6,44E-08 | Up |
| Ift43 | 351,6689502 | 0,693827854 | 0,141011593 | 2,24E-08 | 5,57E-07 | Up |
| Noxred1 | 51,16723147 | 0,757501217 | 0,428965383 | 0,000586942 | 0,003558717 | Up |
| Serpina3f | 222,9316859 | -1,598075134 | 0,383614806 | 5,22E-08 | 1,15E-06 | Down |
| Trmt61a | 4466,011243 | -0,880934455 | 0,121404908 | 5,35E-15 | 7,55E-13 | Down |
| Nudt14 | 1734,516994 | 0,650560268 | 0,110221123 | 1,21E-10 | 6,04E-09 | Up |
| Ighm | 1457,829526 | 1,579856896 | 0,255293671 | 1,54E-12 | 1,23E-10 | Up |
| Gm9260 | 2937,502245 | -0,614251665 | 0,16275754 | 4,52E-06 | 5,39E-05 | Down |
| Akr1c19 | 354,8247835 | 0,844944964 | 0,230688978 | 2,22E-06 | 2,91E-05 | Up |
| Idi1 | 7542,710232 | 1,186096212 | 0,202726314 | 9,90E-12 | 6,40E-10 | Up |
| Heatr1 | 24524,10792 | -0,598987944 | 0,124266287 | 4,75E-08 | 1,06E-06 | Down |
| Zkscan4 | 181,0267623 | 1,069174944 | 0,226511953 | 7,74E-09 | 2,22E-07 | Up |
| Zfp184 | 127,6383561 | 0,61733071 | 0,215684992 | 0,00010145 | 0,000806876 | Up |
| Prss16 | 802,8789257 | 0,636920748 | 0,205783166 | 4,47E-05 | 0,000398626 | Up |
| Cmah | 23,01845165 | 0,94743862 | 0,577096373 | 0,000311867 | 0,002087853 | Up |
| Rpp40 | 2175,583782 | -0,589471067 | 0,09163408 | 4,74E-12 | 3,36E-10 | Down |
| Eef1e1 | 7193,167973 | -0,871942562 | 0,099769611 | 3,64E-20 | 1,11E-17 | Down |
| Mylip | 1392,91633 | 0,792610649 | 0,123485476 | 3,06E-12 | 2,29E-10 | Up |
| Nol8 | 8292,613356 | -0,632651353 | 0,094855131 | 9,63E-13 | 7,97E-11 | Down |
| Simc1 | 1692,816062 | -0,653434757 | 0,111294216 | 1,45E-10 | 6,99E-09 | Down |
| Nop16 | 15847,78597 | -0,616234007 | 0,105467061 | 1,81E-10 | 8,58E-09 | Down |

| | | | | | | |
|---|---|---|---|---|---|---|
| Unc5a | 833,5031363 | 1,114030571 | 0,215143326 | 6,06E-10 | 2,44E-08 | Up |
| Hk3 | 26800,88562 | -0,614460858 | 0,201796429 | 5,88E-05 | 0,000505928 | Down |
| Prr7 | 516,0034562 | -0,662996917 | 0,127922831 | 6,51E-09 | 1,91E-07 | Down |
| Gkap1 | 118,0438576 | -0,735234866 | 0,234338984 | 2,33E-05 | 0,000227704 | Down |
| Ctla2b | 974,1739352 | 0,841496232 | 0,217858938 | 1,08E-06 | 1,57E-05 | Up |
| Platr25 | 316,5512245 | -0,71126802 | 0,17242813 | 7,50E-07 | 1,14E-05 | Down |
| Fbp2 | 13397,45589 | -1,086688975 | 0,197744815 | 1,20E-10 | 5,98E-09 | Down |
| Fbp1 | 506,5409413 | -1,669428943 | 0,328676136 | 9,28E-10 | 3,56E-08 | Down |
| Fancc | 5048,413678 | 0,853341026 | 0,148806023 | 1,36E-10 | 6,61E-09 | Up |
| Mctp1 | 306,1510972 | 0,948059877 | 0,308967155 | 9,51E-06 | 0,000104394 | Up |
| Mblac2 | 543,9052679 | 0,728393626 | 0,136408319 | 2,31E-09 | 7,92E-08 | Up |
| 4833422C13Rik | 219,3540343 | 0,872210289 | 0,235286589 | 1,63E-06 | 2,23E-05 | Up |
| Dimt1 | 2356,402769 | -0,629047949 | 0,148003571 | 6,07E-07 | 9,54E-06 | Down |
| Emb | 15483,94736 | -0,719814531 | 0,162121961 | 1,90E-07 | 3,50E-06 | Down |
| Ndst2 | 7587,748567 | 0,730572087 | 0,106827732 | 2,50E-13 | 2,33E-11 | Up |
| Zcchc24 | 68,62678106 | 1,349950234 | 0,305928829 | 1,66E-08 | 4,32E-07 | Up |
| Sh3bp5 | 733,0241111 | -0,622984393 | 0,219989571 | 0,000107549 | 0,000847905 | Down |
| Eaf1 | 14776,4521 | -0,688747087 | 0,103230931 | 8,75E-13 | 7,32E-11 | Down |
| Hacl1 | 2742,229687 | 0,606404022 | 0,161639599 | 5,08E-06 | 6,00E-05 | Up |
| Gprin2 | 15,52779118 | 1,063429432 | 0,707929912 | 0,000257958 | 0,001783287 | Up |
| Gm7236 | 106,0647051 | -0,627634396 | 0,215966159 | 8,41E-05 | 0,000686949 | Down |
| Cdkn3 | 756,721289 | 0,65812975 | 0,139569702 | 6,74E-08 | 1,43E-06 | Up |
| Rnase10 | 8,856081127 | -1,438716742 | 1,738992104 | 0,000147227 | 0,001103862 | Down |
| Trac | 148,0237605 | 2,100703423 | 0,330466108 | 1,37E-12 | 1,11E-10 | Up |
| Abhd4 | 11072,85963 | 0,636931263 | 0,111382556 | 3,64E-10 | 1,57E-08 | Up |
| Slc7a7 | 7431,349218 | 0,993071353 | 0,177721954 | 1,22E-10 | 6,05E-09 | Up |
| Gm37271 | 94,0022124 | -0,932706036 | 0,379582205 | 5,69E-05 | 0,000491456 | Down |
| Slc22a17 | 1543,873298 | 0,933993414 | 0,11046199 | 2,55E-19 | 6,90E-17 | Up |
| Myh7 | 38,72488183 | -0,746027154 | 0,447746505 | 0,00073603 | 0,004291085 | Down |
| Dhrs1 | 3902,988367 | 0,687317416 | 0,118219836 | 1,89E-10 | 8,88E-09 | Up |
| Cideb | 424,6151422 | 0,686544788 | 0,198945157 | 1,07E-05 | 0,00011576 | Up |
| Ltb4r2 | 305,3376719 | 0,650832158 | 0,263380156 | 0,000244979 | 0,001705917 | Up |
| Adcy4 | 351,8680846 | -0,72976422 | 0,15481076 | 5,24E-08 | 1,15E-06 | Down |
| Ripk3 | 124,2185648 | -2,464548686 | 0,306278049 | 2,91E-18 | 6,65E-16 | Down |
| Mcpt1 | 70,48400238 | 1,306947807 | 0,480550919 | 9,26E-06 | 0,000102019 | Up |
| Mcpt2 | 338,2446608 | 0,989780711 | 0,161384496 | 4,80E-12 | 3,39E-10 | Up |
| Cryl1 | 1527,099161 | 0,603762992 | 0,076699005 | 1,43E-16 | 2,55E-14 | Up |
| Xpo4 | 13478,40596 | -0,654791227 | 0,098677017 | 1,18E-12 | 9,69E-11 | Down |
| Fgf9 | 1434,587232 | -0,809353749 | 0,080454813 | 2,19E-25 | 2,22E-22 | Down |
| Phf11b | 109,8684856 | 0,609327137 | 0,293389982 | 0,000790892 | 0,004568255 | Up |
| Sgcg | 7,308423166 | 0,929166129 | 0,807111253 | 0,00056477 | 0,003450075 | Up |
| Zfp395 | 2940,721273 | 0,646550306 | 0,127550961 | 1,22E-08 | 3,30E-07 | Up |
| Sorbs3 | 66,34212195 | 0,92723865 | 0,391567226 | 7,29E-05 | 0,000607998 | Up |
| Slc39a14 | 6858,21372 | -0,745131033 | 0,10854531 | 1,98E-13 | 1,90E-11 | Down |
| Polr3d | 3655,168708 | -0,885574881 | 0,142446596 | 6,02E-12 | 4,14E-10 | Down |
| Dok2 | 12679,57408 | -0,803311758 | 0,088678329 | 3,46E-21 | 1,40E-18 | Down |
| Gm4285 | 147,8286904 | 0,621006571 | 0,196704121 | 3,97E-05 | 0,000361459 | Up |
| Slain1 | 4991,394592 | 0,629084912 | 0,100520572 | 1,41E-11 | 8,68E-10 | Up |
| Dzip1 | 190,1343369 | 0,729465608 | 0,187894794 | 1,80E-06 | 2,44E-05 | Up |
| Mtmr12 | 10351,46479 | 0,587377696 | 0,091908129 | 6,24E-12 | 4,27E-10 | Up |
| Pdzd2 | 426,2739829 | 1,251032823 | 0,324935283 | 2,11E-07 | 3,82E-06 | Up |

| | | | | | | |
|---|---|---|---|---|---|---|
| Gm2606 | 50,5075093 | 1,048342889 | 0,85082618 | 0,000353549 | 0,002320521 | Up |
| Trio | 13127,54574 | -0,96804499 | 0,074613553 | 1,19E-40 | 9,66E-37 | Down |
| Pop1 | 6346,479171 | -0,803632998 | 0,112652975 | 2,22E-14 | 2,77E-12 | Down |
| Snx31 | 208,4290402 | 0,610758237 | 0,217557056 | 0,00012366 | 0,00095267 | Up |
| Lrp12 | 26105,19814 | -0,889781424 | 0,150482911 | 3,74E-11 | 2,11E-09 | Down |
| Nov | 32,77256783 | -2,224085645 | 0,733968255 | 1,29E-06 | 1,82E-05 | Down |
| Trib1 | 185,8413785 | -0,647655766 | 0,24683748 | 0,000167719 | 0,001226892 | Down |
| Myc | 52082,33665 | -0,752986036 | 0,089880589 | 1,79E-18 | 4,22E-16 | Down |
| Tmem71 | 261,7310985 | 0,708406367 | 0,260872391 | 9,23E-05 | 0,000743944 | Up |
| Sla | 19085,22512 | 1,115330867 | 0,156195817 | 2,36E-15 | 3,52E-13 | Up |
| St3gal1 | 9612,884407 | 0,793262364 | 0,112956467 | 5,20E-14 | 6,01E-12 | Up |
| 9030619P08Rik | 3853,176387 | 1,668859902 | 0,161261276 | 2,48E-27 | 3,35E-24 | Up |
| Kifc2 | 11,06650527 | 0,854991675 | 0,680976078 | 0,000728863 | 0,004261547 | Up |
| Apol10b | 333,8324629 | 0,672610296 | 0,22905527 | 6,02E-05 | 0,000515635 | Up |
| Foxred2 | 326,5017059 | 0,751157104 | 0,28821929 | 9,70E-05 | 0,000776482 | Up |
| Tst | 254,111652 | 0,580695352 | 0,212789884 | 0,000183875 | 0,001326307 | Up |
| Il2rb | 2124,240064 | -0,973899731 | 0,159184751 | 5,89E-12 | 4,07E-10 | Down |
| Mfng | 9836,646962 | -0,822588716 | 0,086496314 | 4,47E-23 | 2,79E-20 | Down |
| Maff | 559,8860758 | -1,68896321 | 0,165022647 | 8,82E-27 | 1,02E-23 | Down |
| Gm16576 | 461,5269038 | 0,712604018 | 0,150711922 | 5,26E-08 | 1,16E-06 | Up |
| Grap2 | 44,71983661 | 1,994896054 | 0,527709925 | 2,12E-07 | 3,83E-06 | Up |
| Mkl1 | 9561,340033 | 0,631182515 | 0,120427112 | 5,16E-09 | 1,57E-07 | Up |
| Ttll1 | 2638,371487 | 0,613424644 | 0,08821158 | 1,38E-13 | 1,42E-11 | Up |
| 5031439G07Rik | 24803,44948 | 0,630163385 | 0,148590361 | 6,30E-07 | 9,86E-06 | Up |
| Creld2 | 9921,911313 | -0,627165369 | 0,109743614 | 3,78E-10 | 1,62E-08 | Down |
| Odf3b | 398,2750906 | 0,630575451 | 0,142539462 | 2,81E-07 | 4,93E-06 | Up |
| Mapk8ip2 | 138,2445756 | 0,987727539 | 0,23179785 | 9,61E-08 | 1,95E-06 | Up |
| Arsa | 6400,399702 | 0,630334147 | 0,101023811 | 1,59E-11 | 9,61E-10 | Up |
| Shank3 | 691,8001706 | -0,87500682 | 0,279493195 | 1,12E-05 | 0,000120596 | Down |
| Prickle1 | 609,2121696 | 1,110329466 | 0,236763836 | 7,48E-09 | 2,15E-07 | Up |
| Pus7l | 4352,454358 | -0,627469261 | 0,09111265 | 2,19E-13 | 2,08E-11 | Down |
| Hdac7 | 1188,666123 | -0,588985742 | 0,314724668 | 0,001355974 | 0,007206247 | Down |
| 1700120C14Rik | 871,9895736 | -0,736386839 | 0,290149445 | 0,000125574 | 0,000965151 | Down |
| Mettl7a1 | 3337,095574 | 1,452381338 | 0,242103812 | 3,64E-12 | 2,61E-10 | Up |
| Pou6f1 | 59,0567969 | 0,744577201 | 0,289416922 | 0,000109716 | 0,00086121 | Up |
| C330013E15Rik | 87,38201403 | 0,719272298 | 0,252860375 | 6,07E-05 | 0,00051926 | Up |
| Galnt6 | 13781,09769 | -0,633225582 | 0,096434113 | 1,92E-12 | 1,50E-10 | Down |
| Acvrl1 | 417,3961895 | -3,220481944 | 0,471643306 | 2,81E-15 | 4,15E-13 | Down |
| Grasp | 173,9788717 | -0,586199373 | 0,206393026 | 0,000128494 | 0,000983355 | Down |
| Tns2 | 900,18026 | 0,76899704 | 0,147884209 | 4,03E-09 | 1,28E-07 | Up |
| Csad | 543,1088425 | 0,808140764 | 0,126256868 | 3,14E-12 | 2,32E-10 | Up |
| Calcoco1 | 1748,419943 | 0,828615893 | 0,158208246 | 2,40E-09 | 8,19E-08 | Up |
| Nfe2 | 33328,36694 | 0,723591926 | 0,152983833 | 4,99E-08 | 1,11E-06 | Up |
| Gpr84 | 315,0824204 | -1,342306674 | 0,347464771 | 1,77E-07 | 3,29E-06 | Down |
| Zfp385a | 4708,771623 | 0,921005255 | 0,153313424 | 1,70E-11 | 1,02E-09 | Up |
| Nlrc3 | 795,6766529 | 1,105652317 | 0,174659493 | 6,71E-13 | 5,76E-11 | Up |
| Sep 12 | 4,559827867 | 0,595752004 | 1,109563897 | 0,00112996 | 0,006189601 | Up |
| Socs1 | 378,5095863 | -0,840594726 | 0,189096195 | 1,00E-07 | 2,01E-06 | Down |
| Snx29 | 528,914144 | 0,89858018 | 0,175072155 | 2,67E-09 | 8,99E-08 | Up |
| Mkl2 | 9248,303095 | 0,730068869 | 0,122995925 | 8,06E-11 | 4,18E-09 | Up |
| Tmem191c | 126,9352272 | 0,595031098 | 0,350725344 | 0,00174996 | 0,008979478 | Up |

| | | | | | | |
|---|---|---|---|---|---|---|
| Klhl24 | 2207,660239 | 0,747601802 | 0,140754534 | 2,48E-09 | 8,40E-08 | Up |
| Fam131a | 401,8423097 | 0,80675443 | 0,185040142 | 1,75E-07 | 3,27E-06 | Up |
| Etv5 | 5632,118427 | -1,54864762 | 0,180474789 | 2,78E-20 | 8,97E-18 | Down |
| St6gal1 | 834,2815111 | -0,663702908 | 0,500810576 | 0,001785865 | 0,009134823 | Down |
| Tfrc | 34594,50296 | -0,88565214 | 0,1954052 | 5,23E-08 | 1,15E-06 | Down |
| Gm15657 | 115,7387431 | 0,708150285 | 0,286798695 | 0,000175505 | 0,001270666 | Up |
| Muc13 | 24196,78658 | 0,598173474 | 0,17915898 | 2,40E-05 | 0,000234942 | Up |
| Stfa2l1 | 107,2059199 | -3,315738728 | 0,71730465 | 1,93E-09 | 6,70E-08 | Down |
| Gm5483 | 452,7039256 | -5,09889227 | 0,672279979 | 2,20E-15 | 3,34E-13 | Down |
| Gm5416 | 22,91861969 | -4,461117314 | 1,495114072 | 7,38E-07 | 1,13E-05 | Down |
| 2010005H15Rik | 294,9231517 | -4,953045274 | 0,71473987 | 1,74E-13 | 1,70E-11 | Down |
| Stfa1 | 435,2804446 | -4,020607221 | 0,868175514 | 5,06E-09 | 1,54E-07 | Down |
| BC100530 | 158,3176464 | -4,070237586 | 0,981450736 | 3,41E-08 | 8,03E-07 | Down |
| Stfa2 | 66,66232615 | -6,77314195 | 1,169173704 | 1,38E-10 | 6,70E-09 | Down |
| Stfa3 | 23,35549363 | -4,810153874 | 1,736366982 | 9,38E-07 | 1,39E-05 | Down |
| Cd80 | 93,93014878 | -0,881911691 | 0,360868555 | 7,40E-05 | 0,000615101 | Down |
| Qtrtd1 | 3150,644814 | -0,613159984 | 0,113072378 | 2,00E-09 | 6,90E-08 | Down |
| Zbtb11os1 | 261,0842517 | 0,78841613 | 0,21653624 | 3,26E-06 | 4,09E-05 | Up |
| Gm9027 | 90,90545374 | -0,763004321 | 0,212568736 | 4,48E-06 | 5,35E-05 | Down |
| Urb1 | 4311,844986 | -0,651006617 | 0,151023458 | 4,34E-07 | 7,13E-06 | Down |
| Wrb | 2437,685251 | -0,902064122 | 0,116152488 | 9,46E-17 | 1,74E-14 | Down |
| Igf2r | 1450,221384 | -0,876085216 | 0,498343742 | 0,000338455 | 0,002235016 | Down |
| Acat2 | 11616,98755 | 0,597964364 | 0,159782875 | 5,45E-06 | 6,36E-05 | Up |
| Hcfc1r1 | 1309,510573 | 0,611292408 | 0,124471367 | 2,94E-08 | 7,05E-07 | Up |
| Tnfrsf12a | 1265,200175 | -0,83626716 | 0,165406504 | 5,82E-09 | 1,74E-07 | Down |
| Abca3 | 75,97703443 | 0,669344785 | 0,283570894 | 0,000287593 | 0,001953623 | Up |
| Haghl | 754,6905472 | 0,745413295 | 0,214503309 | 7,31E-06 | 8,24E-05 | Up |
| Fam173a | 2319,4573 | 0,619981857 | 0,106472008 | 2,03E-10 | 9,44E-09 | Up |
| Wdr90 | 3890,75529 | 0,834495679 | 0,172748549 | 1,77E-08 | 4,54E-07 | Up |
| Nhlrc4 | 124,3654615 | 0,900689275 | 0,338321911 | 3,84E-05 | 0,000352799 | Up |
| Rab11fip3 | 208,3868459 | 1,773620012 | 0,261990471 | 6,70E-14 | 7,40E-12 | Up |
| Crebrf | 888,3180466 | 0,756172868 | 0,21826725 | 7,16E-06 | 8,11E-05 | Up |
| Cpne5 | 368,5817459 | 1,145375246 | 0,256665217 | 1,93E-08 | 4,89E-07 | Up |
| Pim1 | 35421,75936 | -0,943602634 | 0,101883163 | 1,72E-22 | 9,01E-20 | Down |
| Notch3 | 180,1590764 | 0,653911111 | 0,273018638 | 0,000290302 | 0,001965837 | Up |
| H2-Oa | 451,9972525 | 0,657036204 | 0,186933746 | 9,85E-06 | 0,000107354 | Up |
| H2-DMa | 141,6327654 | 0,668951035 | 0,223624995 | 5,20E-05 | 0,000454173 | Up |
| Psmb9 | 6708,266725 | 0,804244317 | 0,145292241 | 5,60E-10 | 2,28E-08 | Up |
| Tap1 | 9558,796644 | 0,791990634 | 0,133067691 | 5,51E-11 | 2,96E-09 | Up |
| Tap2 | 24397,07219 | 0,913286892 | 0,15399154 | 2,85E-11 | 1,65E-09 | Up |
| H2-Ob | 188,701423 | 1,358992105 | 0,409010794 | 1,30E-06 | 1,83E-05 | Up |
| Vwa7 | 200,232371 | 0,661700386 | 0,169246319 | 2,19E-06 | 2,89E-05 | Up |
| Sapcd1 | 21,5082833 | 0,643354582 | 0,451592686 | 0,001831069 | 0,009316115 | Up |
| Msh5 | 205,4298374 | 1,054084106 | 0,375615364 | 1,36E-05 | 0,000143117 | Up |
| Tnf | 575,7083511 | -1,877412909 | 0,40546404 | 9,39E-09 | 2,65E-07 | Down |
| Lta | 13,99742915 | -2,511782101 | 1,00337453 | 3,07E-06 | 3,88E-05 | Down |
| Gm20443 | 18,38509011 | 1,092688088 | 0,552323644 | 0,000100094 | 0,00079805 | Up |
| Gm10499 | 78,20366555 | -1,467508667 | 0,581373929 | 1,21E-05 | 0,000129339 | Down |
| Rnf39 | 26,18132706 | 1,000157518 | 0,437149056 | 6,64E-05 | 0,000560618 | Up |
| Enpp5 | 1931,277406 | 0,712921564 | 0,204801047 | 8,46E-06 | 9,39E-05 | Up |
| Slc29a1 | 6326,353284 | -0,71663221 | 0,124947502 | 2,72E-10 | 1,22E-08 | Down |

| | | | | | | |
|---|---|---|---|---|---|---|
| Klc4 | 2245,837766 | 0,839887259 | 0,130070207 | 1,79E-12 | 1,41E-10 | Up |
| Bysl | 13254,14554 | -0,623330915 | 0,110118926 | 5,19E-10 | 2,12E-08 | Down |
| Apobec2 | 31,47280923 | 2,831777884 | 0,44701304 | 1,51E-13 | 1,52E-11 | Up |
| Adgre4 | 17,11310632 | 0,755258808 | 0,501657387 | 0,000888063 | 0,005055212 | Up |
| Stap2 | 1615,345631 | -0,867526187 | 0,099694326 | 5,25E-20 | 1,58E-17 | Down |
| Vmac | 530,7294514 | 0,818118904 | 0,13044817 | 6,71E-12 | 4,54E-10 | Up |
| Tnfsf14 | 2354,701557 | -1,200619024 | 0,138931866 | 9,54E-21 | 3,44E-18 | Down |
| Adgre1 | 31351,15199 | -1,350283275 | 0,168102249 | 1,53E-18 | 3,66E-16 | Down |
| Ypel5 | 2742,981752 | 0,697707149 | 0,212710184 | 1,80E-05 | 0,000181742 | Up |
| Lbh | 2209,153057 | 0,913084025 | 0,151905192 | 1,76E-11 | 1,05E-09 | Up |
| Xdh | 28,78667226 | 0,765528615 | 0,575305456 | 0,001031745 | 0,0057231 | Up |
| Nlrc4 | 961,5915566 | 0,582030386 | 0,104800682 | 1,01E-09 | 3,86E-08 | Up |
| Thada | 8779,860097 | -0,596477361 | 0,075346251 | 1,00E-16 | 1,80E-14 | Down |
| Cables1 | 446,507358 | 1,339676129 | 0,265720912 | 7,60E-10 | 2,99E-08 | Up |
| Cabyr | 87,62900569 | 0,967339864 | 0,336729307 | 1,57E-05 | 0,000161527 | Up |
| Osbpl1a | 5962,158391 | 0,629427335 | 0,109328804 | 2,94E-10 | 1,29E-08 | Up |
| Slc39a6 | 7023,671152 | -0,62458102 | 0,088173693 | 5,49E-14 | 6,23E-12 | Down |
| Celf4 | 147,4746177 | -1,629399428 | 0,76070531 | 2,18E-05 | 0,000214183 | Down |
| Bin1 | 1945,858156 | 0,641569079 | 0,131068565 | 2,96E-08 | 7,09E-07 | Up |
| Hbegf | 1402,236803 | -1,649965062 | 0,155336634 | 1,29E-28 | 2,09E-25 | Down |
| Wdr55 | 8507,810935 | -0,604435628 | 0,147975416 | 1,34E-06 | 1,88E-05 | Down |
| Spry4 | 227,1957364 | -3,012776149 | 0,275618225 | 1,62E-31 | 5,27E-28 | Down |
| Tubb6 | 18989,66917 | -0,775608479 | 0,073917218 | 3,08E-27 | 3,84E-24 | Down |
| Mc5r | 1407,292664 | 0,927912146 | 0,341150831 | 2,89E-05 | 0,000276511 | Up |
| Katnal2 | 143,4202902 | 0,80525253 | 0,155608672 | 3,80E-09 | 1,21E-07 | Up |
| Cpt1a | 11003,21757 | 0,664196048 | 0,16399642 | 1,24E-06 | 1,76E-05 | Up |
| Acy3 | 548,8331198 | 0,707206132 | 0,162187925 | 2,83E-07 | 4,95E-06 | Up |
| Gstp3 | 31,36297781 | -0,911122168 | 0,601246503 | 0,000436201 | 0,002764257 | Down |
| Coro1b | 28071,18604 | 0,656990902 | 0,078408683 | 2,26E-18 | 5,23E-16 | Up |
| Tbc1d10c | 2904,422825 | 0,777835421 | 0,143100028 | 1,11E-09 | 4,18E-08 | Up |
| Pold4 | 1546,038333 | 0,798722841 | 0,199896795 | 8,17E-07 | 1,23E-05 | Up |
| Ssh3 | 2039,440251 | 0,660210103 | 0,151786319 | 3,55E-07 | 6,00E-06 | Up |
| Slc29a2 | 2462,474498 | -1,003376041 | 0,175184014 | 5,09E-11 | 2,75E-09 | Down |
| Tsga10ip | 1120,594777 | 0,762736782 | 0,137284852 | 6,19E-10 | 2,49E-08 | Up |
| Snx32 | 189,4482655 | 0,760401372 | 0,280334097 | 6,95E-05 | 0,00058298 | Up |
| Tm7sf2 | 449,7034108 | 0,921817559 | 0,230919871 | 4,19E-07 | 6,93E-06 | Up |
| Naaladl1 | 190,7691508 | 0,619736633 | 0,167491258 | 5,87E-06 | 6,80E-05 | Up |
| Cdc42bpg | 102,9241193 | 0,790830929 | 0,288037493 | 5,37E-05 | 0,000467476 | Up |
| Map4k2 | 2216,824695 | 0,593000722 | 0,199093614 | 8,11E-05 | 0,000664799 | Up |
| Fads3 | 83,08442898 | 0,711586191 | 0,328205906 | 0,000346225 | 0,00228168 | Up |
| Cyb561a3 | 2272,109887 | 0,591400406 | 0,106383366 | 9,68E-10 | 3,70E-08 | Up |
| Pga5 | 28,72215418 | 0,984252911 | 0,72059312 | 0,000398307 | 0,002563477 | Up |
| Cd5 | 7,198152361 | 0,74930661 | 0,791868269 | 0,001186615 | 0,006449858 | Up |
| Ccdc86 | 8287,309118 | -0,627748397 | 0,1336347 | 8,00E-08 | 1,67E-06 | Down |
| Ms4a3 | 1878,313563 | -1,872099173 | 0,50845718 | 3,14E-07 | 5,41E-06 | Down |
| Gna14 | 108,4615706 | 1,94172185 | 0,433136776 | 1,75E-08 | 4,50E-07 | Up |
| Trpm6 | 13,08727511 | -2,83328857 | 1,263932907 | 4,27E-06 | 5,12E-05 | Down |
| Kank1 | 118,0666616 | -1,431279068 | 0,355169079 | 8,60E-08 | 1,77E-06 | Down |
| Insl6 | 371,0589277 | 0,74209942 | 0,128172752 | 1,81E-10 | 8,57E-09 | Up |
| Cd274 | 1913,758696 | -0,647334665 | 0,108491989 | 8,31E-11 | 4,28E-09 | Down |
| Pdcd1lg2 | 271,738588 | -0,610547594 | 0,282501248 | 0,000654367 | 0,003907656 | Down |

| | | | | | | |
|---|---|---|---|---|---|---|
| Ifit1bl1 | 24,88496646 | 0,909558719 | 0,519394573 | 0,00029826 | 0,002011336 | Up |
| Rpp30 | 3757,220907 | -0,620656156 | 0,126175001 | 2,77E-08 | 6,67E-07 | Down |
| Frat2 | 3581,670133 | 0,690775703 | 0,149198825 | 9,02E-08 | 1,84E-06 | Up |
| Rrp12 | 16668,34226 | -0,725392555 | 0,11097621 | 1,93E-12 | 1,50E-10 | Down |
| Marveld1 | 4729,305586 | 0,697917691 | 0,075586583 | 1,11E-21 | 5,00E-19 | Up |
| Pyroxd2 | 613,9148453 | 0,70124227 | 0,28946973 | 0,000204905 | 0,001456874 | Up |
| Gm16244 | 65,9803612 | 0,690778969 | 0,253471687 | 9,94E-05 | 0,000794033 | Up |
| Cpn1 | 84,37942997 | 1,034867244 | 0,226204177 | 1,83E-08 | 4,68E-07 | Up |
| Scd2 | 130084,5421 | 0,604677331 | 0,203671994 | 7,86E-05 | 0,000646966 | Up |
| Kazald1 | 80,18685946 | 1,201074006 | 0,355095034 | 1,37E-06 | 1,92E-05 | Up |
| Pdcd11 | 23467,26381 | -0,583941073 | 0,128979261 | 2,01E-07 | 3,68E-06 | Down |
| Obfc1 | 1182,931479 | 0,588820728 | 0,104305738 | 5,95E-10 | 2,41E-08 | Up |
| Gm19557 | 53,97645203 | 1,11284962 | 0,422033236 | 1,83E-05 | 0,000184911 | Up |
| Pdcd4 | 1960,934207 | 1,149386784 | 0,1749988 | 1,14E-13 | 1,20E-11 | Up |
| Slc18a2 | 13240,57545 | 0,669477331 | 0,122096942 | 1,28E-09 | 4,75E-08 | Up |
| Ppp1r3f | 123,3944716 | 0,854558343 | 0,186932306 | 5,27E-08 | 1,16E-06 | Up |
| Pim2 | 2848,311864 | -0,7435958 | 0,115568095 | 3,50E-12 | 2,56E-10 | Down |
| Glod5 | 22,97971437 | 0,970828977 | 0,533783063 | 0,000207686 | 0,001474063 | Up |
| Cfp | 7659,197069 | -0,650516609 | 0,282592827 | 0,000370122 | 0,002407872 | Down |
| Rbmx | 8125,585353 | -0,826573514 | 0,142522718 | 1,10E-10 | 5,56E-09 | Down |
| Pcyt1b | 237,5326495 | 1,415558367 | 0,276441168 | 5,11E-10 | 2,09E-08 | Up |
| Foxo4 | 2174,821687 | 0,752089723 | 0,124966585 | 4,50E-11 | 2,49E-09 | Up |
| Diaph2 | 7804,347423 | -0,668125088 | 0,212834869 | 3,30E-05 | 0,000309852 | Down |
| Timm8a1 | 2632,332527 | -0,620567474 | 0,123956372 | 1,78E-08 | 4,56E-07 | Down |
| Armcx1 | 47,24106716 | 1,656768304 | 0,579264932 | 4,00E-06 | 4,84E-05 | Up |
| Tsc22d3 | 849,7772306 | 1,077814668 | 0,223524903 | 4,50E-09 | 1,40E-07 | Up |

**CM-vs-CM-IL3-72**

| | baseMean | log2FoldChange | lfcSE | pvalue | padj | expression |
|---|---|---|---|---|---|---|
| Paqr8 | 6633,17564 | 0,789489923 | 0,116250989 | 1,99E-13 | 1,41E-11 | Up |
| Tmem14a | 317,0076962 | 0,704367305 | 0,180549232 | 1,14E-06 | 1,67E-05 | Up |
| Gm26580 | 133,7585049 | 0,617917378 | 0,236619115 | 0,000138865 | 0,001058781 | Up |
| Arid5a | 5828,508749 | -0,582277174 | 0,113304609 | 4,72E-09 | 1,28E-07 | Down |
| Fam178b | 26,82373702 | 2,803877596 | 0,786735807 | 2,25E-06 | 3,00E-05 | Up |
| Aff3 | 4375,805027 | 1,032835832 | 0,246801596 | 8,78E-08 | 1,72E-06 | Up |
| Il1rl1 | 109,3455273 | 0,814448953 | 0,250441364 | 7,41E-06 | 8,53E-05 | Up |
| Coq10b | 886,827566 | -0,657645556 | 0,140275746 | 4,08E-08 | 8,76E-07 | Down |
| Aox2 | 25,59447421 | 0,715507977 | 0,485086674 | 0,001116038 | 0,006205524 | Up |
| Ino80dos | 114,8376389 | 1,015146443 | 0,254252563 | 2,10E-07 | 3,76E-06 | Up |
| Kansl1l | 2153,789982 | 0,835358108 | 0,139286403 | 2,60E-11 | 1,22E-09 | Up |
| Slc11a1 | 107,6252626 | -0,960653999 | 0,981305314 | 0,001435992 | 0,007621514 | Down |
| Wnt10a | 66,15698044 | 0,716087012 | 0,299691799 | 0,00014783 | 0,001117156 | Up |
| Gm15433 | 27,06492675 | -2,464493955 | 0,687240497 | 2,19E-06 | 2,95E-05 | Down |
| C130026I21Rik | 154,1012673 | 0,638377798 | 0,32240037 | 0,000603996 | 0,003693812 | Up |
| Ramp1 | 6132,934336 | 0,627867314 | 0,17032056 | 3,46E-06 | 4,41E-05 | Up |
| Hes6 | 2039,661949 | 0,93185422 | 0,10766582 | 4,74E-20 | 9,86E-18 | Up |
| Sned1 | 162,9354004 | -0,642248672 | 0,238064641 | 9,46E-05 | 0,000765251 | Down |
| Gm7889 | 93,02605462 | 0,636175829 | 0,212344092 | 3,91E-05 | 0,000359253 | Up |
| Gm6086 | 66,53827624 | 0,657841971 | 0,448767918 | 0,001477003 | 0,007813618 | Up |

| | | | | | | |
|---|---|---|---|---|---|---|
| Pdcd1 | 212,3635005 | 4,536924276 | 0,58700052 | 3,25E-17 | 4,25E-15 | Up |
| Phlpp1 | 2041,35478 | 0,913158091 | 0,156631886 | 4,48E-11 | 1,94E-09 | Up |
| Tfcp2l1 | 8,265693963 | -0,741647185 | 0,946568284 | 0,00304345 | 0,014280594 | Down |
| Tmem185b | 9566,073431 | -0,731747965 | 0,084477958 | 1,10E-19 | 2,15E-17 | Down |
| Map3k19 | 32,01209998 | 1,429475144 | 0,494606611 | 7,05E-06 | 8,15E-05 | Up |
| Zranb3 | 3159,857756 | 0,64864659 | 0,117237814 | 5,00E-10 | 1,74E-08 | Up |
| Cxcr4 | 7686,079561 | 0,702651501 | 0,117118282 | 3,32E-11 | 1,50E-09 | Up |
| Lax1 | 1870,023799 | 0,654460434 | 0,111737106 | 7,64E-11 | 3,15E-09 | Up |
| Atp2b4 | 17240,5454 | 0,78402228 | 0,083776055 | 2,03E-22 | 5,32E-20 | Up |
| Btg2 | 1120,205726 | 1,58012406 | 0,160227366 | 5,02E-25 | 1,63E-22 | Up |
| Elf3 | 18,41098577 | 0,585181476 | 0,579502168 | 0,004149035 | 0,018428167 | Up |
| Rgs2 | 4394,600716 | 1,337080699 | 0,117701705 | 9,40E-33 | 8,47E-30 | Up |
| Rgs1 | 88,76523314 | 1,60610256 | 0,349684112 | 1,59E-08 | 3,79E-07 | Up |
| BC034090 | 128,6157828 | -0,61166701 | 0,24758362 | 0,000213345 | 0,001528941 | Down |
| Ankrd45 | 18,52547862 | -1,957667676 | 0,850799087 | 3,76E-05 | 0,000347373 | Down |
| Nme7 | 497,7444183 | 0,712204272 | 0,202479813 | 4,71E-06 | 5,75E-05 | Up |
| Tbx19 | 183,1212648 | 1,176316576 | 0,247564745 | 4,00E-09 | 1,10E-07 | Up |
| Fcrlb | 7,982514978 | -1,239432186 | 1,107084463 | 0,000709221 | 0,004224345 | Down |
| Fcgr4 | 32,96658856 | -2,225118836 | 0,763061711 | 9,92E-06 | 0,000109324 | Down |
| Fcgr3 | 445,2897942 | -2,735916338 | 0,48857758 | 1,22E-09 | 3,82E-08 | Down |
| Cd244 | 2297,6769 | 0,834584499 | 0,162036991 | 2,71E-09 | 7,80E-08 | Up |
| Ly9 | 34,8917493 | 0,835942447 | 0,523238192 | 0,000556339 | 0,003436088 | Up |
| A630035G10Rik | 47,50261419 | 0,899601718 | 0,404693333 | 0,000101228 | 0,000810272 | Up |
| Slamf1 | 80,96343175 | -0,683879896 | 0,532915663 | 0,001800623 | 0,009192925 | Down |
| Gm37065 | 26,68573509 | -0,691287973 | 0,944834226 | 0,003631861 | 0,016506146 | Down |
| Igsf8 | 11644,53898 | -0,943249172 | 0,137256209 | 4,80E-14 | 3,91E-12 | Down |
| Slamf9 | 17,06430912 | -3,603352112 | 0,723968443 | 1,04E-08 | 2,58E-07 | Down |
| Ifi207 | 8,319987667 | -2,333274164 | 2,067070303 | 9,71E-05 | 0,000782181 | Down |
| Ifi203 | 624,3994017 | 0,788418524 | 0,236462961 | 6,37E-06 | 7,45E-05 | Up |
| Spta1 | 352,3317916 | 3,620315748 | 0,285670157 | 2,40E-38 | 3,54E-35 | Up |
| Olfr248 | 16,25454946 | 2,238472573 | 0,887460824 | 2,26E-05 | 0,000224544 | Up |
| Olfr414 | 16,69962429 | 2,954051829 | 0,964030301 | 5,77E-06 | 6,84E-05 | Up |
| Kmo | 301,6391008 | 0,622335263 | 0,378897967 | 0,001304349 | 0,007068493 | Up |
| H3f3aos | 23,41425315 | 0,807172899 | 0,447066682 | 0,000388304 | 0,002560257 | Up |
| Enah | 45,52905557 | -0,870464984 | 0,808563219 | 0,001486485 | 0,00784843 | Down |
| Degs1 | 9673,181637 | -1,035830179 | 0,117958888 | 5,62E-21 | 1,34E-18 | Down |
| Slc30a1 | 697,9631187 | -0,650432229 | 0,130015177 | 8,65E-09 | 2,18E-07 | Down |
| Hsd11b1 | 61,53172636 | -0,768043765 | 0,528020068 | 0,000945869 | 0,005399974 | Down |
| Echdc3 | 48,56900599 | 1,392361671 | 0,320403246 | 2,84E-08 | 6,31E-07 | Up |
| Mrc1 | 96,22464949 | -1,126445587 | 0,438823745 | 2,21E-05 | 0,000220614 | Down |
| Il1f9 | 11,44569477 | -1,396530355 | 1,427731792 | 0,000591018 | 0,003621323 | Down |
| Slc34a3 | 164,5404617 | 3,632180773 | 0,429444889 | 6,98E-19 | 1,22E-16 | Up |
| Ptgds | 5,043273106 | 0,647325524 | 0,849682115 | 0,00421649 | 0,018676647 | Up |
| Qsox2 | 6293,675937 | -0,62411198 | 0,060043272 | 4,68E-27 | 1,95E-24 | Down |
| Lcn4 | 258,9932678 | 1,38839703 | 0,442142059 | 3,12E-06 | 4,01E-05 | Up |
| Slc2a6 | 1861,671792 | -0,922949673 | 0,275115529 | 3,26E-06 | 4,18E-05 | Down |
| Spaca9 | 236,2926072 | 1,798178102 | 0,210069902 | 2,94E-19 | 5,49E-17 | Up |
| Ak8 | 47,89398858 | 2,156851693 | 0,438015629 | 1,75E-08 | 4,12E-07 | Up |
| Pkn3 | 3143,026406 | 0,797938481 | 0,199236919 | 5,35E-07 | 8,63E-06 | Up |
| Zer1 | 3262,2588 | 0,882079341 | 0,10783347 | 3,99E-18 | 6,12E-16 | Up |
| Hmcn2 | 28,39534262 | -0,861455806 | 0,689310169 | 0,001084333 | 0,006060389 | Down |

| | | | | | | |
|---|---|---|---|---|---|---|
| Qrfp | 115,3309365 | 0,945913796 | 0,265855716 | 1,46E-06 | 2,06E-05 | Up |
| Lcn2 | 1442,891879 | -2,887597247 | 0,642042253 | 1,54E-07 | 2,86E-06 | Down |
| Sh2d3c | 4594,635707 | 0,580725568 | 0,149976856 | 1,97E-06 | 2,68E-05 | Up |
| Fam129b | 2719,526399 | -0,632227427 | 0,190123387 | 1,30E-05 | 0,000139091 | Down |
| Angptl2 | 299,2275307 | 1,381442328 | 0,267330619 | 4,78E-10 | 1,68E-08 | Up |
| Hspa5 | 166180,2454 | -0,839862263 | 0,145345723 | 9,09E-11 | 3,70E-09 | Down |
| Fmnl2 | 16,35420359 | -0,739333299 | 0,885698619 | 0,002962305 | 0,013972402 | Down |
| Nr4a2 | 1421,172583 | 0,887116279 | 0,143131885 | 5,87E-12 | 3,18E-10 | Up |
| Tanc1 | 179,9517431 | -0,658671686 | 0,580760959 | 0,002546797 | 0,012309141 | Down |
| Galnt3 | 6167,877876 | -0,805500885 | 0,086674539 | 3,54E-22 | 8,99E-20 | Down |
| Dhrs9 | 47,75226993 | -1,771669856 | 0,875635355 | 7,16E-05 | 0,000603028 | Down |
| Ttc30b | 97,98092554 | 0,950649652 | 0,234337455 | 2,15E-07 | 3,84E-06 | Up |
| Plekha3 | 4718,743872 | -0,655960592 | 0,090706064 | 8,48E-15 | 7,86E-13 | Down |
| Cerkl | 138,9130571 | 0,669433241 | 0,204022159 | 1,29E-05 | 0,000138105 | Up |
| Gm13807 | 8,083969344 | -0,707717712 | 0,824576719 | 0,003187366 | 0,014835056 | Down |
| Cd82 | 3036,908916 | -2,164243609 | 0,201516895 | 6,24E-28 | 2,98E-25 | Down |
| Accs | 1531,17446 | 0,708980657 | 0,125514442 | 2,57E-10 | 9,61E-09 | Up |
| Thbs1 | 1984,565179 | 0,629808257 | 0,699410369 | 0,003953413 | 0,017676105 | Up |
| Bmf | 671,855034 | 0,604856738 | 0,165537495 | 4,30E-06 | 5,32E-05 | Up |
| Spint1 | 253,6176392 | 0,614017052 | 0,242153575 | 0,000175897 | 0,001294299 | Up |
| Chac1 | 4,766817882 | -1,776643717 | 1,687775816 | 0,000276119 | 0,001903967 | Down |
| Itpka | 124,4840104 | 0,755238307 | 0,187158937 | 5,61E-07 | 8,94E-06 | Up |
| Gm14002 | 33,59088708 | 0,632232473 | 0,351622406 | 0,000915265 | 0,005251121 | Up |
| Fbn1 | 28,01245812 | -2,132911125 | 0,832322655 | 2,11E-05 | 0,000212325 | Down |
| Atp8b4 | 4650,524735 | -0,620755877 | 0,122973195 | 7,10E-09 | 1,83E-07 | Down |
| Hdc | 51,68643992 | 0,585109637 | 0,410361503 | 0,00233533 | 0,011468138 | Up |
| Dusp2 | 2214,838787 | -0,694909664 | 0,134021217 | 3,23E-09 | 9,14E-08 | Down |
| Slc20a1 | 10384,98151 | -0,772598221 | 0,088355668 | 5,41E-20 | 1,11E-17 | Down |
| Il1b | 14,26609458 | -1,108377234 | 1,119179577 | 0,00105243 | 0,005906493 | Down |
| Ap5s1 | 565,1535008 | 1,000614172 | 0,162327936 | 3,22E-12 | 1,82E-10 | Up |
| Macrod2 | 256,2804725 | 1,267487628 | 0,228127299 | 4,62E-11 | 2,00E-09 | Up |
| Trib3 | 5837,963377 | -1,544278207 | 0,580219265 | 1,44E-05 | 0,000151266 | Down |
| Id1 | 13342,87747 | -1,505859876 | 0,138715845 | 9,34E-30 | 5,61E-27 | Down |
| Bcl2l1 | 12015,67863 | -0,633433622 | 0,106078871 | 3,81E-11 | 1,70E-09 | Down |
| Hck | 112,8469967 | -0,803642931 | 0,246907937 | 7,72E-06 | 8,83E-05 | Down |
| Nol4l | 6164,336407 | 0,92146895 | 0,115422175 | 1,44E-17 | 2,08E-15 | Up |
| Pxmp4 | 2668,970236 | 0,623230218 | 0,182109233 | 9,57E-06 | 0,000105875 | Up |
| Ggt7 | 101,5361539 | 0,705712473 | 0,406819522 | 0,000701118 | 0,004182221 | Up |
| Sla2 | 10507,01329 | 1,001525531 | 0,172594332 | 2,87E-11 | 1,33E-09 | Up |
| Mroh8 | 31,87457383 | 0,688954831 | 0,392817957 | 0,000733013 | 0,00434805 | Up |
| Tgm2 | 10821,87749 | 0,994088868 | 0,362466778 | 1,76E-05 | 0,000180209 | Up |
| Ppp1r16b | 161,7128126 | -1,167621403 | 0,263782407 | 1,96E-08 | 4,51E-07 | Down |
| Fitm2 | 2813,975891 | -0,596872084 | 0,090821167 | 8,09E-13 | 5,07E-11 | Down |
| Pabpc1l | 5,81403127 | 0,835589573 | 0,976120531 | 0,00221731 | 0,010974938 | Up |
| Wfdc12 | 53,19585331 | 1,228867992 | 0,425492095 | 7,29E-06 | 8,42E-05 | Up |
| Svs4 | 22,97847939 | 1,574702553 | 0,888849096 | 0,000136232 | 0,001042626 | Up |
| Slpi | 1213,529228 | -1,159776166 | 0,540660254 | 6,98E-05 | 0,000590564 | Down |
| Mmp9 | 25,13158665 | -0,903607566 | 0,84437316 | 0,001396961 | 0,007463185 | Down |
| Gm11469 | 10,86370356 | -2,788556377 | 1,188590227 | 2,17E-05 | 0,000216893 | Down |
| Gm14267 | 11,2510132 | -3,05340354 | 0,898574493 | 2,80E-06 | 3,64E-05 | Down |
| Gm11473 | 149,3113433 | -0,719098884 | 0,258561669 | 4,98E-05 | 0,000443625 | Down |

| | | | | | | |
|---|---|---|---|---|---|---|
| Gm14236 | 22,32851066 | 0,692037242 | 0,477357883 | 0,001295171 | 0,007032846 | Up |
| Atp9a | 443,3186064 | 1,965687291 | 0,192065363 | 8,46E-26 | 2,98E-23 | Up |
| Zfp831 | 679,8420958 | 1,202064362 | 0,211158282 | 2,17E-11 | 1,03E-09 | Up |
| Edn3 | 84,35801001 | 1,206585098 | 0,236420378 | 6,10E-10 | 2,08E-08 | Up |
| 4921531C22Rik | 144,5384733 | 0,788313246 | 0,185362016 | 2,04E-07 | 3,67E-06 | Up |
| Col9a3 | 253,833036 | 1,026157561 | 0,296379936 | 1,52E-06 | 2,13E-05 | Up |
| Slc17a9 | 2603,04439 | -0,660466338 | 0,157508409 | 3,88E-07 | 6,51E-06 | Down |
| Pde7a | 6154,783926 | -0,60955687 | 0,079410297 | 2,70E-16 | 3,11E-14 | Down |
| Cp | 259,6809462 | -2,352171974 | 0,807099901 | 9,98E-06 | 0,000109832 | Down |
| Fndc3b | 2971,696078 | -0,735299496 | 0,133994455 | 6,15E-10 | 2,09E-08 | Down |
| Gm2011 | 65,66743088 | 0,857984616 | 0,23113504 | 1,24E-06 | 1,79E-05 | Up |
| Slc7a11 | 5980,104493 | -0,787411349 | 0,234162208 | 5,82E-06 | 6,89E-05 | Down |
| 5031434O11Rik | 165,0372201 | 0,719872067 | 0,172564019 | 3,61E-07 | 6,09E-06 | Up |
| Mgst2 | 3909,610169 | 1,030209294 | 0,204937333 | 1,72E-09 | 5,16E-08 | Up |
| Maml3 | 4177,416488 | 0,870504714 | 0,117817726 | 2,06E-15 | 2,05E-13 | Up |
| Wwtr1 | 82,48549328 | -3,210742886 | 0,419398412 | 1,09E-15 | 1,19E-13 | Down |
| Rnf13 | 3586,596236 | -0,61855472 | 0,127606972 | 1,99E-08 | 4,58E-07 | Down |
| Serp1 | 26709,77499 | -0,582485837 | 0,201263447 | 7,17E-05 | 0,000603992 | Down |
| Siah2 | 5297,132062 | -0,617728673 | 0,092840282 | 4,68E-13 | 3,03E-11 | Down |
| Gpr171 | 2450,094039 | 0,877779807 | 0,225522708 | 5,78E-07 | 9,18E-06 | Up |
| P2ry14 | 6297,454053 | -0,974088848 | 0,204990226 | 9,05E-09 | 2,27E-07 | Down |
| Ctso | 607,02333 | 0,60155962 | 0,211700115 | 7,63E-05 | 0,000637035 | Up |
| Tlr2 | 2738,950673 | -0,60005103 | 0,135532258 | 1,59E-07 | 2,94E-06 | Down |
| Muc1 | 13,32333098 | 0,91846873 | 0,775224272 | 0,001086704 | 0,006069456 | Up |
| Dcst1 | 28,3747227 | 0,744007304 | 0,4837457 | 0,000877815 | 0,005087834 | Up |
| Selenbp2 | 107,7579436 | 0,845514841 | 0,33068965 | 5,13E-05 | 0,000454428 | Up |
| Selenbp1 | 1431,108593 | 0,954765623 | 0,271716594 | 1,65E-06 | 2,29E-05 | Up |
| Sema6c | 26,40754602 | 1,655880743 | 0,57120416 | 7,84E-06 | 8,95E-05 | Up |
| Ctss | 663,8275755 | -0,92913606 | 0,432426136 | 0,000111468 | 0,000877207 | Down |
| Fcgr1 | 5289,287716 | -0,894957815 | 0,150115189 | 2,32E-11 | 1,10E-09 | Down |
| Pias3 | 2678,101871 | 0,601560703 | 0,08585691 | 3,98E-14 | 3,29E-12 | Up |
| Pde4dip | 5539,840031 | -0,600578735 | 0,131639983 | 8,39E-08 | 1,66E-06 | Down |
| Gm5544 | 8,013562118 | -0,971674602 | 0,967443632 | 0,00135319 | 0,007267987 | Down |
| Gm5853 | 29,29279593 | 0,756302637 | 0,844846565 | 0,002640855 | 0,012710728 | Up |
| Slc16a1 | 34988,48471 | -0,688873247 | 0,109319583 | 5,10E-12 | 2,78E-10 | Down |
| Cd53 | 2647,867443 | -2,048619198 | 0,627666311 | 4,00E-06 | 4,99E-05 | Down |
| Prok1 | 7,228139486 | -2,118438646 | 1,452056437 | 0,00014006 | 0,001066886 | Down |
| Csf1 | 1974,845064 | -0,793095273 | 0,234400332 | 5,28E-06 | 6,35E-05 | Down |
| Gm42776 | 16,6969338 | -0,912056953 | 0,962056709 | 0,001664387 | 0,008641497 | Down |
| 5730508B09Rik | 697,5044354 | -0,587779262 | 0,14326594 | 7,16E-07 | 1,11E-05 | Down |
| Bank1 | 390,3070533 | -0,96675039 | 0,293928004 | 3,41E-06 | 4,35E-05 | Down |
| Dapp1 | 12758,11943 | 0,625849494 | 0,091566351 | 1,36E-13 | 9,85E-12 | Up |
| 9530052C20Rik | 67,58235858 | 0,649177209 | 0,294814032 | 0,000340736 | 0,00229016 | Up |
| Ddah1 | 12,46966775 | -0,63212935 | 0,581787547 | 0,003049954 | 0,01430217 | Down |
| Dnajb4 | 3052,467561 | 0,683267397 | 0,154900793 | 1,41E-07 | 2,64E-06 | Up |
| Lrriq3 | 41,48503189 | 0,628852939 | 0,301067746 | 0,000501359 | 0,003161361 | Up |
| Cth | 7,41930163 | -1,814296196 | 1,567602784 | 0,000265694 | 0,001841472 | Down |
| Tmem68 | 1812,880651 | -0,59570892 | 0,118630834 | 8,52E-09 | 2,16E-07 | Down |
| Gem | 254,5110708 | -4,829762261 | 1,570491711 | 7,79E-07 | 1,20E-05 | Down |
| Cdh17 | 137,6528549 | -5,471201184 | 0,990256762 | 3,65E-11 | 1,64E-09 | Down |
| Pdp1 | 2184,299773 | -1,455500957 | 0,236521179 | 1,97E-12 | 1,15E-10 | Down |

| | | | | | | |
|---|---|---|---|---|---|---|
| Osgin2 | 3112,211715 | -0,735125312 | 0,102814304 | 1,73E-14 | 1,50E-12 | Down |
| Bach2os | 16,43915049 | -2,358160464 | 0,955380771 | 2,40E-05 | 0,000236248 | Down |
| Cfap206 | 10,11146459 | 2,895863083 | 1,102639327 | 1,28E-05 | 0,000136913 | Up |
| AI464131 | 14,48568737 | -0,595431712 | 0,636714182 | 0,004344089 | 0,019137344 | Down |
| Cd72 | 402,1646575 | 1,172458736 | 0,164055467 | 1,40E-15 | 1,47E-13 | Up |
| Glipr2 | 6207,664248 | -1,058010649 | 0,242430317 | 3,62E-08 | 7,90E-07 | Down |
| Dnajc25 | 1972,880386 | -0,605915426 | 0,095471475 | 3,58E-12 | 2,00E-10 | Down |
| Tnfsf8 | 2093,575493 | -1,441764003 | 0,133038894 | 6,78E-30 | 4,23E-27 | Down |
| Tlr4 | 3138,593152 | -0,820772234 | 0,187045756 | 1,01E-07 | 1,96E-06 | Down |
| Elavl2 | 986,5023764 | -1,109984576 | 0,119105956 | 2,13E-23 | 6,06E-21 | Down |
| Nfia | 299,0884067 | 0,725685569 | 0,194692964 | 2,07E-06 | 2,81E-05 | Up |
| Lepr | 34,45425309 | -0,817746166 | 0,501961583 | 0,000551667 | 0,003411124 | Down |
| Pde4b | 1526,193179 | 0,936749521 | 0,131480798 | 8,57E-15 | 7,86E-13 | Up |
| Lrp8 | 9688,957132 | -0,70361587 | 0,083742468 | 9,60E-19 | 1,66E-16 | Down |
| Echdc2 | 151,1693368 | -0,812022598 | 0,208541879 | 7,75E-07 | 1,20E-05 | Down |
| Cdkn2c | 6893,065099 | 1,067393433 | 0,091803514 | 7,16E-34 | 7,26E-31 | Up |
| Mob3c | 849,361509 | -0,777033494 | 0,117439934 | 6,54E-13 | 4,18E-11 | Down |
| Eif2b3 | 7067,412782 | -0,601981988 | 0,104448074 | 1,34E-10 | 5,24E-09 | Down |
| Slc2a1 | 11722,19401 | -0,806222174 | 0,11978893 | 2,86E-13 | 1,97E-11 | Down |
| Hivep3 | 636,3806459 | -0,973702353 | 0,181111236 | 3,82E-10 | 1,38E-08 | Down |
| Yrdc | 6107,586385 | -0,663786431 | 0,102820266 | 1,84E-12 | 1,09E-10 | Down |
| Csf3r | 399,1508416 | -1,147934242 | 0,504536156 | 4,95E-05 | 0,0004417 | Down |
| Gjb3 | 19,38929696 | -1,556427452 | 0,684227959 | 4,06E-05 | 0,000372219 | Down |
| Tlr12 | 147,3951441 | 0,653378407 | 0,25875389 | 0,000144988 | 0,001098239 | Up |
| Zfp362 | 5602,70999 | 0,677833597 | 0,124326238 | 7,79E-10 | 2,56E-08 | Up |
| Ccdc28b | 668,8239146 | 0,741366394 | 0,116294594 | 3,24E-12 | 1,82E-10 | Up |
| Spocd1 | 103,4212202 | 0,864333523 | 0,372131228 | 8,89E-05 | 0,000725228 | Up |
| Cd52 | 1011,541979 | 1,158263746 | 0,244877158 | 4,64E-09 | 1,26E-07 | Up |
| Zfp593 | 158,6262685 | -0,636352202 | 0,226461634 | 6,99E-05 | 0,000590692 | Down |
| Extl1 | 477,9996045 | -0,85255049 | 0,602537212 | 0,000785912 | 0,004620082 | Down |
| Selenon | 8822,329824 | -0,999801242 | 0,08097804 | 2,51E-37 | 3,13E-34 | Down |
| Man1c1 | 2246,77068 | -2,600738141 | 0,20184059 | 6,76E-39 | 1,10E-35 | Down |
| Clic4 | 37244,65662 | -0,613065992 | 0,168450441 | 4,40E-06 | 5,42E-05 | Down |
| Cnr2 | 639,2081342 | -1,007831347 | 0,252675931 | 2,20E-07 | 3,92E-06 | Down |
| Gm13012 | 1059,186717 | 0,635418129 | 0,14730749 | 2,43E-07 | 4,28E-06 | Up |
| Sh2d5 | 9049,580851 | 0,580264102 | 0,20819099 | 0,000101759 | 0,000814126 | Up |
| Kif17 | 1815,628705 | 0,682933729 | 0,336047886 | 0,000417852 | 0,002724941 | Up |
| Pla2g2e | 33,9934475 | -1,816113566 | 0,93649954 | 8,43E-05 | 0,000693236 | Down |
| Padi6 | 31,51235846 | 0,92630617 | 0,495119817 | 0,000225489 | 0,001599721 | Up |
| Crocc | 1849,853189 | 0,635038697 | 0,125696767 | 6,82E-09 | 1,77E-07 | Up |
| B330016D10Rik | 305,906685 | 0,669005677 | 0,127033547 | 2,14E-09 | 6,27E-08 | Up |
| Kazn | 3608,778112 | 0,666216037 | 0,136280754 | 1,51E-08 | 3,61E-07 | Up |
| Dhrs3 | 6629,037815 | 0,581204548 | 0,297219964 | 0,000907909 | 0,005216531 | Up |
| Zfp986 | 19,52119009 | -0,812469115 | 1,031348528 | 0,002448385 | 0,011929443 | Down |
| Casz1 | 137,4757495 | -0,719587983 | 0,304128424 | 0,000154086 | 0,001158144 | Down |
| Slc25a33 | 1909,241325 | -1,085111103 | 0,102720255 | 9,16E-29 | 4,64E-26 | Down |
| Gm13091 | 135,4061648 | 1,641641185 | 0,38342211 | 6,72E-08 | 1,36E-06 | Up |
| Plekhg5 | 3224,120707 | 0,672477172 | 0,14535233 | 5,33E-08 | 1,11E-06 | Up |
| Abcb1b | 4149,851263 | -1,022250828 | 0,114758808 | 2,06E-21 | 5,06E-19 | Down |
| Crot | 3100,332009 | 0,614524283 | 0,124078694 | 1,17E-08 | 2,87E-07 | Up |
| Napepld | 977,2137184 | 0,673284154 | 0,090024413 | 1,41E-15 | 1,47E-13 | Up |

| | | | | | | |
|---|---|---|---|---|---|---|
| Kcnh2 | 15,80891239 | -1,741423186 | 0,95147365 | 0,00010981 | 0,000867413 | Down |
| Nos3 | 60,47437923 | -2,219266802 | 0,272401646 | 3,04E-17 | 4,08E-15 | Down |
| Khk | 614,7936558 | 0,848114094 | 0,136816968 | 7,26E-12 | 3,77E-10 | Up |
| AI839979 | 54,48895441 | -0,78635527 | 0,56013117 | 0,000982802 | 0,00557551 | Down |
| Plb1 | 16,8012613 | -0,634509173 | 0,548805073 | 0,002724719 | 0,013014 | Down |
| Gm15614 | 21,15205568 | 0,767791445 | 0,58516845 | 0,001252209 | 0,006833868 | Up |
| Mxd4 | 4469,061136 | 0,600059131 | 0,394889857 | 0,001840442 | 0,00936968 | Up |
| Acox3 | 2910,678591 | 0,625265704 | 0,233779936 | 0,000111054 | 0,000875532 | Up |
| Sh3tc1 | 912,8728443 | -0,807375882 | 0,16135032 | 6,28E-09 | 1,64E-07 | Down |
| Ppp2r2c | 9,036617986 | 2,403519364 | 1,621538501 | 9,22E-05 | 0,00074804 | Up |
| Bst1 | 33,67837635 | -5,481207718 | 1,53383925 | 1,47E-07 | 2,74E-06 | Down |
| Apbb2 | 43,3166444 | 1,918153346 | 0,523858083 | 1,12E-06 | 1,64E-05 | Up |
| Ppbp | 98,37811165 | -2,003772793 | 1,24856529 | 0,000134807 | 0,001033182 | Down |
| Pf4 | 352,6255095 | -0,896814976 | 0,398082881 | 9,46E-05 | 0,000765251 | Down |
| Cxcl3 | 14,84452677 | -2,360473437 | 1,768647318 | 0,000103567 | 0,000828179 | Down |
| Cxcl2 | 57,55468698 | -3,962523399 | 0,68011576 | 5,73E-11 | 2,43E-09 | Down |
| Mthfd2l | 1151,184745 | -0,748416243 | 0,100555164 | 2,06E-15 | 2,05E-13 | Down |
| Epgn | 8,228003661 | -0,74580198 | 0,772368446 | 0,002469023 | 0,012008363 | Down |
| Cxcl10 | 6,839317694 | -0,984365868 | 1,044749348 | 0,00142883 | 0,007595513 | Down |
| Ccng2 | 1329,31746 | 0,706681877 | 0,228847725 | 2,07E-05 | 0,000208918 | Up |
| Anxa3 | 8980,662929 | -0,67276977 | 0,290694308 | 0,000226018 | 0,001602775 | Down |
| Agpat9 | 20,36580129 | -1,058963361 | 0,88049217 | 0,00080848 | 0,00473947 | Down |
| Spp1 | 14166,01247 | -1,494270079 | 0,214066788 | 9,87E-15 | 8,90E-13 | Down |
| Tgfbr3 | 7,699380303 | -0,81573246 | 1,010112858 | 0,002412899 | 0,011799061 | Down |
| Gm42517 | 916,9586079 | 1,125125772 | 0,146662875 | 3,15E-17 | 4,18E-15 | Up |
| Gm10419 | 2401,713468 | 0,991067138 | 0,106335982 | 6,31E-23 | 1,71E-20 | Up |
| Gm18555 | 102,3025521 | 0,752228786 | 0,310871676 | 0,000113691 | 0,000891728 | Up |
| Ulk1 | 5417,08254 | 0,755611961 | 0,104149861 | 8,14E-15 | 7,59E-13 | Up |
| Hscb | 2015,219503 | 0,625729495 | 0,083040844 | 8,24E-16 | 9,03E-14 | Up |
| Ttc28 | 1354,800972 | 0,702700983 | 0,162510591 | 1,97E-07 | 3,57E-06 | Up |
| Ssh1 | 468,8903113 | 1,424874962 | 0,251245308 | 3,22E-11 | 1,47E-09 | Up |
| 4930515G01Rik | 137,6154348 | 0,763463416 | 0,183652741 | 3,32E-07 | 5,67E-06 | Up |
| Tesc | 31,63473976 | -0,734457928 | 0,497179382 | 0,001027833 | 0,005780448 | Down |
| Sdsl | 2168,067081 | 0,59422355 | 0,106369002 | 3,82E-10 | 1,38E-08 | Up |
| 2510016D11Rik | 486,9729743 | 0,668696073 | 0,140399972 | 2,79E-08 | 6,22E-07 | Up |
| Oas2 | 153,550936 | -3,288039028 | 0,328127296 | 6,90E-25 | 2,20E-22 | Down |
| Oas3 | 341,8636315 | -2,396834037 | 0,461579645 | 7,70E-09 | 1,97E-07 | Down |
| Acad12 | 811,1184419 | 0,85004271 | 0,141506819 | 2,29E-11 | 1,09E-09 | Up |
| P2rx7 | 32,61155604 | -0,803917448 | 0,736185683 | 0,001703081 | 0,008788962 | Down |
| Tmem120b | 909,7917846 | -0,764433559 | 0,123845793 | 1,12E-11 | 5,54E-10 | Down |
| Hcar2 | 29,20565238 | -3,088733834 | 0,627640167 | 2,65E-08 | 5,93E-07 | Down |
| Stx2 | 1745,612562 | 0,583628662 | 0,064671158 | 2,88E-21 | 6,98E-19 | Up |
| Clip2 | 3968,004608 | 0,828187905 | 0,079758145 | 6,90E-27 | 2,80E-24 | Up |
| Rfc2 | 25282,33014 | 0,980905354 | 0,067934227 | 1,82E-49 | 7,37E-46 | Up |
| Lat2 | 4766,608057 | 0,889884313 | 0,151403486 | 3,93E-11 | 1,75E-09 | Up |
| Cux1 | 58335,39229 | 0,965069326 | 0,123010808 | 2,92E-17 | 3,94E-15 | Up |
| Myl10 | 156,465505 | 3,022451972 | 0,424584192 | 7,61E-14 | 5,96E-12 | Up |
| 9130604C24Rik | 380,1543433 | 0,639928692 | 0,12526673 | 5,05E-09 | 1,36E-07 | Up |
| Actl6b | 4,255639881 | 0,670201036 | 0,962239383 | 0,003880731 | 0,017427306 | Up |
| Tfr2 | 6268,32924 | 1,576970822 | 0,20684195 | 1,48E-16 | 1,80E-14 | Up |
| Fbxo24 | 42,22236255 | 0,693028878 | 0,378554136 | 0,000612904 | 0,003740608 | Up |

| | | | | | | |
|---|---|---|---|---|---|---|
| Gm36266 | 28,60635371 | 0,596146582 | 0,636018852 | 0,004320128 | 0,019057659 | Up |
| Zcwpw1 | 1245,961138 | 0,631512355 | 0,155355058 | 7,29E-07 | 1,13E-05 | Up |
| Pilra | 72,16270277 | -1,72941361 | 0,422365324 | 1,76E-07 | 3,22E-06 | Down |
| Pilrb1 | 241,8391833 | -2,046304538 | 0,287395928 | 5,03E-14 | 4,08E-12 | Down |
| Gm36551 | 78,80995591 | -1,614294963 | 0,327625183 | 3,45E-09 | 9,66E-08 | Down |
| Prkar1b | 85,97825731 | 1,063870533 | 0,227444954 | 8,26E-09 | 2,10E-07 | Up |
| Micall2 | 18999,42081 | 0,782783383 | 0,098601193 | 4,42E-17 | 5,69E-15 | Up |
| Lfng | 13135,06574 | -0,589731468 | 0,080815886 | 4,76E-15 | 4,57E-13 | Down |
| 8430423G03Rik | 88,71111414 | -0,737751768 | 0,259394808 | 3,77E-05 | 0,00034851 | Down |
| Asns | 30785,17086 | -0,649872303 | 0,163280854 | 9,85E-07 | 1,48E-05 | Down |
| 2610001J05Rik | 9642,694611 | -0,9344585 | 0,109856009 | 1,67E-19 | 3,23E-17 | Down |
| Met | 4940,381982 | -0,606890814 | 0,162460951 | 3,09E-06 | 3,98E-05 | Down |
| Wnt2 | 10,91640668 | 1,247380957 | 0,899954999 | 0,000423039 | 0,002749922 | Up |
| Asz1 | 351,0637868 | 1,738843074 | 0,366353084 | 1,23E-08 | 3,00E-07 | Up |
| Podxl | 4629,984652 | -0,627847118 | 0,097565021 | 2,03E-12 | 1,18E-10 | Down |
| Akr1b7 | 8,54091321 | -0,792253021 | 1,173336193 | 0,002365375 | 0,011602583 | Down |
| 1810058I24Rik | 8858,372937 | 0,615635121 | 0,09109306 | 2,29E-13 | 1,60E-11 | Up |
| Kdm7a | 753,3487724 | -0,6410025 | 0,118474373 | 9,93E-10 | 3,16E-08 | Down |
| Slc37a3 | 345,3545136 | -1,481788776 | 0,385216683 | 2,69E-07 | 4,69E-06 | Down |
| Clec5a | 2770,782305 | -1,735628879 | 0,363510747 | 1,07E-08 | 2,64E-07 | Down |
| Mgam | 41,70671067 | -1,998396879 | 0,526596106 | 8,32E-07 | 1,27E-05 | Down |
| Gm43889 | 34,29366836 | -1,853991644 | 0,936880446 | 7,64E-05 | 0,000637035 | Down |
| Pdia4 | 29604,59528 | -0,707148652 | 0,141335242 | 8,07E-09 | 2,06E-07 | Down |
| Gimap6 | 107,6610722 | 0,926615605 | 0,325969037 | 1,64E-05 | 0,000168836 | Up |
| Gimap5 | 1384,344771 | 0,658566052 | 0,167023671 | 1,12E-06 | 1,64E-05 | Up |
| Ccdc129 | 6,049398883 | 1,539473848 | 1,72273622 | 0,000382522 | 0,002532199 | Up |
| Tnip3 | 175,1734794 | -0,660128539 | 0,306428357 | 0,000358452 | 0,002395342 | Down |
| Gadd45a | 450,8678506 | -0,961941948 | 0,375913218 | 3,32E-05 | 0,000312889 | Down |
| Hk2 | 19649,01962 | -1,023828177 | 0,10177309 | 3,19E-26 | 1,18E-23 | Down |
| Gm17034 | 20,10568958 | 1,204677163 | 0,733831728 | 0,000244177 | 0,001714308 | Up |
| Gm43878 | 55,94813608 | -0,822190941 | 0,423206693 | 0,000267587 | 0,001852592 | Down |
| Mthfd2 | 15492,30892 | -0,766562952 | 0,113123149 | 2,29E-13 | 1,60E-11 | Down |
| Rab11fip5 | 155,2319165 | -2,579486649 | 0,40085003 | 9,54E-12 | 4,83E-10 | Down |
| Gm44198 | 184,4808107 | -0,593415495 | 0,158053391 | 3,02E-06 | 3,90E-05 | Down |
| Gm43904 | 7,527783735 | 0,936999116 | 1,191745926 | 0,001696116 | 0,008761378 | Up |
| Gm26588 | 43,18107596 | -0,61148955 | 0,305998569 | 0,000682711 | 0,004091979 | Down |
| Abtb1 | 2440,507243 | 0,695919219 | 0,160422917 | 1,89E-07 | 3,44E-06 | Up |
| Aldh1l1 | 4,678236003 | -0,867582957 | 1,214094343 | 0,001941467 | 0,009816237 | Down |
| Slc41a3 | 349,8240344 | 0,614300711 | 0,227663898 | 0,000110056 | 0,000868935 | Up |
| Frmd4b | 176,0736522 | 1,947872866 | 0,324311945 | 4,35E-11 | 1,91E-09 | Up |
| Bhlhe40 | 21345,23936 | -0,743641647 | 0,127744612 | 9,33E-11 | 3,78E-09 | Down |
| Srgap3 | 2102,686749 | 0,993593941 | 0,09728711 | 9,21E-27 | 3,64E-24 | Up |
| Il17re | 269,1383942 | 1,043508769 | 0,276857762 | 4,57E-07 | 7,51E-06 | Up |
| Mkrn2os | 29,64026423 | 1,00168345 | 0,423600559 | 5,11E-05 | 0,000452922 | Up |
| Cand2 | 1623,639553 | 1,792999169 | 0,233583711 | 3,47E-16 | 3,94E-14 | Up |
| Gm8213 | 184,2105679 | 1,110555744 | 0,212953932 | 4,34E-10 | 1,54E-08 | Up |
| Cacna2d4 | 144,6023226 | 1,037459606 | 0,225515946 | 1,34E-08 | 3,25E-07 | Up |
| Tuba8 | 21444,23503 | -0,610250152 | 0,168746941 | 4,86E-06 | 5,90E-05 | Down |
| Usp18 | 164,2414255 | -1,51110539 | 0,234614011 | 4,13E-13 | 2,72E-11 | Down |
| Phc1 | 2601,639079 | 0,869731655 | 0,104475629 | 1,32E-18 | 2,22E-16 | Up |
| Slc2a3 | 28352,06815 | -0,850656606 | 0,12714232 | 3,09E-13 | 2,11E-11 | Down |

| | | | | | | |
|---|---|---|---|---|---|---|
| Clec4n | 64,08773858 | -1,270624298 | 1,091133005 | 0,000631365 | 0,003842423 | Down |
| Clec4e | 2450,530282 | -0,838500723 | 0,225286532 | 1,31E-06 | 1,87E-05 | Down |
| C1ra | 714,1074443 | -0,663298396 | 0,276135113 | 0,000190438 | 0,001388073 | Down |
| Ptpn6 | 45882,45776 | 0,643121175 | 0,184164698 | 6,81E-06 | 7,91E-05 | Up |
| Grcc10 | 710,8403106 | 0,788798454 | 0,172340369 | 4,99E-08 | 1,04E-06 | Up |
| Lag3 | 151,5211733 | 1,150375746 | 0,225655093 | 7,11E-10 | 2,37E-08 | Up |
| Ptms | 5382,522805 | 0,688212798 | 0,12702455 | 9,35E-10 | 3,01E-08 | Up |
| Fgf23 | 169,6018244 | -3,199076527 | 0,546141773 | 2,06E-10 | 7,77E-09 | Down |
| Tigar | 2775,434518 | -0,688344534 | 0,13813019 | 9,14E-09 | 2,29E-07 | Down |
| Tulp3 | 484,9721266 | 1,149451804 | 0,93791757 | 0,00067229 | 0,004042959 | Up |
| Gm26770 | 159,7417486 | 1,138904489 | 0,225396264 | 9,35E-10 | 3,01E-08 | Up |
| Rhno1 | 8719,928351 | 1,061708719 | 0,108044438 | 2,29E-25 | 7,74E-23 | Up |
| Foxm1 | 28001,67483 | 0,683423017 | 0,082397286 | 2,24E-18 | 3,57E-16 | Up |
| Gm43965 | 93,62503784 | 0,893631733 | 0,338111058 | 3,32E-05 | 0,000312949 | Up |
| Cd69 | 4378,454413 | -2,221133302 | 0,289902084 | 1,39E-15 | 1,47E-13 | Down |
| Clec12a | 1081,153479 | -0,820257545 | 0,782192629 | 0,001762773 | 0,009025241 | Down |
| Klra2 | 7,263873726 | -0,862662842 | 1,307299994 | 0,001705431 | 0,008798291 | Down |
| Bcl2l14 | 2146,088438 | -1,036224267 | 0,147305536 | 7,10E-15 | 6,65E-13 | Down |
| Fam60a | 3785,475364 | -0,720051248 | 0,081087802 | 1,58E-20 | 3,62E-18 | Down |
| Dennd5b | 69,37463206 | -0,626943966 | 0,423744166 | 0,001691904 | 0,008745188 | Down |
| Gm44160 | 2666,084895 | -0,686554808 | 0,24179918 | 4,94E-05 | 0,000440751 | Down |
| Tarm1 | 1208,628513 | -1,11213753 | 0,280346966 | 1,68E-07 | 3,10E-06 | Down |
| Oscar | 78,10024709 | -0,793220548 | 0,298999035 | 4,98E-05 | 0,000443625 | Down |
| Rdh13 | 4023,991193 | -0,587025352 | 0,138170428 | 3,76E-07 | 6,33E-06 | Down |
| Zfp580 | 1035,542264 | 0,678327265 | 0,124652047 | 8,24E-10 | 2,69E-08 | Up |
| 1810019N24Rik | 33,00560074 | 0,586524893 | 0,461473771 | 0,002912651 | 0,013769743 | Up |
| Lig1 | 46515,767 | 0,584759833 | 0,060193352 | 4,16E-24 | 1,23E-21 | Up |
| Pla2g4c | 248,4843085 | 1,691489208 | 0,371818117 | 2,51E-08 | 5,66E-07 | Up |
| Ehd2 | 210,8835274 | -1,684286571 | 0,338843374 | 3,61E-09 | 1,00E-07 | Down |
| C5ar1 | 80,99254291 | -2,839462286 | 0,358161058 | 2,01E-16 | 2,35E-14 | Down |
| Slc1a5 | 12790,89028 | -0,701845782 | 0,086301826 | 8,96E-18 | 1,33E-15 | Down |
| Gng8 | 26,98188007 | -0,658445889 | 0,577422092 | 0,002524733 | 0,012220702 | Down |
| Ptgir | 1602,251641 | -1,296294902 | 0,223601807 | 1,12E-11 | 5,54E-10 | Down |
| Psg17 | 14,23180398 | 3,026801373 | 1,604495728 | 2,91E-05 | 0,000279879 | Up |
| Mill2 | 26,68541156 | 1,676397437 | 0,7791933 | 5,46E-05 | 0,000479498 | Up |
| Ccdc61 | 2143,291749 | 0,61722786 | 0,113159334 | 7,89E-10 | 2,59E-08 | Up |
| Dmwd | 5860,363685 | -0,772462998 | 0,083022382 | 3,43E-22 | 8,83E-20 | Down |
| Relb | 527,0632685 | -1,445896351 | 0,236347374 | 2,37E-12 | 1,37E-10 | Down |
| Pvr | 729,3870502 | -1,063736254 | 0,182255172 | 1,57E-11 | 7,64E-10 | Down |
| Plaur | 2252,858594 | -1,280295119 | 0,163327794 | 6,08E-18 | 9,21E-16 | Down |
| Cd177 | 3314,386718 | -0,746974837 | 0,211961552 | 3,98E-06 | 4,98E-05 | Down |
| Numbl | 178,9177678 | 0,873168577 | 0,373124273 | 8,21E-05 | 0,000678598 | Up |
| Ltbp4 | 28,60761441 | 1,289810596 | 0,528804036 | 2,73E-05 | 0,000264836 | Up |
| Gm44685 | 125,6007902 | 0,958458695 | 0,264967363 | 1,12E-06 | 1,64E-05 | Up |
| 9530053A07Rik | 88,21856048 | -1,950933118 | 0,455521998 | 1,45E-07 | 2,71E-06 | Down |
| Gm45010 | 81,40064451 | 0,666662636 | 0,21829758 | 2,80E-05 | 0,000270294 | Up |
| Zfp36 | 1356,414366 | -1,133252881 | 0,19889364 | 2,55E-11 | 1,20E-09 | Down |
| Rasgrp4 | 12,14188763 | -0,703111364 | 0,763769497 | 0,002996116 | 0,014105421 | Down |
| Proser3 | 945,1244147 | 0,710286975 | 0,167318116 | 2,72E-07 | 4,73E-06 | Up |
| Ffar2 | 2584,762228 | 0,776570425 | 0,532682441 | 0,00091157 | 0,005231773 | Up |
| Mag | 736,0866341 | -0,627920904 | 0,150185646 | 4,47E-07 | 7,37E-06 | Down |

| | | | | | | |
|---|---|---|---|---|---|---|
| Rhpn2 | 178,5529272 | -1,696244675 | 0,926083441 | 0,000112051 | 0,000880826 | Down |
| Siglecf | 2157,366372 | 2,976242543 | 0,312603857 | 1,50E-22 | 3,99E-20 | Up |
| Klk1b11 | 4,513679802 | -1,044695388 | 1,4452997 | 0,001063892 | 0,005960516 | Down |
| Kcnc3 | 605,3869334 | -0,681937265 | 0,279497808 | 0,000155665 | 0,001164434 | Down |
| Gm44832 | 38,63610838 | -0,770268661 | 0,410982447 | 0,000387051 | 0,002553843 | Down |
| Fut1 | 6,061453696 | -0,987010863 | 1,2282032 | 0,001485853 | 0,007847644 | Down |
| Kcnj14 | 93,30567782 | -0,721014124 | 0,352804569 | 0,000333248 | 0,002244478 | Down |
| Syngr4 | 51,83139779 | 0,675964749 | 0,272637346 | 0,000144863 | 0,001098072 | Up |
| Saa4 | 11,52120595 | -5,413260484 | 1,35568692 | 4,72E-08 | 9,98E-07 | Down |
| Tjp1 | 79,21729523 | 0,594208902 | 0,483935878 | 0,002972822 | 0,014013375 | Up |
| 1810008I18Rik | 31,5009811 | -3,105597469 | 0,547790685 | 6,66E-10 | 2,24E-08 | Down |
| Selenos | 9797,488967 | -0,817616136 | 0,168457663 | 1,25E-08 | 3,03E-07 | Down |
| Aldh1a3 | 9,407886034 | -0,619835179 | 0,668857534 | 0,003977707 | 0,017769356 | Down |
| Akap13 | 34518,55022 | 1,029679625 | 0,128280151 | 3,72E-18 | 5,81E-16 | Up |
| Isg20 | 612,3464215 | -0,590372744 | 0,466428089 | 0,002874425 | 0,01362874 | Down |
| Gm15880 | 241,0916252 | 0,638557234 | 0,278299656 | 0,000288748 | 0,001978437 | Up |
| Zscan2 | 576,059414 | 0,7471578 | 0,116538082 | 2,56E-12 | 1,48E-10 | Up |
| Stard5 | 2301,646709 | -0,59967292 | 0,077610852 | 1,80E-16 | 2,14E-14 | Down |
| Abhd17c | 18929,1166 | -1,552940772 | 0,124908234 | 1,41E-37 | 1,91E-34 | Down |
| Arnt2 | 849,7298932 | -2,797219517 | 0,363857631 | 1,36E-15 | 1,45E-13 | Down |
| Gm44556 | 10,50485091 | -0,847402624 | 1,008615188 | 0,002173259 | 0,01078322 | Down |
| Rab38 | 29661,05411 | 0,776843645 | 0,091768385 | 5,95E-19 | 1,07E-16 | Up |
| Kctd14 | 149,1826973 | -0,944560887 | 0,277806228 | 2,55E-06 | 3,36E-05 | Down |
| Aamdc | 625,5213932 | 0,674672044 | 0,140968975 | 2,47E-08 | 5,60E-07 | Up |
| Aqp11 | 254,6430333 | 0,614127483 | 0,146936777 | 4,68E-07 | 7,65E-06 | Up |
| Lrrc32 | 9,104524139 | -0,759589048 | 0,775393871 | 0,002323102 | 0,011425382 | Down |
| Dgat2 | 116,9439592 | 0,613194938 | 0,381380888 | 0,001464041 | 0,007752632 | Up |
| Slco2b1 | 117,650939 | 0,645480754 | 0,371770068 | 0,00095799 | 0,00545288 | Up |
| Usp17ld | 816,2852816 | -3,453044494 | 0,19599971 | 7,94E-71 | 1,29E-66 | Down |
| Usp17lc | 25,65547678 | -4,082741419 | 0,774179726 | 9,67E-10 | 3,09E-08 | Down |
| Hbb-bs | 200,9586584 | 1,051143499 | 1,312772517 | 0,001233315 | 0,006751193 | Up |
| Hbb-y | 20,29731936 | 2,336458198 | 1,702728947 | 0,000106976 | 0,000848335 | Up |
| Gm4070 | 372,7722627 | -1,898029961 | 0,236072219 | 3,24E-17 | 4,25E-15 | Down |
| 4930431P19Rik | 74,04228373 | 0,709122405 | 0,281542926 | 0,000109194 | 0,000863391 | Up |
| Sbf2 | 14,00933974 | 0,895859734 | 0,801290332 | 0,001300174 | 0,007050577 | Up |
| Cdr2 | 4249,818112 | -0,669934629 | 0,076987523 | 6,59E-20 | 1,34E-17 | Down |
| Gm20274 | 275,0519533 | -0,646187065 | 0,239803198 | 9,35E-05 | 0,00075756 | Down |
| Sbk1 | 2234,883186 | 0,644070054 | 0,109027294 | 5,63E-11 | 2,40E-09 | Up |
| Apobr | 3806,992693 | -0,629448721 | 0,114984048 | 7,00E-10 | 2,34E-08 | Down |
| Nupr1 | 765,1048752 | -1,315798456 | 0,737140944 | 0,000154608 | 0,001160815 | Down |
| Ypel3 | 612,7755544 | 0,782690735 | 0,426188814 | 0,000399388 | 0,002623513 | Up |
| Mylpf | 586,579511 | 0,716013284 | 0,135718604 | 1,96E-09 | 5,77E-08 | Up |
| Gm44728 | 15,19890685 | 0,964033397 | 0,655789427 | 0,000532412 | 0,003310996 | Up |
| Itgam | 2870,78632 | -1,266398341 | 0,236095122 | 1,39E-10 | 5,39E-09 | Down |
| Itgax | 157,6541338 | 1,468839442 | 0,393453912 | 4,08E-07 | 6,79E-06 | Up |
| Cox6a2 | 153,6528533 | -2,432575895 | 0,722710551 | 3,70E-06 | 4,68E-05 | Down |
| Plekha1 | 5,765030263 | -0,615437537 | 1,103556484 | 0,003586973 | 0,016352525 | Down |
| Lrrc27 | 225,731834 | -0,6331052 | 0,185669391 | 9,60E-06 | 0,000106149 | Down |
| Adam8 | 724,7217499 | -2,318532849 | 0,415022425 | 1,01E-09 | 3,21E-08 | Down |
| Ifitm1 | 142241,0703 | -0,796189711 | 0,169742166 | 2,90E-08 | 6,41E-07 | Down |
| Gm7514 | 58,25403277 | -2,350085154 | 0,367704428 | 1,08E-11 | 5,38E-10 | Down |

| | | | | | | |
|---|---|---|---|---|---|---|
| Ifitm3 | 93618,87326 | -1,891937545 | 0,279894698 | 3,47E-13 | 2,34E-11 | Down |
| Ifitm6 | 396,4509885 | -4,072040991 | 0,711953831 | 8,43E-11 | 3,44E-09 | Down |
| Lrrc56 | 288,3536262 | 1,014960989 | 0,170994173 | 1,18E-11 | 5,84E-10 | Up |
| Irf7 | 224,653155 | -0,910240008 | 0,268068272 | 2,99E-06 | 3,87E-05 | Down |
| Muc6 | 10,9192414 | 1,011030442 | 0,838672323 | 0,000871796 | 0,005060781 | Up |
| Dusp8 | 13,96681384 | -3,68540273 | 0,907059639 | 2,92E-07 | 5,06E-06 | Down |
| Osbpl5 | 159,6143773 | 0,884003696 | 0,465951877 | 0,00023986 | 0,001688384 | Up |
| Cttn | 155,5479725 | 1,718752286 | 0,385520869 | 4,01E-08 | 8,66E-07 | Up |
| A430078G23Rik | 2138,522 | 0,659466269 | 0,126931565 | 3,14E-09 | 8,93E-08 | Up |
| Arhgef18 | 7935,683872 | 0,680198166 | 0,101567529 | 3,80E-13 | 2,53E-11 | Up |
| Retn | 803,0730145 | -0,653755009 | 0,280867906 | 0,000243317 | 0,001710303 | Down |
| Irs2 | 33,97500087 | -0,72903868 | 0,495941096 | 0,001065337 | 0,005966548 | Down |
| Col4a2 | 1922,424638 | -1,706286237 | 0,173365124 | 1,43E-24 | 4,29E-22 | Down |
| Mcf2l | 271,8840191 | -1,340535663 | 0,884199852 | 0,000287625 | 0,001972411 | Down |
| Adprhl1 | 11,71580451 | 2,497255992 | 1,135399479 | 3,49E-05 | 0,000326748 | Up |
| Rbpms | 4998,03979 | 0,676526427 | 0,172162622 | 1,11E-06 | 1,64E-05 | Up |
| Nwd1 | 3134,380359 | 1,109958616 | 0,343730459 | 2,75E-06 | 3,59E-05 | Up |
| F2rl3 | 623,7087597 | -0,699530043 | 0,177965842 | 1,03E-06 | 1,54E-05 | Down |
| Gm4890 | 153,8798741 | 0,586040506 | 0,160342152 | 4,63E-06 | 5,68E-05 | Up |
| Smad1 | 116,2128646 | 1,091367158 | 0,328267489 | 2,06E-06 | 2,80E-05 | Up |
| Il15 | 77,27478044 | 1,024724623 | 0,468860643 | 7,87E-05 | 0,000655439 | Up |
| Gm10644 | 47,0296698 | 1,325137411 | 0,359899274 | 4,33E-07 | 7,17E-06 | Up |
| Zswim4 | 2740,83809 | 0,643201409 | 0,126364174 | 5,55E-09 | 1,47E-07 | Up |
| Ier2 | 10364,82877 | -0,619875175 | 0,108124211 | 1,59E-10 | 6,04E-09 | Down |
| G430095P16Rik | 558,6620452 | 0,77797326 | 0,107888415 | 1,09E-14 | 9,70E-13 | Up |
| Klf1 | 812,5738791 | 0,940493256 | 0,125888407 | 6,54E-16 | 7,32E-14 | Up |
| Junb | 6396,057393 | -0,695007255 | 0,104333646 | 4,92E-13 | 3,17E-11 | Down |
| Gm42031 | 152,1581822 | 0,715442398 | 0,276911833 | 8,86E-05 | 0,000723486 | Up |
| Gpt2 | 11447,41538 | -1,056824899 | 0,196120594 | 2,16E-10 | 8,13E-09 | Down |
| Gm45425 | 35,99974872 | -0,67777914 | 0,409583179 | 0,000943802 | 0,005391968 | Down |
| Nod2 | 319,3092233 | 0,737140248 | 0,148013063 | 8,60E-09 | 2,17E-07 | Up |
| Bbs2 | 490,3657757 | 0,718432897 | 0,113310063 | 4,05E-12 | 2,23E-10 | Up |
| Drc7 | 87,03787323 | 2,936656999 | 0,410513064 | 6,48E-14 | 5,16E-12 | Up |
| Kifc3 | 1942,482142 | 1,142352721 | 0,212085058 | 1,50E-10 | 5,74E-09 | Up |
| Lrrc29 | 453,0647245 | 0,688749445 | 0,095355098 | 9,68E-15 | 8,78E-13 | Up |
| Lrrc36 | 24,50060593 | 1,113552193 | 0,475077585 | 4,30E-05 | 0,000390898 | Up |
| Gm38250 | 55,17544284 | 0,665726166 | 0,301639289 | 0,000306194 | 0,002081272 | Up |
| Fam65a | 21843,04541 | 1,019811642 | 0,15706801 | 3,38E-13 | 2,29E-11 | Up |
| Carmil2 | 2179,795262 | 0,785301543 | 0,155358002 | 5,26E-09 | 1,41E-07 | Up |
| Smpd3 | 56,30965067 | 0,774362989 | 0,396884045 | 0,000319715 | 0,002162307 | Up |
| Tldc1 | 1064,837869 | 0,6021897 | 0,109312522 | 5,90E-10 | 2,02E-08 | Up |
| Cotl1 | 639,3924189 | -0,668826568 | 0,242832036 | 6,97E-05 | 0,000589582 | Down |
| Gse1 | 29886,72165 | 0,763248691 | 0,113893713 | 3,80E-13 | 2,53E-11 | Up |
| 1700016A09Rik | 37,75277305 | 2,062516963 | 0,384201862 | 1,82E-09 | 5,43E-08 | Up |
| Irf8 | 26,07030485 | 0,970531165 | 0,775538064 | 0,000852954 | 0,004969184 | Up |
| RP24-365B9,2 | 30,38660638 | -0,718225192 | 0,732166391 | 0,00262132 | 0,012620451 | Down |
| Slc7a5 | 89433,40736 | -0,715304876 | 0,120159471 | 4,38E-11 | 1,92E-09 | Down |
| Gm27011 | 34,78370778 | 1,565941549 | 0,437192379 | 8,22E-07 | 1,26E-05 | Up |
| Gm45353 | 43,3470747 | 1,60063898 | 0,401474258 | 1,97E-07 | 3,57E-06 | Up |
| RP23-115O21,3 | 359,7888741 | 3,163194816 | 2,850419621 | 1,61E-05 | 0,000166175 | Up |
| RP23-312B17,1 | 6,252076908 | -0,600480429 | 0,799225326 | 0,005051098 | 0,021686709 | Down |

| | | | | | | |
|---|---|---|---|---|---|---|
| RP23-312B17,3 | 98,75736627 | 0,698926985 | 0,283230726 | 0,000132007 | 0,001016041 | Up |
| Pard3 | 118,9275553 | 0,834884835 | 0,324230814 | 5,11E-05 | 0,000452922 | Up |
| Maml2 | 1577,826102 | 0,769664186 | 0,156076574 | 1,01E-08 | 2,52E-07 | Up |
| RP24-496O17,7 | 26,46044764 | 1,197242763 | 0,488737369 | 2,85E-05 | 0,000274739 | Up |
| Ccdc151 | 5,124378888 | 0,604981707 | 0,842876182 | 0,005013199 | 0,021546797 | Up |
| Pknox2 | 56,51895352 | -1,30515453 | 0,661063154 | 9,62E-05 | 0,000776394 | Down |
| Robo3 | 29,5167106 | -2,020009956 | 0,550755783 | 1,26E-06 | 1,82E-05 | Down |
| Vsig2 | 48,79650489 | 0,80401565 | 0,376421148 | 0,000181869 | 0,001332806 | Up |
| Nrgn | 5176,970255 | 0,883560022 | 0,174321139 | 3,22E-09 | 9,13E-08 | Up |
| Hyou1 | 86610,38977 | -0,800137861 | 0,149696682 | 1,14E-09 | 3,57E-08 | Down |
| Gm32742 | 44,04069002 | 0,603174394 | 0,498295126 | 0,002912089 | 0,013769743 | Up |
| Lman1l | 22,96843767 | -0,796731946 | 0,445232042 | 0,000419005 | 0,002728343 | Down |
| Cd276 | 489,4175603 | -2,825723105 | 0,249023422 | 8,18E-31 | 6,03E-28 | Down |
| Tle3 | 102,443084 | -1,969679047 | 0,37816393 | 3,05E-09 | 8,72E-08 | Down |
| Slc24a1 | 31,36208682 | 0,756767898 | 0,464592549 | 0,000693333 | 0,004141873 | Up |
| Parp16 | 265,9383329 | 0,674661649 | 0,148032956 | 7,33E-08 | 1,47E-06 | Up |
| Car12 | 3838,001602 | -0,802703818 | 0,115080962 | 5,45E-14 | 4,40E-12 | Down |
| Aqp9 | 2236,450141 | -2,814612135 | 0,379785032 | 1,09E-14 | 9,70E-13 | Down |
| Gm37234 | 51,52794593 | -0,867879064 | 0,535804173 | 0,000480215 | 0,003049508 | Down |
| Ccpg1os | 184,6730959 | 0,744974806 | 0,142522003 | 2,43E-09 | 7,03E-08 | Up |
| Gsta4 | 475,055393 | 0,823090698 | 0,151440192 | 6,49E-10 | 2,19E-08 | Up |
| Sh3bgrl2 | 22,9565093 | -0,615745957 | 0,658587095 | 0,004001056 | 0,017868742 | Down |
| Bcl2a1d | 151,2978307 | -1,849189363 | 0,292259841 | 4,52E-12 | 2,48E-10 | Down |
| Bcl2a1a | 3219,401871 | -2,094100515 | 0,317443104 | 1,88E-12 | 1,11E-10 | Down |
| Bcl2a1b | 1443,482025 | -2,080056648 | 0,238864832 | 2,13E-19 | 4,01E-17 | Down |
| Pcolce2 | 191,0624723 | 1,23953177 | 0,268610057 | 7,26E-09 | 1,87E-07 | Up |
| Pls1 | 3170,196103 | 2,285737567 | 0,200339222 | 4,15E-31 | 3,20E-28 | Up |
| 1700065D16Rik | 152,2227638 | 1,890207325 | 0,234217895 | 2,47E-17 | 3,37E-15 | Up |
| Rasa2 | 3208,881562 | -0,590672052 | 0,124612117 | 3,62E-08 | 7,90E-07 | Down |
| Gm26767 | 319,5603008 | -2,216263632 | 1,384314381 | 0,000106755 | 0,000846993 | Down |
| Ky | 967,2072409 | -2,269350693 | 0,146088754 | 2,42E-55 | 1,31E-51 | Down |
| Cep63 | 5322,413822 | -0,615725421 | 0,153137735 | 9,25E-07 | 1,40E-05 | Down |
| Srprb | 8688,194461 | -0,837686116 | 0,166015056 | 4,51E-09 | 1,23E-07 | Down |
| Bfsp2 | 56,3096011 | -1,868232268 | 0,310617004 | 3,11E-11 | 1,42E-09 | Down |
| Tlr9 | 120,2603843 | -1,471577437 | 0,64126883 | 3,88E-05 | 0,00035669 | Down |
| Alas1 | 5364,66239 | -0,948788383 | 0,121517051 | 4,55E-17 | 5,81E-15 | Down |
| Dusp7 | 2854,224814 | -0,995068419 | 0,085849221 | 2,40E-33 | 2,29E-30 | Down |
| Rrp9 | 13275,934 | -0,63819409 | 0,100409497 | 3,42E-12 | 1,91E-10 | Down |
| Manf | 21620,3964 | -0,645353906 | 0,193935793 | 1,22E-05 | 0,000131307 | Down |
| Mapkapk3 | 18249,63982 | -1,128966193 | 0,182431134 | 1,25E-12 | 7,55E-11 | Down |
| Cish | 14896,70043 | -2,458972797 | 0,150488188 | 6,67E-61 | 5,41E-57 | Down |
| Hemk1 | 5617,126081 | -1,221774104 | 0,158504769 | 1,70E-17 | 2,38E-15 | Down |
| 6430571L13Rik | 340,745674 | -2,047172816 | 0,192156728 | 1,28E-27 | 5,63E-25 | Down |
| Gm42432 | 31,82440315 | -0,791726431 | 0,556125713 | 0,000927673 | 0,005316672 | Down |
| Kif9 | 100,6737366 | 0,596721206 | 0,225313064 | 0,000140741 | 0,001070571 | Up |
| Gm42762 | 74,11714831 | 1,713268238 | 0,382675489 | 3,65E-08 | 7,95E-07 | Up |
| Ltf | 278,6434042 | -1,294764084 | 0,429744692 | 4,71E-06 | 5,75E-05 | Down |
| Ccr4 | 3470,273775 | -0,753732097 | 0,171625827 | 1,27E-07 | 2,42E-06 | Down |
| Csrnp1 | 4757,737723 | -0,718775078 | 0,125364457 | 1,58E-10 | 6,01E-09 | Down |
| RP24-233B16,1 | 11,75484017 | -0,760263603 | 0,722025357 | 0,002058383 | 0,010308002 | Down |
| Pomgnt2 | 40,24995299 | 0,90577671 | 0,437756924 | 0,000146749 | 0,001110026 | Up |

| | | | | | | |
|---|---|---|---|---|---|---|
| Abhd5 | 4995,859859 | -0,640415106 | 0,120398005 | 1,64E-09 | 4,95E-08 | Down |
| Ccr1 | 520,2067066 | -2,818410591 | 0,604455298 | 8,60E-08 | 1,69E-06 | Down |
| Grm1 | 15,35233286 | 0,968423327 | 0,821625239 | 0,000995407 | 0,005631269 | Up |
| Aig1 | 5,711943907 | -0,884214101 | 1,284298535 | 0,00171949 | 0,008856738 | Down |
| Raet1d | 35,58970094 | -3,031398862 | 0,795675725 | 1,12E-06 | 1,64E-05 | Down |
| Gm10825 | 50,07323976 | -7,028703539 | 1,254434552 | 1,91E-11 | 9,24E-10 | Down |
| Cdk19 | 4869,431134 | 0,77818424 | 0,11265539 | 9,21E-14 | 7,05E-12 | Up |
| Scml4 | 61,52170318 | 3,178792584 | 0,418619275 | 1,85E-15 | 1,88E-13 | Up |
| Lilrb4a | 4829,529236 | -4,099517616 | 0,579815975 | 1,23E-14 | 1,07E-12 | Down |
| Cdh23 | 245,9054833 | 0,983528527 | 0,370019504 | 2,34E-05 | 0,000231153 | Up |
| Sgpl1 | 5661,291754 | -0,59670433 | 0,125658544 | 3,42E-08 | 7,51E-07 | Down |
| Hkdc1 | 599,7391645 | -1,378953142 | 0,24977835 | 6,69E-11 | 2,80E-09 | Down |
| Rhobtb1 | 961,7149619 | -0,647742176 | 0,094345646 | 1,14E-13 | 8,45E-12 | Down |
| Adora2a | 4036,259336 | 1,164084987 | 0,136744993 | 2,42E-20 | 5,38E-18 | Up |
| Ggt5 | 50,42020708 | 0,841701485 | 0,352329421 | 8,18E-05 | 0,000677308 | Up |
| Gm10941 | 15,40124955 | 0,908008229 | 0,605130324 | 0,000563717 | 0,003473724 | Up |
| Agpat3 | 3052,747228 | -0,952280008 | 0,129570708 | 1,46E-15 | 1,51E-13 | Down |
| Plppr3 | 5055,720158 | -0,592972509 | 0,207092187 | 7,47E-05 | 0,000625685 | Down |
| Gamt | 53,96634743 | 0,690410163 | 0,296488444 | 0,000197878 | 0,001435215 | Up |
| Apc2 | 208,6627662 | 0,914071696 | 0,259956584 | 1,94E-06 | 2,65E-05 | Up |
| Reep6 | 194,4615902 | -0,699181848 | 0,156804026 | 1,09E-07 | 2,11E-06 | Down |
| Btbd2 | 3933,672217 | 0,730342641 | 0,139275571 | 2,28E-09 | 6,63E-08 | Up |
| Izumo4 | 21,48224716 | 0,610134296 | 0,606059803 | 0,00373473 | 0,016897935 | Up |
| Smim24 | 924,2397926 | 0,909169623 | 0,212071366 | 1,02E-07 | 1,97E-06 | Up |
| Celf5 | 35,77832373 | 0,82259083 | 0,448293718 | 0,000342782 | 0,002302004 | Up |
| RP23-132J19,1 | 72,17940298 | -2,758568236 | 0,322987507 | 1,33E-18 | 2,23E-16 | Down |
| Bpifc | 1080,100301 | 0,994535858 | 0,939767128 | 0,001187004 | 0,006535168 | Up |
| A230060F14Rik | 81,9251504 | 2,212063968 | 0,325758483 | 6,82E-13 | 4,31E-11 | Up |
| Fbxo7 | 10180,18259 | 1,315372673 | 0,191004774 | 8,84E-15 | 8,05E-13 | Up |
| Syn3 | 40,9513894 | 2,592353614 | 0,49909875 | 8,90E-09 | 2,24E-07 | Up |
| Gm26546 | 34,28311744 | 0,644104068 | 0,387223667 | 0,001109665 | 0,00617958 | Up |
| Gas2l3 | 16653,24652 | 0,999668457 | 0,195316087 | 1,27E-09 | 3,95E-08 | Up |
| Nr1h4 | 81,58779119 | 3,934318449 | 0,636274186 | 7,00E-12 | 3,66E-10 | Up |
| Slc17a8 | 23,32408474 | 2,202919976 | 1,023621029 | 4,62E-05 | 0,000414902 | Up |
| 1500026H17Rik | 14,69804799 | 1,053174758 | 0,851559348 | 0,000760911 | 0,004491072 | Up |
| RP23-381H23,1 | 442,2506705 | 0,812922125 | 0,180746811 | 6,47E-08 | 1,32E-06 | Up |
| Gm15915 | 150,4032388 | -1,108560352 | 0,214524045 | 5,66E-10 | 1,94E-08 | Down |
| Cep83os | 549,2854676 | 0,730006997 | 0,220462475 | 8,95E-06 | 0,000100345 | Up |
| Plxnc1 | 718,5796189 | -1,428555133 | 0,19099368 | 1,78E-16 | 2,14E-14 | Down |
| Dusp6 | 5131,240224 | -0,686876644 | 0,151742376 | 8,25E-08 | 1,63E-06 | Down |
| Gm15663 | 429,8583652 | -0,67135465 | 0,145836772 | 5,95E-08 | 1,22E-06 | Down |
| Phlda1 | 371,3231829 | -0,597895095 | 0,175020827 | 1,09E-05 | 0,00011914 | Down |
| Tmem5 | 4017,668629 | -0,623055996 | 0,104551182 | 4,10E-11 | 1,82E-09 | Down |
| Avil | 247,8839592 | -1,356073481 | 1,069617376 | 0,000471942 | 0,003001669 | Down |
| Kif5a | 352,4981487 | 0,886593685 | 0,197379489 | 4,73E-08 | 9,98E-07 | Up |
| Lrp1 | 58,39551314 | -0,592388994 | 0,344646543 | 0,001357614 | 0,007284155 | Down |
| Gm26847 | 35,45656891 | 0,61014851 | 0,3774581 | 0,001467763 | 0,007769805 | Up |
| Il23a | 78,30081049 | -0,797736606 | 0,328528696 | 8,95E-05 | 0,000729157 | Down |
| A430046D13Rik | 300,764932 | 0,604150882 | 0,174825559 | 9,18E-06 | 0,000102757 | Up |
| Suox | 2564,30032 | 0,677435062 | 0,087438366 | 1,82E-16 | 2,14E-14 | Up |
| Bloc1s1 | 493,9946311 | 0,626075882 | 0,128322461 | 1,68E-08 | 3,95E-07 | Up |

| | | | | | | |
|---|---|---|---|---|---|---|
| Ccdc157 | 239,7664646 | 0,949595647 | 0,196461564 | 7,02E-09 | 1,81E-07 | Up |
| Gatsl3 | 234,9136018 | -2,490726218 | 0,220065678 | 1,29E-30 | 9,07E-28 | Down |
| Osm | 2398,355282 | -1,62140132 | 0,136621593 | 2,32E-34 | 2,51E-31 | Down |
| Lif | 95,69670836 | -1,963438355 | 0,289500453 | 3,88E-13 | 2,56E-11 | Down |
| Znrf3 | 2485,085134 | -0,614924576 | 0,092335313 | 4,49E-13 | 2,91E-11 | Down |
| Gm11963 | 137,8353194 | -0,820284174 | 0,265404309 | 1,21E-05 | 0,000129806 | Down |
| Xbp1 | 46769,59394 | -0,94746844 | 0,106918877 | 6,60E-21 | 1,55E-18 | Down |
| Polm | 1099,511717 | 0,863859182 | 0,160983215 | 7,76E-10 | 2,56E-08 | Up |
| H2afv | 5771,250671 | 0,76017987 | 0,145275521 | 2,28E-09 | 6,64E-08 | Up |
| Upp1 | 1663,281966 | 1,049569872 | 0,202326798 | 6,70E-10 | 2,25E-08 | Up |
| Abca13 | 258,6533689 | -0,979193649 | 0,274335109 | 1,22E-06 | 1,78E-05 | Down |
| Plek | 48649,19621 | 1,154687024 | 0,162375177 | 1,94E-15 | 1,97E-13 | Up |
| Wdpcp | 417,7205719 | 0,934201543 | 0,161076209 | 4,64E-11 | 2,00E-09 | Up |
| Hba-x | 4,178230899 | 0,937447905 | 1,5132402 | 0,000998943 | 0,005645238 | Up |
| Hba-a1 | 127,3128889 | 3,78005082 | 1,203712486 | 2,21E-06 | 2,97E-05 | Up |
| Olfr56 | 109,8057391 | -1,214756673 | 0,255082382 | 3,57E-09 | 9,97E-08 | Down |
| Ifi47 | 1481,469323 | -0,693809333 | 0,149793307 | 5,00E-08 | 1,04E-06 | Down |
| N4bp3 | 3027,908061 | 0,628035484 | 0,114790498 | 7,13E-10 | 2,37E-08 | Up |
| D930048N14Rik | 299,0755494 | 0,964090473 | 0,199690613 | 6,65E-09 | 1,73E-07 | Up |
| Tcf7 | 7,876019983 | 0,586219341 | 0,633048746 | 0,004564583 | 0,019940861 | Up |
| Gm12216 | 52,18333005 | 0,974039388 | 0,288510711 | 2,47E-06 | 3,27E-05 | Up |
| Acsl6 | 220,3004438 | 1,185435799 | 0,235679704 | 9,38E-10 | 3,01E-08 | Up |
| Slc36a2 | 101,0701759 | 1,026692127 | 0,501338159 | 0,000112898 | 0,000886626 | Up |
| Fat2 | 185,720143 | -0,899076507 | 0,397283007 | 9,14E-05 | 0,000743152 | Down |
| Gm12247 | 49,05827899 | 0,810811903 | 0,369649895 | 0,000153419 | 0,001154558 | Up |
| Trim58 | 2091,574301 | 1,033295663 | 0,12864985 | 3,46E-18 | 5,45E-16 | Up |
| Gm12258 | 129,3155738 | 0,77077119 | 0,237906261 | 9,43E-06 | 0,000104786 | Up |
| Natd1 | 989,8476047 | 0,752226507 | 0,159474743 | 2,95E-08 | 6,51E-07 | Up |
| Specc1 | 7087,684873 | 0,93604938 | 0,133823163 | 2,17E-14 | 1,85E-12 | Up |
| Lrrc75a | 1127,904021 | 0,702681781 | 0,177217977 | 8,89E-07 | 1,35E-05 | Up |
| Trim16 | 1370,837268 | -0,739980368 | 0,492100355 | 0,000954727 | 0,005440972 | Down |
| Hs3st3b1 | 2950,344531 | -1,000277655 | 0,098278467 | 1,23E-26 | 4,74E-24 | Down |
| Slc25a35 | 67,47216633 | 1,037502245 | 0,59447626 | 0,000239003 | 0,001683816 | Up |
| 9330160F10Rik | 24,12097283 | 0,73893583 | 0,479039744 | 0,000888095 | 0,005131397 | Up |
| Alox8 | 110,5484431 | -0,6636965 | 0,262212673 | 0,00013473 | 0,001033078 | Down |
| Kdm6b | 6599,126144 | -1,312502502 | 0,120348524 | 1,37E-30 | 9,24E-28 | Down |
| Plscr3 | 1690,338622 | 0,632417281 | 0,074514214 | 3,73E-19 | 6,80E-17 | Up |
| 2810408A11Rik | 34,14067 | 0,976054822 | 0,398168068 | 4,31E-05 | 0,000391208 | Up |
| Gp1ba | 496,4015227 | 1,240617605 | 0,202879443 | 1,52E-12 | 9,10E-11 | Up |
| Rnf167 | 5108,691452 | 0,917585601 | 0,091366755 | 1,16E-25 | 4,01E-23 | Up |
| Aipl1 | 422,8715118 | 0,706734337 | 0,214945785 | 1,07E-05 | 0,000117256 | Up |
| Fam64a | 7453,227251 | 0,777152332 | 0,170672067 | 5,77E-08 | 1,19E-06 | Up |
| Xaf1 | 63,53803563 | -0,731976862 | 0,296362229 | 0,000110442 | 0,000871281 | Down |
| Spns2 | 32535,43036 | 0,684350911 | 0,106833118 | 2,61E-12 | 1,49E-10 | Up |
| Gm45606 | 553,0133456 | 0,722343245 | 0,166616583 | 1,78E-07 | 3,25E-06 | Up |
| Gm12341 | 96,30970412 | -0,824585679 | 0,233047364 | 2,69E-06 | 3,51E-05 | Down |
| Nek8 | 533,294894 | 0,59214214 | 0,162142411 | 4,56E-06 | 5,62E-05 | Up |
| Tmem97 | 13701,63084 | -0,624355182 | 0,091305959 | 1,33E-13 | 9,76E-12 | Down |
| Gm11417 | 102,8043685 | 0,669734226 | 0,205021104 | 1,36E-05 | 0,000144116 | Up |
| Slfn5 | 587,8125687 | -1,429118801 | 0,279369272 | 7,12E-10 | 2,37E-08 | Down |
| Slfn10-ps | 1497,875807 | -0,769146903 | 0,094443553 | 8,65E-18 | 1,30E-15 | Down |

| | | | | | | |
|---|---|---|---|---|---|---|
| Slfn2 | 1045,664031 | -3,548984217 | 0,333915674 | 7,27E-28 | 3,37E-25 | Down |
| Slfn1 | 8,722753994 | -3,501728999 | 1,758725062 | 1,45E-05 | 0,000152202 | Down |
| Ccl6 | 1119,856899 | -2,687568433 | 0,902125061 | 7,97E-06 | 9,09E-05 | Down |
| Ccl3 | 441,8749966 | -2,444564875 | 0,453184144 | 3,09E-09 | 8,81E-08 | Down |
| Ccl4 | 335,2517241 | -3,121291542 | 0,354273586 | 8,72E-20 | 1,75E-17 | Down |
| Acsf2 | 171,4211119 | 1,050104663 | 0,228204278 | 1,26E-08 | 3,06E-07 | Up |
| Sgca | 18,65168968 | -0,854006685 | 0,582329096 | 0,000700711 | 0,004181332 | Down |
| Nfe2l1 | 20498,0326 | -0,602009069 | 0,075340219 | 2,19E-17 | 3,02E-15 | Down |
| Tbkbp1 | 9540,603635 | -0,625151783 | 0,103868646 | 2,85E-11 | 1,33E-09 | Down |
| Plxdc1 | 19,07902451 | 0,78111829 | 0,529554535 | 0,000866398 | 0,00503125 | Up |
| Cacnb1 | 130,9913445 | 0,660995978 | 0,270683375 | 0,000173756 | 0,001280288 | Up |
| Fbxl20 | 4941,19867 | 0,746641436 | 0,170337024 | 1,35E-07 | 2,55E-06 | Up |
| Rara | 1249,45101 | -1,401794776 | 0,132493479 | 7,97E-29 | 4,17E-26 | Down |
| Cnp | 6752,069168 | 0,669516593 | 0,139104542 | 2,18E-08 | 5,00E-07 | Up |
| Plekhh3 | 34,01354151 | -1,505799002 | 0,571628784 | 1,54E-05 | 0,000159588 | Down |
| Cntnap1 | 15,48734515 | -0,879377878 | 0,708679907 | 0,001057476 | 0,005927111 | Down |
| Etv4 | 842,4524741 | -1,813415555 | 0,181981064 | 7,43E-25 | 2,32E-22 | Down |
| Myl4 | 105,1560188 | 0,749395537 | 0,23965869 | 1,50E-05 | 0,000156448 | Up |
| Itgb3 | 28796,22896 | 1,71422014 | 0,128448593 | 3,37E-42 | 7,81E-39 | Up |
| Milr1 | 749,3495395 | -1,850926782 | 0,250982731 | 4,20E-15 | 4,08E-13 | Down |
| Slc16a6 | 11528,65552 | -0,648110237 | 0,133097638 | 1,70E-08 | 4,00E-07 | Down |
| Cd300ld | 194,4014277 | -0,629832897 | 0,152971585 | 5,86E-07 | 9,29E-06 | Down |
| Cd300ld2 | 45,67226224 | -0,657604282 | 0,335092595 | 0,000561444 | 0,003463339 | Down |
| Cd300lf | 3294,058118 | -0,707920671 | 0,108402198 | 1,18E-12 | 7,19E-11 | Down |
| Fads6 | 493,9930731 | 0,878662049 | 0,211025688 | 1,97E-07 | 3,57E-06 | Up |
| Hid1 | 20,88227253 | 0,741366108 | 0,709454 | 0,002206243 | 0,010930165 | Up |
| Slc16a5 | 242,7657574 | 0,635542433 | 0,146078017 | 2,06E-07 | 3,70E-06 | Up |
| Smim5 | 8823,774637 | 1,021238378 | 0,135337729 | 1,81E-16 | 2,14E-14 | Up |
| Rhbdf2 | 7965,901881 | -0,601214946 | 0,123573514 | 1,88E-08 | 4,38E-07 | Down |
| Tha1 | 1141,932535 | 0,669600028 | 0,156454582 | 2,60E-07 | 4,55E-06 | Up |
| C1qtnf1 | 9,56653061 | -1,099301332 | 0,8596269 | 0,00064793 | 0,003922638 | Down |
| Sgsh | 2347,24797 | 1,460625317 | 0,233847179 | 1,13E-12 | 6,86E-11 | Up |
| Slc26a11 | 69,14630225 | 1,201759411 | 0,477694804 | 2,34E-05 | 0,000231207 | Up |
| Gm11767 | 51,92071233 | 0,835071663 | 0,33612851 | 6,53E-05 | 0,00055841 | Up |
| Cep131 | 3246,244364 | 0,725921623 | 0,096757439 | 1,30E-15 | 1,39E-13 | Up |
| 2900052L18Rik | 562,9252861 | 0,581849541 | 0,229732045 | 0,00021428 | 0,001533478 | Up |
| 0610009L18Rik | 162,2274992 | 0,713351717 | 0,232170277 | 2,11E-05 | 0,00021209 | Up |
| Oxld1 | 699,1327809 | 0,582825174 | 0,08649943 | 2,63E-13 | 1,82E-11 | Up |
| Narf | 2989,659917 | 0,867954275 | 0,177261452 | 8,09E-09 | 2,06E-07 | Up |
| Metrnl | 4887,976873 | -0,893661335 | 0,108221386 | 1,93E-18 | 3,13E-16 | Down |
| 1110002L01Rik | 454,7787592 | -0,663883365 | 0,13730012 | 1,97E-08 | 4,54E-07 | Down |
| Fkbp1b | 190,2514802 | 0,916004722 | 0,218656415 | 1,48E-07 | 2,76E-06 | Up |
| Mycn | 8714,752032 | -0,65905234 | 0,212489944 | 2,48E-05 | 0,000243418 | Down |
| Greb1 | 5300,365898 | 7,225006399 | 0,537535015 | 1,35E-43 | 3,66E-40 | Up |
| Gm4419 | 81,5986721 | 0,895651758 | 0,278269016 | 5,74E-06 | 6,83E-05 | Up |
| Dus4l | 2202,641229 | -0,701408051 | 0,119205854 | 6,60E-11 | 2,77E-09 | Down |
| Hdac9 | 275,840004 | -1,969181877 | 0,284753802 | 1,63E-13 | 1,17E-11 | Down |
| Ifrd1 | 11870,87126 | -1,043229307 | 0,156929041 | 1,00E-13 | 7,60E-12 | Down |
| Mia2,1 | 18210,05399 | -0,6042741 | 0,091930164 | 8,02E-13 | 5,04E-11 | Down |
| Prkch | 22395,79984 | 0,799106316 | 0,079774714 | 3,29E-25 | 1,09E-22 | Up |
| Rhoj | 8706,867025 | -1,809327125 | 0,539625518 | 2,45E-06 | 3,24E-05 | Down |

| | | | | | | |
|---|---|---|---|---|---|---|
| 2310015A10Rik | 599,3068281 | 0,670557455 | 0,11456009 | 7,86E-11 | 3,23E-09 | Up |
| Sipa1l1 | 98257,06086 | 0,673571461 | 0,116149334 | 1,08E-10 | 4,35E-09 | Up |
| Prox2 | 519,098261 | 0,728270422 | 0,105711526 | 1,08E-13 | 8,10E-12 | Up |
| Tgfb3 | 214,763744 | 2,138052033 | 0,318193401 | 9,43E-13 | 5,86E-11 | Up |
| Ift43 | 351,6689502 | 0,589612537 | 0,144581894 | 7,92E-07 | 1,22E-05 | Up |
| Ston2 | 4047,202282 | 0,659878616 | 0,137271653 | 2,28E-08 | 5,18E-07 | Up |
| Ifi27 | 6938,423981 | -1,503097754 | 0,236250777 | 6,58E-13 | 4,19E-11 | Down |
| Serpina3f | 222,9316859 | -2,214115587 | 0,348419078 | 1,11E-11 | 5,50E-10 | Down |
| Zfp839 | 686,5180136 | 0,595456355 | 0,146758224 | 8,47E-07 | 1,30E-05 | Up |
| Cdc42bpb | 66,57133961 | 2,570279584 | 1,15267409 | 3,13E-05 | 0,000297751 | Up |
| Trmt61a | 4466,011243 | -0,61707903 | 0,124877437 | 1,24E-08 | 3,02E-07 | Down |
| Nudt14 | 1734,516994 | 0,686232609 | 0,111633763 | 1,34E-11 | 6,56E-10 | Up |
| Ighm | 1457,829526 | 0,587878714 | 0,255002781 | 0,000381161 | 0,002524219 | Up |
| Idi1 | 7542,710232 | 0,946801693 | 0,203617565 | 1,69E-08 | 3,98E-07 | Up |
| Lyst | 43,14978485 | -1,785848431 | 0,946961085 | 9,56E-05 | 0,000772257 | Down |
| Gpr141 | 1956,96167 | -1,475161479 | 0,652501292 | 4,24E-05 | 0,000386359 | Down |
| Zkscan4 | 181,0267623 | 1,161901511 | 0,23422098 | 1,42E-09 | 4,33E-08 | Up |
| Mboat1 | 3788,958972 | -0,612852223 | 0,225514935 | 0,000104767 | 0,000836123 | Down |
| Serpinb9g | 19,49516199 | -2,336312711 | 0,628390589 | 1,47E-06 | 2,08E-05 | Down |
| Slc22a23 | 6460,465171 | 0,726895964 | 0,12149423 | 3,65E-11 | 1,64E-09 | Up |
| Ppp1r3g | 145,2577066 | -1,365776677 | 0,590620699 | 3,78E-05 | 0,000348704 | Down |
| Eef1e1 | 7193,167973 | -0,856910847 | 0,099919296 | 1,70E-19 | 3,25E-17 | Down |
| Mylip | 1392,91633 | 0,679068675 | 0,12613224 | 1,13E-09 | 3,56E-08 | Up |
| S1pr3 | 2148,774827 | 0,744123982 | 0,190874634 | 1,00E-06 | 1,50E-05 | Up |
| Nfil3 | 7839,401382 | -0,649954971 | 0,109960654 | 5,53E-11 | 2,36E-09 | Down |
| Hrh2 | 19,27366988 | -0,633069412 | 0,688314284 | 0,003814836 | 0,01720281 | Down |
| Unc5a | 833,5031363 | 0,860678349 | 0,21633451 | 4,49E-07 | 7,39E-06 | Up |
| Hk3 | 26800,88562 | -1,0890173 | 0,210480804 | 5,94E-10 | 2,03E-08 | Down |
| Prr7 | 516,0034562 | -0,761569771 | 0,128397191 | 4,79E-11 | 2,06E-09 | Down |
| 4930486L24Rik | 154,5300492 | 1,403491135 | 0,369671552 | 2,93E-07 | 5,08E-06 | Up |
| Ctla2b | 974,1739352 | 2,258993031 | 0,211757493 | 1,51E-27 | 6,44E-25 | Up |
| Ctla2a | 2671,589779 | 2,271430315 | 0,174764632 | 1,45E-39 | 2,94E-36 | Up |
| Fbp2 | 13397,45589 | -0,849437908 | 0,198840238 | 1,45E-07 | 2,71E-06 | Down |
| Fbp1 | 506,5409413 | -2,370403785 | 0,305517511 | 7,80E-16 | 8,61E-14 | Down |
| Fancc | 5048,413678 | 1,036976895 | 0,152443888 | 3,63E-14 | 3,04E-12 | Up |
| Gm28557 | 187,792033 | 0,627897128 | 0,157158175 | 9,90E-07 | 1,49E-05 | Up |
| Mctp1 | 306,1510972 | 1,050337231 | 0,328021009 | 3,49E-06 | 4,44E-05 | Up |
| Tmem161b | 1137,04339 | -0,595523656 | 0,116221053 | 4,96E-09 | 1,33E-07 | Down |
| F2r | 100569,7073 | 1,084247868 | 0,232064409 | 7,80E-09 | 2,00E-07 | Up |
| Rnf180 | 15,47976487 | 0,918296344 | 0,678084739 | 0,000760922 | 0,004491072 | Up |
| Emb | 15483,94736 | -1,543304706 | 0,16347519 | 2,22E-23 | 6,21E-21 | Down |
| Acox2 | 9,330206791 | -2,377221375 | 1,866985612 | 0,00010002 | 0,000801793 | Down |
| Slc4a7 | 6190,52347 | -0,595211685 | 0,062646042 | 3,36E-23 | 9,25E-21 | Down |
| Ndst2 | 7587,748567 | 0,862551746 | 0,107396286 | 1,52E-17 | 2,19E-15 | Up |
| Plau | 7,995041561 | -0,761289384 | 0,738124391 | 0,002127769 | 0,010589896 | Down |
| Samd8 | 2754,828224 | -0,646906519 | 0,086776422 | 1,59E-15 | 1,63E-13 | Down |
| Kcnma1 | 68,5027925 | -1,520125609 | 1,226843567 | 0,000398996 | 0,002621996 | Down |
| Chdh | 23,90098117 | -0,720996331 | 0,568578541 | 0,001600352 | 0,008346422 | Down |
| Itih4 | 63,65729192 | 1,706438393 | 0,663045036 | 1,93E-05 | 0,000195342 | Up |
| Sh3bp5 | 733,0241111 | -0,919771074 | 0,229351683 | 2,99E-07 | 5,17E-06 | Down |
| Eaf1 | 14776,4521 | -0,651653931 | 0,105233805 | 9,81E-12 | 4,94E-10 | Down |

| | | | | | | |
|---|---|---|---|---|---|---|
| Colq | 46,55451075 | -0,952714055 | 0,335925816 | 1,52E-05 | 0,000158467 | Down |
| Gch1 | 3248,718667 | -0,71749891 | 0,172644345 | 3,88E-07 | 6,51E-06 | Down |
| Gm26782 | 363,4083126 | -0,612518554 | 0,162123342 | 2,55E-06 | 3,36E-05 | Down |
| Rnase10 | 8,856081127 | -0,817205962 | 1,242939144 | 0,00203863 | 0,010221499 | Down |
| Arhgef40 | 1049,080184 | 0,712466643 | 0,634850347 | 0,002127264 | 0,010589896 | Up |
| Zfp219 | 5413,154234 | 0,616777179 | 0,073351736 | 7,00E-19 | 1,22E-16 | Up |
| Olfr1509 | 53,89086866 | 0,611405859 | 0,289847493 | 0,000531524 | 0,003306741 | Up |
| B230359F08Rik | 15,86769749 | 1,018463137 | 0,84654494 | 0,000864829 | 0,005023933 | Up |
| Trac | 148,0237605 | 3,631816843 | 0,356102339 | 5,42E-26 | 1,95E-23 | Up |
| Abhd4 | 11072,85963 | 1,209979605 | 0,113482681 | 1,65E-29 | 9,22E-27 | Up |
| Oxa1l | 19222,95985 | 0,621296683 | 0,084739612 | 3,79E-15 | 3,72E-13 | Up |
| Slc7a7 | 7431,349218 | 1,540203868 | 0,177818478 | 2,51E-20 | 5,50E-18 | Up |
| Haus4 | 5870,874913 | 0,857740459 | 0,075283388 | 9,33E-32 | 7,57E-29 | Up |
| 4931414P19Rik | 1814,180241 | 0,740527236 | 0,091400164 | 1,22E-17 | 1,79E-15 | Up |
| Gm37271 | 94,0022124 | -0,894361335 | 0,402756625 | 0,000103687 | 0,000828731 | Down |
| Adcy4 | 351,8680846 | -0,869636376 | 0,156130772 | 2,51E-10 | 9,42E-09 | Down |
| Ripk3 | 124,2185648 | -2,416061458 | 0,300866478 | 9,43E-17 | 1,16E-14 | Down |
| Mcpt1 | 70,48400238 | 4,496897082 | 0,523288614 | 2,68E-20 | 5,79E-18 | Up |
| Mcpt2 | 338,2446608 | 1,73723753 | 0,164519541 | 1,15E-27 | 5,16E-25 | Up |
| Gzmc | 158,6019186 | 3,299513239 | 1,07980558 | 4,31E-06 | 5,33E-05 | Up |
| Gzmb | 5878,048741 | 1,498183636 | 0,418154663 | 7,41E-07 | 1,15E-05 | Up |
| Fgf9 | 1434,587232 | -1,106515757 | 0,080962832 | 2,52E-45 | 8,19E-42 | Down |
| Sgcg | 7,308423166 | 1,956674176 | 1,100373587 | 0,000106047 | 0,000843545 | Up |
| Nuggc | 871,5562984 | 0,592120918 | 0,203592146 | 6,50E-05 | 0,000556132 | Up |
| Slc39a14 | 6858,21372 | -0,634855849 | 0,111378908 | 1,92E-10 | 7,25E-09 | Down |
| Polr3d | 3655,168708 | -0,677774628 | 0,146021827 | 4,92E-08 | 1,03E-06 | Down |
| Dok2 | 12679,57408 | -0,788998391 | 0,088983775 | 1,79E-20 | 4,04E-18 | Down |
| Tbc1d4 | 1398,062824 | 0,682300402 | 0,172007426 | 9,42E-07 | 1,43E-05 | Up |
| Acod1 | 37,89133951 | -1,59707911 | 0,551606569 | 7,60E-06 | 8,73E-05 | Down |
| Dzip1 | 190,1343369 | 1,192135192 | 0,200526714 | 4,74E-12 | 2,59E-10 | Up |
| Mtmr12 | 10351,46479 | 0,739192324 | 0,091673574 | 1,67E-17 | 2,35E-15 | Up |
| Pdzd2 | 426,2739829 | 1,604202368 | 0,320720081 | 2,31E-09 | 6,70E-08 | Up |
| Gm2606 | 50,5075093 | 1,409890159 | 1,007706485 | 0,000345655 | 0,002319375 | Up |
| Trio | 13127,54574 | -0,594235287 | 0,075644922 | 6,44E-17 | 8,11E-15 | Down |
| Mtdh | 38220,90968 | -0,659427121 | 0,098770645 | 4,23E-13 | 2,77E-11 | Down |
| Slc25a32 | 3606,511636 | -0,615456494 | 0,089262484 | 8,85E-14 | 6,81E-12 | Down |
| Lrp12 | 26105,19814 | -0,899917746 | 0,151984046 | 2,86E-11 | 1,33E-09 | Down |
| Nov | 32,77256783 | -3,247460469 | 0,59083247 | 1,41E-09 | 4,30E-08 | Down |
| Mtss1 | 4672,466355 | 0,746417139 | 0,109916519 | 2,11E-13 | 1,49E-11 | Up |
| Trib1 | 185,8413785 | -0,612266114 | 0,252968625 | 0,000243395 | 0,001710303 | Down |
| Sla | 19085,22512 | 1,229687482 | 0,158702502 | 1,23E-17 | 1,79E-15 | Up |
| Ndrg1 | 5030,416346 | -0,719479916 | 0,159960295 | 8,75E-08 | 1,72E-06 | Down |
| St3gal1 | 9612,884407 | 0,717987109 | 0,114627093 | 6,57E-12 | 3,48E-10 | Up |
| Ly6g | 18,38853675 | -3,352446569 | 1,351717357 | 9,94E-06 | 0,000109534 | Down |
| 9030619P08Rik | 3853,176387 | 1,664391867 | 0,161790326 | 1,26E-26 | 4,74E-24 | Up |
| Kifc2 | 11,06650527 | 0,917813247 | 0,765339763 | 0,001056866 | 0,005927111 | Up |
| Apol10b | 333,8324629 | 0,692745777 | 0,2375061 | 3,79E-05 | 0,000349573 | Up |
| Il2rb | 2124,240064 | -1,805069725 | 0,155715457 | 1,72E-32 | 1,47E-29 | Down |
| Maff | 559,8860758 | -1,806940062 | 0,164125731 | 1,27E-29 | 7,33E-27 | Down |
| Gm16576 | 461,5269038 | 0,648153464 | 0,154863653 | 4,15E-07 | 6,89E-06 | Up |
| Atf4 | 54340,73496 | -0,654055518 | 0,156529444 | 4,20E-07 | 6,98E-06 | Down |

| | | | | | | |
|---|---|---|---|---|---|---|
| Grap2 | 44,71983661 | 2,477601856 | 0,491485337 | 1,65E-08 | 3,90E-07 | Up |
| Mkl1 | 9561,340033 | 0,703684011 | 0,121608153 | 1,17E-10 | 4,64E-09 | Up |
| Sep 03 | 1811,656838 | 0,718885943 | 0,188316863 | 1,50E-06 | 2,11E-05 | Up |
| Ttll1 | 2638,371487 | 0,652781368 | 0,089071018 | 4,13E-15 | 4,03E-13 | Up |
| Parvb | 447,153434 | -0,998799697 | 0,338704563 | 9,24E-06 | 0,000103258 | Down |
| 5031439G07Rik | 24803,44948 | 0,863645796 | 0,148995682 | 7,29E-11 | 3,03E-09 | Up |
| Creld2 | 9921,911313 | -0,644562827 | 0,111489198 | 1,19E-10 | 4,71E-09 | Down |
| Pim3 | 6,282194326 | -0,814240338 | 0,927129655 | 0,002297728 | 0,011316554 | Down |
| Odf3b | 398,2750906 | 0,586083392 | 0,145881017 | 1,04E-06 | 1,55E-05 | Up |
| Mapk8ip2 | 138,2445756 | 0,641373682 | 0,230209144 | 7,32E-05 | 0,000615558 | Up |
| Shank3 | 691,8001706 | -1,46330541 | 0,286998796 | 8,63E-10 | 2,79E-08 | Down |
| Prickle1 | 609,2121696 | 1,104831144 | 0,243219996 | 1,34E-08 | 3,25E-07 | Up |
| Mettl7a1 | 3337,095574 | 1,691347466 | 0,237350971 | 1,12E-14 | 9,90E-13 | Up |
| Galnt6 | 13781,09769 | -0,800704163 | 0,096188524 | 1,85E-18 | 3,03E-16 | Down |
| Acvrl1 | 417,3961895 | -1,099472845 | 0,576145842 | 0,000142384 | 0,001081544 | Down |
| Csad | 543,1088425 | 0,743790725 | 0,127698054 | 9,15E-11 | 3,72E-09 | Up |
| Calcoco1 | 1748,419943 | 1,06004916 | 0,16319977 | 2,48E-13 | 1,72E-11 | Up |
| Gpr84 | 315,0824204 | -2,684406062 | 0,303855956 | 1,06E-19 | 2,10E-17 | Down |
| Zfp385a | 4708,771623 | 1,161898589 | 0,157951532 | 3,04E-16 | 3,47E-14 | Up |
| Nlrc3 | 795,6766529 | 0,672793636 | 0,17534609 | 1,63E-06 | 2,27E-05 | Up |
| Coro7 | 8024,278426 | -0,746553237 | 0,138427588 | 1,01E-09 | 3,21E-08 | Down |
| Socs1 | 378,5095863 | -1,541324213 | 0,189027601 | 1,81E-18 | 2,99E-16 | Down |
| Snx29 | 528,914144 | 0,843756753 | 0,177398194 | 1,76E-08 | 4,13E-07 | Up |
| Mkl2 | 9248,303095 | 0,774514023 | 0,124214142 | 7,48E-12 | 3,85E-10 | Up |
| Cebpd | 1386,681711 | -0,684851902 | 0,132714014 | 3,71E-09 | 1,03E-07 | Down |
| Vpreb1 | 21,9782659 | 0,742671032 | 0,540554856 | 0,00121449 | 0,006659378 | Up |
| Rimbp3 | 353,1258945 | 0,93862141 | 0,164595236 | 7,87E-11 | 3,23E-09 | Up |
| Klhl24 | 2207,660239 | 0,971563716 | 0,143429809 | 7,49E-14 | 5,90E-12 | Up |
| Etv5 | 5632,118427 | -1,917104271 | 0,176502563 | 9,71E-29 | 4,77E-26 | Down |
| Gm15651 | 39,44863053 | -0,587429654 | 0,430405405 | 0,00252789 | 0,012232331 | Down |
| St6gal1 | 834,2815111 | -2,634710841 | 0,499340851 | 6,14E-09 | 1,61E-07 | Down |
| Gm15657 | 115,7387431 | 0,721925256 | 0,300715171 | 0,000139295 | 0,001061559 | Up |
| Mylk | 5865,78619 | 0,632742015 | 0,230399962 | 8,63E-05 | 0,000706902 | Up |
| Stfa2l1 | 107,2059199 | -3,817820253 | 0,619144808 | 1,01E-11 | 5,08E-10 | Down |
| Gm5483 | 452,7039256 | -4,956125357 | 0,698379173 | 2,03E-15 | 2,04E-13 | Down |
| Gm5416 | 22,91861969 | -3,359591106 | 1,484096729 | 1,30E-05 | 0,000138559 | Down |
| 2010005H15Rik | 294,9231517 | -4,513978446 | 0,732374878 | 2,33E-12 | 1,35E-10 | Down |
| Stfa1 | 435,2804446 | -3,899059336 | 0,774891759 | 4,70E-09 | 1,27E-07 | Down |
| BC100530 | 158,3176464 | -4,136955888 | 0,85984184 | 8,53E-09 | 2,16E-07 | Down |
| Stfa2 | 66,66232615 | -6,494059633 | 1,294976492 | 3,99E-10 | 1,43E-08 | Down |
| Stfa3 | 23,35549363 | -4,176779124 | 1,6215691 | 3,41E-06 | 4,35E-05 | Down |
| Cd80 | 93,93014878 | -1,872216378 | 0,336057954 | 3,61E-10 | 1,33E-08 | Down |
| Tmem39a | 7637,017843 | -0,676935656 | 0,147310002 | 6,12E-08 | 1,25E-06 | Down |
| Wrb | 2437,685251 | -0,811510784 | 0,116452154 | 5,55E-14 | 4,46E-12 | Down |
| Tulp4 | 2056,861606 | -0,581284414 | 0,145381222 | 1,15E-06 | 1,69E-05 | Down |
| Gm16702 | 9,818284983 | -0,595814275 | 0,623486334 | 0,004232114 | 0,018735623 | Down |
| Abca3 | 75,97703443 | 0,700974166 | 0,298620291 | 0,00017827 | 0,00131057 | Up |
| D330041H03Rik | 116,2073629 | 0,740867341 | 0,285385655 | 7,52E-05 | 0,000629111 | Up |
| Ift140 | 5718,39304 | 0,597230887 | 0,071559212 | 1,15E-18 | 1,96E-16 | Up |
| Tpsab1 | 9,015693058 | -0,732688106 | 1,021797986 | 0,003123493 | 0,014579594 | Down |
| Haghl | 754,6905472 | 0,867603417 | 0,221952591 | 5,68E-07 | 9,05E-06 | Up |

| | | | | | | |
|---|---|---|---|---|---|---|
| Wdr90 | 3890,75529 | 0,816331246 | 0,176086364 | 3,44E-08 | 7,54E-07 | Up |
| Rab11fip3 | 208,3868459 | 1,391318548 | 0,278678723 | 1,23E-09 | 3,84E-08 | Up |
| Crebrf | 888,3180466 | 0,598616995 | 0,222016757 | 0,000120871 | 0,000941947 | Up |
| E230001N04Rik | 814,5500931 | 0,643326705 | 0,105985132 | 2,09E-11 | 1,00E-09 | Up |
| Pim1 | 35421,75936 | -1,12396368 | 0,103273833 | 2,11E-30 | 1,37E-27 | Down |
| Glp1r | 48,06580039 | -0,725367665 | 0,646100111 | 0,002035552 | 0,010209221 | Down |
| Pdxk-ps | 22,29220025 | 0,719197735 | 0,498866475 | 0,001170502 | 0,006457461 | Up |
| Notch3 | 180,1590764 | 0,94878536 | 0,301055389 | 5,75E-06 | 6,84E-05 | Up |
| Cyp4f40 | 4,755755275 | -1,270602525 | 1,750143225 | 0,000483465 | 0,003064148 | Down |
| Cyp4f13 | 251,5320698 | 0,631754769 | 0,185291846 | 9,66E-06 | 0,000106673 | Up |
| Psmb9 | 6708,266725 | 0,924365088 | 0,147408143 | 2,91E-12 | 1,66E-10 | Up |
| Tap1 | 9558,796644 | 0,88773305 | 0,134440218 | 4,43E-13 | 2,89E-11 | Up |
| Psmb8 | 18413,12469 | 0,617401625 | 0,138246285 | 1,27E-07 | 2,41E-06 | Up |
| Tap2 | 24397,07219 | 1,003804831 | 0,156679083 | 6,70E-13 | 4,25E-11 | Up |
| Gm15821 | 68,2584575 | 1,049171816 | 0,605211963 | 0,000240944 | 0,001695283 | Up |
| H2-Ob | 188,701423 | 2,255715865 | 0,366010063 | 3,76E-11 | 1,69E-09 | Up |
| Btnl2 | 13,35834957 | -0,601587006 | 1,080026457 | 0,003867357 | 0,017376867 | Down |
| Msh5 | 205,4298374 | 0,988864044 | 0,397135528 | 3,73E-05 | 0,000345258 | Up |
| Ltb | 2723,481728 | -0,589397638 | 0,321546248 | 0,001109849 | 0,00617958 | Down |
| Tnf | 575,7083511 | -2,205109694 | 0,386979136 | 4,66E-10 | 1,65E-08 | Down |
| Lta | 13,99742915 | -3,338088358 | 0,746476165 | 1,20E-07 | 2,30E-06 | Down |
| Gm20443 | 18,38509011 | 2,705629307 | 0,589486696 | 1,13E-07 | 2,18E-06 | Up |
| H2-T24 | 6998,46961 | -0,645834557 | 0,117609733 | 6,31E-10 | 2,13E-08 | Down |
| Crisp1 | 11,46684021 | -3,771513184 | 1,560416892 | 6,65E-06 | 7,74E-05 | Down |
| Vegfa | 4784,933326 | -0,74745315 | 0,162356971 | 5,03E-08 | 1,05E-06 | Down |
| Klc4 | 2245,837766 | 0,672064794 | 0,133097583 | 6,65E-09 | 1,73E-07 | Up |
| Trem3 | 390,164595 | -2,222165603 | 0,276678269 | 7,76E-17 | 9,61E-15 | Down |
| Apobec2 | 31,47280923 | 2,720704757 | 0,403286846 | 1,25E-12 | 7,55E-11 | Up |
| Stap2 | 1615,345631 | -0,754842821 | 0,100347404 | 1,13E-15 | 1,23E-13 | Down |
| Mpnd | 8047,801588 | -0,773844125 | 0,101027307 | 3,94E-16 | 4,44E-14 | Down |
| Vmac | 530,7294514 | 0,778160153 | 0,131698798 | 5,30E-11 | 2,27E-09 | Up |
| Tnfsf14 | 2354,701557 | -0,817308307 | 0,137840082 | 4,17E-11 | 1,85E-09 | Down |
| C3 | 3832,766568 | -2,466544059 | 0,379313904 | 5,93E-12 | 3,20E-10 | Down |
| Trip10 | 4583,055595 | -0,637160505 | 0,102895061 | 9,71E-12 | 4,91E-10 | Down |
| Adgre1 | 31351,15199 | -1,779265459 | 0,163570229 | 4,71E-29 | 2,55E-26 | Down |
| Lbh | 2209,153057 | 1,237983051 | 0,156961334 | 4,00E-18 | 6,12E-16 | Up |
| Prkd3 | 7,102428607 | -2,577900204 | 1,442928262 | 5,63E-05 | 0,000491016 | Down |
| 1110020A21Rik | 81,41906703 | 0,937878581 | 0,291416159 | 4,83E-06 | 5,86E-05 | Up |
| Rhoq | 4659,410248 | -0,584437291 | 0,151826036 | 2,13E-06 | 2,88E-05 | Down |
| Mcfd2 | 6634,911829 | -0,911394682 | 0,132997758 | 7,02E-14 | 5,56E-12 | Down |
| Cables1 | 446,507358 | 1,335988933 | 0,273223594 | 1,89E-09 | 5,61E-08 | Up |
| Cabyr | 87,62900569 | 0,91997064 | 0,352960594 | 3,34E-05 | 0,000314432 | Up |
| Slc39a6 | 7023,671152 | -0,771455379 | 0,087953076 | 4,28E-20 | 9,13E-18 | Down |
| Celf4 | 147,4746177 | -1,784346401 | 0,803021061 | 4,57E-05 | 0,000410453 | Down |
| Bin1 | 1945,858156 | 0,830920482 | 0,131371492 | 3,60E-12 | 2,00E-10 | Up |
| Cxxc5 | 8768,051505 | 0,694180831 | 0,097710771 | 2,30E-14 | 1,95E-12 | Up |
| Hbegf | 1402,236803 | -1,533042031 | 0,15756167 | 1,28E-24 | 3,92E-22 | Down |
| Tmco6 | 2528,610177 | 0,5999354 | 0,133935637 | 1,25E-07 | 2,38E-06 | Up |
| Spry4 | 227,1957364 | -3,307922105 | 0,264217349 | 3,30E-37 | 3,83E-34 | Down |
| Cd74 | 1766,084264 | -2,523500115 | 0,453338117 | 1,38E-09 | 4,22E-08 | Down |
| Afap1l1 | 2169,408454 | -0,857900765 | 0,138426435 | 6,96E-12 | 3,66E-10 | Down |

| | | | | | | |
|---|---|---|---|---|---|---|
| Tubb6 | 18989,66917 | -0,65377964 | 0,074837524 | 4,61E-20 | 9,71E-18 | Down |
| Mc5r | 1407,292664 | 1,535441863 | 0,348931996 | 3,05E-08 | 6,72E-07 | Up |
| Smad7 | 2163,350201 | 0,69332553 | 0,153208217 | 8,19E-08 | 1,62E-06 | Up |
| Katnal2 | 143,4202902 | 0,596544391 | 0,15769937 | 2,66E-06 | 3,48E-05 | Up |
| St8sia5 | 12,72983078 | -0,614041209 | 0,642468964 | 0,003924925 | 0,017577122 | Down |
| A330094K24Rik | 12,80696012 | 2,533445242 | 0,729188168 | 2,90E-06 | 3,76E-05 | Up |
| Aldh3b1 | 123,1433698 | -0,685447212 | 0,394018294 | 0,000768002 | 0,00452299 | Down |
| Tbc1d10c | 2904,422825 | 0,662186189 | 0,147339099 | 1,01E-07 | 1,96E-06 | Up |
| Pold4 | 1546,038333 | 0,61305616 | 0,204981239 | 4,46E-05 | 0,000402783 | Up |
| Slc29a2 | 2462,474498 | -0,63312259 | 0,178883539 | 5,95E-06 | 7,02E-05 | Down |
| Tsga10ip | 1120,594777 | 0,775210686 | 0,139208792 | 3,67E-10 | 1,34E-08 | Up |
| Atg2a | 6409,884903 | 0,585347793 | 0,082684353 | 2,36E-14 | 1,99E-12 | Up |
| Pygm | 82,27583861 | 1,811489168 | 0,499760216 | 1,06E-06 | 1,58E-05 | Up |
| Gm6445 | 648,3199505 | -0,813285363 | 0,239446078 | 4,61E-06 | 5,66E-05 | Down |
| Slc15a3 | 17,59272379 | -1,849628023 | 1,560286725 | 0,000248811 | 0,001743071 | Down |
| Ms4a3 | 1878,313563 | -2,834996417 | 0,445342643 | 1,40E-11 | 6,86E-10 | Down |
| Pfpl | 22,55191131 | -0,814661926 | 0,70853728 | 0,001487824 | 0,007852941 | Down |
| Psat1 | 15548,68537 | -0,800296312 | 0,161756601 | 8,48E-09 | 2,15E-07 | Down |
| Gna14 | 108,4615706 | 1,592337741 | 0,478529792 | 2,01E-06 | 2,74E-05 | Up |
| Trpm6 | 13,08727511 | -2,532377683 | 1,1639727 | 3,52E-05 | 0,000328556 | Down |
| Mamdc2 | 17,24135101 | -1,038005307 | 0,748566818 | 0,000561802 | 0,003463339 | Down |
| Kank1 | 118,0666616 | -1,824633186 | 0,335275049 | 5,92E-10 | 2,03E-08 | Down |
| Slc1a1 | 77,54256669 | -3,213397836 | 0,907632278 | 1,82E-06 | 2,49E-05 | Down |
| Cd274 | 1913,758696 | -0,640314256 | 0,11048989 | 1,10E-10 | 4,41E-09 | Down |
| Pdcd1lg2 | 271,738588 | -0,728006473 | 0,304451412 | 0,000138321 | 0,001056622 | Down |
| Ankrd22 | 50,41664981 | -1,989377055 | 0,957023557 | 5,89E-05 | 0,000511278 | Down |
| Stambpl1 | 4519,680499 | -0,658007668 | 0,068020574 | 7,86E-24 | 2,28E-21 | Down |
| Ifit1 | 34,68047528 | -1,468494843 | 0,516872086 | 8,30E-06 | 9,42E-05 | Down |
| Frat2 | 3581,670133 | 0,942487684 | 0,15114594 | 3,18E-12 | 1,80E-10 | Up |
| Cpn1 | 84,37942997 | 1,010914787 | 0,230624333 | 4,08E-08 | 8,76E-07 | Up |
| Kazald1 | 80,18685946 | 0,765038481 | 0,344924097 | 0,000177631 | 0,001306462 | Up |
| Pdcd4 | 1960,934207 | 1,242307879 | 0,178132684 | 4,38E-15 | 4,23E-13 | Up |
| Ccdc120 | 1294,409318 | 0,715685067 | 0,094505057 | 7,55E-16 | 8,39E-14 | Up |
| Pim2 | 2848,311864 | -0,955643206 | 0,116756376 | 2,04E-18 | 3,28E-16 | Down |
| Cfp | 7659,197069 | -1,645461837 | 0,292296917 | 1,04E-10 | 4,19E-09 | Down |
| Gm4764 | 15,87551447 | -2,529831483 | 0,612822253 | 5,04E-07 | 8,17E-06 | Down |
| Plac1 | 7,696839202 | -0,644431529 | 0,863576168 | 0,004287433 | 0,018939179 | Down |
| 3830403N18Rik | 1450,474543 | 0,916084188 | 0,234577552 | 4,59E-07 | 7,53E-06 | Up |
| 4933407K13Rik | 225,7767179 | 0,848547827 | 0,415675457 | 0,000191425 | 0,001393174 | Up |
| Gm7153 | 20,38094683 | 0,839080015 | 0,612502892 | 0,000897153 | 0,005169141 | Up |
| Foxo4 | 2174,821687 | 0,73116102 | 0,126932069 | 1,34E-10 | 5,24E-09 | Up |
| Nhsl2 | 4186,278228 | -0,628606117 | 0,156292198 | 8,83E-07 | 1,35E-05 | Down |
| Gm14848 | 8,060289595 | 0,604194654 | 0,784115759 | 0,004926837 | 0,021260089 | Up |
| C77370 | 8,093194279 | 0,653246792 | 0,780174815 | 0,00389853 | 0,017478212 | Up |
| Tlr13 | 26,2013956 | -1,98367742 | 0,891576238 | 4,39E-05 | 0,000397122 | Down |
| Diaph2 | 7804,347423 | -0,616295937 | 0,218312827 | 7,47E-05 | 0,000625685 | Down |
| Phka2 | 5427,570464 | 0,60269821 | 0,075919574 | 3,34E-17 | 4,34E-15 | Up |
| Piga | 3237,514279 | -0,706966905 | 0,094096624 | 1,17E-15 | 1,26E-13 | Down |
| Arhgap6 | 8355,513282 | 1,01302803 | 0,220981312 | 1,62E-08 | 3,86E-07 | Up |

| CM-vs-IL3-CM-72 | | | | | | |
|---|---|---|---|---|---|---|
| | baseMean | log2FoldChange | lfcSE | pvalue | padj | expression |
| Prdm14 | 31,29544993 | -6,342549656 | 2,275629861 | 1,75E-06 | 9,50E-05 | Down |
| Gm29228 | 300,9864628 | -7,082942213 | 1,751233356 | 2,60E-07 | 2,29E-05 | Down |
| Fam178b | 26,82373702 | -0,68817982 | 0,637531943 | 0,000412093 | 0,005784659 | Down |
| Aff3 | 4375,805027 | -0,661327647 | 0,297479706 | 1,53E-05 | 0,000498384 | Down |
| Il1r1 | 3429,923903 | 0,626500468 | 0,143864632 | 4,50E-09 | 8,76E-07 | Up |
| Fhl2 | 24,97946615 | 2,547446486 | 0,582087565 | 2,99E-08 | 4,39E-06 | Up |
| Ica1l | 337,8059389 | 2,190567419 | 0,253975505 | 9,20E-20 | 2,25E-16 | Up |
| Slc11a1 | 107,6252626 | 0,817235621 | 1,138582211 | 0,000400345 | 0,005679412 | Up |
| Wnt10a | 66,15698044 | 0,776335784 | 0,3853391 | 2,44E-05 | 0,000712844 | Up |
| Gm29284 | 18,07175565 | -0,727693729 | 0,804492607 | 0,000472688 | 0,006440487 | Down |
| C130026I21Rik | 154,1012673 | -0,788653478 | 0,406216703 | 3,08E-05 | 0,000842564 | Down |
| Rab17 | 452,0251321 | 1,324710966 | 0,337028859 | 3,23E-07 | 2,64E-05 | Up |
| Per2 | 2201,555183 | 0,580271914 | 0,141578858 | 3,22E-08 | 4,59E-06 | Up |
| Gm6170 | 289,1940133 | -2,578732791 | 1,51165394 | 1,66E-05 | 0,000533367 | Down |
| Chil1 | 11,05716765 | -0,683121491 | 1,234711688 | 0,000613735 | 0,007816351 | Down |
| 5730559C18Rik | 313,4442831 | 1,193137901 | 1,072131998 | 0,000134757 | 0,002498067 | Up |
| Slamf1 | 80,96343175 | -1,507326269 | 0,655508173 | 1,48E-05 | 0,000486188 | Down |
| Armc3 | 30,58335862 | -0,873716921 | 0,601554061 | 0,000125428 | 0,002371643 | Down |
| 9330198N18Rik | 33,82811219 | -0,912392109 | 0,509869543 | 4,88E-05 | 0,001190573 | Down |
| Fam129b | 2719,526399 | 0,7044544 | 0,230576401 | 6,02E-07 | 4,25E-05 | Up |
| C230014O12Rik | 99,5449943 | 1,747882992 | 0,466150174 | 8,50E-07 | 5,36E-05 | Up |
| Dhrs9 | 47,75226993 | 4,473585758 | 1,379583077 | 1,27E-06 | 7,41E-05 | Up |
| Tfpi | 171,5966274 | -0,861839438 | 0,401191243 | 1,58E-05 | 0,000513085 | Down |
| Platr14 | 1385,087422 | -0,815680901 | 0,204858246 | 9,30E-09 | 1,61E-06 | Down |
| Gm14208 | 10,5253319 | -1,641103301 | 1,007831623 | 3,63E-05 | 0,000957161 | Down |
| Ckmt1 | 1388,774942 | 1,544072957 | 0,15497949 | 5,10E-24 | 1,87E-20 | Up |
| Cd93 | 362,579175 | 8,972195343 | 1,750411914 | 5,92E-09 | 1,10E-06 | Up |
| Src | 22,70018317 | 0,793350643 | 0,551424443 | 0,000141447 | 0,002605066 | Up |
| Ppp1r16b | 161,7128126 | 0,639560908 | 0,338571413 | 4,88E-05 | 0,001190573 | Up |
| Gm14393 | 181,2155506 | -1,254894846 | 0,218731411 | 2,71E-10 | 9,66E-08 | Down |
| Gm14439 | 115,6200027 | -1,050659154 | 0,309897426 | 4,42E-07 | 3,35E-05 | Down |
| Gm14387 | 53,8286656 | -1,28550461 | 0,486623777 | 7,27E-06 | 0,000287895 | Down |
| Trim59 | 22,57472393 | 0,785126786 | 0,90479646 | 0,000407607 | 0,005740599 | Up |
| Rab13 | 171,6098962 | 0,929418641 | 0,807556779 | 0,000215392 | 0,003589138 | Up |
| Npr1 | 165,0698062 | 2,918139867 | 0,331811513 | 7,74E-22 | 2,28E-18 | Up |
| Col11a1 | 20,31507519 | 4,81603409 | 2,069742414 | 4,87E-06 | 0,000209892 | Up |
| Bdh2 | 50,70983699 | 0,625521148 | 0,561242687 | 0,000449635 | 0,006218279 | Up |
| Bank1 | 390,3070533 | 1,269972757 | 0,307504029 | 1,43E-07 | 1,42E-05 | Up |
| Atp8b5 | 128,6979359 | 1,304714821 | 0,288111338 | 4,83E-08 | 6,45E-06 | Up |
| Cd52 | 1011,541979 | -0,616959292 | 0,281152627 | 2,10E-05 | 0,00063181 | Down |
| Zfp980 | 377,3244265 | 0,587369162 | 0,254848245 | 1,84E-05 | 0,000571296 | Up |
| Casz1 | 137,4757495 | -0,590785672 | 0,35907901 | 0,000123383 | 0,002342846 | Down |
| Samd11 | 115,4863289 | 0,897879351 | 0,486777523 | 4,15E-05 | 0,001052144 | Up |
| Actr3b | 14,58669815 | 4,922454562 | 2,505583451 | 7,42E-06 | 0,000290769 | Up |
| Slc5a6 | 1836,307114 | 0,593795845 | 0,136627654 | 8,40E-09 | 1,47E-06 | Up |
| Ccng2 | 1329,31746 | -0,682587048 | 0,279287867 | 6,58E-06 | 0,000265539 | Down |
| Tmem119 | 172,4539261 | 0,678180186 | 0,220779431 | 6,44E-07 | 4,47E-05 | Up |

| | | | | | | |
|---|---|---|---|---|---|---|
| Acacb | 858,2365049 | 0,654122254 | 0,212531873 | 7,44E-07 | 4,86E-05 | Up |
| Nos1 | 525,2454348 | 1,660828391 | 0,23646478 | 3,13E-13 | 2,56E-10 | Up |
| Wbscr27 | 680,0789105 | -0,621671832 | 0,204801729 | 1,17E-06 | 6,91E-05 | Down |
| Gm15753 | 62,44242772 | -0,740052127 | 0,411576918 | 5,11E-05 | 0,001235231 | Down |
| Prkar1b | 85,97825731 | 1,017429012 | 0,249493262 | 2,84E-08 | 4,25E-06 | Up |
| Flt1 | 206,4400117 | 1,18122599 | 0,234302235 | 2,99E-09 | 6,47E-07 | Up |
| Mtus2 | 474,9184921 | -5,403840558 | 0,970647982 | 3,06E-09 | 6,52E-07 | Down |
| 8430423G03Rik | 88,71111414 | 0,626626584 | 0,347235785 | 6,65E-05 | 0,001504618 | Up |
| Flnc | 18,83963871 | -3,337716463 | 0,599911513 | 5,33E-11 | 2,30E-08 | Down |
| Cpa5 | 32,44097674 | -0,759553013 | 0,466534442 | 8,48E-05 | 0,00177955 | Down |
| Dennd2a | 730,3801737 | 1,470627964 | 0,266565448 | 2,18E-09 | 5,01E-07 | Up |
| 8030453O22Rik | 369,0514289 | 1,534149302 | 0,241383296 | 2,87E-11 | 1,45E-08 | Up |
| Trbv12-2 | 42,5808743 | -1,042710679 | 0,366794348 | 2,49E-06 | 0,000125871 | Down |
| Trbv13-2 | 116,371446 | -1,304434983 | 0,224590951 | 2,76E-10 | 9,66E-08 | Down |
| Ccser1 | 22,01476401 | 1,978578769 | 1,225220731 | 2,64E-05 | 0,000749826 | Up |
| Wdr54 | 120,5076728 | 0,962371928 | 0,583636769 | 7,12E-05 | 0,001568221 | Up |
| Frmd4b | 176,0736522 | 0,960144852 | 0,448946038 | 1,79E-05 | 0,000559412 | Up |
| Mitf | 799,4542622 | -0,583345029 | 0,202139415 | 3,00E-06 | 0,000144926 | Down |
| Il5ra | 247,2943963 | -0,815444396 | 0,489386435 | 7,20E-05 | 0,001576519 | Down |
| Ret | 239,5945892 | -0,627253124 | 0,324373561 | 4,48E-05 | 0,001116117 | Down |
| B4galnt3 | 240,1792339 | -0,774586025 | 0,445835334 | 5,97E-05 | 0,001387145 | Down |
| Gm26770 | 159,7417486 | -0,604406981 | 0,230845412 | 5,88E-06 | 0,000242652 | Down |
| Rhno1 | 8719,928351 | -0,63575658 | 0,11460133 | 7,26E-12 | 4,64E-09 | Down |
| Sbk2 | 129,9005395 | -0,768182099 | 0,283253735 | 2,04E-06 | 0,00010594 | Down |
| Vmn1r65 | 32,69716163 | -0,888212674 | 0,54122144 | 7,48E-05 | 0,001618949 | Down |
| Gm44685 | 125,6007902 | -0,593691367 | 0,274597827 | 2,70E-05 | 0,000759518 | Down |
| Zbtb32 | 139,278689 | 0,748523898 | 0,295572204 | 4,06E-06 | 0,000181282 | Up |
| Siglecg | 42,41160238 | 1,215301163 | 0,318343079 | 2,79E-07 | 2,37E-05 | Up |
| Zfp658 | 25,82002918 | 1,273183983 | 1,130855703 | 0,000116448 | 0,002237165 | Up |
| Adamts17 | 142,2581383 | -0,628919471 | 0,392046926 | 0,000119451 | 0,002278462 | Down |
| Mfge8 | 989,6536238 | 1,119574627 | 0,166142657 | 2,50E-13 | 2,29E-10 | Up |
| Usp17ld | 816,2852816 | 0,604624943 | 0,274582131 | 2,22E-05 | 0,000660467 | Up |
| Trim30a | 2281,284289 | -0,629538108 | 0,163261858 | 4,06E-08 | 5,63E-06 | Down |
| Trim30d | 91,68218026 | -1,946624233 | 0,33454727 | 2,51E-10 | 9,46E-08 | Down |
| Lat | 308,9352463 | -0,756600328 | 0,289795434 | 3,01E-06 | 0,000144926 | Down |
| 1500002F19Rik | 43,92224363 | 0,605963242 | 0,489357572 | 0,000352094 | 0,005128575 | Up |
| Tnfrsf23 | 77,78628846 | 1,857055388 | 0,326399983 | 7,10E-10 | 2,09E-07 | Up |
| B020031H02Rik | 68,36503267 | -7,166485567 | 2,381328328 | 8,88E-07 | 5,53E-05 | Down |
| Gm29243 | 387,7691212 | 0,812253986 | 0,213095893 | 2,04E-08 | 3,27E-06 | Up |
| Vegfc | 2024,754752 | -0,594039602 | 0,179519142 | 5,90E-07 | 4,19E-05 | Down |
| Drc7 | 87,03787323 | -0,839781452 | 0,502026334 | 6,97E-05 | 0,001552744 | Down |
| Cmtm3 | 208,2524223 | 1,014606397 | 0,199778595 | 3,51E-10 | 1,17E-07 | Up |
| Pard3 | 118,9275553 | -0,593846275 | 0,341778152 | 9,37E-05 | 0,001905506 | Down |
| RP23-134M7,2 | 102,1844425 | -0,663585604 | 0,326713565 | 2,85E-05 | 0,000791913 | Down |
| Pknox2 | 56,51895352 | 2,172531666 | 0,773362143 | 3,25E-06 | 0,000152596 | Up |
| Stra6 | 22,08416683 | 0,83945387 | 1,241314422 | 0,000393116 | 0,005603899 | Up |
| Cd276 | 489,4175603 | 1,091017019 | 0,353264191 | 1,40E-06 | 7,98E-05 | Up |
| Celf6 | 135,8293714 | 1,36085044 | 0,213576421 | 1,90E-11 | 1,12E-08 | Up |
| Tle3 | 102,443084 | -1,279812219 | 0,488068487 | 7,55E-06 | 0,000292108 | Down |
| Pls1 | 3170,196103 | -0,884463703 | 0,260584373 | 1,73E-07 | 1,63E-05 | Down |
| 1700080E11Rik | 70,29486181 | 1,102796494 | 0,280841919 | 9,92E-08 | 1,11E-05 | Up |

| | | | | | | |
|---|---|---|---|---|---|---|
| Cish | 14896,70043 | 0,588944223 | 0,173299208 | 4,55E-07 | 3,41E-05 | Up |
| Bsn | 1393,889029 | 0,641458919 | 0,107553036 | 5,27E-13 | 4,08E-10 | Up |
| Gm42762 | 74,11714831 | -0,689637744 | 0,4169892 | 8,74E-05 | 0,001821297 | Down |
| Ltf | 278,6434042 | -1,454239378 | 0,409940341 | 1,18E-06 | 6,93E-05 | Down |
| Arfgef3 | 2395,371319 | -0,749741391 | 0,165264864 | 5,24E-10 | 1,60E-07 | Down |
| Il22ra2 | 15,26979953 | 6,687886753 | 1,477909212 | 1,23E-07 | 1,28E-05 | Up |
| Aldh8a1 | 263,4931932 | 5,588880849 | 1,034463293 | 7,11E-09 | 1,28E-06 | Up |
| Enpp1 | 21,06440165 | -2,117125148 | 0,595140871 | 8,25E-07 | 5,25E-05 | Down |
| Slc19a1 | 5856,627741 | 0,665381938 | 0,126736215 | 2,27E-11 | 1,28E-08 | Up |
| Elane | 81,15160536 | -0,887334436 | 0,560941619 | 8,81E-05 | 0,001829351 | Down |
| Reep6 | 194,4615902 | 1,121381331 | 0,180087777 | 4,99E-12 | 3,34E-09 | Up |
| Syn3 | 40,9513894 | -1,298763703 | 0,492087906 | 7,41E-06 | 0,000290769 | Down |
| Anks1b | 52,58431522 | -2,07543123 | 0,341340279 | 2,97E-11 | 1,45E-08 | Down |
| Mgat4c | 92,55360561 | 2,514247245 | 0,334085578 | 1,11E-16 | 1,81E-13 | Up |
| Tph2 | 11,66720527 | 1,973193178 | 0,836631867 | 8,43E-06 | 0,000316957 | Up |
| 1700025F24Rik | 51,88385947 | 0,718562459 | 0,583721077 | 0,000273989 | 0,004325257 | Up |
| Mettl7b | 11,56976992 | 4,096607485 | 2,471792618 | 1,39E-05 | 0,000465815 | Up |
| Gck | 66,43713324 | 0,607502099 | 0,327817513 | 6,27E-05 | 0,001438953 | Up |
| Tgtp2 | 68,61877603 | -1,939616992 | 0,305505851 | 1,07E-11 | 6,57E-09 | Down |
| Grap | 708,8283831 | 1,316086224 | 0,173403248 | 5,52E-15 | 7,38E-12 | Up |
| Mgl2 | 17,80481755 | 2,227906126 | 2,905864344 | 9,01E-05 | 0,001857651 | Up |
| Rph3al | 784,6910796 | -0,831198908 | 0,39794357 | 1,89E-05 | 0,000582449 | Down |
| Gm10392 | 35,00273646 | -0,699365826 | 0,519312457 | 0,000210115 | 0,003536235 | Down |
| Epx | 30,09912387 | -4,182617791 | 1,411918029 | 1,91E-06 | 0,000100806 | Down |
| Nr1d1 | 398,4579192 | 0,893969878 | 0,294373586 | 7,33E-07 | 4,81E-05 | Up |
| Plekhh3 | 34,01354151 | 3,367622538 | 1,192517927 | 2,04E-06 | 0,00010594 | Up |
| Ccdc103 | 19,73196061 | -0,617304291 | 0,537681976 | 0,000425604 | 0,00594592 | Down |
| Cd300a | 305,8562464 | 0,741093331 | 0,340298522 | 1,45E-05 | 0,000481027 | Up |
| Cd300e | 351,0290318 | 0,591700683 | 0,213943645 | 4,08E-06 | 0,000181648 | Up |
| Evpl | 79,20208204 | -0,645404781 | 0,436093788 | 0,000162882 | 0,002880712 | Down |
| Engase | 48,20369037 | 1,347973684 | 0,981119582 | 7,65E-05 | 0,00164218 | Up |
| Zfp750 | 44,7215211 | -0,78350131 | 0,446266307 | 5,60E-05 | 0,001324543 | Down |
| Grhl1 | 381,2159414 | 0,736645473 | 0,399148795 | 4,41E-05 | 0,001105985 | Up |
| Arl4a | 2052,309251 | -0,952686823 | 0,087569255 | 2,22E-30 | 1,47E-26 | Down |
| Rtn1 | 324,6254748 | 1,122165171 | 0,159230605 | 3,75E-14 | 4,23E-11 | Up |
| 2310015A10Rik | 599,3068281 | -0,617547898 | 0,114434776 | 2,54E-11 | 1,34E-08 | Down |
| Ifi27l2a | 191,4777903 | -1,670890552 | 0,430294942 | 6,41E-07 | 4,46E-05 | Down |
| Trmt61a | 4466,011243 | 0,758119168 | 0,140978361 | 4,42E-12 | 3,09E-09 | Up |
| Gpr132 | 72,34824397 | -0,921420066 | 0,472935593 | 3,07E-05 | 0,000842564 | Down |
| Akr1c14 | 101,2916035 | 4,941613023 | 1,165154558 | 2,19E-07 | 1,98E-05 | Up |
| Akr1c18 | 358,3630148 | 0,96810517 | 0,868751451 | 0,000208592 | 0,003531889 | Up |
| F13a1 | 33,96566027 | 2,179702087 | 2,808824281 | 9,46E-05 | 0,001914197 | Up |
| Hk3 | 26800,88562 | -0,61526364 | 0,243934756 | 7,38E-06 | 0,000290769 | Down |
| 4933425L06Rik | 18,09099876 | 2,101419053 | 0,571556412 | 6,49E-07 | 4,47E-05 | Up |
| Rnf180 | 15,47976487 | 4,088937461 | 0,727132486 | 3,24E-10 | 1,11E-07 | Up |
| Gm10248 | 75,04419213 | -0,915653605 | 0,369824457 | 5,74E-06 | 0,000237472 | Down |
| Chdh | 23,90098117 | 2,394746291 | 0,887949375 | 3,22E-06 | 0,00015173 | Up |
| Stab1 | 573,217639 | -0,749876863 | 0,588131718 | 0,000231907 | 0,003808195 | Down |
| Ccnb1ip1 | 3382,037082 | 1,203979884 | 0,280951921 | 5,94E-08 | 7,66E-06 | Up |
| Ripk3 | 124,2185648 | 0,739524939 | 0,525786464 | 0,000166206 | 0,002934809 | Up |
| Mcpt8 | 2841,911022 | -0,598121967 | 0,203390678 | 2,10E-06 | 0,000108649 | Down |

| | | | | | | |
|---|---|---|---|---|---|---|
| Gzmb | 5878,048741 | -1,585186481 | 0,378737038 | 2,91E-07 | 2,44E-05 | Down |
| Olfm4 | 16,80107504 | -0,76450667 | 1,321830752 | 0,000501931 | 0,006724591 | Down |
| Farp1 | 33,18368729 | -2,170048009 | 0,424497035 | 3,87E-09 | 7,80E-07 | Down |
| Adm2 | 23,83837331 | 1,376793623 | 1,819576522 | 0,000175557 | 0,003071613 | Up |
| Mettl7a1 | 3337,095574 | -1,011662712 | 0,292910239 | 2,84E-07 | 2,40E-05 | Down |
| Krt87 | 31,74655852 | 1,008439528 | 0,70110031 | 0,00011287 | 0,002182707 | Up |
| Calcoco1 | 1748,419943 | -0,629600967 | 0,181688803 | 2,06E-07 | 1,88E-05 | Down |
| Myh11 | 103,5531616 | -1,165109903 | 0,988400145 | 0,000131653 | 0,00246214 | Down |
| Klhl24 | 2207,660239 | -0,624685917 | 0,157614235 | 2,73E-08 | 4,18E-06 | Down |
| Etv5 | 5632,118427 | 1,320074453 | 0,176529926 | 1,26E-14 | 1,54E-11 | Up |
| Gm15694 | 57,10517763 | 0,814230086 | 0,400413798 | 2,27E-05 | 0,000673306 | Up |
| Gm15658 | 25,83763082 | -0,694652074 | 0,531123786 | 0,000235538 | 0,003859201 | Down |
| Stfa2 | 66,66232615 | -3,319930439 | 2,371166931 | 2,48E-05 | 0,000719753 | Down |
| Brwd1 | 11,88811841 | 1,124517837 | 1,609348697 | 0,000247268 | 0,003989124 | Up |
| Fam3b | 42,49105394 | -2,357334028 | 0,742977474 | 1,34E-06 | 7,70E-05 | Down |
| Tmem181a | 243,6252396 | -0,608133739 | 0,246120515 | 9,23E-06 | 0,000336346 | Down |
| Igfals | 346,6645581 | 1,848450031 | 0,227474028 | 3,77E-17 | 6,93E-14 | Up |
| Syngap1 | 324,9439682 | 2,462326133 | 0,763353697 | 1,08E-06 | 6,52E-05 | Up |
| 1700097N02Rik | 188,4621016 | 4,520454863 | 0,39950301 | 3,00E-30 | 1,47E-26 | Up |
| Dnah8 | 445,878478 | 4,067620612 | 0,287633239 | 2,69E-46 | 3,95E-42 | Up |
| Glp1r | 48,06580039 | 0,586550644 | 0,741624411 | 0,000819287 | 0,009702711 | Up |
| H2-M3 | 532,7257739 | -0,581390384 | 0,302119167 | 5,89E-05 | 0,001375609 | Down |
| Trem1 | 776,7634029 | 1,492133467 | 0,265029138 | 1,35E-09 | 3,55E-07 | Up |
| Xdh | 28,78667226 | 2,635985021 | 0,661698565 | 1,14E-07 | 1,21E-05 | Up |
| Dync2li1 | 41,62070145 | 4,192308504 | 0,971257641 | 1,01E-07 | 1,12E-05 | Up |
| Cystm1 | 190,5066499 | 0,867733972 | 0,508605908 | 6,26E-05 | 0,001438953 | Up |
| Cd14 | 159,6812712 | 1,720584167 | 0,242513356 | 1,62E-13 | 1,58E-10 | Up |
| Spry4 | 227,1957364 | 1,948319745 | 0,361822059 | 2,50E-09 | 5,56E-07 | Up |
| Mc5r | 1407,292664 | -1,006694883 | 0,441893853 | 1,26E-05 | 0,000427251 | Down |
| Slc29a2 | 2462,474498 | 0,726364572 | 0,215959123 | 1,54E-07 | 1,50E-05 | Up |
| Rab3il1 | 21,18357849 | 1,3149572 | 0,72061134 | 3,96E-05 | 0,001015343 | Up |
| Ccdc86 | 8287,309118 | 0,595676295 | 0,146068307 | 2,66E-08 | 4,11E-06 | Up |
| Psat1 | 15548,68537 | 1,194036585 | 0,158881408 | 4,10E-15 | 6,03E-12 | Up |
| Gna14 | 108,4615706 | -0,595255713 | 0,489145174 | 0,000384115 | 0,005505122 | Down |
| Emx2os | 20,80782657 | 6,672331761 | 1,233414992 | 6,96E-09 | 1,26E-06 | Up |
| Emx2 | 40,39203557 | 7,877365626 | 1,596166713 | 2,49E-08 | 3,89E-06 | Up |
| Gata1 | 227,7361185 | -1,255692184 | 0,834757367 | 7,30E-05 | 0,001589388 | Down |
| Glod5 | 22,97971437 | 1,811857152 | 0,539514703 | 1,82E-06 | 9,74E-05 | Up |
| Gria3 | 443,711147 | 1,068278577 | 0,910758647 | 0,000158841 | 0,002840007 | Up |
| 3830403N18Rik | 1450,474543 | -1,507454897 | 0,209616999 | 1,22E-13 | 1,29E-10 | Down |
| Nap1l3 | 2142,146348 | 0,989445123 | 0,181019432 | 3,44E-11 | 1,63E-08 | Up |
| Nxf7 | 20,08543614 | -1,441055087 | 0,94868867 | 5,50E-05 | 0,00131012 | Down |
| Trpc5os | 22,13987163 | 6,930936035 | 1,506680012 | 9,58E-08 | 1,09E-05 | Up |

| IL3-vs-IL3-CM-12 | | | | | | |
|---|---|---|---|---|---|---|
| | baseMean | log2FoldChange | lfcSE | pvalue | padj | expression |
| Aff3 | 4375,805027 | -0,753147125 | 0,261043575 | 1,01E-05 | 0,000367977 | Down |
| Il1rl1 | 109,3455273 | -0,599848712 | 0,26327065 | 7,59E-05 | 0,001851375 | Down |
| Ino80dos | 114,8376389 | -1,223228699 | 0,248933235 | 1,44E-09 | 2,04E-07 | Down |
| Klf7 | 1705,119904 | 0,733420846 | 0,122160081 | 2,13E-11 | 5,34E-09 | Up |
| Kansl1l | 2153,789982 | -0,690753112 | 0,141378333 | 1,09E-08 | 1,10E-06 | Down |
| Glrp1 | 2495,064469 | 0,597516328 | 0,239833454 | 4,80E-05 | 0,001313741 | Up |
| Ramp1 | 6132,934336 | -0,82382004 | 0,169759805 | 6,94E-09 | 7,81E-07 | Down |
| Pdcd1 | 212,3635005 | -1,24453777 | 0,853889332 | 6,68E-05 | 0,001678016 | Down |
| Rgs2 | 4394,600716 | -0,81441586 | 0,116587519 | 2,30E-14 | 1,55E-11 | Down |
| BC034090 | 128,6157828 | 0,839388655 | 0,256951091 | 2,63E-06 | 0,000118162 | Up |
| Nme7 | 497,7444183 | -0,646173336 | 0,217063413 | 1,21E-05 | 0,000423687 | Down |
| Spta1 | 352,3317916 | -1,96983131 | 0,32349585 | 3,68E-11 | 8,60E-09 | Down |
| Gm13292 | 42,42987693 | 1,823197625 | 0,986257815 | 1,58E-05 | 0,000512473 | Up |
| Slc34a3 | 164,5404617 | -2,877011696 | 0,464618989 | 1,13E-11 | 3,27E-09 | Down |
| Ak8 | 47,89398858 | -0,772352776 | 0,516028979 | 0,000159876 | 0,003321314 | Down |
| Gm13431 | 129,5364897 | 0,649471305 | 0,234505944 | 2,02E-05 | 0,000628403 | Up |
| Cd82 | 3036,908916 | 0,891709483 | 0,219005564 | 1,47E-07 | 1,05E-05 | Up |
| Macrod2 | 256,2804725 | -0,779237409 | 0,22876742 | 2,21E-06 | 0,000103458 | Down |
| Trib3 | 5837,963377 | 2,002491254 | 0,483007894 | 1,91E-07 | 1,33E-05 | Up |
| Nol4l | 6164,336407 | -0,589185594 | 0,117532007 | 8,56E-09 | 9,29E-07 | Down |
| Tgm2 | 10821,87749 | -1,575725241 | 0,335779423 | 1,02E-08 | 1,05E-06 | Down |
| Ppp1r16b | 161,7128126 | 0,870860422 | 0,27878125 | 3,51E-06 | 0,000148186 | Up |
| Wfdc12 | 53,19585331 | -1,830095392 | 0,372523381 | 8,91E-09 | 9,50E-07 | Down |
| Col9a3 | 253,833036 | -0,642011264 | 0,321633503 | 0,000109894 | 0,002450903 | Down |
| Mgst2 | 3909,610169 | -1,02842286 | 0,207004595 | 1,52E-09 | 2,08E-07 | Down |
| Wwtr1 | 82,48549328 | 2,404666078 | 0,38566935 | 2,17E-11 | 5,34E-09 | Up |
| Siah2 | 5297,132062 | 0,597814554 | 0,092021046 | 1,80E-12 | 6,84E-10 | Up |
| I830077J02Rik | 350,8043149 | 0,580618282 | 0,293754902 | 0,000152473 | 0,003205898 | Up |
| Gstm1 | 2134,720948 | -1,206675953 | 0,452346079 | 6,10E-06 | 0,00023909 | Down |
| Cdh17 | 137,6528549 | 0,889080519 | 1,036085728 | 0,000199114 | 0,003928299 | Up |
| Pdp1 | 2184,299773 | 0,870409926 | 0,245989883 | 1,03E-06 | 5,40E-05 | Up |
| Glipr2 | 6207,664248 | 0,851833374 | 0,249970904 | 1,66E-06 | 8,15E-05 | Up |
| Tnfsf8 | 2093,575493 | 0,764496467 | 0,130961628 | 4,93E-11 | 1,11E-08 | Up |
| Elavl2 | 986,5023764 | 0,627556344 | 0,11676499 | 1,21E-09 | 1,73E-07 | Up |
| Cdkn2c | 6893,065099 | -0,626518213 | 0,091585019 | 1,56E-13 | 7,88E-11 | Down |
| Slc6a9 | 1220,144269 | 1,410920048 | 0,265321867 | 2,59E-10 | 4,43E-08 | Up |
| Extl1 | 477,9996045 | 1,122261349 | 0,737734018 | 7,37E-05 | 0,001812604 | Up |
| Clic4 | 37244,65662 | 0,977692523 | 0,168572982 | 2,00E-11 | 5,16E-09 | Up |
| Slc25a33 | 1909,241325 | 0,586841351 | 0,10192849 | 1,72E-10 | 3,13E-08 | Up |
| Nos3 | 60,47437923 | 1,387546872 | 0,279415752 | 1,49E-09 | 2,06E-07 | Up |
| Plac8 | 44329,63147 | -0,777336686 | 0,152058239 | 2,44E-09 | 3,22E-07 | Down |
| Rfc2 | 25282,33014 | -0,595242964 | 0,067958752 | 5,01E-20 | 2,03E-16 | Down |
| Myl10 | 156,465505 | -2,554768892 | 0,43714013 | 1,73E-10 | 3,13E-08 | Down |
| Tfr2 | 6268,32924 | -0,689180823 | 0,225025462 | 8,22E-06 | 0,000309373 | Down |
| Asns | 30785,17086 | 1,050561445 | 0,163854017 | 3,14E-13 | 1,41E-10 | Up |
| Slc37a3 | 345,3545136 | 0,842531984 | 0,444894689 | 6,34E-05 | 0,001616295 | Up |
| 9430076C15Rik | 3288,824048 | -0,633137969 | 0,186811164 | 4,04E-06 | 0,000168622 | Down |
| Mthfd2 | 15492,30892 | 0,966797169 | 0,114391554 | 1,03E-19 | 3,14E-16 | Up |

| | | | | | | |
|---|---|---|---|---|---|---|
| Rab11fip5 | 155,2319165 | 1,31973294 | 0,540166492 | 9,67E-06 | 0,000355586 | Up |
| Gm26588 | 43,18107596 | 0,645516319 | 0,340519744 | 0,000130147 | 0,002819388 | Up |
| Il5ra | 247,2943963 | -0,803925496 | 0,447105783 | 8,52E-05 | 0,002006409 | Down |
| Cand2 | 1623,639553 | -0,845585179 | 0,261932791 | 2,87E-06 | 0,000126959 | Down |
| Ret | 239,5945892 | -0,648240643 | 0,301381455 | 7,92E-05 | 0,001891956 | Down |
| Slc2a3 | 28352,06815 | 0,690627471 | 0,128727339 | 9,79E-10 | 1,45E-07 | Up |
| Fgf23 | 169,6018244 | 0,870593595 | 0,742978716 | 0,000205616 | 0,004017445 | Up |
| Gm26770 | 159,7417486 | -0,95457721 | 0,229812384 | 8,24E-08 | 6,27E-06 | Down |
| Rhno1 | 8719,928351 | -0,869726211 | 0,107676116 | 4,25E-18 | 6,46E-15 | Down |
| Gm43965 | 93,62503784 | -0,658523883 | 0,377049029 | 0,000159625 | 0,003321314 | Down |
| 2310001H17Rik | 762,8386363 | -0,635036287 | 0,211049167 | 1,16E-05 | 0,000411035 | Down |
| Pla2g4c | 248,4843085 | -1,597052072 | 0,372036264 | 5,64E-08 | 4,51E-06 | Down |
| Ehd2 | 210,8835274 | 1,583204716 | 0,332572819 | 7,75E-09 | 8,53E-07 | Up |
| C5ar1 | 80,99254291 | 0,645900222 | 0,372741654 | 0,000172722 | 0,003510187 | Up |
| Dmwd | 5860,363685 | 0,675221464 | 0,083361974 | 9,05E-18 | 1,22E-14 | Up |
| Relb | 527,0632685 | 1,154177216 | 0,241861033 | 3,01E-09 | 3,85E-07 | Up |
| Plaur | 2252,858594 | 0,644863873 | 0,164316072 | 6,82E-07 | 3,79E-05 | Up |
| Zfp36 | 1356,414366 | 0,770702776 | 0,199229824 | 5,16E-07 | 3,03E-05 | Up |
| Siglecf | 2157,366372 | -1,066257217 | 0,405066865 | 7,75E-06 | 0,000296124 | Down |
| Gm44832 | 38,63610838 | 1,487096628 | 0,412810567 | 4,95E-07 | 2,97E-05 | Up |
| Atf5 | 8887,322386 | 0,852507962 | 0,231486952 | 6,94E-07 | 3,83E-05 | Up |
| Kcnj14 | 93,30567782 | 0,608297623 | 0,39985833 | 0,000287921 | 0,005230349 | Up |
| Gm44509 | 276,7010006 | -0,607621235 | 0,204747316 | 1,44E-05 | 0,000472695 | Down |
| Akap13 | 34518,55022 | -0,60667384 | 0,129762147 | 3,87E-08 | 3,25E-06 | Down |
| Abhd17c | 18929,1166 | 0,865508816 | 0,126168135 | 4,19E-14 | 2,55E-11 | Up |
| Arnt2 | 849,7298932 | 1,270441359 | 0,479188256 | 6,12E-06 | 0,000239343 | Up |
| Usp17ld | 816,2852816 | 1,513254251 | 0,191217269 | 1,48E-17 | 1,80E-14 | Up |
| Xylt1 | 3634,687618 | -0,678418643 | 0,15947019 | 1,77E-07 | 1,25E-05 | Down |
| Igsf6 | 1312,914639 | -0,646889736 | 0,237409939 | 2,27E-05 | 0,00069358 | Down |
| Lat | 308,9352463 | 0,858987762 | 0,245763636 | 1,24E-06 | 6,35E-05 | Up |
| Nupr1 | 765,1048752 | 1,854453144 | 0,692001934 | 4,85E-06 | 0,000196726 | Up |
| Irf7 | 224,653155 | 0,76768033 | 0,293510939 | 1,81E-05 | 0,00057627 | Up |
| Cars | 13173,21404 | 0,914815442 | 0,138723646 | 1,92E-13 | 9,33E-11 | Up |
| Eif4ebp1 | 15868,55686 | 0,741062932 | 0,118215212 | 3,99E-12 | 1,35E-09 | Up |
| Gpt2 | 11447,41538 | 1,460610665 | 0,192263241 | 1,21E-16 | 1,23E-13 | Up |
| Herpud1 | 11810,58671 | 0,619753458 | 0,201360328 | 1,02E-05 | 0,000371526 | Up |
| Drc7 | 87,03787323 | -1,080908388 | 0,512497412 | 2,49E-05 | 0,000753445 | Down |
| 1700016A09Rik | 37,75277305 | -1,037005104 | 0,469076397 | 2,14E-05 | 0,000659104 | Down |
| Slc7a5 | 89433,40736 | 0,69150559 | 0,120303726 | 1,17E-10 | 2,29E-08 | Up |
| Cd276 | 489,4175603 | 1,815583152 | 0,250913296 | 1,43E-14 | 1,08E-11 | Up |
| Parp16 | 265,9383329 | -0,622737713 | 0,154239115 | 4,87E-07 | 2,93E-05 | Down |
| Aqp9 | 2236,450141 | 1,816172891 | 0,415516406 | 7,14E-08 | 5,54E-06 | Up |
| Gsta4 | 475,055393 | -0,617989216 | 0,165204738 | 1,40E-06 | 7,05E-05 | Down |
| Bcl2a1b | 1443,482025 | 0,879222245 | 0,272862594 | 2,59E-06 | 0,000117111 | Up |
| Pls1 | 3170,196103 | -1,036048349 | 0,219517622 | 5,05E-09 | 5,84E-07 | Down |
| Gm26767 | 319,5603008 | 1,310911559 | 1,330228338 | 4,15E-05 | 0,001163215 | Up |
| Ky | 967,2072409 | 1,51748516 | 0,145036444 | 1,00E-27 | 1,22E-23 | Up |
| Cish | 14896,70043 | 0,881116255 | 0,157018741 | 9,87E-11 | 2,00E-08 | Up |
| Hemk1 | 5617,126081 | 0,678279246 | 0,158531123 | 1,61E-07 | 1,14E-05 | Up |
| 6430571L13Rik | 340,745674 | 0,940864039 | 0,193742941 | 3,84E-09 | 4,74E-07 | Up |
| Nbeal2 | 37849,0394 | -0,622867097 | 0,125361048 | 9,52E-09 | 1,01E-06 | Down |

| | | | | | | |
|---|---|---|---|---|---|---|
| Gm42762 | 74,11714831 | -0,837531327 | 0,421632185 | 5,37E-05 | 0,001436197 | Down |
| Scml4 | 61,52170318 | -1,893096637 | 0,45295134 | 1,53E-07 | 1,10E-05 | Down |
| Hkdc1 | 599,7391645 | 1,223261985 | 0,252142345 | 1,98E-09 | 2,67E-07 | Up |
| A930033H14Rik | 224,3071726 | 0,9540232 | 0,149300039 | 5,89E-13 | 2,38E-10 | Up |
| Agpat3 | 3052,747228 | 0,811640196 | 0,130179874 | 3,48E-12 | 1,21E-09 | Up |
| Fgf22 | 43,48951939 | 0,643118105 | 0,499952955 | 0,000346287 | 0,005966441 | Up |
| Apc2 | 208,6627662 | -0,62463606 | 0,28343417 | 7,92E-05 | 0,001891956 | Down |
| Bpifc | 1080,100301 | -0,627633351 | 0,857444293 | 0,000509471 | 0,008168341 | Down |
| A230060F14Rik | 81,9251504 | -1,377071447 | 0,38766178 | 5,12E-07 | 3,02E-05 | Down |
| Fbxo7 | 10180,18259 | -0,792010744 | 0,192039385 | 1,85E-07 | 1,30E-05 | Down |
| Syn3 | 40,9513894 | -1,669272599 | 0,614177765 | 4,93E-06 | 0,000198909 | Down |
| Gnptab | 53833,2608 | -0,666115354 | 0,11441019 | 8,53E-11 | 1,82E-08 | Down |
| Gas2l3 | 16653,24652 | -0,717786103 | 0,201946192 | 1,80E-06 | 8,73E-05 | Down |
| Nr1h4 | 81,58779119 | -3,050609049 | 0,568148951 | 6,21E-10 | 9,56E-08 | Down |
| Ntn4 | 72,88246259 | 0,64822717 | 0,293519189 | 7,06E-05 | 0,001754719 | Up |
| Gm15663 | 429,8583652 | 0,71668826 | 0,142425037 | 4,81E-09 | 5,68E-07 | Up |
| Ddit3 | 1598,432583 | 0,792772594 | 0,383600128 | 5,29E-05 | 0,001418293 | Up |
| Il23a | 78,30081049 | 1,458534463 | 0,339069106 | 3,79E-08 | 3,20E-06 | Up |
| Osm | 2398,355282 | 0,844419814 | 0,138109837 | 6,20E-12 | 1,93E-09 | Up |
| Slc1a4 | 6577,979338 | 0,920317578 | 0,147630218 | 1,93E-12 | 7,10E-10 | Up |
| Specc1 | 7087,684873 | -0,747838818 | 0,134721877 | 2,74E-10 | 4,63E-08 | Down |
| Kdm6b | 6599,126144 | 0,968412507 | 0,11990215 | 2,34E-18 | 4,07E-15 | Up |
| Slfn2 | 1045,664031 | 0,948562488 | 0,442053825 | 2,92E-05 | 0,000865357 | Up |
| Ccl4 | 335,2517241 | 1,188214346 | 0,44225406 | 5,90E-06 | 0,000232442 | Up |
| Plekhh3 | 34,01354151 | 1,721705337 | 0,633625999 | 4,86E-06 | 0,000196726 | Up |
| Etv4 | 842,4524741 | 1,236076672 | 0,185737929 | 4,40E-14 | 2,55E-11 | Up |
| Itgb3 | 28796,22896 | -0,678556016 | 0,131460794 | 2,96E-09 | 3,83E-07 | Down |
| Tex2 | 17789,82602 | -0,631863223 | 0,248458368 | 3,69E-05 | 0,001055352 | Down |
| Gm11714 | 341,4437219 | 0,617227287 | 0,669176563 | 0,000573296 | 0,00886148 | Up |
| Smim5 | 8823,774637 | -0,7432208 | 0,135359928 | 3,91E-10 | 6,42E-08 | Down |
| Trim47 | 130,4935779 | -0,984867666 | 0,204538987 | 3,86E-09 | 4,74E-07 | Down |
| Sgsh | 2347,24797 | -0,759743869 | 0,249029418 | 6,44E-06 | 0,000250868 | Down |
| Metrnl | 4887,976873 | 0,740200615 | 0,108841666 | 1,20E-13 | 6,32E-11 | Up |
| Mycn | 8714,752032 | 0,820275829 | 0,218744587 | 6,31E-07 | 3,58E-05 | Up |
| Greb1 | 5300,365898 | -3,005972335 | 0,520942927 | 8,81E-11 | 1,85E-08 | Down |
| Ifrd1 | 11870,87126 | 0,67134634 | 0,158947161 | 2,05E-07 | 1,42E-05 | Up |
| Sipa1l1 | 98257,06086 | -0,706449552 | 0,115532167 | 1,23E-11 | 3,47E-09 | Down |
| Otub2 | 105,5808246 | 0,626219534 | 0,263823136 | 5,51E-05 | 0,001464613 | Up |
| Cdc42bpb | 66,57133961 | -1,381179986 | 1,233942382 | 4,87E-05 | 0,001327676 | Down |
| Gm20658 | 466,5322882 | -1,204532288 | 0,562272788 | 2,01E-05 | 0,00062835 | Down |
| Prr7 | 516,0034562 | 0,60655334 | 0,125506182 | 1,90E-08 | 1,72E-06 | Up |
| Ctla2b | 974,1739352 | -1,04271323 | 0,231862842 | 1,40E-08 | 1,38E-06 | Down |
| Ctla2a | 2671,589779 | -0,880399188 | 0,184592309 | 7,79E-09 | 8,53E-07 | Down |
| Kcnma1 | 68,5027925 | 0,677751485 | 0,97241124 | 0,00033325 | 0,005827327 | Up |
| Gch1 | 3248,718667 | 1,193197852 | 0,175279553 | 1,51E-14 | 1,08E-11 | Up |
| Trac | 148,0237605 | -0,774354475 | 0,470894016 | 0,000123274 | 0,002704238 | Down |
| Pck2 | 22785,40009 | 0,851004977 | 0,229446859 | 6,39E-07 | 3,60E-05 | Up |
| Adcy4 | 351,8680846 | 0,746274061 | 0,156461742 | 1,48E-08 | 1,43E-06 | Up |
| Ripk3 | 124,2185648 | 0,770539075 | 0,348228759 | 4,27E-05 | 0,001183854 | Up |
| Mcpt1 | 70,48400238 | -2,124081819 | 0,533677379 | 3,13E-07 | 2,03E-05 | Down |
| Mcpt2 | 338,2446608 | -0,803233431 | 0,179000574 | 4,01E-08 | 3,34E-06 | Down |

| | | | | | | |
|---|---|---|---|---|---|---|
| Dok2 | 12679,57408 | 0,598429632 | 0,089644107 | 5,49E-13 | 2,30E-10 | Up |
| Bora | 18311,78373 | -0,61022666 | 0,118668179 | 4,25E-09 | 5,11E-07 | Down |
| Sla | 19085,22512 | -0,632514625 | 0,161130369 | 7,08E-07 | 3,89E-05 | Down |
| 9030619P08Rik | 3853,176387 | -1,063569002 | 0,168018932 | 5,07E-13 | 2,20E-10 | Down |
| Il2rb | 2124,240064 | 0,64582713 | 0,163257416 | 6,08E-07 | 3,49E-05 | Up |
| Maff | 559,8860758 | 0,977297908 | 0,168046142 | 1,82E-11 | 4,92E-09 | Up |
| Atf4 | 54340,73496 | 0,899925294 | 0,155920649 | 3,56E-11 | 8,48E-09 | Up |
| Mettl7a1 | 3337,095574 | -1,256066053 | 0,251951529 | 1,02E-09 | 1,48E-07 | Down |
| Acvrl1 | 417,3961895 | 1,054337695 | 0,623919049 | 6,03E-05 | 0,001559613 | Up |
| Rarg | 671,1207684 | 0,601900818 | 0,208783166 | 1,83E-05 | 0,000579774 | Up |
| Gpr84 | 315,0824204 | 1,611906991 | 0,324036292 | 3,23E-09 | 4,09E-07 | Up |
| Socs1 | 378,5095863 | 0,99965092 | 0,188326839 | 2,89E-10 | 4,81E-08 | Up |
| Mkl2 | 9248,303095 | -0,701326373 | 0,125215546 | 2,58E-10 | 4,43E-08 | Down |
| Klhl24 | 2207,660239 | -0,582368514 | 0,148036756 | 7,93E-07 | 4,28E-05 | Down |
| Etv5 | 5632,118427 | 1,217764086 | 0,186907132 | 1,12E-13 | 6,18E-11 | Up |
| Stfa2 | 66,66232615 | 0,598313208 | 0,591763664 | 0,000573849 | 0,00886148 | Up |
| Phf10 | 12548,7236 | 0,616330852 | 0,125379439 | 1,25E-08 | 1,25E-06 | Up |
| Gm16194 | 2143,354735 | -0,643968897 | 0,22806275 | 1,80E-05 | 0,0005757 | Down |
| Psmb9 | 6708,266725 | -0,595630893 | 0,153552902 | 9,30E-07 | 4,89E-05 | Down |
| Tap2 | 24397,07219 | -0,746457744 | 0,157635881 | 1,73E-08 | 1,61E-06 | Down |
| Gm15821 | 68,2584575 | -2,03792104 | 0,518059048 | 3,47E-07 | 2,17E-05 | Down |
| H2-Ob | 188,701423 | -0,813604234 | 0,481853385 | 0,000101181 | 0,002277136 | Down |
| Trem3 | 390,164595 | 1,257362836 | 0,306869842 | 6,29E-08 | 4,96E-06 | Up |
| Cables1 | 446,507358 | -0,807945423 | 0,294274622 | 1,15E-05 | 0,000408334 | Down |
| Bin1 | 1945,858156 | -0,663987233 | 0,133063436 | 7,37E-09 | 8,21E-07 | Down |
| Hbegf | 1402,236803 | 0,761353579 | 0,157264539 | 9,88E-09 | 1,03E-06 | Up |
| Cd14 | 159,6812712 | -0,68795672 | 0,287920521 | 4,09E-05 | 0,001150521 | Down |
| Spry4 | 227,1957364 | 1,976487476 | 0,244064114 | 3,69E-17 | 4,08E-14 | Up |
| Afap1l1 | 2169,408454 | 0,743664365 | 0,139924502 | 1,01E-09 | 1,47E-07 | Up |
| Nars | 63894,03596 | 0,66713468 | 0,098770186 | 2,32E-13 | 1,08E-10 | Up |
| Mc5r | 1407,292664 | -1,874359729 | 0,314387141 | 5,71E-11 | 1,26E-08 | Down |
| Slc1a1 | 77,54256669 | 1,753760647 | 1,205677879 | 2,08E-05 | 0,000645852 | Up |
| Ifit1 | 34,68047528 | 0,745125062 | 0,56070436 | 0,000227771 | 0,004373664 | Up |
| Aldh18a1 | 35196,84936 | 0,641948857 | 0,108056078 | 4,72E-11 | 1,08E-08 | Up |
| Frat2 | 3581,670133 | -0,668137143 | 0,15438342 | 1,37E-07 | 9,93E-06 | Down |

| **IL3-vs-IL3-CM-72** | **baseMean** | **log2FoldChange** | **lfcSE** | **pvalue** | **padj** | **expression** |
|---|---|---|---|---|---|---|
| Pcmtd1 | 1173,701955 | -0,832367821 | 0,124127335 | 3,87E-13 | 8,94E-12 | Down |
| Rrs1 | 14742,20323 | 0,634059233 | 0,11523087 | 3,98E-09 | 4,77E-08 | Up |
| Ncoa2 | 12570,35725 | -0,814271128 | 0,08449971 | 1,25E-23 | 1,21E-21 | Down |
| Paqr8 | 6633,17564 | -1,174669838 | 0,116599388 | 4,06E-26 | 5,15E-24 | Down |
| Tmem14a | 317,0076962 | -0,956391605 | 0,172642227 | 2,60E-10 | 3,80E-09 | Down |
| Gm38224 | 29,05670913 | 0,900203079 | 0,442117895 | 0,000582747 | 0,002531427 | Up |
| Gm26580 | 133,7585049 | -0,713953087 | 0,210603967 | 3,17E-05 | 0,000182895 | Down |
| Gm28439 | 101,9243743 | -0,621496949 | 0,406452805 | 0,006353704 | 0,021169739 | Down |
| 1700001G17Rik | 10,83578211 | -0,952646324 | 0,704274931 | 0,00186613 | 0,007151995 | Down |
| Gm33280 | 28,37326226 | 0,809336908 | 0,34912445 | 0,000449146 | 0,002004516 | Up |

| | | | | | | |
|---|---|---|---|---|---|---|
| Arid5a | 5828,508749 | 0,872526175 | 0,110903535 | 4,75E-17 | 1,93E-15 | Up |
| Fam178b | 26,82373702 | -2,107106655 | 0,570409287 | 5,52E-07 | 4,60E-06 | Down |
| Inpp4a | 8,562766052 | -2,99069248 | 0,956117426 | 5,93E-06 | 3,99E-05 | Down |
| 2010300C02Rik | 49,72523763 | -0,604960091 | 0,343581706 | 0,004866678 | 0,01670588 | Down |
| Aff3 | 4375,805027 | -1,795760628 | 0,232377608 | 3,95E-17 | 1,63E-15 | Down |
| Map4k4 | 1065,531247 | 1,124235837 | 0,278676293 | 3,20E-07 | 2,76E-06 | Up |
| Il1r2 | 11,32800349 | 1,293315211 | 0,736153746 | 0,000341995 | 0,001565291 | Up |
| Il1rl2 | 37,86316982 | 0,925519563 | 0,379686679 | 0,000188544 | 0,000917693 | Up |
| Wdr75 | 13568,03793 | 0,896170848 | 0,094020829 | 1,58E-23 | 1,50E-21 | Up |
| Nabp1 | 2345,527233 | 0,611594143 | 0,130293833 | 3,09E-07 | 2,67E-06 | Up |
| Gm28178 | 10,16725339 | 0,728275749 | 0,588063714 | 0,005198915 | 0,017726398 | Up |
| Nemp2 | 1176,525608 | -0,698977166 | 0,088243487 | 1,63E-16 | 6,26E-15 | Down |
| Stk17b | 4957,975189 | 0,631168538 | 0,145856108 | 1,45E-06 | 1,10E-05 | Up |
| Coq10b | 886,827566 | 0,779100757 | 0,135621468 | 2,90E-10 | 4,22E-09 | Up |
| Hspd1 | 145235,0668 | 0,705364777 | 0,099767122 | 9,92E-14 | 2,52E-12 | Up |
| Rftn2 | 532,1447096 | -0,907930667 | 0,126981337 | 8,85E-15 | 2,69E-13 | Down |
| Mars2 | 6118,293221 | 0,738298823 | 0,101414032 | 1,58E-14 | 4,54E-13 | Up |
| Plcl1 | 6,102168433 | 0,855023755 | 1,054141694 | 0,005012466 | 0,017154732 | Up |
| Orc2 | 9910,453522 | 0,813485461 | 0,073549875 | 4,35E-30 | 8,44E-28 | Up |
| Nop58 | 36117,51578 | 0,734940331 | 0,081611026 | 1,08E-20 | 7,18E-19 | Up |
| Ica1l | 337,8059389 | 0,59681905 | 0,248086892 | 0,001315344 | 0,005244721 | Up |
| Ino80d | 5951,488508 | -0,588472387 | 0,107241334 | 5,91E-09 | 6,93E-08 | Down |
| Ino80dos | 114,8376389 | -1,474110616 | 0,231124837 | 1,08E-12 | 2,32E-11 | Down |
| Gm20342 | 172,0280904 | -0,9906341 | 0,313617102 | 1,50E-05 | 9,31E-05 | Down |
| Fastkd2 | 5356,931692 | 0,707995519 | 0,098799356 | 4,84E-14 | 1,29E-12 | Up |
| Klf7 | 1705,119904 | 0,699475321 | 0,118486015 | 2,30E-10 | 3,38E-09 | Up |
| Unc80 | 7,647501364 | 0,809017706 | 0,779534306 | 0,004725874 | 0,016296054 | Up |
| Kansl1l | 2153,789982 | -1,156360535 | 0,139599593 | 6,32E-19 | 3,28E-17 | Down |
| Gm38162 | 3,734625736 | -0,621795779 | 0,784553039 | 0,011689545 | 0,035656334 | Down |
| Cxcr2 | 77,55932955 | 1,002154017 | 0,513673109 | 0,000477205 | 0,002118824 | Up |
| Slc11a1 | 107,6252626 | 3,943013758 | 0,88680504 | 1,63E-07 | 1,47E-06 | Up |
| Stk36 | 69,79589087 | -1,35876886 | 0,263456685 | 1,23E-09 | 1,60E-08 | Down |
| Wnt10a | 66,15698044 | -0,600015779 | 0,284370177 | 0,002541876 | 0,009427242 | Down |
| Tuba4a | 73308,85271 | 0,768989919 | 0,086124196 | 1,53E-20 | 9,97E-19 | Up |
| Resp18 | 25,97294118 | 0,70426876 | 0,380977875 | 0,002286568 | 0,008584972 | Up |
| Slc4a3 | 706,4850726 | 1,162515107 | 0,22907989 | 2,05E-09 | 2,58E-08 | Up |
| Farsb | 15628,87625 | 0,676189154 | 0,069940065 | 3,61E-23 | 3,33E-21 | Up |
| Dock10 | 33536,79182 | -0,767913406 | 0,158793869 | 4,50E-08 | 4,52E-07 | Down |
| Gm15433 | 27,06492675 | 2,857546194 | 0,706086748 | 3,94E-07 | 3,35E-06 | Up |
| A530040E14Rik | 79,45801135 | -0,661418444 | 0,287753064 | 0,001115928 | 0,004522903 | Down |
| Gm16341 | 10,38379541 | 0,903914361 | 0,64700862 | 0,002038203 | 0,007743221 | Up |
| Ncl | 316680,2043 | 0,625130311 | 0,098818973 | 3,01E-11 | 5,08E-10 | Up |
| Gm37485 | 18,20905006 | 1,36513389 | 0,583212071 | 6,70E-05 | 0,000359466 | Up |
| Sag | 31,77199352 | 0,819117698 | 0,352264475 | 0,000419344 | 0,001882049 | Up |
| Ramp1 | 6132,934336 | -0,862922758 | 0,165653478 | 3,00E-09 | 3,65E-08 | Down |
| Gm26683 | 25,34783889 | 0,60588274 | 0,447159483 | 0,008716286 | 0,02774676 | Up |
| Hes6 | 2039,661949 | -0,991138861 | 0,107766143 | 2,09E-22 | 1,67E-20 | Down |
| Per2 | 2201,555183 | 0,827413262 | 0,130744275 | 5,04E-12 | 9,60E-11 | Up |
| 5033417F24Rik | 113,0355414 | -0,624266156 | 0,204072671 | 0,000177803 | 0,000870285 | Down |
| D2hgdh | 1369,304957 | -0,834313633 | 0,092048814 | 2,30E-21 | 1,64E-19 | Down |
| Gm7889 | 93,02605462 | -0,977398441 | 0,206717038 | 1,92E-08 | 2,06E-07 | Down |

| | | | | | | |
|---|---|---|---|---|---|---|
| Pdcd1 | 212,3635005 | -3,573000566 | 0,552012862 | 8,16E-12 | 1,52E-10 | Down |
| D1Ertd622e | 3231,772624 | 0,739568275 | 0,094107741 | 1,83E-16 | 6,98E-15 | Up |
| 1810006J02Rik | 1663,580782 | 0,663307902 | 0,141637821 | 2,23E-07 | 1,97E-06 | Up |
| Phlpp1 | 2041,35478 | -0,823112216 | 0,154441378 | 2,12E-09 | 2,65E-08 | Down |
| Bcl2 | 6392,698425 | 0,644808278 | 0,120647235 | 8,79E-09 | 9,97E-08 | Up |
| Nifk | 20066,37479 | 0,627238609 | 0,085966638 | 3,63E-14 | 9,94E-13 | Up |
| Tfcp2l1 | 8,265693963 | 1,009264764 | 0,793008384 | 0,001814901 | 0,006986451 | Up |
| Tmem185b | 9566,073431 | 1,031153275 | 0,084940656 | 3,02E-36 | 1,08E-33 | Up |
| Ddx18 | 21063,86072 | 0,856995826 | 0,085400441 | 1,54E-25 | 1,87E-23 | Up |
| Slc35f5 | 747,7812834 | -0,763900674 | 0,142078189 | 2,71E-09 | 3,33E-08 | Down |
| Map3k19 | 32,01209998 | -1,390597908 | 0,455349898 | 7,57E-06 | 5,00E-05 | Down |
| Zranb3 | 3159,857756 | -0,860134523 | 0,115007482 | 1,11E-15 | 3,73E-14 | Down |
| Gm6170 | 289,1940133 | -1,571819338 | 1,161772001 | 0,000577739 | 0,002512062 | Down |
| Cxcr4 | 7686,079561 | -1,073021555 | 0,11675392 | 1,88E-22 | 1,52E-20 | Down |
| Gm26686 | 12,38670003 | -0,645121288 | 0,42218688 | 0,005534265 | 0,01870221 | Down |
| Pik3c2b | 2670,127644 | 0,702599439 | 0,142103869 | 4,52E-08 | 4,54E-07 | Up |
| Gm26616 | 28,73629189 | -0,687565903 | 0,33327655 | 0,001621784 | 0,006314928 | Down |
| Lax1 | 1870,023799 | -0,608159499 | 0,107151309 | 1,79E-09 | 2,27E-08 | Down |
| Atp2b4 | 17240,5454 | -0,800633106 | 0,083650695 | 2,71E-23 | 2,51E-21 | Down |
| Btg2 | 1120,205726 | -1,812043751 | 0,157118845 | 2,94E-33 | 7,90E-31 | Down |
| Ppfia4 | 21,39133961 | -0,61881858 | 0,385288077 | 0,005759072 | 0,01937586 | Down |
| Cyb5r1 | 4606,312619 | 0,828999141 | 0,102210466 | 9,79E-18 | 4,40E-16 | Up |
| Timm17a | 13053,33836 | 0,80207682 | 0,09759531 | 5,23E-18 | 2,44E-16 | Up |
| 9230116N13Rik | 101,8740312 | -0,628065002 | 0,206182189 | 0,000180285 | 0,00088078 | Down |
| Gm28556 | 19,47096576 | -0,598861606 | 0,371582924 | 0,006459392 | 0,021478774 | Down |
| Rgs2 | 4394,600716 | -1,075033522 | 0,1165202 | 1,35E-22 | 1,13E-20 | Down |
| Rgs1 | 88,76523314 | -2,203341634 | 0,342543512 | 5,04E-13 | 1,14E-11 | Down |
| 1700025G04Rik | 12748,56337 | -0,582705236 | 0,087662197 | 4,80E-12 | 9,18E-11 | Down |
| Gm7278 | 111,0920879 | 1,05185442 | 0,272104967 | 7,93E-07 | 6,38E-06 | Up |
| Gm28960 | 66,24102451 | 0,760386673 | 0,340388716 | 0,00072058 | 0,003063137 | Up |
| Rnasel | 9275,783115 | -0,798671818 | 0,117026664 | 2,32E-13 | 5,58E-12 | Down |
| BC034090 | 128,6157828 | 1,552721912 | 0,238687275 | 4,36E-13 | 9,97E-12 | Up |
| Soat1 | 16,62526506 | 0,950219677 | 0,601223998 | 0,001248098 | 0,005004998 | Up |
| Ankrd45 | 18,52547862 | 2,287567486 | 0,806779318 | 9,84E-06 | 6,36E-05 | Up |
| Kifap3 | 2702,007157 | -0,581118863 | 0,090762082 | 2,45E-11 | 4,19E-10 | Down |
| Sell | 30736,62868 | -1,080813716 | 0,17586325 | 4,34E-12 | 8,36E-11 | Down |
| Selp | 1150,37954 | -0,808653777 | 0,20410673 | 1,79E-06 | 1,34E-05 | Down |
| Gm16587 | 15,37589313 | 1,551382628 | 1,133023425 | 0,000571585 | 0,00248886 | Up |
| Nme7 | 497,7444183 | -1,382858841 | 0,195237787 | 9,92E-15 | 2,98E-13 | Down |
| Tbx19 | 183,1212648 | -1,284138298 | 0,233776918 | 2,07E-10 | 3,06E-09 | Down |
| Gm15853 | 67,60233372 | -0,716628446 | 0,239556387 | 0,000116037 | 0,000588792 | Down |
| Mgst3 | 1096,111012 | -0,580640782 | 0,185440357 | 0,00018992 | 0,000924144 | Down |
| Gm7694 | 1519,455987 | -0,913719122 | 0,157961506 | 7,80E-11 | 1,24E-09 | Down |
| Nos1ap | 689,4264596 | -0,977036584 | 0,156813659 | 3,38E-12 | 6,61E-11 | Down |
| Dusp12 | 1364,685485 | 0,816391907 | 0,116132984 | 4,61E-14 | 1,23E-12 | Up |
| Fcrlb | 7,982514978 | 3,296709539 | 0,990023007 | 3,82E-06 | 2,67E-05 | Up |
| Fcgr2b | 3341,950652 | 0,679175108 | 0,296050614 | 0,001008118 | 0,004130637 | Up |
| Fcgr4 | 32,96658856 | 2,517392823 | 0,635690672 | 3,37E-07 | 2,90E-06 | Up |
| Fcgr3 | 445,2897942 | 4,00973343 | 0,47697896 | 3,65E-18 | 1,75E-16 | Up |
| Cfap126 | 33,52725716 | -1,555123382 | 0,384557994 | 1,63E-07 | 1,48E-06 | Down |
| Gm9765 | 10,17440408 | -0,842526001 | 0,615707385 | 0,002683115 | 0,009880832 | Down |

| | | | | | | |
|---|---|---|---|---|---|---|
| Itln1 | 144,5123685 | -0,633354888 | 0,406374485 | 0,005670638 | 0,019113544 | Down |
| Cd244 | 2297,6769 | -1,114628649 | 0,161808694 | 2,90E-14 | 8,07E-13 | Down |
| Ly9 | 34,8917493 | -1,892568513 | 0,474813666 | 1,64E-07 | 1,48E-06 | Down |
| Slamf7 | 17,07327356 | -1,956181701 | 0,723191266 | 1,36E-05 | 8,50E-05 | Down |
| A630035G10Rik | 47,50261419 | -0,984929308 | 0,354373429 | 5,41E-05 | 0,000296545 | Down |
| Slamf1 | 80,96343175 | 1,263440555 | 0,489141112 | 4,21E-05 | 0,000236426 | Up |
| Gm37065 | 26,68573509 | 2,6095789 | 1,371066734 | 8,74E-05 | 0,000456114 | Up |
| Igsf8 | 11644,53898 | 1,805151236 | 0,135928837 | 9,84E-43 | 5,45E-40 | Up |
| Kcnj9 | 14,56965211 | 1,004867539 | 0,687488681 | 0,001306359 | 0,005213445 | Up |
| Slamf9 | 17,06430912 | 4,51862226 | 0,87131365 | 4,86E-09 | 5,74E-08 | Up |
| Gm37125 | 34,42922649 | -0,596656168 | 0,284680605 | 0,002677807 | 0,009865301 | Down |
| Vsig8 | 10,39295152 | 0,794256989 | 0,627646724 | 0,003773899 | 0,013364155 | Up |
| Ifi207 | 8,319987667 | 3,040791772 | 1,680163871 | 7,22E-05 | 0,000384845 | Up |
| Ifi204 | 148,3465776 | -1,515862232 | 0,238895925 | 1,27E-12 | 2,70E-11 | Down |
| Ifi203-ps | 17,54352426 | -0,963512294 | 0,455021977 | 0,000371174 | 0,001683651 | Down |
| Mndal | 929,3119738 | -1,339083379 | 0,2015597 | 1,94E-13 | 4,70E-12 | Down |
| Ifi211 | 27,8513269 | -0,865399028 | 0,385684287 | 0,00041156 | 0,001848449 | Down |
| Ifi203 | 624,3994017 | -1,690389637 | 0,224658446 | 2,52E-16 | 9,50E-15 | Down |
| Spta1 | 352,3317916 | -5,494013408 | 0,310782341 | 3,89E-72 | 1,42E-68 | Down |
| Olfr417 | 8,060975743 | -3,348768763 | 1,009578118 | 3,93E-06 | 2,74E-05 | Down |
| Olfr248 | 16,25454946 | -3,644257751 | 0,607111405 | 1,36E-10 | 2,06E-09 | Down |
| Olfr414 | 16,69962429 | -4,766587453 | 0,855812415 | 4,72E-10 | 6,60E-09 | Down |
| H3f3aos | 23,41425315 | -0,733987824 | 0,357404628 | 0,001265593 | 0,005070665 | Down |
| Pycr2 | 11216,90091 | 0,599411808 | 0,089125775 | 2,54E-12 | 5,13E-11 | Up |
| 2210411M09Rik | 9,656999446 | -0,885945852 | 0,661202793 | 0,002383923 | 0,008910193 | Down |
| Ephx1 | 11,28576173 | 0,813901218 | 0,759131664 | 0,00446514 | 0,01549352 | Up |
| Cnih3 | 27,56962464 | 0,810417154 | 0,372667387 | 0,000637457 | 0,002738473 | Up |
| Enah | 45,52905557 | 2,373542132 | 0,769284882 | 5,00E-06 | 3,42E-05 | Up |
| Degs1 | 9673,181637 | 1,310685228 | 0,116358701 | 1,88E-31 | 4,04E-29 | Up |
| Gm37800 | 94,49497482 | 1,266431118 | 0,330777804 | 5,79E-07 | 4,79E-06 | Up |
| Rrp15 | 11326,24982 | 0,691191823 | 0,091359148 | 2,85E-15 | 9,17E-14 | Up |
| Tmem206 | 1860,342544 | 0,663845627 | 0,083759083 | 2,14E-16 | 8,09E-15 | Up |
| Gm38037 | 90,14570145 | -0,606384552 | 0,217936707 | 0,0004564 | 0,002034232 | Down |
| Slc30a1 | 697,9631187 | 0,639030933 | 0,124316832 | 2,73E-08 | 2,85E-07 | Up |
| Hsd11b1 | 61,53172636 | 1,818606637 | 0,515194386 | 9,39E-07 | 7,43E-06 | Up |
| Rpp38 | 1625,662356 | 0,703356629 | 0,098729692 | 6,85E-14 | 1,79E-12 | Up |
| Sec61a2 | 454,7876257 | 0,676987686 | 0,206317063 | 5,99E-05 | 0,000326454 | Up |
| Echdc3 | 48,56900599 | -0,935480891 | 0,283912905 | 1,17E-05 | 7,40E-05 | Down |
| Gm13261 | 115,9308044 | 1,035785228 | 0,294091889 | 3,34E-06 | 2,36E-05 | Up |
| Gm13292 | 42,42987693 | 1,526937264 | 0,879178649 | 0,000250899 | 0,001185491 | Up |
| Il2ra | 1536,200824 | 1,382735306 | 0,490030214 | 1,60E-05 | 9,83E-05 | Up |
| Tmem236 | 6,621621767 | 0,724977226 | 0,718186961 | 0,006668395 | 0,022109337 | Up |
| Mrc1 | 96,22464949 | 1,139693671 | 0,390962578 | 2,10E-05 | 0,000125863 | Up |
| Plxdc2 | 16129,59241 | -0,69651044 | 0,109910979 | 1,58E-11 | 2,81E-10 | Down |
| Skida1 | 318,9769318 | -0,604947109 | 0,206646002 | 0,00030251 | 0,001401075 | Down |
| Armc3 | 30,58335862 | -1,084428315 | 0,450822957 | 0,000114226 | 0,000580734 | Down |
| Gm13375 | 364,7283636 | -0,890859508 | 0,165317958 | 9,11E-10 | 1,20E-08 | Down |
| Pdss1 | 2541,459306 | 0,664613461 | 0,091000568 | 2,58E-14 | 7,20E-13 | Up |
| Il1f9 | 11,44569477 | 2,803015554 | 1,64831937 | 0,000104091 | 0,000533514 | Up |
| Slc34a3 | 164,5404617 | -5,821441343 | 0,477412829 | 1,06E-36 | 3,87E-34 | Down |
| Rnf224 | 1,785951605 | -0,968409489 | 1,495344661 | 0,002730682 | 0,010015609 | Down |

| | | | | | | |
|---|---|---|---|---|---|---|
| Cysrt1 | 3,523306404 | -0,874658716 | 1,233847568 | 0,004585086 | 0,015867507 | Down |
| Ptgds | 5,043273106 | -0,62446206 | 0,818180939 | 0,01159801 | 0,035448176 | Down |
| Nacc2 | 20,53282361 | -0,58658574 | 0,362473094 | 0,006908316 | 0,022788959 | Down |
| Qsox2 | 6293,675937 | 0,845932357 | 0,059541551 | 1,28E-47 | 9,33E-45 | Up |
| Egfl7 | 3346,789809 | 0,823430459 | 0,116538055 | 3,33E-14 | 9,17E-13 | Up |
| Fam69b | 4177,57228 | 0,621855479 | 0,090029812 | 6,21E-13 | 1,39E-11 | Up |
| Lcn4 | 258,9932678 | -1,770827163 | 0,391935355 | 1,74E-08 | 1,88E-07 | Down |
| Slc2a6 | 1861,671792 | 0,738316396 | 0,247725029 | 0,000103951 | 0,000532946 | Up |
| Gm13398 | 9,319686238 | 0,66134825 | 0,590426082 | 0,00806558 | 0,025982295 | Up |
| Sardh | 12,83693174 | -0,922325173 | 0,572030754 | 0,001299773 | 0,005188297 | Down |
| Spaca9 | 236,2926072 | -1,955430394 | 0,203871819 | 2,21E-24 | 2,40E-22 | Down |
| Ak8 | 47,89398858 | -1,758996712 | 0,413597805 | 5,63E-08 | 5,55E-07 | Down |
| Ddx31 | 4878,407981 | 0,657719976 | 0,096545318 | 9,15E-13 | 2,00E-11 | Up |
| Cercam | 1490,254638 | -0,763689374 | 0,181555526 | 8,84E-07 | 7,05E-06 | Down |
| Zer1 | 3262,2588 | -1,042303811 | 0,108429401 | 3,44E-24 | 3,63E-22 | Down |
| Crat | 3788,761325 | -0,680167977 | 0,145000685 | 1,89E-07 | 1,68E-06 | Down |
| Ntmt1 | 4542,77234 | 0,708536844 | 0,090251943 | 2,63E-16 | 9,86E-15 | Up |
| Hmcn2 | 28,39534262 | 1,710231808 | 0,676605795 | 2,52E-05 | 0,000148312 | Up |
| Lcn2 | 1442,891879 | 2,993375524 | 0,670950574 | 1,08E-07 | 1,01E-06 | Up |
| Slc25a25 | 2916,462257 | 0,735936839 | 0,079462767 | 1,00E-21 | 7,51E-20 | Up |
| St6galnac4 | 10520,07696 | 0,862115281 | 0,090542897 | 2,36E-23 | 2,22E-21 | Up |
| Fpgs | 7260,704439 | 0,681888934 | 0,097157555 | 1,77E-13 | 4,30E-12 | Up |
| Sh2d3c | 4594,635707 | -0,784052235 | 0,144080711 | 1,58E-09 | 2,02E-08 | Down |
| Fam129b | 2719,526399 | 0,983385616 | 0,188257413 | 1,37E-09 | 1,77E-08 | Up |
| Gm38299 | 22,19414092 | 1,011039357 | 0,754707867 | 0,001604752 | 0,006264639 | Up |
| Ralgps1 | 728,9853817 | -0,787125489 | 0,116056221 | 3,48E-13 | 8,11E-12 | Down |
| Morn5 | 4,135342961 | -0,616717456 | 0,837779579 | 0,011996188 | 0,036451646 | Down |
| Gm13431 | 129,5364897 | 1,072276504 | 0,216543031 | 4,47E-09 | 5,32E-08 | Up |
| Ptgs1 | 17804,70589 | 0,685061526 | 0,094905923 | 4,06E-14 | 1,10E-12 | Up |
| Nek6 | 9623,426214 | 0,870102439 | 0,086112318 | 6,71E-26 | 8,39E-24 | Up |
| Gm13496 | 34,17031884 | 0,939893525 | 0,362669194 | 0,000114534 | 0,000582138 | Up |
| Gm13502 | 42,7656329 | -0,982609987 | 0,441789094 | 0,000264219 | 0,001239764 | Down |
| Nr4a2 | 1421,172583 | -0,691470159 | 0,138910246 | 4,18E-08 | 4,23E-07 | Down |
| Tanc1 | 179,9517431 | 1,013840721 | 0,601636171 | 0,000818145 | 0,003433127 | Up |
| Galnt3 | 6167,877876 | 0,928631076 | 0,087128758 | 1,23E-28 | 2,00E-26 | Up |
| Dhrs9 | 47,75226993 | 3,918592753 | 1,156924139 | 2,77E-06 | 1,98E-05 | Up |
| Fastkd1 | 2957,576253 | 0,649883748 | 0,077456882 | 5,17E-18 | 2,41E-16 | Up |
| Ccdc173 | 148,0710595 | -0,72167231 | 0,199638131 | 1,33E-05 | 8,33E-05 | Down |
| Itga6 | 7790,840068 | -0,641125647 | 0,134359919 | 1,73E-07 | 1,55E-06 | Down |
| Pdk1 | 4301,739563 | -0,724850613 | 0,118837355 | 5,53E-11 | 8,98E-10 | Down |
| Ttc30b | 97,98092554 | -1,325067393 | 0,213427638 | 3,14E-12 | 6,24E-11 | Down |
| Prkra | 1574,834489 | -0,652001976 | 0,107660913 | 1,34E-10 | 2,04E-09 | Down |
| Plekha3 | 4718,743872 | 0,889739395 | 0,089906045 | 4,61E-25 | 5,40E-23 | Up |
| Cerkl | 138,9130571 | -1,015508845 | 0,191067449 | 7,31E-10 | 9,86E-09 | Down |
| Tfpi | 171,5966274 | 0,694165397 | 0,261150462 | 0,000358341 | 0,001633553 | Up |
| Selenoh | 4614,859462 | 0,587202805 | 0,125451527 | 3,87E-07 | 3,29E-06 | Up |
| Smtnl1 | 6,099863943 | 1,743407233 | 1,289808601 | 0,000481811 | 0,002137195 | Up |
| Timm10 | 2883,452495 | 0,761717416 | 0,119390533 | 6,60E-12 | 1,24E-10 | Up |
| Ptprj | 12041,99226 | -0,721142528 | 0,100724349 | 4,49E-14 | 1,21E-12 | Down |
| Prdm11 | 3830,057943 | -0,684884915 | 0,124028861 | 2,39E-09 | 2,96E-08 | Down |
| Gm13807 | 8,083969344 | 0,884969644 | 0,740566363 | 0,002980496 | 0,010831805 | Up |

| | | | | | | |
|---|---|---|---|---|---|---|
| Cd82 | 3036,908916 | 2,541383982 | 0,202971528 | 1,42E-37 | 5,64E-35 | Up |
| Accs | 1531,17446 | -0,675978556 | 0,120483389 | 1,56E-09 | 2,00E-08 | Down |
| Hsd17b12 | 16725,86354 | 0,654041725 | 0,117674714 | 2,51E-09 | 3,08E-08 | Up |
| Ldlrad3 | 3238,467396 | 0,707687929 | 0,098293553 | 3,80E-14 | 1,03E-12 | Up |
| Nat10 | 18285,97154 | 0,681742982 | 0,095583934 | 7,80E-14 | 2,02E-12 | Up |
| Slc12a6 | 1609,908685 | -0,723737089 | 0,138294767 | 8,39E-09 | 9,58E-08 | Down |
| Spred1 | 8673,256827 | 0,606591038 | 0,102044205 | 3,69E-10 | 5,29E-09 | Up |
| Thbs1 | 1984,565179 | -1,599641065 | 0,90633418 | 0,000216023 | 0,001035772 | Down |
| Bmf | 671,855034 | -0,635070369 | 0,153920572 | 3,35E-06 | 2,37E-05 | Down |
| Rpusd2 | 3129,977401 | 0,695967893 | 0,102151666 | 6,60E-13 | 1,48E-11 | Up |
| Spint1 | 253,6176392 | -0,833067958 | 0,230069717 | 5,99E-06 | 4,03E-05 | Down |
| Itpka | 124,4840104 | -0,639780249 | 0,164819861 | 8,76E-06 | 5,71E-05 | Down |
| Ganc | 2970,092092 | -0,773083622 | 0,117719266 | 1,72E-12 | 3,59E-11 | Down |
| Map1a | 7522,903493 | -0,974793427 | 0,238736593 | 4,10E-07 | 3,47E-06 | Down |
| Frmd5 | 14,20375734 | -0,873034958 | 0,523089361 | 0,001417544 | 0,005611789 | Down |
| Eif3j1 | 5425,686807 | 0,582035808 | 0,076097127 | 3,40E-15 | 1,08E-13 | Up |
| Gm14002 | 33,59088708 | -2,084330386 | 0,386650729 | 2,25E-10 | 3,31E-09 | Down |
| Fbn1 | 28,01245812 | 3,688181346 | 0,633102944 | 3,64E-10 | 5,23E-09 | Up |
| Itpripl1 | 10452,18108 | 0,586356906 | 0,09900398 | 4,79E-10 | 6,69E-09 | Up |
| Dusp2 | 2214,838787 | 0,947567969 | 0,133500092 | 1,00E-14 | 3,01E-13 | Up |
| Nphp1 | 1221,814138 | -0,681071759 | 0,115227255 | 2,57E-10 | 3,76E-09 | Down |
| Zc3h8 | 810,6321501 | 0,679953719 | 0,248417098 | 0,000315594 | 0,001454662 | Up |
| Ttl | 207,8065823 | -0,978403323 | 0,148052679 | 2,72E-13 | 6,48E-12 | Down |
| Polr1b | 10878,04045 | 0,984135679 | 0,122366508 | 5,45E-18 | 2,53E-16 | Up |
| Slc20a1 | 10384,98151 | 0,991458076 | 0,089075168 | 4,74E-31 | 1,01E-28 | Up |
| Il1b | 14,26609458 | 0,883264563 | 0,75529394 | 0,003109591 | 0,011260292 | Up |
| Ptpra | 12891,58638 | -0,581580684 | 0,098968554 | 6,51E-10 | 8,86E-09 | Down |
| Fastkd5 | 1720,723459 | 0,679167021 | 0,096071596 | 1,26E-13 | 3,14E-12 | Up |
| Hspa12b | 1291,193093 | -0,669796409 | 0,118058013 | 1,14E-09 | 1,49E-08 | Down |
| Cdc25b | 10853,36342 | -0,649265439 | 0,138276673 | 2,34E-07 | 2,06E-06 | Down |
| Ap5s1 | 565,1535008 | -1,256259204 | 0,160534572 | 3,25E-17 | 1,36E-15 | Down |
| Trmt6 | 8511,311653 | 0,603722985 | 0,107727035 | 2,77E-09 | 3,39E-08 | Up |
| Crls1 | 3860,085247 | 0,601127532 | 0,091994486 | 9,09E-12 | 1,68E-10 | Up |
| Esf1 | 8841,339853 | 0,74947663 | 0,099621056 | 2,28E-15 | 7,44E-14 | Up |
| Macrod2 | 256,2804725 | -1,094852806 | 0,21508889 | 2,06E-09 | 2,58E-08 | Down |
| Gm11688 | 57,51989578 | 0,782677561 | 0,288890265 | 0,000178673 | 0,000874078 | Up |
| Gm561 | 1103,035226 | 0,663672372 | 0,090382127 | 1,94E-14 | 5,51E-13 | Up |
| Rin2 | 3,310630952 | 2,365454425 | 2,024782712 | 0,000251237 | 0,001186484 | Up |
| Naa20 | 5878,205865 | 0,580963221 | 0,082943782 | 4,09E-13 | 9,40E-12 | Up |
| Trib3 | 5837,963377 | 1,22531441 | 0,525777687 | 9,38E-05 | 0,000485357 | Up |
| H13 | 23152,75005 | 0,643331628 | 0,150642717 | 1,70E-06 | 1,28E-05 | Up |
| Id1 | 13342,87747 | 2,32394012 | 0,136755864 | 6,74E-67 | 1,76E-63 | Up |
| Cox4i2 | 37,51901582 | 1,112848358 | 0,346023112 | 8,18E-06 | 5,36E-05 | Up |
| Bcl2l1 | 12015,67863 | 0,929577558 | 0,104836433 | 6,07E-21 | 4,17E-19 | Up |
| Ttll9 | 22,18053213 | 1,365570632 | 0,565906759 | 5,48E-05 | 0,000300371 | Up |
| Hck | 112,8469967 | 1,064034849 | 0,224342814 | 1,33E-08 | 1,47E-07 | Up |
| Nol4l | 6164,336407 | -1,36274874 | 0,114642641 | 1,55E-34 | 4,65E-32 | Down |
| Gm21850 | 111,8616401 | 2,962482777 | 0,730154774 | 4,29E-07 | 3,62E-06 | Up |
| Pxmp4 | 2668,970236 | -1,133825723 | 0,179257188 | 1,32E-12 | 2,79E-11 | Down |
| Ggt7 | 101,5361539 | -2,269596483 | 0,386873954 | 2,06E-11 | 3,59E-10 | Down |
| Acss2 | 1823,739378 | -0,978159331 | 0,205608307 | 1,65E-08 | 1,79E-07 | Down |

| | | | | | | |
|---|---|---|---|---|---|---|
| Sla2 | 10507,01329 | -0,77913236 | 0,166720146 | 9,12E-08 | 8,73E-07 | Down |
| Mroh8 | 31,87457383 | -0,945674755 | 0,373725211 | 0,000133424 | 0,000669025 | Down |
| Tgm2 | 10821,87749 | -1,37121099 | 0,334632999 | 1,66E-07 | 1,50E-06 | Down |
| D630003M21Rik | 639,0190827 | -1,477239344 | 0,235399147 | 2,05E-12 | 4,20E-11 | Down |
| Bpi | 137,6985846 | -0,654390042 | 0,307589691 | 0,001726589 | 0,006678874 | Down |
| Snhg11 | 12,70100956 | 0,628611952 | 0,629856634 | 0,01037018 | 0,032213 | Up |
| Slc32a1 | 35,61746957 | 1,053258398 | 0,342467864 | 1,60E-05 | 9,83E-05 | Up |
| Ppp1r16b | 161,7128126 | 1,757502097 | 0,257050585 | 3,10E-14 | 8,57E-13 | Up |
| Zhx3 | 21,63512516 | -0,596102666 | 0,372626587 | 0,006692557 | 0,022177367 | Down |
| Fitm2 | 2813,975891 | 0,813686792 | 0,089101416 | 1,51E-21 | 1,10E-19 | Up |
| Wfdc5 | 2,34860713 | -1,571822895 | 1,710452989 | 0,000939017 | 0,003880591 | Down |
| Wfdc12 | 53,19585331 | -1,687444353 | 0,392109727 | 4,80E-08 | 4,79E-07 | Down |
| Slpi | 1213,529228 | 1,521575902 | 0,484440705 | 4,63E-06 | 3,19E-05 | Up |
| Mmp9 | 25,13158665 | 2,927819153 | 0,761612973 | 8,16E-07 | 6,54E-06 | Up |
| Gm11469 | 10,86370356 | 1,432473869 | 1,09128271 | 0,000746718 | 0,003158074 | Up |
| Gm14267 | 11,2510132 | 2,548763151 | 1,003761942 | 2,26E-05 | 0,000134146 | Up |
| Kcnb1 | 4,347393697 | 1,105866939 | 1,461875879 | 0,002405119 | 0,008974721 | Up |
| Gm11473 | 149,3113433 | 2,051070799 | 0,250602055 | 7,68E-19 | 3,93E-17 | Up |
| B4galt5 | 12736,83706 | 0,718849804 | 0,100598115 | 5,08E-14 | 1,35E-12 | Up |
| Gm14236 | 22,32851066 | -0,760885964 | 0,40502854 | 0,001585669 | 0,006202946 | Down |
| Kcng1 | 18,44932235 | -0,590753758 | 0,45373287 | 0,010100486 | 0,031514619 | Down |
| Atp9a | 443,3186064 | -1,990744365 | 0,19633227 | 8,98E-27 | 1,20E-24 | Down |
| Gm17619 | 213,8174021 | -0,693120423 | 0,183954341 | 9,21E-06 | 5,98E-05 | Down |
| Pfdn4 | 3611,725278 | 0,787044783 | 0,093284268 | 9,60E-19 | 4,86E-17 | Up |
| Ppp4r1l-ps | 511,7770638 | -0,652760277 | 0,182128372 | 2,47E-05 | 0,000145384 | Down |
| Ctsz | 32776,91395 | 1,002346968 | 0,095330634 | 3,86E-28 | 5,97E-26 | Up |
| Zfp831 | 679,8420958 | -1,590897965 | 0,197603102 | 5,42E-18 | 2,52E-16 | Down |
| Edn3 | 84,35801001 | -2,150255328 | 0,232531489 | 7,42E-23 | 6,55E-21 | Down |
| Gm14296 | 24,96905536 | 1,036721352 | 0,441792963 | 0,000157303 | 0,000778925 | Up |
| Gm14327 | 90,33729027 | 0,661366802 | 0,284097569 | 0,001040399 | 0,004254314 | Up |
| 4921531C22Rik | 144,5384733 | -0,615684074 | 0,164516404 | 1,78E-05 | 0,000108426 | Down |
| B230312C02Rik | 601,4097964 | 1,124949777 | 0,266320246 | 1,38E-07 | 1,27E-06 | Up |
| Col9a3 | 253,833036 | -1,860378445 | 0,269471274 | 1,61E-14 | 4,62E-13 | Down |
| Dido1 | 22447,29191 | -0,646864366 | 0,084281229 | 1,77E-15 | 5,85E-14 | Down |
| Slc17a9 | 2603,04439 | 0,746745767 | 0,150737025 | 2,96E-08 | 3,08E-07 | Up |
| Ptk6 | 2,172040014 | 0,630022567 | 1,041790609 | 0,010097424 | 0,031514619 | Up |
| Zbtb46 | 16,81810674 | 0,768133472 | 0,517236314 | 0,003093131 | 0,011205467 | Up |
| Pcmtd2 | 2150,375499 | -0,741518648 | 0,1286043 | 3,59E-10 | 5,16E-09 | Down |
| E2f5 | 524,6423764 | 0,660738417 | 0,136459256 | 1,05E-07 | 9,96E-07 | Up |
| Car13 | 1555,814043 | 0,602406402 | 0,09550303 | 3,95E-11 | 6,55E-10 | Up |
| Mtfr1 | 7788,576125 | 0,586601633 | 0,093854402 | 6,31E-11 | 1,01E-09 | Up |
| Pde7a | 6154,783926 | 0,604177847 | 0,0770509 | 6,53E-16 | 2,30E-14 | Up |
| Cp | 259,6809462 | 3,361270813 | 0,673602915 | 1,93E-08 | 2,07E-07 | Up |
| Cpa3 | 212,7372238 | -0,990357444 | 0,330725066 | 2,64E-05 | 0,000154574 | Down |
| 7530428D23Rik | 4,34023333 | 0,675800163 | 0,795145076 | 0,009111247 | 0,028837238 | Up |
| Gm43672 | 2744,10837 | -0,942241013 | 0,194471304 | 1,25E-08 | 1,38E-07 | Down |
| Mrpl47 | 3789,224476 | 0,620765474 | 0,0973369 | 2,23E-11 | 3,84E-10 | Up |
| Gm43080 | 11,91841822 | -0,656693165 | 0,553825868 | 0,00779022 | 0,025197483 | Down |
| Cetn4 | 78,19373463 | -0,604794605 | 0,200893338 | 0,000235716 | 0,001119543 | Down |
| Bbs12 | 659,7257054 | 0,941984718 | 0,198176545 | 2,00E-08 | 2,14E-07 | Up |
| Spata5 | 4856,692355 | 0,619562761 | 0,074274946 | 9,80E-18 | 4,40E-16 | Up |

| | | | | | | |
|---|---|---|---|---|---|---|
| Gm2011 | 65,66743088 | -0,851450532 | 0,219071939 | 1,84E-06 | 1,37E-05 | Down |
| Gm6209 | 5,265119697 | 1,616339252 | 1,695010474 | 0,000880199 | 0,003666561 | Up |
| 4930577N17Rik | 144,423629 | -0,719050121 | 0,22176441 | 5,07E-05 | 0,000279432 | Down |
| Naa15 | 21743,0851 | 0,603850436 | 0,075640222 | 2,10E-16 | 7,99E-15 | Up |
| 5031434O11Rik | 165,0372201 | -0,904360894 | 0,16620686 | 6,19E-10 | 8,49E-09 | Down |
| Mgst2 | 3909,610169 | -1,07231692 | 0,201274575 | 5,85E-10 | 8,06E-09 | Down |
| Gm37261 | 9,632809236 | -2,458538597 | 1,056549418 | 3,74E-05 | 0,000212796 | Down |
| Maml3 | 4177,416488 | -1,237835276 | 0,117945912 | 6,50E-28 | 9,65E-26 | Down |
| Gm37170 | 7,982678 | -1,203253073 | 0,997775414 | 0,00132727 | 0,005286501 | Down |
| Alg5 | 6018,366977 | 0,716324434 | 0,10412505 | 3,46E-13 | 8,08E-12 | Up |
| Wwtr1 | 82,48549328 | 2,022176377 | 0,431410977 | 7,88E-09 | 9,05E-08 | Up |
| Rnf13 | 3586,596236 | 0,632694112 | 0,120872875 | 1,74E-08 | 1,88E-07 | Up |
| Gpr171 | 2450,094039 | -0,661805624 | 0,206308676 | 8,52E-05 | 0,000445739 | Down |
| P2ry14 | 6297,454053 | 1,536354684 | 0,19684887 | 4,30E-17 | 1,76E-15 | Up |
| P2ry13 | 20,34428268 | 2,515665618 | 0,699619478 | 1,17E-06 | 9,04E-06 | Up |
| Gm37035 | 10,23322782 | -0,760196558 | 0,543172883 | 0,003643468 | 0,012965122 | Down |
| Gm37696 | 93,53146176 | 0,77078006 | 0,267033637 | 0,000112988 | 0,000574917 | Up |
| Gm5540 | 29,31223885 | -0,826947 | 0,382406696 | 0,000604424 | 0,002612544 | Down |
| Gm18588 | 40,49121756 | -0,892760421 | 0,293813037 | 3,50E-05 | 0,000200165 | Down |
| Nmd3 | 8492,234769 | 0,800835276 | 0,080835011 | 9,90E-25 | 1,13E-22 | Up |
| Ppid | 11918,65474 | 0,865936658 | 0,143200044 | 2,19E-11 | 3,79E-10 | Up |
| Ctso | 607,02333 | -0,632581795 | 0,193497174 | 8,54E-05 | 0,000446902 | Down |
| Tlr2 | 2738,950673 | 0,674203239 | 0,128403133 | 1,16E-08 | 1,28E-07 | Up |
| Gpatch4 | 8405,066357 | 0,633375899 | 0,099473817 | 2,17E-11 | 3,76E-10 | Up |
| Lmna | 40,41698306 | 1,247126971 | 0,541638901 | 9,59E-05 | 0,000494979 | Up |
| Adam15 | 13988,43619 | 0,669191275 | 0,102991412 | 7,01E-12 | 1,31E-10 | Up |
| Dcst1 | 28,3747227 | -0,771345151 | 0,382923914 | 0,001127082 | 0,00456482 | Down |
| Pmvk | 4873,250219 | -0,703124315 | 0,128680033 | 2,84E-09 | 3,47E-08 | Down |
| Il6ra | 221,0480683 | 1,277602076 | 0,18352291 | 2,08E-14 | 5,89E-13 | Up |
| Creb3l4 | 41,44301925 | 0,649070773 | 0,322475013 | 0,002280817 | 0,008565142 | Up |
| Gm45477 | 22,79961156 | 1,032843944 | 0,428762477 | 0,000134556 | 0,000674333 | Up |
| S100a5 | 68,6835382 | -0,706930786 | 0,259834808 | 0,000276734 | 0,001291851 | Down |
| S100a9 | 1176,461046 | 0,82354398 | 1,114575144 | 0,005548921 | 0,018741335 | Up |
| Selenbp2 | 107,7579436 | -1,237413864 | 0,300502685 | 1,83E-07 | 1,64E-06 | Down |
| Selenbp1 | 1431,108593 | -1,425112218 | 0,255434915 | 1,26E-10 | 1,92E-09 | Down |
| Rfx5 | 3474,945428 | -0,597574946 | 0,10095123 | 4,55E-10 | 6,37E-09 | Down |
| 4930481B07Rik | 48,0674704 | -0,601471313 | 0,240447251 | 0,001003802 | 0,004115722 | Down |
| Sema6c | 26,40754602 | -1,3135681 | 0,434787228 | 9,78E-06 | 6,33E-05 | Down |
| 6330562C20Rik | 160,5213133 | -0,769757243 | 0,181250512 | 7,06E-07 | 5,72E-06 | Down |
| Ctss | 663,8275755 | 2,005391266 | 0,38819821 | 7,07E-10 | 9,54E-09 | Up |
| Golph3l | 5937,956436 | -0,659992029 | 0,154896383 | 1,57E-06 | 1,19E-05 | Down |
| Mtmr11 | 23,88170608 | -0,728965279 | 0,355444361 | 0,001309796 | 0,005224877 | Down |
| Hist2h2be | 164,9131176 | -0,716146692 | 0,168145351 | 1,02E-06 | 8,00E-06 | Down |
| Fcgr1 | 5289,287716 | 1,611815007 | 0,148050705 | 1,00E-29 | 1,85E-27 | Up |
| Pias3 | 2678,101871 | -0,785461905 | 0,083608745 | 1,72E-22 | 1,42E-20 | Down |
| Fmo5 | 307,3533099 | -0,79757347 | 0,150449385 | 3,07E-09 | 3,72E-08 | Down |
| Wdr3 | 15977,57698 | 0,625003232 | 0,065490963 | 1,82E-22 | 1,47E-20 | Up |
| Slc16a1 | 34988,48471 | 0,934198249 | 0,10899771 | 8,15E-20 | 4,80E-18 | Up |
| Fam212b | 327,4581697 | 0,726611027 | 0,171673365 | 1,06E-06 | 8,23E-06 | Up |
| Adora3 | 15,85920336 | 0,950334142 | 0,709695562 | 0,001923763 | 0,007340525 | Up |
| Wdr77 | 5436,562739 | 0,924378463 | 0,099373769 | 1,14E-22 | 9,69E-21 | Up |

| | | | | | | |
|---|---|---|---|---|---|---|
| Cd53 | 2647,867443 | 3,713826819 | 0,516591006 | 6,43E-14 | 1,69E-12 | Up |
| Prok1 | 7,228139486 | 4,715772473 | 1,536446776 | 2,42E-06 | 1,75E-05 | Up |
| Csf1 | 1974,845064 | 1,707175474 | 0,224052548 | 1,16E-16 | 4,51E-15 | Up |
| Gstm1 | 2134,720948 | -0,904989557 | 0,3701412 | 0,000202079 | 0,000974997 | Down |
| Gstm4 | 2477,900815 | -0,733688976 | 0,140836666 | 8,75E-09 | 9,94E-08 | Down |
| Celsr2 | 444,6304532 | -0,585822679 | 0,290162148 | 0,003374527 | 0,012104815 | Down |
| Cdc14a | 2958,192025 | -0,594118464 | 0,094043406 | 3,92E-11 | 6,50E-10 | Down |
| Frrs1 | 11187,99111 | 0,671109692 | 0,090072755 | 8,12E-15 | 2,47E-13 | Up |
| Dnttip2 | 14593,69658 | 0,762773258 | 0,095588329 | 5,51E-17 | 2,23E-15 | Up |
| 5730508B09Rik | 697,5044354 | 0,852269882 | 0,138718576 | 1,33E-11 | 2,39E-10 | Up |
| Pla2g12a | 4144,408242 | 0,698649095 | 0,090933713 | 1,07E-15 | 3,63E-14 | Up |
| Casp6 | 37,76323369 | 0,679223514 | 0,302646324 | 0,001134049 | 0,004587315 | Up |
| Gm43254 | 228,5329306 | -0,624954589 | 0,18763709 | 7,33E-05 | 0,000389316 | Down |
| Bdh2 | 50,70983699 | 0,667747725 | 0,404103801 | 0,004004149 | 0,014070756 | Up |
| Bank1 | 390,3070533 | 1,796969375 | 0,280088798 | 4,86E-13 | 1,10E-11 | Up |
| Dapp1 | 12758,11943 | -1,063063733 | 0,090739084 | 4,84E-34 | 1,40E-31 | Down |
| Gbp5 | 9,891779657 | 0,670753412 | 0,672795349 | 0,008497764 | 0,027164751 | Up |
| 9530052C20Rik | 67,58235858 | -0,624203659 | 0,252798621 | 0,000936468 | 0,00387181 | Down |
| Znhit6 | 3789,010644 | 0,605825754 | 0,085797911 | 2,33E-13 | 5,60E-12 | Up |
| Ddah1 | 12,46966775 | 2,686261629 | 0,648280686 | 2,23E-07 | 1,97E-06 | Up |
| Samd13 | 18,74532017 | -1,642881308 | 0,614514126 | 1,74E-05 | 0,00010612 | Down |
| Prkacb | 17503,93465 | -0,583727184 | 0,081206659 | 1,07E-13 | 2,69E-12 | Down |
| Dnajb4 | 3052,467561 | -1,337601865 | 0,150826622 | 6,27E-21 | 4,29E-19 | Down |
| St6galnac5 | 54,68535395 | -0,713929668 | 0,278474746 | 0,000408368 | 0,00183685 | Down |
| Cth | 7,41930163 | 2,062234099 | 1,419449078 | 0,000295429 | 0,001371061 | Up |
| Tmem68 | 1812,880651 | 0,620734906 | 0,112520075 | 4,05E-09 | 4,85E-08 | Up |
| A830012C17Rik | 10,27996963 | 4,817567 | 1,606170449 | 2,41E-06 | 1,74E-05 | Up |
| Cyp7a1 | 157,4428304 | -0,599346397 | 0,19436729 | 0,000195038 | 0,000944023 | Down |
| Gem | 254,5110708 | 8,778926099 | 1,767983773 | 5,71E-11 | 9,25E-10 | Up |
| Cdh17 | 137,6528549 | 9,256848505 | 1,436298613 | 1,17E-14 | 3,43E-13 | Up |
| Pdp1 | 2184,299773 | 1,735384082 | 0,226663023 | 8,10E-17 | 3,22E-15 | Up |
| Osgin2 | 3112,211715 | 0,861454495 | 0,102512406 | 6,18E-19 | 3,22E-17 | Up |
| Ttpa | 282,5969096 | 0,776084094 | 0,139992615 | 9,54E-10 | 1,25E-08 | Up |
| Ndufaf4 | 4332,278384 | 0,97157665 | 0,122930801 | 1,80E-17 | 7,86E-16 | Up |
| Epha7 | 24,0133338 | 0,710832564 | 0,497560803 | 0,004448383 | 0,015447113 | Up |
| Bach2os | 16,43915049 | 0,858003621 | 0,811772042 | 0,003924107 | 0,013815179 | Up |
| Mdn1 | 16966,43841 | 0,647330217 | 0,197112164 | 7,33E-05 | 0,000389316 | Up |
| Pnrc1 | 2319,610728 | -0,905020507 | 0,203513816 | 1,10E-07 | 1,03E-06 | Down |
| Gm11934 | 121,5598978 | 1,156445976 | 0,317481011 | 1,49E-06 | 1,13E-05 | Up |
| Cfap206 | 10,11146459 | -3,05002938 | 0,902287982 | 3,23E-06 | 2,29E-05 | Down |
| Sigmar1 | 15012,46882 | 0,666381846 | 0,084094219 | 2,12E-16 | 8,03E-15 | Up |
| Dnajb5 | 333,5590156 | 0,610091834 | 0,214046626 | 0,000366506 | 0,001665787 | Up |
| Fancg | 3664,885987 | -0,623477727 | 0,070099708 | 7,73E-20 | 4,59E-18 | Down |
| Cd72 | 402,1646575 | -1,461075287 | 0,152670657 | 1,05E-23 | 1,04E-21 | Down |
| Hint2 | 2775,708215 | -0,59859951 | 0,086896209 | 8,28E-13 | 1,82E-11 | Down |
| Tmem8b | 62,56912016 | -0,888581562 | 0,360459894 | 0,00020591 | 0,000992433 | Down |
| Glipr2 | 6207,664248 | 0,931736117 | 0,232100916 | 6,77E-07 | 5,51E-06 | Up |
| Polr1e | 6301,415042 | 0,668441634 | 0,096491001 | 3,76E-13 | 8,72E-12 | Up |
| Tomm5 | 13985,63141 | 0,600357244 | 0,108792059 | 4,60E-09 | 5,47E-08 | Up |
| Shb | 5,460236759 | -0,812855833 | 0,83305637 | 0,00499142 | 0,017098726 | Down |
| Trim14 | 1073,518607 | -0,620046621 | 0,126706779 | 1,10E-07 | 1,04E-06 | Down |

| | | | | | | |
|---|---|---|---|---|---|---|
| Baat | 9,027667035 | 1,392109451 | 0,956315545 | 0,000580179 | 0,002521472 | Up |
| Fktn | 1312,111317 | 0,682936939 | 0,127227121 | 5,72E-09 | 6,73E-08 | Up |
| Ikbkap | 10546,3223 | 0,676169874 | 0,082641463 | 2,38E-17 | 1,03E-15 | Up |
| Dnajc25 | 1972,880386 | 0,651995323 | 0,092113361 | 1,47E-13 | 3,67E-12 | Up |
| Gm12596 | 11,31078166 | -0,95674719 | 0,767418181 | 0,002199753 | 0,008286856 | Down |
| Snx30 | 5780,075166 | -0,707561767 | 0,107481978 | 2,85E-12 | 5,67E-11 | Down |
| Tnfsf15 | 4,46901965 | -0,741490382 | 0,999747075 | 0,007338111 | 0,023972914 | Down |
| Tnfsf8 | 2093,575493 | 2,351246451 | 0,130206247 | 6,07E-75 | 3,69E-71 | Up |
| Tlr4 | 3138,593152 | 0,706896656 | 0,177306498 | 3,47E-06 | 2,44E-05 | Up |
| Hacd4 | 6567,581851 | -0,738446912 | 0,224245248 | 3,73E-05 | 0,000212435 | Down |
| Mtap | 12636,51083 | 0,949492797 | 0,13265932 | 6,41E-15 | 1,98E-13 | Up |
| Elavl2 | 986,5023764 | 1,199729431 | 0,116778991 | 5,73E-27 | 7,93E-25 | Up |
| Tek | 20,72839217 | 0,587150651 | 0,366205669 | 0,007042513 | 0,023131362 | Up |
| Nfia | 299,0884067 | -1,505655801 | 0,185476467 | 3,76E-18 | 1,80E-16 | Down |
| Patj | 4,73228747 | -1,843490746 | 1,321032741 | 0,000401232 | 0,00180698 | Down |
| Gm12796 | 25,86428539 | -0,699558815 | 0,31675786 | 0,001088593 | 0,004429566 | Down |
| Lepr | 34,45425309 | 1,875571107 | 0,519638058 | 6,96E-07 | 5,64E-06 | Up |
| Pde4b | 1526,193179 | -1,256072708 | 0,130429382 | 4,33E-24 | 4,47E-22 | Down |
| Lrrc42 | 3447,548639 | 0,763337988 | 0,149080709 | 1,09E-08 | 1,21E-07 | Up |
| Ndc1 | 17390,36667 | 0,645328289 | 0,091358718 | 1,72E-13 | 4,22E-12 | Up |
| Lrp8 | 9688,957132 | 1,035814917 | 0,084033475 | 3,01E-37 | 1,17E-34 | Up |
| Echdc2 | 151,1693368 | 1,201566635 | 0,214332656 | 1,08E-10 | 1,67E-09 | Up |
| Coa7 | 2953,548371 | 0,713063223 | 0,104628782 | 5,58E-13 | 1,26E-11 | Up |
| Zfyve9 | 604,7746878 | 1,021576292 | 0,243814796 | 2,13E-07 | 1,88E-06 | Up |
| Osbpl9 | 8859,641117 | -0,606891818 | 0,098505645 | 9,72E-11 | 1,52E-09 | Down |
| Cdkn2c | 6893,065099 | -1,304140515 | 0,090679457 | 7,15E-49 | 5,93E-46 | Down |
| Foxd2 | 9,982544839 | -0,756321364 | 0,548563099 | 0,003819504 | 0,013512548 | Down |
| Pdzk1ip1 | 14,65678721 | -0,969208355 | 0,440394783 | 0,000295187 | 0,001370285 | Down |
| Tex38 | 12,37776897 | -0,65337801 | 0,497907901 | 0,006968238 | 0,022953456 | Down |
| Mob3c | 849,361509 | 0,986748061 | 0,117734582 | 3,15E-19 | 1,72E-17 | Up |
| Pomgnt1 | 6037,73283 | 0,624510725 | 0,105691314 | 4,04E-10 | 5,71E-09 | Up |
| Eif2b3 | 7067,412782 | 0,99425713 | 0,10336662 | 3,68E-24 | 3,86E-22 | Up |
| Tctex1d4 | 29,84910437 | -0,67725276 | 0,332754452 | 0,001831069 | 0,007036822 | Down |
| Plk3 | 911,0619477 | 1,094648814 | 0,228280533 | 9,56E-09 | 1,08E-07 | Up |
| Slc6a9 | 1220,144269 | 0,640450615 | 0,237018258 | 0,000453828 | 0,00202438 | Up |
| Ebna1bp2 | 23574,15937 | 0,676015146 | 0,099939594 | 1,10E-12 | 2,35E-11 | Up |
| Olfr1342 | 762,4197633 | -0,703520949 | 0,203070111 | 2,63E-05 | 0,000153938 | Down |
| Gm12863 | 3,749195886 | 2,986220281 | 1,860987993 | 9,41E-05 | 0,00048652 | Up |
| Slc2a1 | 11722,19401 | 0,848708608 | 0,119846061 | 2,36E-14 | 6,61E-13 | Up |
| Hivep3 | 636,3806459 | 0,964980187 | 0,181250596 | 8,53E-10 | 1,13E-08 | Up |
| Ctps | 21694,67565 | 0,758371912 | 0,09078154 | 2,60E-18 | 1,27E-16 | Up |
| Yrdc | 6107,586385 | 0,857842405 | 0,101793293 | 5,21E-19 | 2,75E-17 | Up |
| Csf3r | 399,1508416 | 1,641411125 | 0,446413529 | 6,14E-07 | 5,04E-06 | Up |
| Oscp1 | 235,554472 | -0,754992496 | 0,183228612 | 1,37E-06 | 1,05E-05 | Down |
| Tmem35b | 1067,155059 | -0,693181668 | 0,100216519 | 3,28E-13 | 7,68E-12 | Down |
| Gjb3 | 19,38929696 | 1,965869476 | 0,58710083 | 1,77E-06 | 1,32E-05 | Up |
| Tlr12 | 147,3951441 | -0,711406354 | 0,249471041 | 0,000183756 | 0,000896543 | Down |
| Zfp362 | 5602,70999 | -0,899603731 | 0,122851984 | 2,62E-15 | 8,48E-14 | Down |
| Fndc5 | 39,53355946 | -0,607324156 | 0,31119055 | 0,003360093 | 0,012062517 | Down |
| Lck | 92,78525319 | -0,778758128 | 0,401063298 | 0,001268578 | 0,005079318 | Down |
| Ccdc28b | 668,8239146 | -0,699115349 | 0,110511383 | 1,66E-11 | 2,93E-10 | Down |

| | | | | | | |
|---|---|---|---|---|---|---|
| Spocd1 | 103,4212202 | -2,805760443 | 0,32415477 | 2,57E-19 | 1,42E-17 | Down |
| 1700003M07Rik | 6,073520345 | 0,759033554 | 0,769624504 | 0,005967855 | 0,020004583 | Up |
| Matn1 | 423,3702018 | -0,670715333 | 0,247455506 | 0,000361817 | 0,001647754 | Down |
| Xkr8 | 1201,331124 | 0,616864714 | 0,093261914 | 4,80E-12 | 9,18E-11 | Up |
| Ahdc1 | 3993,025044 | -0,611408217 | 0,093457963 | 8,07E-12 | 1,50E-10 | Down |
| Pigv | 1139,638197 | -0,672919376 | 0,094442119 | 8,86E-14 | 2,26E-12 | Down |
| Cd52 | 1011,541979 | -0,946627848 | 0,23037646 | 4,15E-07 | 3,52E-06 | Down |
| Zfp593 | 158,6262685 | 0,714714849 | 0,217368282 | 4,48E-05 | 0,000249951 | Up |
| Extl1 | 477,9996045 | 0,670020717 | 0,460844209 | 0,005335338 | 0,018150882 | Up |
| Selenon | 8822,329824 | 1,051976357 | 0,080799337 | 4,24E-41 | 2,09E-38 | Up |
| Man1c1 | 2246,77068 | 2,489478919 | 0,204609114 | 8,38E-36 | 2,78E-33 | Up |
| Nipal3 | 3909,165676 | -0,743666413 | 0,11898793 | 1,81E-11 | 3,18E-10 | Down |
| Cnr2 | 639,2081342 | 1,277254908 | 0,241337584 | 6,18E-10 | 8,49E-09 | Up |
| E2f2 | 6364,647203 | -0,634403019 | 0,098912186 | 1,59E-11 | 2,82E-10 | Down |
| Gm13012 | 1059,186717 | -0,682156914 | 0,13736047 | 4,78E-08 | 4,78E-07 | Down |
| Hp1bp3 | 34369,36417 | -0,662499992 | 0,070015109 | 2,96E-22 | 2,35E-20 | Down |
| Sh2d5 | 9049,580851 | -0,618252949 | 0,192754458 | 0,000115527 | 0,000586695 | Down |
| Kif17 | 1815,628705 | -0,877132193 | 0,321406875 | 0,00010127 | 0,000520517 | Down |
| Pink1 | 1030,605435 | -1,187319002 | 0,371210217 | 7,16E-06 | 4,74E-05 | Down |
| Ubxn10 | 2,636420443 | 1,114595018 | 1,488824139 | 0,002307699 | 0,008650082 | Up |
| Pla2g2e | 33,9934475 | 2,714369112 | 0,792570359 | 2,37E-06 | 1,72E-05 | Up |
| Mrto4 | 19176,00783 | 0,842672965 | 0,124367411 | 2,18E-13 | 5,25E-12 | Up |
| Padi6 | 31,51235846 | -2,223836991 | 0,407247539 | 1,92E-10 | 2,85E-09 | Down |
| Crocc | 1849,853189 | -0,730802605 | 0,120736619 | 7,00E-11 | 1,12E-09 | Down |
| Tmem82 | 2,304810132 | -1,465336092 | 1,63423595 | 0,001160593 | 0,004688069 | Down |
| Gm13053 | 19,79336848 | -0,65634515 | 0,456140027 | 0,005873454 | 0,019709852 | Down |
| Kazn | 3608,778112 | -1,158904142 | 0,134540435 | 3,76E-20 | 2,33E-18 | Down |
| Gm13080 | 3,787561136 | 0,661209433 | 0,80330845 | 0,009786378 | 0,030666427 | Up |
| Dhrs3 | 6629,037815 | -1,092284999 | 0,308712818 | 2,64E-06 | 1,89E-05 | Down |
| Zfp933 | 1642,954443 | 0,682718505 | 0,141584112 | 9,78E-08 | 9,33E-07 | Up |
| Srm | 41624,26871 | 0,936271981 | 0,086182307 | 1,24E-29 | 2,22E-27 | Up |
| Gm13205 | 187,7671106 | 1,026436626 | 0,171717923 | 1,41E-11 | 2,51E-10 | Up |
| Slc25a33 | 1909,241325 | 1,559044869 | 0,101905245 | 8,10E-55 | 9,25E-52 | Up |
| Rere | 21079,32033 | -0,863860857 | 0,109012951 | 3,24E-17 | 1,36E-15 | Down |
| Gm13091 | 135,4061648 | -1,684601422 | 0,3565899 | 7,28E-09 | 8,40E-08 | Down |
| Tnfrsf9 | 961,1425314 | 0,9290053 | 0,188216482 | 8,40E-09 | 9,58E-08 | Up |
| Nol9 | 19987,10714 | 0,623181827 | 0,108369091 | 1,04E-09 | 1,36E-08 | Up |
| Plekhg5 | 3224,120707 | -0,921584094 | 0,143060767 | 1,15E-12 | 2,44E-11 | Down |
| Chd5 | 261,9581367 | -0,64439185 | 0,428167861 | 0,005744091 | 0,019329464 | Down |
| Atad3a | 19498,28592 | 0,649591127 | 0,102912528 | 2,74E-11 | 4,65E-10 | Up |
| B3galt6 | 1677,791476 | 0,597614576 | 0,101992623 | 6,52E-10 | 8,87E-09 | Up |
| Rnf223 | 55,244501 | -1,065862788 | 0,278957227 | 9,17E-07 | 7,27E-06 | Down |
| Samd11 | 115,4863289 | 1,6493704 | 0,352229979 | 9,19E-09 | 1,04E-07 | Up |
| Cdk14 | 64,29446486 | -0,677597459 | 0,241598886 | 0,000264385 | 0,001240226 | Down |
| Gm26825 | 144,5899417 | 0,640499759 | 0,215389413 | 0,000206641 | 0,000995695 | Up |
| Abcb1b | 4149,851263 | 1,599022385 | 0,113053125 | 1,76E-47 | 1,24E-44 | Up |
| Crot | 3100,332009 | -0,752270782 | 0,11932225 | 1,18E-11 | 2,12E-10 | Down |
| A630072M18Rik | 81,10294426 | -0,686880756 | 0,219026118 | 9,01E-05 | 0,000468476 | Down |
| Napepld | 977,2137184 | -0,764123156 | 0,08768232 | 1,09E-19 | 6,20E-18 | Down |
| 5031425E22Rik | 1281,159012 | -0,746734422 | 0,093108864 | 4,66E-17 | 1,90E-15 | Down |
| Gm25219 | 3,321012091 | -0,772825392 | 0,948021869 | 0,006484928 | 0,021551913 | Down |

| | | | | | | |
|---|---|---|---|---|---|---|
| AI506816 | 33460,89859 | 0,770927077 | 0,093187643 | 4,54E-18 | 2,14E-16 | Up |
| 2700038G22Rik | 1350,78502 | 0,622917674 | 0,209279582 | 0,000231167 | 0,001099656 | Up |
| Kcnh2 | 15,80891239 | 1,041303177 | 0,717902347 | 0,001200282 | 0,004833417 | Up |
| Nos3 | 60,47437923 | 2,545825427 | 0,286167903 | 1,06E-20 | 7,08E-19 | Up |
| Otof | 5,02105147 | -1,109753299 | 1,413564858 | 0,002476778 | 0,009211999 | Down |
| Khk | 614,7936558 | -1,030784321 | 0,14134961 | 1,69E-15 | 5,58E-14 | Down |
| Slc5a6 | 1836,307114 | 0,694739358 | 0,12367212 | 1,28E-09 | 1,66E-08 | Up |
| Ift172 | 1798,684001 | -0,809202999 | 0,085667463 | 8,26E-23 | 7,19E-21 | Down |
| AI839979 | 54,48895441 | 2,097815742 | 0,543629928 | 2,95E-07 | 2,56E-06 | Up |
| Plb1 | 16,8012613 | 2,150984357 | 0,69696 | 4,43E-06 | 3,07E-05 | Up |
| Gm15614 | 21,15205568 | -2,576411179 | 0,6086115 | 1,33E-07 | 1,23E-06 | Down |
| Gm42851 | 17,19997602 | 0,706578692 | 0,58290662 | 0,00592513 | 0,019865011 | Up |
| Mxd4 | 4469,061136 | -1,321855892 | 0,429925223 | 8,07E-06 | 5,29E-05 | Down |
| Sh3bp2 | 2111,114239 | 0,690191555 | 0,077306512 | 3,32E-20 | 2,09E-18 | Up |
| Nop14 | 11133,79299 | 0,753271736 | 0,086270339 | 1,05E-19 | 6,00E-18 | Up |
| Msantd1 | 3,267509621 | -0,599270788 | 0,783453057 | 0,012983479 | 0,039042093 | Down |
| Rgs12 | 20,80970257 | 1,760182095 | 0,65141021 | 1,46E-05 | 9,08E-05 | Up |
| Acox3 | 2910,678591 | -0,773583316 | 0,222156169 | 1,50E-05 | 9,29E-05 | Down |
| Htra3 | 66,85045509 | -1,15652827 | 0,579534309 | 0,000265539 | 0,001244678 | Down |
| Sh3tc1 | 912,8728443 | 1,320112431 | 0,158940969 | 7,65E-19 | 3,92E-17 | Up |
| Ppp2r2c | 9,036617986 | -1,310139974 | 0,831820527 | 0,000505213 | 0,002225861 | Down |
| Jakmip1 | 3673,307213 | -0,692790766 | 0,125585845 | 2,31E-09 | 2,87E-08 | Down |
| Lyar | 22318,58928 | 0,634405326 | 0,129647639 | 9,96E-08 | 9,47E-07 | Up |
| Clnk | 240,0028538 | 1,059483151 | 0,214338688 | 4,83E-09 | 5,72E-08 | Up |
| Bst1 | 33,67837635 | 3,562549846 | 0,892357149 | 7,10E-07 | 5,75E-06 | Up |
| Prom1 | 9,441801031 | 3,805962342 | 1,388094611 | 9,81E-06 | 6,35E-05 | Up |
| Lap3 | 21729,48932 | 0,614824634 | 0,086330027 | 1,42E-13 | 3,53E-12 | Up |
| Wdr19 | 376,459108 | -1,082153656 | 0,163026989 | 1,68E-13 | 4,13E-12 | Down |
| N4bp2os | 74,35762309 | -0,788618971 | 0,25079423 | 4,42E-05 | 0,000246859 | Down |
| Apbb2 | 43,3166444 | -2,043549609 | 0,435751349 | 7,92E-09 | 9,09E-08 | Down |
| Cnga1 | 150,1498631 | -1,139686301 | 0,307108269 | 1,18E-06 | 9,15E-06 | Down |
| Spata18 | 3,064164479 | 2,184643813 | 1,998585035 | 0,000327765 | 0,001506579 | Up |
| Gm6051 | 22,05936603 | -0,737899726 | 0,368232804 | 0,001380018 | 0,0054751 | Down |
| 2310040G07Rik | 3,379569122 | 0,678654504 | 0,918381941 | 0,009277782 | 0,029293024 | Up |
| 5830473C10Rik | 3008,949964 | 0,857874005 | 0,094889346 | 2,26E-21 | 1,62E-19 | Up |
| Ppbp | 98,37811165 | 6,181363865 | 0,982176476 | 4,94E-13 | 1,12E-11 | Up |
| Pf4 | 352,6255095 | 2,562793395 | 0,341231992 | 9,47E-16 | 3,23E-14 | Up |
| Cxcl3 | 14,84452677 | 3,555877577 | 1,800110262 | 3,50E-05 | 0,000200348 | Up |
| Cxcl2 | 57,55468698 | 4,587414314 | 0,695586786 | 1,31E-12 | 2,78E-11 | Up |
| Mthfd2l | 1151,184745 | 0,90474124 | 0,099832856 | 1,23E-21 | 9,05E-20 | Up |
| Sdad1 | 12175,14052 | 0,947134694 | 0,102661175 | 2,00E-22 | 1,61E-20 | Up |
| Shroom3 | 2,898188565 | 0,817397762 | 1,263146027 | 0,005002648 | 0,017127551 | Up |
| Gm43173 | 16,72317913 | 0,724520089 | 0,438815062 | 0,002936641 | 0,010691573 | Up |
| Ccng2 | 1329,31746 | -1,441665791 | 0,218867771 | 2,88E-13 | 6,84E-12 | Down |
| Gm7993 | 11509,51679 | 1,144598723 | 0,091379136 | 2,84E-38 | 1,23E-35 | Up |
| Anxa3 | 8980,662929 | 0,784549668 | 0,273520408 | 0,000109809 | 0,000559935 | Up |
| Bmp2k | 5934,617466 | -0,727526984 | 0,121387622 | 1,04E-10 | 1,61E-09 | Down |
| Agpat9 | 20,36580129 | 1,361676785 | 0,824765856 | 0,000388557 | 0,001755959 | Up |
| Spp1 | 14166,01247 | 1,760787164 | 0,2079212 | 9,98E-20 | 5,75E-18 | Up |
| Gm43819 | 34,74630871 | -0,804027072 | 0,441692969 | 0,001437375 | 0,005682906 | Down |
| Tmed5 | 8412,151858 | 0,765120183 | 0,113222898 | 5,09E-13 | 1,15E-11 | Up |

| | | | | | | |
|---|---|---|---|---|---|---|
| Gm42517 | 916,9586079 | -1,786335425 | 0,143341923 | 4,28E-38 | 1,82E-35 | Down |
| Gm10419 | 2401,713468 | -1,412508531 | 0,105035939 | 4,07E-43 | 2,32E-40 | Down |
| Gm18555 | 102,3025521 | -1,509512082 | 0,289270944 | 7,97E-10 | 1,06E-08 | Down |
| 1010001B22Rik | 41,51545778 | -1,374682946 | 0,462916782 | 1,02E-05 | 6,59E-05 | Down |
| Gm15787 | 34,12570586 | -0,72102044 | 0,320293398 | 0,000866128 | 0,003612065 | Down |
| Noc4l | 16149,99359 | 0,593259307 | 0,07703018 | 2,10E-15 | 6,91E-14 | Up |
| Ulk1 | 5417,08254 | -0,986388391 | 0,104610808 | 2,38E-23 | 2,22E-21 | Down |
| Hscb | 2015,219503 | -0,917613749 | 0,08092655 | 6,91E-32 | 1,60E-29 | Down |
| Ttc28 | 1354,800972 | -1,154857682 | 0,159895594 | 2,67E-15 | 8,63E-14 | Down |
| Tpst2 | 5661,792817 | 0,786766012 | 0,07575101 | 8,48E-27 | 1,14E-24 | Up |
| Tmem119 | 172,4539261 | 1,586769056 | 0,173130556 | 3,67E-22 | 2,89E-20 | Up |
| Ssh1 | 468,8903113 | -1,488792993 | 0,234793626 | 1,35E-12 | 2,84E-11 | Down |
| Svop | 52,53891762 | 1,497605888 | 0,32944173 | 2,06E-08 | 2,20E-07 | Up |
| Mvk | 3925,470077 | -0,634696295 | 0,179104992 | 3,25E-05 | 0,000187097 | Down |
| 4930515G01Rik | 137,6154348 | -0,871892451 | 0,167517874 | 2,88E-09 | 3,52E-08 | Down |
| 2410131K14Rik | 2866,547795 | 0,784333973 | 0,086377357 | 3,28E-21 | 2,32E-19 | Up |
| Rbm19 | 8907,972912 | 0,608078941 | 0,121396364 | 6,69E-08 | 6,54E-07 | Up |
| Sdsl | 2168,067081 | -0,674822889 | 0,10215506 | 3,25E-12 | 6,42E-11 | Down |
| Cfap73 | 9,059072128 | 0,698970514 | 0,668659235 | 0,007211535 | 0,023610085 | Up |
| Oas2 | 153,550936 | 2,191042495 | 0,362820003 | 6,02E-12 | 1,14E-10 | Up |
| Oas3 | 341,8636315 | 0,948476678 | 0,491003527 | 0,000607928 | 0,002624583 | Up |
| Naa25 | 16322,7553 | 0,752031031 | 0,078979747 | 7,21E-23 | 6,39E-21 | Up |
| Acad12 | 811,1184419 | -0,839923811 | 0,138711741 | 2,58E-11 | 4,40E-10 | Down |
| Atp2a2 | 60609,48706 | 0,611982088 | 0,077556963 | 4,18E-16 | 1,53E-14 | Up |
| P2rx7 | 32,61155604 | 2,828885179 | 0,722847938 | 5,93E-07 | 4,89E-06 | Up |
| Gm43684 | 112,7906376 | 0,853382856 | 0,264318397 | 2,25E-05 | 0,000133734 | Up |
| Gm43412 | 41,97313152 | -0,712803079 | 0,319726354 | 0,00095749 | 0,003951567 | Down |
| Tmem120b | 909,7917846 | 1,145655484 | 0,124460831 | 1,78E-22 | 1,45E-20 | Up |
| 4932422M17Rik | 25,70504106 | -0,645001858 | 0,473738255 | 0,006896974 | 0,022759767 | Down |
| Hcar2 | 29,20565238 | 4,602391971 | 0,706533544 | 2,18E-12 | 4,45E-11 | Up |
| Gm43661 | 410,5188138 | -0,667221192 | 0,126614438 | 1,11E-08 | 1,23E-07 | Down |
| Ncor2 | 30036,39979 | 0,58594854 | 0,102069666 | 1,41E-09 | 1,82E-08 | Up |
| Stx2 | 1745,612562 | -0,608297584 | 0,061859411 | 1,20E-23 | 1,17E-21 | Down |
| Cct6a | 67014,70203 | 0,618766041 | 0,081241109 | 3,43E-15 | 1,09E-13 | Up |
| Kctd7 | 181,009892 | -0,746755181 | 0,335235924 | 0,00078973 | 0,003325256 | Down |
| Wbscr16 | 7068,951236 | 0,61924173 | 0,089820536 | 6,93E-13 | 1,55E-11 | Up |
| Clip2 | 3968,004608 | -1,112724822 | 0,080077996 | 3,23E-46 | 2,18E-43 | Down |
| Syna | 5,872461978 | -2,363631778 | 1,647193763 | 0,000225812 | 0,00107811 | Down |
| Rfc2 | 25282,33014 | -0,97373528 | 0,068010461 | 9,20E-49 | 7,31E-46 | Down |
| Lat2 | 4766,608057 | -1,078985473 | 0,152090012 | 6,74E-15 | 2,07E-13 | Down |
| Wbscr27 | 680,0789105 | -0,810184904 | 0,172799398 | 6,61E-08 | 6,46E-07 | Down |
| Fkbp6 | 271,5532543 | -0,830070313 | 0,130214033 | 3,64E-12 | 7,08E-11 | Down |
| Gm43123 | 31,46551854 | 0,96065403 | 0,590390378 | 0,001099816 | 0,004468145 | Up |
| Ssc4d | 446,6429888 | -0,645353363 | 0,13400844 | 1,35E-07 | 1,24E-06 | Down |
| Sh2b2 | 4920,838545 | 0,998461966 | 0,137651897 | 2,50E-15 | 8,11E-14 | Up |
| Cux1 | 58335,39229 | -1,204005069 | 0,122753652 | 6,40E-25 | 7,45E-23 | Down |
| Myl10 | 156,465505 | -4,720959477 | 0,44374011 | 5,75E-28 | 8,68E-26 | Down |
| Serpine1 | 9,953018613 | 1,654901907 | 1,215445118 | 0,000518019 | 0,002277885 | Up |
| Ufsp1 | 671,7375763 | 0,836858413 | 0,150867241 | 5,56E-10 | 7,67E-09 | Up |
| Zan | 30,89876746 | 0,616340258 | 0,313883136 | 0,003097254 | 0,011218179 | Up |
| Gm8066 | 348,4924925 | -0,659795844 | 0,2262475 | 0,000215049 | 0,001032391 | Down |

| | | | | | | |
|---|---|---|---|---|---|---|
| Tfr2 | 6268,32924 | -2,209306757 | 0,202967607 | 5,18E-30 | 9,85E-28 | Down |
| Mospd3 | 5783,974927 | -0,722575348 | 0,096003613 | 2,88E-15 | 9,25E-14 | Down |
| Zcwpw1 | 1245,961138 | -0,76355338 | 0,149247924 | 1,11E-08 | 1,23E-07 | Down |
| Pilra | 72,16270277 | 3,226861403 | 0,354442443 | 1,16E-20 | 7,69E-19 | Up |
| Pilrb1 | 241,8391833 | 3,330280478 | 0,279061954 | 1,35E-33 | 3,75E-31 | Up |
| Gm36551 | 78,80995591 | 2,91228713 | 0,290845308 | 1,20E-24 | 1,36E-22 | Up |
| Pilrb2 | 19,10539615 | 2,034885032 | 0,897237702 | 4,54E-05 | 0,000253059 | Up |
| Zscan21 | 3787,406892 | -0,718247245 | 0,072118805 | 1,37E-24 | 1,53E-22 | Down |
| Nxpe5 | 888,0046898 | 1,046250812 | 0,674669543 | 0,000972635 | 0,004008637 | Up |
| BC037034 | 9528,791345 | 0,81856131 | 0,197583429 | 7,73E-07 | 6,23E-06 | Up |
| Fam20c | 3,118193038 | 2,048195001 | 1,953376379 | 0,00040643 | 0,001829484 | Up |
| 3110082I17Rik | 13716,31976 | 0,657530615 | 0,098354055 | 2,19E-12 | 4,47E-11 | Up |
| Gpr146 | 6276,57489 | -0,841538969 | 0,086992437 | 6,57E-24 | 6,59E-22 | Down |
| Micall2 | 18999,42081 | -1,018420848 | 0,099491408 | 6,83E-27 | 9,37E-25 | Down |
| Snx8 | 11856,6436 | 0,720830159 | 0,115704163 | 2,53E-11 | 4,32E-10 | Up |
| Lfng | 13135,06574 | 1,092273649 | 0,080009548 | 9,03E-45 | 5,49E-42 | Up |
| Spdye4b | 5,901235266 | -0,598145931 | 0,655478195 | 0,012599499 | 0,038000337 | Down |
| Aimp2 | 10114,03915 | 0,663828014 | 0,085743809 | 9,17E-16 | 3,15E-14 | Up |
| Zkscan14 | 2037,216129 | -0,601372334 | 0,107772054 | 3,23E-09 | 3,90E-08 | Down |
| Cdk8 | 14946,15952 | 0,637554236 | 0,079382468 | 1,12E-16 | 4,37E-15 | Up |
| D5Ertd605e | 7495,92732 | -0,69990849 | 0,090479573 | 7,05E-16 | 2,45E-14 | Down |
| Gm43460 | 384,162468 | -0,645495687 | 0,126817335 | 3,38E-08 | 3,47E-07 | Down |
| Flt1 | 206,4400117 | 1,250061926 | 0,222696677 | 1,05E-10 | 1,63E-09 | Up |
| Slc7a1 | 102909,5287 | 0,747845583 | 0,109242839 | 3,26E-13 | 7,63E-12 | Up |
| 8430423G03Rik | 88,71111414 | 1,208123792 | 0,273826314 | 5,16E-08 | 5,12E-07 | Up |
| Gm43197 | 35,47681764 | -0,892767407 | 0,368733685 | 0,000227897 | 0,001086078 | Down |
| Asns | 30785,17086 | 0,616150973 | 0,152637362 | 5,55E-06 | 3,76E-05 | Up |
| 2610001J05Rik | 9642,694611 | 0,902486216 | 0,11025053 | 2,74E-18 | 1,33E-16 | Up |
| Met | 4940,381982 | 1,165442687 | 0,159656892 | 1,53E-15 | 5,09E-14 | Up |
| Wnt2 | 10,91640668 | -0,664894109 | 0,576000822 | 0,007689767 | 0,024907861 | Down |
| Asz1 | 351,0637868 | -2,48085515 | 0,349342978 | 1,32E-14 | 3,84E-13 | Down |
| Impdh1 | 18681,09057 | 0,617982557 | 0,07657262 | 9,39E-17 | 3,70E-15 | Up |
| Hilpda | 1219,348805 | 0,65661083 | 0,101345567 | 8,77E-12 | 1,62E-10 | Up |
| Flnc | 18,83963871 | -0,933792839 | 0,457161942 | 0,000499459 | 0,002204087 | Down |
| Cpa5 | 32,44097674 | -1,536821308 | 0,367084768 | 9,30E-08 | 8,89E-07 | Down |
| Mkln1os | 45,96211869 | -1,137724608 | 0,268523011 | 1,27E-07 | 1,18E-06 | Down |
| Akr1b8 | 20,47584585 | 2,101868976 | 0,604600104 | 1,20E-06 | 9,24E-06 | Up |
| Akr1b7 | 8,54091321 | 0,722305678 | 0,870549028 | 0,007687568 | 0,024906348 | Up |
| 1810058I24Rik | 8858,372937 | -0,718561449 | 0,088602744 | 2,94E-17 | 1,24E-15 | Down |
| Ttc26 | 602,2881173 | -0,668447343 | 0,166868079 | 4,31E-06 | 2,99E-05 | Down |
| Slc37a3 | 345,3545136 | 2,042615717 | 0,360449361 | 4,51E-11 | 7,42E-10 | Up |
| Rab19 | 13,70314557 | 2,533719634 | 1,035062166 | 2,80E-05 | 0,00016334 | Up |
| Gm10244 | 14,42052626 | -1,103549724 | 0,601391178 | 0,000449156 | 0,002004516 | Down |
| Olfr460 | 2,085048759 | 0,66248933 | 1,09774786 | 0,008641008 | 0,027539632 | Up |
| Clec5a | 2770,782305 | 2,863229522 | 0,332121296 | 4,15E-19 | 2,22E-17 | Up |
| Mgam | 41,70671067 | 0,763559793 | 0,404512328 | 0,001535396 | 0,006027386 | Up |
| Trbv12-2 | 42,5808743 | -0,866788943 | 0,275519547 | 2,78E-05 | 0,000162254 | Down |
| Trbv13-2 | 116,371446 | -0,977999639 | 0,226549175 | 1,40E-07 | 1,29E-06 | Down |
| Gm43889 | 34,29366836 | 3,894133005 | 0,708357501 | 1,92E-09 | 2,42E-08 | Up |
| A930035D04Rik | 54,44041785 | -0,680949065 | 0,261945586 | 0,000458278 | 0,002040739 | Down |
| Sspo | 6,647129153 | 1,188678945 | 1,398260542 | 0,002111763 | 0,007999381 | Up |

| | | | | | | |
|---|---|---|---|---|---|---|
| Atp6v0e2 | 73,35517351 | 0,7145538 | 0,286902262 | 0,000494027 | 0,002182781 | Up |
| Gimap5 | 1384,344771 | -0,76168585 | 0,16027285 | 7,16E-08 | 6,96E-07 | Down |
| Tmem176a | 11316,0182 | -0,761966117 | 0,187324069 | 1,62E-06 | 1,22E-05 | Down |
| Mpp6 | 13443,06018 | 0,656043988 | 0,094461492 | 3,68E-13 | 8,55E-12 | Up |
| 9430076C15Rik | 3288,824048 | -0,993927623 | 0,1766822 | 1,33E-10 | 2,02E-09 | Down |
| Ggct | 684,1492907 | 0,961851911 | 0,142518282 | 1,12E-13 | 2,80E-12 | Up |
| Ccdc129 | 6,049398883 | -1,905162152 | 1,324053083 | 0,00034863 | 0,001593263 | Down |
| Gm10209 | 7,663127667 | -0,780084707 | 0,679094969 | 0,004626812 | 0,015990696 | Down |
| Lancl2 | 7269,827452 | -0,628059997 | 0,102346003 | 9,77E-11 | 1,52E-09 | Down |
| Vopp1 | 17950,10709 | -0,657746444 | 0,098298759 | 2,10E-12 | 4,30E-11 | Down |
| Ccser1 | 22,01476401 | 3,263997419 | 0,789277111 | 4,25E-07 | 3,59E-06 | Up |
| Hpgds | 21,68390694 | 0,898798152 | 0,511180715 | 0,00107111 | 0,004364258 | Up |
| Tnip3 | 175,1734794 | 0,869021389 | 0,301871772 | 6,63E-05 | 0,000356555 | Up |
| Gm6786 | 14,5831849 | 1,960270128 | 0,843832778 | 3,99E-05 | 0,000225377 | Up |
| Smyd1 | 43,64976419 | -0,757604072 | 0,250302208 | 7,92E-05 | 0,000418216 | Down |
| Rmnd5a | 17235,20124 | -0,60127766 | 0,113015814 | 1,36E-08 | 1,50E-07 | Down |
| Gm44186 | 18,00696885 | 0,707492398 | 0,517889956 | 0,004918142 | 0,016860353 | Up |
| Polr1a | 20692,40902 | 0,584677278 | 0,107387136 | 7,67E-09 | 8,82E-08 | Up |
| Gm45051 | 35,65395658 | -0,597734839 | 0,285997652 | 0,002692444 | 0,009909193 | Down |
| Tmem150a | 223,9175703 | 1,261044784 | 0,169130376 | 5,61E-16 | 2,01E-14 | Up |
| Vamp8 | 23722,97101 | -0,763619434 | 0,077935819 | 4,32E-24 | 4,47E-22 | Down |
| Mat2a | 30186,51922 | 0,732412853 | 0,09472284 | 5,34E-16 | 1,91E-14 | Up |
| Particl | 170,8602644 | -0,705261874 | 0,150058578 | 1,47E-07 | 1,35E-06 | Down |
| Capg | 17969,49395 | 0,661744655 | 0,151959211 | 1,03E-06 | 8,03E-06 | Up |
| Mrpl19 | 11280,89539 | 0,604635859 | 0,080236305 | 7,03E-15 | 2,15E-13 | Up |
| Tacr1 | 5,078795076 | 0,739610647 | 1,123538003 | 0,006957868 | 0,022923432 | Up |
| Hk2 | 19649,01962 | 1,068597399 | 0,101793994 | 4,13E-28 | 6,31E-26 | Up |
| Gm17034 | 20,10568958 | -1,188570691 | 0,518209791 | 0,00011394 | 0,000579439 | Down |
| Gm43878 | 55,94813608 | 1,469445812 | 0,378255469 | 3,43E-07 | 2,94E-06 | Up |
| Wbp1 | 1559,880948 | -0,820289085 | 0,139560698 | 9,09E-11 | 1,43E-09 | Down |
| Gm44287 | 2,948203565 | -0,663742296 | 1,139579387 | 0,008044935 | 0,025924934 | Down |
| Mthfd2 | 15492,30892 | 0,832556434 | 0,112845079 | 3,06E-15 | 9,78E-14 | Up |
| Dguok | 2663,947094 | -0,633392136 | 0,093917777 | 1,77E-12 | 3,68E-11 | Down |
| Gm21284 | 114,1338114 | -0,597291444 | 0,247355959 | 0,001282515 | 0,0051295 | Down |
| Paip2b | 144,6099157 | 0,709213571 | 0,198721858 | 1,73E-05 | 0,000105579 | Up |
| Exoc6b | 1711,101604 | -0,584371909 | 0,111889867 | 2,56E-08 | 2,69E-07 | Down |
| Rab11fip5 | 155,2319165 | 3,24328075 | 0,410634464 | 3,10E-16 | 1,15E-14 | Up |
| Smyd5 | 18393,46982 | 0,682633292 | 0,103004049 | 2,64E-12 | 5,28E-11 | Up |
| Add2 | 176,2583898 | -1,168011953 | 0,35529736 | 5,49E-06 | 3,72E-05 | Down |
| Fam136a | 15246,30518 | 0,744615343 | 0,102596692 | 1,75E-14 | 5,01E-13 | Up |
| Gm44198 | 184,4808107 | 0,698791773 | 0,14181474 | 5,07E-08 | 5,05E-07 | Up |
| Gm43904 | 7,527783735 | -2,396784593 | 1,191026306 | 7,73E-05 | 0,000409018 | Down |
| Gp9 | 601,7823474 | -1,010178512 | 0,264368255 | 1,10E-06 | 8,55E-06 | Down |
| Gm26588 | 43,18107596 | 1,642685113 | 0,306969611 | 3,34E-10 | 4,82E-09 | Up |
| Mgll | 53,88534522 | 2,126653481 | 1,175293587 | 0,000136016 | 0,000681088 | Up |
| Abtb1 | 2440,507243 | -0,790389734 | 0,15448664 | 8,82E-09 | 1,00E-07 | Down |
| Txnrd3 | 1087,985947 | 0,759714009 | 0,13873963 | 1,62E-09 | 2,06E-08 | Up |
| Aldh1l1 | 4,678236003 | 0,913109005 | 0,936760751 | 0,003689992 | 0,013120452 | Up |
| Chchd4 | 12949,78282 | 0,76606397 | 0,098639001 | 2,93E-16 | 1,10E-14 | Up |
| Kbtbd8 | 2457,735476 | 0,623522293 | 0,097831265 | 2,24E-11 | 3,86E-10 | Up |
| Frmd4b | 176,0736522 | -0,834958978 | 0,332775236 | 0,000236749 | 0,001123867 | Down |

| | | | | | | |
|---|---|---|---|---|---|---|
| Il5ra | 247,2943963 | -1,295974852 | 0,370049958 | 1,92E-06 | 1,42E-05 | Down |
| Gm16433 | 130,5902321 | 1,079249133 | 0,215091256 | 3,12E-09 | 3,78E-08 | Up |
| Bhlhe40 | 21345,23936 | 1,021525919 | 0,128319225 | 9,33E-18 | 4,22E-16 | Up |
| Mtmr14 | 5290,174629 | -0,80989284 | 0,101441163 | 3,33E-17 | 1,39E-15 | Down |
| Cpne9 | 16,51521114 | 0,645230713 | 0,572286719 | 0,008697178 | 0,027694483 | Up |
| Jagn1 | 6669,605259 | 0,746658444 | 0,088641329 | 1,62E-18 | 8,07E-17 | Up |
| Il17re | 269,1383942 | -1,481252572 | 0,256228092 | 3,88E-11 | 6,45E-10 | Down |
| Tatdn2 | 13269,80128 | 0,609850428 | 0,10309109 | 4,29E-10 | 6,05E-09 | Up |
| Cand2 | 1623,639553 | -2,650260423 | 0,228694001 | 1,92E-32 | 4,80E-30 | Down |
| H1foo | 5,167665808 | 1,531920361 | 1,5532881 | 0,001039804 | 0,004252835 | Up |
| Gm8213 | 184,2105679 | -1,095963933 | 0,194569914 | 9,81E-11 | 1,53E-09 | Down |
| Zfand4 | 842,5799436 | -0,662094455 | 0,10432064 | 1,99E-11 | 3,46E-10 | Down |
| Fxyd4 | 283,3115694 | 0,644256346 | 0,222616311 | 0,000255565 | 0,001203488 | Up |
| Gm23908 | 19,19493465 | -1,443784087 | 0,596469631 | 4,67E-05 | 0,000259524 | Down |
| Ret | 239,5945892 | -1,00698016 | 0,261934975 | 1,01E-06 | 7,92E-06 | Down |
| Bms1 | 25595,54079 | 0,583106898 | 0,086239833 | 2,20E-12 | 4,48E-11 | Up |
| Cacna2d4 | 144,6023226 | -1,214152355 | 0,209982268 | 3,90E-11 | 6,47E-10 | Down |
| Lrtm2 | 28,31569374 | -1,009258288 | 0,509138347 | 0,000432858 | 0,001936988 | Down |
| Tuba8 | 21444,23503 | 0,60362799 | 0,156883862 | 1,29E-05 | 8,09E-05 | Up |
| Usp18 | 164,2414255 | 0,734212411 | 0,214545633 | 2,45E-05 | 0,000144459 | Up |
| Phc1 | 2601,639079 | -1,285435462 | 0,104311098 | 6,27E-37 | 2,34E-34 | Down |
| Slc2a3 | 28352,06815 | 0,774531429 | 0,125894493 | 2,52E-11 | 4,31E-10 | Up |
| Foxj2 | 9,183841608 | 0,603802357 | 0,51455287 | 0,010496794 | 0,032552216 | Up |
| Clec4a2 | 3,0192714 | 0,856370411 | 1,341282068 | 0,004157794 | 0,014529205 | Up |
| Clec4n | 64,08773858 | 3,562654089 | 0,835554082 | 3,14E-07 | 2,71E-06 | Up |
| Clec4d | 1846,380056 | 0,631750357 | 0,201028164 | 0,000129768 | 0,000652124 | Up |
| Clec4e | 2450,530282 | 0,969159881 | 0,220265565 | 9,88E-08 | 9,41E-07 | Up |
| C1ra | 714,1074443 | 1,424765067 | 0,269345738 | 5,95E-10 | 8,19E-09 | Up |
| Grcc10 | 710,8403106 | -0,768718688 | 0,164952404 | 1,06E-07 | 9,99E-07 | Down |
| Spsb2 | 1070,785801 | -0,686760676 | 0,10318377 | 2,10E-12 | 4,30E-11 | Down |
| Lag3 | 151,5211733 | -1,667465023 | 0,211752618 | 1,79E-17 | 7,84E-16 | Down |
| Ptms | 5382,522805 | -0,835234272 | 0,125011244 | 4,46E-13 | 1,02E-11 | Down |
| Nop2 | 29253,70513 | 0,882289324 | 0,129170419 | 1,07E-13 | 2,69E-12 | Up |
| Gm28809 | 390,8351635 | -0,837917755 | 0,269937219 | 3,76E-05 | 0,000213825 | Down |
| Fgf23 | 169,6018244 | 4,108655638 | 0,533145804 | 9,34E-16 | 3,20E-14 | Up |
| Tigar | 2775,434518 | 1,140946233 | 0,13734311 | 5,04E-19 | 2,67E-17 | Up |
| Ccnd2 | 58882,22585 | 0,761282698 | 0,077447522 | 3,24E-24 | 3,44E-22 | Up |
| Gm26770 | 159,7417486 | -2,198496379 | 0,207782329 | 1,34E-28 | 2,17E-26 | Down |
| Rhno1 | 8719,928351 | -1,795326885 | 0,106320707 | 1,87E-66 | 4,26E-63 | Down |
| Foxm1 | 28001,67483 | -0,980326583 | 0,082367163 | 6,33E-35 | 1,96E-32 | Down |
| Gm43965 | 93,62503784 | -1,244350261 | 0,319315692 | 4,56E-07 | 3,84E-06 | Down |
| Cd69 | 4378,454413 | 2,355563778 | 0,299701731 | 2,64E-17 | 1,14E-15 | Up |
| 4922502D21Rik | 2,495827171 | 1,204904777 | 1,56006073 | 0,001862162 | 0,007138288 | Up |
| Clec12a | 1081,153479 | 0,60787862 | 0,56380115 | 0,010989548 | 0,033843361 | Up |
| Klra2 | 7,263873726 | 1,009289047 | 0,926534452 | 0,002455031 | 0,009140423 | Up |
| Magohb | 745,1752379 | 0,614071551 | 0,104112617 | 4,63E-10 | 6,48E-09 | Up |
| Styk1 | 7,943591511 | 1,172046261 | 1,077272257 | 0,001731183 | 0,006693809 | Up |
| Ybx3 | 69223,30953 | 0,616042924 | 0,09545265 | 1,40E-11 | 2,50E-10 | Up |
| Bcl2l14 | 2146,088438 | 1,55326407 | 0,141488446 | 4,50E-30 | 8,66E-28 | Up |
| Cdkn1b | 2508,115179 | -0,793950231 | 0,138479011 | 2,71E-10 | 3,95E-09 | Down |
| Lockd | 845,699717 | -0,682109084 | 0,095770772 | 8,36E-14 | 2,15E-12 | Down |

| | | | | | | |
|---|---|---|---|---|---|---|
| Emp1 | 7,454085829 | 4,251732208 | 1,835161789 | 1,12E-05 | 7,12E-05 | Up |
| Gm15707 | 291,8809628 | -0,921005811 | 0,190479553 | 1,48E-08 | 1,62E-07 | Down |
| Caprin2 | 238,6661571 | -0,807669304 | 0,158105042 | 7,96E-09 | 9,12E-08 | Down |
| Fam60a | 3785,475364 | 0,860948366 | 0,081163056 | 3,77E-28 | 5,89E-26 | Up |
| Dennd5b | 69,37463206 | 1,001016304 | 0,431368391 | 0,000191742 | 0,000930778 | Up |
| Tarm1 | 1208,628513 | 0,594634285 | 0,239896684 | 0,001112932 | 0,00451452 | Up |
| Oscar | 78,10024709 | 1,703918402 | 0,287799724 | 1,24E-11 | 2,23E-10 | Up |
| Rdh13 | 4023,991193 | 0,997209716 | 0,136294636 | 1,56E-15 | 5,19E-14 | Up |
| Isoc2b | 147,8203905 | -0,692048516 | 0,275342692 | 0,000536267 | 0,002348515 | Down |
| Zfp580 | 1035,542264 | -0,878863432 | 0,122446049 | 9,05E-15 | 2,74E-13 | Down |
| Zfp128 | 642,141913 | -0,828979401 | 0,113275189 | 4,95E-15 | 1,54E-13 | Down |
| 1810019N24Rik | 33,00560074 | -0,917740336 | 0,458892588 | 0,000588706 | 0,002554276 | Down |
| Pla2g4c | 248,4843085 | -2,067547808 | 0,356725011 | 2,16E-11 | 3,74E-10 | Down |
| Ehd2 | 210,8835274 | 2,797630411 | 0,315394543 | 3,87E-20 | 2,36E-18 | Up |
| C5ar1 | 80,99254291 | 3,257848211 | 0,328291221 | 4,89E-24 | 4,99E-22 | Up |
| Gm45509 | 70,84509336 | 0,736866244 | 0,24013578 | 7,94E-05 | 0,000419115 | Up |
| Slc1a5 | 12790,89028 | 0,91130631 | 0,086323535 | 4,14E-28 | 6,31E-26 | Up |
| Prkd2 | 8148,275064 | 0,858188933 | 0,090722597 | 4,45E-23 | 4,08E-21 | Up |
| Gng8 | 26,98188007 | 2,677438757 | 0,636016896 | 1,72E-07 | 1,55E-06 | Up |
| Ptgir | 1602,251641 | 1,465446234 | 0,211009181 | 2,58E-14 | 7,20E-13 | Up |
| Pnmal1 | 16,19989722 | -0,782228396 | 0,431043218 | 0,001616703 | 0,006303209 | Down |
| Psg17 | 14,23180398 | -2,003390993 | 1,681800961 | 0,000445462 | 0,001989977 | Down |
| Mill2 | 26,68541156 | -1,720469222 | 0,721831052 | 3,78E-05 | 0,000214604 | Down |
| Gm45138 | 8,552773256 | -0,596292687 | 0,60420135 | 0,012316721 | 0,03727058 | Down |
| Ccdc61 | 2143,291749 | -0,720793955 | 0,108897157 | 1,98E-12 | 4,08E-11 | Down |
| Irf2bp1 | 4849,034545 | -0,596429258 | 0,109577145 | 7,19E-09 | 8,30E-08 | Down |
| Rsph6a | 18,19852447 | 1,013542233 | 0,589631382 | 0,000763375 | 0,003221806 | Up |
| Dmwd | 5860,363685 | 0,892230573 | 0,08331506 | 9,46E-29 | 1,57E-26 | Up |
| Cd3eap | 8986,04644 | 0,752975737 | 0,147106692 | 1,20E-08 | 1,33E-07 | Up |
| Zfp296 | 879,7415816 | 0,674419108 | 0,099877985 | 1,21E-12 | 2,57E-11 | Up |
| Relb | 527,0632685 | 0,988266741 | 0,230201473 | 1,50E-07 | 1,37E-06 | Up |
| Tomm40 | 21036,04782 | 0,630032144 | 0,109308199 | 9,18E-10 | 1,21E-08 | Up |
| Pvr | 729,3870502 | 1,208863185 | 0,179394328 | 8,81E-14 | 2,25E-12 | Up |
| Plaur | 2252,858594 | 1,457015487 | 0,156467517 | 1,21E-22 | 1,02E-20 | Up |
| Cd177 | 3314,386718 | 1,166115553 | 0,208798233 | 1,23E-10 | 1,89E-09 | Up |
| Ccdc97 | 5506,470297 | -0,775348893 | 0,086124657 | 7,33E-21 | 4,98E-19 | Down |
| Cyp2b10 | 20,59787078 | 1,517255366 | 0,948897876 | 0,000353855 | 0,00161593 | Up |
| Rab4b | 333,2527098 | -0,63775669 | 0,123258681 | 2,31E-08 | 2,44E-07 | Down |
| Numbl | 178,9177678 | -1,503697204 | 0,338941342 | 3,33E-08 | 3,42E-07 | Down |
| Ltbp4 | 28,60761441 | -1,281986177 | 0,452592821 | 1,91E-05 | 0,000115297 | Down |
| Pgam1-ps2 | 138,2521283 | 0,936517419 | 0,293296753 | 1,67E-05 | 0,000102243 | Up |
| Sertad3 | 879,2062655 | -0,62819988 | 0,106325504 | 3,95E-10 | 5,61E-09 | Down |
| Prx | 249,3020605 | -0,604058853 | 0,177976635 | 6,81E-05 | 0,000364873 | Down |
| Gm44685 | 125,6007902 | -1,07946761 | 0,233763104 | 2,39E-08 | 2,53E-07 | Down |
| Gm10046 | 23,86213808 | -1,1776622 | 0,421782603 | 2,79E-05 | 0,000162724 | Down |
| Zfp974 | 168,5219009 | -0,591862047 | 0,145267368 | 5,59E-06 | 3,78E-05 | Down |
| 9530053A07Rik | 88,21856048 | 3,994781757 | 0,418339255 | 1,22E-22 | 1,03E-20 | Up |
| Gm45010 | 81,40064451 | -0,733583634 | 0,198688605 | 9,10E-06 | 5,91E-05 | Down |
| Zfp36 | 1356,414366 | 1,118821496 | 0,193028595 | 3,63E-11 | 6,06E-10 | Up |
| Rasgrp4 | 12,14188763 | 2,959877573 | 0,858305696 | 2,60E-06 | 1,88E-05 | Up |
| Zfp383 | 209,7850735 | -0,712519131 | 0,147657691 | 7,53E-08 | 7,28E-07 | Down |

| | | | | | | |
|---|---|---|---|---|---|---|
| Cox7a1 | 131,1184365 | -0,723897024 | 0,190668721 | 6,53E-06 | 4,35E-05 | Down |
| Tyrobp | 8140,496444 | 0,674251954 | 0,178567975 | 1,03E-05 | 6,62E-05 | Up |
| Proser3 | 945,1244147 | -1,027935965 | 0,165430948 | 3,13E-12 | 6,22E-11 | Down |
| Hspb6 | 1201,830233 | -0,786316798 | 0,128803777 | 3,04E-11 | 5,13E-10 | Down |
| Tmem147 | 10504,35498 | 0,634006573 | 0,095754724 | 4,05E-12 | 7,84E-11 | Up |
| Dmkn | 61,86508586 | 1,472420212 | 0,508742757 | 1,10E-05 | 7,00E-05 | Up |
| Ffar2 | 2584,762228 | -1,130956261 | 0,519157078 | 0,00018097 | 0,000883705 | Down |
| Ffar3 | 7,045440282 | -0,87153697 | 0,682420548 | 0,002764127 | 0,010126076 | Down |
| Ffar1 | 8,299548859 | -1,886873409 | 1,096790903 | 0,000191386 | 0,000929938 | Down |
| Mag | 736,0866341 | 0,60449659 | 0,140426344 | 1,93E-06 | 1,43E-05 | Up |
| 2310043P16Rik | 228,3786929 | -0,753741734 | 0,159782448 | 9,07E-08 | 8,69E-07 | Down |
| Faap24 | 7631,35049 | 0,596096431 | 0,095208519 | 5,55E-11 | 9,01E-10 | Up |
| Cep89 | 2843,62441 | -0,797670214 | 0,175146445 | 1,39E-07 | 1,28E-06 | Down |
| Rgs9bp | 50,43011054 | -0,980656968 | 0,272527577 | 3,04E-06 | 2,16E-05 | Down |
| Ccne1 | 10878,0054 | 0,722352511 | 0,143314441 | 2,34E-08 | 2,48E-07 | Up |
| 2610021A01Rik | 2849,921361 | 0,665799634 | 0,081522552 | 2,94E-17 | 1,24E-15 | Up |
| Gm9268 | 4,783999837 | 0,762828372 | 0,947594347 | 0,006737811 | 0,022295599 | Up |
| Siglecf | 2157,366372 | -1,038865456 | 0,34432999 | 2,05E-05 | 0,000123272 | Down |
| Cd33 | 317,7481236 | 2,4192394 | 0,573683674 | 1,09E-07 | 1,02E-06 | Up |
| Nr1h2 | 9268,898884 | -0,588308379 | 0,067496694 | 4,76E-19 | 2,53E-17 | Down |
| Napsa | 27704,49886 | 0,624822561 | 0,114272907 | 5,17E-09 | 6,10E-08 | Up |
| Kcnc3 | 605,3869334 | 1,486676609 | 0,270941542 | 1,98E-10 | 2,93E-09 | Up |
| Gm44832 | 38,63610838 | 2,155485374 | 0,390150206 | 1,18E-10 | 1,81E-09 | Up |
| Vrk3 | 6681,944923 | -0,596747995 | 0,082225985 | 5,95E-14 | 1,57E-12 | Down |
| Rras | 783,3356231 | -0,663222631 | 0,20549756 | 7,91E-05 | 0,000417797 | Down |
| Ntf5 | 118,2839326 | -0,846541345 | 0,203865964 | 6,11E-07 | 5,02E-06 | Down |
| Gm10252 | 100,012985 | 0,828872614 | 0,229123752 | 6,23E-06 | 4,18E-05 | Up |
| Dhdh | 20,26135402 | 1,367682704 | 0,895902918 | 0,000513 | 0,002257447 | Up |
| Ppp1r15a | 9279,917644 | -0,750363274 | 0,099886107 | 2,46E-15 | 8,00E-14 | Down |
| Fut1 | 6,061453696 | 1,479751985 | 1,385022671 | 0,001065534 | 0,004344448 | Up |
| A030001D20Rik | 31,10105575 | -1,079946622 | 0,387358463 | 3,76E-05 | 0,000213825 | Down |
| Fut2 | 16,08528333 | 1,341752388 | 0,654469509 | 0,000151354 | 0,000751095 | Up |
| Kcnj14 | 93,30567782 | 1,997926631 | 0,352958143 | 4,54E-11 | 7,47E-10 | Up |
| Grwd1 | 15240,31023 | 0,868513546 | 0,134846934 | 1,70E-12 | 3,55E-11 | Up |
| Syngr4 | 51,83139779 | -0,5848584 | 0,221178193 | 0,000771296 | 0,003252981 | Down |
| Tmem143 | 1692,831018 | -0,780184358 | 0,085147538 | 1,60E-21 | 1,17E-19 | Down |
| Nomo1 | 42213,93297 | 0,844059786 | 0,111330089 | 5,82E-16 | 2,07E-14 | Up |
| Saa3 | 230,6534583 | -1,220173906 | 0,18060821 | 7,94E-14 | 2,05E-12 | Down |
| Saa4 | 11,52120595 | 4,98897334 | 1,299202746 | 3,68E-07 | 3,14E-06 | Up |
| Gtf2h1 | 19111,93618 | 0,893541931 | 0,119028468 | 6,76E-16 | 2,37E-14 | Up |
| Ldhc | 153,8158164 | -1,215262903 | 0,227212863 | 4,59E-10 | 6,42E-09 | Down |
| Uevld | 1935,29791 | -0,637812525 | 0,10338038 | 7,40E-11 | 1,18E-09 | Down |
| Zdhhc13 | 4032,751175 | 0,624819534 | 0,058369826 | 1,30E-27 | 1,88E-25 | Up |
| Gm2788 | 162,2014534 | -0,939412733 | 0,266647701 | 4,88E-06 | 3,35E-05 | Down |
| Gm44913 | 6,936059497 | 0,625729019 | 0,617382307 | 0,010422839 | 0,032351081 | Up |
| A230056P14Rik | 457,7275246 | -0,650770301 | 0,111550832 | 5,20E-10 | 7,20E-09 | Down |
| Gm44509 | 276,7010006 | -0,762797233 | 0,181395354 | 8,94E-07 | 7,12E-06 | Down |
| Herc2 | 17779,69212 | -0,9152638 | 0,128878937 | 1,20E-14 | 3,51E-13 | Down |
| Mphosph10 | 9516,09996 | 0,765523291 | 0,125088716 | 3,35E-11 | 5,61E-10 | Up |
| 1810008I18Rik | 31,5009811 | 3,575216476 | 0,507150807 | 1,80E-13 | 4,37E-12 | Up |
| Cers3 | 46,22677859 | 0,615665058 | 0,297722321 | 0,002511246 | 0,00932311 | Up |

| | | | | | | |
|---|---|---|---|---|---|---|
| Ttc23 | 62,53814804 | -0,688501443 | 0,251545707 | 0,0002982 | 0,001382517 | Down |
| Gm44686 | 19,61594527 | -0,777970846 | 0,45582922 | 0,002028381 | 0,007707512 | Down |
| Akap13 | 34518,55022 | -1,404243803 | 0,125911476 | 7,85E-31 | 1,65E-28 | Down |
| Gm44694 | 2,246424632 | -0,731635763 | 1,212699575 | 0,006274765 | 0,02093346 | Down |
| Aen | 10887,20713 | 1,099718773 | 0,124088062 | 3,82E-21 | 2,67E-19 | Up |
| Mfge8 | 989,6536238 | 0,744643492 | 0,159420588 | 1,22E-07 | 1,14E-06 | Up |
| Rhcg | 7,387391096 | 2,415140698 | 1,231027383 | 8,56E-05 | 0,000447765 | Up |
| Plin1 | 54,95853274 | 0,813062776 | 0,300157849 | 0,000150416 | 0,000747048 | Up |
| Pex11a | 937,1379236 | 0,673054077 | 0,151876775 | 6,67E-07 | 5,44E-06 | Up |
| Gm44974 | 16,87263608 | 1,368467676 | 0,552076618 | 4,53E-05 | 0,000252482 | Up |
| Gm21057 | 58,34856749 | -0,717836499 | 0,237974801 | 0,000106467 | 0,000544777 | Down |
| Furin | 16110,97693 | -0,602146883 | 0,160271579 | 1,85E-05 | 0,000112059 | Down |
| Gm15880 | 241,0916252 | -0,848210135 | 0,261707291 | 2,22E-05 | 0,000132144 | Down |
| Zscan2 | 576,059414 | -0,992107055 | 0,114210636 | 2,16E-20 | 1,39E-18 | Down |
| 1700010L04Rik | 60,69247392 | 0,828866365 | 0,233950186 | 8,29E-06 | 5,42E-05 | Up |
| Stard5 | 2301,646709 | 0,648563742 | 0,075477413 | 9,18E-19 | 4,66E-17 | Up |
| Mesdc2 | 12840,0392 | 0,644672429 | 0,082742562 | 7,25E-16 | 2,52E-14 | Up |
| Abhd17c | 18929,1166 | 1,561322607 | 0,123888558 | 1,97E-38 | 9,00E-36 | Up |
| Arnt2 | 849,7298932 | 3,206584928 | 0,36410878 | 1,62E-19 | 9,08E-18 | Up |
| Gm44556 | 10,50485091 | 1,760680513 | 1,183457709 | 0,000357988 | 0,001632354 | Up |
| Rab38 | 29661,05411 | -1,038033743 | 0,092542612 | 1,58E-31 | 3,44E-29 | Down |
| Gm45221 | 24,24808269 | -0,608842063 | 0,446802632 | 0,008489563 | 0,027151733 | Down |
| Ankrd42 | 38,73378453 | -0,897206586 | 0,35689348 | 0,000172705 | 0,000847376 | Down |
| Usp35 | 183,9917536 | -0,779557553 | 0,229791017 | 1,99E-05 | 0,000120045 | Down |
| Kctd21 | 361,1427177 | -0,846116355 | 0,128328816 | 7,38E-13 | 1,64E-11 | Down |
| Kctd14 | 149,1826973 | 0,747419606 | 0,246842203 | 8,43E-05 | 0,000441578 | Up |
| Aamdc | 625,5213932 | -0,797929146 | 0,134881854 | 8,79E-11 | 1,39E-09 | Down |
| Aqp11 | 254,6430333 | -0,651981632 | 0,134185442 | 1,04E-07 | 9,82E-07 | Down |
| Lrrc32 | 9,104524139 | 2,612125533 | 1,195133874 | 4,97E-05 | 0,00027453 | Up |
| Dgat2 | 116,9439592 | -0,711427328 | 0,343083777 | 0,001377188 | 0,005466249 | Down |
| Map6 | 2,803051762 | 2,3821112 | 1,998642603 | 0,000247555 | 0,001170599 | Up |
| Gdpd5 | 21,89444491 | 1,734324559 | 0,511150153 | 1,59E-06 | 1,20E-05 | Up |
| Kcne3 | 7,950813995 | 1,371286829 | 0,948913375 | 0,000612498 | 0,002643064 | Up |
| P2ry6 | 4,885679306 | 0,627705257 | 0,66715501 | 0,010754911 | 0,033239066 | Up |
| Stard10 | 87,27593361 | -0,650375862 | 0,252335262 | 0,000594894 | 0,002576839 | Down |
| Gm45620 | 11,67618121 | -0,732146706 | 0,507759687 | 0,00392284 | 0,013814665 | Down |
| Gm45380 | 74,23343444 | 0,653090215 | 0,237990034 | 0,000370602 | 0,001681475 | Up |
| Folr1 | 1260,518609 | 0,713550783 | 0,24664712 | 0,000160101 | 0,000790422 | Up |
| Numa1 | 38235,15433 | -0,639554304 | 0,084357558 | 3,86E-15 | 1,22E-13 | Down |
| Pgap2 | 8400,764806 | 0,604566438 | 0,106778362 | 1,98E-09 | 2,50E-08 | Up |
| Trim21 | 770,4738521 | -0,790206245 | 0,11423603 | 1,32E-13 | 3,28E-12 | Down |
| Usp17ld | 816,2852816 | 4,766806726 | 0,20287019 | 1,26E-123 | 2,31E-119 | Up |
| Usp17lc | 25,65547678 | 6,556436753 | 1,265392082 | 1,83E-10 | 2,73E-09 | Up |
| Hbb-y | 20,29731936 | -2,043444604 | 1,435780386 | 0,000316396 | 0,001457991 | Down |
| Trim34a | 208,7301542 | -0,917414254 | 0,14363696 | 1,69E-12 | 3,54E-11 | Down |
| Trim34b | 146,991542 | -0,710243984 | 0,164741548 | 8,51E-07 | 6,80E-06 | Down |
| Trim30b | 36,33819081 | -0,605786642 | 0,316472596 | 0,003650788 | 0,012988639 | Down |
| Trim30d | 91,68218026 | -0,600916579 | 0,307339291 | 0,00347668 | 0,012434614 | Down |
| Rrp8 | 10068,89848 | 0,718683225 | 0,106123213 | 7,13E-13 | 1,59E-11 | Up |
| Gm4070 | 372,7722627 | 1,601133809 | 0,23877142 | 1,07E-13 | 2,69E-12 | Up |
| Gm18852 | 8,760042199 | 1,175493791 | 0,867033987 | 0,00104641 | 0,004276021 | Up |

| | | | | | | |
|---|---|---|---|---|---|---|
| 4930431P19Rik | 74,04228373 | -0,780672255 | 0,255321301 | 6,19E-05 | 0,00033612 | Down |
| RP23-462D5,5 | 106,5448752 | -0,723736239 | 0,228776686 | 6,38E-05 | 0,000345261 | Down |
| Mrvi1 | 556,50059 | 1,170070209 | 0,540050557 | 0,000167033 | 0,000822641 | Up |
| Gm4353 | 904,7235362 | 0,615973535 | 0,094141111 | 7,77E-12 | 1,45E-10 | Up |
| Knop1 | 19099,91644 | 0,67688994 | 0,095419719 | 1,07E-13 | 2,69E-12 | Up |
| Lyrm1 | 176,1040581 | 0,744887895 | 0,164549591 | 2,42E-07 | 2,12E-06 | Up |
| Dnah3 | 9,939820027 | 2,148615421 | 0,944798789 | 4,38E-05 | 0,000244968 | Up |
| Eef2k | 13910,61727 | -0,58224956 | 0,151746652 | 1,54E-05 | 9,53E-05 | Down |
| Cdr2 | 4249,818112 | 0,88056153 | 0,076915192 | 2,69E-32 | 6,56E-30 | Up |
| Igsf6 | 1312,914639 | 0,631654201 | 0,201725761 | 0,000134638 | 0,000674557 | Up |
| Gm20274 | 275,0519533 | 1,47206898 | 0,228994229 | 7,86E-13 | 1,74E-11 | Up |
| Il4ra | 15006,45332 | 0,768273695 | 0,110867309 | 1,49E-13 | 3,69E-12 | Up |
| Apobr | 3806,992693 | 0,830691654 | 0,112248567 | 2,63E-15 | 8,51E-14 | Up |
| Nupr1 | 765,1048752 | 1,910409205 | 0,663306519 | 8,07E-06 | 5,29E-05 | Up |
| Ypel3 | 612,7755544 | -0,956904661 | 0,393397548 | 0,000167854 | 0,000826017 | Down |
| Kctd13 | 2741,270034 | 0,680235422 | 0,08042864 | 2,26E-18 | 1,11E-16 | Up |
| D830044I16Rik | 223,7556018 | -0,789759002 | 0,186603975 | 6,46E-07 | 5,27E-06 | Down |
| Mylpf | 586,579511 | -0,99104951 | 0,133352612 | 6,69E-16 | 2,35E-14 | Down |
| Gm44728 | 15,19890685 | -0,60089255 | 0,407790773 | 0,007771918 | 0,025156109 | Down |
| Itgal | 16604,85612 | 0,603758704 | 0,143630268 | 3,00E-06 | 2,14E-05 | Up |
| Ccdc189 | 34,1889125 | -0,93437308 | 0,33343416 | 6,20E-05 | 0,000336504 | Down |
| Gm45205 | 5,15658734 | 0,695391413 | 0,924154872 | 0,008696509 | 0,027694483 | Up |
| Itgam | 2870,78632 | 1,330024037 | 0,223901457 | 1,61E-11 | 2,85E-10 | Up |
| Itgax | 157,6541338 | -1,721325671 | 0,348434169 | 2,47E-09 | 3,05E-08 | Down |
| Cox6a2 | 153,6528533 | 2,028208758 | 0,80068187 | 2,21E-05 | 0,000132053 | Up |
| Hmx2 | 1383,316542 | -0,58367049 | 0,104269375 | 3,27E-09 | 3,94E-08 | Down |
| 1500002F19Rik | 43,92224363 | 1,370941326 | 0,405908439 | 2,62E-06 | 1,89E-05 | Up |
| Lrrc27 | 225,731834 | 0,7585154 | 0,169074873 | 2,61E-07 | 2,28E-06 | Up |
| Gm45329 | 50,42146339 | 0,592571743 | 0,331686063 | 0,005032364 | 0,01720993 | Up |
| Cfap46 | 27,52282962 | -0,842380444 | 0,404658476 | 0,000679348 | 0,002904085 | Down |
| Adam8 | 724,7217499 | 3,230032263 | 0,39987839 | 7,44E-17 | 2,97E-15 | Up |
| Ifitm2 | 7555,364692 | 0,590277751 | 0,107455856 | 5,65E-09 | 6,65E-08 | Up |
| Ifitm1 | 142241,0703 | 1,080532449 | 0,169923133 | 1,09E-12 | 2,35E-11 | Up |
| Gm7514 | 58,25403277 | 2,850705442 | 0,33270377 | 6,26E-19 | 3,25E-17 | Up |
| Ifitm3 | 93618,87326 | 2,715553665 | 0,273878324 | 1,63E-24 | 1,81E-22 | Up |
| Ifitm6 | 396,4509885 | 5,209447133 | 0,695440447 | 7,04E-16 | 2,45E-14 | Up |
| Lrrc56 | 288,3536262 | -1,379474294 | 0,164308166 | 3,94E-19 | 2,12E-17 | Down |
| Cdhr5 | 60,10094114 | 0,714926983 | 0,322146517 | 0,000968422 | 0,003992172 | Up |
| Slc25a22 | 4130,3119 | 0,998785793 | 0,145359699 | 4,02E-14 | 1,09E-12 | Up |
| Cracr2b | 146,9201885 | -0,713459532 | 0,218248475 | 4,82E-05 | 0,000267155 | Down |
| Tspan4 | 10535,26859 | 1,059182005 | 0,09978331 | 1,19E-28 | 1,95E-26 | Up |
| Muc6 | 10,9192414 | -2,049167251 | 0,846902517 | 3,02E-05 | 0,000174959 | Down |
| Dusp8 | 13,96681384 | 3,454101417 | 0,985396695 | 2,52E-06 | 1,81E-05 | Up |
| Tnni2 | 70,68094249 | 0,895015999 | 0,273642128 | 1,55E-05 | 9,55E-05 | Up |
| Trpm5 | 32,60771993 | 1,111267275 | 0,363372362 | 1,42E-05 | 8,84E-05 | Up |
| E230032D23Rik | 70,81691303 | -0,722430185 | 0,310094484 | 0,000709436 | 0,003018573 | Down |
| Tnfrsf23 | 77,78628846 | 1,155905564 | 0,372085294 | 1,06E-05 | 6,81E-05 | Up |
| Osbpl5 | 159,6143773 | -0,991179215 | 0,425792255 | 0,000195086 | 0,000944023 | Down |
| Dhcr7 | 10305,01746 | -0,600961276 | 0,161918588 | 2,21E-05 | 0,000132007 | Down |
| Cttn | 155,5479725 | -0,962885501 | 0,3505375 | 6,53E-05 | 0,000351471 | Down |
| A430078G23Rik | 2138,522 | -0,963467297 | 0,125298465 | 1,03E-16 | 4,05E-15 | Down |

| | | | | | | |
|---|---|---|---|---|---|---|
| Arhgef18 | 7935,683872 | -0,94784198 | 0,101139808 | 5,02E-23 | 4,56E-21 | Down |
| Pcp2 | 5,360082991 | -0,739090271 | 0,812184279 | 0,00686197 | 0,022664731 | Down |
| Retn | 803,0730145 | 1,189655116 | 0,282275588 | 1,29E-07 | 1,20E-06 | Up |
| Cd209d | 4,226322041 | 0,598056068 | 0,731404475 | 0,01297078 | 0,039010602 | Up |
| 3930402G23Rik | 4,265417362 | 1,010055211 | 1,292443204 | 0,00324041 | 0,011690315 | Up |
| Irs2 | 33,97500087 | 1,274984435 | 0,447635573 | 1,84E-05 | 0,000111906 | Up |
| Col4a2 | 1922,424638 | 2,451435697 | 0,170668781 | 1,38E-48 | 1,05E-45 | Up |
| Mcf2l | 271,8840191 | 1,758591523 | 0,827001893 | 7,44E-05 | 0,00039457 | Up |
| F7 | 73,06081911 | 1,69524542 | 0,69450108 | 3,27E-05 | 0,000188086 | Up |
| Adprhl1 | 11,71580451 | -2,757357506 | 0,79801308 | 2,25E-06 | 1,64E-05 | Down |
| 9530085L11Rik | 133,4593061 | -0,674708543 | 0,1524801 | 6,78E-07 | 5,52E-06 | Down |
| Gm7676 | 30,2888419 | 1,383289948 | 0,393785225 | 1,58E-06 | 1,20E-05 | Up |
| Agpat5 | 11830,5872 | 0,699723995 | 0,103143246 | 7,81E-13 | 1,73E-11 | Up |
| Htra4 | 3,06300829 | 1,012721056 | 1,377809461 | 0,003065188 | 0,011110852 | Up |
| Adgra2 | 117,7847931 | -0,804802745 | 0,210314718 | 3,21E-06 | 2,28E-05 | Down |
| Mak16 | 6174,166411 | 0,713980757 | 0,08479956 | 2,30E-18 | 1,12E-16 | Up |
| Rbpms | 4998,03979 | -0,754990815 | 0,164825484 | 1,73E-07 | 1,55E-06 | Down |
| Gm29243 | 387,7691212 | 0,849159846 | 0,172131599 | 1,44E-08 | 1,57E-07 | Up |
| Gm45347 | 8,981262367 | -0,597079848 | 0,551697667 | 0,011632478 | 0,03553378 | Down |
| Tnks | 13087,15931 | -0,723057231 | 0,073041795 | 2,38E-24 | 2,58E-22 | Down |
| Dctd | 5876,214349 | 0,694311142 | 0,098042659 | 1,01E-13 | 2,56E-12 | Up |
| Asb5 | 24,23544199 | -0,786646085 | 0,37834229 | 0,000900247 | 0,003742389 | Down |
| Tma16 | 3261,515626 | 0,949456579 | 0,133833566 | 1,02E-14 | 3,04E-13 | Up |
| RP23-451G9,2 | 2,933032362 | 0,775817887 | 1,184119393 | 0,0060041 | 0,020118692 | Up |
| Gm9755 | 218,8304035 | 0,601372975 | 0,157831256 | 1,51E-05 | 9,37E-05 | Up |
| Slc18a1 | 40,47516133 | -0,61511681 | 0,291290085 | 0,00229557 | 0,008615227 | Down |
| Zfp963 | 93,20162276 | -0,735440912 | 0,198086412 | 8,29E-06 | 5,42E-05 | Down |
| Rfxank | 1188,777364 | -0,605456283 | 0,080524515 | 7,94E-15 | 2,43E-13 | Down |
| Crtc1 | 11770,38755 | -0,613022216 | 0,093591644 | 7,57E-12 | 1,42E-10 | Down |
| Gdf15 | 14,31957586 | 0,582857201 | 0,518986794 | 0,012289548 | 0,037206851 | Up |
| Jund | 3265,448104 | 0,700378558 | 0,203247333 | 2,87E-05 | 0,000166888 | Up |
| C430049E01Rik | 6,525434205 | 1,120138551 | 0,897681159 | 0,001450239 | 0,005730044 | Up |
| Bst2 | 8503,342442 | 0,802542666 | 0,115289785 | 8,58E-14 | 2,20E-12 | Up |
| Olfr373 | 18,79907821 | 0,602131487 | 0,421213878 | 0,008161999 | 0,026251219 | Up |
| Tpm4 | 35,46563035 | 0,762016368 | 0,390527165 | 0,001356022 | 0,005390439 | Up |
| Tmem38a | 930,6816207 | -0,631589914 | 0,134775309 | 2,79E-07 | 2,43E-06 | Down |
| Nwd1 | 3134,380359 | -1,815461058 | 0,310905766 | 1,70E-11 | 3,00E-10 | Down |
| F2rl3 | 623,7087597 | 0,796093269 | 0,170493629 | 8,11E-08 | 7,81E-07 | Up |
| Hmox1 | 11159,22222 | 0,612890963 | 0,195793245 | 0,000153608 | 0,00076166 | Up |
| Gm7984 | 118,695457 | 0,688456038 | 0,179488499 | 7,33E-06 | 4,85E-05 | Up |
| Smad1 | 116,2128646 | -1,122616938 | 0,319174477 | 2,63E-06 | 1,89E-05 | Down |
| Abce1 | 41006,83942 | 0,624721741 | 0,067229491 | 1,98E-21 | 1,43E-19 | Up |
| Il15 | 77,27478044 | -1,443699335 | 0,413122328 | 1,54E-06 | 1,16E-05 | Down |
| Gm10644 | 47,0296698 | -1,252626523 | 0,342229899 | 1,15E-06 | 8,90E-06 | Down |
| Prkaca | 16556,11722 | -0,587121995 | 0,100238068 | 7,05E-10 | 9,53E-09 | Down |
| C330011M18Rik | 85,67927948 | -0,618321118 | 0,20271429 | 0,00019033 | 0,000925892 | Down |
| Ier2 | 10364,82877 | 0,633373903 | 0,103833053 | 1,18E-10 | 1,82E-09 | Up |
| G430095P16Rik | 558,6620452 | -0,682098674 | 0,102928146 | 2,65E-12 | 5,31E-11 | Down |
| Klf1 | 812,5738791 | -1,453788611 | 0,125841541 | 8,31E-33 | 2,11E-30 | Down |
| Junb | 6396,057393 | 0,629758125 | 0,101233215 | 5,69E-11 | 9,22E-10 | Up |
| Gm42031 | 152,1581822 | -0,620960229 | 0,239250011 | 0,00069039 | 0,00294646 | Down |

| | | | | | | |
|---|---|---|---|---|---|---|
| Gpt2 | 11447,41538 | 0,876706281 | 0,19012755 | 5,96E-08 | 5,85E-07 | Up |
| Gm45425 | 35,99974872 | 1,637897951 | 0,367722732 | 2,62E-08 | 2,75E-07 | Up |
| Heatr3 | 19515,88278 | 0,737277844 | 0,090912162 | 2,46E-17 | 1,06E-15 | Up |
| Nod2 | 319,3092233 | -0,896077319 | 0,142135732 | 3,38E-12 | 6,61E-11 | Down |
| Rbl2 | 4418,67709 | -0,63360542 | 0,098530573 | 1,44E-11 | 2,56E-10 | Down |
| Bbs2 | 490,3657757 | -0,694942261 | 0,10934015 | 1,42E-11 | 2,53E-10 | Down |
| Adgrg1 | 224934,7665 | -0,813323281 | 0,099288086 | 5,85E-18 | 2,70E-16 | Down |
| Drc7 | 87,03787323 | -3,061836484 | 0,36053953 | 1,98E-18 | 9,73E-17 | Down |
| Katnb1 | 3828,962268 | -0,623228202 | 0,060536581 | 9,97E-26 | 1,22E-23 | Down |
| Kifc3 | 1942,482142 | -1,38316257 | 0,201706048 | 4,74E-14 | 1,27E-12 | Down |
| Ces2g | 341,3859819 | 1,602366564 | 0,647770433 | 3,25E-05 | 0,000186927 | Up |
| Lrrc29 | 453,0647245 | -0,891410281 | 0,091423924 | 1,94E-24 | 2,12E-22 | Down |
| Slc9a5 | 593,7840745 | -0,638259638 | 0,142697403 | 7,19E-07 | 5,82E-06 | Down |
| Lrrc36 | 24,50060593 | -0,989235068 | 0,368762876 | 7,16E-05 | 0,000382063 | Down |
| Fam65a | 21843,04541 | -0,944929576 | 0,155883151 | 1,17E-11 | 2,11E-10 | Down |
| Carmil2 | 2179,795262 | -0,796279483 | 0,151720931 | 4,14E-09 | 4,94E-08 | Down |
| Ctrl | 102,9559872 | 0,903885695 | 0,356975445 | 0,000159161 | 0,000786842 | Up |
| Prmt7 | 12831,48533 | 0,583095554 | 0,071170594 | 4,30E-17 | 1,76E-15 | Up |
| Cirh1a | 18575,52838 | 0,986213521 | 0,090917868 | 1,13E-29 | 2,05E-27 | Up |
| Nip7 | 12254,68966 | 0,897507561 | 0,108738173 | 1,61E-18 | 8,04E-17 | Up |
| Nqo1 | 88,86985884 | -0,708624067 | 0,309891828 | 0,000856243 | 0,003575741 | Down |
| Pmfbp1 | 2,589408726 | -0,781032343 | 1,234733736 | 0,005563538 | 0,018786658 | Down |
| Cmtr2 | 3786,491526 | 0,594414141 | 0,096989667 | 1,29E-10 | 1,97E-09 | Up |
| Ldhd | 15,42549047 | -2,243071031 | 0,696168861 | 3,04E-06 | 2,16E-05 | Down |
| Adat1 | 2595,152039 | 0,614263121 | 0,085107794 | 7,10E-14 | 1,85E-12 | Up |
| Nudt7 | 3037,555588 | -0,721953258 | 0,110344721 | 3,27E-12 | 6,44E-11 | Down |
| 1700030J22Rik | 40,17257116 | -0,929429968 | 0,320434407 | 4,63E-05 | 0,000257736 | Down |
| Gcsh | 7466,543028 | 0,627191369 | 0,114273693 | 4,53E-09 | 5,39E-08 | Up |
| Mlycd | 2455,726043 | -0,580541716 | 0,134074969 | 2,01E-06 | 1,48E-05 | Down |
| Osgin1 | 362,5784168 | 1,186682462 | 0,228681306 | 1,10E-09 | 1,44E-08 | Up |
| Tldc1 | 1064,837869 | -0,628336089 | 0,103136409 | 1,28E-10 | 1,96E-09 | Down |
| Cotl1 | 639,3924189 | 1,410587595 | 0,234845175 | 1,09E-11 | 1,98E-10 | Up |
| Gse1 | 29886,72165 | -1,116987896 | 0,114504146 | 8,63E-25 | 9,98E-23 | Down |
| 1700016A09Rik | 37,75277305 | -1,848528659 | 0,368428923 | 1,52E-09 | 1,95E-08 | Down |
| Gm10614 | 22,11297397 | -0,613513296 | 0,359903505 | 0,005077227 | 0,017347112 | Down |
| Irf8 | 26,07030485 | -0,725762242 | 0,556547478 | 0,004863041 | 0,016696531 | Down |
| 1700030M09Rik | 6,085666463 | 0,80848612 | 1,079484468 | 0,005844117 | 0,019622223 | Up |
| Gm26812 | 25,95583854 | 1,636972224 | 0,492965789 | 2,21E-06 | 1,61E-05 | Up |
| Slc7a5 | 89433,40736 | 0,875596403 | 0,119500122 | 3,06E-15 | 9,78E-14 | Up |
| Gm27011 | 34,78370778 | -1,366419348 | 0,364344632 | 6,71E-07 | 5,47E-06 | Down |
| Gm17709 | 80,27379135 | 1,034253714 | 0,272858404 | 1,15E-06 | 8,90E-06 | Up |
| Zfpm1 | 21567,51357 | -0,90322091 | 0,119850217 | 5,01E-16 | 1,81E-14 | Down |
| Gm45353 | 43,3470747 | -2,449704783 | 0,397996052 | 6,18E-12 | 1,17E-10 | Down |
| Cyba | 21002,84296 | 0,793688636 | 0,143628353 | 9,04E-10 | 1,19E-08 | Up |
| Rnf166 | 3759,37705 | -0,718130173 | 0,108286958 | 1,87E-12 | 3,86E-11 | Down |
| Piezo1 | 10592,40236 | 0,58299451 | 0,078356288 | 1,66E-14 | 4,75E-13 | Up |
| Cpne7 | 3,727745369 | 0,627569638 | 0,870705246 | 0,011419036 | 0,034994871 | Up |
| 4732419C18Rik | 13,91751017 | 0,904218371 | 0,606505746 | 0,001734693 | 0,006703124 | Up |
| Rab4a | 2004,414781 | -0,758265089 | 0,07935133 | 4,88E-23 | 4,45E-21 | Down |
| RP24-246F6,2 | 76,25263971 | 0,685218577 | 0,419478921 | 0,003742261 | 0,013289584 | Up |
| Urb2 | 19357,13824 | 0,784244955 | 0,091829314 | 4,07E-19 | 2,18E-17 | Up |

| | | | | | | |
|---|---|---|---|---|---|---|
| Arv1 | 875,0851008 | 0,591413281 | 0,117330143 | 6,35E-08 | 6,22E-07 | Up |
| RP23-312B17,3 | 98,75736627 | -0,790897243 | 0,253102337 | 4,65E-05 | 0,000258577 | Down |
| Mmp8 | 19,57039496 | 1,016550867 | 0,939097365 | 0,002437386 | 0,009080756 | Up |
| Maml2 | 1577,826102 | -1,387025185 | 0,153058141 | 1,19E-21 | 8,80E-20 | Down |
| Ppan | 19983,65447 | 0,615984469 | 0,159169688 | 1,09E-05 | 6,93E-05 | Up |
| Icam1 | 2167,985081 | 0,714612491 | 0,123784122 | 4,37E-10 | 6,14E-09 | Up |
| RP23-186O3,13 | 19,89286091 | 0,81103121 | 0,548533827 | 0,002573727 | 0,009522165 | Up |
| RP24-496O17,7 | 26,46044764 | -0,770408725 | 0,354788262 | 0,000791824 | 0,003332634 | Down |
| Ecsit | 6807,228046 | 0,66968304 | 0,065876981 | 2,63E-25 | 3,14E-23 | Up |
| Zfp599 | 1511,601738 | 0,628960267 | 0,112936921 | 2,84E-09 | 3,47E-08 | Up |
| RP23-187B11,11 | 34,10967491 | -0,767291435 | 0,386345756 | 0,001224528 | 0,004914796 | Down |
| 9530077C05Rik | 70,95159107 | -0,959035013 | 0,305706162 | 1,84E-05 | 0,000111891 | Down |
| Bbs9 | 1720,563298 | -0,762218469 | 0,094596158 | 2,94E-17 | 1,24E-15 | Down |
| Fam118b | 4373,43634 | 0,627035734 | 0,076608654 | 3,44E-17 | 1,43E-15 | Up |
| Ddx25 | 19,39925116 | -2,317174342 | 1,073891039 | 5,68E-05 | 0,000310831 | Down |
| Pknox2 | 56,51895352 | 3,777708185 | 0,571542468 | 3,21E-12 | 6,34E-11 | Up |
| Robo3 | 29,5167106 | 4,072451678 | 0,529116203 | 1,05E-15 | 3,57E-14 | Up |
| Vsig2 | 48,79650489 | -1,143037773 | 0,359035291 | 8,44E-06 | 5,51E-05 | Down |
| Nrgn | 5176,970255 | -1,489635894 | 0,170292566 | 1,91E-20 | 1,24E-18 | Down |
| Spa17 | 115,0619206 | -0,677268058 | 0,169558849 | 4,23E-06 | 2,94E-05 | Down |
| Hspa8 | 409222,1768 | 0,723564302 | 0,076430144 | 1,62E-22 | 1,34E-20 | Up |
| RP24-175C20,15 | 71,69740683 | 1,108133183 | 0,383239954 | 2,48E-05 | 0,000145973 | Up |
| RP24-175C20,19 | 351,9516438 | -0,659429059 | 0,156166101 | 1,86E-06 | 1,38E-05 | Down |
| Cxcr5 | 38,76873695 | -0,899481046 | 0,354324767 | 0,000159394 | 0,00078778 | Down |
| RP24-282C4,4 | 30,73858901 | -1,126285588 | 0,522284233 | 0,000194055 | 0,00094001 | Down |
| Dmxl2 | 12820,73825 | 0,638679561 | 0,151326566 | 2,19E-06 | 1,60E-05 | Up |
| Idh3a | 38695,69638 | 0,675477732 | 0,104973099 | 1,00E-11 | 1,83E-10 | Up |
| Acsbg1 | 13,55163834 | 1,708849421 | 0,834456578 | 9,49E-05 | 0,000490647 | Up |
| Rcn2 | 6205,014351 | 0,657818322 | 0,100383397 | 5,32E-12 | 1,01E-10 | Up |
| Cspg4 | 24,4024247 | 2,078877162 | 0,532748176 | 2,46E-07 | 2,15E-06 | Up |
| Ptpn9 | 8439,030459 | 0,682482477 | 0,084098313 | 3,91E-17 | 1,62E-15 | Up |
| Gm10658 | 113,396517 | -0,813318814 | 0,177616884 | 1,09E-07 | 1,03E-06 | Down |
| 2700012I20Rik | 49,62703122 | -1,02192756 | 0,321046498 | 1,23E-05 | 7,76E-05 | Down |
| Lman1l | 22,96843767 | 2,337976819 | 0,454105353 | 1,31E-09 | 1,70E-08 | Up |
| Stra6 | 22,08416683 | 3,87175297 | 0,998257081 | 9,05E-07 | 7,20E-06 | Up |
| 1600029O15Rik | 4,967432649 | 0,639237745 | 0,915306566 | 0,01079443 | 0,033338047 | Up |
| Cd276 | 489,4175603 | 3,621151802 | 0,25798389 | 1,45E-45 | 9,11E-43 | Up |
| Adpgk | 8387,369047 | 0,90062174 | 0,130292745 | 5,20E-14 | 1,38E-12 | Up |
| Gramd2 | 15,68022229 | 1,053025915 | 0,598512217 | 0,000617594 | 0,002661607 | Up |
| Uaca | 470,6194136 | 0,832300307 | 0,386609332 | 0,000602636 | 0,002606047 | Up |
| RP23-9A12,4 | 5,547729581 | 2,141435831 | 1,626379764 | 0,000332607 | 0,001526359 | Up |
| RP23-116C19,1 | 234,4387363 | 1,107534648 | 0,322564832 | 3,76E-06 | 2,64E-05 | Up |
| Tle3 | 102,443084 | 1,217848643 | 0,354489357 | 2,86E-06 | 2,05E-05 | Up |
| Hacd3 | 17050,06922 | 0,741298032 | 0,108092671 | 3,19E-13 | 7,50E-12 | Up |
| Parp16 | 265,9383329 | -0,88909331 | 0,146739083 | 1,71E-11 | 3,01E-10 | Down |
| Car12 | 3838,001602 | 1,304603381 | 0,115251659 | 9,84E-32 | 2,19E-29 | Up |
| C2cd4a | 4,008515755 | 0,869085786 | 1,037709248 | 0,004746898 | 0,016350029 | Up |
| Gm26849 | 2,566556236 | 1,746265029 | 1,842152468 | 0,000666157 | 0,002854378 | Up |
| Gm10642 | 31,73537749 | -1,812025437 | 0,546311758 | 1,99E-06 | 1,47E-05 | Down |
| Aqp9 | 2236,450141 | 3,044468218 | 0,3833366 | 1,67E-16 | 6,39E-15 | Up |
| RP23-2F19,2 | 3,19749501 | 0,903035167 | 1,065192957 | 0,004284084 | 0,014927871 | Up |

| | | | | | | |
|---|---|---|---|---|---|---|
| Gm37234 | 51,52794593 | 1,764452554 | 0,530614664 | 1,97E-06 | 1,46E-05 | Up |
| Ccpg1os | 184,6730959 | -0,830364222 | 0,13570252 | 1,88E-11 | 3,29E-10 | Down |
| Rsl24d1 | 11831,58297 | 0,651537568 | 0,093808674 | 3,80E-13 | 8,79E-12 | Up |
| Gm38215 | 5,683671131 | 0,656663826 | 0,75767616 | 0,009790526 | 0,030673286 | Up |
| RP24-365N15,9 | 171,8654634 | 0,690171788 | 0,669631637 | 0,007589905 | 0,024641217 | Up |
| Gsta4 | 475,055393 | -1,338381791 | 0,153671353 | 2,54E-20 | 1,62E-18 | Down |
| Sh3bgrl2 | 22,9565093 | 2,661323549 | 0,644607283 | 2,27E-07 | 2,00E-06 | Up |
| Bcl2a1d | 151,2978307 | 2,394487853 | 0,273393459 | 1,71E-20 | 1,11E-18 | Up |
| Gm2396 | 8,488661895 | 0,87813447 | 0,718011566 | 0,002927208 | 0,010665736 | Up |
| Bcl2a1a | 3219,401871 | 2,388790872 | 0,324866971 | 1,47E-15 | 4,92E-14 | Up |
| Gm37833 | 2,307976081 | 1,427148669 | 1,649804448 | 0,001223883 | 0,004913287 | Up |
| Bcl2a1b | 1443,482025 | 2,397807816 | 0,242356988 | 4,17E-25 | 4,91E-23 | Up |
| Pcolce2 | 191,0624723 | -1,40714899 | 0,249896907 | 9,50E-11 | 1,49E-09 | Down |
| Pls1 | 3170,196103 | -3,334799728 | 0,199345027 | 1,55E-63 | 2,36E-60 | Down |
| 1700065D16Rik | 152,2227638 | -2,265157666 | 0,23508037 | 2,75E-24 | 2,95E-22 | Down |
| Rasa2 | 3208,881562 | 1,139590045 | 0,122626129 | 7,68E-23 | 6,74E-21 | Up |
| 4930579K19Rik | 40,89656886 | -0,583756247 | 0,286140376 | 0,00327522 | 0,011792599 | Down |
| Gm37621 | 38,92001387 | 0,832011527 | 0,417846217 | 0,000876707 | 0,003653679 | Up |
| Ky | 967,2072409 | 2,109723165 | 0,145708698 | 4,11E-50 | 3,76E-47 | Up |
| Ryk | 47,51712129 | -1,36713552 | 0,380185099 | 1,20E-06 | 9,25E-06 | Down |
| Srprb | 8688,194461 | 0,832645358 | 0,163527104 | 7,11E-09 | 8,22E-08 | Up |
| Cdv3 | 39299,34711 | 0,788662698 | 0,080912894 | 5,51E-24 | 5,56E-22 | Up |
| Bfsp2 | 56,3096011 | 3,041472735 | 0,310418524 | 1,33E-23 | 1,28E-21 | Up |
| Tlr9 | 120,2603843 | 2,079838807 | 0,542047485 | 3,16E-07 | 2,73E-06 | Up |
| Alas1 | 5364,66239 | 1,461998622 | 0,119854491 | 3,80E-36 | 1,33E-33 | Up |
| Dusp7 | 2854,224814 | 1,342119138 | 0,08582206 | 4,95E-57 | 6,46E-54 | Up |
| Acy1 | 10122,32125 | 0,905206304 | 0,079154891 | 2,51E-32 | 6,18E-30 | Up |
| Rrp9 | 13275,934 | 1,202124057 | 0,099215252 | 5,65E-36 | 1,91E-33 | Up |
| Mapkapk3 | 18249,63982 | 1,475306581 | 0,174725671 | 2,64E-19 | 1,45E-17 | Up |
| Gm17040 | 22,76947498 | 1,139672741 | 0,577587065 | 0,000293638 | 0,001364134 | Up |
| Cish | 14896,70043 | 3,160114065 | 0,150625832 | 1,99E-98 | 1,82E-94 | Up |
| Hemk1 | 5617,126081 | 1,648516536 | 0,152737235 | 2,40E-29 | 4,17E-27 | Up |
| 6430571L13Rik | 340,745674 | 2,110663329 | 0,191847093 | 1,02E-30 | 2,09E-28 | Up |
| Hyal2 | 3828,837013 | 0,887436524 | 0,094413698 | 6,05E-23 | 5,47E-21 | Up |
| Nat6 | 3196,68408 | -0,794490049 | 0,189637767 | 7,52E-07 | 6,07E-06 | Down |
| Ifrd2 | 22895,94146 | 0,805744882 | 0,076822182 | 2,36E-27 | 3,37E-25 | Up |
| Lsmem2 | 133,6128646 | 0,791902094 | 0,316465735 | 0,000303564 | 0,001405601 | Up |
| Gm38150 | 138,0026428 | 0,998203824 | 0,313213487 | 1,32E-05 | 8,28E-05 | Up |
| Bsn | 1393,889029 | 0,935945823 | 0,10031361 | 8,03E-23 | 7,02E-21 | Up |
| 4930447F24Rik | 32,12314476 | 0,656358993 | 0,319158306 | 0,001988592 | 0,00756734 | Up |
| Nicn1 | 1482,958208 | -0,870364219 | 0,107820349 | 9,15E-18 | 4,15E-16 | Down |
| 4833445I07Rik | 115,5348381 | -0,613258611 | 0,21647933 | 0,000377248 | 0,00170993 | Down |
| Nme6 | 4117,500967 | 0,734417585 | 0,087554406 | 2,47E-18 | 1,20E-16 | Up |
| Cdc25a | 15366,18591 | 0,644995866 | 0,084471446 | 2,44E-15 | 7,96E-14 | Up |
| Gm42432 | 31,82440315 | 1,189979121 | 0,538836387 | 0,000142111 | 0,000709077 | Up |
| Gm42762 | 74,11714831 | -2,145988723 | 0,330674082 | 3,04E-13 | 7,19E-12 | Down |
| Ltf | 278,6434042 | 1,101768005 | 0,370770438 | 1,95E-05 | 0,00011772 | Up |
| Arpp21 | 45,49730575 | -1,386419285 | 0,300418778 | 1,70E-08 | 1,84E-07 | Down |
| 2310075C17Rik | 28,0468072 | -0,747255012 | 0,405247793 | 0,001825335 | 0,007020696 | Down |
| Bcl2a1c | 7,99021949 | 2,064619008 | 1,114086303 | 0,000126778 | 0,000638153 | Up |
| Ccr4 | 3470,273775 | 1,175987696 | 0,170289793 | 2,66E-14 | 7,42E-13 | Up |

| | | | | | | |
|---|---|---|---|---|---|---|
| Gm9888 | 933,0397109 | -0,641234118 | 0,152644974 | 2,34E-06 | 1,70E-05 | Down |
| Cx3cr1 | 25,89732091 | 0,672264265 | 0,58104412 | 0,007394576 | 0,024122857 | Up |
| Ttc21a | 268,5701923 | 0,793267246 | 0,165580456 | 4,57E-08 | 4,59E-07 | Up |
| Csrnp1 | 4757,737723 | 0,702077032 | 0,121617677 | 4,88E-10 | 6,80E-09 | Up |
| RP24-233B16,1 | 11,75484017 | 1,091732758 | 0,859855996 | 0,001484285 | 0,00584433 | Up |
| RP23-366E1,3 | 77,63750656 | -0,667095094 | 0,210509974 | 9,31E-05 | 0,000482438 | Down |
| RP24-264F17,3 | 10,5827608 | -0,786710064 | 0,661444101 | 0,004290293 | 0,014940759 | Down |
| RP23-242K3,2 | 9,736395035 | -0,750581214 | 0,579094295 | 0,004377106 | 0,015222759 | Down |
| Pomgnt2 | 40,24995299 | -0,92212987 | 0,388164931 | 0,000226483 | 0,001080186 | Down |
| Ano10 | 47,04066066 | 1,398566486 | 0,572170432 | 4,72E-05 | 0,000262059 | Up |
| Abhd5 | 4995,859859 | 0,923575007 | 0,118807274 | 6,74E-17 | 2,71E-15 | Up |
| Ccr1 | 520,2067066 | 4,253091822 | 0,585656313 | 2,16E-14 | 6,09E-13 | Up |
| Ccr2 | 80,94767328 | 2,171545223 | 0,96983861 | 4,78E-05 | 0,000264937 | Up |
| Ccr5 | 30,1303597 | 1,68921309 | 0,902498268 | 0,000152061 | 0,000754191 | Up |
| Syne1 | 2049,991481 | -0,628204897 | 0,175099362 | 2,90E-05 | 0,000168576 | Down |
| Nup43 | 4381,001673 | 0,672398581 | 0,082214394 | 2,53E-17 | 1,09E-15 | Up |
| Grm1 | 15,35233286 | -1,638123866 | 0,772964883 | 8,35E-05 | 0,000437926 | Down |
| Gm9797 | 77,38427558 | 0,61175682 | 0,209406098 | 0,000294175 | 0,001365999 | Up |
| Utrn | 2061,625378 | -0,81272963 | 0,239040194 | 1,56E-05 | 9,60E-05 | Down |
| Ltv1 | 9793,295526 | 0,617146464 | 0,09132986 | 1,81E-12 | 3,77E-11 | Up |
| Aig1 | 5,711943907 | 1,738401292 | 1,831495757 | 0,000678823 | 0,002902522 | Up |
| Cited2 | 55471,44831 | -0,763088194 | 0,178887465 | 6,84E-07 | 5,56E-06 | Down |
| Ect2l | 6,596117478 | 0,713748612 | 0,718671879 | 0,007080547 | 0,023222873 | Up |
| Ccdc28a | 620,3424095 | -0,890175477 | 0,124256639 | 9,13E-15 | 2,77E-13 | Down |
| Nhsl1 | 1298,074484 | 0,822509115 | 0,133750828 | 1,66E-11 | 2,93E-10 | Up |
| Myb | 33614,8568 | -0,694998606 | 0,089318205 | 5,15E-16 | 1,85E-14 | Down |
| Gm34267 | 25,99560008 | 1,222756456 | 0,478623579 | 5,04E-05 | 0,000278159 | Up |
| Raet1e | 2,561981638 | 0,774082705 | 1,213296985 | 0,005812356 | 0,019529945 | Up |
| Gm26581 | 4,75514491 | 2,237943654 | 1,84615024 | 0,000313352 | 0,00144579 | Up |
| Raet1d | 35,58970094 | 5,01101227 | 0,838062427 | 2,97E-11 | 5,03E-10 | Up |
| Gm10825 | 50,07323976 | 5,301398825 | 0,657740343 | 6,65E-18 | 3,06E-16 | Up |
| Gm15137 | 53,27674523 | -0,633237902 | 0,243680628 | 0,000630515 | 0,002711837 | Down |
| Enpp1 | 21,06440165 | -1,910153101 | 0,615000022 | 3,95E-06 | 2,76E-05 | Down |
| Akap7 | 340,5561466 | -0,604239768 | 0,108939024 | 3,83E-09 | 4,60E-08 | Down |
| Arhgap18 | 4949,269253 | -0,784188234 | 0,155844603 | 1,45E-08 | 1,58E-07 | Down |
| RP23-331E5,10 | 95,31111914 | -0,816935809 | 0,270956214 | 5,70E-05 | 0,00031143 | Down |
| RP23-128C4,4 | 544,2426585 | -0,763107006 | 0,157640828 | 4,57E-08 | 4,59E-07 | Down |
| Traf3ip2 | 784,1149802 | 0,882508444 | 0,114328048 | 1,42E-16 | 5,48E-15 | Up |
| Rpf2 | 6166,928577 | 0,622033447 | 0,096537876 | 1,44E-11 | 2,56E-10 | Up |
| Cdk19 | 4869,431134 | -1,247351804 | 0,112282757 | 8,60E-31 | 1,78E-28 | Down |
| Sesn1 | 3095,315968 | -0,65883709 | 0,119612198 | 3,20E-09 | 3,87E-08 | Down |
| Scml4 | 61,52170318 | -2,408117219 | 0,443793274 | 3,64E-10 | 5,23E-09 | Down |
| Lilrb4a | 4829,529236 | 5,938227714 | 0,580629999 | 3,76E-27 | 5,32E-25 | Up |
| Nus1 | 12718,49947 | 0,585439108 | 0,071171464 | 3,23E-17 | 1,36E-15 | Up |
| Slc35f1 | 4,00465791 | 0,660091771 | 1,005693597 | 0,009594134 | 0,030171874 | Up |
| P4ha1 | 8154,44816 | 0,604416169 | 0,102488123 | 4,97E-10 | 6,90E-09 | Up |
| Oit3 | 35,99690478 | 0,751108304 | 0,432698851 | 0,00219954 | 0,008286856 | Up |
| Cdh23 | 245,9054833 | -1,252647347 | 0,340346689 | 1,06E-06 | 8,28E-06 | Down |
| Sgpl1 | 5661,291754 | 0,693221095 | 0,119976683 | 5,11E-10 | 7,10E-09 | Up |
| Rpl27a-ps1 | 58,33750587 | -1,428662747 | 0,251074633 | 6,75E-11 | 1,08E-09 | Down |
| Pald1 | 3,718416667 | 0,960053836 | 1,455689498 | 0,002982589 | 0,010837255 | Up |

| | | | | | | |
|---|---|---|---|---|---|---|
| Ppa1 | 20164,62417 | 0,876640604 | 0,123132557 | 1,41E-14 | 4,08E-13 | Up |
| Hkdc1 | 599,7391645 | 1,355377376 | 0,238305756 | 6,99E-11 | 1,12E-09 | Up |
| Supv3l1 | 5157,006582 | 0,603139836 | 0,081101855 | 1,50E-14 | 4,33E-13 | Up |
| Ddx21 | 82809,58928 | 0,630962755 | 0,116902178 | 7,30E-09 | 8,42E-08 | Up |
| Rhobtb1 | 961,7149619 | 1,055291043 | 0,093507303 | 7,12E-32 | 1,62E-29 | Up |
| Adora2a | 4036,259336 | -1,112781304 | 0,134553218 | 6,66E-19 | 3,45E-17 | Down |
| Ggt1 | 87,28368687 | 0,889986821 | 0,310239952 | 6,17E-05 | 0,000335297 | Up |
| Ggt5 | 50,42020708 | -1,440887869 | 0,327148737 | 4,13E-08 | 4,19E-07 | Down |
| Gstt3 | 2123,004728 | -0,779902038 | 0,150912291 | 7,33E-09 | 8,45E-08 | Down |
| Gstt4 | 4,653169103 | 3,220778389 | 2,108870458 | 7,09E-05 | 0,000378918 | Up |
| Slc19a1 | 5856,627741 | 1,180470364 | 0,114817053 | 4,95E-27 | 6,96E-25 | Up |
| Gm10941 | 15,40124955 | -0,910210915 | 0,504472983 | 0,000930931 | 0,003853278 | Down |
| Itgb2 | 23012,70608 | 0,667926519 | 0,095686086 | 2,60E-13 | 6,23E-12 | Up |
| Trpm2 | 18,47986833 | 2,032999683 | 0,623279567 | 2,41E-06 | 1,74E-05 | Up |
| Dnmt3l | 1127,374534 | 0,893689408 | 0,212495378 | 3,58E-07 | 3,06E-06 | Up |
| Icosl | 2230,391057 | 0,712372092 | 0,081216111 | 1,10E-19 | 6,23E-18 | Up |
| Pwp2 | 12485,74247 | 0,795343158 | 0,112469611 | 4,16E-14 | 1,12E-12 | Up |
| Agpat3 | 3052,747228 | 0,952789613 | 0,129908467 | 1,69E-15 | 5,58E-14 | Up |
| Pdxk | 17750,85048 | 0,624489383 | 0,102950278 | 1,55E-10 | 2,33E-09 | Up |
| 2610008E11Rik | 615,8191732 | -0,949985591 | 0,111367136 | 1,05E-19 | 6,02E-18 | Down |
| Plpp2 | 1736,273981 | 0,868935195 | 0,090827948 | 1,43E-23 | 1,37E-21 | Up |
| Cdc34 | 18221,21722 | 0,611408846 | 0,117581838 | 2,43E-08 | 2,56E-07 | Up |
| Plppr3 | 5055,720158 | 0,919348051 | 0,203833574 | 7,49E-08 | 7,25E-07 | Up |
| R3hdm4 | 15754,64639 | -0,695873809 | 0,107944779 | 7,81E-12 | 1,45E-10 | Down |
| Kiss1r | 39,32940063 | -0,73926952 | 0,417455992 | 0,002191576 | 0,008267415 | Down |
| Wdr18 | 15478,93575 | 0,580389159 | 0,086544787 | 3,26E-12 | 6,43E-11 | Up |
| Cirbp | 1994,063706 | -0,739173436 | 0,178450299 | 1,41E-06 | 1,08E-05 | Down |
| Gamt | 53,96634743 | -0,756176413 | 0,275300021 | 0,000187854 | 0,000914707 | Down |
| Apc2 | 208,6627662 | -1,788197864 | 0,239471249 | 3,00E-16 | 1,12E-14 | Down |
| Pcsk4 | 779,9053235 | 0,634157459 | 0,174204274 | 2,30E-05 | 0,000136166 | Up |
| Reep6 | 194,4615902 | 1,52689982 | 0,167735226 | 7,54E-22 | 5,71E-20 | Up |
| Klf16 | 6895,414822 | 0,645308765 | 0,123784004 | 1,77E-08 | 1,92E-07 | Up |
| Btbd2 | 3933,672217 | -0,759466814 | 0,135624938 | 8,03E-10 | 1,07E-08 | Down |
| Thop1 | 16835,18282 | 0,648902558 | 0,128944197 | 4,43E-08 | 4,46E-07 | Up |
| Tjp3 | 623,5626684 | -1,127321256 | 0,274931484 | 2,39E-07 | 2,09E-06 | Down |
| Gm16315 | 89,02980554 | 0,680634918 | 0,299526164 | 0,001052095 | 0,004294446 | Up |
| Smim24 | 924,2397926 | -1,304912624 | 0,20435776 | 1,02E-12 | 2,19E-11 | Down |
| Ncln | 25067,2087 | 0,651187716 | 0,053265633 | 2,52E-35 | 8,09E-33 | Up |
| RP23-132J19,1 | 72,17940298 | 3,546492722 | 0,371194925 | 1,77E-22 | 1,45E-20 | Up |
| Pwp1 | 14333,54235 | 0,639916426 | 0,068772495 | 1,57E-21 | 1,15E-19 | Up |
| Bpifc | 1080,100301 | -1,668065105 | 0,99857687 | 0,000254925 | 0,001201092 | Down |
| A230060F14Rik | 81,9251504 | -2,936800158 | 0,315278188 | 1,06E-21 | 7,90E-20 | Down |
| Fbxo7 | 10180,18259 | -2,007737891 | 0,183359724 | 1,57E-30 | 3,15E-28 | Down |
| Syn3 | 40,9513894 | -3,99559039 | 0,446890651 | 3,50E-20 | 2,19E-18 | Down |
| Gnptab | 53833,2608 | -0,70059888 | 0,11148998 | 2,15E-11 | 3,73E-10 | Down |
| Gm26546 | 34,28311744 | -1,489002607 | 0,382379764 | 3,22E-07 | 2,77E-06 | Down |
| Utp20 | 17353,32244 | 0,726630594 | 0,166030838 | 5,58E-07 | 4,64E-06 | Up |
| Gas2l3 | 16653,24652 | -1,672889989 | 0,188189457 | 3,36E-21 | 2,37E-19 | Down |
| Nr1h4 | 81,58779119 | -4,47097185 | 0,584263667 | 8,21E-16 | 2,84E-14 | Down |
| Slc17a8 | 23,32408474 | -4,216901694 | 0,774164555 | 1,92E-09 | 2,42E-08 | Down |
| 1500026H17Rik | 14,69804799 | -1,588549778 | 0,791651252 | 0,000118133 | 0,00059893 | Down |

| | | | | | | |
|---|---|---|---|---|---|---|
| Cfap54 | 359,0175003 | -0,830787637 | 0,201307777 | 7,60E-07 | 6,13E-06 | Down |
| RP23-337J18,2 | 10,41117979 | 0,917306295 | 0,620609541 | 0,001691625 | 0,006554733 | Up |
| Elk3 | 7839,28634 | -0,750830939 | 0,124253831 | 6,20E-11 | 9,99E-10 | Down |
| RP23-381H23,1 | 442,2506705 | -1,263010517 | 0,175789988 | 4,15E-15 | 1,31E-13 | Down |
| Amdhd1 | 239,6937976 | 0,936952792 | 0,212427494 | 1,10E-07 | 1,03E-06 | Up |
| Gm15915 | 150,4032388 | 1,70111509 | 0,198985109 | 6,07E-20 | 3,66E-18 | Up |
| Cep83os | 549,2854676 | -1,135541875 | 0,215139999 | 7,03E-10 | 9,51E-09 | Down |
| Plxnc1 | 718,5796189 | 1,856079909 | 0,185499585 | 4,25E-26 | 5,35E-24 | Up |
| Cradd | 420,2247501 | -0,693697874 | 0,119204803 | 3,99E-10 | 5,66E-09 | Down |
| Dusp6 | 5131,240224 | 1,308116774 | 0,148335936 | 9,16E-21 | 6,18E-19 | Up |
| Mgat4c | 92,55360561 | 1,172388418 | 0,3678819 | 7,74E-06 | 5,09E-05 | Up |
| Tmtc2 | 3411,412567 | 0,730614767 | 0,109857939 | 1,47E-12 | 3,09E-11 | Up |
| Gm15663 | 429,8583652 | 0,748959439 | 0,137179915 | 1,95E-09 | 2,46E-08 | Up |
| Phlda1 | 371,3231829 | 0,72475005 | 0,166149393 | 6,05E-07 | 4,98E-06 | Up |
| Tmem5 | 4017,668629 | 0,884516143 | 0,103166999 | 1,16E-19 | 6,56E-18 | Up |
| Ctdsp2 | 16834,36362 | -0,781323667 | 0,129540586 | 5,02E-11 | 8,20E-10 | Down |
| Avil | 247,8839592 | 2,280314428 | 1,029129982 | 4,98E-05 | 0,000274618 | Up |
| Slc26a10 | 228,3319397 | 0,691566914 | 0,145416527 | 1,26E-07 | 1,17E-06 | Up |
| Kif5a | 352,4981487 | -1,344471078 | 0,188497783 | 6,78E-15 | 2,08E-13 | Down |
| Lrp1 | 58,39551314 | 1,850004266 | 0,322615094 | 3,04E-11 | 5,13E-10 | Up |
| Apof | 33,50506624 | -0,945797034 | 0,500164532 | 0,000672385 | 0,002877686 | Down |
| Stat2 | 5653,784203 | -0,980829238 | 0,136535409 | 4,50E-15 | 1,41E-13 | Down |
| Mir8105 | 15,00918783 | -0,603132156 | 0,446904645 | 0,008914263 | 0,028296937 | Down |
| Myl6b | 4879,902041 | -0,644742523 | 0,105159637 | 8,93E-11 | 1,41E-09 | Down |
| A430046D13Rik | 300,764932 | -0,757601675 | 0,160827436 | 9,08E-08 | 8,69E-07 | Down |
| Pa2g4 | 86246,25837 | 0,699835211 | 0,093897284 | 6,20E-15 | 1,92E-13 | Up |
| Ikzf4 | 2431,949517 | 0,781271044 | 0,0896882 | 9,43E-20 | 5,47E-18 | Up |
| Suox | 2564,30032 | -0,680940058 | 0,085044226 | 9,60E-17 | 3,78E-15 | Down |
| Bloc1s1 | 493,9946311 | -0,785463975 | 0,120683325 | 2,27E-12 | 4,60E-11 | Down |
| Gm9770 | 961,5193827 | -0,770614742 | 0,144038031 | 2,96E-09 | 3,60E-08 | Down |
| Patz1 | 12103,05685 | -0,748635723 | 0,076034386 | 3,14E-24 | 3,35E-22 | Down |
| Gm11944 | 195,4232213 | -0,701968716 | 0,208569499 | 3,76E-05 | 0,000213825 | Down |
| Pik3ip1 | 544,4473269 | -1,427609215 | 0,490777753 | 1,13E-05 | 7,18E-05 | Down |
| Gm11948 | 5,77167277 | -0,663931529 | 0,733968306 | 0,009301493 | 0,029348159 | Down |
| Ccdc157 | 239,7664646 | -1,09164019 | 0,190796996 | 5,84E-11 | 9,45E-10 | Down |
| Gatsl3 | 234,9136018 | 2,852044026 | 0,225104634 | 8,51E-38 | 3,45E-35 | Up |
| Osm | 2398,355282 | 1,54675385 | 0,135630597 | 3,84E-32 | 8,99E-30 | Up |
| Lif | 95,69670836 | 1,065505508 | 0,286189271 | 1,37E-06 | 1,05E-05 | Up |
| Znrf3 | 2485,085134 | 0,85867503 | 0,091470998 | 8,81E-23 | 7,62E-21 | Up |
| Gm11963 | 137,8353194 | 0,923801972 | 0,244216472 | 1,87E-06 | 1,39E-05 | Up |
| Xbp1 | 46769,59394 | 0,952455634 | 0,107460416 | 5,42E-21 | 3,74E-19 | Up |
| Polm | 1099,511717 | -1,558493247 | 0,157599116 | 3,95E-25 | 4,69E-23 | Down |
| H2afv | 5771,250671 | -1,256484525 | 0,143819684 | 1,65E-20 | 1,07E-18 | Down |
| Upp1 | 1663,281966 | -1,371136561 | 0,194129694 | 1,13E-14 | 3,34E-13 | Down |
| Abca13 | 258,6533689 | 1,469128908 | 0,2596131 | 7,81E-11 | 1,24E-09 | Up |
| Ddc | 1088,133481 | 0,899029013 | 0,13513402 | 3,22E-13 | 7,56E-12 | Up |
| Grb10 | 260,5275999 | 0,709950948 | 0,473575883 | 0,004039731 | 0,014181769 | Up |
| Plek | 48649,19621 | -1,192879963 | 0,158934562 | 3,40E-16 | 1,26E-14 | Down |
| Pno1 | 8850,649921 | 0,974499197 | 0,102017151 | 7,65E-24 | 7,59E-22 | Up |
| Gm12020 | 8,167842106 | 0,869650369 | 0,665167571 | 0,002658356 | 0,009803494 | Up |
| Spred2 | 9083,74591 | 0,687520869 | 0,098090443 | 1,80E-13 | 4,37E-12 | Up |

| | | | | | | |
|---|---|---|---|---|---|---|
| Gm12031 | 350,0440717 | 1,085960237 | 0,186003333 | 2,91E-11 | 4,93E-10 | Up |
| Wdpcp | 417,7205719 | -1,349610274 | 0,155982912 | 4,26E-20 | 2,59E-18 | Down |
| Gm12056 | 6,028267217 | 0,966379298 | 1,073938482 | 0,003478796 | 0,012439746 | Up |
| Zrsr1 | 692,0371414 | -0,642209235 | 0,125331144 | 2,90E-08 | 3,02E-07 | Down |
| Pnpt1 | 7087,756857 | 0,626322203 | 0,119188064 | 1,63E-08 | 1,78E-07 | Up |
| Gm12097 | 2,996864391 | 0,655046967 | 1,073034427 | 0,00909886 | 0,028812793 | Up |
| Hba-x | 4,178230899 | -2,045047966 | 2,031412644 | 0,000384379 | 0,001740522 | Down |
| Hba-a1 | 127,3128889 | -1,304422761 | 1,195263347 | 0,001361918 | 0,005410743 | Down |
| Rars | 28763,49294 | 0,602539598 | 0,072308396 | 1,18E-17 | 5,26E-16 | Up |
| Ccng1 | 36169,2802 | 0,626704008 | 0,176264603 | 3,30E-05 | 0,000189497 | Up |
| Hspd1-ps3 | 318,2352883 | 0,683683985 | 0,160441535 | 1,31E-06 | 1,00E-05 | Up |
| Atp10b | 18,6628345 | 0,680403664 | 0,573281247 | 0,006903127 | 0,022775954 | Up |
| Olfr56 | 109,8057391 | 1,81267979 | 0,244768732 | 4,52E-16 | 1,64E-14 | Up |
| Gm12187 | 3,637023547 | 0,60381365 | 0,847719892 | 0,012652982 | 0,038130158 | Up |
| Ifi47 | 1481,469323 | 1,09035793 | 0,148179771 | 9,48E-16 | 3,23E-14 | Up |
| Rasgef1c | 6,811438603 | -0,790211131 | 0,79936361 | 0,005320847 | 0,018108333 | Down |
| 3010026O09Rik | 675,4498137 | -0,583296044 | 0,0971144 | 2,94E-10 | 4,27E-09 | Down |
| Ltc4s | 612,1824125 | 0,954589897 | 0,1574019 | 1,09E-11 | 1,98E-10 | Up |
| Adamts2 | 2,515732767 | 0,630729633 | 1,123576136 | 0,008783859 | 0,02793154 | Up |
| N4bp3 | 3027,908061 | -0,597901545 | 0,109336696 | 6,18E-09 | 7,23E-08 | Down |
| D930048N14Rik | 299,0755494 | -0,824702396 | 0,189252907 | 2,83E-07 | 2,47E-06 | Down |
| Tcf7 | 7,876019983 | -0,882728741 | 0,696948514 | 0,00270065 | 0,009933388 | Down |
| Zcchc10 | 2783,333201 | 0,635035344 | 0,089714079 | 1,68E-13 | 4,12E-12 | Up |
| Sep 08 | 18118,04691 | -0,696325333 | 0,076497468 | 6,40E-21 | 4,36E-19 | Down |
| A430108G06Rik | 12,9641637 | -0,596199188 | 0,429035608 | 0,008881056 | 0,028211139 | Down |
| Irf1 | 3337,094619 | -0,648988696 | 0,112509165 | 7,76E-10 | 1,04E-08 | Down |
| Acsl6 | 220,3004438 | -1,578459545 | 0,214028878 | 1,02E-15 | 3,47E-14 | Down |
| Slc36a2 | 101,0701759 | -1,061849231 | 0,419261521 | 8,59E-05 | 0,000449316 | Down |
| Slc36a1os | 60,40644346 | -1,270218233 | 0,283246669 | 3,45E-08 | 3,54E-07 | Down |
| Fat2 | 185,720143 | 1,458496302 | 0,351156042 | 1,18E-07 | 1,10E-06 | Up |
| 2010001A14Rik | 248,5656826 | -0,814887418 | 0,195480354 | 7,09E-07 | 5,74E-06 | Down |
| Gm12247 | 49,05827899 | -2,15033872 | 0,327047279 | 1,73E-13 | 4,23E-12 | Down |
| Gemin5 | 19951,2193 | 0,651097216 | 0,096893357 | 1,82E-12 | 3,78E-11 | Up |
| Trim58 | 2091,574301 | -1,21089921 | 0,127352011 | 1,21E-23 | 1,18E-21 | Down |
| Gm12258 | 129,3155738 | -0,683586224 | 0,210879274 | 6,53E-05 | 0,000351471 | Down |
| Nlrp3 | 104,4376434 | 1,019940579 | 0,530412557 | 0,000478343 | 0,002123359 | Up |
| Med9os | 88,52164496 | -0,699198265 | 0,217630345 | 6,43E-05 | 0,000347325 | Down |
| Shmt1 | 21665,39839 | 0,715860275 | 0,139828661 | 1,64E-08 | 1,78E-07 | Up |
| Natd1 | 989,8476047 | -1,316807492 | 0,156917285 | 3,70E-19 | 2,00E-17 | Down |
| Map2k3 | 20175,43432 | -0,723839098 | 0,108397001 | 1,30E-12 | 2,76E-11 | Down |
| Grap | 708,8283831 | 0,823337113 | 0,178261699 | 8,38E-08 | 8,07E-07 | Up |
| Specc1 | 7087,684873 | -0,714340726 | 0,130452529 | 2,41E-09 | 2,98E-08 | Down |
| Lrrc75a | 1127,904021 | -0,991696605 | 0,174171283 | 8,94E-11 | 1,41E-09 | Down |
| Trim16 | 1370,837268 | 0,726126218 | 0,420536751 | 0,002549265 | 0,009442883 | Up |
| Hs3st3b1 | 2950,344531 | 1,144545475 | 0,098240257 | 1,20E-33 | 3,36E-31 | Up |
| Slc25a35 | 67,47216633 | -1,142871207 | 0,503257011 | 0,000137402 | 0,000687841 | Down |
| Alox8 | 110,5484431 | 1,009449011 | 0,247761575 | 3,72E-07 | 3,18E-06 | Up |
| Kdm6b | 6599,126144 | 1,14084305 | 0,119109402 | 5,09E-24 | 5,16E-22 | Up |
| Cd68 | 1730,234601 | 1,385702473 | 0,240935504 | 4,85E-11 | 7,93E-10 | Up |
| Eif4a1 | 225385,2986 | 0,589127627 | 0,066552989 | 1,42E-19 | 7,99E-18 | Up |
| Tnfsf13 | 62,71437089 | -0,662106975 | 0,283143291 | 0,001009644 | 0,00413596 | Down |

| | | | | | | |
|---|---|---|---|---|---|---|
| Zbtb4 | 21,65918398 | -1,283368985 | 0,518314953 | 5,46E-05 | 0,000299405 | Down |
| Nlgn2 | 91,9016118 | -0,645552637 | 0,218350043 | 0,000210212 | 0,001010765 | Down |
| Plscr3 | 1690,338622 | -0,637693882 | 0,071732809 | 7,24E-20 | 4,33E-18 | Down |
| 2810408A11Rik | 34,14067 | -0,859225585 | 0,312944159 | 0,000105346 | 0,00053949 | Down |
| Arrb2 | 15836,62708 | 0,955093538 | 0,090367631 | 2,68E-28 | 4,30E-26 | Up |
| Gp1ba | 496,4015227 | -1,463184334 | 0,189048803 | 7,80E-17 | 3,10E-15 | Down |
| Rnf167 | 5108,691452 | -1,156468668 | 0,091360486 | 5,53E-39 | 2,66E-36 | Down |
| Eno3 | 95,25925351 | -0,621761314 | 0,275529612 | 0,001590808 | 0,006218181 | Down |
| Scimp | 7,686062567 | 3,130746137 | 1,37552061 | 3,34E-05 | 0,000191917 | Up |
| C1qbp | 42245,23443 | 0,586906905 | 0,091456395 | 2,15E-11 | 3,73E-10 | Up |
| Nlrp1a | 84,63719684 | 0,693801812 | 0,206559661 | 4,08E-05 | 0,000230006 | Up |
| Aipl1 | 422,8715118 | -0,972880799 | 0,209149026 | 2,87E-08 | 2,99E-07 | Down |
| Fam64a | 7453,227251 | -0,898666168 | 0,168743845 | 1,23E-09 | 1,60E-08 | Down |
| Mybbp1a | 122111,9091 | 0,73729268 | 0,128146935 | 4,01E-10 | 5,69E-09 | Up |
| Spns2 | 32535,43036 | -0,912907102 | 0,106304308 | 8,17E-20 | 4,80E-18 | Down |
| Spns3 | 7776,20801 | -0,830235743 | 0,309907052 | 0,000149512 | 0,000742758 | Down |
| Mettl16 | 15985,67294 | 0,795054761 | 0,123831591 | 3,71E-12 | 7,20E-11 | Up |
| Tsr1 | 12877,14543 | 0,732736452 | 0,094213933 | 3,73E-16 | 1,37E-14 | Up |
| Serpinf2 | 2,989043846 | 0,710708237 | 1,127150845 | 0,007437063 | 0,024248463 | Up |
| Tlcd2 | 25,01158449 | 0,633254551 | 0,52319258 | 0,008626904 | 0,027513887 | Up |
| Gm45606 | 553,0133456 | -0,886737968 | 0,164098347 | 8,62E-10 | 1,14E-08 | Down |
| Inpp5k | 2879,57959 | -0,721352537 | 0,105375639 | 4,19E-13 | 9,61E-12 | Down |
| Gm12341 | 96,30970412 | 1,240173251 | 0,201720954 | 4,30E-12 | 8,29E-11 | Up |
| Dhrs13 | 3086,77933 | 0,634687765 | 0,087443855 | 4,55E-14 | 1,22E-12 | Up |
| Traf4 | 4242,227827 | 0,757447563 | 0,090300774 | 1,97E-18 | 9,71E-17 | Up |
| Nek8 | 533,294894 | -0,658499342 | 0,149327089 | 8,22E-07 | 6,58E-06 | Down |
| BC030499 | 57,65991066 | 0,912181154 | 0,348555512 | 0,00011996 | 0,000606844 | Up |
| Slc46a1 | 294,4818785 | 0,669892505 | 0,174304774 | 8,18E-06 | 5,36E-05 | Up |
| Tmem199 | 3546,194249 | 0,814653572 | 0,133078419 | 2,12E-11 | 3,69E-10 | Up |
| Tmem97 | 13701,63084 | 0,840963478 | 0,090003325 | 1,58E-22 | 1,31E-20 | Up |
| Lgals9 | 16272,47577 | -1,02522201 | 0,18888504 | 3,75E-10 | 5,36E-09 | Down |
| Rab11fip4 | 393,4511642 | 0,765012425 | 0,266456852 | 0,000122753 | 0,000619941 | Up |
| Gm11417 | 102,8043685 | -0,730310572 | 0,192601475 | 6,34E-06 | 4,24E-05 | Down |
| Nle1 | 10053,96248 | 0,712086931 | 0,118976134 | 1,25E-10 | 1,92E-09 | Up |
| Slfn5 | 587,8125687 | 1,416921066 | 0,265704237 | 4,77E-10 | 6,67E-09 | Up |
| Slfn2 | 1045,664031 | 4,256054246 | 0,333255687 | 1,61E-38 | 7,53E-36 | Up |
| Slfn1 | 8,722753994 | 5,117889438 | 1,349731658 | 3,55E-07 | 3,04E-06 | Up |
| Ccl5 | 57,8624051 | 1,076015954 | 1,006315526 | 0,002169956 | 0,00819771 | Up |
| Ccl6 | 1119,856899 | 2,727729313 | 0,929838545 | 8,90E-06 | 5,79E-05 | Up |
| Ccl3 | 441,8749966 | 3,084276967 | 0,447182807 | 3,54E-13 | 8,23E-12 | Up |
| Ccl4 | 335,2517241 | 3,53857997 | 0,345732994 | 2,06E-25 | 2,49E-23 | Up |
| Wfdc17 | 1,731702877 | 0,804519407 | 1,312620296 | 0,004685821 | 0,016173212 | Up |
| Wfdc21 | 16,19669521 | 1,883533159 | 0,99339032 | 0,000125197 | 0,00063089 | Up |
| Pigw | 629,5883303 | 0,764105597 | 0,108989519 | 8,71E-14 | 2,23E-12 | Up |
| Myo19 | 4985,160782 | 1,114365511 | 0,162586024 | 3,70E-14 | 1,01E-12 | Up |
| Znhit3 | 3027,402432 | 0,591953581 | 0,08423742 | 3,25E-13 | 7,62E-12 | Up |
| Bcas3 | 2232,20399 | -0,606252741 | 0,0930614 | 1,00E-11 | 1,83E-10 | Down |
| Sep 04 | 11,63961559 | -0,957719064 | 0,52524789 | 0,000744194 | 0,003149588 | Down |
| Akap1 | 13656,01915 | 0,581023814 | 0,094786232 | 1,39E-10 | 2,10E-09 | Up |
| Utp18 | 8967,047736 | 0,734894447 | 0,085491829 | 4,12E-19 | 2,20E-17 | Up |
| Wfikkn2 | 10,9105815 | 0,736756542 | 0,458395068 | 0,002967279 | 0,010788066 | Up |

| | | | | | | |
|---|---|---|---|---|---|---|
| Acsf2 | 171,4211119 | -1,370746846 | 0,217758026 | 1,87E-12 | 3,87E-11 | Down |
| Sgca | 18,65168968 | 0,610845975 | 0,40405387 | 0,006943464 | 0,022888366 | Up |
| Dlx4os | 173,3534353 | 0,737106148 | 0,174301231 | 9,87E-07 | 7,77E-06 | Up |
| Gm11515 | 57,49730154 | 0,690075372 | 0,270626978 | 0,000493252 | 0,002180528 | Up |
| Nfe2l1 | 20498,0326 | 0,676893923 | 0,073698916 | 3,57E-21 | 2,51E-19 | Up |
| Lrrc46 | 65,64838443 | -1,325847524 | 0,235028104 | 9,08E-11 | 1,43E-09 | Down |
| Tbkbp1 | 9540,603635 | 1,258744181 | 0,102052876 | 4,93E-37 | 1,88E-34 | Up |
| Plxdc1 | 19,07902451 | -1,256205673 | 0,519706182 | 6,86E-05 | 0,00036732 | Down |
| Arl5c | 2861,647057 | 0,71339821 | 0,40000103 | 0,002460078 | 0,009157346 | Up |
| Cacnb1 | 130,9913445 | -0,637301987 | 0,241992649 | 0,000558857 | 0,002439254 | Down |
| Fbxl20 | 4941,19867 | -0,868819295 | 0,167142564 | 3,06E-09 | 3,71E-08 | Down |
| Rara | 1249,45101 | 1,575427947 | 0,12941687 | 3,90E-36 | 1,34E-33 | Up |
| Igfbp4 | 49594,14105 | 0,613381957 | 0,114140837 | 9,39E-09 | 1,06E-07 | Up |
| Tns4 | 359,6303733 | 0,880628975 | 0,333685045 | 0,000130174 | 0,000653983 | Up |
| Jup | 235,9404248 | -1,301685338 | 0,427686363 | 9,32E-06 | 6,04E-05 | Down |
| Ttc25 | 969,533925 | 1,107025439 | 0,165117591 | 1,05E-13 | 2,65E-12 | Up |
| Cnp | 6752,069168 | -0,689040796 | 0,13286802 | 1,45E-08 | 1,59E-07 | Down |
| Stat5a | 4232,933386 | -0,659860091 | 0,121981975 | 5,49E-09 | 6,47E-08 | Down |
| Plekhh3 | 34,01354151 | 4,582199816 | 0,810680967 | 4,03E-10 | 5,71E-09 | Up |
| Aoc2 | 1152,772416 | 0,703333893 | 0,253660665 | 0,000244155 | 0,001156316 | Up |
| Aoc3 | 84,85756428 | 1,464570058 | 0,501822327 | 1,03E-05 | 6,62E-05 | Up |
| Etv4 | 842,4524741 | 2,994161166 | 0,178754373 | 9,06E-64 | 1,50E-60 | Up |
| Dusp3 | 9406,05736 | 0,827925406 | 0,102803548 | 1,58E-17 | 6,97E-16 | Up |
| Ccdc103 | 19,73196061 | -1,402070913 | 0,486215325 | 1,28E-05 | 8,02E-05 | Down |
| Acbd4 | 2574,740235 | -0,789787008 | 0,126705937 | 1,31E-11 | 2,35E-10 | Down |
| Hexim2 | 3738,663849 | -0,637152368 | 0,110182617 | 7,80E-10 | 1,04E-08 | Down |
| Gm20511 | 9,213551052 | -0,623626856 | 0,704360722 | 0,011255556 | 0,03455766 | Down |
| Myl4 | 105,1560188 | -1,376161933 | 0,230158312 | 1,29E-11 | 2,31E-10 | Down |
| Itgb3 | 28796,22896 | -2,169833516 | 0,128764759 | 3,15E-66 | 6,39E-63 | Down |
| Gm11672 | 9,081971331 | 0,84676874 | 0,780163832 | 0,003926783 | 0,013819795 | Up |
| Ftsj3 | 39137,72399 | 0,895374241 | 0,08834462 | 3,91E-26 | 4,99E-24 | Up |
| Cd79b | 160,9966712 | -0,701869488 | 0,313886606 | 0,001006406 | 0,004124547 | Down |
| Icam2 | 2298,673176 | 0,582633049 | 0,105335842 | 4,79E-09 | 5,68E-08 | Up |
| Ern1 | 10008,65217 | -0,682327557 | 0,145114079 | 1,76E-07 | 1,58E-06 | Down |
| Tex2 | 17789,82602 | -0,620920334 | 0,206337706 | 0,000212115 | 0,00101938 | Down |
| Milr1 | 749,3495395 | 2,896025661 | 0,243998423 | 1,79E-33 | 4,89E-31 | Up |
| Nol11 | 17281,1567 | 0,64051819 | 0,100393669 | 1,89E-11 | 3,32E-10 | Up |
| Slc16a6 | 11528,65552 | 0,892805171 | 0,130828662 | 1,03E-13 | 2,60E-12 | Up |
| Ttyh2 | 6117,857672 | -0,671048143 | 0,098157035 | 6,95E-13 | 1,55E-11 | Down |
| Dnaic2 | 3,271629048 | -0,694613451 | 0,863695894 | 0,008625195 | 0,027513242 | Down |
| Gpr142 | 10,70709448 | 0,603988102 | 0,664586482 | 0,012254151 | 0,037124304 | Up |
| Cd300ld | 194,4014277 | 1,291414554 | 0,147171293 | 1,29E-20 | 8,53E-19 | Up |
| Cd300ld2 | 45,67226224 | 1,377094753 | 0,33321342 | 1,43E-07 | 1,31E-06 | Up |
| Rab37 | 416,2420846 | -0,84438534 | 0,203261905 | 6,15E-07 | 5,04E-06 | Down |
| Cd300lf | 3294,058118 | 1,261897829 | 0,107402028 | 5,87E-34 | 1,67E-31 | Up |
| Fads6 | 493,9930731 | -0,644389228 | 0,192405006 | 6,01E-05 | 0,000327132 | Down |
| Hid1 | 20,88227253 | -0,611886245 | 0,530681905 | 0,010190857 | 0,031774873 | Down |
| Myo15b | 803,2103227 | 0,587415449 | 0,237459257 | 0,00118227 | 0,004766138 | Up |
| Smim5 | 8823,774637 | -1,534796673 | 0,132401665 | 4,55E-33 | 1,19E-30 | Down |
| Smim6 | 358,5801824 | -0,860856599 | 0,200215153 | 2,87E-07 | 2,50E-06 | Down |
| Itgb4 | 428,4791319 | 0,792976696 | 0,199196604 | 1,85E-06 | 1,38E-05 | Up |

| | | | | | | |
|---|---|---|---|---|---|---|
| Unk | 7649,837446 | -0,605037956 | 0,133772031 | 7,21E-07 | 5,84E-06 | Down |
| Wbp2 | 31706,53821 | -0,621607727 | 0,104727205 | 3,51E-10 | 5,06E-09 | Down |
| Cdk3-ps | 19,10087672 | -0,648178629 | 0,535604422 | 0,007961332 | 0,02569179 | Down |
| Evpl | 79,20208204 | -1,725754039 | 0,353323979 | 3,24E-09 | 3,91E-08 | Down |
| Rhbdf2 | 7965,901881 | 0,791002851 | 0,119663924 | 1,09E-12 | 2,34E-11 | Up |
| Jmjd6 | 12636,99834 | 0,669651721 | 0,126745037 | 1,01E-08 | 1,13E-07 | Up |
| Tha1 | 1141,932535 | -0,661119752 | 0,147401842 | 5,73E-07 | 4,75E-06 | Down |
| Socs3 | 3350,218104 | -0,629589493 | 0,134294435 | 2,81E-07 | 2,45E-06 | Down |
| Usp36 | 15953,13352 | 0,896823929 | 0,09821745 | 6,96E-22 | 5,30E-20 | Up |
| C1qtnf1 | 9,56653061 | 1,526629132 | 0,851434083 | 0,000218433 | 0,001045342 | Up |
| Sgsh | 2347,24797 | -1,322828611 | 0,22701769 | 3,12E-11 | 5,26E-10 | Down |
| Slc26a11 | 69,14630225 | -2,050229511 | 0,40209918 | 1,04E-09 | 1,36E-08 | Down |
| Gm11767 | 51,92071233 | -0,871529875 | 0,280211689 | 3,06E-05 | 0,000177051 | Down |
| Cep131 | 3246,244364 | -0,850258567 | 0,095967266 | 1,25E-20 | 8,30E-19 | Down |
| 0610009L18Rik | 162,2274992 | -1,088702795 | 0,218495179 | 3,68E-09 | 4,43E-08 | Down |
| Ccdc137 | 5048,72837 | 0,820327864 | 0,23797648 | 1,25E-05 | 7,89E-05 | Up |
| Ppp1r27 | 19,09673643 | -0,76359023 | 0,451258716 | 0,00223635 | 0,008415459 | Down |
| Tex19,2 | 27,7198312 | 1,409046134 | 0,476779524 | 1,01E-05 | 6,49E-05 | Up |
| Narf | 2989,659917 | -1,272560419 | 0,174865047 | 2,15E-15 | 7,07E-14 | Down |
| Metrnl | 4887,976873 | 0,805379715 | 0,107628966 | 1,79E-15 | 5,90E-14 | Up |
| Gm12589 | 33,59654821 | 1,094175508 | 0,401061868 | 4,33E-05 | 0,000242277 | Up |
| 1110002L01Rik | 454,7787592 | 0,862748636 | 0,133989591 | 1,80E-12 | 3,75E-11 | Up |
| Fkbp1b | 190,2514802 | -1,098665051 | 0,212650696 | 1,36E-09 | 1,75E-08 | Down |
| Mycn | 8714,752032 | 0,901287016 | 0,20723574 | 1,78E-07 | 1,59E-06 | Up |
| Lpin1 | 5191,723591 | -0,965681509 | 0,219341389 | 9,91E-08 | 9,43E-07 | Down |
| Greb1 | 5300,365898 | -7,74953795 | 0,547492154 | 5,96E-49 | 5,18E-46 | Down |
| Nol10 | 12006,45166 | 0,745162346 | 0,092162997 | 2,78E-17 | 1,19E-15 | Up |
| Odc1 | 90651,84054 | 1,07775954 | 0,116868064 | 1,40E-22 | 1,17E-20 | Up |
| Adam17 | 20460,03915 | -0,70608989 | 0,11805614 | 1,36E-10 | 2,06E-09 | Down |
| Rrm2 | 48359,56412 | 0,916511511 | 0,114288369 | 9,77E-18 | 4,40E-16 | Up |
| Mboat2 | 3794,461036 | 0,64561413 | 0,102005995 | 2,53E-11 | 4,32E-10 | Up |
| Sh3yl1 | 223,6600349 | -0,607530149 | 0,144437204 | 2,89E-06 | 2,06E-05 | Down |
| Bcap29 | 12417,3914 | 0,644254169 | 0,106122005 | 1,30E-10 | 1,98E-09 | Up |
| Dus4l | 2202,641229 | 1,025137799 | 0,119741635 | 5,86E-20 | 3,54E-18 | Up |
| Nampt | 13993,26218 | 0,580429675 | 0,066286617 | 3,50E-19 | 1,89E-17 | Up |
| Sypl | 25108,83645 | 0,735765187 | 0,130458879 | 7,88E-10 | 1,05E-08 | Up |
| F730043M19Rik | 218,2844722 | 0,639123515 | 0,174775563 | 2,08E-05 | 0,000125143 | Up |
| Hdac9 | 275,840004 | 2,262529795 | 0,291091292 | 3,67E-17 | 1,52E-15 | Up |
| Arl4a | 2052,309251 | -1,182973557 | 0,085519525 | 1,00E-45 | 6,52E-43 | Down |
| Scin | 4842,191563 | -1,217650733 | 0,666635259 | 0,000340794 | 0,001560187 | Down |
| Ifrd1 | 11870,87126 | 1,153560629 | 0,155411662 | 6,00E-16 | 2,13E-14 | Up |
| Mia2,1 | 18210,05399 | 0,787525269 | 0,090060052 | 6,58E-20 | 3,95E-18 | Up |
| Abhd12b | 4,786448649 | 0,611175872 | 0,717420488 | 0,012081522 | 0,036674329 | Up |
| F730035M05Rik | 15,82309441 | 0,585879173 | 0,473698913 | 0,011060873 | 0,034034314 | Up |
| Timm9 | 7560,531239 | 0,750157395 | 0,115848011 | 3,94E-12 | 7,62E-11 | Up |
| Prkch | 22395,79984 | -0,883884951 | 0,080156749 | 3,11E-30 | 6,11E-28 | Down |
| Rhoj | 8706,867025 | 3,630387928 | 0,442594757 | 2,84E-17 | 1,21E-15 | Up |
| Mthfd1 | 58043,86861 | 0,589064506 | 0,098952903 | 3,92E-10 | 5,58E-09 | Up |
| Hspa2 | 331,253434 | -0,790328161 | 0,154437633 | 8,76E-09 | 9,95E-08 | Down |
| Eif2s1 | 36611,28724 | 0,583571472 | 0,086369478 | 2,27E-12 | 4,60E-11 | Up |
| Rdh12 | 730,9090474 | -0,593130498 | 0,172650426 | 6,34E-05 | 0,000343481 | Down |

| | | | | | | |
|---|---|---|---|---|---|---|
| 2310015A10Rik | 599,3068281 | -1,192984249 | 0,109265435 | 5,78E-30 | 1,09E-27 | Down |
| Sipa1l1 | 98257,06086 | -1,225042503 | 0,114751174 | 9,07E-29 | 1,52E-26 | Down |
| Zfyve1 | 3976,992005 | -0,751165645 | 0,103133845 | 1,36E-14 | 3,95E-13 | Down |
| Aldh6a1 | 1572,594812 | -0,592504058 | 0,126724347 | 3,83E-07 | 3,26E-06 | Down |
| Ltbp2 | 184,5277466 | -0,760148579 | 0,24940682 | 7,28E-05 | 0,000387455 | Down |
| Prox2 | 519,098261 | -0,925132066 | 0,102873246 | 2,00E-21 | 1,44E-19 | Down |
| Gm38355 | 9,496150694 | -0,77047719 | 0,562957337 | 0,003631411 | 0,012927256 | Down |
| Tgfb3 | 214,763744 | -3,549057886 | 0,317020913 | 6,65E-30 | 1,24E-27 | Down |
| Ston2 | 4047,202282 | -0,814271085 | 0,133115273 | 2,19E-11 | 3,79E-10 | Down |
| 9030617O03Rik | 9762,931775 | 0,819011791 | 0,089027605 | 7,64E-22 | 5,76E-20 | Up |
| Rin3 | 19394,08306 | -0,683914122 | 0,136390484 | 3,69E-08 | 3,77E-07 | Down |
| Tmem251 | 1735,090556 | 0,698035225 | 0,113107135 | 4,48E-11 | 7,38E-10 | Up |
| Ifi27 | 6938,423981 | 0,685818233 | 0,219260185 | 9,32E-05 | 0,000482792 | Up |
| Serpina3f | 222,9316859 | 2,543202499 | 0,34426275 | 2,17E-15 | 7,11E-14 | Up |
| Zfp839 | 686,5180136 | -0,611386123 | 0,135946242 | 7,77E-07 | 6,26E-06 | Down |
| Cdc42bpb | 66,57133961 | -4,279306073 | 0,840637163 | 1,00E-08 | 1,13E-07 | Down |
| Trmt61a | 4466,011243 | 1,245132617 | 0,122647125 | 2,34E-26 | 3,05E-24 | Up |
| Pld4 | 17833,91103 | 1,076170486 | 0,168073411 | 8,23E-13 | 1,81E-11 | Up |
| Nudt14 | 1734,516994 | -0,70550673 | 0,108232264 | 4,46E-12 | 8,57E-11 | Down |
| Pacs2 | 10669,39289 | 0,59444101 | 0,072227449 | 2,94E-17 | 1,24E-15 | Up |
| Ighm | 1457,829526 | -1,740000527 | 0,252275638 | 2,12E-14 | 6,00E-13 | Down |
| Gm9260 | 2937,502245 | 0,865417941 | 0,162670194 | 1,60E-09 | 2,05E-08 | Up |
| Gm20658 | 466,5322882 | -0,927944193 | 0,432433299 | 0,000399161 | 0,001798982 | Down |
| Akr1c18 | 358,3630148 | 2,607020858 | 0,678020704 | 5,70E-07 | 4,73E-06 | Up |
| Idi1 | 7542,710232 | -1,464256783 | 0,19651291 | 6,90E-16 | 2,41E-14 | Down |
| Idi2 | 7,461024338 | -0,710571079 | 0,763690895 | 0,00753745 | 0,024505769 | Down |
| Heatr1 | 24524,10792 | 0,716285321 | 0,121617854 | 2,15E-10 | 3,18E-09 | Up |
| Lyst | 43,14978485 | 1,567952422 | 0,759232098 | 0,000103637 | 0,000531784 | Up |
| Sugct | 17,07323454 | -1,040656368 | 0,579159596 | 0,000591877 | 0,00256586 | Down |
| Gpr141 | 1956,96167 | 0,739816599 | 0,477556096 | 0,003210982 | 0,011595602 | Up |
| Zkscan4 | 181,0267623 | -1,41320594 | 0,215282701 | 3,36E-13 | 7,85E-12 | Down |
| Zfp184 | 127,6383561 | -0,935697023 | 0,214599041 | 1,40E-07 | 1,28E-06 | Down |
| Gm11335 | 2,238161616 | -0,599448702 | 1,009390552 | 0,011432714 | 0,035019159 | Down |
| Fam65b | 9612,90591 | -0,580840682 | 0,114351055 | 5,55E-08 | 5,48E-07 | Down |
| Mboat1 | 3788,958972 | 0,799428301 | 0,216859943 | 5,80E-06 | 3,91E-05 | Up |
| Serpinb6b | 1801,582066 | -0,859449387 | 0,143412188 | 3,24E-11 | 5,43E-10 | Down |
| Serpinb9 | 774,4030164 | -0,777989743 | 0,171378593 | 1,74E-07 | 1,56E-06 | Down |
| Serpinb9f | 2,374515574 | 0,656958679 | 1,140748067 | 0,008122399 | 0,026137665 | Up |
| Serpinb9g | 19,49516199 | 3,096016013 | 0,659527587 | 5,00E-08 | 4,97E-07 | Up |
| Slc22a23 | 6460,465171 | -1,129474956 | 0,121352635 | 6,60E-23 | 5,90E-21 | Down |
| Rpp40 | 2175,583782 | 0,956973659 | 0,091966971 | 1,50E-27 | 2,16E-25 | Up |
| Ppp1r3g | 145,2577066 | 0,602495952 | 0,395603008 | 0,007194373 | 0,02355812 | Up |
| Fars2 | 3576,941012 | -0,788017106 | 0,085799452 | 1,21E-21 | 8,94E-20 | Down |
| Eef1e1 | 7193,167973 | 1,189677304 | 0,10058038 | 1,72E-34 | 5,08E-32 | Up |
| Mylip | 1392,91633 | -0,953311988 | 0,124058043 | 1,13E-16 | 4,41E-15 | Down |
| Gmpr | 95,63828154 | 1,091629555 | 0,253092732 | 9,92E-08 | 9,43E-07 | Up |
| Nhlrc1 | 2682,416734 | 0,709408331 | 0,088394489 | 6,39E-17 | 2,57E-15 | Up |
| A830005F24Rik | 84,9503461 | -0,682231921 | 0,216809381 | 9,00E-05 | 0,000467827 | Down |
| Nol8 | 8292,613356 | 0,774910806 | 0,094523921 | 8,15E-18 | 3,71E-16 | Up |
| S1pr3 | 2148,774827 | -0,709201958 | 0,179544727 | 3,97E-06 | 2,76E-05 | Down |
| Sema4d | 29089,31025 | -0,626502737 | 0,128793472 | 1,22E-07 | 1,13E-06 | Down |

| | | | | | | |
|---|---|---|---|---|---|---|
| Nfil3 | 7839,401382 | 0,861966958 | 0,108399908 | 2,64E-17 | 1,14E-15 | Up |
| Simc1 | 1692,816062 | 0,749407211 | 0,110311065 | 4,61E-13 | 1,05E-11 | Up |
| Nop16 | 15847,78597 | 0,712563692 | 0,103935856 | 4,23E-13 | 9,69E-12 | Up |
| Unc5a | 833,5031363 | -1,482314373 | 0,208658689 | 8,38E-15 | 2,55E-13 | Down |
| Hk3 | 26800,88562 | 0,941985582 | 0,204244166 | 4,05E-08 | 4,11E-07 | Up |
| Prr7 | 516,0034562 | 1,379602548 | 0,12502584 | 2,85E-30 | 5,65E-28 | Up |
| Ctla2b | 974,1739352 | -1,558063547 | 0,216186844 | 3,62E-15 | 1,15E-13 | Down |
| Ctla2a | 2671,589779 | -1,18513269 | 0,18050931 | 2,77E-13 | 6,59E-12 | Down |
| Fbp2 | 13397,45589 | 1,75034596 | 0,192577547 | 4,18E-22 | 3,28E-20 | Up |
| Fbp1 | 506,5409413 | 2,765643773 | 0,307067138 | 1,03E-20 | 6,90E-19 | Up |
| Fancc | 5048,413678 | -1,364328122 | 0,149016669 | 4,93E-22 | 3,80E-20 | Down |
| Gm28557 | 187,792033 | -0,79963785 | 0,148894327 | 2,06E-09 | 2,58E-08 | Down |
| Zfp457 | 109,6044326 | 0,591351004 | 0,210496287 | 0,000467496 | 0,002078241 | Up |
| Zfp273 | 168,4897827 | -0,644005327 | 0,173213171 | 1,60E-05 | 9,83E-05 | Down |
| Mtrr | 6019,783425 | 0,649596748 | 0,052778485 | 9,21E-36 | 3,00E-33 | Up |
| Fastkd3 | 3201,382893 | 0,661771223 | 0,108927354 | 1,10E-10 | 1,70E-09 | Up |
| Nsun2 | 57730,03831 | 0,681647362 | 0,09795598 | 2,70E-13 | 6,45E-12 | Up |
| Clptm1l | 25078,75182 | 0,659021082 | 0,095793756 | 5,69E-13 | 1,28E-11 | Up |
| Trip13 | 10381,32846 | 0,722039017 | 0,092918654 | 4,36E-16 | 1,59E-14 | Up |
| Gpr150 | 2,049347692 | 2,200073069 | 1,949212916 | 0,00032791 | 0,001506869 | Up |
| Mctp1 | 306,1510972 | -1,329253073 | 0,301682689 | 4,64E-08 | 4,65E-07 | Down |
| Mblac2 | 543,9052679 | -0,685330331 | 0,132027021 | 1,46E-08 | 1,59E-07 | Down |
| Tmem161b | 1137,04339 | 0,691948949 | 0,11160293 | 3,94E-11 | 6,53E-10 | Up |
| Gm4117 | 181,6761914 | 0,701181049 | 0,191179898 | 1,27E-05 | 7,99E-05 | Up |
| Ssbp2 | 17549,14385 | -0,788286279 | 0,077633932 | 9,27E-26 | 1,15E-23 | Down |
| 4833422C13Rik | 219,3540343 | -1,091136542 | 0,231829292 | 1,50E-08 | 1,64E-07 | Down |
| F2r | 100569,7073 | -0,785341595 | 0,216552404 | 8,05E-06 | 5,28E-05 | Down |
| Gm6169 | 158,8930841 | 0,607349115 | 0,171635882 | 4,00E-05 | 0,00022599 | Up |
| Utp15 | 12195,55508 | 0,728876684 | 0,086544356 | 1,95E-18 | 9,67E-17 | Up |
| Tmem171 | 65,6614268 | 0,931799394 | 0,325190015 | 5,13E-05 | 0,000282665 | Up |
| Gm807 | 4,91661878 | 2,324662936 | 1,70008446 | 0,000253909 | 0,001197545 | Up |
| Smn1 | 7924,807772 | 0,651020557 | 0,089537265 | 3,66E-14 | 9,99E-13 | Up |
| Dimt1 | 2356,402769 | 0,974317367 | 0,14976124 | 5,56E-13 | 1,26E-11 | Up |
| Elovl7 | 1709,176708 | 0,710085838 | 0,161496731 | 5,81E-07 | 4,81E-06 | Up |
| Emb | 15483,94736 | 1,783782179 | 0,161821033 | 1,08E-30 | 2,20E-28 | Up |
| Ccl28 | 4,695982991 | 1,303856011 | 1,118454515 | 0,001201617 | 0,00483666 | Up |
| Hmgcs1 | 16890,43542 | -0,615185377 | 0,186548726 | 8,76E-05 | 0,000456924 | Down |
| Acox2 | 9,330206791 | 0,903904751 | 1,033651296 | 0,0041925 | 0,014639279 | Up |
| Slc4a7 | 6190,52347 | 0,819597885 | 0,061788953 | 7,68E-42 | 3,90E-39 | Up |
| Ndst2 | 7587,748567 | -1,125389824 | 0,108218251 | 1,24E-27 | 1,82E-25 | Down |
| Samd8 | 2754,828224 | 0,599936769 | 0,084324123 | 1,65E-13 | 4,06E-12 | Up |
| Kcnma1 | 68,5027925 | 0,815480493 | 0,842279455 | 0,004987487 | 0,017088461 | Up |
| Erc2 | 100,901895 | -0,614408183 | 0,41931327 | 0,007237873 | 0,023683574 | Down |
| Chdh | 23,90098117 | 1,444063047 | 0,855107683 | 0,000313711 | 0,001446716 | Up |
| Itih4 | 63,65729192 | -1,382566584 | 0,604562253 | 7,51E-05 | 0,000398261 | Down |
| Itih1 | 5,409161159 | 0,659342272 | 0,763754542 | 0,00969485 | 0,030436197 | Up |
| Gnl3 | 16777,88997 | 0,820396609 | 0,16079735 | 7,42E-09 | 8,54E-08 | Up |
| Sh3bp5 | 733,0241111 | 1,232997514 | 0,222536647 | 1,58E-10 | 2,37E-09 | Up |
| Eaf1 | 14776,4521 | 0,928993235 | 0,10436913 | 4,48E-21 | 3,11E-19 | Up |
| Colq | 46,55451075 | 1,522181721 | 0,310323869 | 3,74E-09 | 4,50E-08 | Up |
| Hacl1 | 2742,229687 | -0,620706088 | 0,153795346 | 5,39E-06 | 3,66E-05 | Down |

| | | | | | | |
|---|---|---|---|---|---|---|
| Timm23 | 17407,03583 | 0,677685933 | 0,107461613 | 2,26E-11 | 3,89E-10 | Up |
| 1810011H11Rik | 2581,982718 | 0,61463819 | 0,11820162 | 2,37E-08 | 2,51E-07 | Up |
| Gm7236 | 106,0647051 | 0,77715531 | 0,212959912 | 7,82E-06 | 5,15E-05 | Up |
| Txndc16 | 11,87922943 | -0,845336201 | 0,61327625 | 0,002613157 | 0,009648498 | Down |
| Cdkn3 | 756,721289 | -0,630063267 | 0,132917001 | 2,18E-07 | 1,92E-06 | Down |
| Ccnb1ip1 | 3382,037082 | 1,280949859 | 0,259409288 | 3,88E-09 | 4,66E-08 | Up |
| Gm26782 | 363,4083126 | 1,199346579 | 0,159809507 | 3,45E-16 | 1,28E-14 | Up |
| Rnase10 | 8,856081127 | 4,256973448 | 1,282628602 | 2,47E-06 | 1,78E-05 | Up |
| Rnase6 | 3,026418327 | 1,12161688 | 1,516911579 | 0,002210441 | 0,008324826 | Up |
| Zfp219 | 5413,154234 | -0,609587471 | 0,071403232 | 1,97E-18 | 9,72E-17 | Down |
| Gm4742 | 94,39653872 | 0,601386856 | 0,229797177 | 0,000743218 | 0,003146188 | Up |
| Olfr1509 | 53,89086866 | -0,849188772 | 0,265441036 | 2,55E-05 | 0,000149679 | Down |
| B230359F08Rik | 15,86769749 | -1,099215715 | 0,743573289 | 0,000973905 | 0,004012964 | Down |
| Traj61 | 2,525021882 | -0,810358089 | 1,268150264 | 0,005026823 | 0,017200639 | Down |
| Traj60 | 2,449826244 | -0,928845323 | 1,405142146 | 0,003386583 | 0,012136143 | Down |
| Traj59 | 3,331204144 | -2,66823082 | 1,995938005 | 0,000161933 | 0,000798169 | Down |
| Trac | 148,0237605 | -3,692993884 | 0,332973548 | 1,90E-29 | 3,38E-27 | Down |
| Abhd4 | 11072,85963 | -1,257409209 | 0,1113315 | 1,11E-31 | 2,45E-29 | Down |
| Slc7a7 | 7431,349218 | -1,700621742 | 0,175026386 | 1,31E-24 | 1,46E-22 | Down |
| Haus4 | 5870,874913 | -0,8027983 | 0,075205883 | 3,34E-28 | 5,25E-26 | Down |
| 4931414P19Rik | 1814,180241 | -0,771216317 | 0,08992314 | 3,39E-19 | 1,84E-17 | Down |
| Gm37271 | 94,0022124 | 1,063976417 | 0,364748368 | 2,62E-05 | 0,000153743 | Up |
| Slc22a17 | 1543,873298 | -0,610918572 | 0,107919409 | 1,91E-09 | 2,41E-08 | Down |
| Myh7 | 38,72488183 | 1,027412155 | 0,450496399 | 0,000193738 | 0,000939115 | Up |
| Cideb | 424,6151422 | -0,616464595 | 0,183790704 | 7,17E-05 | 0,000382764 | Down |
| Ltb4r2 | 305,3376719 | -0,726439817 | 0,250023437 | 0,000141563 | 0,000706538 | Down |
| Adcy4 | 351,8680846 | 1,214373003 | 0,155740804 | 3,70E-17 | 1,53E-15 | Up |
| Ripk3 | 124,2185648 | 3,856922134 | 0,333719655 | 7,89E-32 | 1,78E-29 | Up |
| Khnyn | 2851,77498 | -0,721931906 | 0,088428882 | 1,82E-17 | 7,96E-16 | Down |
| Mcpt1 | 70,48400238 | -4,102722763 | 0,476831025 | 5,95E-19 | 3,11E-17 | Down |
| Mcpt2 | 338,2446608 | -2,542296781 | 0,163755431 | 6,74E-56 | 8,20E-53 | Down |
| Mcpt4 | 8,384654525 | -2,887700725 | 0,927310991 | 5,98E-06 | 4,02E-05 | Down |
| Gzmc | 158,6019186 | -0,591301745 | 0,728448188 | 0,013411101 | 0,040136599 | Down |
| Gzmb | 5878,048741 | -2,168958429 | 0,377296391 | 3,32E-11 | 5,56E-10 | Down |
| 2410022M11Rik | 165,2247897 | -0,632046467 | 0,201864157 | 0,00013436 | 0,000673535 | Down |
| Cryl1 | 1527,099161 | -0,723059369 | 0,076087764 | 1,16E-22 | 9,81E-21 | Down |
| Xpo4 | 13478,40596 | 0,726853914 | 0,097891855 | 5,97E-15 | 1,85E-13 | Up |
| Fgf9 | 1434,587232 | 1,184203457 | 0,079387929 | 1,62E-52 | 1,64E-49 | Up |
| Nupl1 | 9943,899575 | 0,632038672 | 0,071097511 | 7,56E-20 | 4,50E-18 | Up |
| Fdft1 | 7085,519987 | -0,590063187 | 0,187429834 | 0,000169105 | 0,000831726 | Down |
| Gm15918 | 25,26934198 | -1,100797814 | 0,551652808 | 0,000311524 | 0,001438444 | Down |
| Zfp395 | 2940,721273 | -0,898311748 | 0,127700392 | 2,20E-14 | 6,19E-13 | Down |
| Nuggc | 871,5562984 | -0,722554852 | 0,19288422 | 8,03E-06 | 5,28E-05 | Down |
| Dpysl2 | 3918,68976 | -0,729462023 | 0,094003832 | 4,42E-16 | 1,61E-14 | Down |
| Bnip3l | 10745,29701 | -0,585243586 | 0,092780196 | 4,40E-11 | 7,26E-10 | Down |
| Egr3 | 20,35365811 | 1,005930239 | 0,524126284 | 0,000506257 | 0,002229388 | Up |
| Slc39a14 | 6858,21372 | 1,121917458 | 0,110152912 | 1,14E-26 | 1,51E-24 | Up |
| Piwil2 | 1039,508185 | 0,68730659 | 0,124300587 | 2,25E-09 | 2,80E-08 | Up |
| Polr3d | 3655,168708 | 1,1079118 | 0,144999466 | 1,09E-16 | 4,25E-15 | Up |
| Dok2 | 12679,57408 | 0,782238982 | 0,088785129 | 3,86E-20 | 2,36E-18 | Up |
| Gm4285 | 147,8286904 | -0,674943913 | 0,180213245 | 1,15E-05 | 7,31E-05 | Down |

| | | | | | | |
|---|---|---|---|---|---|---|
| Tnfsf11 | 578,5572351 | 0,582909667 | 0,165905811 | 5,15E-05 | 0,000283352 | Up |
| Bora | 18311,78373 | -0,890945495 | 0,117343168 | 3,56E-16 | 1,31E-14 | Down |
| Dis3 | 11397,47899 | 0,732418402 | 0,109110127 | 9,44E-13 | 2,05E-11 | Up |
| Tbc1d4 | 1398,062824 | -0,63061565 | 0,160062484 | 7,43E-06 | 4,91E-05 | Down |
| Uchl3 | 8828,901475 | 0,644883397 | 0,086931173 | 1,28E-14 | 3,73E-13 | Up |
| Acod1 | 37,89133951 | 3,545995395 | 0,48627212 | 3,25E-14 | 8,97E-13 | Up |
| Scel | 103,6608156 | 0,705619671 | 0,260226205 | 0,000286416 | 0,001332958 | Up |
| Slain1 | 4991,394592 | -0,709570849 | 0,098935812 | 4,57E-14 | 1,22E-12 | Down |
| Dzip1 | 190,1343369 | -1,344373293 | 0,193994883 | 2,81E-14 | 7,82E-13 | Down |
| Mbnl2 | 4902,590424 | -0,782823173 | 0,187990456 | 9,17E-07 | 7,27E-06 | Down |
| Fyb | 1338,586815 | -0,958024743 | 0,394823261 | 0,000169864 | 0,000835232 | Down |
| Gm7666 | 993,1694886 | 0,626208111 | 0,134188073 | 3,18E-07 | 2,74E-06 | Up |
| Tars | 30479,99523 | 0,607702518 | 0,077525454 | 6,51E-16 | 2,30E-14 | Up |
| Mtmr12 | 10351,46479 | -0,988819283 | 0,092082881 | 3,65E-29 | 6,24E-27 | Down |
| Trio | 13127,54574 | 0,864395526 | 0,074589235 | 6,16E-33 | 1,59E-30 | Up |
| Dap | 10614,41672 | -0,589318811 | 0,137729762 | 2,38E-06 | 1,73E-05 | Down |
| Ropn1l | 180,3430499 | -0,892165869 | 0,226414337 | 1,15E-06 | 8,90E-06 | Down |
| Mtdh | 38220,90968 | 0,72363495 | 0,096571454 | 3,67E-15 | 1,16E-13 | Up |
| Pop1 | 6346,479171 | 0,607851678 | 0,110823664 | 5,28E-09 | 6,23E-08 | Up |
| Grhl2 | 2,202783119 | 0,705599985 | 1,175321239 | 0,007004128 | 0,023030137 | Up |
| Slc25a32 | 3606,511636 | 0,797073411 | 0,087840291 | 3,05E-21 | 2,17E-19 | Up |
| Lrp12 | 26105,19814 | 0,837098124 | 0,150569638 | 5,16E-10 | 7,16E-09 | Up |
| Oxr1 | 1720,685016 | -0,653771915 | 0,122342522 | 8,24E-09 | 9,41E-08 | Down |
| Nov | 32,77256783 | 4,739883924 | 0,852754163 | 5,24E-10 | 7,25E-09 | Up |
| Deptor | 7619,750823 | 0,732926866 | 0,114784571 | 8,34E-12 | 1,54E-10 | Up |
| Mtss1 | 4672,466355 | -0,782151999 | 0,108562792 | 1,80E-14 | 5,13E-13 | Down |
| Myc | 52082,33665 | 0,662753849 | 0,089210407 | 1,02E-14 | 3,05E-13 | Up |
| Gsdmc4 | 20,8131417 | 0,82076055 | 0,395265293 | 0,000762951 | 0,00322076 | Up |
| Sla | 19085,22512 | -1,366844647 | 0,153300794 | 4,30E-21 | 3,00E-19 | Down |
| St3gal1 | 9612,884407 | -1,229416528 | 0,113899342 | 2,57E-29 | 4,44E-27 | Down |
| Ptp4a3 | 19190,23997 | 0,773276804 | 0,131719827 | 1,44E-10 | 2,17E-09 | Up |
| Ly6a | 28,32201796 | 3,597937969 | 0,695983765 | 9,72E-09 | 1,09E-07 | Up |
| Ly6g | 18,38853675 | 1,641063363 | 1,019047721 | 0,000301758 | 0,001397947 | Up |
| 9030619P08Rik | 3853,176387 | -2,12595254 | 0,161464014 | 3,67E-42 | 1,92E-39 | Down |
| Top1mt | 1309,388148 | 0,877822594 | 0,174430571 | 6,96E-09 | 8,07E-08 | Up |
| Tigd5 | 83,52348364 | -0,639744738 | 0,226387122 | 0,000321541 | 0,001480157 | Down |
| Maf1 | 7023,752595 | -0,774921624 | 0,100895794 | 5,27E-16 | 1,89E-14 | Down |
| Bop1 | 41096,09342 | 0,698510799 | 0,083713365 | 5,04E-18 | 2,37E-16 | Up |
| Apol10b | 333,8324629 | -0,899454611 | 0,225814458 | 9,19E-07 | 7,29E-06 | Down |
| Csf2rb2 | 86409,2453 | -0,648982992 | 0,163303494 | 5,67E-06 | 3,83E-05 | Down |
| Tst | 254,111652 | -0,64750592 | 0,197134989 | 7,31E-05 | 0,000388639 | Down |
| Kctd17 | 16,47963295 | 0,831560296 | 0,563049444 | 0,002364055 | 0,008841363 | Up |
| Il2rb | 2124,240064 | 2,103332061 | 0,156995339 | 1,57E-43 | 9,25E-41 | Up |
| Mfng | 9836,646962 | 0,919073204 | 0,087392884 | 5,99E-28 | 8,97E-26 | Up |
| 1700088E04Rik | 18,59398645 | -0,709383261 | 0,446097811 | 0,003519875 | 0,012569408 | Down |
| Maff | 559,8860758 | 1,593819489 | 0,163773042 | 1,67E-24 | 1,83E-22 | Up |
| Gm16576 | 461,5269038 | -0,723419051 | 0,14530487 | 3,19E-08 | 3,29E-07 | Down |
| Nptxr | 14,70139706 | -0,665995808 | 0,606012862 | 0,008025277 | 0,025870717 | Down |
| Atf4 | 54340,73496 | 0,704690219 | 0,148825642 | 1,25E-07 | 1,16E-06 | Up |
| Grap2 | 44,71983661 | -1,757000052 | 0,539491423 | 2,50E-06 | 1,81E-05 | Down |
| Tnrc6b | 9484,276762 | -0,691560988 | 0,107662767 | 9,43E-12 | 1,73E-10 | Down |

| | | | | | | |
|---|---|---|---|---|---|---|
| Mkl1 | 9561,340033 | -1,044554653 | 0,121382963 | 3,83E-20 | 2,36E-18 | Down |
| 1110025M09Rik | 53,80687314 | -0,967616247 | 0,254918392 | 1,45E-06 | 1,10E-05 | Down |
| Gm17597 | 14,44893181 | 0,635688414 | 0,548592267 | 0,008908875 | 0,028289667 | Up |
| Nhp2l1 | 19911,6971 | 0,620582087 | 0,083600504 | 1,48E-14 | 4,26E-13 | Up |
| Sep 03 | 1811,656838 | -0,724800663 | 0,177515906 | 2,03E-06 | 1,49E-05 | Down |
| Fam109b | 8,589926927 | -0,668618057 | 0,660884061 | 0,008488985 | 0,027151733 | Down |
| Cyb5r3 | 8163,948955 | -0,704401645 | 0,125419331 | 1,19E-09 | 1,54E-08 | Down |
| Ttll1 | 2638,371487 | -0,650896961 | 0,086239862 | 4,59E-15 | 1,43E-13 | Down |
| 5031439G07Rik | 24803,44948 | -0,85597685 | 0,148312579 | 1,28E-10 | 1,96E-09 | Down |
| Ttc38 | 117,9557906 | -0,634440425 | 0,33722991 | 0,003246065 | 0,011706089 | Down |
| Creld2 | 9921,911313 | 0,59142759 | 0,106840366 | 4,42E-09 | 5,27E-08 | Up |
| Pim3 | 6,282194326 | 1,995442028 | 1,601576155 | 0,000429128 | 0,001921709 | Up |
| Adm2 | 23,83837331 | 3,249121343 | 1,508828004 | 3,75E-05 | 0,000213582 | Up |
| Odf3b | 398,2750906 | -0,645025909 | 0,137952103 | 2,66E-07 | 2,32E-06 | Down |
| Shank3 | 691,8001706 | 2,569582015 | 0,265180939 | 7,54E-24 | 7,52E-22 | Up |
| Prickle1 | 609,2121696 | -1,503436158 | 0,227063266 | 2,19E-13 | 5,28E-12 | Down |
| Pus7l | 4352,454358 | 0,78397579 | 0,091092321 | 2,30E-19 | 1,28E-17 | Up |
| Hdac7 | 1188,666123 | 0,752682442 | 0,316493907 | 0,000525806 | 0,002308238 | Up |
| Vdr | 1600,252529 | 0,848701013 | 0,156903683 | 1,11E-09 | 1,45E-08 | Up |
| 1700120C14Rik | 871,9895736 | 1,146703858 | 0,298686293 | 6,94E-07 | 5,63E-06 | Up |
| Mettl7a1 | 3337,095574 | -2,641334656 | 0,229023807 | 3,50E-32 | 8,30E-30 | Down |
| C330013E15Rik | 87,38201403 | -0,867183309 | 0,237755489 | 4,39E-06 | 3,04E-05 | Down |
| Galnt6 | 13781,09769 | 1,212314396 | 0,096500293 | 2,33E-38 | 1,04E-35 | Up |
| Acvrl1 | 417,3961895 | 3,851479419 | 0,428183677 | 2,54E-20 | 1,62E-18 | Up |
| Grasp | 173,9788717 | 0,852914166 | 0,205572138 | 5,94E-07 | 4,90E-06 | Up |
| Krt87 | 31,74655852 | 0,808742138 | 0,480593395 | 0,001830802 | 0,007036822 | Up |
| Krt86 | 5,385063062 | 1,156478075 | 1,087559973 | 0,001854967 | 0,007115187 | Up |
| Tns2 | 900,18026 | -0,648580046 | 0,14244047 | 4,60E-07 | 3,87E-06 | Down |
| Csad | 543,1088425 | -1,145514617 | 0,125938927 | 4,87E-22 | 3,77E-20 | Down |
| Sp7 | 10,34123389 | 1,281945789 | 0,824759098 | 0,00055686 | 0,002432284 | Up |
| Prr13 | 7754,026831 | -0,611755007 | 0,180098998 | 6,40E-05 | 0,000346349 | Down |
| Calcoco1 | 1748,419943 | -1,487823099 | 0,156633362 | 2,03E-23 | 1,92E-21 | Down |
| Nfe2 | 33328,36694 | -0,941554363 | 0,15526955 | 1,18E-11 | 2,12E-10 | Down |
| Gpr84 | 315,0824204 | 3,686265646 | 0,311081111 | 3,03E-33 | 8,03E-31 | Up |
| Zfp385a | 4708,771623 | -1,397744158 | 0,152414026 | 4,45E-22 | 3,46E-20 | Down |
| Coro7 | 8024,278426 | 0,609671952 | 0,132413344 | 4,79E-07 | 4,02E-06 | Up |
| 4930451G09Rik | 58,58546048 | -0,674526318 | 0,282111368 | 0,000819744 | 0,003438255 | Down |
| Ciita | 35,0702241 | 0,860446445 | 0,438379092 | 0,000818906 | 0,00343553 | Up |
| Socs1 | 378,5095863 | 2,194662056 | 0,185690222 | 1,11E-34 | 3,37E-32 | Up |
| Rsl1d1 | 31354,00282 | 0,766609492 | 0,093862587 | 1,14E-17 | 5,10E-16 | Up |
| Snx29 | 528,914144 | -1,272069204 | 0,17293815 | 1,21E-15 | 4,07E-14 | Down |
| Mkl2 | 9248,303095 | -0,956025985 | 0,124938793 | 1,44E-16 | 5,54E-15 | Down |
| Gm15738 | 9,261497828 | -0,583933881 | 0,503743034 | 0,011909304 | 0,03623264 | Down |
| Rrn3 | 15526,28427 | 0,643675018 | 0,081135025 | 2,36E-16 | 8,92E-15 | Up |
| Cebpd | 1386,681711 | 0,716033135 | 0,127227056 | 1,00E-09 | 1,32E-08 | Up |
| Gm15714 | 84,59786768 | 0,832579362 | 0,257250062 | 2,47E-05 | 0,00014521 | Up |
| Vpreb1 | 21,9782659 | -1,356699672 | 0,534895109 | 3,93E-05 | 0,000222602 | Down |
| Rimbp3 | 353,1258945 | -0,681603562 | 0,155112515 | 7,38E-07 | 5,96E-06 | Down |
| Tmem191c | 126,9352272 | -0,726990034 | 0,345822166 | 0,001181881 | 0,004765622 | Down |
| Vpreb2 | 4,469127453 | -0,93334413 | 1,283479382 | 0,003878864 | 0,013688079 | Down |
| B3gnt5 | 2155,836412 | -0,879075451 | 0,103085292 | 1,81E-19 | 1,02E-17 | Down |

| | | | | | | |
|---|---|---|---|---|---|---|
| Klhl24 | 2207,660239 | -1,328856065 | 0,140290561 | 2,41E-23 | 2,25E-21 | Down |
| Alg3 | 6360,774689 | 0,770747705 | 0,094645037 | 1,33E-17 | 5,91E-16 | Up |
| Ece2 | 5586,088202 | 0,621221314 | 0,086314115 | 7,85E-14 | 2,03E-12 | Up |
| Fam131a | 401,8423097 | -0,878656289 | 0,181554458 | 1,88E-08 | 2,03E-07 | Down |
| Tmem41a | 1718,671098 | 0,688212533 | 0,086721468 | 1,60E-16 | 6,13E-15 | Up |
| Etv5 | 5632,118427 | 3,14224839 | 0,175690228 | 2,93E-72 | 1,34E-68 | Up |
| Gm15651 | 39,44863053 | 0,890358465 | 0,448188604 | 0,000680853 | 0,002908478 | Up |
| St6gal1 | 834,2815111 | 2,958217172 | 0,506877244 | 1,75E-10 | 2,61E-09 | Up |
| Rtp4 | 8,506784885 | 1,758677258 | 1,551030255 | 0,000658719 | 0,00282516 | Up |
| Gp5 | 13,93274348 | -0,614998155 | 0,459618707 | 0,008414969 | 0,026969843 | Down |
| Atp13a3 | 17060,59201 | 0,743444689 | 0,055193129 | 1,11E-42 | 5,95E-40 | Up |
| Gm15694 | 57,10517763 | 0,995359604 | 0,295019857 | 6,78E-06 | 4,51E-05 | Up |
| Bex6 | 86,47031393 | -0,793818121 | 0,40818356 | 0,001168078 | 0,004714129 | Down |
| Heg1 | 3249,865821 | -1,044065047 | 0,202487875 | 1,61E-09 | 2,05E-08 | Down |
| Gm15658 | 25,83763082 | -0,779197912 | 0,376423272 | 0,000955538 | 0,003945295 | Down |
| Gm15657 | 115,7387431 | -1,541635847 | 0,280453416 | 1,76E-10 | 2,63E-09 | Down |
| Muc13 | 24196,78658 | -1,065607853 | 0,1817703 | 2,60E-11 | 4,44E-10 | Down |
| Mylk | 5865,78619 | -0,889010126 | 0,224804131 | 1,10E-06 | 8,57E-06 | Down |
| Hacd2 | 5490,494879 | 0,665270085 | 0,138481308 | 1,23E-07 | 1,14E-06 | Up |
| Stfa2l1 | 107,2059199 | 5,283502109 | 0,512103909 | 5,56E-27 | 7,76E-25 | Up |
| Gm5483 | 452,7039256 | 5,790382001 | 0,641238929 | 5,71E-22 | 4,38E-20 | Up |
| Gm5416 | 22,91861969 | 4,361846674 | 0,772711128 | 5,72E-10 | 7,89E-09 | Up |
| 2010005H15Rik | 294,9231517 | 6,561445292 | 0,723620917 | 1,09E-22 | 9,36E-21 | Up |
| Stfa1 | 435,2804446 | 5,045021572 | 0,7327471 | 7,94E-14 | 2,05E-12 | Up |
| BC117090 | 4,993770928 | 0,605654317 | 0,84509182 | 0,012562213 | 0,037900403 | Up |
| BC100530 | 158,3176464 | 5,143477407 | 0,737389655 | 3,49E-14 | 9,57E-13 | Up |
| Stfa2 | 66,66232615 | 4,450804112 | 0,519569599 | 4,83E-19 | 2,56E-17 | Up |
| Stfa3 | 23,35549363 | 4,280147735 | 0,714903236 | 9,24E-11 | 1,45E-09 | Up |
| Fbxo40 | 950,1072435 | -0,662000914 | 0,097363949 | 9,69E-13 | 2,10E-11 | Down |
| Maats1 | 2,649988089 | 1,308680974 | 1,564042893 | 0,001572291 | 0,00615766 | Up |
| Pla1a | 16,4083965 | 1,572731107 | 0,930446805 | 0,00026692 | 0,001250831 | Up |
| Cd80 | 93,93014878 | 3,230992392 | 0,316169528 | 2,45E-25 | 2,94E-23 | Up |
| B4galt4 | 217,0944193 | 0,754706011 | 0,142360123 | 4,44E-09 | 5,29E-08 | Up |
| Qtrtd1 | 3150,644814 | 0,795785988 | 0,112530193 | 4,14E-14 | 1,12E-12 | Up |
| Gramd1c | 174,3401074 | -0,845480217 | 0,245140444 | 1,06E-05 | 6,81E-05 | Down |
| Cd200r4 | 1682,79966 | -0,912241945 | 0,243391804 | 2,29E-06 | 1,67E-05 | Down |
| Cd200r2 | 30,52187736 | 0,629905339 | 0,556676924 | 0,00939351 | 0,029607229 | Up |
| Btla | 870,6273683 | 1,141717265 | 0,207594093 | 2,02E-10 | 3,00E-09 | Up |
| Nxpe3 | 5796,237967 | -0,863214271 | 0,135207552 | 2,58E-12 | 5,19E-11 | Down |
| Zbtb11os1 | 261,0842517 | -0,902882833 | 0,213219422 | 2,98E-07 | 2,58E-06 | Down |
| Trmt10c | 4211,665144 | 0,626941799 | 0,080797287 | 1,06E-15 | 3,60E-14 | Up |
| Senp7 | 2356,591948 | -0,662870189 | 0,061281391 | 2,85E-28 | 4,53E-26 | Down |
| Gpr15 | 254,8754746 | 0,665337805 | 0,280823912 | 0,000917373 | 0,003804919 | Up |
| Gm9027 | 90,90545374 | 0,673616484 | 0,198399918 | 4,16E-05 | 0,000233998 | Up |
| 1700063K16Rik | 4,080562336 | -1,007056208 | 1,219336499 | 0,003284349 | 0,011813824 | Down |
| Jam2 | 79,40043214 | 0,657630979 | 0,249645706 | 0,000486855 | 0,002156425 | Up |
| Cct8 | 74362,13212 | 0,620267928 | 0,069018636 | 3,42E-20 | 2,15E-18 | Up |
| 2310061N02Rik | 28,73899619 | 2,013264348 | 0,488935429 | 9,88E-08 | 9,41E-07 | Up |
| Krtap13 | 7,096016804 | 1,933706024 | 1,16816894 | 0,000215609 | 0,001034811 | Up |
| 2310034C09Rik | 9,532623147 | 2,364249519 | 1,018774851 | 3,82E-05 | 0,000216856 | Up |
| Krtap15 | 8,747682322 | 0,60112258 | 0,576368335 | 0,011649632 | 0,035564289 | Up |

| | | | | | | |
|---|---|---|---|---|---|---|
| Urb1 | 4311,844986 | 0,592275386 | 0,145059891 | 5,38E-06 | 3,65E-05 | Up |
| A930006K02Rik | 72,34929999 | -0,652913492 | 0,252872884 | 0,000577928 | 0,00251229 | Down |
| Ifngr2 | 4510,296374 | 0,613092435 | 0,110860179 | 3,95E-09 | 4,74E-08 | Up |
| Dnajc28 | 41,83256624 | 0,688746054 | 0,316651826 | 0,001255214 | 0,00503243 | Up |
| Gart | 43874,58988 | 0,806903705 | 0,117833406 | 1,83E-13 | 4,43E-12 | Up |
| Itsn1 | 572,1132072 | -0,634312948 | 0,49831179 | 0,008072845 | 0,025996889 | Down |
| Slc5a3 | 1303,674779 | 0,616869422 | 0,101601099 | 1,58E-10 | 2,38E-09 | Up |
| Wrb | 2437,685251 | 0,975580416 | 0,118055135 | 8,98E-19 | 4,57E-17 | Up |
| Arid1b | 10266,32432 | -0,592095727 | 0,094781281 | 6,22E-11 | 1,00E-09 | Down |
| Tulp4 | 2056,861606 | 0,997485673 | 0,143351654 | 2,17E-14 | 6,11E-13 | Up |
| Sytl3 | 33,67731622 | 1,814075922 | 0,492889755 | 5,41E-07 | 4,51E-06 | Up |
| Igf2r | 1450,221384 | 1,782024433 | 0,51277477 | 1,16E-06 | 8,96E-06 | Up |
| Acat2 | 11616,98755 | -0,610367912 | 0,152502818 | 6,64E-06 | 4,42E-05 | Down |
| Fpr1 | 2,360319834 | 1,802504367 | 1,927251819 | 0,000573962 | 0,002496948 | Up |
| Hcfc1r1 | 1309,510573 | -0,763894228 | 0,121005124 | 9,99E-12 | 1,83E-10 | Down |
| Tnfrsf12a | 1265,200175 | 1,05252685 | 0,169988201 | 3,40E-12 | 6,63E-11 | Up |
| Amdhd2 | 1756,621887 | -0,582786152 | 0,178945782 | 0,000124582 | 0,000628549 | Down |
| D330041H03Rik | 116,2073629 | -0,77179886 | 0,261386069 | 9,14E-05 | 0,00047451 | Down |
| Slc9a3r2 | 17,79518977 | 0,830442263 | 0,424962257 | 0,000958216 | 0,003953487 | Up |
| Syngr3 | 17,62875654 | 0,607020099 | 0,444174119 | 0,008543646 | 0,027276935 | Up |
| Tbl3 | 16620,00934 | 0,763997506 | 0,099473339 | 5,87E-16 | 2,09E-14 | Up |
| Rnf151 | 143,8642516 | 0,595381011 | 0,256836419 | 0,001639269 | 0,006372144 | Up |
| Ift140 | 5718,39304 | -0,759770146 | 0,070034203 | 7,98E-29 | 1,35E-26 | Down |
| Tpsab1 | 9,015693058 | 2,86442018 | 1,295692047 | 4,31E-05 | 0,000241538 | Up |
| Haghl | 754,6905472 | -0,793807945 | 0,208928966 | 3,87E-06 | 2,71E-05 | Down |
| Wdr90 | 3890,75529 | -0,624451818 | 0,165653639 | 1,51E-05 | 9,36E-05 | Down |
| Fam195a | 3222,516533 | 0,714079599 | 0,114112964 | 2,25E-11 | 3,88E-10 | Up |
| Nhlrc4 | 124,3654615 | -0,908803759 | 0,311620892 | 4,84E-05 | 0,000267895 | Down |
| Rab11fip3 | 208,3868459 | -1,457669265 | 0,260070016 | 1,06E-10 | 1,65E-09 | Down |
| Crebrf | 888,3180466 | -1,144181866 | 0,220272924 | 1,10E-09 | 1,44E-08 | Down |
| Pacsin1 | 98,84726718 | 1,314959286 | 0,445537775 | 1,22E-05 | 7,72E-05 | Up |
| Gm20109 | 23,66754949 | -0,796187541 | 0,366153391 | 0,000685942 | 0,002928847 | Down |
| E230001N04Rik | 814,5500931 | -0,58265754 | 0,097237075 | 3,22E-10 | 4,66E-09 | Down |
| Cdkn1a | 27392,66737 | 0,876368539 | 0,136377321 | 1,76E-12 | 3,67E-11 | Up |
| Cpne5 | 368,5817459 | -0,826040595 | 0,23984748 | 1,22E-05 | 7,72E-05 | Down |
| Pim1 | 35421,75936 | 0,623450034 | 0,099808311 | 5,06E-11 | 8,25E-10 | Up |
| 1700097N02Rik | 188,4621016 | 2,333002096 | 0,444634839 | 7,79E-10 | 1,04E-08 | Up |
| Dnah8 | 445,878478 | 2,416553634 | 0,3003507 | 7,77E-18 | 3,55E-16 | Up |
| Glp1r | 48,06580039 | 2,667195104 | 0,717216608 | 9,29E-07 | 7,36E-06 | Up |
| Abcg1 | 18716,36396 | -0,585043198 | 0,152998639 | 1,60E-05 | 9,81E-05 | Down |
| Wdr4 | 8949,026876 | 0,712787695 | 0,140780387 | 2,26E-08 | 2,40E-07 | Up |
| Cbs | 5,549943991 | 1,090654081 | 1,030394405 | 0,002137594 | 0,008087163 | Up |
| Pdxk-ps | 22,29220025 | -0,598304866 | 0,386296 | 0,007117132 | 0,02333448 | Down |
| Notch3 | 180,1590764 | -1,294814179 | 0,274248036 | 1,11E-08 | 1,24E-07 | Down |
| Cyp4f39 | 17,51273588 | 1,356931678 | 0,699991929 | 0,000195142 | 0,000944023 | Up |
| Cyp4f40 | 4,755755275 | 0,944436353 | 1,357057197 | 0,003543913 | 0,012647828 | Up |
| Cd320 | 3100,200885 | 0,70473127 | 0,102521512 | 4,00E-13 | 9,22E-12 | Up |
| Psmb9 | 6708,266725 | -1,221264296 | 0,146810846 | 5,44E-19 | 2,86E-17 | Down |
| Tap1 | 9558,796644 | -1,339974904 | 0,133461398 | 9,60E-26 | 1,18E-23 | Down |
| Psmb8 | 18413,12469 | -0,912430256 | 0,135688906 | 1,79E-13 | 4,34E-12 | Down |
| Tap2 | 24397,07219 | -1,569345023 | 0,15272592 | 7,67E-27 | 1,04E-24 | Down |

| | | | | | | |
|---|---|---|---|---|---|---|
| Gm15821 | 68,2584575 | -3,140982546 | 0,471375011 | 1,80E-12 | 3,73E-11 | Down |
| H2-Ob | 188,701423 | -2,281531501 | 0,372360925 | 4,36E-12 | 8,37E-11 | Down |
| Fkbpl | 952,7969669 | -0,859078094 | 0,151928291 | 2,50E-10 | 3,66E-09 | Down |
| C2 | 114,3745269 | 0,705367825 | 0,252466005 | 0,000226581 | 0,001080373 | Up |
| Msh5 | 205,4298374 | -1,182028659 | 0,358935269 | 5,20E-06 | 3,54E-05 | Down |
| Aif1 | 158,4295468 | 0,776616669 | 0,205023079 | 4,57E-06 | 3,15E-05 | Up |
| Tnf | 575,7083511 | 2,472554113 | 0,398539516 | 4,94E-12 | 9,44E-11 | Up |
| Gm20443 | 18,38509011 | -2,705757525 | 0,503295323 | 1,07E-09 | 1,40E-08 | Down |
| Mrps18b | 4295,98223 | 0,662870437 | 0,078708176 | 3,56E-18 | 1,71E-16 | Up |
| 2410017I17Rik | 6,007788131 | -1,761179709 | 1,109227508 | 0,000286004 | 0,00133138 | Down |
| Crisp1 | 11,46684021 | 4,384748863 | 1,582663925 | 5,47E-06 | 3,71E-05 | Up |
| Enpp5 | 1931,277406 | -0,929099825 | 0,205573537 | 6,79E-08 | 6,63E-07 | Down |
| Slc29a1 | 6326,353284 | 0,985555384 | 0,127257752 | 6,01E-17 | 2,43E-15 | Up |
| Klc4 | 2245,837766 | -0,794590999 | 0,128950719 | 1,97E-11 | 3,44E-10 | Down |
| Bysl | 13254,14554 | 0,887592222 | 0,110492037 | 1,09E-17 | 4,88E-16 | Up |
| A530064D06Rik | 15,41685562 | 1,634756918 | 0,871493463 | 0,000157974 | 0,000781823 | Up |
| Trem1 | 776,7634029 | 2,095024524 | 0,275711409 | 9,23E-17 | 3,66E-15 | Up |
| Trem3 | 390,164595 | 3,4913904 | 0,27462945 | 8,35E-38 | 3,45E-35 | Up |
| Treml4 | 45,07301518 | 1,843786235 | 0,463647193 | 1,71E-07 | 1,53E-06 | Up |
| Trem2 | 69,74837705 | 3,90127114 | 0,796929884 | 3,06E-08 | 3,17E-07 | Up |
| Tbc1d5 | 11056,20514 | -0,738375153 | 0,093216563 | 1,13E-16 | 4,38E-15 | Down |
| Mrps36-ps1 | 34,22580135 | -0,594727522 | 0,284090307 | 0,002725878 | 0,009999997 | Down |
| Stap2 | 1615,345631 | 1,651503544 | 0,098384681 | 2,13E-65 | 3,89E-62 | Up |
| Mpnd | 8047,801588 | 0,840223679 | 0,100939444 | 1,48E-18 | 7,40E-17 | Up |
| Sema6b | 3648,920805 | 0,620089904 | 0,115414287 | 9,03E-09 | 1,02E-07 | Up |
| Kdm4b | 2232,26203 | -0,594842834 | 0,113920418 | 2,41E-08 | 2,55E-07 | Down |
| Vmac | 530,7294514 | -1,053391719 | 0,130403582 | 3,35E-18 | 1,62E-16 | Down |
| Rfx2 | 198,2124184 | -0,956324471 | 0,236286197 | 5,21E-07 | 4,35E-06 | Down |
| Tnfsf14 | 2354,701557 | 1,278351484 | 0,13740596 | 1,04E-22 | 8,92E-21 | Up |
| C3 | 3832,766568 | 2,968743695 | 0,380916325 | 4,48E-16 | 1,63E-14 | Up |
| Trip10 | 4583,055595 | 0,778142102 | 0,10068199 | 3,50E-16 | 1,29E-14 | Up |
| Adgre1 | 31351,15199 | 2,467901075 | 0,162970645 | 1,51E-53 | 1,62E-50 | Up |
| Lpin2 | 12522,65665 | -0,762358186 | 0,086096279 | 3,19E-20 | 2,01E-18 | Down |
| Wdr43 | 29434,32735 | 0,742444491 | 0,082222576 | 8,01E-21 | 5,42E-19 | Up |
| Ypel5 | 2742,981752 | -0,814055238 | 0,209311697 | 2,33E-06 | 1,70E-05 | Down |
| Lbh | 2209,153057 | -1,032383316 | 0,153536573 | 1,02E-13 | 2,57E-12 | Down |
| Rasgrp3 | 128,5222732 | 1,499407694 | 0,964380931 | 0,000401066 | 0,001806675 | Up |
| Zfp36l2 | 36259,45751 | -0,734323427 | 0,100215669 | 1,15E-14 | 3,40E-13 | Down |
| Lrpprc | 21501,77826 | 0,589292626 | 0,07653802 | 2,19E-15 | 7,17E-14 | Up |
| 1110020A21Rik | 81,41906703 | -0,948588374 | 0,26902911 | 4,63E-06 | 3,19E-05 | Down |
| Rhoq | 4659,410248 | 0,679366096 | 0,143363526 | 1,51E-07 | 1,38E-06 | Up |
| Pigf | 1045,799675 | 0,628868002 | 0,113135577 | 3,02E-09 | 3,67E-08 | Up |
| Socs5 | 4691,970377 | 0,603530444 | 0,076798168 | 5,71E-16 | 2,04E-14 | Up |
| Mcfd2 | 6634,911829 | 0,697750966 | 0,12960285 | 4,64E-09 | 5,50E-08 | Up |
| Nrxn1 | 1027,814435 | -1,010798055 | 0,171944061 | 2,72E-11 | 4,62E-10 | Down |
| Svil | 19366,24402 | -0,646022426 | 0,109923923 | 4,17E-10 | 5,88E-09 | Down |
| Cables1 | 446,507358 | -1,438072206 | 0,257903297 | 1,27E-10 | 1,94E-09 | Down |
| Lama3 | 18,10368665 | 0,617613584 | 0,479041449 | 0,008728414 | 0,027769737 | Up |
| Ttc39c | 3820,98171 | -0,72640665 | 0,1365132 | 5,10E-09 | 6,03E-08 | Down |
| Cabyr | 87,62900569 | -1,586973441 | 0,314264601 | 1,70E-09 | 2,16E-08 | Down |
| Osbpl1a | 5962,158391 | -0,906348607 | 0,109573898 | 1,30E-18 | 6,50E-17 | Down |

| | | | | | | |
|---|---|---|---|---|---|---|
| Gm6457 | 90,24875753 | 0,706765784 | 0,218565516 | 5,69E-05 | 0,000311197 | Up |
| Taf4b | 3543,417242 | 0,708557761 | 0,080738195 | 1,09E-19 | 6,21E-18 | Up |
| Kctd1 | 1682,27025 | -0,686753758 | 0,13474708 | 2,36E-08 | 2,49E-07 | Down |
| Slc39a6 | 7023,671152 | 1,056414663 | 0,088510491 | 3,52E-35 | 1,11E-32 | Up |
| Celf4 | 147,4746177 | 3,35944794 | 0,588903987 | 6,32E-10 | 8,63E-09 | Up |
| Bin1 | 1945,858156 | -0,639600533 | 0,127080341 | 4,74E-08 | 4,74E-07 | Down |
| Gypc | 10769,16681 | -0,747662252 | 0,103709136 | 2,43E-14 | 6,81E-13 | Down |
| Wdr36 | 17354,67748 | 0,608286187 | 0,060302318 | 9,35E-25 | 1,07E-22 | Up |
| Hspa9 | 115364,2864 | 0,65472097 | 0,061002701 | 7,60E-28 | 1,12E-25 | Up |
| Paip2 | 8834,786945 | -0,684368209 | 0,111137861 | 5,49E-11 | 8,93E-10 | Down |
| 1700066B19Rik | 786,8202738 | -0,650030054 | 0,167057763 | 7,84E-06 | 5,16E-05 | Down |
| Cxxc5 | 8768,051505 | -0,898060146 | 0,097305818 | 2,76E-22 | 2,20E-20 | Down |
| Gm29417 | 51,81299151 | -0,832117831 | 0,281912907 | 6,41E-05 | 0,000346582 | Down |
| Hbegf | 1402,236803 | 2,648822659 | 0,154553035 | 4,13E-67 | 1,26E-63 | Up |
| Eif4ebp3 | 6,552997067 | -0,864466919 | 0,823642582 | 0,00388058 | 0,013691491 | Down |
| Cd14 | 159,6812712 | 0,690235007 | 0,250161586 | 0,000276844 | 0,001292034 | Up |
| Tmco6 | 2528,610177 | -0,86113439 | 0,130280477 | 5,81E-13 | 1,31E-11 | Down |
| Wdr55 | 8507,810935 | 0,815103212 | 0,146654391 | 6,26E-10 | 8,57E-09 | Up |
| Spry4 | 227,1957364 | 5,32546848 | 0,338313076 | 7,07E-58 | 9,93E-55 | Up |
| Lars | 36330,13331 | 0,598194253 | 0,082947584 | 8,24E-14 | 2,13E-12 | Up |
| Eif3j2 | 2441,93661 | 0,60069228 | 0,081305308 | 2,20E-14 | 6,19E-13 | Up |
| Trim36 | 974,7020316 | 0,584448454 | 0,11178236 | 2,48E-08 | 2,62E-07 | Up |
| Srfbp1 | 5244,07598 | 0,695295061 | 0,089795617 | 6,93E-16 | 2,42E-14 | Up |
| Ndst1 | 8177,205955 | 0,726917478 | 0,093674926 | 4,52E-16 | 1,64E-14 | Up |
| Cd74 | 1766,084264 | 3,349516574 | 0,445991202 | 6,38E-15 | 1,97E-13 | Up |
| Camk2a | 28,86836888 | 0,911344029 | 0,396613925 | 0,000291527 | 0,00135571 | Up |
| Csf1r | 781,9611532 | 1,248767574 | 0,375294256 | 3,96E-06 | 2,76E-05 | Up |
| Pcyox1l | 10082,03128 | 0,662298194 | 0,074100216 | 3,87E-20 | 2,36E-18 | Up |
| Grpel2 | 4103,900192 | 0,604102737 | 0,060015861 | 1,20E-24 | 1,36E-22 | Up |
| Afap1l1 | 2169,408454 | 1,082375955 | 0,139711126 | 4,72E-17 | 1,93E-15 | Up |
| Nedd4l | 11741,17331 | -0,865898011 | 0,178225162 | 1,87E-08 | 2,01E-07 | Down |
| Alpk2 | 20,42225976 | 0,88186487 | 0,453223366 | 0,000773691 | 0,003262328 | Up |
| Tubb6 | 18989,66917 | 1,085830797 | 0,074587726 | 2,35E-50 | 2,26E-47 | Up |
| Mc5r | 1407,292664 | -3,399219233 | 0,30491186 | 1,20E-29 | 2,17E-27 | Down |
| Smad7 | 2163,350201 | -0,628659964 | 0,144854154 | 1,40E-06 | 1,07E-05 | Down |
| St8sia5 | 12,72983078 | 0,740188972 | 0,60355155 | 0,004995093 | 0,017108099 | Up |
| A330094K24Rik | 12,80696012 | -2,021690099 | 0,768786668 | 1,68E-05 | 0,000102992 | Down |
| Slc14a1 | 40,32534834 | 0,792101881 | 0,469135962 | 0,001959226 | 0,007466484 | Up |
| Mbp | 13931,65298 | -0,742084994 | 0,091749077 | 2,80E-17 | 1,20E-15 | Down |
| Cndp2 | 20280,98025 | 0,624893979 | 0,110710368 | 1,90E-09 | 2,41E-08 | Up |
| Cpt1a | 11003,21757 | -0,595870199 | 0,156522253 | 1,59E-05 | 9,77E-05 | Down |
| 1810055G02Rik | 2185,449402 | 0,797016014 | 0,092918489 | 2,58E-19 | 1,42E-17 | Up |
| Aldh3b1 | 123,1433698 | 1,494739272 | 0,385245731 | 3,38E-07 | 2,90E-06 | Up |
| Unc93b1 | 4614,648566 | 0,729838988 | 0,151948166 | 7,27E-08 | 7,06E-07 | Up |
| Acy3 | 548,8331198 | -0,662891456 | 0,152887364 | 1,11E-06 | 8,60E-06 | Down |
| Gstp3 | 31,36297781 | 1,510589521 | 0,699490924 | 8,66E-05 | 0,000452242 | Up |
| Coro1b | 28071,18604 | -0,702372117 | 0,077875341 | 1,30E-20 | 8,53E-19 | Down |
| Tbc1d10c | 2904,422825 | -0,805758239 | 0,142535713 | 3,92E-10 | 5,58E-09 | Down |
| Pold4 | 1546,038333 | -0,826878932 | 0,19708975 | 5,78E-07 | 4,79E-06 | Down |
| Pcx | 5514,382933 | -0,689984345 | 0,14514002 | 1,29E-07 | 1,19E-06 | Down |
| Slc29a2 | 2462,474498 | 1,391537914 | 0,173640735 | 9,07E-18 | 4,12E-16 | Up |

| | | | | | | |
|---|---|---|---|---|---|---|
| Eif1ad | 21104,54562 | 0,77423799 | 0,088082141 | 5,03E-20 | 3,05E-18 | Up |
| Tsga10ip | 1120,594777 | -0,854719041 | 0,137624755 | 8,54E-12 | 1,58E-10 | Down |
| Gm31166 | 139,3756314 | -0,587671197 | 0,348288485 | 0,006148872 | 0,020562298 | Down |
| Sssca1 | 3459,198731 | 0,685143087 | 0,109417853 | 2,83E-11 | 4,79E-10 | Up |
| Capn1 | 10560,33282 | -0,747094762 | 0,100238464 | 3,97E-15 | 1,26E-13 | Down |
| Gm10814 | 5,612445307 | -0,886498196 | 0,865010811 | 0,003743381 | 0,013289584 | Down |
| Tm7sf2 | 449,7034108 | -0,896867444 | 0,225223458 | 9,37E-07 | 7,41E-06 | Down |
| Atg2a | 6409,884903 | -0,697876508 | 0,080367599 | 2,73E-19 | 1,50E-17 | Down |
| Pygm | 82,27583861 | -1,357049769 | 0,519196808 | 3,14E-05 | 0,000181388 | Down |
| Nudt22 | 527,1842647 | -0,592407141 | 0,107981387 | 5,80E-09 | 6,81E-08 | Down |
| Rcor2 | 6595,707282 | 0,71820352 | 0,120302643 | 1,30E-10 | 1,98E-09 | Up |
| Wdr74 | 187,4925613 | 0,826823299 | 0,182000345 | 1,17E-07 | 1,10E-06 | Up |
| Rom1 | 416,3077795 | -0,807566131 | 0,118860161 | 2,64E-13 | 6,31E-12 | Down |
| Ahnak | 13,95710064 | 0,846409747 | 0,609131382 | 0,00255755 | 0,009469989 | Up |
| Gm6445 | 648,3199505 | 0,993233089 | 0,233455385 | 1,76E-07 | 1,58E-06 | Up |
| Fads3 | 83,08442898 | -0,872822111 | 0,311044843 | 8,17E-05 | 0,000429946 | Down |
| Cyb561a3 | 2272,109887 | -0,641835692 | 0,102847281 | 4,58E-11 | 7,52E-10 | Down |
| Pga5 | 28,72215418 | -1,308672742 | 0,713146525 | 0,000275377 | 0,001285845 | Down |
| Cd5 | 7,198152361 | -0,997867903 | 0,801673284 | 0,001964071 | 0,007480263 | Down |
| Slc15a3 | 17,59272379 | 5,243285566 | 1,626776005 | 1,01E-06 | 7,93E-06 | Up |
| Ccdc86 | 8287,309118 | 0,989105974 | 0,135098733 | 1,56E-15 | 5,18E-14 | Up |
| Ms4a2 | 7,863810689 | 4,475898911 | 1,513218481 | 3,75E-06 | 2,63E-05 | Up |
| Ms4a3 | 1878,313563 | 3,295744724 | 0,44497387 | 1,32E-14 | 3,84E-13 | Up |
| Stx3 | 660,5510656 | 0,73653012 | 0,088749868 | 5,14E-18 | 2,41E-16 | Up |
| Pfpl | 22,55191131 | 0,847144079 | 0,56828639 | 0,002166457 | 0,008186188 | Up |
| Mpeg1 | 526,0471994 | 0,689459019 | 0,239864532 | 0,00019897 | 0,000961013 | Up |
| Glyat | 7,185423152 | 2,113632364 | 1,326372081 | 0,000217386 | 0,001040606 | Up |
| Psat1 | 15548,68537 | 1,101599179 | 0,162407918 | 6,10E-14 | 1,60E-12 | Up |
| Gna14 | 108,4615706 | -2,754700488 | 0,391014542 | 5,55E-14 | 1,47E-12 | Down |
| Trpm6 | 13,08727511 | 2,776044083 | 0,79308681 | 2,00E-06 | 1,47E-05 | Up |
| Cbwd1 | 1816,919383 | 0,643176818 | 0,094170726 | 9,10E-13 | 1,99E-11 | Up |
| Kank1 | 118,0666616 | 3,727263791 | 0,32018894 | 3,44E-32 | 8,26E-30 | Up |
| Pum3 | 17485,55816 | 0,581120974 | 0,054971271 | 7,18E-27 | 9,78E-25 | Up |
| Slc1a1 | 77,54256669 | 4,279193533 | 0,837129 | 9,11E-09 | 1,03E-07 | Up |
| 4430402I18Rik | 89,91632962 | -1,167712859 | 0,239256338 | 5,58E-09 | 6,58E-08 | Down |
| Cd274 | 1913,758696 | 0,75874227 | 0,10720394 | 5,69E-14 | 1,50E-12 | Up |
| Pdcd1lg2 | 271,738588 | 1,453516076 | 0,289888036 | 2,37E-09 | 2,93E-08 | Up |
| Pten | 24135,26811 | -0,714782554 | 0,125136229 | 6,24E-10 | 8,55E-09 | Down |
| Ankrd22 | 50,41664981 | 2,219220537 | 0,845628389 | 1,74E-05 | 0,000105925 | Up |
| Stambpl1 | 4519,680499 | 0,737499382 | 0,06711466 | 2,15E-29 | 3,77E-27 | Up |
| Ifit1bl1 | 24,88496646 | -0,761707095 | 0,387829202 | 0,001321911 | 0,005267454 | Down |
| Ifit1 | 34,68047528 | 0,588703072 | 0,360525698 | 0,006663564 | 0,022101344 | Up |
| Htr7 | 2,847157638 | 1,410141744 | 1,639861921 | 0,001265862 | 0,005070665 | Up |
| Rpp30 | 3757,220907 | 0,641324223 | 0,122685388 | 1,69E-08 | 1,84E-07 | Up |
| Mar 05 | 7060,303215 | 0,58149397 | 0,068619801 | 4,06E-18 | 1,92E-16 | Up |
| Noc3l | 5446,023547 | 0,686567567 | 0,109264099 | 2,43E-11 | 4,17E-10 | Up |
| Pdlim1 | 3,632629155 | 0,615423576 | 0,800594517 | 0,012064534 | 0,036634937 | Up |
| Frat2 | 3581,670133 | -1,068354493 | 0,151677502 | 9,85E-15 | 2,96E-13 | Down |
| Rrp12 | 16668,34226 | 0,85482877 | 0,111952913 | 3,47E-16 | 1,28E-14 | Up |
| Exosc1 | 6112,409885 | 0,698507125 | 0,067171041 | 1,81E-26 | 2,38E-24 | Up |
| Golga7b | 11,0846472 | 0,637677586 | 0,494165644 | 0,00778042 | 0,025179165 | Up |

| | | | | | | |
|---|---|---|---|---|---|---|
| Cpn1 | 84,37942997 | -1,389578736 | 0,21660281 | 8,78E-13 | 1,92E-11 | Down |
| Sema4g | 21,70962321 | 0,606102427 | 0,383288118 | 0,00646548 | 0,021495102 | Up |
| Kazald1 | 80,18685946 | -0,697741196 | 0,281960524 | 0,00057222 | 0,002490441 | Down |
| Pprc1 | 26554,08726 | 0,636503583 | 0,095132738 | 2,49E-12 | 5,02E-11 | Up |
| Nolc1 | 64907,13545 | 0,848384457 | 0,130764027 | 1,47E-12 | 3,09E-11 | Up |
| Pdcd11 | 23467,26381 | 0,716276868 | 0,125920401 | 7,07E-10 | 9,54E-09 | Up |
| Obfc1 | 1182,931479 | -0,831104714 | 0,102969325 | 1,33E-17 | 5,89E-16 | Down |
| Gm19557 | 53,97645203 | -1,980699285 | 0,366408461 | 1,91E-10 | 2,84E-09 | Down |
| Gsto1 | 18625,6743 | 0,597301285 | 0,105595623 | 2,14E-09 | 2,68E-08 | Up |
| Rbm20 | 11,33176987 | -0,599679247 | 0,550624326 | 0,011415565 | 0,034990106 | Down |
| Pdcd4 | 1960,934207 | -1,631655249 | 0,169657263 | 4,45E-24 | 4,56E-22 | Down |
| Ablim1 | 17995,92626 | 0,801674638 | 0,182419433 | 2,84E-07 | 2,47E-06 | Up |
| Slc18a2 | 13240,57545 | -0,76198767 | 0,121026998 | 1,13E-11 | 2,06E-10 | Down |
| Nanos1 | 185,1394648 | -0,850105752 | 0,331091967 | 0,000186008 | 0,000906074 | Down |
| Csf2ra | 799,8728441 | 0,817472505 | 0,278037015 | 7,23E-05 | 0,000385072 | Up |
| 2010204K13Rik | 763,3757843 | 0,794075762 | 0,143700501 | 9,01E-10 | 1,19E-08 | Up |
| Plp2 | 16419,96745 | 0,67774493 | 0,118681115 | 8,63E-10 | 1,14E-08 | Up |
| Wdr45 | 898,4736451 | -0,700961492 | 0,136966538 | 1,88E-08 | 2,02E-07 | Down |
| Ccdc120 | 1294,409318 | -0,90460915 | 0,09366397 | 4,31E-24 | 4,47E-22 | Down |
| Pim2 | 2848,311864 | 0,834954098 | 0,115724483 | 1,01E-14 | 3,01E-13 | Up |
| Gata1 | 227,7361185 | -1,269014421 | 0,665734595 | 0,000251239 | 0,001186484 | Down |
| Cfp | 7659,197069 | 2,393540649 | 0,279636146 | 9,62E-20 | 5,56E-18 | Up |
| Gm4764 | 15,87551447 | 1,600048905 | 0,645983979 | 3,23E-05 | 0,000185832 | Up |
| Sep 06 | 14428,92337 | -0,703575616 | 0,104489323 | 1,07E-12 | 2,30E-11 | Down |
| C1galt1c1 | 1795,553176 | 0,758026347 | 0,10215631 | 4,58E-15 | 1,43E-13 | Up |
| Ap3s1-ps1 | 33,2590392 | 0,604850877 | 0,337531456 | 0,00460897 | 0,0159411 | Up |
| Sash3 | 19663,7804 | -0,847021291 | 0,127922684 | 6,06E-13 | 1,36E-11 | Down |
| Plac1 | 7,696839202 | 2,62507871 | 0,911771619 | 9,89E-06 | 6,40E-05 | Up |
| Rbmx | 8125,585353 | 0,620322365 | 0,138613466 | 8,08E-07 | 6,48E-06 | Up |
| Gm364 | 50,56655528 | 1,17234204 | 0,404193858 | 2,01E-05 | 0,000121259 | Up |
| Slc6a8 | 15,69526122 | 0,638782689 | 0,457356542 | 0,0068471 | 0,022619706 | Up |
| Plxnb3 | 28,31240364 | -0,883017643 | 0,468036533 | 0,00087564 | 0,003650898 | Down |
| L1cam | 1191,237261 | 0,817683321 | 0,247469042 | 2,13E-05 | 0,000127542 | Up |
| Atp6ap1 | 13335,74287 | 0,629244383 | 0,076937189 | 3,56E-17 | 1,48E-15 | Up |
| Gab3 | 199,6674587 | -0,740586565 | 0,25286192 | 0,000120164 | 0,000607711 | Down |
| Dkc1 | 25427,24761 | 0,628824638 | 0,067803416 | 2,28E-21 | 1,63E-19 | Up |
| F8 | 128,8397527 | -0,651583766 | 0,26115585 | 0,000731603 | 0,003103492 | Down |
| 4933407K13Rik | 225,7767179 | -1,669259832 | 0,385710653 | 4,40E-08 | 4,44E-07 | Down |
| Gm7153 | 20,38094683 | -1,489028846 | 0,560653423 | 2,21E-05 | 0,000131849 | Down |
| Pcyt1b | 237,5326495 | -1,53674149 | 0,260642765 | 1,88E-11 | 3,29E-10 | Down |
| Zc4h2 | 578,4890475 | -0,663034168 | 0,124101428 | 7,71E-09 | 8,86E-08 | Down |
| Eda2r | 159,2036293 | 1,551966397 | 0,23825475 | 4,13E-13 | 9,48E-12 | Up |
| Foxo4 | 2174,821687 | -1,019549805 | 0,126402712 | 3,97E-18 | 1,89E-16 | Down |
| Il2rg | 4447,634033 | 0,662446888 | 0,098056729 | 1,30E-12 | 2,76E-11 | Up |
| Nlgn3 | 6,930095257 | 0,769032256 | 0,657657981 | 0,004720734 | 0,016281407 | Up |
| Zmym3 | 6712,818186 | -0,683390703 | 0,097065006 | 1,49E-13 | 3,69E-12 | Down |
| Nhsl2 | 4186,278228 | 0,820952435 | 0,150986076 | 1,18E-09 | 1,54E-08 | Up |
| Gm9115 | 48,01441101 | 0,953655829 | 0,340825637 | 5,78E-05 | 0,000315586 | Up |
| C77370 | 8,093194279 | -0,786436965 | 0,723455209 | 0,00484915 | 0,016658234 | Down |
| Tlr13 | 26,2013956 | 2,94150812 | 0,689248306 | 2,04E-07 | 1,81E-06 | Up |
| Fam46d | 48,93126473 | 0,675206512 | 0,326351979 | 0,001724962 | 0,006673995 | Up |

| | | | | | | |
|---|---|---|---|---|---|---|
| Nap1l3 | 2142,146348 | 0,739330264 | 0,15912099 | 1,44E-07 | 1,32E-06 | Up |
| Nox1 | 50,65757819 | 1,149936417 | 0,470375092 | 8,39E-05 | 0,000440211 | Up |
| Timm8a1 | 2632,332527 | 0,934827155 | 0,125034351 | 6,37E-16 | 2,25E-14 | Up |
| Armcx1 | 47,24106716 | -1,465033985 | 0,495115184 | 9,05E-06 | 5,88E-05 | Down |
| Zmat1 | 166,7915089 | -0,600850056 | 0,196870715 | 0,000213037 | 0,001023273 | Down |
| Prps1 | 13576,95867 | 0,605789504 | 0,071634229 | 4,04E-18 | 1,92E-16 | Up |
| Tsc22d3 | 849,7772306 | -0,650811736 | 0,204913883 | 0,000102046 | 0,00052421 | Down |
| Iqsec2 | 1484,928006 | -0,716189318 | 0,084272571 | 1,15E-18 | 5,79E-17 | Down |
| Kantr | 79,91398355 | -0,663498882 | 0,212376368 | 0,000110294 | 0,000561997 | Down |
| Mbtps2 | 3218,912179 | 0,623098399 | 0,071096688 | 2,47E-19 | 1,37E-17 | Up |
| Phka2 | 5427,570464 | -0,690829657 | 0,073880741 | 6,67E-22 | 5,10E-20 | Down |
| Mir3473a | 688,7237014 | 1,034028087 | 0,155242843 | 1,57E-13 | 3,87E-12 | Up |
| Siah1b | 1638,159652 | 0,586469132 | 0,092243035 | 3,17E-11 | 5,33E-10 | Up |
| Piga | 3237,514279 | 0,676497049 | 0,092205619 | 1,82E-14 | 5,19E-13 | Up |
| Fancb | 1400,655085 | 0,660611052 | 0,088940329 | 1,05E-14 | 3,13E-13 | Up |
| Gpm6b | 624,9046539 | 0,604462817 | 0,205583947 | 0,000292165 | 0,001357986 | Up |
| Tceanc | 692,770873 | -0,641041342 | 0,081486708 | 4,10E-16 | 1,50E-14 | Down |
| Arhgap6 | 8355,513282 | -1,435695728 | 0,211255017 | 7,21E-14 | 1,88E-12 | Down |
| Amelx | 3,170263177 | -0,60698821 | 1,027552213 | 0,010929124 | 0,033668635 | Down |

| **CM-IL3-12-vs-CM-IL3-72** | **baseMean** | **log2FoldChange** | **lfcSE** | **pvalue** | **padj** | **expression** |
|---|---|---|---|---|---|---|
| Gm5526 | 377,8448003 | 1,127826052 | 0,357585379 | 2,99E-06 | 0,000232324 | Up |
| Aox1 | 412,1174204 | 0,649762895 | 0,166063519 | 6,76E-07 | 7,33E-05 | Up |
| Idh1 | 5830,96915 | -0,601758347 | 0,222711748 | 2,08E-05 | 0,001003098 | Down |
| Gm15433 | 27,06492675 | -3,630763034 | 0,669198129 | 8,00E-10 | 2,96E-07 | Down |
| Sned1 | 162,9354004 | -0,920291121 | 0,242098907 | 1,04E-06 | 0,000103393 | Down |
| Gm15850 | 27,76454983 | -1,774109989 | 0,562025068 | 1,61E-06 | 0,000146275 | Down |
| Gm37065 | 26,68573509 | -0,892845491 | 1,264780537 | 0,000201116 | 0,005561931 | Down |
| Slamf9 | 17,06430912 | -4,012771465 | 0,84732927 | 6,21E-09 | 1,60E-06 | Down |
| Enah | 45,52905557 | 0,654136403 | 0,764718793 | 0,000503215 | 0,010563093 | Up |
| Mrc1 | 96,22464949 | -0,678608424 | 0,470558 | 0,000249374 | 0,006510376 | Down |
| Stkld1 | 148,4656588 | -0,606197568 | 0,365592397 | 0,000231027 | 0,006209138 | Down |
| Slc2a6 | 1861,671792 | -0,726767515 | 0,311755998 | 4,54E-05 | 0,001796362 | Down |
| Lcn2 | 1442,891879 | -1,478563485 | 1,004104343 | 4,98E-05 | 0,001884053 | Down |
| Angptl2 | 299,2275307 | 2,518651867 | 0,236605641 | 2,76E-27 | 1,89E-23 | Up |
| Traf1 | 456,240135 | -0,631252199 | 0,542086207 | 0,000417359 | 0,009268616 | Down |
| Ube2l6 | 580,9973168 | -0,884542394 | 0,211119777 | 3,46E-07 | 4,22E-05 | Down |
| Rapsn | 937,8298717 | -0,671470609 | 0,13887132 | 2,49E-08 | 5,16E-06 | Down |
| Cd82 | 3036,908916 | -1,409703295 | 0,213423901 | 5,72E-14 | 8,69E-11 | Down |
| Ckmt1 | 1388,774942 | -0,586981215 | 0,187502765 | 6,03E-06 | 0,000408505 | Down |
| Hck | 112,8469967 | -1,000130253 | 0,242944909 | 3,06E-07 | 3,84E-05 | Down |
| Wfdc12 | 53,19585331 | 0,711034547 | 0,47047027 | 0,000201469 | 0,005561931 | Up |
| Svs4 | 22,97847939 | 0,81320711 | 0,889580844 | 0,000306178 | 0,007519787 | Up |
| Cp | 259,6809462 | -1,568515524 | 1,05758199 | 4,42E-05 | 0,001764325 | Down |
| Wwtr1 | 82,48549328 | -2,792765354 | 0,402793356 | 3,56E-13 | 4,42E-10 | Down |
| Tlr2 | 2738,950673 | -0,622830466 | 0,141922575 | 1,07E-07 | 1,81E-05 | Down |
| I830077J02Rik | 350,8043149 | -0,610281951 | 0,317196026 | 0,000135114 | 0,004162969 | Down |

| | | | | | | |
|---|---|---|---|---|---|---|
| Sec24d | 17207,24135 | -0,6043849 | 0,276239569 | 7,57E-05 | 0,002641935 | Down |
| Bach2os | 16,43915049 | -1,316327506 | 1,148267528 | 9,22E-05 | 0,003047806 | Down |
| Glipr2 | 6207,664248 | -0,923478928 | 0,235794519 | 7,40E-07 | 7,85E-05 | Down |
| Tlr4 | 3138,593152 | -0,637253987 | 0,214834507 | 1,07E-05 | 0,00061198 | Down |
| Ror1 | 227,5447606 | -1,017549222 | 0,185431413 | 6,77E-10 | 2,57E-07 | Down |
| Echdc2 | 151,1693368 | -0,734919074 | 0,224376657 | 5,19E-06 | 0,000365674 | Down |
| Slc6a9 | 1220,144269 | -0,831220058 | 0,286255315 | 1,13E-05 | 0,000631832 | Down |
| Spocd1 | 103,4212202 | -0,959542251 | 0,491600106 | 4,85E-05 | 0,001858863 | Down |
| Sytl1 | 1608,937679 | -0,581601699 | 0,315613586 | 0,000172219 | 0,004949481 | Down |
| Extl1 | 477,9996045 | -1,932053934 | 0,620522058 | 1,89E-06 | 0,000168077 | Down |
| Man1c1 | 2246,77068 | -1,264403261 | 0,211589159 | 6,97E-12 | 5,61E-09 | Down |
| Clic4 | 37244,65662 | -0,744053946 | 0,168563904 | 1,68E-07 | 2,47E-05 | Down |
| Pla2g2e | 33,9934475 | -0,702879183 | 0,867136064 | 0,00044286 | 0,009677842 | Down |
| Iffo2 | 56,46667505 | -1,449963303 | 0,36432788 | 1,15E-07 | 1,87E-05 | Down |
| Casz1 | 137,4757495 | -0,591609354 | 0,362684691 | 0,000254832 | 0,006619865 | Down |
| Chd5 | 261,9581367 | -0,678159795 | 0,62847996 | 0,000382171 | 0,008742639 | Down |
| Kcnh2 | 15,80891239 | -0,671811473 | 0,845354103 | 0,000490876 | 0,010403752 | Down |
| Nos3 | 60,47437923 | -1,855988438 | 0,275485643 | 7,35E-14 | 1,01E-10 | Down |
| Insig1 | 4185,584737 | -0,81306744 | 0,248282494 | 4,98E-06 | 0,000356824 | Down |
| Bst1 | 33,67837635 | -1,104689427 | 1,205978097 | 0,000151991 | 0,004539824 | Down |
| Oasl2 | 42,41555797 | -2,096830667 | 0,469861007 | 4,40E-08 | 8,15E-06 | Down |
| Tesc | 31,63473976 | -1,184526249 | 0,658303244 | 4,04E-05 | 0,001660105 | Down |
| Oas2 | 153,550936 | -3,589918002 | 0,331620266 | 7,89E-29 | 1,08E-24 | Down |
| Oas3 | 341,8636315 | -2,621957975 | 0,448726944 | 2,55E-10 | 1,28E-07 | Down |
| Oas1b | 92,37012937 | -1,07141475 | 0,259346389 | 2,16E-07 | 3,02E-05 | Down |
| Oas1a | 3224,377083 | -0,809420863 | 0,110605951 | 1,84E-14 | 3,60E-11 | Down |
| Hvcn1 | 2889,337632 | -0,68344398 | 0,187745219 | 1,78E-06 | 0,00015875 | Down |
| Morn3 | 17,93221644 | -0,648813498 | 0,562639179 | 0,000394255 | 0,008940168 | Down |
| Gm43661 | 410,5188138 | 0,749203321 | 0,1301505 | 4,22E-10 | 1,80E-07 | Up |
| Zbed5 | 301,228206 | 0,670493748 | 0,216796397 | 7,96E-06 | 0,000502443 | Up |
| Myl10 | 156,465505 | 0,985821699 | 0,639967883 | 8,92E-05 | 0,002975997 | Up |
| Card11 | 8704,093875 | 0,738016514 | 0,139154924 | 4,09E-09 | 1,14E-06 | Up |
| Asns | 30785,17086 | -0,788914852 | 0,158755159 | 1,98E-08 | 4,37E-06 | Down |
| Tnip3 | 175,1734794 | -0,978123797 | 0,337629472 | 8,00E-06 | 0,000502443 | Down |
| Gadd45a | 450,8678506 | -1,178483546 | 0,375809834 | 2,74E-06 | 0,000216352 | Down |
| Igkv14-111 | 84,3055212 | 0,887977926 | 0,382684386 | 3,11E-05 | 0,001383655 | Up |
| Gm25436 | 45,25769782 | 0,642710443 | 0,551531064 | 0,000398619 | 0,008998533 | Up |
| Rab11fip5 | 155,2319165 | -1,743102885 | 0,470061669 | 3,32E-07 | 4,13E-05 | Down |
| Gm44198 | 184,4808107 | -0,64276539 | 0,170310574 | 1,03E-06 | 0,000102561 | Down |
| Lrig1 | 86,48851571 | -0,966992012 | 0,225846423 | 1,97E-07 | 2,81E-05 | Down |
| Rasgef1a | 67,70702736 | -0,588801394 | 0,400352298 | 0,000338255 | 0,008118126 | Down |
| Usp18 | 164,2414255 | -0,745207982 | 0,252064766 | 1,13E-05 | 0,000631832 | Down |
| Tmem150b | 60,43056541 | -0,594897164 | 0,408261459 | 0,00033719 | 0,008106789 | Down |
| Sbk2 | 129,9005395 | -0,767052707 | 0,28460356 | 1,97E-05 | 0,000969987 | Down |
| Relb | 527,0632685 | -0,76364054 | 0,255997349 | 1,04E-05 | 0,000605474 | Down |
| Cd177 | 3314,386718 | -0,964183789 | 0,203326991 | 3,36E-08 | 6,86E-06 | Down |
| Pou2f2 | 282,5412565 | 0,624745333 | 0,161619854 | 7,00E-07 | 7,54E-05 | Up |
| Mag | 736,0866341 | -0,662660067 | 0,156030157 | 2,29E-07 | 3,14E-05 | Down |
| Rhpn2 | 178,5529272 | -1,778370268 | 1,031279834 | 2,53E-05 | 0,001165463 | Down |
| Nkg7 | 1081,063626 | -0,642353193 | 0,180208206 | 1,97E-06 | 0,000171707 | Down |
| Kcnc3 | 605,3869334 | -0,623678242 | 0,338176594 | 0,000153718 | 0,004571427 | Down |

| | | | | | | |
|---|---|---|---|---|---|---|
| Atf5 | 8887,322386 | -0,769374294 | 0,240007774 | 6,13E-06 | 0,000413167 | Down |
| 1810008I18Rik | 31,5009811 | -3,09981574 | 0,541605823 | 4,44E-10 | 1,84E-07 | Down |
| Selenos | 9797,488967 | -0,588171563 | 0,197585658 | 9,50E-06 | 0,000559649 | Down |
| Isg20 | 612,3464215 | -0,598019931 | 0,583191431 | 0,000541923 | 0,011034768 | Down |
| Il16 | 1681,260088 | -1,085405935 | 0,257855937 | 1,54E-07 | 2,31E-05 | Down |
| Stard10 | 87,27593361 | -0,674115213 | 0,374765673 | 0,000144408 | 0,004390003 | Down |
| Folr1 | 1260,518609 | -0,786519112 | 0,303422041 | 2,37E-05 | 0,001108366 | Down |
| Usp17ld | 816,2852816 | -1,344306907 | 0,195343868 | 1,06E-14 | 2,42E-11 | Down |
| Usp17lc | 25,65547678 | -0,90106612 | 0,710461645 | 0,000162104 | 0,004739034 | Down |
| Lyve1 | 54,82085729 | -0,697760149 | 0,506680921 | 0,000253349 | 0,006601545 | Down |
| Lat | 308,9352463 | -0,637485446 | 0,28241766 | 6,18E-05 | 0,002224008 | Down |
| Cox6a2 | 153,6528533 | -1,94336269 | 0,927884505 | 1,27E-05 | 0,000691602 | Down |
| Lrrc27 | 225,731834 | -0,712013064 | 0,197554202 | 2,09E-06 | 0,000178693 | Down |
| Ifitm3 | 93618,87326 | -0,919764773 | 0,324314084 | 1,06E-05 | 0,000608992 | Down |
| Irf7 | 224,653155 | -1,666772725 | 0,265076339 | 5,67E-13 | 5,97E-10 | Down |
| Cttn | 155,5479725 | 1,223056241 | 0,434446415 | 5,21E-06 | 0,000365802 | Up |
| Gm7676 | 30,2888419 | -0,650984881 | 0,545176741 | 0,000374599 | 0,008611901 | Down |
| Gpt2 | 11447,41538 | -1,131376192 | 0,189132245 | 1,71E-11 | 1,12E-08 | Down |
| Rpl10-ps2 | 16,17777613 | 1,151065945 | 0,756217372 | 6,85E-05 | 0,002416846 | Up |
| Gm16116 | 19,46580365 | -1,187956304 | 0,659449799 | 4,00E-05 | 0,001659581 | Down |
| Zcchc14 | 64,29279821 | -1,080093705 | 0,347304502 | 3,78E-06 | 0,000290744 | Down |
| Gm26812 | 25,95583854 | -1,052451795 | 0,643728396 | 6,69E-05 | 0,002378574 | Down |
| Slc35f3 | 594,0832717 | -0,924939786 | 0,182815803 | 9,78E-09 | 2,35E-06 | Down |
| Col5a3 | 19,69508524 | -0,711267429 | 0,736877012 | 0,000384813 | 0,008759139 | Down |
| Lman1l | 22,96843767 | -0,8193283 | 0,534912385 | 0,000137653 | 0,004231666 | Down |
| Cd276 | 489,4175603 | -1,162664679 | 0,261057541 | 4,04E-08 | 7,85E-06 | Down |
| Aqp9 | 2236,450141 | -1,919138036 | 0,427360148 | 2,49E-08 | 5,16E-06 | Down |
| Bcl2a1b | 1443,482025 | -1,073826205 | 0,249785495 | 1,16E-07 | 1,87E-05 | Down |
| Gm26767 | 319,5603008 | -0,707451398 | 1,120843966 | 0,000349248 | 0,008293998 | Down |
| Entpd3 | 586,3407412 | -0,613574047 | 0,187627264 | 4,32E-06 | 0,000324949 | Down |
| Map7 | 40,43709115 | -0,784930275 | 0,482375336 | 0,000132998 | 0,004124704 | Down |
| Ddit4 | 6755,504962 | -0,672050259 | 0,274116926 | 3,81E-05 | 0,00160086 | Down |
| Hkdc1 | 599,7391645 | -0,886084015 | 0,250149836 | 2,27E-06 | 0,000189663 | Down |
| Agpat3 | 3052,747228 | -0,762139661 | 0,127225104 | 1,17E-10 | 6,65E-08 | Down |
| Slc41a2 | 107,4646155 | -0,941133971 | 0,27526569 | 2,71E-06 | 0,000216352 | Down |
| Ikbip | 1651,302565 | -0,713822862 | 0,094136346 | 2,93E-15 | 8,02E-12 | Down |
| Gm3571 | 49,55293037 | -0,85398591 | 0,385001485 | 4,09E-05 | 0,001676427 | Down |
| Il23a | 78,30081049 | -1,922265541 | 0,326405804 | 2,15E-11 | 1,34E-08 | Down |
| Gatsl3 | 234,9136018 | -1,095612884 | 0,215067007 | 3,11E-09 | 8,94E-07 | Down |
| Lif | 95,69670836 | -0,885794948 | 0,319332641 | 1,31E-05 | 0,000702903 | Down |
| Abca13 | 258,6533689 | -0,862997649 | 0,284640418 | 8,01E-06 | 0,000502443 | Down |
| Plek | 48649,19621 | 0,620046912 | 0,179252262 | 2,51E-06 | 0,000201739 | Up |
| Slc1a4 | 6577,979338 | -0,671916634 | 0,152488357 | 1,35E-07 | 2,12E-05 | Down |
| Igtp | 608,3891546 | -0,772650115 | 0,173071187 | 1,46E-07 | 2,24E-05 | Down |
| Irgm2 | 601,167991 | -0,810351845 | 0,16891861 | 4,08E-08 | 7,85E-06 | Down |
| Xaf1 | 63,53803563 | -1,875144662 | 0,31703769 | 1,47E-11 | 1,00E-08 | Down |
| Serpinf1 | 509,0275843 | -0,734592154 | 0,365969342 | 8,31E-05 | 0,002843667 | Down |
| Rhbdl3 | 31,14154638 | -0,936969123 | 0,471804072 | 4,86E-05 | 0,001858863 | Down |
| Slfn5 | 587,8125687 | -1,110171457 | 0,270541648 | 2,01E-07 | 2,84E-05 | Down |
| Scpep1 | 1322,873176 | -0,586433953 | 0,247978182 | 4,99E-05 | 0,001884053 | Down |
| Itgb3 | 28796,22896 | 0,584707499 | 0,143670759 | 2,66E-07 | 3,50E-05 | Up |

| | | | | | | |
|---|---|---|---|---|---|---|
| Pycr1 | 1511,586998 | -0,800574142 | 0,39254111 | 6,51E-05 | 0,002318844 | Down |
| Tex19,2 | 27,7198312 | 0,679581491 | 0,516263974 | 0,000292237 | 0,007308599 | Up |
| Greb1 | 5300,365898 | 2,877187724 | 0,530195729 | 2,29E-09 | 7,47E-07 | Up |
| Dnajb9 | 2451,619579 | -0,775259225 | 0,285028284 | 1,85E-05 | 0,000929094 | Down |
| Slc8a3 | 59,93000479 | -1,002219726 | 0,299510569 | 2,72E-06 | 0,000216352 | Down |
| Ifi27 | 6938,423981 | -1,742825023 | 0,232729394 | 1,72E-16 | 5,88E-13 | Down |
| Akr1c19 | 354,8247835 | -0,709145903 | 0,271209951 | 2,54E-05 | 0,001166615 | Down |
| Gpr141 | 1956,96167 | -0,945118987 | 0,722173087 | 0,000138839 | 0,004258567 | Down |
| Hist1h1c | 938,9348709 | -0,720178565 | 0,206852441 | 2,98E-06 | 0,000232324 | Down |
| 4930486L24Rik | 154,5300492 | 0,796557353 | 0,422059701 | 8,52E-05 | 0,002878395 | Up |
| Ctla2b | 974,1739352 | 1,147424315 | 0,222151461 | 1,54E-09 | 5,28E-07 | Up |
| Ctla2a | 2671,589779 | 1,727559851 | 0,181388894 | 3,91E-24 | 1,78E-20 | Up |
| Emb | 15483,94736 | -0,647875858 | 0,181574764 | 1,99E-06 | 0,000171944 | Down |
| Myoz1 | 318,4189518 | -0,829187351 | 0,153396618 | 2,51E-09 | 7,94E-07 | Down |
| Kcnma1 | 68,5027925 | -2,447489622 | 1,350742734 | 9,23E-06 | 0,000553759 | Down |
| Chdh | 23,90098117 | -2,563503635 | 0,562301696 | 6,54E-08 | 1,18E-05 | Down |
| Trac | 148,0237605 | 0,919549743 | 0,518309187 | 7,19E-05 | 0,00251695 | Up |
| Slc22a17 | 1543,873298 | -0,668583486 | 0,111809197 | 9,20E-11 | 5,47E-08 | Down |
| Pck2 | 22785,40009 | -1,090633001 | 0,213882727 | 3,14E-09 | 8,94E-07 | Down |
| Mcpt1 | 70,48400238 | 2,665350696 | 0,53175153 | 1,34E-08 | 3,12E-06 | Up |
| Gzmc | 158,6019186 | 1,985004311 | 1,468038157 | 2,43E-05 | 0,001122861 | Up |
| Gzmb | 5878,048741 | 1,909146287 | 0,383730122 | 2,68E-09 | 8,15E-07 | Up |
| Farp1 | 33,18368729 | 0,666486036 | 0,740427879 | 0,000468034 | 0,010083006 | Up |
| Pop1 | 6346,479171 | 0,616486937 | 0,118354555 | 3,10E-09 | 8,94E-07 | Up |
| Snx31 | 208,4290402 | -0,58598295 | 0,273094408 | 8,57E-05 | 0,002886398 | Down |
| Foxred2 | 326,5017059 | -0,722007077 | 0,352735231 | 8,00E-05 | 0,002749603 | Down |
| Il2rb | 2124,240064 | -0,674599305 | 0,168495177 | 5,55E-07 | 6,44E-05 | Down |
| Arfgap3 | 3891,6953 | -0,733871556 | 0,239379083 | 8,74E-06 | 0,000529281 | Down |
| Parvb | 447,153434 | -0,90609934 | 0,359586399 | 2,05E-05 | 0,000995897 | Down |
| Pou6f1 | 59,0567969 | -0,780551146 | 0,3465653 | 4,68E-05 | 0,001829446 | Down |
| Acvrl1 | 417,3961895 | 1,182520175 | 0,619413096 | 3,33E-05 | 0,001464872 | Up |
| Igfbp6 | 168,7036312 | -0,747068353 | 0,273420425 | 1,87E-05 | 0,000931155 | Down |
| Rarg | 671,1207684 | -0,798312639 | 0,197172452 | 6,00E-07 | 6,84E-05 | Down |
| Gpr84 | 315,0824204 | -0,765048879 | 0,392307974 | 8,42E-05 | 0,002867172 | Down |
| Coro7 | 8024,278426 | -0,790149457 | 0,135193444 | 2,62E-10 | 1,28E-07 | Down |
| Rimbp3 | 353,1258945 | 0,787224149 | 0,163662775 | 3,87E-08 | 7,68E-06 | Up |
| Phf10 | 12548,7236 | -0,693200086 | 0,123973827 | 8,44E-10 | 3,04E-07 | Down |
| Tnxb | 660,7082365 | -0,979417816 | 0,305248068 | 4,03E-06 | 0,000306172 | Down |
| Aif1 | 158,4295468 | -0,774068137 | 0,231821835 | 4,39E-06 | 0,000326637 | Down |
| Trem3 | 390,164595 | -1,185556535 | 0,29762995 | 2,24E-07 | 3,10E-05 | Down |
| Mpnd | 8047,801588 | -0,596006307 | 0,104938595 | 2,36E-10 | 1,24E-07 | Down |
| C3 | 3832,766568 | -1,134217671 | 0,516544539 | 2,15E-05 | 0,001023167 | Down |
| Cd74 | 1766,084264 | -1,215200199 | 0,663812475 | 3,65E-05 | 0,001562232 | Down |
| Afap1l1 | 2169,408454 | -0,67403688 | 0,143354062 | 4,36E-08 | 8,15E-06 | Down |
| Zdhhc24 | 1666,941351 | -0,678074963 | 0,094089838 | 4,09E-14 | 7,00E-11 | Down |
| Psat1 | 15548,68537 | -0,816032511 | 0,158345282 | 8,61E-09 | 2,14E-06 | Down |
| Mamdc2 | 17,24135101 | -0,838533205 | 0,830610027 | 0,000262139 | 0,006715469 | Down |
| Ifit1 | 34,68047528 | -1,371132232 | 0,551186872 | 8,51E-06 | 0,000520653 | Down |
| Scd2 | 130084,5421 | -0,791725273 | 0,217374568 | 1,97E-06 | 0,000171707 | Down |
| Shtn1 | 50,71352077 | 0,676725839 | 0,446738911 | 0,000226706 | 0,006117033 | Up |
| 3830403N18Rik | 1450,474543 | 1,11788328 | 0,224822924 | 4,93E-09 | 1,32E-06 | Up |

| | | | | | | |
|---|---|---|---|---|---|---|
| Tlr13 | 26,2013956 | -1,65502156 | 1,055825794 | 3,58E-05 | 0,001534002 | Down |
| Tsc22d3 | 849,7772306 | -0,639195503 | 0,267483196 | 4,54E-05 | 0,001796362 | Down |

| **IL3-CM-12-vs-IL3-CM-72** | | | | | | |
|---|---|---|---|---|---|---|
| | **baseMean** | **log2FoldChange** | **lfcSE** | **pvalue** | **padj** | **expression** |
| Tmem14a | 317,0076962 | -0,712690189 | 0,16815456 | 2,82E-07 | 4,28E-06 | Down |
| Gm33280 | 28,37326226 | 1,152482902 | 0,40640959 | 8,12E-06 | 8,04E-05 | Up |
| D430040D24Rik | 97,31340253 | -1,229369574 | 0,266422796 | 6,48E-09 | 1,49E-07 | Down |
| Fam178b | 26,82373702 | -0,724850406 | 0,513838849 | 0,001257576 | 0,006427336 | Down |
| Inpp4a | 8,562766052 | -2,011885986 | 1,019263799 | 4,82E-05 | 0,000384228 | Down |
| Aff3 | 4375,805027 | -0,658904362 | 0,229386275 | 6,29E-05 | 0,0004837 | Down |
| Map4k4 | 1065,531247 | 1,283341403 | 0,292340135 | 2,23E-08 | 4,38E-07 | Up |
| Il1r1 | 3429,923903 | 0,713925418 | 0,128573491 | 4,23E-10 | 1,38E-08 | Up |
| Fhl2 | 24,97946615 | 1,682916207 | 0,686225852 | 2,13E-05 | 0,000189134 | Up |
| Wdr75 | 13568,03793 | 0,692565086 | 0,09268184 | 1,93E-15 | 2,26E-13 | Up |
| Rftn2 | 532,1447096 | -0,700834163 | 0,123550726 | 2,58E-10 | 8,87E-09 | Down |
| Orc2 | 9910,453522 | 0,667930063 | 0,072927242 | 1,99E-21 | 8,23E-19 | Up |
| Nop58 | 36117,51578 | 0,5885416 | 0,080764937 | 2,48E-14 | 2,26E-12 | Up |
| Ica1l | 337,8059389 | 0,802801541 | 0,285278816 | 2,36E-05 | 0,000205075 | Up |
| Gm20342 | 172,0280904 | -0,748115391 | 0,319641952 | 0,00013398 | 0,000925956 | Down |
| Fastkd2 | 5356,931692 | 0,609333518 | 0,097877915 | 2,83E-11 | 1,28E-09 | Up |
| D630023F18Rik | 24,6894906 | 0,669745119 | 0,39557947 | 0,000963394 | 0,005085893 | Up |
| Cxcr2 | 77,55932955 | 0,806774628 | 0,539235307 | 0,000773019 | 0,004218323 | Up |
| Slc11a1 | 107,6252626 | 3,173936783 | 1,023413272 | 2,74E-06 | 3,05E-05 | Up |
| Stk36 | 69,79589087 | -1,313410013 | 0,273213794 | 3,71E-09 | 9,24E-08 | Down |
| Tuba4a | 73308,85271 | 0,796696387 | 0,087159363 | 2,69E-22 | 1,21E-19 | Up |
| D2hgdh | 1369,304957 | -0,748423431 | 0,091653897 | 3,30E-18 | 6,66E-16 | Down |
| Pdcd1 | 212,3635005 | -0,593358418 | 0,535931728 | 0,003671479 | 0,015769599 | Down |
| 1810006J02Rik | 1663,580782 | 0,864205212 | 0,151579455 | 2,23E-11 | 1,04E-09 | Up |
| Phlpp1 | 2041,35478 | -0,595059812 | 0,149290376 | 2,87E-06 | 3,19E-05 | Down |
| Tmem185b | 9566,073431 | 0,668321265 | 0,083478757 | 4,22E-17 | 6,70E-15 | Up |
| Epb41l5 | 3300,791017 | -0,697215264 | 0,100542178 | 8,97E-14 | 7,23E-12 | Down |
| Ddx18 | 21063,86072 | 0,671229498 | 0,084278327 | 5,67E-17 | 8,62E-15 | Up |
| Map3k19 | 32,01209998 | -1,392676163 | 0,484572839 | 7,26E-06 | 7,29E-05 | Down |
| Gm6170 | 289,1940133 | -2,117268466 | 1,243867196 | 6,91E-05 | 0,000522413 | Down |
| Btg2 | 1120,205726 | -1,266773825 | 0,159699205 | 8,15E-18 | 1,58E-15 | Down |
| Ppfia4 | 21,39133961 | -0,939890541 | 0,528620275 | 0,000268519 | 0,001700262 | Down |
| Elf3 | 18,41098577 | -1,56543465 | 0,814202212 | 7,76E-05 | 0,000576133 | Down |
| 9230116N13Rik | 101,8740312 | -0,725978749 | 0,222405581 | 9,96E-06 | 9,59E-05 | Down |
| Rgs1 | 88,76523314 | -1,441765664 | 0,34976999 | 1,09E-07 | 1,80E-06 | Down |
| Rgl1 | 163,2359952 | -1,005597915 | 0,226666084 | 1,04E-08 | 2,25E-07 | Down |
| Soat1 | 16,62526506 | 0,663044638 | 0,57448708 | 0,002478425 | 0,011432517 | Up |
| Selp | 1150,37954 | -0,774234495 | 0,212440931 | 1,59E-06 | 1,91E-05 | Down |
| Tbx19 | 183,1212648 | -1,391176339 | 0,235847571 | 2,01E-11 | 9,50E-10 | Down |
| Dusp12 | 1364,685485 | 0,632202319 | 0,113852303 | 1,19E-09 | 3,42E-08 | Up |
| Fcgr4 | 32,96658856 | 2,146859303 | 0,664284431 | 2,86E-06 | 3,19E-05 | Up |
| Fcgr3 | 445,2897942 | 3,116090839 | 0,489635606 | 3,90E-12 | 2,21E-10 | Up |
| Cfap126 | 33,52725716 | -0,926885112 | 0,396835239 | 5,99E-05 | 0,000463727 | Down |

| | | | | | | |
|---|---|---|---|---|---|---|
| Cd244 | 2297,6769 | -0,644378232 | 0,15499224 | 8,75E-07 | 1,14E-05 | Down |
| A630035G10Rik | 47,50261419 | -0,678473037 | 0,343528203 | 0,000508127 | 0,002952209 | Down |
| Igsf8 | 11644,53898 | 1,234034816 | 0,139127701 | 2,25E-21 | 9,06E-19 | Up |
| Ifi204 | 148,3465776 | -0,606190227 | 0,22234411 | 0,000153003 | 0,001043661 | Down |
| Mndal | 929,3119738 | -0,627374488 | 0,193852622 | 2,89E-05 | 0,000245124 | Down |
| Ifi211 | 27,8513269 | -0,712729459 | 0,399069197 | 0,000626671 | 0,003514901 | Down |
| Ifi203 | 624,3994017 | -0,857389472 | 0,239227886 | 1,01E-06 | 1,29E-05 | Down |
| Spta1 | 352,3317916 | -3,391140665 | 0,271470905 | 3,55E-37 | 1,12E-33 | Down |
| Olfr417 | 8,060975743 | -1,207142091 | 0,928235828 | 0,000436048 | 0,002581661 | Down |
| Olfr248 | 16,25454946 | -2,217792914 | 0,566409862 | 4,12E-07 | 5,94E-06 | Down |
| Olfr414 | 16,69962429 | -1,930663291 | 0,701931685 | 9,66E-06 | 9,33E-05 | Down |
| B230369F24Rik | 135,9585883 | -0,670086693 | 0,206936389 | 1,88E-05 | 0,000168736 | Down |
| H3f3aos | 23,41425315 | -0,833749399 | 0,424487882 | 0,000234755 | 0,001507111 | Down |
| Enah | 45,52905557 | 1,076127562 | 0,810477587 | 0,000533511 | 0,003070178 | Up |
| Degs1 | 9673,181637 | 0,665276571 | 0,11459979 | 1,93E-10 | 6,93E-09 | Up |
| Gm37800 | 94,49497482 | 1,608033816 | 0,319847735 | 3,79E-09 | 9,43E-08 | Up |
| Rrp15 | 11326,24982 | 0,601675958 | 0,090646631 | 2,11E-12 | 1,28E-10 | Up |
| Hsd11b1 | 61,53172636 | 0,864011166 | 0,53249098 | 0,000484591 | 0,002828013 | Up |
| Rpp38 | 1625,662356 | 0,594814779 | 0,097790382 | 7,80E-11 | 3,11E-09 | Up |
| Echdc3 | 48,56900599 | -0,839554164 | 0,300221269 | 2,01E-05 | 0,00017907 | Down |
| Gm13261 | 115,9308044 | 0,810522557 | 0,307813786 | 3,97E-05 | 0,000323909 | Up |
| Il2ra | 1536,200824 | 1,602922257 | 0,501873848 | 3,05E-06 | 3,36E-05 | Up |
| Gm13375 | 364,7283636 | -1,049590834 | 0,176066731 | 1,78E-12 | 1,10E-10 | Down |
| Il1f9 | 11,44569477 | 2,261894548 | 1,678446983 | 7,49E-05 | 0,000557878 | Up |
| Slc34a3 | 164,5404617 | -2,571392156 | 0,405041198 | 3,11E-12 | 1,79E-10 | Down |
| Clic3 | 77,12869887 | -0,646749532 | 0,320772356 | 0,00057719 | 0,003275953 | Down |
| Fam69b | 4177,57228 | 0,598810196 | 0,089728192 | 1,70E-12 | 1,05E-10 | Up |
| Spaca9 | 236,2926072 | -1,444098138 | 0,199283943 | 5,65E-15 | 5,84E-13 | Down |
| Cercam | 1490,254638 | -0,645345402 | 0,180145907 | 7,66E-06 | 7,67E-05 | Down |
| Zer1 | 3262,2588 | -0,763547842 | 0,107511648 | 9,50E-15 | 9,33E-13 | Down |
| Kyat1 | 320,4710276 | -0,654110515 | 0,141987483 | 1,12E-07 | 1,86E-06 | Down |
| Hmcn2 | 28,39534262 | 0,584336716 | 0,529450078 | 0,003864818 | 0,016451648 | Up |
| Qrfp | 115,3309365 | -0,638867251 | 0,233004394 | 0,000108564 | 0,00076994 | Down |
| Lcn2 | 1442,891879 | 1,017010592 | 0,790645677 | 0,000653492 | 0,003647115 | Up |
| St6galnac4 | 10520,07696 | 0,750701672 | 0,090331189 | 9,51E-19 | 2,17E-16 | Up |
| Fam129b | 2719,526399 | 0,635314146 | 0,180103827 | 1,03E-05 | 9,86E-05 | Up |
| Gm38299 | 22,19414092 | 0,709045314 | 0,705299828 | 0,002445964 | 0,011327894 | Up |
| Gm13496 | 34,17031884 | 0,637876505 | 0,354015843 | 0,000949322 | 0,005023416 | Up |
| Gm13502 | 42,7656329 | -1,231905902 | 0,516136596 | 3,16E-05 | 0,000265192 | Down |
| Dhrs9 | 47,75226993 | 0,895720975 | 1,128843945 | 0,001394477 | 0,007016664 | Up |
| Ttc30b | 97,98092554 | -1,151680482 | 0,224325307 | 3,31E-10 | 1,10E-08 | Down |
| Plekha3 | 4718,743872 | 0,618860171 | 0,088077312 | 1,24E-13 | 9,59E-12 | Up |
| Cerkl | 138,9130571 | -0,786984668 | 0,189785871 | 1,77E-07 | 2,82E-06 | Down |
| Ube2l6 | 580,9973168 | -0,58252692 | 0,195860692 | 9,73E-05 | 0,000698667 | Down |
| Timm10 | 2883,452495 | 0,709738923 | 0,119463553 | 4,73E-11 | 1,98E-09 | Up |
| Cd82 | 3036,908916 | 1,451073464 | 0,208087705 | 3,82E-14 | 3,39E-12 | Up |
| Nat10 | 18285,97154 | 0,61049926 | 0,094937795 | 7,59E-12 | 3,88E-10 | Up |
| Ganc | 2970,092092 | -0,636032161 | 0,1157745 | 1,59E-09 | 4,36E-08 | Down |
| Map1a | 7522,903493 | -1,121227379 | 0,257761225 | 1,65E-08 | 3,35E-07 | Down |
| Gm14002 | 33,59088708 | -1,008222836 | 0,38755174 | 2,10E-05 | 0,000186537 | Down |
| Fbn1 | 28,01245812 | 3,154430772 | 0,662350637 | 2,13E-08 | 4,22E-07 | Up |

| | | | | | | |
|---|---|---|---|---|---|---|
| Atp8b4 | 4650,524735 | 0,631757153 | 0,116933666 | 2,73E-09 | 7,00E-08 | Up |
| Ttl | 207,8065823 | -1,093627231 | 0,154344253 | 9,65E-16 | 1,16E-13 | Down |
| Polr1b | 10878,04045 | 0,765402316 | 0,121761956 | 2,38E-12 | 1,42E-10 | Up |
| Cdc25b | 10853,36342 | -0,62938776 | 0,138532055 | 1,97E-07 | 3,09E-06 | Down |
| Ap5s1 | 565,1535008 | -1,325602876 | 0,161263218 | 1,50E-18 | 3,37E-16 | Down |
| Cd93 | 362,579175 | 2,829824594 | 2,410967627 | 9,11E-06 | 8,90E-05 | Up |
| Id1 | 13342,87747 | 1,723549097 | 0,136015987 | 3,40E-38 | 1,33E-34 | Up |
| Cox4i2 | 37,51901582 | 1,239841838 | 0,373263302 | 1,51E-06 | 1,84E-05 | Up |
| Ttll9 | 22,18053213 | 0,679554378 | 0,512949888 | 0,001785152 | 0,008661764 | Up |
| Hck | 112,8469967 | 0,718606453 | 0,223051416 | 1,23E-05 | 0,000115866 | Up |
| Nol4l | 6164,336407 | -0,671243487 | 0,112857831 | 7,71E-11 | 3,09E-09 | Down |
| Gm21850 | 111,8616401 | 2,088148944 | 0,876284524 | 2,04E-05 | 0,000181644 | Up |
| Pxmp4 | 2668,970236 | -1,018511987 | 0,189864621 | 6,70E-11 | 2,73E-09 | Down |
| Ggt7 | 101,5361539 | -2,015371195 | 0,371276708 | 7,61E-10 | 2,31E-08 | Down |
| Acss2 | 1823,739378 | -0,965986124 | 0,21955386 | 1,39E-08 | 2,88E-07 | Down |
| D630003M21Rik | 639,0190827 | -1,309869788 | 0,246422686 | 2,92E-10 | 9,87E-09 | Down |
| Gm16316 | 586,0465185 | 0,704092782 | 0,117990247 | 4,38E-11 | 1,86E-09 | Up |
| Slpi | 1213,529228 | 1,04121509 | 0,524045927 | 0,000124005 | 0,000866547 | Up |
| Gm11473 | 149,3113433 | 1,773794141 | 0,244951904 | 1,14E-14 | 1,10E-12 | Up |
| Atp9a | 443,3186064 | -1,683614586 | 0,190292702 | 3,10E-20 | 9,75E-18 | Down |
| Ctsz | 32776,91395 | 0,8267942 | 0,095958606 | 1,74E-20 | 6,37E-18 | Up |
| Zfp831 | 679,8420958 | -0,932858345 | 0,21305473 | 1,79E-08 | 3,58E-07 | Down |
| Edn3 | 84,35801001 | -1,839402042 | 0,217410829 | 7,53E-19 | 1,77E-16 | Down |
| Gm14296 | 24,96905536 | 0,628505975 | 0,416523818 | 0,001714466 | 0,0083861 | Up |
| Gm14418 | 426,865861 | 0,633998606 | 0,120254525 | 5,34E-09 | 1,26E-07 | Up |
| Gm14327 | 90,33729027 | 0,769052343 | 0,321077983 | 0,000101531 | 0,000725299 | Up |
| Col9a3 | 253,833036 | -0,679308336 | 0,263633914 | 0,000118186 | 0,000830316 | Down |
| Cp | 259,6809462 | 1,019864712 | 0,830926239 | 0,00071888 | 0,003958624 | Up |
| Cetn4 | 78,19373463 | -0,600624161 | 0,204859655 | 9,24E-05 | 0,000669968 | Down |
| Bbs12 | 659,7257054 | 0,589559985 | 0,188014939 | 5,68E-05 | 0,000442965 | Up |
| 1700017G19Rik | 81,01020673 | -0,601201915 | 0,210236355 | 0,000112272 | 0,000793376 | Down |
| Gm2011 | 65,66743088 | -0,700030203 | 0,21563749 | 1,36E-05 | 0,000126885 | Down |
| Gm6209 | 5,265119697 | 1,145395922 | 1,502771965 | 0,000512923 | 0,002973856 | Up |
| 5031434O11Rik | 165,0372201 | -0,613042766 | 0,155806446 | 2,94E-06 | 3,25E-05 | Down |
| Maml3 | 4177,416488 | -0,669222693 | 0,115283573 | 1,84E-10 | 6,65E-09 | Down |
| P2ry14 | 6297,454053 | 1,240994857 | 0,207958139 | 4,89E-12 | 2,67E-10 | Up |
| P2ry13 | 20,34428268 | 1,868830256 | 0,794166783 | 2,44E-05 | 0,00021086 | Up |
| Nmd3 | 8492,234769 | 0,594007143 | 0,079699378 | 6,90E-15 | 6,95E-13 | Up |
| Ppid | 11918,65474 | 0,882581803 | 0,149046849 | 4,70E-12 | 2,58E-10 | Up |
| Gpatch4 | 8405,066357 | 0,66613617 | 0,099801071 | 8,16E-13 | 5,44E-11 | Up |
| Fdps | 2752,942083 | -1,228309297 | 0,249029777 | 1,35E-09 | 3,80E-08 | Down |
| Dcst1 | 28,3747227 | -1,097200096 | 0,484144562 | 5,15E-05 | 0,000407067 | Down |
| Pmvk | 4873,250219 | -1,046018408 | 0,13788678 | 1,71E-17 | 2,99E-15 | Down |
| Il6ra | 221,0480683 | 1,730685408 | 0,176173437 | 3,21E-24 | 1,74E-21 | Up |
| Selenbp2 | 107,7579436 | -0,708050485 | 0,29252255 | 0,000142361 | 0,000977863 | Down |
| Selenbp1 | 1431,108593 | -0,657727126 | 0,250151116 | 0,000123596 | 0,000864072 | Down |
| Ctss | 663,8275755 | 1,323026603 | 0,434200637 | 4,13E-06 | 4,40E-05 | Up |
| Gm17690 | 58,03328332 | -0,68070525 | 0,244890974 | 6,72E-05 | 0,00050986 | Down |
| Hist2h2be | 164,9131176 | -0,651970612 | 0,167602851 | 2,32E-06 | 2,65E-05 | Down |
| Fcgr1 | 5289,287716 | 1,05887229 | 0,158057193 | 1,31E-14 | 1,24E-12 | Up |
| I830077J02Rik | 350,8043149 | -0,643017947 | 0,246095377 | 0,00014679 | 0,001004336 | Down |

| | | | | | | |
|---|---|---|---|---|---|---|
| Wdr77 | 5436,562739 | 0,821732376 | 0,100204862 | 6,69E-19 | 1,64E-16 | Up |
| Cd53 | 2647,867443 | 3,005099805 | 0,535603366 | 3,17E-10 | 1,06E-08 | Up |
| Csf1 | 1974,845064 | 1,271975004 | 0,237712551 | 1,95E-10 | 6,99E-09 | Up |
| Celsr2 | 444,6304532 | -0,941900567 | 0,367680786 | 2,83E-05 | 0,000240162 | Down |
| Gm42776 | 16,6969338 | 0,723625414 | 0,614643785 | 0,001863999 | 0,008988831 | Up |
| Dnttip2 | 14593,69658 | 0,642046527 | 0,09453244 | 4,94E-13 | 3,47E-11 | Up |
| 5730508B09Rik | 697,5044354 | 0,764097434 | 0,139667535 | 3,24E-10 | 1,08E-08 | Up |
| Casp6 | 37,76323369 | 0,617045308 | 0,313405409 | 0,000791876 | 0,004310744 | Up |
| Bank1 | 390,3070533 | 0,903480132 | 0,310751761 | 1,01E-05 | 9,71E-05 | Up |
| 9530052C20Rik | 67,58235858 | -0,589147758 | 0,259824772 | 0,000534182 | 0,003072913 | Down |
| Dnajb4 | 3052,467561 | -0,692823725 | 0,148799174 | 5,59E-08 | 9,87E-07 | Down |
| A830012C17Rik | 10,27996963 | 2,694766703 | 1,957544845 | 2,98E-05 | 0,000250757 | Up |
| Gem | 254,5110708 | 3,230923671 | 2,256565574 | 7,86E-06 | 7,85E-05 | Up |
| Cdh17 | 137,6528549 | 5,479043513 | 1,392533402 | 2,05E-08 | 4,07E-07 | Up |
| Ttpa | 282,5969096 | 0,718414693 | 0,140453335 | 4,20E-09 | 1,02E-07 | Up |
| Ndufaf4 | 4332,278384 | 0,884659014 | 0,125951938 | 2,59E-15 | 2,89E-13 | Up |
| Epha7 | 24,0133338 | 0,739374423 | 0,574929151 | 0,001476614 | 0,007362456 | Up |
| Mdn1 | 16966,43841 | 0,611247549 | 0,199751544 | 5,78E-05 | 0,000450066 | Up |
| 4933421O10Rik | 340,2676421 | -0,669049445 | 0,149318753 | 1,66E-07 | 2,67E-06 | Down |
| Cd72 | 402,1646575 | -1,225679089 | 0,156434055 | 1,12E-17 | 2,06E-15 | Down |
| Polr1e | 6301,415042 | 0,582469646 | 0,095785324 | 8,72E-11 | 3,43E-09 | Up |
| 4933430I17Rik | 79,94493225 | 0,632590087 | 0,296879138 | 0,000501693 | 0,002921305 | Up |
| Tnfsf8 | 2093,575493 | 1,502513487 | 0,130403202 | 4,12E-32 | 6,47E-29 | Up |
| Mtap | 12636,51083 | 0,803858397 | 0,133757585 | 7,51E-12 | 3,86E-10 | Up |
| Tek | 20,72839217 | 1,21327381 | 0,489553927 | 2,45E-05 | 0,00021086 | Up |
| Gm12708 | 12,12002274 | 0,841041364 | 0,657547063 | 0,001050756 | 0,005478992 | Up |
| Nfia | 299,0884067 | -1,105347909 | 0,197412724 | 1,95E-11 | 9,28E-10 | Down |
| Gm12796 | 25,86428539 | -0,777712305 | 0,366082046 | 0,000199689 | 0,001309813 | Down |
| Lepr | 34,45425309 | 1,777676477 | 0,520606978 | 1,72E-06 | 2,05E-05 | Up |
| Pde4b | 1526,193179 | -0,752234459 | 0,129764614 | 5,92E-11 | 2,44E-09 | Down |
| Echdc2 | 151,1693368 | 0,615896159 | 0,207155379 | 7,11E-05 | 0,000535128 | Up |
| Zfyve9 | 604,7746878 | 1,030672964 | 0,263949626 | 1,26E-07 | 2,07E-06 | Up |
| Cdkn2c | 6893,065099 | -0,615712969 | 0,088813941 | 2,47E-13 | 1,81E-11 | Down |
| Foxd2 | 9,982544839 | -0,71231915 | 0,609319199 | 0,00197239 | 0,009396205 | Down |
| Eif2b3 | 7067,412782 | 0,624755911 | 0,100430171 | 2,50E-11 | 1,15E-09 | Up |
| Mpl | 618,1466412 | -0,622346543 | 0,465131719 | 0,002331261 | 0,010871839 | Down |
| Olfr1342 | 762,4197633 | -0,832275152 | 0,222511733 | 6,44E-07 | 8,80E-06 | Down |
| Csf3r | 399,1508416 | 1,227332327 | 0,494264758 | 2,39E-05 | 0,00020686 | Up |
| Oscp1 | 235,554472 | -0,636412807 | 0,180825367 | 1,04E-05 | 9,99E-05 | Down |
| Tmem35b | 1067,155059 | -0,607468804 | 0,099065769 | 5,14E-11 | 2,14E-09 | Down |
| Gjb3 | 19,38929696 | 1,412501845 | 0,652299309 | 5,02E-05 | 0,000397484 | Up |
| Fndc5 | 39,53355946 | -1,297010797 | 0,414686261 | 3,11E-06 | 3,42E-05 | Down |
| Lck | 92,78525319 | -1,558220253 | 0,474386854 | 2,28E-06 | 2,61E-05 | Down |
| Spocd1 | 103,4212202 | -1,590947728 | 0,328693782 | 8,37E-09 | 1,86E-07 | Down |
| Ahdc1 | 3993,025044 | -0,625637761 | 0,09339264 | 1,11E-12 | 7,23E-11 | Down |
| Pigv | 1139,638197 | -0,597863261 | 0,093505995 | 1,08E-11 | 5,36E-10 | Down |
| Man1c1 | 2246,77068 | 1,885989915 | 0,204071383 | 6,60E-22 | 2,80E-19 | Up |
| Ece1 | 103,7080613 | -0,927217832 | 0,356634511 | 2,62E-05 | 0,000224082 | Down |
| Pla2g2e | 33,9934475 | 1,670646906 | 0,941081547 | 9,55E-05 | 0,000688998 | Up |
| Mrto4 | 19176,00783 | 0,650943696 | 0,121634318 | 2,89E-09 | 7,39E-08 | Up |
| Padi6 | 31,51235846 | -1,784154846 | 0,38703911 | 2,92E-08 | 5,53E-07 | Down |

| | | | | | | |
|---|---|---|---|---|---|---|
| B330016D10Rik | 305,906685 | -0,847211142 | 0,127147485 | 5,51E-14 | 4,69E-12 | Down |
| Kazn | 3608,778112 | -0,699081804 | 0,131550083 | 1,92E-09 | 5,14E-08 | Down |
| Tnfrsf1b | 5944,912803 | 0,584529197 | 0,115916373 | 2,81E-08 | 5,34E-07 | Up |
| Zfp988 | 236,19846 | 0,702085427 | 0,198268131 | 4,86E-06 | 5,06E-05 | Up |
| Clcn6 | 2648,135738 | -0,877707506 | 0,18062347 | 2,26E-09 | 5,97E-08 | Down |
| Srm | 41624,26871 | 0,690430464 | 0,084919339 | 1,13E-17 | 2,06E-15 | Up |
| Gm13205 | 187,7671106 | 0,896719165 | 0,179064463 | 8,68E-10 | 2,61E-08 | Up |
| Slc25a33 | 1909,241325 | 0,897075362 | 0,106128564 | 2,44E-20 | 8,15E-18 | Up |
| Gm13073 | 34,71055071 | -1,243221536 | 0,547198337 | 4,33E-05 | 0,000349991 | Down |
| Gm13091 | 135,4061648 | -0,686589854 | 0,345692718 | 0,000470619 | 0,002754655 | Down |
| Plekhg5 | 3224,120707 | -0,60094022 | 0,136970067 | 5,20E-07 | 7,30E-06 | Down |
| Chd5 | 261,9581367 | -2,045975479 | 0,513987759 | 3,27E-07 | 4,85E-06 | Down |
| Mib2 | 2229,653776 | -0,717634282 | 0,224645083 | 1,36E-05 | 0,000126885 | Down |
| Atad3a | 19498,28592 | 0,588418443 | 0,102321405 | 5,92E-10 | 1,86E-08 | Up |
| Pusl1 | 1596,946224 | 0,586978936 | 0,185810936 | 5,42E-05 | 0,000426014 | Up |
| Rnf223 | 55,244501 | -0,728653143 | 0,270663613 | 5,79E-05 | 0,00045073 | Down |
| Samd11 | 115,4863289 | 0,772093006 | 0,370612902 | 0,000228356 | 0,001472042 | Up |
| Cyp51 | 12639,3396 | -0,668370176 | 0,17788016 | 3,19E-06 | 3,50E-05 | Down |
| Abcb1b | 4149,851263 | 1,145037198 | 0,116832152 | 1,18E-25 | 7,13E-23 | Up |
| 5031425E22Rik | 1281,159012 | -0,799205056 | 0,094652547 | 1,28E-19 | 3,44E-17 | Down |
| Nos3 | 60,47437923 | 0,703980938 | 0,324100634 | 0,000273096 | 0,001725766 | Up |
| Insig1 | 4185,584737 | -0,784519202 | 0,236290917 | 4,85E-06 | 5,05E-05 | Down |
| Khk | 614,7936558 | -0,678272344 | 0,135290572 | 1,19E-08 | 2,53E-07 | Down |
| Slc5a6 | 1836,307114 | 0,703362969 | 0,12471261 | 3,00E-10 | 1,01E-08 | Up |
| Ift172 | 1798,684001 | -0,702976137 | 0,084842261 | 2,59E-18 | 5,57E-16 | Down |
| AI839979 | 54,48895441 | 1,848696954 | 0,553315934 | 2,15E-06 | 2,47E-05 | Up |
| Plb1 | 16,8012613 | 2,185609995 | 0,667073811 | 2,55E-06 | 2,88E-05 | Up |
| Gm15614 | 21,15205568 | -0,744221331 | 0,530009084 | 0,001174352 | 0,006055156 | Down |
| Gm9903 | 32,41987642 | -0,615559976 | 0,328307397 | 0,00096273 | 0,005084094 | Down |
| Rgs12 | 20,80970257 | 1,176528372 | 0,691802614 | 0,000202471 | 0,001324743 | Up |
| Hgfac | 155,0153472 | 0,670244634 | 0,210324925 | 2,20E-05 | 0,000194536 | Up |
| Sh3tc1 | 912,8728443 | 0,874630774 | 0,167410205 | 3,20E-10 | 1,07E-08 | Up |
| Ppp2r2c | 9,036617986 | -0,586499892 | 0,594726651 | 0,004255894 | 0,017912596 | Down |
| Clnk | 240,0028538 | 1,125666232 | 0,226782213 | 7,51E-10 | 2,29E-08 | Up |
| Bst1 | 33,67837635 | 2,731042112 | 1,076490207 | 1,03E-05 | 9,89E-05 | Up |
| Prom1 | 9,441801031 | 0,867386089 | 1,174819961 | 0,001411688 | 0,007085943 | Up |
| Wdr19 | 376,459108 | -0,845936949 | 0,165641906 | 8,26E-10 | 2,49E-08 | Down |
| N4bp2os | 74,35762309 | -0,832881748 | 0,275077446 | 9,56E-06 | 9,26E-05 | Down |
| Apbb2 | 43,3166444 | -0,668708622 | 0,398741615 | 0,00099999 | 0,005249113 | Down |
| Ppbp | 98,37811165 | 4,446625985 | 0,951350652 | 1,37E-08 | 2,85E-07 | Up |
| Pf4 | 352,6255095 | 1,865863793 | 0,345923495 | 9,06E-10 | 2,71E-08 | Up |
| Cxcl2 | 57,55468698 | 2,265030154 | 0,945794071 | 1,80E-05 | 0,000163184 | Up |
| Thap6 | 687,4740247 | -0,59547924 | 0,10086598 | 2,28E-10 | 8,06E-09 | Down |
| Sdad1 | 12175,14052 | 0,804308721 | 0,103102495 | 2,35E-17 | 3,93E-15 | Up |
| Ccng2 | 1329,31746 | -0,76855244 | 0,225506038 | 4,06E-06 | 4,33E-05 | Down |
| Gm7993 | 11509,51679 | 0,745691326 | 0,090944514 | 2,62E-18 | 5,57E-16 | Up |
| Anxa3 | 8980,662929 | 0,779390453 | 0,296277954 | 4,84E-05 | 0,00038543 | Up |
| Spp1 | 14166,01247 | 1,354980421 | 0,213332429 | 1,11E-12 | 7,23E-11 | Up |
| Gm43819 | 34,74630871 | -0,776005475 | 0,494938881 | 0,000746846 | 0,004096835 | Down |
| Gm42517 | 916,9586079 | -1,456516066 | 0,141124161 | 1,70E-26 | 1,22E-23 | Down |
| Gm10419 | 2401,713468 | -1,044112814 | 0,108534084 | 2,66E-25 | 1,55E-22 | Down |

| | | | | | | |
|---|---|---|---|---|---|---|
| Gm18555 | 102,3025521 | -1,180853091 | 0,313402861 | 2,47E-07 | 3,81E-06 | Down |
| Ttc28 | 1354,800972 | -0,611378866 | 0,152030525 | 2,13E-06 | 2,46E-05 | Down |
| Tmem119 | 172,4539261 | 0,847428787 | 0,186964088 | 1,58E-08 | 3,23E-07 | Up |
| Ssh1 | 468,8903113 | -0,667717421 | 0,230530422 | 5,42E-05 | 0,000425591 | Down |
| Mmab | 3784,298737 | -0,593783613 | 0,115738206 | 1,66E-08 | 3,36E-07 | Down |
| Mvk | 3925,470077 | -0,987658498 | 0,202985273 | 1,16E-09 | 3,36E-08 | Down |
| Rbm19 | 8907,972912 | 0,63065162 | 0,121813313 | 9,09E-09 | 2,00E-07 | Up |
| Slc8b1 | 1370,302668 | 0,686376933 | 0,145913697 | 4,85E-08 | 8,73E-07 | Up |
| Oas2 | 153,550936 | 1,280076991 | 0,417888922 | 3,83E-06 | 4,12E-05 | Up |
| P2rx7 | 32,61155604 | 2,124565511 | 0,83576665 | 1,43E-05 | 0,000133159 | Up |
| Tmem120b | 909,7917846 | 0,633014675 | 0,121272947 | 7,11E-09 | 1,61E-07 | Up |
| Hcar2 | 29,20565238 | 3,824299699 | 0,714103058 | 1,31E-09 | 3,71E-08 | Up |
| Clip2 | 3968,004608 | -0,8653501 | 0,080658016 | 8,68E-30 | 1,07E-26 | Down |
| Wbscr27 | 680,0789105 | -0,832074584 | 0,181311221 | 1,40E-08 | 2,88E-07 | Down |
| Fkbp6 | 271,5532543 | -0,695234112 | 0,127202882 | 8,90E-10 | 2,67E-08 | Down |
| Ssc4d | 446,6429888 | -0,826757398 | 0,142486633 | 1,93E-11 | 9,24E-10 | Down |
| Sh2b2 | 4920,838545 | 0,675070797 | 0,133560829 | 1,01E-08 | 2,19E-07 | Up |
| Cux1 | 58335,39229 | -0,637918021 | 0,119335111 | 3,47E-09 | 8,73E-08 | Down |
| Myl10 | 156,465505 | -1,740209055 | 0,455248685 | 4,83E-07 | 6,86E-06 | Down |
| Tfr2 | 6268,32924 | -1,284161952 | 0,21247842 | 4,27E-12 | 2,37E-10 | Down |
| Nyap1 | 219,7785756 | -0,666954275 | 0,203093212 | 1,67E-05 | 0,000152271 | Down |
| Pilra | 72,16270277 | 2,1058125 | 0,375078139 | 2,76E-10 | 9,40E-09 | Up |
| Pilrb1 | 241,8391833 | 2,672169105 | 0,285109617 | 1,18E-22 | 5,60E-20 | Up |
| Gm36551 | 78,80995591 | 2,14504223 | 0,300368269 | 1,60E-14 | 1,50E-12 | Up |
| Pilrb2 | 19,10539615 | 0,65317089 | 0,683335095 | 0,003203331 | 0,014093938 | Up |
| Nxpe5 | 888,0046898 | 1,299434996 | 0,794603246 | 0,000195642 | 0,001287569 | Up |
| BC037034 | 9528,791345 | 0,617301053 | 0,192561957 | 3,51E-05 | 0,000290711 | Up |
| 3110082I17Rik | 13716,31976 | 0,782517782 | 0,100660788 | 4,28E-17 | 6,73E-15 | Up |
| Gpr146 | 6276,57489 | -0,760106007 | 0,086958784 | 1,97E-20 | 6,73E-18 | Down |
| Micall2 | 18999,42081 | -0,585920495 | 0,096369545 | 8,54E-11 | 3,36E-09 | Down |
| Snx8 | 11856,6436 | 0,598540368 | 0,11420489 | 8,97E-09 | 1,98E-07 | Up |
| Lfng | 13135,06574 | 0,855261657 | 0,080798309 | 4,88E-29 | 4,79E-26 | Up |
| Cyth3 | 32,66380779 | 0,984929898 | 0,487730896 | 0,000127716 | 0,000888021 | Up |
| Gm43460 | 384,162468 | -0,808493852 | 0,133649725 | 5,49E-12 | 2,92E-10 | Down |
| Flt1 | 206,4400117 | 1,234194769 | 0,233184153 | 2,14E-10 | 7,59E-09 | Up |
| 8430423G03Rik | 88,71111414 | 0,691763404 | 0,273029466 | 0,000119729 | 0,000839654 | Up |
| Hsph1 | 59171,53977 | 0,631019126 | 0,129936806 | 4,50E-08 | 8,13E-07 | Up |
| Gm5873 | 16,5346979 | -1,002558686 | 0,622134131 | 0,000344512 | 0,002109215 | Down |
| 2610001J05Rik | 9642,694611 | 0,617774567 | 0,107102998 | 4,09E-10 | 1,34E-08 | Up |
| Tfec | 6794,885541 | 0,724519222 | 0,098183814 | 2,38E-15 | 2,73E-13 | Up |
| Met | 4940,381982 | 1,218097174 | 0,163005717 | 1,64E-16 | 2,26E-14 | Up |
| Asz1 | 351,0637868 | -1,049751097 | 0,406380893 | 2,07E-05 | 0,000184241 | Down |
| Hilpda | 1219,348805 | 0,62459654 | 0,100917405 | 3,05E-11 | 1,38E-09 | Up |
| Cpa5 | 32,44097674 | -1,484025112 | 0,373918389 | 2,12E-07 | 3,31E-06 | Down |
| Rab19 | 13,70314557 | 1,903448561 | 1,120167545 | 8,71E-05 | 0,000637862 | Up |
| Clec5a | 2770,782305 | 2,213885668 | 0,338235374 | 9,31E-13 | 6,17E-11 | Up |
| Trbv12-2 | 42,5808743 | -1,024049837 | 0,31364286 | 1,88E-06 | 2,21E-05 | Down |
| Trbv13-2 | 116,371446 | -0,866493601 | 0,236862053 | 7,01E-07 | 9,41E-06 | Down |
| Gm43889 | 34,29366836 | 3,269367438 | 0,750117783 | 1,02E-07 | 1,69E-06 | Up |
| A930035D04Rik | 54,44041785 | -0,732666062 | 0,288183218 | 8,58E-05 | 0,000630586 | Down |
| Gimap8 | 581,8541394 | -0,895071086 | 0,266823599 | 2,01E-06 | 2,33E-05 | Down |

| | | | | | | |
|---|---|---|---|---|---|---|
| Gimap5 | 1384,344771 | -0,727763536 | 0,162718025 | 8,36E-08 | 1,41E-06 | Down |
| Tmem176a | 11316,0182 | -0,625273527 | 0,185113046 | 1,88E-05 | 0,000168913 | Down |
| Ggct | 684,1492907 | 0,817728478 | 0,14458084 | 5,24E-11 | 2,18E-09 | Up |
| Ccser1 | 22,01476401 | 2,783244 | 0,869127935 | 2,70E-06 | 3,03E-05 | Up |
| Hpgds | 21,68390694 | 0,792089607 | 0,551555197 | 0,000919907 | 0,004897444 | Up |
| Gm6786 | 14,5831849 | 1,632998279 | 0,885527113 | 8,62E-05 | 0,000632114 | Up |
| Smyd1 | 43,64976419 | -0,677563716 | 0,255446983 | 9,80E-05 | 0,000703 | Down |
| Tmem150a | 223,9175703 | 0,781321882 | 0,174212229 | 4,08E-08 | 7,47E-07 | Up |
| Mat2a | 30186,51922 | 0,737195564 | 0,095326652 | 1,29E-16 | 1,85E-14 | Up |
| Particl | 170,8602644 | -0,636130511 | 0,148406271 | 5,60E-07 | 7,79E-06 | Down |
| Gm17034 | 20,10568958 | -0,948610213 | 0,545343577 | 0,000288761 | 0,00181454 | Down |
| Rab11fip5 | 155,2319165 | 1,073010039 | 0,535495952 | 0,000111839 | 0,00079067 | Up |
| Add2 | 176,2583898 | -0,875052452 | 0,376979719 | 7,47E-05 | 0,000557189 | Down |
| Fam136a | 15246,30518 | 0,585594231 | 0,101064246 | 4,71E-10 | 1,52E-08 | Up |
| Gm43904 | 7,527783735 | -1,53395413 | 1,141199912 | 0,000241968 | 0,001551517 | Down |
| Gp9 | 601,7823474 | -0,89669564 | 0,282619259 | 3,98E-06 | 4,26E-05 | Down |
| Mgll | 53,88534522 | 0,733276692 | 0,865565636 | 0,002483883 | 0,011449383 | Up |
| Chst13 | 19,81342215 | 0,622290409 | 0,517793284 | 0,00282517 | 0,012722169 | Up |
| Chchd4 | 12949,78282 | 0,684529343 | 0,098061241 | 7,78E-14 | 6,38E-12 | Up |
| Gm16433 | 130,5902321 | 0,621076995 | 0,206447987 | 6,11E-05 | 0,00047195 | Up |
| Bhlhe40 | 21345,23936 | 0,721165599 | 0,126054993 | 1,46E-10 | 5,49E-09 | Up |
| Mtmr14 | 5290,174629 | -0,674610947 | 0,100140943 | 4,81E-13 | 3,39E-11 | Down |
| Il17re | 269,1383942 | -1,482229147 | 0,257024405 | 6,54E-11 | 2,68E-09 | Down |
| Cand2 | 1623,639553 | -1,539164521 | 0,232683776 | 5,77E-13 | 3,99E-11 | Down |
| Gm8213 | 184,2105679 | -0,607676225 | 0,180071046 | 2,26E-05 | 0,000198064 | Down |
| Fxyd4 | 283,3115694 | 0,722907651 | 0,239241248 | 2,25E-05 | 0,000197704 | Up |
| Phc1 | 2601,639079 | -0,96751085 | 0,107656419 | 1,09E-22 | 5,34E-20 | Down |
| C3ar1 | 122,548269 | 0,67570723 | 0,3600777 | 0,000639356 | 0,003577115 | Up |
| Clec4n | 64,08773858 | 2,528252479 | 1,050368869 | 1,50E-05 | 0,000138422 | Up |
| C1ra | 714,1074443 | 0,698977979 | 0,270464099 | 9,87E-05 | 0,000707157 | Up |
| Lag3 | 151,5211733 | -1,644643602 | 0,206791868 | 5,44E-17 | 8,38E-15 | Down |
| Nop2 | 29253,70513 | 0,771489785 | 0,129660133 | 1,77E-11 | 8,54E-10 | Up |
| Fgf23 | 169,6018244 | 1,935846926 | 0,681497718 | 7,78E-06 | 7,78E-05 | Up |
| Tigar | 2775,434518 | 0,831508254 | 0,139997751 | 7,84E-12 | 4,00E-10 | Up |
| Gm26770 | 159,7417486 | -1,000797076 | 0,21186371 | 2,41E-09 | 6,28E-08 | Down |
| Rhno1 | 8719,928351 | -0,847295721 | 0,108567454 | 1,12E-17 | 2,06E-15 | Down |
| Cd69 | 4378,454413 | 1,955245244 | 0,294403919 | 6,20E-13 | 4,26E-11 | Up |
| Clec12a | 1081,153479 | 0,657104754 | 0,66663714 | 0,003072613 | 0,013617931 | Up |
| Styk1 | 7,943591511 | 2,301663515 | 1,257318927 | 4,71E-05 | 0,000377482 | Up |
| Bcl2l14 | 2146,088438 | 1,136609488 | 0,148853669 | 2,10E-17 | 3,55E-15 | Up |
| Mansc1 | 318,8005865 | 0,900913918 | 0,168695141 | 1,30E-10 | 4,94E-09 | Up |
| Emp1 | 7,454085829 | 0,8191927 | 1,303782094 | 0,001074749 | 0,005592967 | Up |
| Eps8 | 255,8963776 | -0,726087104 | 0,2356251 | 1,81E-05 | 0,000163575 | Down |
| Gm15707 | 291,8809628 | -0,583323617 | 0,178731248 | 4,06E-05 | 0,000330629 | Down |
| Fam60a | 3785,475364 | 0,692956612 | 0,080407283 | 1,77E-19 | 4,57E-17 | Up |
| Lair1 | 2092,32175 | 0,66366172 | 0,145073398 | 1,16E-07 | 1,92E-06 | Up |
| Rdh13 | 4023,991193 | 0,859743204 | 0,139338936 | 1,26E-12 | 8,07E-11 | Up |
| Zfp580 | 1035,542264 | -0,821593807 | 0,12420183 | 1,12E-13 | 8,84E-12 | Down |
| Zfp128 | 642,141913 | -0,610981398 | 0,109541272 | 1,29E-09 | 3,66E-08 | Down |
| Ehd2 | 210,8835274 | 0,693240879 | 0,34654312 | 0,000436969 | 0,002584645 | Up |
| C5ar1 | 80,99254291 | 2,090309807 | 0,346172199 | 2,47E-11 | 1,14E-09 | Up |

| | | | | | | |
|---|---|---|---|---|---|---|
| Gng8 | 26,98188007 | 1,025507475 | 0,743053981 | 0,00053754 | 0,003087714 | Up |
| Ptgir | 1602,251641 | 0,708850468 | 0,213002556 | 9,49E-06 | 9,21E-05 | Up |
| Psg17 | 14,23180398 | -1,182551838 | 1,396523119 | 0,000600997 | 0,003392695 | Down |
| Cd3eap | 8986,04644 | 0,614533766 | 0,144692788 | 8,27E-07 | 1,08E-05 | Up |
| Plaur | 2252,858594 | 0,627660868 | 0,153358978 | 1,36E-06 | 1,67E-05 | Up |
| Cd177 | 3314,386718 | 0,589659535 | 0,196624678 | 8,48E-05 | 0,000624251 | Up |
| Grik5 | 7642,925011 | -0,592375642 | 0,093028247 | 1,34E-11 | 6,56E-10 | Down |
| Ccdc97 | 5506,470297 | -0,666127077 | 0,085306428 | 2,09E-16 | 2,78E-14 | Down |
| Numbl | 178,9177678 | -0,673781846 | 0,327398553 | 0,000436162 | 0,002581661 | Down |
| Prx | 249,3020605 | -0,804918308 | 0,195468278 | 1,66E-07 | 2,67E-06 | Down |
| Gm44685 | 125,6007902 | -1,159847931 | 0,248441963 | 3,85E-09 | 9,52E-08 | Down |
| 9530053A07Rik | 88,21856048 | 2,720897435 | 0,438747703 | 8,35E-12 | 4,23E-10 | Up |
| Rasgrp4 | 12,14188763 | 0,638469753 | 0,73599607 | 0,003594806 | 0,015486833 | Up |
| Tyrobp | 8140,496444 | 0,604276996 | 0,178641572 | 2,28E-05 | 0,00019915 | Up |
| Proser3 | 945,1244147 | -0,722268313 | 0,161732254 | 9,23E-08 | 1,55E-06 | Down |
| Hspb6 | 1201,830233 | -0,723907504 | 0,128913239 | 2,57E-10 | 8,86E-09 | Down |
| Dmkn | 61,86508586 | 2,081055947 | 0,457901072 | 4,28E-08 | 7,80E-07 | Up |
| Ffar1 | 8,299548859 | -0,581733062 | 0,674857529 | 0,004671046 | 0,019330291 | Down |
| Fxyd5 | 29060,09297 | 0,623318218 | 0,124868193 | 2,51E-08 | 4,84E-07 | Up |
| Cep89 | 2843,62441 | -0,709893388 | 0,17620991 | 6,98E-07 | 9,39E-06 | Down |
| Rgs9bp | 50,43011054 | -0,898285856 | 0,290415002 | 5,31E-06 | 5,51E-05 | Down |
| 2610021A01Rik | 2849,921361 | 0,59394277 | 0,081064494 | 1,77E-14 | 1,64E-12 | Up |
| Nkg7 | 1081,063626 | 0,751174189 | 0,159334451 | 2,00E-08 | 3,98E-07 | Up |
| Cd33 | 317,7481236 | 2,039739702 | 0,592420772 | 1,64E-06 | 1,97E-05 | Up |
| Kcnc3 | 605,3869334 | 0,70341732 | 0,273889595 | 9,96E-05 | 0,000712937 | Up |
| Atf5 | 8887,322386 | -0,708408549 | 0,216991927 | 1,18E-05 | 0,000111129 | Down |
| Rras | 783,3356231 | -0,66554446 | 0,212485172 | 2,74E-05 | 0,000233385 | Down |
| Ntf5 | 118,2839326 | -0,599776608 | 0,193537823 | 5,75E-05 | 0,000448307 | Down |
| Tulp2 | 34,43347811 | -0,613449562 | 0,407724185 | 0,001890095 | 0,009078462 | Down |
| A030001D20Rik | 31,10105575 | -0,945552063 | 0,418814894 | 7,15E-05 | 0,000537136 | Down |
| Grwd1 | 15240,31023 | 0,779048011 | 0,136010049 | 5,96E-11 | 2,45E-09 | Up |
| Nomo1 | 42213,93297 | 0,628251656 | 0,108738839 | 3,48E-10 | 1,15E-08 | Up |
| Saa3 | 230,6534583 | -0,617018175 | 0,168418613 | 7,70E-06 | 7,70E-05 | Down |
| Saa4 | 11,52120595 | 2,724103382 | 1,760835393 | 3,21E-05 | 0,000269045 | Up |
| Gtf2h1 | 19111,93618 | 0,745036431 | 0,118377113 | 3,13E-12 | 1,79E-10 | Up |
| Gm2788 | 162,2014534 | -0,747241236 | 0,271103229 | 4,16E-05 | 0,000337691 | Down |
| Gm7367 | 52,18244903 | -0,661570716 | 0,283249112 | 0,0002505 | 0,001599046 | Down |
| Mphosph10 | 9516,09996 | 0,633205322 | 0,123195556 | 1,07E-08 | 2,30E-07 | Up |
| 1810008I18Rik | 31,5009811 | 0,97720686 | 0,670586853 | 0,000512988 | 0,002973856 | Up |
| Akap13 | 34518,55022 | -0,674728682 | 0,124375027 | 1,46E-09 | 4,03E-08 | Down |
| Aen | 10887,20713 | 0,981630811 | 0,128586259 | 1,15E-17 | 2,07E-15 | Up |
| Gm21057 | 58,34856749 | -0,766333286 | 0,258810427 | 1,91E-05 | 0,000171085 | Down |
| Zscan2 | 576,059414 | -0,585564801 | 0,108154363 | 3,99E-09 | 9,78E-08 | Down |
| Alpk3 | 1330,12285 | -0,658274102 | 0,15011715 | 2,87E-07 | 4,34E-06 | Down |
| 1700010L04Rik | 60,69247392 | 0,841522805 | 0,250418914 | 2,70E-06 | 3,03E-05 | Up |
| Abhd17c | 18929,1166 | 0,589610577 | 0,121081477 | 6,32E-08 | 1,10E-06 | Up |
| Arnt2 | 849,7298932 | 1,336308322 | 0,452703817 | 5,62E-06 | 5,79E-05 | Up |
| Rab38 | 29661,05411 | -0,598966798 | 0,090005705 | 1,92E-12 | 1,18E-10 | Down |
| Usp35 | 183,9917536 | -0,716698827 | 0,237137214 | 2,38E-05 | 0,000205967 | Down |
| Lrrc32 | 9,104524139 | 1,407478434 | 1,20867366 | 0,000370891 | 0,002249681 | Up |
| Dgat2 | 116,9439592 | -0,642552834 | 0,362182249 | 0,000971482 | 0,005119992 | Down |

| | | | | | | |
|---|---|---|---|---|---|---|
| Gdpd5 | 21,89444491 | 1,859154582 | 0,48909326 | 5,51E-07 | 7,69E-06 | Up |
| Stard10 | 87,27593361 | -0,581287842 | 0,255386901 | 0,000561552 | 0,00320805 | Down |
| Folr1 | 1260,518609 | 0,778510961 | 0,270058326 | 2,24E-05 | 0,000196628 | Up |
| Usp17ld | 816,2852816 | 3,192081699 | 0,204826206 | 3,50E-56 | 5,50E-52 | Up |
| Usp17lc | 25,65547678 | 3,770762201 | 1,525698836 | 3,42E-06 | 3,73E-05 | Up |
| Trim30d | 91,68218026 | -0,762651136 | 0,367110178 | 0,000243413 | 0,001559513 | Down |
| Gm4070 | 372,7722627 | 1,034694154 | 0,266882647 | 1,43E-07 | 2,31E-06 | Up |
| Knop1 | 19099,91644 | 0,625147765 | 0,094930143 | 2,38E-12 | 1,42E-10 | Up |
| 9030407P20Rik | 153,6849365 | -0,642217681 | 0,147611041 | 3,99E-07 | 5,79E-06 | Down |
| 2210406H18Rik | 45,35380974 | 0,959206927 | 0,398617092 | 4,40E-05 | 0,00035533 | Up |
| Igsf6 | 1312,914639 | 1,581228562 | 0,20835196 | 8,06E-16 | 9,82E-14 | Up |
| Lat | 308,9352463 | -1,576960392 | 0,220601287 | 1,89E-14 | 1,73E-12 | Down |
| Mylpf | 586,579511 | -0,641915968 | 0,12711482 | 1,54E-08 | 3,15E-07 | Down |
| Ccdc189 | 34,1889125 | -0,759472055 | 0,342805711 | 0,000174039 | 0,001168889 | Down |
| Itgam | 2870,78632 | 1,059874233 | 0,24436188 | 1,64E-08 | 3,34E-07 | Up |
| Itgax | 157,6541338 | -1,008095242 | 0,384986172 | 1,99E-05 | 0,000177535 | Down |
| 1500002F19Rik | 43,92224363 | 1,371715619 | 0,433054839 | 2,88E-06 | 3,20E-05 | Up |
| Cfap46 | 27,52282962 | -0,730587705 | 0,429887219 | 0,000689066 | 0,003815246 | Down |
| Adam8 | 724,7217499 | 2,842951783 | 0,408720719 | 6,07E-14 | 5,10E-12 | Up |
| Ifitm1 | 142241,0703 | 0,962743024 | 0,178707308 | 6,70E-11 | 2,73E-09 | Up |
| Gm7514 | 58,25403277 | 2,290126646 | 0,341504489 | 3,03E-13 | 2,19E-11 | Up |
| Ifitm3 | 93618,87326 | 1,907982177 | 0,276466849 | 1,13E-13 | 8,84E-12 | Up |
| Ifitm6 | 396,4509885 | 3,814629386 | 0,69177822 | 5,89E-10 | 1,85E-08 | Up |
| Lrrc56 | 288,3536262 | -0,760677946 | 0,162359136 | 2,04E-08 | 4,05E-07 | Down |
| Slc25a22 | 4130,3119 | 0,900232151 | 0,15029263 | 2,56E-12 | 1,51E-10 | Up |
| Tspan4 | 10535,26859 | 0,76902874 | 0,099414317 | 7,34E-17 | 1,09E-14 | Up |
| Muc6 | 10,9192414 | -1,291597105 | 0,815957059 | 0,000221085 | 0,001429864 | Down |
| Tnni2 | 70,68094249 | 1,011929906 | 0,303158983 | 1,44E-06 | 1,76E-05 | Up |
| Tnfrsf23 | 77,78628846 | 0,967057444 | 0,407365437 | 4,76E-05 | 0,00038117 | Up |
| Dhcr7 | 10305,01746 | -0,995604704 | 0,181844388 | 3,60E-11 | 1,59E-09 | Down |
| Camsap3 | 26,6459717 | -1,024999533 | 0,448769942 | 5,50E-05 | 0,000430827 | Down |
| Col4a2 | 1922,424638 | 1,876283066 | 0,170265241 | 8,81E-30 | 1,07E-26 | Up |
| Mcf2l | 271,8840191 | 1,476544505 | 0,863351977 | 0,000133771 | 0,000925738 | Up |
| F7 | 73,06081911 | 2,197755231 | 0,632543902 | 1,51E-06 | 1,83E-05 | Up |
| Adprhl1 | 11,71580451 | -2,758706082 | 0,777975759 | 1,19E-06 | 1,49E-05 | Down |
| Gm7676 | 30,2888419 | 1,108768879 | 0,437633041 | 2,25E-05 | 0,000197605 | Up |
| Adgra2 | 117,7847931 | -0,834173767 | 0,224947063 | 7,24E-07 | 9,69E-06 | Down |
| Gm29243 | 387,7691212 | 0,751441496 | 0,174060718 | 1,25E-07 | 2,05E-06 | Up |
| Dctd | 5876,214349 | 0,663825704 | 0,097821946 | 3,95E-13 | 2,80E-11 | Up |
| Msmo1 | 8605,116594 | -0,8083412 | 0,234518035 | 2,50E-06 | 2,83E-05 | Down |
| Tma16 | 3261,515626 | 0,788995168 | 0,134415335 | 2,21E-11 | 1,03E-09 | Up |
| Tmem59l | 691,2130217 | -0,624027433 | 0,143235971 | 4,72E-07 | 6,72E-06 | Down |
| Bst2 | 8503,342442 | 0,580102821 | 0,112538302 | 1,64E-08 | 3,34E-07 | Up |
| Nwd1 | 3134,380359 | -0,675338851 | 0,309348994 | 0,000324141 | 0,002006383 | Down |
| Gm7984 | 118,695457 | 0,605896214 | 0,178941142 | 2,22E-05 | 0,000195405 | Up |
| Gm9725 | 12,45647287 | -0,818782205 | 0,712581197 | 0,001393073 | 0,007012664 | Down |
| Gm10644 | 47,0296698 | -0,728957762 | 0,325704982 | 0,000197313 | 0,001297075 | Down |
| Gm10643 | 38,12254391 | -0,672602593 | 0,294590246 | 0,000261013 | 0,001658747 | Down |
| Klf1 | 812,5738791 | -1,06030823 | 0,130450566 | 2,24E-19 | 5,67E-17 | Down |
| Heatr3 | 19515,88278 | 0,58848073 | 0,089805889 | 4,14E-12 | 2,33E-10 | Up |
| Nod2 | 319,3092233 | -0,942645829 | 0,148744768 | 1,88E-13 | 1,41E-11 | Down |

| | | | | | | |
|---|---|---|---|---|---|---|
| Drc7 | 87,03787323 | -1,331604885 | 0,40704606 | 1,97E-06 | 2,30E-05 | Down |
| Kifc3 | 1942,482142 | -0,759319704 | 0,205767383 | 1,54E-06 | 1,87E-05 | Down |
| Ces2g | 341,3859819 | 0,999987591 | 0,656590183 | 0,000422669 | 0,002513308 | Up |
| Lrrc29 | 453,0647245 | -0,734253644 | 0,089346705 | 2,73E-18 | 5,73E-16 | Down |
| Lrrc36 | 24,50060593 | -0,585939217 | 0,3267486 | 0,001394761 | 0,007016664 | Down |
| Ctrl | 102,9559872 | 0,692714779 | 0,363279856 | 0,000539803 | 0,003098448 | Up |
| Cirh1a | 18575,52838 | 0,726133209 | 0,089891772 | 9,92E-18 | 1,90E-15 | Up |
| Nip7 | 12254,68966 | 0,709340473 | 0,107260324 | 6,73E-13 | 4,56E-11 | Up |
| Hp | 192,1068908 | 0,718015375 | 0,579525226 | 0,001735287 | 0,008464238 | Up |
| Ldhd | 15,42549047 | -1,556683317 | 0,731485098 | 4,93E-05 | 0,000391046 | Down |
| Cotl1 | 639,3924189 | 1,080522425 | 0,258734648 | 3,58E-08 | 6,65E-07 | Up |
| Gm27011 | 34,78370778 | -1,017590313 | 0,388105649 | 1,93E-05 | 0,000172782 | Down |
| Gm17709 | 80,27379135 | 0,725438935 | 0,274902241 | 6,91E-05 | 0,000522231 | Up |
| Zfpm1 | 21567,51357 | -0,935379104 | 0,124044663 | 3,09E-17 | 5,09E-15 | Down |
| Gm45353 | 43,3470747 | -1,728787981 | 0,378432052 | 3,26E-08 | 6,10E-07 | Down |
| Cyba | 21002,84296 | 0,689922917 | 0,142741682 | 2,50E-08 | 4,83E-07 | Up |
| Mvd | 5967,227248 | -0,967321407 | 0,207601931 | 3,70E-09 | 9,24E-08 | Down |
| Urb2 | 19357,13824 | 0,632844493 | 0,090610967 | 1,44E-13 | 1,10E-11 | Up |
| Kcnk1 | 553,4308129 | 0,917768855 | 0,166122552 | 3,87E-11 | 1,68E-09 | Up |
| Mmp8 | 19,57039496 | 0,616946486 | 0,778924785 | 0,003996922 | 0,016968023 | Up |
| Maml2 | 1577,826102 | -0,893501797 | 0,160889674 | 3,99E-11 | 1,71E-09 | Down |
| Ppan | 19983,65447 | 0,586519488 | 0,15966749 | 9,99E-06 | 9,61E-05 | Up |
| Ldlr | 23218,52126 | -0,641739048 | 0,195826859 | 2,20E-05 | 0,000194498 | Down |
| RP23-187B11,11 | 34,10967491 | -0,650678119 | 0,401187735 | 0,00123278 | 0,006317043 | Down |
| Pknox2 | 56,51895352 | 3,12197683 | 0,596301709 | 2,39E-09 | 6,26E-08 | Up |
| Robo3 | 29,5167106 | 3,286370591 | 0,546617714 | 3,71E-11 | 1,62E-09 | Up |
| Nrgn | 5176,970255 | -0,916830453 | 0,182431144 | 6,77E-10 | 2,09E-08 | Down |
| Hspa8 | 409222,1768 | 0,788845289 | 0,077302275 | 9,49E-27 | 7,10E-24 | Up |
| RP24-175C20,19 | 351,9516438 | -0,632858576 | 0,156635188 | 1,59E-06 | 1,91E-05 | Down |
| RP24-282C4,4 | 30,73858901 | -0,642815949 | 0,464681186 | 0,001946987 | 0,009300561 | Down |
| 4833427G06Rik | 25,96191716 | 0,623417664 | 0,423029034 | 0,001872852 | 0,009012167 | Up |
| Dmxl2 | 12820,73825 | 0,703919989 | 0,155482059 | 8,80E-08 | 1,48E-06 | Up |
| Idh3a | 38695,69638 | 0,583823659 | 0,104092889 | 1,38E-09 | 3,87E-08 | Up |
| Pstpip1 | 6890,017773 | 0,637051956 | 0,109313267 | 2,36E-10 | 8,25E-09 | Up |
| Cspg4 | 24,4024247 | 0,597751886 | 0,479960509 | 0,003039109 | 0,013499898 | Up |
| Stra6 | 22,08416683 | 1,765069076 | 1,341101779 | 0,000177581 | 0,001186595 | Up |
| Cd276 | 489,4175603 | 1,660420257 | 0,272488767 | 1,79E-11 | 8,62E-10 | Up |
| Gramd2 | 15,68022229 | 1,322310048 | 0,692284032 | 0,000101764 | 0,000726305 | Up |
| RP23-9A12,4 | 5,547729581 | 1,05948287 | 1,317773332 | 0,000818375 | 0,004435025 | Up |
| RP23-116C19,1 | 234,4387363 | 0,680503461 | 0,315096472 | 0,000330041 | 0,002034885 | Up |
| Hacd3 | 17050,06922 | 0,599534615 | 0,106521658 | 1,08E-09 | 3,15E-08 | Up |
| Car12 | 3838,001602 | 1,236450098 | 0,116830898 | 1,42E-28 | 1,24E-25 | Up |
| Gm10642 | 31,73537749 | -1,582772275 | 0,56445756 | 9,01E-06 | 8,81E-05 | Down |
| Ccpg1os | 184,6730959 | -0,68647635 | 0,131664809 | 3,85E-09 | 9,52E-08 | Down |
| Sh3bgrl2 | 22,9565093 | 2,412334759 | 0,651767859 | 7,97E-07 | 1,05E-05 | Up |
| Bcl2a1d | 151,2978307 | 1,42697283 | 0,294087651 | 4,84E-09 | 1,16E-07 | Up |
| Bcl2a1a | 3219,401871 | 1,46108583 | 0,352121875 | 1,02E-07 | 1,70E-06 | Up |
| Bcl2a1b | 1443,482025 | 1,194873336 | 0,270754413 | 1,48E-08 | 3,04E-07 | Up |
| Pcolce2 | 191,0624723 | -1,143446105 | 0,27052784 | 3,09E-08 | 5,81E-07 | Down |
| Pls1 | 3170,196103 | -2,139890873 | 0,200812132 | 2,95E-28 | 2,44E-25 | Down |
| 1700065D16Rik | 152,2227638 | -1,955248082 | 0,225456369 | 1,00E-19 | 2,77E-17 | Down |

| | | | | | | |
|---|---|---|---|---|---|---|
| Rasa2 | 3208,881562 | 0,694722715 | 0,120347722 | 1,58E-10 | 5,86E-09 | Up |
| Ryk | 47,51712129 | -0,940667062 | 0,396005917 | 5,09E-05 | 0,000403404 | Down |
| Bfsp2 | 56,3096011 | 2,230631004 | 0,32445044 | 9,82E-14 | 7,79E-12 | Up |
| Tlr9 | 120,2603843 | 1,766421486 | 0,562552353 | 3,68E-06 | 3,99E-05 | Up |
| Alas1 | 5364,66239 | 1,565932931 | 0,11921477 | 9,90E-41 | 5,19E-37 | Up |
| Dusp7 | 2854,224814 | 0,774112663 | 0,086808551 | 3,15E-21 | 1,21E-18 | Up |
| Acy1 | 10122,32125 | 0,845310731 | 0,079862289 | 5,88E-29 | 5,44E-26 | Up |
| Rrp9 | 13275,934 | 0,927818396 | 0,102025826 | 5,37E-23 | 2,72E-20 | Up |
| Mapkapk3 | 18249,63982 | 1,308741825 | 0,179297324 | 1,42E-15 | 1,68E-13 | Up |
| Gm17040 | 22,76947498 | 0,996482902 | 0,624806531 | 0,000362803 | 0,002206582 | Up |
| Cish | 14896,70043 | 2,174768872 | 0,151822325 | 2,65E-48 | 2,08E-44 | Up |
| Hemk1 | 5617,126081 | 0,798213292 | 0,157593181 | 1,84E-09 | 4,98E-08 | Up |
| 6430571L13Rik | 340,745674 | 0,950048295 | 0,212603803 | 1,05E-08 | 2,27E-07 | Up |
| Hyal2 | 3828,837013 | 0,806011395 | 0,095028348 | 8,14E-20 | 2,29E-17 | Up |
| Ifrd2 | 22895,94146 | 0,600192043 | 0,075799116 | 1,79E-16 | 2,44E-14 | Up |
| Gm38150 | 138,0026428 | 0,773746907 | 0,32498883 | 0,000102748 | 0,000732995 | Up |
| Bsn | 1393,889029 | 0,686248336 | 0,09872599 | 9,35E-14 | 7,50E-12 | Up |
| Nicn1 | 1482,958208 | -0,872696497 | 0,110286625 | 3,17E-18 | 6,47E-16 | Down |
| Gm42762 | 74,11714831 | -0,606657552 | 0,297683182 | 0,000746713 | 0,004096835 | Down |
| Arpp21 | 45,49730575 | -1,137938574 | 0,320842563 | 5,83E-07 | 8,04E-06 | Down |
| Ccr4 | 3470,273775 | 0,698597174 | 0,166886511 | 4,17E-07 | 6,00E-06 | Up |
| Ttc21a | 268,5701923 | 0,605440239 | 0,161832749 | 6,53E-06 | 6,67E-05 | Up |
| Ano10 | 47,04066066 | 0,9978569 | 0,599508423 | 0,000307651 | 0,001919432 | Up |
| Ccr1 | 520,2067066 | 3,707985651 | 0,589411671 | 7,40E-12 | 3,81E-10 | Up |
| Ccr2 | 80,94767328 | 3,460130095 | 0,734796752 | 2,69E-08 | 5,16E-07 | Up |
| Ccr5 | 30,1303597 | 3,065938875 | 0,707074995 | 1,10E-07 | 1,82E-06 | Up |
| Utrn | 2061,625378 | -0,616795651 | 0,234280398 | 0,000176629 | 0,001181735 | Down |
| Ccdc28a | 620,3424095 | -0,665653679 | 0,12029179 | 8,99E-10 | 2,69E-08 | Down |
| Nhsl1 | 1298,074484 | 0,620027305 | 0,130427672 | 8,23E-08 | 1,39E-06 | Up |
| Arfgef3 | 2395,371319 | -0,606756792 | 0,135434454 | 3,28E-07 | 4,86E-06 | Down |
| 4933406P04Rik | 52,5340851 | -0,747151751 | 0,269899336 | 4,02E-05 | 0,00032738 | Down |
| Raet1d | 35,58970094 | 4,464212643 | 0,82625368 | 5,29E-10 | 1,69E-08 | Up |
| Gm10825 | 50,07323976 | 4,713597436 | 0,647692 | 3,12E-15 | 3,46E-13 | Up |
| Cdk19 | 4869,431134 | -0,816313399 | 0,113799209 | 2,33E-15 | 2,70E-13 | Down |
| Ak9 | 6,828321323 | -0,654748508 | 0,836263666 | 0,003383564 | 0,01472614 | Down |
| Smpd2 | 190,4245367 | -0,740982389 | 0,173889705 | 1,82E-07 | 2,88E-06 | Down |
| F930017D23Rik | 36,30408448 | -1,306684503 | 0,441899269 | 5,48E-06 | 5,65E-05 | Down |
| Lilrb4a | 4829,529236 | 4,599051805 | 0,567728835 | 7,04E-18 | 1,38E-15 | Up |
| Oit3 | 35,99690478 | 0,964546824 | 0,535106903 | 0,000237233 | 0,001521777 | Up |
| Rpl27a-ps1 | 58,33750587 | -0,936113965 | 0,260647951 | 6,35E-07 | 8,69E-06 | Down |
| Ppa1 | 20164,62417 | 0,674820094 | 0,120576492 | 5,63E-10 | 1,77E-08 | Up |
| Rhobtb1 | 961,7149619 | 0,677792701 | 0,092401208 | 6,62E-15 | 6,71E-13 | Up |
| Adora2a | 4036,259336 | -0,660800971 | 0,130087564 | 1,06E-08 | 2,28E-07 | Down |
| Ggt1 | 87,28368687 | 0,638338455 | 0,305194008 | 0,00052258 | 0,003020547 | Up |
| Ggt5 | 50,42020708 | -0,595577047 | 0,286336222 | 0,000749745 | 0,004109867 | Down |
| Lss | 8927,411891 | -0,99699822 | 0,207854254 | 1,65E-09 | 4,51E-08 | Down |
| Slc19a1 | 5856,627741 | 0,886795942 | 0,118050599 | 6,37E-17 | 9,53E-15 | Up |
| Trpm2 | 18,47986833 | 2,267661895 | 0,565016123 | 3,01E-07 | 4,51E-06 | Up |
| Dnmt3l | 1127,374534 | 1,384290584 | 0,213758422 | 6,31E-13 | 4,29E-11 | Up |
| Pdxk | 17750,85048 | 0,614465151 | 0,102760299 | 1,22E-10 | 4,65E-09 | Up |
| Plppr3 | 5055,720158 | 1,138906411 | 0,217222305 | 1,73E-10 | 6,32E-09 | Up |

| | | | | | | |
|---|---|---|---|---|---|---|
| Prtn3 | 25868,27904 | 0,831362799 | 0,235629584 | 1,52E-06 | 1,85E-05 | Up |
| Apc2 | 208,6627662 | -0,728895563 | 0,237142588 | 1,81E-05 | 0,000163575 | Down |
| Reep6 | 194,4615902 | 1,245235556 | 0,176471586 | 4,37E-15 | 4,67E-13 | Up |
| Adamtsl5 | 24,20268873 | -0,679795873 | 0,461504415 | 0,001382225 | 0,006969216 | Down |
| 4930442H23Rik | 28,96264306 | -0,635568676 | 0,36051761 | 0,001038892 | 0,005426128 | Down |
| Tjp3 | 623,5626684 | -1,006643458 | 0,298468197 | 1,27E-06 | 1,57E-05 | Down |
| Smim24 | 924,2397926 | -0,668162812 | 0,19909776 | 1,30E-05 | 0,000122279 | Down |
| Ncln | 25067,2087 | 0,624395601 | 0,053161261 | 5,07E-33 | 8,85E-30 | Up |
| RP23-132J19,1 | 72,17940298 | 2,570273651 | 0,391630502 | 7,65E-13 | 5,12E-11 | Up |
| A230060F14Rik | 81,9251504 | -1,155921099 | 0,337903058 | 9,71E-07 | 1,25E-05 | Down |
| Fbxo7 | 10180,18259 | -1,007989305 | 0,198570087 | 3,51E-10 | 1,16E-08 | Down |
| Syn3 | 40,9513894 | -1,811220373 | 0,396559244 | 3,54E-08 | 6,58E-07 | Down |
| Gm26546 | 34,28311744 | -1,22497728 | 0,408725178 | 4,68E-06 | 4,90E-05 | Down |
| Utp20 | 17353,32244 | 0,646806031 | 0,16573745 | 2,34E-06 | 2,67E-05 | Up |
| Gas2l3 | 16653,24652 | -0,696086292 | 0,188152227 | 2,92E-06 | 3,23E-05 | Down |
| RP23-337J18,2 | 10,41117979 | 0,754756852 | 0,639486086 | 0,001650859 | 0,00811285 | Up |
| RP23-381H23,1 | 442,2506705 | -1,095674608 | 0,185809777 | 3,07E-12 | 1,78E-10 | Down |
| Gm15915 | 150,4032388 | 1,264148349 | 0,211360513 | 5,35E-12 | 2,86E-10 | Up |
| Gm3571 | 49,55293037 | -1,196165178 | 0,33172644 | 4,88E-07 | 6,92E-06 | Down |
| Plxnc1 | 718,5796189 | 1,362853182 | 0,188639031 | 3,81E-15 | 4,16E-13 | Up |
| Dusp6 | 5131,240224 | 0,7073029 | 0,147300684 | 2,33E-08 | 4,54E-07 | Up |
| Ctdsp2 | 16834,36362 | -0,612063332 | 0,12682529 | 6,32E-08 | 1,10E-06 | Down |
| Avil | 247,8839592 | 0,667575826 | 0,758464831 | 0,003164548 | 0,01394673 | Up |
| Kif5a | 352,4981487 | -0,724883454 | 0,186272003 | 1,02E-06 | 1,30E-05 | Down |
| Inhbe | 22,80795408 | -1,452707098 | 0,761609136 | 8,94E-05 | 0,000651048 | Down |
| Lrp1 | 58,39551314 | 1,717183321 | 0,315494761 | 6,39E-10 | 1,98E-08 | Up |
| Stat2 | 5653,784203 | -0,72067497 | 0,134008929 | 1,01E-09 | 2,97E-08 | Down |
| Pa2g4 | 86246,25837 | 0,612817893 | 0,093145566 | 2,81E-12 | 1,65E-10 | Up |
| Ccdc157 | 239,7664646 | -0,835544262 | 0,193853448 | 5,13E-08 | 9,19E-07 | Down |
| Gatsl3 | 234,9136018 | 2,008787764 | 0,230193399 | 5,72E-20 | 1,63E-17 | Up |
| Lif | 95,69670836 | 0,624477516 | 0,272862593 | 0,00037742 | 0,002281356 | Up |
| Gm11963 | 137,8353194 | 1,272294076 | 0,254950469 | 1,25E-09 | 3,57E-08 | Up |
| Xbp1 | 46769,59394 | 0,843153972 | 0,10884611 | 1,89E-17 | 3,27E-15 | Up |
| Polm | 1099,511717 | -1,302733697 | 0,160303982 | 2,54E-18 | 5,55E-16 | Down |
| H2afv | 5771,250671 | -0,663804048 | 0,140269063 | 5,59E-08 | 9,87E-07 | Down |
| Upp1 | 1663,281966 | -0,974123993 | 0,210031152 | 4,04E-09 | 9,88E-08 | Down |
| Abca13 | 258,6533689 | 1,151329593 | 0,286807529 | 8,20E-08 | 1,39E-06 | Up |
| Ddc | 1088,133481 | 0,663933552 | 0,131764663 | 1,26E-08 | 2,64E-07 | Up |
| Grb10 | 260,5275999 | 0,701270692 | 0,534434023 | 0,001653426 | 0,008120386 | Up |
| Plek | 48649,19621 | -0,790730453 | 0,16089706 | 4,43E-09 | 1,08E-07 | Down |
| Pno1 | 8850,649921 | 0,772377144 | 0,101694573 | 2,06E-16 | 2,77E-14 | Up |
| Wdpcp | 417,7205719 | -1,047699029 | 0,164331596 | 1,18E-13 | 9,19E-12 | Down |
| Slit3 | 276,9446631 | -0,698965001 | 0,38717568 | 0,000648282 | 0,003621896 | Down |
| Olfr56 | 109,8057391 | 1,2792041 | 0,265763019 | 3,08E-09 | 7,84E-08 | Up |
| Ltc4s | 612,1824125 | 0,678441707 | 0,153181719 | 1,87E-07 | 2,96E-06 | Up |
| Acsl6 | 220,3004438 | -1,136581011 | 0,229541198 | 8,31E-10 | 2,50E-08 | Down |
| Slc36a1os | 60,40644346 | -0,727685031 | 0,270260262 | 5,83E-05 | 0,000453096 | Down |
| Gm12247 | 49,05827899 | -1,626054311 | 0,310373053 | 1,48E-09 | 4,09E-08 | Down |
| Nlrp3 | 104,4376434 | 2,182782546 | 0,510116456 | 1,19E-07 | 1,96E-06 | Up |
| Med9os | 88,52164496 | -0,699839607 | 0,227308425 | 2,32E-05 | 0,000202557 | Down |
| Shmt1 | 21665,39839 | 0,626080063 | 0,138630852 | 2,30E-07 | 3,56E-06 | Up |

| | | | | | | |
|---|---|---|---|---|---|---|
| Natd1 | 989,8476047 | -0,819051254 | 0,160683113 | 1,19E-09 | 3,42E-08 | Down |
| Grap | 708,8283831 | 0,632002337 | 0,174214307 | 7,51E-06 | 7,53E-05 | Up |
| Pik3r5 | 3647,102947 | 1,009047574 | 0,147827963 | 4,97E-15 | 5,24E-13 | Up |
| Cd68 | 1730,234601 | 1,492008135 | 0,239367823 | 4,92E-12 | 2,68E-10 | Up |
| Nlgn2 | 91,9016118 | -0,594058454 | 0,220399582 | 0,00018555 | 0,001230946 | Down |
| Arrb2 | 15836,62708 | 0,683308652 | 0,088831982 | 4,11E-16 | 5,21E-14 | Up |
| Gp1ba | 496,4015227 | -1,034851653 | 0,204276251 | 3,65E-10 | 1,20E-08 | Down |
| Aipl1 | 422,8715118 | -0,918949247 | 0,221672736 | 5,90E-08 | 1,03E-06 | Down |
| Fam64a | 7453,227251 | -0,696599523 | 0,165780847 | 3,99E-07 | 5,79E-06 | Down |
| Mybbp1a | 122111,9091 | 0,60092856 | 0,126136275 | 9,45E-08 | 1,58E-06 | Up |
| Mettl16 | 15985,67294 | 0,706333326 | 0,123325391 | 1,74E-10 | 6,35E-09 | Up |
| Tsr1 | 12877,14543 | 0,604999161 | 0,093140799 | 5,23E-12 | 2,82E-10 | Up |
| Serpinf1 | 509,0275843 | -0,829451829 | 0,316232986 | 3,69E-05 | 0,000303936 | Down |
| Gm12341 | 96,30970412 | 0,824679396 | 0,211846971 | 3,64E-07 | 5,33E-06 | Up |
| Rab11fip4 | 393,4511642 | 1,09063415 | 0,307732993 | 5,81E-07 | 8,02E-06 | Up |
| Gm11417 | 102,8043685 | -0,801531738 | 0,206502074 | 4,72E-07 | 6,72E-06 | Down |
| Slfn2 | 1045,664031 | 2,861203358 | 0,339959609 | 7,51E-19 | 1,77E-16 | Up |
| Slfn1 | 8,722753994 | 2,471468464 | 1,828351646 | 4,86E-05 | 0,000386345 | Up |
| Slfn14 | 100,757885 | -0,605794716 | 0,365653469 | 0,0015369 | 0,007612329 | Down |
| Ccl6 | 1119,856899 | 1,994339611 | 1,050343337 | 5,62E-05 | 0,000438715 | Up |
| Ccl3 | 441,8749966 | 1,48408231 | 0,566821992 | 1,50E-05 | 0,000138602 | Up |
| Ccl4 | 335,2517241 | 1,918397743 | 0,368225223 | 2,26E-09 | 5,96E-08 | Up |
| Wfdc21 | 16,19669521 | 2,158417222 | 0,984796865 | 2,86E-05 | 0,000242425 | Up |
| Myo19 | 4985,160782 | 0,776519481 | 0,163096566 | 1,15E-08 | 2,44E-07 | Up |
| Sep 04 | 11,63961559 | -0,691771315 | 0,509202743 | 0,00159557 | 0,00787009 | Down |
| Utp18 | 8967,047736 | 0,585431822 | 0,084553033 | 3,42E-13 | 2,43E-11 | Up |
| Acsf2 | 171,4211119 | -0,971259226 | 0,232161041 | 3,92E-08 | 7,21E-07 | Down |
| Dlx4os | 173,3534353 | 0,648598671 | 0,173951777 | 4,39E-06 | 4,65E-05 | Up |
| Gm11515 | 57,49730154 | 0,74213741 | 0,294943246 | 8,63E-05 | 0,00063288 | Up |
| Lrrc46 | 65,64838443 | -0,743615247 | 0,223995572 | 6,98E-06 | 7,07E-05 | Down |
| Tbkbp1 | 9540,603635 | 0,73682426 | 0,101864635 | 5,71E-15 | 5,87E-13 | Up |
| Plxdc1 | 19,07902451 | -1,202460392 | 0,568282858 | 6,89E-05 | 0,000521625 | Down |
| Rara | 1249,45101 | 0,955437509 | 0,137515832 | 2,41E-15 | 2,74E-13 | Up |
| Igfbp4 | 49594,14105 | 0,585635722 | 0,113843744 | 1,66E-08 | 3,36E-07 | Up |
| Tns4 | 359,6303733 | 0,909312733 | 0,375151768 | 4,85E-05 | 0,000385782 | Up |
| Plekhh3 | 34,01354151 | 1,195104425 | 1,102964222 | 0,00060365 | 0,003406448 | Up |
| Etv4 | 842,4524741 | 1,652729714 | 0,181928045 | 3,75E-21 | 1,40E-18 | Up |
| Ccdc103 | 19,73196061 | -0,812292385 | 0,465228073 | 0,00043598 | 0,002581661 | Down |
| Acbd4 | 2574,740235 | -0,670789865 | 0,125102441 | 2,15E-09 | 5,72E-08 | Down |
| Itgb3 | 28796,22896 | -1,404650824 | 0,12851458 | 2,04E-29 | 2,14E-26 | Down |
| Ftsj3 | 39137,72399 | 0,759879518 | 0,088102615 | 5,47E-20 | 1,59E-17 | Up |
| Cd79b | 160,9966712 | -0,824094846 | 0,366168618 | 0,000113501 | 0,000800258 | Down |
| Milr1 | 749,3495395 | 2,883342274 | 0,244973577 | 1,24E-33 | 2,44E-30 | Up |
| 1810010H24Rik | 75,91311996 | -0,673531479 | 0,302368249 | 0,000295913 | 0,001855031 | Down |
| Gpr142 | 10,70709448 | 0,637032545 | 0,760686153 | 0,003649455 | 0,015696454 | Up |
| Cd300ld | 194,4014277 | 0,774761257 | 0,152209149 | 2,20E-09 | 5,84E-08 | Up |
| Cd300ld2 | 45,67226224 | 0,962895109 | 0,369557634 | 2,31E-05 | 0,000201391 | Up |
| Rab37 | 416,2420846 | -0,775761589 | 0,209543822 | 1,25E-06 | 1,55E-05 | Down |
| Cd300lf | 3294,058118 | 0,904080003 | 0,110997764 | 3,10E-19 | 7,73E-17 | Up |
| Smim5 | 8823,774637 | -0,662711996 | 0,130422204 | 1,03E-08 | 2,22E-07 | Down |
| Smim6 | 358,5801824 | -0,715347828 | 0,200050974 | 3,73E-06 | 4,03E-05 | Down |

| | | | | | | |
|---|---|---|---|---|---|---|
| Itgb4 | 428,4791319 | 0,695184499 | 0,201060436 | 7,00E-06 | 7,08E-05 | Up |
| Wbp2 | 31706,53821 | -0,587398693 | 0,104350929 | 1,20E-09 | 3,42E-08 | Down |
| Trim47 | 130,4935779 | 0,615598105 | 0,180917198 | 1,91E-05 | 0,000171085 | Up |
| Evpl | 79,20208204 | -0,93343511 | 0,378043153 | 3,89E-05 | 0,000318743 | Down |
| Ubald2 | 7913,679561 | -0,586454592 | 0,120932421 | 7,16E-08 | 1,23E-06 | Down |
| Gm44935 | 184,9425406 | -0,626863269 | 0,151377815 | 1,13E-06 | 1,43E-05 | Down |
| Ccdc137 | 5048,72837 | 0,671345578 | 0,2389526 | 6,71E-05 | 0,000509697 | Up |
| Narf | 2989,659917 | -0,903613805 | 0,184689428 | 1,53E-09 | 4,23E-08 | Down |
| Gm12589 | 33,59654821 | 0,73614104 | 0,40480211 | 0,000516592 | 0,002991436 | Up |
| Lpin1 | 5191,723591 | -1,153549591 | 0,233551821 | 9,43E-10 | 2,80E-08 | Down |
| Greb1 | 5300,365898 | -4,153028659 | 0,523562267 | 5,40E-17 | 8,38E-15 | Down |
| Nol10 | 12006,45166 | 0,611520847 | 0,091084322 | 1,15E-12 | 7,47E-11 | Up |
| Odc1 | 90651,84054 | 0,664935727 | 0,113665901 | 1,49E-10 | 5,58E-09 | Up |
| Rrm2 | 48359,56412 | 0,597988804 | 0,110536824 | 3,67E-09 | 9,17E-08 | Up |
| Bcap29 | 12417,3914 | 0,650168538 | 0,106232214 | 3,52E-11 | 1,57E-09 | Up |
| Dus4l | 2202,641229 | 0,700195859 | 0,1173434 | 4,64E-11 | 1,95E-09 | Up |
| Hdac9 | 275,840004 | 1,551238335 | 0,299239188 | 1,59E-09 | 4,37E-08 | Up |
| Arl4a | 2052,309251 | -0,775549247 | 0,084892569 | 4,19E-22 | 1,83E-19 | Down |
| Mipol1 | 719,4501988 | 0,610913066 | 0,134931951 | 2,55E-07 | 3,91E-06 | Up |
| Trim9 | 416,7026228 | 0,614786341 | 0,115981428 | 5,61E-09 | 1,31E-07 | Up |
| Timm9 | 7560,531239 | 0,646067118 | 0,114600947 | 6,36E-10 | 1,98E-08 | Up |
| Rhoj | 8706,867025 | 3,102059228 | 0,452308971 | 1,48E-13 | 1,12E-11 | Up |
| 2310015A10Rik | 599,3068281 | -0,912349903 | 0,110968521 | 1,48E-19 | 3,87E-17 | Down |
| Ltbp2 | 184,5277466 | -0,587691548 | 0,24388442 | 0,000393867 | 0,00236531 | Down |
| Tgfb3 | 214,763744 | -3,13418151 | 0,314161849 | 5,63E-25 | 3,16E-22 | Down |
| 9030617O03Rik | 9762,931775 | 0,610153682 | 0,087521397 | 1,98E-13 | 1,47E-11 | Up |
| Serpina3f | 222,9316859 | 1,371778017 | 0,398166189 | 1,13E-06 | 1,43E-05 | Up |
| Exoc3l4 | 11,54447877 | -0,67166553 | 0,556449083 | 0,002217035 | 0,010407086 | Down |
| Trmt61a | 4466,011243 | 1,064471077 | 0,127673138 | 3,89E-20 | 1,20E-17 | Up |
| Pld4 | 17833,91103 | 0,945790526 | 0,176304953 | 8,30E-11 | 3,28E-09 | Up |
| Ahnak2 | 117,9360797 | 0,58774919 | 0,285492059 | 0,000834494 | 0,004511492 | Up |
| Ighm | 1457,829526 | -1,621986569 | 0,247779458 | 1,10E-12 | 7,23E-11 | Down |
| Akr1c18 | 358,3630148 | 0,742167651 | 0,654159241 | 0,00184735 | 0,008918447 | Up |
| Idi1 | 7542,710232 | -1,358842481 | 0,200550644 | 7,79E-14 | 6,38E-12 | Down |
| Heatr1 | 24524,10792 | 0,598790947 | 0,120071592 | 3,27E-08 | 6,11E-07 | Up |
| Lyst | 43,14978485 | 2,149943774 | 0,717197771 | 5,09E-06 | 5,30E-05 | Up |
| Zfp184 | 127,6383561 | -1,114973397 | 0,23210354 | 1,69E-09 | 4,61E-08 | Down |
| Mboat1 | 3788,958972 | 0,686950959 | 0,219082647 | 2,20E-05 | 0,000194248 | Up |
| Serpinb6b | 1801,582066 | -1,143263942 | 0,149356674 | 1,93E-17 | 3,30E-15 | Down |
| Serpinb9 | 774,4030164 | -0,663972159 | 0,169895839 | 1,90E-06 | 2,23E-05 | Down |
| Serpinb9g | 19,49516199 | 1,410929761 | 0,882195456 | 0,000181809 | 0,001212781 | Up |
| Rpp40 | 2175,583782 | 0,87022243 | 0,093234069 | 1,19E-23 | 6,21E-21 | Up |
| Eef1e1 | 7193,167973 | 0,761637309 | 0,100607845 | 3,01E-16 | 3,91E-14 | Up |
| Gmpr | 95,63828154 | 0,946854642 | 0,272267345 | 9,76E-07 | 1,25E-05 | Up |
| Nol8 | 8292,613356 | 0,698735875 | 0,094114472 | 2,53E-15 | 2,84E-13 | Up |
| Simc1 | 1692,816062 | 0,624372888 | 0,108888255 | 4,65E-10 | 1,50E-08 | Up |
| Nop16 | 15847,78597 | 0,680244221 | 0,103831027 | 1,54E-12 | 9,59E-11 | Up |
| Hk3 | 26800,88562 | 0,958296625 | 0,218722536 | 1,57E-08 | 3,21E-07 | Up |
| Prr7 | 516,0034562 | 0,653819584 | 0,124834428 | 5,12E-09 | 1,22E-07 | Up |
| Fbp2 | 13397,45589 | 1,476741124 | 0,191899765 | 2,58E-16 | 3,41E-14 | Up |
| Fbp1 | 506,5409413 | 2,170207375 | 0,311070534 | 5,07E-14 | 4,40E-12 | Up |

| | | | | | | |
|---|---|---|---|---|---|---|
| Fancc | 5048,413678 | -0,871862003 | 0,155956713 | 3,98E-11 | 1,71E-09 | Down |
| Nsun2 | 57730,03831 | 0,595864322 | 0,097184288 | 5,73E-11 | 2,37E-09 | Up |
| Trip13 | 10381,32846 | 0,625347843 | 0,092104251 | 6,02E-13 | 4,15E-11 | Up |
| Polr3g | 3044,392191 | 0,679810096 | 0,093208703 | 8,76E-15 | 8,66E-13 | Up |
| Mblac2 | 543,9052679 | -0,762996968 | 0,13625533 | 1,59E-10 | 5,86E-09 | Down |
| Gm4117 | 181,6761914 | 0,95013526 | 0,211898816 | 9,74E-09 | 2,12E-07 | Up |
| Hmgcr | 33087,04773 | -0,77004783 | 0,138003296 | 1,60E-10 | 5,90E-09 | Down |
| Dimt1 | 2356,402769 | 0,740739412 | 0,148205487 | 5,63E-09 | 1,32E-07 | Up |
| Emb | 15483,94736 | 1,752362639 | 0,159495226 | 1,61E-29 | 1,81E-26 | Up |
| Hmgcs1 | 16890,43542 | -0,907392102 | 0,211014154 | 3,03E-08 | 5,71E-07 | Down |
| Ndst2 | 7587,748567 | -0,605826271 | 0,104705526 | 4,14E-10 | 1,35E-08 | Down |
| Chdh | 23,90098117 | 0,634319445 | 0,66311766 | 0,003474768 | 0,015068832 | Up |
| Gnl3 | 16777,88997 | 0,726365123 | 0,161459189 | 7,55E-08 | 1,29E-06 | Up |
| Sh3bp5 | 733,0241111 | 0,919254946 | 0,238128847 | 2,16E-07 | 3,37E-06 | Up |
| Eaf1 | 14776,4521 | 0,655761148 | 0,101887161 | 4,48E-12 | 2,48E-10 | Up |
| 1810011H11Rik | 2581,982718 | 0,777438079 | 0,122397086 | 1,29E-12 | 8,17E-11 | Up |
| Txndc16 | 11,87922943 | -1,210015871 | 0,816238681 | 0,000308446 | 0,001923627 | Down |
| Cdkn3 | 756,721289 | -0,749215355 | 0,137757495 | 4,81E-10 | 1,55E-08 | Down |
| Ccnb1ip1 | 3382,037082 | 1,462720883 | 0,257965981 | 1,01E-10 | 3,91E-09 | Up |
| Gm26782 | 363,4083126 | 0,586237851 | 0,154362987 | 6,37E-06 | 6,53E-05 | Up |
| Rnase10 | 8,856081127 | 0,769224434 | 1,110732133 | 0,001884783 | 0,009063486 | Up |
| Trac | 148,0237605 | -2,331872015 | 0,321323404 | 5,98E-15 | 6,10E-13 | Down |
| Abhd4 | 11072,85963 | -0,681433899 | 0,109540591 | 1,27E-11 | 6,26E-10 | Down |
| Slc7a7 | 7431,349218 | -1,54007734 | 0,173265815 | 1,93E-20 | 6,73E-18 | Down |
| Cebpe | 1625,059056 | 0,747125132 | 0,369056801 | 0,000301461 | 0,00188455 | Up |
| Pck2 | 22785,40009 | -0,710009308 | 0,215505727 | 1,05E-05 | 0,000100226 | Down |
| Cideb | 424,6151422 | -0,74696366 | 0,196705868 | 1,16E-06 | 1,45E-05 | Down |
| Ltb4r2 | 305,3376719 | -0,586160266 | 0,247436604 | 0,000438114 | 0,00258876 | Down |
| Ripk3 | 124,2185648 | 2,694558174 | 0,347727445 | 1,50E-16 | 2,08E-14 | Up |
| Mcpt1 | 70,48400238 | -1,392795485 | 0,499086739 | 9,29E-06 | 9,05E-05 | Down |
| Mcpt2 | 338,2446608 | -1,599282745 | 0,154935917 | 2,28E-26 | 1,55E-23 | Down |
| Mcpt4 | 8,384654525 | -0,754508146 | 0,691324501 | 0,001872465 | 0,009012167 | Down |
| Gzmb | 5878,048741 | -1,63331767 | 0,38893745 | 1,18E-07 | 1,94E-06 | Down |
| Cryl1 | 1527,099161 | -0,600196018 | 0,075001317 | 9,08E-17 | 1,33E-14 | Down |
| Fgf9 | 1434,587232 | 0,670815789 | 0,079169309 | 8,44E-19 | 1,95E-16 | Up |
| Fdft1 | 7085,519987 | -0,952744042 | 0,21515283 | 1,27E-08 | 2,67E-07 | Down |
| Zfp395 | 2940,721273 | -0,696610851 | 0,12516922 | 5,02E-10 | 1,61E-08 | Down |
| Dpysl2 | 3918,68976 | -0,730377056 | 0,094463723 | 1,46E-16 | 2,07E-14 | Down |
| Entpd4 | 6,820500441 | 0,766815962 | 1,285869908 | 0,001125705 | 0,005829188 | Up |
| Slc39a14 | 6858,21372 | 0,632597151 | 0,107107074 | 1,56E-10 | 5,82E-09 | Up |
| Polr3d | 3655,168708 | 0,775747471 | 0,145172869 | 5,56E-10 | 1,76E-08 | Up |
| Lpar6 | 217,3238701 | 0,729664872 | 0,173947236 | 2,77E-07 | 4,20E-06 | Up |
| Dis3 | 11397,47899 | 0,584505263 | 0,107514979 | 3,56E-09 | 8,95E-08 | Up |
| Acod1 | 37,89133951 | 1,95823094 | 0,600439088 | 2,68E-06 | 3,02E-05 | Up |
| Scel | 103,6608156 | 1,080366227 | 0,303319945 | 5,41E-07 | 7,56E-06 | Up |
| Dzip1 | 190,1343369 | -0,9004671 | 0,201423162 | 1,38E-08 | 2,86E-07 | Down |
| Uggt2 | 68,39542675 | -0,614983836 | 0,232059365 | 0,00017179 | 0,001155266 | Down |
| Mtmr12 | 10351,46479 | -0,581007999 | 0,089420038 | 6,32E-12 | 3,32E-10 | Down |
| Trio | 13127,54574 | 0,588163531 | 0,073413519 | 9,30E-17 | 1,35E-14 | Up |
| Pop1 | 6346,479171 | 0,593569348 | 0,110601109 | 4,81E-09 | 1,16E-07 | Up |
| Nov | 32,77256783 | 2,862189414 | 1,117544455 | 8,99E-06 | 8,81E-05 | Up |

| | | | | | | |
|---|---|---|---|---|---|---|
| Deptor | 7619,750823 | 0,584840792 | 0,113028827 | 1,43E-08 | 2,95E-07 | Up |
| Myc | 52082,33665 | 0,690681909 | 0,089535213 | 3,12E-16 | 4,02E-14 | Up |
| St3gal1 | 9612,884407 | -0,599512573 | 0,110521898 | 3,34E-09 | 8,45E-08 | Down |
| Ly6a | 28,32201796 | 3,170025483 | 0,728713918 | 1,05E-07 | 1,75E-06 | Up |
| Ly6g | 18,38853675 | 1,361169954 | 1,032093815 | 0,000326682 | 0,002017214 | Up |
| 9030619P08Rik | 3853,176387 | -0,905548126 | 0,170857589 | 1,58E-10 | 5,86E-09 | Down |
| Tigd5 | 83,52348364 | -0,640580075 | 0,235400072 | 0,000112936 | 0,000797353 | Down |
| Bop1 | 41096,09342 | 0,595435597 | 0,083053172 | 5,52E-14 | 4,69E-12 | Up |
| Tst | 254,111652 | -0,704164408 | 0,207341708 | 7,89E-06 | 7,86E-05 | Down |
| Il2rb | 2124,240064 | 1,324819341 | 0,159512816 | 7,46E-19 | 1,77E-16 | Up |
| Mfng | 9836,646962 | 0,758256775 | 0,087054941 | 2,67E-20 | 8,75E-18 | Up |
| Gm16576 | 461,5269038 | -0,625910676 | 0,143381733 | 4,46E-07 | 6,39E-06 | Down |
| 1110025M09Rik | 53,80687314 | -0,775997722 | 0,255954791 | 1,40E-05 | 0,000130764 | Down |
| Srebf2 | 59719,23704 | -0,646126004 | 0,126815957 | 1,17E-08 | 2,48E-07 | Down |
| Odf3b | 398,2750906 | -0,591087909 | 0,136963251 | 7,76E-07 | 1,03E-05 | Down |
| Shank3 | 691,8001706 | 1,687144416 | 0,269185874 | 6,69E-12 | 3,47E-10 | Up |
| Prickle1 | 609,2121696 | -0,826851927 | 0,239475149 | 2,11E-06 | 2,44E-05 | Down |
| Pus7l | 4352,454358 | 0,716192867 | 0,090878958 | 5,70E-17 | 8,62E-15 | Up |
| 2610037D02Rik | 30,96875883 | -0,608466996 | 0,418229697 | 0,002109717 | 0,009962833 | Down |
| 1700120C14Rik | 871,9895736 | 0,765552316 | 0,304496716 | 7,43E-05 | 0,00055413 | Up |
| Mettl7a1 | 3337,095574 | -1,149310809 | 0,258376905 | 1,10E-08 | 2,35E-07 | Down |
| Galnt6 | 13781,09769 | 0,820280548 | 0,097674231 | 1,29E-19 | 3,44E-17 | Up |
| Acvrl1 | 417,3961895 | 2,023778994 | 0,475988003 | 1,28E-07 | 2,09E-06 | Up |
| Tns2 | 900,18026 | -0,589449659 | 0,141721292 | 1,51E-06 | 1,84E-05 | Down |
| Calcoco1 | 1748,419943 | -0,932722145 | 0,166745483 | 2,30E-11 | 1,07E-09 | Down |
| Gpr84 | 315,0824204 | 1,848406618 | 0,329749195 | 3,04E-10 | 1,02E-08 | Up |
| Zfp385a | 4708,771623 | -1,333852559 | 0,154046453 | 4,26E-20 | 1,29E-17 | Down |
| Nlrc3 | 795,6766529 | -0,741844684 | 0,171124291 | 1,30E-07 | 2,12E-06 | Down |
| Socs1 | 378,5095863 | 1,00015532 | 0,206391683 | 1,26E-09 | 3,58E-08 | Up |
| Rimbp3 | 353,1258945 | -0,580431729 | 0,152967073 | 6,80E-06 | 6,89E-05 | Down |
| Ccdc188 | 24,5331844 | -1,071115015 | 0,452316475 | 3,95E-05 | 0,000322127 | Down |
| Gnb1l | 2229,290271 | 0,594004242 | 0,099649548 | 1,64E-10 | 6,02E-09 | Up |
| Klhl24 | 2207,660239 | -0,596462471 | 0,13538752 | 5,02E-07 | 7,08E-06 | Down |
| Etv5 | 5632,118427 | 1,820425289 | 0,177523962 | 3,57E-26 | 2,25E-23 | Up |
| St6gal1 | 834,2815111 | 1,954504199 | 0,581355319 | 2,05E-06 | 2,38E-05 | Up |
| Gm15694 | 57,10517763 | 0,607592589 | 0,282001778 | 0,000582101 | 0,003301446 | Up |
| Hacd2 | 5490,494879 | 0,651692529 | 0,13906226 | 7,99E-08 | 1,37E-06 | Up |
| Stfa2l1 | 107,2059199 | 4,167588161 | 0,506992228 | 5,22E-18 | 1,04E-15 | Up |
| Gm5483 | 452,7039256 | 4,380211638 | 0,625664396 | 4,01E-14 | 3,54E-12 | Up |
| Gm5416 | 22,91861969 | 3,117300096 | 0,893567633 | 1,25E-06 | 1,55E-05 | Up |
| 2010005H15Rik | 294,9231517 | 4,754531032 | 0,699413767 | 8,23E-14 | 6,69E-12 | Up |
| Stfa1 | 435,2804446 | 4,227018288 | 0,722522004 | 6,78E-11 | 2,75E-09 | Up |
| BC100530 | 158,3176464 | 3,951692919 | 0,732178036 | 9,84E-10 | 2,90E-08 | Up |
| Stfa2 | 66,66232615 | 2,775680917 | 0,554023706 | 6,23E-09 | 1,44E-07 | Up |
| Stfa3 | 23,35549363 | 3,479051163 | 0,745546779 | 3,17E-08 | 5,95E-07 | Up |
| Fbxo40 | 950,1072435 | -0,644328799 | 0,096978696 | 1,31E-12 | 8,25E-11 | Down |
| Cd80 | 93,93014878 | 2,289118841 | 0,329457455 | 5,67E-14 | 4,79E-12 | Up |
| Qtrtd1 | 3150,644814 | 0,65084984 | 0,110845199 | 1,56E-10 | 5,81E-09 | Up |
| Btla | 870,6273683 | 1,097078482 | 0,221001296 | 6,91E-10 | 2,12E-08 | Up |
| 2310061N02Rik | 28,73899619 | 0,889650724 | 0,535371784 | 0,000409271 | 0,002445461 | Up |
| 2310034C09Rik | 9,532623147 | 1,275736315 | 1,033386641 | 0,000428907 | 0,002544619 | Up |

| | | | | | | |
|---|---|---|---|---|---|---|
| A930006K02Rik | 72,34929999 | -0,874569392 | 0,298697981 | 1,07E-05 | 0,000102197 | Down |
| Gart | 43874,58988 | 0,725586127 | 0,117674355 | 9,45E-12 | 4,77E-10 | Up |
| Wrb | 2437,685251 | 0,853708036 | 0,120068132 | 2,08E-15 | 2,42E-13 | Up |
| Sytl3 | 33,67731622 | 1,661655555 | 0,505814956 | 2,39E-06 | 2,72E-05 | Up |
| Igf2r | 1450,221384 | 1,52286276 | 0,545083644 | 9,27E-06 | 9,04E-05 | Up |
| Acat2 | 11616,98755 | -0,800824216 | 0,16289999 | 3,87E-09 | 9,56E-08 | Down |
| Tnfrsf12a | 1265,200175 | 0,664650936 | 0,164072569 | 1,07E-06 | 1,36E-05 | Up |
| Tbl3 | 16620,00934 | 0,637329981 | 0,098260894 | 4,01E-12 | 2,27E-10 | Up |
| Tpsab1 | 9,015693058 | 0,973806414 | 1,116545289 | 0,001145821 | 0,005923591 | Up |
| Fam195a | 3222,516533 | 0,708664746 | 0,114744167 | 1,14E-11 | 5,66E-10 | Up |
| Rab11fip3 | 208,3868459 | -1,199699241 | 0,28080502 | 2,88E-08 | 5,47E-07 | Down |
| Pacsin1 | 98,84726718 | 1,170021031 | 0,491111746 | 3,36E-05 | 0,000280388 | Up |
| 1700097N02Rik | 188,4621016 | 2,372316303 | 0,422896151 | 2,61E-10 | 8,93E-09 | Up |
| Dnah8 | 445,878478 | 2,343968459 | 0,295268578 | 3,11E-17 | 5,09E-15 | Up |
| Glp1r | 48,06580039 | 0,683326005 | 0,660867492 | 0,002627734 | 0,0119981 | Up |
| Wdr4 | 8949,026876 | 0,711967258 | 0,142687003 | 8,67E-09 | 1,92E-07 | Up |
| Pdxk-ps | 22,29220025 | -0,638829618 | 0,447118409 | 0,001847701 | 0,008918447 | Down |
| Notch3 | 180,1590764 | -0,806145063 | 0,282312807 | 2,02E-05 | 0,000180204 | Down |
| Angptl4 | 2264,452623 | 0,600016468 | 0,085418742 | 1,50E-13 | 1,13E-11 | Up |
| Tap1 | 9558,796644 | -0,670516707 | 0,131222359 | 8,09E-09 | 1,81E-07 | Down |
| Tap2 | 24397,07219 | -0,653212875 | 0,150255316 | 3,58E-07 | 5,27E-06 | Down |
| Btnl2 | 13,35834957 | 0,999479386 | 0,972462849 | 0,00099163 | 0,005213601 | Up |
| Aif1 | 158,4295468 | 0,779473836 | 0,215064919 | 1,63E-06 | 1,96E-05 | Up |
| Tnf | 575,7083511 | 1,065002471 | 0,476872475 | 5,95E-05 | 0,000461803 | Up |
| Gm20443 | 18,38509011 | -0,632283258 | 0,397799267 | 0,001458827 | 0,007287643 | Down |
| 2410017I17Rik | 6,007788131 | -1,019603384 | 0,945620707 | 0,00089328 | 0,004779974 | Down |
| Crisp1 | 11,46684021 | 0,672349639 | 1,113871491 | 0,002137191 | 0,010077434 | Up |
| Slc29a1 | 6326,353284 | 0,81634238 | 0,128455094 | 6,89E-13 | 4,65E-11 | Up |
| Bysl | 13254,14554 | 0,656304865 | 0,107975631 | 4,23E-11 | 1,80E-09 | Up |
| A530064D06Rik | 15,41685562 | 1,419515927 | 0,90560805 | 0,000191609 | 0,001263139 | Up |
| Trem1 | 776,7634029 | 1,809444049 | 0,268060314 | 3,34E-13 | 2,39E-11 | Up |
| Trem3 | 390,164595 | 2,001252109 | 0,283785242 | 3,56E-14 | 3,18E-12 | Up |
| Treml4 | 45,07301518 | 2,126051004 | 0,424364657 | 5,79E-09 | 1,35E-07 | Up |
| Trem2 | 69,74837705 | 3,950684104 | 0,78788726 | 6,01E-09 | 1,39E-07 | Up |
| Apobec2 | 31,47280923 | -0,930350155 | 0,480838897 | 0,000184161 | 0,001222761 | Down |
| Stap2 | 1615,345631 | 1,220147705 | 0,100135433 | 1,37E-36 | 3,58E-33 | Up |
| Vmac | 530,7294514 | -0,732492135 | 0,126898283 | 9,20E-11 | 3,60E-09 | Down |
| Tnfsf14 | 2354,701557 | 0,915654542 | 0,144150049 | 2,08E-13 | 1,54E-11 | Up |
| C3 | 3832,766568 | 2,326262509 | 0,390995879 | 3,72E-11 | 1,63E-09 | Up |
| Adgre1 | 31351,15199 | 1,970374976 | 0,162599959 | 2,03E-35 | 4,55E-32 | Up |
| Xdh | 28,78667226 | 1,298972472 | 0,780113114 | 0,000183544 | 0,001220558 | Up |
| Dync2li1 | 41,62070145 | 1,273955344 | 1,130953998 | 0,000493821 | 0,002878669 | Up |
| Ppp1r21 | 389,9340194 | 1,461928442 | 0,635413021 | 3,46E-05 | 0,000287063 | Up |
| Svil | 19366,24402 | -0,716491192 | 0,111509806 | 2,08E-12 | 1,27E-10 | Down |
| Cabyr | 87,62900569 | -0,80000838 | 0,321033094 | 6,44E-05 | 0,000491969 | Down |
| Osbpl1a | 5962,158391 | -0,612064388 | 0,106197036 | 4,44E-10 | 1,44E-08 | Down |
| Slc39a6 | 7023,671152 | 0,696054973 | 0,087238824 | 3,56E-17 | 5,76E-15 | Up |
| Celf4 | 147,4746177 | 2,33240973 | 0,684812815 | 1,80E-06 | 2,13E-05 | Up |
| 1700066B19Rik | 786,8202738 | -0,707556534 | 0,172885734 | 5,52E-07 | 7,70E-06 | Down |
| Hbegf | 1402,236803 | 1,767831842 | 0,155518173 | 2,18E-31 | 3,12E-28 | Up |
| Cd14 | 159,6812712 | 1,798707996 | 0,249012186 | 1,29E-14 | 1,22E-12 | Up |

| | | | | | | |
|---|---|---|---|---|---|---|
| Wdr55 | 8507,810935 | 0,682624217 | 0,144992742 | 4,99E-08 | 8,96E-07 | Up |
| Spry4 | 227,1957364 | 3,25150434 | 0,346430064 | 1,99E-22 | 9,19E-20 | Up |
| Ap3s1 | 11592,20357 | 0,717924016 | 0,154028738 | 3,97E-08 | 7,27E-07 | Up |
| Cd74 | 1766,084264 | 1,999766275 | 0,500421902 | 3,07E-07 | 4,58E-06 | Up |
| Csf1r | 781,9611532 | 2,030317154 | 0,345109009 | 6,35E-11 | 2,61E-09 | Up |
| Pcyox1l | 10082,03128 | 0,581033591 | 0,073719213 | 2,78E-16 | 3,64E-14 | Up |
| Tubb6 | 18989,66917 | 0,756180716 | 0,074306933 | 2,37E-26 | 1,55E-23 | Up |
| Mc5r | 1407,292664 | -1,202741067 | 0,366942048 | 1,72E-06 | 2,05E-05 | Down |
| Rab27b | 2275,746957 | -0,629161204 | 0,180987973 | 1,30E-05 | 0,000122242 | Down |
| St8sia5 | 12,72983078 | 0,666021004 | 0,64268679 | 0,002816514 | 0,012694101 | Up |
| A330094K24Rik | 12,80696012 | -1,933244344 | 0,765977907 | 1,62E-05 | 0,000148468 | Down |
| Aldh3b1 | 123,1433698 | 1,286950895 | 0,421391714 | 3,96E-06 | 4,24E-05 | Up |
| Acy3 | 548,8331198 | -0,61211881 | 0,152523695 | 2,19E-06 | 2,51E-05 | Down |
| Gstp3 | 31,36297781 | 0,763741662 | 0,627731494 | 0,001510366 | 0,007511679 | Up |
| Pcx | 5514,382933 | -0,724016592 | 0,148480714 | 1,30E-08 | 2,72E-07 | Down |
| Slc29a2 | 2462,474498 | 1,188943843 | 0,182947744 | 1,17E-13 | 9,11E-12 | Up |
| Gm8034 | 63,1254761 | -0,69975479 | 0,240789468 | 3,94E-05 | 0,000321804 | Down |
| Tm7sf2 | 449,7034108 | -0,90472246 | 0,24239275 | 4,06E-07 | 5,87E-06 | Down |
| Rom1 | 416,3077795 | -0,603135065 | 0,114689057 | 7,83E-09 | 1,75E-07 | Down |
| Fads3 | 83,08442898 | -1,168168312 | 0,357459387 | 1,75E-06 | 2,08E-05 | Down |
| Fads2 | 31051,18057 | -0,995969241 | 0,205696316 | 1,30E-09 | 3,67E-08 | Down |
| Fads1 | 31775,70524 | -0,818159229 | 0,117773841 | 1,17E-14 | 1,12E-12 | Down |
| Slc15a3 | 17,59272379 | 1,737193047 | 1,841093292 | 0,000134998 | 0,000931765 | Up |
| Ccdc86 | 8287,309118 | 0,767897375 | 0,134592923 | 8,02E-11 | 3,19E-09 | Up |
| Ms4a2 | 7,863810689 | 1,954331514 | 1,779495747 | 0,000115705 | 0,000814337 | Up |
| Ms4a3 | 1878,313563 | 3,256790385 | 0,448539387 | 9,79E-15 | 9,55E-13 | Up |
| Stx3 | 660,5510656 | 0,623948566 | 0,08803628 | 7,62E-14 | 6,32E-12 | Up |
| Mpeg1 | 526,0471994 | 1,415045704 | 0,257412274 | 1,99E-10 | 7,12E-09 | Up |
| Dtx4 | 6304,999064 | 0,703014103 | 0,123020775 | 1,97E-10 | 7,05E-09 | Up |
| Glyat | 7,185423152 | 1,071648019 | 1,143457458 | 0,000877926 | 0,004720315 | Up |
| Gna14 | 108,4615706 | -1,119048499 | 0,463848913 | 3,24E-05 | 0,000271139 | Down |
| Gm8250 | 25,68361481 | -0,96966506 | 0,426149384 | 6,37E-05 | 0,00048807 | Down |
| Kank1 | 118,0666616 | 3,392950766 | 0,322587761 | 2,78E-27 | 2,19E-24 | Up |
| 4430402I18Rik | 89,91632962 | -0,770135686 | 0,234621325 | 6,26E-06 | 6,43E-05 | Down |
| Cd274 | 1913,758696 | 0,65525535 | 0,106150533 | 2,39E-11 | 1,11E-09 | Up |
| Pdcd1lg2 | 271,738588 | 0,670728346 | 0,287347618 | 0,000234081 | 0,001504011 | Up |
| Ankrd22 | 50,41664981 | 2,977884673 | 0,693563982 | 1,27E-07 | 2,07E-06 | Up |
| Acta2 | 609,0612816 | 0,633043967 | 0,199587522 | 3,34E-05 | 0,000278875 | Up |
| Cpeb3 | 150,1406492 | -0,639307045 | 0,199973206 | 2,90E-05 | 0,00024587 | Down |
| Noc3l | 5446,023547 | 0,595122183 | 0,108284875 | 2,35E-09 | 6,17E-08 | Up |
| Rrp12 | 16668,34226 | 0,710941517 | 0,110823815 | 2,42E-12 | 1,43E-10 | Up |
| Exosc1 | 6112,409885 | 0,593601154 | 0,066797505 | 5,18E-20 | 1,54E-17 | Up |
| Gm16244 | 65,9803612 | -0,59920804 | 0,236877557 | 0,000267643 | 0,001696763 | Down |
| Cpn1 | 84,37942997 | -0,64647426 | 0,197123421 | 2,08E-05 | 0,00018513 | Down |
| Scd2 | 130084,5421 | -0,844148551 | 0,215393441 | 2,75E-07 | 4,19E-06 | Down |
| Nolc1 | 64907,13545 | 0,686797429 | 0,128725681 | 2,02E-09 | 5,39E-08 | Up |
| Obfc1 | 1182,931479 | -0,615030606 | 0,100132756 | 4,49E-11 | 1,90E-09 | Down |
| Gm19557 | 53,97645203 | -0,984753439 | 0,389917418 | 2,83E-05 | 0,000240447 | Down |
| Pdcd4 | 1960,934207 | -0,959632478 | 0,182295564 | 1,39E-10 | 5,25E-09 | Down |
| Ablim1 | 17995,92626 | 0,830985595 | 0,192566163 | 5,24E-08 | 9,32E-07 | Up |
| Emx2 | 40,39203557 | 1,149471754 | 1,638394554 | 0,00031956 | 0,001981929 | Up |

| | | | | | | |
|---|---|---|---|---|---|---|
| Csf2ra | 799,8728441 | 1,127059451 | 0,317020143 | 5,60E-07 | 7,79E-06 | Up |
| Ccdc120 | 1294,409318 | -0,589471835 | 0,090439517 | 5,17E-12 | 2,79E-10 | Down |
| Gata1 | 227,7361185 | -0,614942964 | 0,547623125 | 0,00323172 | 0,014198968 | Down |
| Cfp | 7659,197069 | 1,697590528 | 0,279693145 | 2,16E-11 | 1,02E-09 | Up |
| Gm4764 | 15,87551447 | 0,808431372 | 0,611243268 | 0,001078784 | 0,005612107 | Up |
| Ap3s1-ps1 | 33,2590392 | 0,590282812 | 0,360570612 | 0,00176747 | 0,0085985 | Up |
| Plac1 | 7,696839202 | 1,338023068 | 1,006912293 | 0,000333467 | 0,002052785 | Up |
| Gm364 | 50,56655528 | 1,255075228 | 0,443215238 | 8,10E-06 | 8,04E-05 | Up |
| Nsdhl | 6551,291164 | -0,925857605 | 0,201930721 | 6,60E-09 | 1,51E-07 | Down |
| 4933407K13Rik | 225,7767179 | -1,37760125 | 0,414294532 | 1,72E-06 | 2,05E-05 | Down |
| Gm7153 | 20,38094683 | -0,709202149 | 0,485374554 | 0,001227531 | 0,00629836 | Down |
| Pcyt1b | 237,5326495 | -1,685669727 | 0,254290681 | 7,15E-13 | 4,80E-11 | Down |
| Rpl17-ps8 | 110,639247 | -0,642178537 | 0,195306115 | 2,12E-05 | 0,000187706 | Down |
| Eda2r | 159,2036293 | 0,934071911 | 0,264755833 | 8,33E-07 | 1,09E-05 | Up |
| Foxo4 | 2174,821687 | -0,591190155 | 0,120754238 | 5,48E-08 | 9,70E-07 | Down |
| Nhsl2 | 4186,278228 | 0,62869008 | 0,147206845 | 6,48E-07 | 8,82E-06 | Up |
| Tlr13 | 26,2013956 | 2,203428975 | 0,801878798 | 8,89E-06 | 8,72E-05 | Up |
| Gm7429 | 22,8735666 | -0,607340573 | 0,367139536 | 0,001527573 | 0,007580465 | Down |
| Nap1l3 | 2142,146348 | 0,619150905 | 0,157184725 | 2,72E-06 | 3,04E-05 | Up |
| Diaph2 | 7804,347423 | 0,654776578 | 0,208072098 | 2,91E-05 | 0,000246108 | Up |
| Timm8a1 | 2632,332527 | 0,76001678 | 0,124558965 | 8,30E-12 | 4,22E-10 | Up |
| Gla | 2434,275006 | 0,601500494 | 0,148338198 | 2,06E-06 | 2,38E-05 | Up |
| Armcx1 | 47,24106716 | -1,310519853 | 0,534199127 | 2,49E-05 | 0,000214458 | Down |
| Tsc22d3 | 849,7772306 | -0,69105617 | 0,21490736 | 1,64E-05 | 0,000149562 | Down |
| Trpc5os | 22,13987163 | 1,200695834 | 1,590862317 | 0,000385946 | 0,00232307 | Up |
| Fancb | 1400,655085 | 0,735443238 | 0,089653192 | 3,03E-18 | 6,27E-16 | Up |
| Arhgap6 | 8355,513282 | -0,716138565 | 0,21306633 | 7,89E-06 | 7,86E-05 | Down |
